# Metastasis founder cells activate immunosuppression early in human melanoma metastatic colonization

**DOI:** 10.1101/2024.05.08.591627

**Authors:** Severin Guetter, Courtney König, Huiqin Koerkel-Qu, Aleksandra Markiewicz, Sebastian Scheitler, Marie Katzer, Mark Berneburg, Philipp Renner, Beatrix Cucuruz, Leonhard Guttenberger, Veronika Naimer, Kathrin Weidele, Steffi Treitschke, Christian Werno, Hanna Jaser, Tonia Bargmann, Armin Braun, Florian Weber, Reinhard Rachel, Felix Baumann, Lisa Schmidleithner, Kathrin Schambeck, Parvaneh Mohammadi, Anja Ulmer, Sebastian Haferkamp, Christoph A. Klein, Melanie Werner-Klein

**Author notes:** equal contribution (Severin Guetter, Courtney König, Huiqin Koerkel-Qu, Christoph A. Klein, Melanie Werner-Klein).

## Abstract

The earliest steps of lethal metastasis in patients are incompletely understood. To dissect them, we prospectively searched for the earliest detectable disseminated cancer cells (DCC) in sentinel lymph node biopsies of 492 stage I-III patients. By visually-controlled, micromanipulator-assisted isolation and single cell transcriptome analysis of these extremely rare DCC, we identified MCSP^+^ melanoma cells as strong candidates for metastasis founder cells (MFC) in lymph nodes. Based on a median follow-up time of 6 years, their detection was the strongest predictor of systemic metastasis and death upon multivariable analysis. During transition from single cells to metastasis-initiating clusters, melanoma DCC were exposed to CD8 T cell attack, activated the extracellular vesicular exosomal pathway, and expressed the immunomodulatory proteins CD155 and CD276, but rarely PD-L1. CD155 and CD276-positive extracellular vesicles from patient-derived DCC models exhibited an immunosuppressive activity on CD8 T cells. Our data indicate that either direct targeting of MFC employing MCSP or their immune escape mechanisms might be key for cure of early-stage melanoma.

## Introduction

Despite recent progress in immunotherapy of melanoma, more than 40 % of patients developing metastasis will inevitably die from the disease ^1^. Early prevention of metastasis formation therefore is an important medical goal. At diagnosis, however, almost 50 % of melanomas have disseminated to regional or distant sites even from tumour stages measuring ≤ 1 mm thickness (T1 stage) ^2^. We previously found that (i) most of these early disseminated cancer cells (DCC) are not (yet) able to form a metastatic colony as they are lacking important genetic changes and (ii) that such changes are acquired during cell divisions within the metastatic site ^2^. We therefore concluded that melanomas (and other cancers such as non-small-cell-lung cancer as well ^3^) undergo molecular evolution at distant sites in parallel to the primary tumour ^4^. Yet, the metastasis founder cells and the molecular features enabling and driving this evolution are currently unknown.

Experiments to identify a cancer stem cell phenotype in melanoma have resulted in controversy ^5–9^. A major conclusion from these experiments, however, was that stemness in melanoma is not strictly hierarchical but associated with a plastic state that has been postulated (but not demonstrated) to be relevant for metastasis in patients. As these experiments used primary melanomas or macroscopic metastases, i.e. melanoma cells from evolutionary advanced stages as opposed to more adequate representatives of early stages of systemically spread cancer cells, we attempted to dissect phenotypic progression in early human metastasis formation from its earliest beginnings after homing to an ectopic site (here: lymph node, LN) over incipient colony- and subsequent micro- and macrometastasis formation. We chose LN metastasis because samples are easily available, LN status informs directly about the risk to die from melanoma and because exposure to the lymphatic environment protects melanoma cells from ferroptosis and increases their ability to survive during subsequent metastasis through the blood ^10^. We exploited a quantitative assay to measure lymph node invasion, i.e. quantitative immunocytology based on gp100-staining of melanoma DCC ^11^, which determines the exact number of gp100-positive cells within one million lymph node cells and is expressed as disseminated cancer cell density (DCCD). This assay (i) provided best accuracy for DCC detection as single marker assay ^12^; (ii) demonstrated that every detected cancer cell increases the risk to die from melanoma ^11^; (iii) established that a DCCD around 100 marks the formation of microscopic metastasis ^2^; and (iv) revealed that specific genetic alterations are acquired at exactly this stage and DCCD ^2^. However, an intracellular marker such as gp100 precludes the isolation of viable cells for transcriptomic analyses. We therefore used the gp100-assay as reference to rank the phenotypic progression and microenvironmental responses during early metastasis formation and compared it with melanoma cell detection using MCSP (melanoma associated chondroitin sulfate proteoglycan). We chose MCSP for two reasons, first, because as a cell surface marker MCSP enables further molecular characterization of isolated cells by assessing their transcriptome and second, because published data as well as our own work had indicated that MCSP is expressed by cancer stem-like cells ^2,13^. Importantly, unlike scRNA-seq studies before, we aimed to capture the earliest metastatic steps from single invading melanoma cells to early micrometastatic colonies. High-throughput scRNA-seq approaches require target cell concentrations of at least 1 in 1000 per sample. However, since at colonization DCCD values are much lower than this, ranging from 1-200 cells per million lymph node cells, we performed scRNA-seq after microscopic inspection and micromanipulator-assisted manual isolation of candidate MFC.

## Results

### MCSP as surface marker for melanoma DCC detection

Staining for the transmembrane glycoprotein MCSP (*CSPG4*) in 625 lymph nodes of 492 early-stage melanoma patients (Fig. 1a, ED Fig. 1a) identified two morphologically distinct MCSP^+^ cell populations, large cells with a diameter of about 20 µm and bright fluorescence and small cells with a diameter of half the size (10 µm) and often weaker fluorescent staining (Fig. 1b, ED Fig. 1b). MCSP-positive lymph nodes (n = 477) with only small cells were more frequently recorded (79 %; 378/477) than samples with a mixture of small and large cells (12 % (58/477)), or large cells only (9 %; 41/477; ED Fig. 1c). The median DCCD_MCSP_ increased with the presence of large cells (Fig. 1c, *P* < 0.0001, Wilcoxon test) and direct comparison of the DCCD_gp100_ and DCCD_MCSP_ from the same lymph nodes (n = 542), revealed a significant correlation for samples containing MCSP^+^ cells of the large but not the small phenotype (Fig. 1d, *P* < 0.0001, Spearman’s correlation). Of note, 51 % (275/542) of the gp100-negative sentinel lymph node (SLN) samples harboured MCSP^+^ cells, whereas only 3 % (17/542) of the MCSP-negative SLN were gp100-positive (ED Fig. 1d). Since this discrepancy was predominantly caused by cells with small-cell morphology, we asked whether all small cells are indeed melanoma cells.

**Figure 1:**
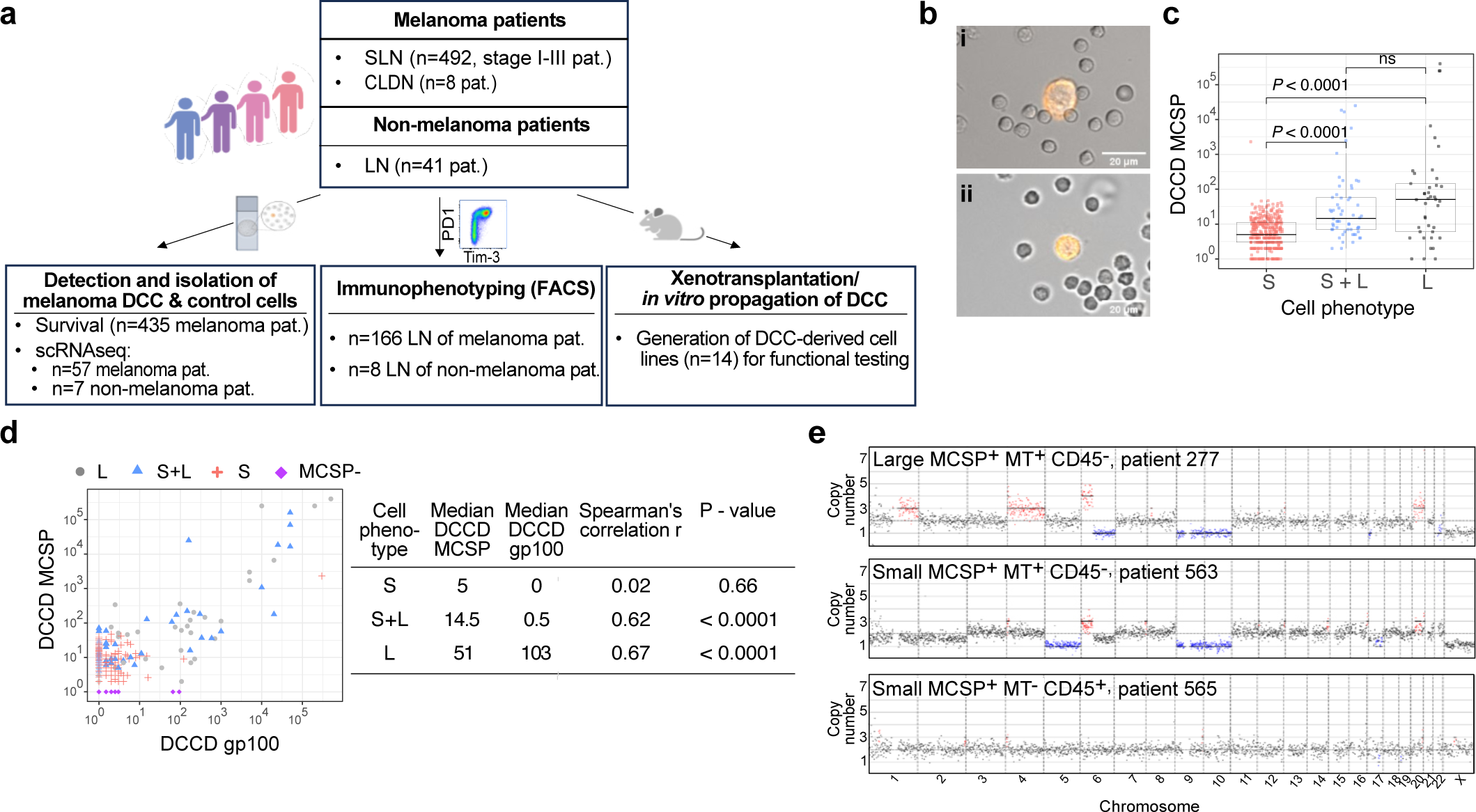
Identification of DCC using melanoma markers gp100 and MCSP. **a,** Study design. SLN = sentinel lymph node, LN = lymph node, CLND = complete lymphadenectomy, pat. = patients. Created with BioRender.com. **b**, MCSP-staining intensity and diameter of MCSP-positive cells in SLN of melanoma patients with large, intensely stained cells (i) and small cells (ii). Scale bars as indicated on merged images of fluorescence and brightfield channels. **c**, DCCD_MCSP_ of lymph nodes with MCSP^+^ cells (n = 477 SLN from 392 patients) separated according to detected cellular phenotypes (diameter) into small (S, n = 378), small and large (S+L, n = 58) and large (L, n = 41). **d**, Correlation of DCCD_MCSP_ and DCCDgp100 in SLN stained for both MCSP and gp100 (n = 542 SLN, 430 patients). Displayed are phenotypes of MCSP^+^ cells (S, n = 335; S+L, n = 54; L, n = 37) and SLN negative for MCSP (no MCSP^+^ cells, n = 116). **e**, Representative CNA profiles of small and large MCSP^+^MT^+^CD45^-^ and small MCSP^+^MT^-^CD45^+^ cells. *P* values in **c,** Wilcoxon test, and **d,** Spearman’s rank correlation. Boxes mark the median, lower-quartile, and upper-quartile, and lines connecting the extremes.

### Identification of MCSP^+^ melanoma cells

We found small MCSP^+^ cells in 85 % (35/41) of non-melanoma patients, albeit at low frequency (median DCCD = 4, range 0-15) ED Fig. 1b, ED Fig. 3a). We then isolated 1606 MCSP^+^ cells from 477 SLN samples of 392 melanoma patients, performed whole transcriptome amplification ^14,15^ and tested all high-quality single cells (n = 1026) for the melanoma-associated transcripts (MT) *PMEL* (gp100), *DCT* (dopachrome tautomerase) and *MLANA* (melan A). Large MCSP^+^ cells (n = 237) predominantly expressed all three investigated MT and were obviously of melanoma origin (ED Fig. 1e). Of small MCSP^+^ cells, 9 % (74/789) expressed at least one of the MT, indicating that a small but significant population of these cells are indeed of melanocytic origin.

We next tested the leukocyte marker CD45 and found that 50 % (360/715) and 53 % (32/61) of small MCSP^+^MT^-^ cells from melanoma patients and non-melanoma patients, respectively, expressed the *CD45* transcript (ED Fig. 1f). In summary, of the 789 small MCSP^+^ cells, we identified 9 % (74/789) as putative melanoma cells based on their expression of MT (ED Fig. 1e) 46 % (360/789) as CD45 positive lymphocytes, and 45 % (355/789) as cells of currently unknown non-melanoma lineage (ED Fig. 1f). Since copy number alterations (CNA) have been shown to not only differentiate between normal and malignant cells but also between malignant melanoma cells and rare lymph node-residing nevus cells ^16,5,17^, we tested small (n = 15) and large (n = 13) MCSP*^+^*MT*^+^*CD45^-^ and MCSP^+^MT^-^CD45^+^ (n = 15) for CNA after whole genome amplification of the single cell DNA ^15^. None of the small MCSP^+^MT^-^CD45^+^ cells harboured CNA, in contrast to 100 % of small (n = 15) and large (n = 13) MCSP^+^MT^+^CD45^-^ cells (Fig. 1e). Comparing the cumulative profiles of CNA in small and large MCSP^+^MT^+^CD45^-^ cells, we found a high similarity (ED Fig. 1g). Thus, genomic analysis confirmed the malignant origin of small and large MCSP^+^MT^+^ cells. A side-by-side comparison of patient SLN for which both the MCSP/MT and gp100 status was available (n = 380, ED Fig. 2a) revealed that the positivity rate was significantly higher for gp100 than for MCSP^+^MT^+^ (MCSP^+^MT^+^ = 86/380 (22 %) vs. gp100^+^ 142/380 (37 %), P < 0.0001 Fisher’s exact test). This difference prompted us to investigate the prognostic impact of MCSP^+^MT^+^ cells on progression-free (PFS), melanoma-specific (MSS) and overall survival (OS).

**Figure 2:**
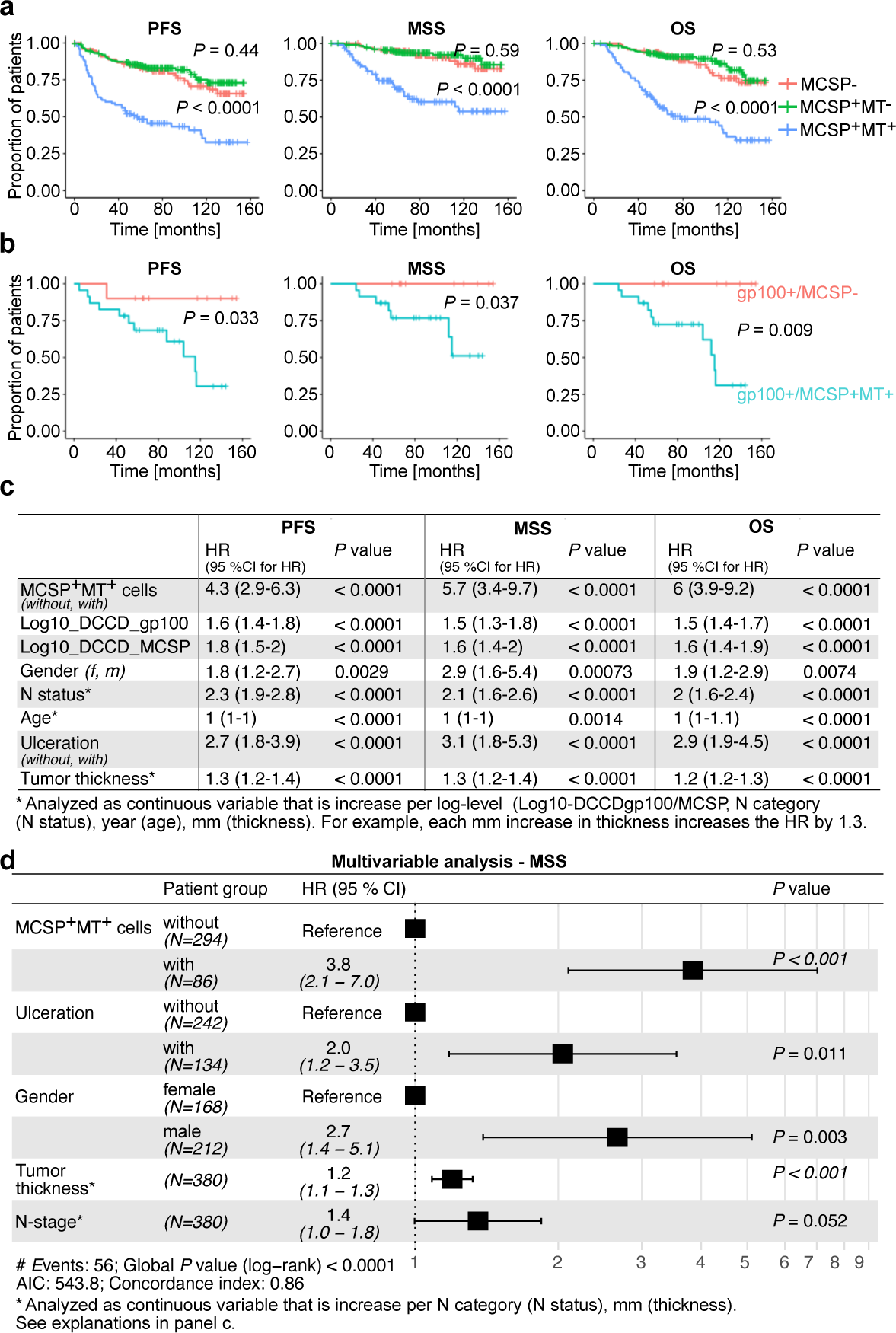
Patient survival in relation to DCCD_gp100_ and DCCD_MCSP_. **a, b,** Kaplan Meier curves of progression-free (PFS), melanoma-specific (MSS) and overall survival (OS) of patients stratified according to their DCCD_MCSP_, DCCD_gp100_ and MT expression of MCSP positive cells. Patients with lymph nodes without MCSP-cells (n = 99), positive for MCSP^+^/MT^-^ cells (n = 238) or positive for MCSP^+^/MT^+^ cells (n = 98) (a) and patients with gp100 positive lymph nodes but MCSP negative (n = 10) or gp100 negative lymph nodes containing MCSP^+^MT^+^ cells (n = 23) (b). **c**, Univariable analysis for PFS, MSS and OS (n = 380). Ulceration group “yes, no” denotes patients without and with ulceration, gender group “female, male” patients and MCSP^+^MT^+^ group “yes, no” categorizes patients with and without MCSP^+^MT^+^ cells. **d,** Multivariable Cox regression analysis for MSS comprising the most informative, backward selected features (n = 380). Ulceration group a and b denotes patients without and with ulceration, gender group f and m female and male patients and MCSP^+^MT^+^ group 0 and 1 categorize patients without and with MCSP^+^MT^+^ cells. Group 0 MCSP^+^MT^+^ patients, male patients and patients without ulceration were used as reference. *P* values in **a, b,** Log-rank test. **c, d,** Wald-test. See Supplementary Table 1 for baseline characteristics of patients.

### MCSP^+^ melanoma cells predict poor outcome

We compared three groups: patients with lymph nodes (i) negative for MCSP expressing cells (n = 99); (ii) positive for MCSP^+^MT^-^ cells (n = 238) and (iii) positive for MCSP^+^MT^+^ cells (n = 98), summing up to 435 patients for whom follow-up information for PFS, MSS and OS was available (ED Fig. 2a, Supplementary Table 1). The strongest impact for all endpoints was found for MCSP^+^MT^+^ samples (for all *P* < 0.001, Log-rank test, (Fig. 2a)). In contrast, MCSP^+^MT^-^ patients displayed an almost identical outcome as MCSP-negative patients, consistent with a classification of MCSP^+^MT^-^ cells as non-malignant cells. Our standardized single cell isolation protocol of MCSP^+^ cells for RNA analysis stops after isolation of 5–10 cells and hence, the MT status is not available for additionally detected and counted, but not isolated MCSP^+^ cells. Therefore, the DCCD_MCSP_ overestimates the number of MCSP^+^MT^+^ cells (because DCCD_MCSP_ = DCCD_MCSP+MT+_ + DCCD_MCSP+MT-_), particularly when more than 10 cells were detected. We could, however, directly compare how single positivity (i.e. either for gp100 or MCSP^+^MT^+^ within a sample) impacts on survival. Of the 380 patients, for which gp100 and MCSP staining results were available, we identified 23 patients with only MCSP^+^MT^+^ cells in their SLN and 10 patients lacking MCSP^+^MT^+^ cells, but harboring gp100^+^ cells (ED Fig. 2a). Strikingly, patients harboring only MCSP^+^MT^+^ cells and lacking gp100^+^ cells had a significantly worse outcome for all endpoints (Fig. 2b, *P* = 0.033 for PFS, *P* = 0.037 for MSS and *P* = 0.009 for OS, Log-rank test) than the MCSP-negative patients with gp100^+^ cells. The fact that MCSP^+^ DCC carry the prognostic information was further supported by splitting patients with gp100^+^ DCC into those with and without MCSP^+^ DCC. For all endpoints poor outcome was observed for patients with MCSP^+^ DCC (*P* < 0.0001; ED Fig. 2d). Therefore, using only gp100 as detection marker failed to detect melanoma DCC with metastasis founder potential in about 6 % (23/380) of patients. Equally important, exclusive gp100-positivity will overestimate the patient’s risk for reduced PFS, MSS and OS in about every second gp100^+^ patient, as 55 % (79/142) of them are lacking MCSP^+^MT^+^ DCC (ED Fig. 2a). This pointed to a strong independent risk contribution by MCSP^+^MT^+^ DCC, prompting us to assess the impact of DCCD_MCSP_ and MT status of the 380 patients together with known clinical risk factors, such as Breslow thickness, N status, ulceration, gp100, sex and age. Upon univariable analysis, both gp100 and MCSP counts had a strong impact on PFS, MSS and OS (all *P* < 0.0001, Wald-test), as had the detection of MT in MCSP^+^ cells (Fig. 2c). However, multivariable analysis revealed that detection of MCSP^+^MT^+^ melanoma cells in SLN was the strongest risk factor together with melanoma thickness, outperforming all other variables, including DCCD_gp100_ for PFS, MSS and OS (Fig. 2d and ED Fig. 2b, c). Given their recurrent detection in gp100 negative samples and impact on patient survival, (small and large) MCSP^+^MT^+^ cells may comprise the so far earliest known harbingers of metachronous metastasis and death in malignant melanoma. Since cutaneous melanoma patients do not die from lymph node metastasis but from distant metastasis, we directly evaluated the risk for distant metastasis of patients that showed loco-regional lymphatic spread. Here, patients with SLN-involvement show a four-fold higher frequency of distant metastasis than patients without evidence for local lymph node involvement during the course of the disease (ED Fig. 2e; *P* < 0.0001). Thus, the vast majority of patients (78 %, 32/41) develop systemic melanoma with lymphatic involvement whereas in few cases (12 %, 9/41) melanoma cells bypass the draining lymph node basin leading to exclusive haematogenous spread.

### Molecular characterization of melanoma DCC

To gain insight into the molecular characteristics of candidate MFC, we performed scRNA-seq. We isolated lymph node-derived small and large MCSP^+^MT^+^ cells (n = 170) and MCSP^+^MT^-^ cells (n = 23) from a total of 77 melanoma patients and lymph node-derived MCSP^+^MT^-^ cells (n = 9) from 7 non-melanoma patients. These cell numbers were obtained after screening of 492 melanoma and 41 non-melanoma patients and applying stringent quality controls (ED Fig. 3a). In addition, we tested cultured human melanocytes (n = 14 cells, 1 donor; ED Fig. 3a). Expression based dimension reduction separated DCC (MCSP^+^MT^+^), non-melanoma cells (MCSP^+^MT^-^) and melanocytes (ED Fig. 3b). Transcript-based CNA analysis and transcript expression of CD45 (*PTPRC*) and Melan A (*MLANA*) largely matched uniform manifold and projection (UMAP) dimension reduction (ED Fig. 3c, d): the DCC group comprised 139/164 (85 %) cells with inferred genomic aberrations, whereas melanocytes or immune cells had none (0/14) or only 5 % (2/38), respectively. Integration of the cells into the Human cell atlas confirmed their cell types (ED Fig. 3e). Dimension reduction of the DCC cluster resolved transcriptomes into five clusters (Fig. 3a). All five clusters overlapped with previously identified melanoma transcriptional signatures in human ^18–22^ and mouse ^23,24^ data sets (ED Fig. 3f). Of note, MCSP-expression was not only detected across all clusters in our DCC collective, but also abundantly in published data sets on fetal development of human epidermal melanocytes ^25^, mouse NRAS/BRAF-mutated melanoma models ^23^ and primary and metastatic cell lines from the Cancer Cell Line Encyclopedia (CCLE ^26^; ED Fig. 4a), further validating MCSP as relevant marker for DCC detection. We adopted the melanoma subtype nomenclature proposed by Tsoi et al., derived from an *in vitro* differentiation assay of human embryonic stem cells into melanocytes that was validated using human melanoma cell lines and patient samples ^21^. Each of the five DCC clusters was significantly associated with one of these subtypes (transitory (cluster 0 and 3), neural crest-like (cluster 1) and melanocytic (cluster 4) all *P* < 0.001, undifferentiated (cluster 2), *P* = 0.011; ANOVA with post-hoc analysis, ED Fig. 3f, Fig. 3b, Supplementary Table 2). Most strikingly, cluster annotation was associated with DCCD_gp100_ values (*P* < 0.0001, Fig. 3c, ANOVA with post-hoc analysis, Supplementary Table 2), with median DCCD_gp100_ values rising from 15 (early-transitory), over 61 (NC-like), 334 (undifferentiated), 20,000 (late-transitory) to 37,500 (melanocytic) for Seurat groups 0-4, respectively. This strongly indicates that melanoma cells dynamically switch their phenotype during lymph node colonization.

**Figure 3:**
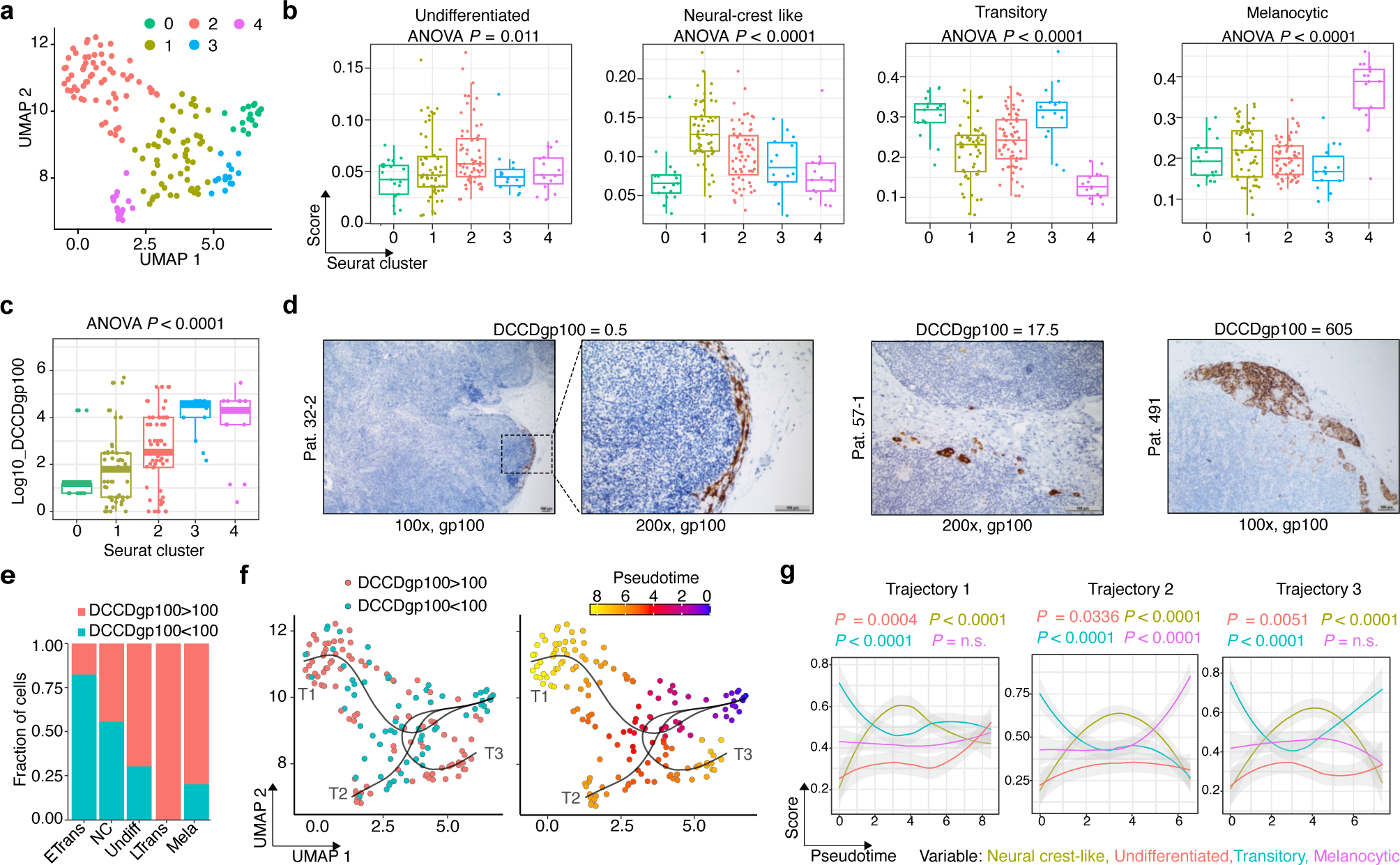
Molecular characterization of melanoma DCC. **a,** Louvain clustering of single cell RNA-seq data of the DCC cluster (n = 164 cells, see ED Figure 3d) with Seurat based on dimension reduction result of UMAP. **b,** Signature scores of the four melanoma phenotypes ^21^ for DCC annotated by Seurat cluster label. **c,** DCCD_gp100_ annotated by Seurat cluster label. **d,** Representative examples of SLN analyzed by histopathology and immunocytology, using gp100 staining. The examples illustrate that a DCCD_gp100_ < 100 in immunocytology corresponds to isolated tumour cells in histopathology, while a DCCD_gp100_ > 100 corresponds to micrometastasis ^2^ . **e,** Percentages of the four melanoma phenotypes ^21^ among DCC before (DCCD_gp100_ < 100) and after metastatic colony formation (DCCD_gp100_ > 100). **f,** Inferred trajectories (T1, T2, T3) with slingshot. Left: each cell is colored according to its DCCD (DCCD_gp100_ < 100 blue, DCCD_gp100_ > 100 red). Right: each cell is colored according to its pseudotime. **g,** Melanocytic, neural-crest-like, transitory and undifferentiated signature scores ^21^ with AUCell along pseudotime (slingshot) of trajectory 1-3. *P* values in **b, g,** one-way ANOVA; results of post-hoc analysis are either indicated (f) or found in Supplementary Table 3 as well as *P* values for **f.** Boxes mark the median, lower-quartile, and upper-quartile, and lines connecting the extremes.

**Figure 4:**
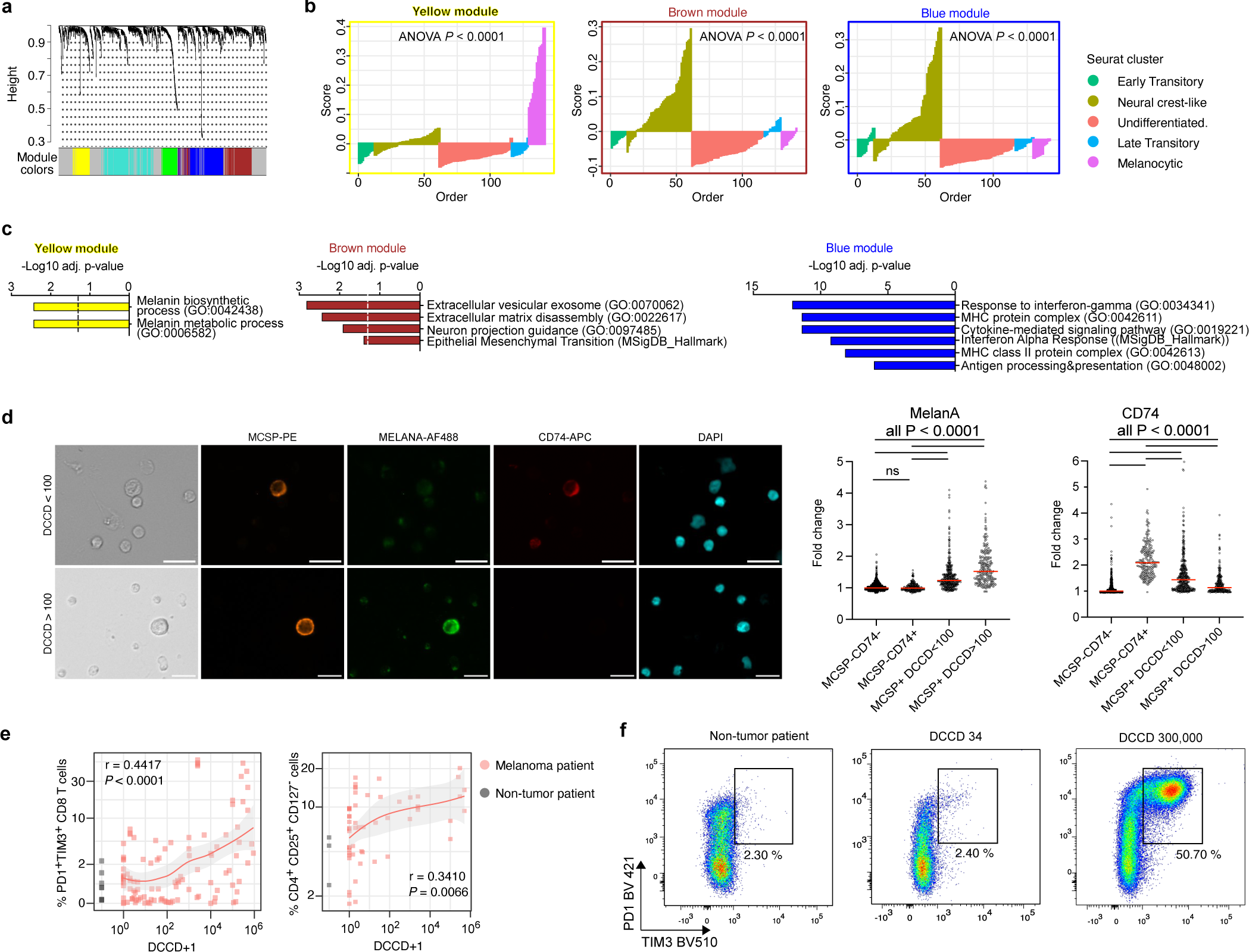
Identification of differentially expressed gene-signatures in melanoma DCC and flow cytometric analysis of lymph node cells. **a, b,** Single cell weighted gene co-expression network analysis using top variable genes. Cluster dendrogram showing genes grouped into distinct modules (a). Module score summaries are shown with values of the components of the module versus DCC samples of distinct Seurat clusters (x-axis) (b). **c,** Enrichr gene set enrichment analysis. Enriched GO terms and terms of the Hallmark collection in the Molecular Signature Database assigned to genes in modules brown, blue and yellow. **d,** Immunofluorescent staining and quantification of CD74 and MelanA fluorescence intensity in MCSP^+^, MCSP^-^ cells in SLN with DCCD < 100 (n = 28 patients/SLN) and DCCD > 100 (n=13 patients/SLN). Images: representative staining of MCSP, CD74, MelanA and DAPI with scale bars, 20 μm. Plots depict fold change in gray value of MelanA or CD74 immunofluorescence of MCSP^+^ (DCCD < 100: n = 402 cells; DCCD > 100: n = 230 cells) and MCSP^-^CD74^+^ cells (n = 366 cells) over the respective median gray value of MCSP^-^ CD74^-^ cells (n = 765 cells). **e,** Percentage of PD1^+^TIM3^+^ CD8 T cells or CD4^+^CD25^+^CD127^-^ regulatory T cells versus the DCCD of lymph nodes from melanoma (red, n = 116 for PD1^+^TIM3^+^ CD8 T cells; n = 57 for regulatory T cells) and non-melanoma patients (grey, n = 8 for PD1^+^TIM3^+^ CD8 T cells and n = 3 for regulatory T cells)). Red and grey symbols indicate detected values of individual lymph nodes from melanoma and non-melanoma patients, respectively. The grey area indicates the standard error of the smoothed curve (red line). **f,** Representative flow cytometric analysis for PD1 and TIM3 expression in CD3^+^CD8^+^ T cells in lymph nodes of a non-tumour patient or two lymph nodes of the same regional bed of a melanoma patient. *P* values in **b,** one-way ANOVA, results of post-hoc analysis see Supplementary Table 3. **c,** Hypergeometric test **e,** Pearson’s correlation.

We had previously documented that a DCCD of around 100 marks the transition from individual cells to detectable colony-formation as assessed by histopathology ^2,12^ (Fig. 3d). Interestingly, we found that the early-transitory and NC-like phenotypes dominated DCCD values < 100, whereas undifferentiated, late-transitory and melanocytic phenotypes were mostly found at DCCD > 100 (Fig. 3e). Slingshot and ELPIGraph-based pseudotime-analysis algorithms ^27,28^, two methods specifically validated for data sets of dozens to few hundreds samples, consistently identified trajectories from early-transitory cells entering the lymph nodes over an intermediate phenotypic NC-like stage, from which colony-expanding DCC further progress to either late-transitory, undifferentiated or melanocytic phenotypes (Fig. 3f, g; ED Fig. 4b-d).

### NC-like DCC activate exosomal vesicle production and interferon γ response

To identify which pathways are characteristic for the different phenotypes, we performed single cell weighted gene co-expression network analysis (scWGCNA; Fig. 4a), which revealed gene modules that were significantly differentially activated between the Seurat clusters and comprised significantly enriched pathways (all *P* < 0.0001, ANOVA with post-hoc analysis Fig. 4b, Supplementary Table 2). For example, DCC with a melanocytic signature showed enrichment for GO terms for melanin biosynthetic and metabolic processes (yellow module; Seurat group 4; Fig. 4b, c). In contrast, DCC with a NC-like signature (Seurat group 1) were enriched in GO terms associated with extracellular vesicular exosome production and epithelial mesenchymal transition (brown module) or response to interferon gamma and antigen processing/presentation (blue module) (Fig. 4c), which was also supported by gene regulatory network analysis (ED Fig. 5a). Of note, the activation of the blue module in NC-like cells was reminiscent of a recent analysis of lymph node metastasis in a mouse model ^24^. Here, B16 melanoma cells were selected for lymph node metastasis by iterative injection and lymph node isolation and switched from early-transitory to a NC-like gene expression phenotype after 6-8 (!) rounds (ED Fig 3f; Reticker-Flynn Parentalup vs. LNup (LNup associated with NC-like DCC cluster1). Apparently, forced adaptation in the mouse resulted in an upregulation of immune pathways similar to human NC-like DCC at incipient metastatic colonization. To corroborate our finding that DCC encompass an intermediate phenotypic NC-like stage before colony-expansion, we stained an independent set of patient samples (n = 41 patients/SLN) for MCSP (DCC detection marker), MelanA (melanocytic marker) and CD74, the invariant light chain of the HLA Class II Histocompatibility complex. CD74 was selected, as it was found to be significantly upregulated in the NC-like Seurat cluster 1 and is part of the antigen-presenting pathway (Supplementary Table 3). In line with our findings from pseudotime analysis, CD74 expression was significantly higher in MCSP^+^ cells in the DCCD < 100, NC-like group than in the DCCD > 100 group, which included further differentiated cells with higher MelanA expression (Fig. 4d).

**Figure 5:**
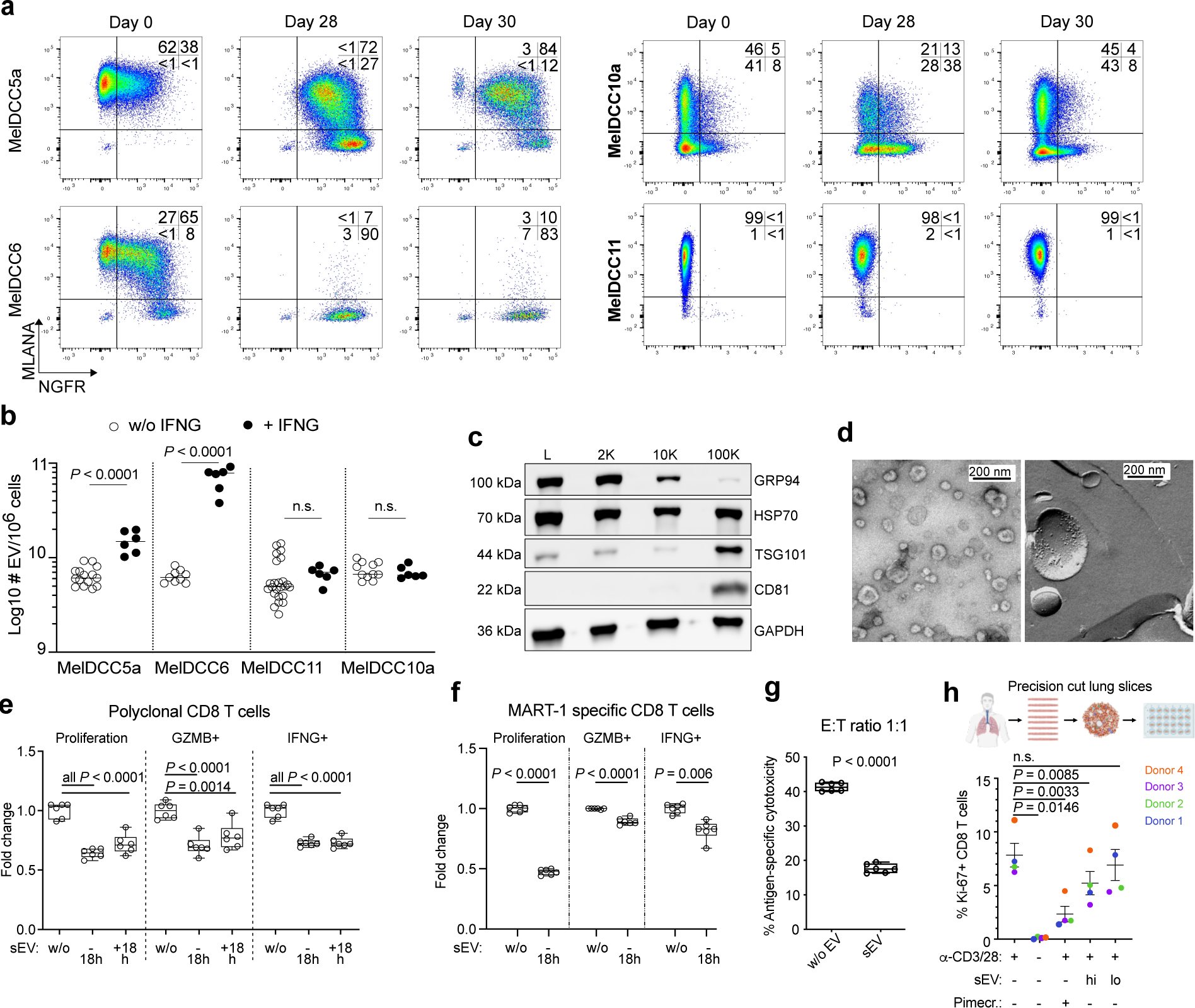
Characterization and impact of DCC-derived sEV on CD8 T cell function. **a,** Flow cytometric analysis of MelDCC 5a, 6, 10a and 11 for MelanA and NGFR expression before IFNG-treatment (d0), after 4 weeks of IFNG treatment (d28) or after 4 weeks of IFNG treatment followed by culture for 48 h in IFNG-free EV-production medium (d30). **b,** number of sEV secreted per million of MelDCC within 48 h. MelDCC were either untreated or treated with 500 U IFNG for 28 days before the medium was replaced for 48 h by EV-production medium. **c,** Western blot analysis of successive pellets (2K, 10K, 100K) of EV isolated from MelDCC10a using antibodies to proteins marking small (CD81, TSG101) and large EV (GRP94) or as pan EV-marker (HSP70). **d,** Transmission electron microscopy of 100K preparations. Left: negative staining, right: freeze-etching. **e, f,** Flow cytometric analysis of proliferation and production of effector cytokines IFNG and GZMB of polyclonal (e) or MART1_27L26-35_-specific CD8 T cells (f) exposed to sEV of MelDCC 10a (n = 6) or PBS (ctrl), n = 6) 18 h before or after anti-CD3/CD28 stimulation. **g,** Cytotoxic activity of MART1_27L26-35_-specific CD8 T cells exposed (n = 6) or non-exposed (n = 6) to sEV of MelDCC 10a prior to anti-CD3/CD28 stimulation. CD8 T cells were harvested at day 4 and added at an effector:target ratio of 1:1 to MART1_27L26-35_-loaded CFSE-labeled T2 cells. Non-loaded, CellTrace Violet labeled T2 cells served a non-target reference population. **h,** Flow cytometric analysis of the % of Ki67^+^ CD8 T cells in human PCLS 5 days after anti-CD3/CD28 stimulation and exposure to sEV of MelDCC 10a at high or low dose (sEV produced by 6.25×10^6^ and 1.25×10^6^ MelDCC 10a, respectively) or in the presence of 5 µM of the immunosuppressive pimecrolimus. Non-stimulated PCLS served as control. sEV or the pimecrolimus were added 18 h after anti-CD3/CD28 stimulation. Illustration created with BioRender.com. *P* values in **e, h** one-way ANOVA with Dunnett’s post hoc multiple comparison test and **f, g** Student’s *t* test. Boxes mark the median, lower-quartile, and upper-quartile, and lines connecting the extremes.

The activation of the interferon gamma pathway in NC-like DCC pointed towards an early interaction with T cells after arrival. Since extracellular vesicles (EV) are known as mediators of cell-to-cell communication and because of their concomitantly up-regulated production in NC-like DCC (brown module in Fig. 4c), we searched for candidate recipient cells of EV-transmitted signals. Performing flow cytometric analysis of sentinel and regional lymph nodes of melanoma patients we found that the number of exhausted PD1^+^TIM3^+^ CD8 T cells and CD127^-^CD25^+^ regulatory T cells (Treg) steadily increased with rising DCCD_gp100_ (*P* < 0.0001 for CD8 T cells and *P* < 0.0066 for Treg, Pearson’s correlation, Fig. 4e, f). Analysis of patients from whom we had obtained several lymph nodes of the same regional bed with both low and high DCCD_gp100_ (n = 5-9; ED Fig. 5b, c) indicated that specifically the increase of exhausted CD8 T cells was not a systemic effect but associated with local metastatic colonization. This prompted us to investigate whether DCC-derived EV affect CD8 T cell function during metastatic lymph node colonization.

### DCC-derived cell lines with NC-like phenotype secrete T cell suppressing sEV

We used cell lines that we had previously established lymph node-derived DCC (MelDCC) and that encompassed the different DCC-phenotypes (ED Fig. 6a, b). We first tested whether NC-like cells produce more small extracellular vesicles (sEV, previously called exosomes) as our pathway analysis of patient-DCC had suggested. Exposure to IFNG for 28 days induced the dedifferentiation of the plastic MelDCC 5a and MelDCC 6 lines to NC-like, NGFR-expressing cells (Fig. 5a). To enable vesicle marker-independent detection of EV and separation from contaminating non-vesicular particles in nanoparticle tracking analysis (NTA), we used the lipid membrane dye CellMaskGreen (CMG). As IFNG/BSA interfered with CMG-based NTA, sEV secretion into the culture supernatant was measured 48 h after their withdrawal (day 28-30). As controls we used MelDCC 10a and 11, as these cell lines did either not acquire a NC-like phenotype in response to IFNG stimulation (MelDCC 11) or quickly lost it (MelDCC 10a) during the IFNG-free EV production phase (Fig. 5a). NTA of culture supernatants of NC-enriched, IFNG-treated MelDCC 5a and 6 cultures clearly showed that the NC-phenotype is associated with an increased sEV-production (Fig. 5b).

**Figure 6:**
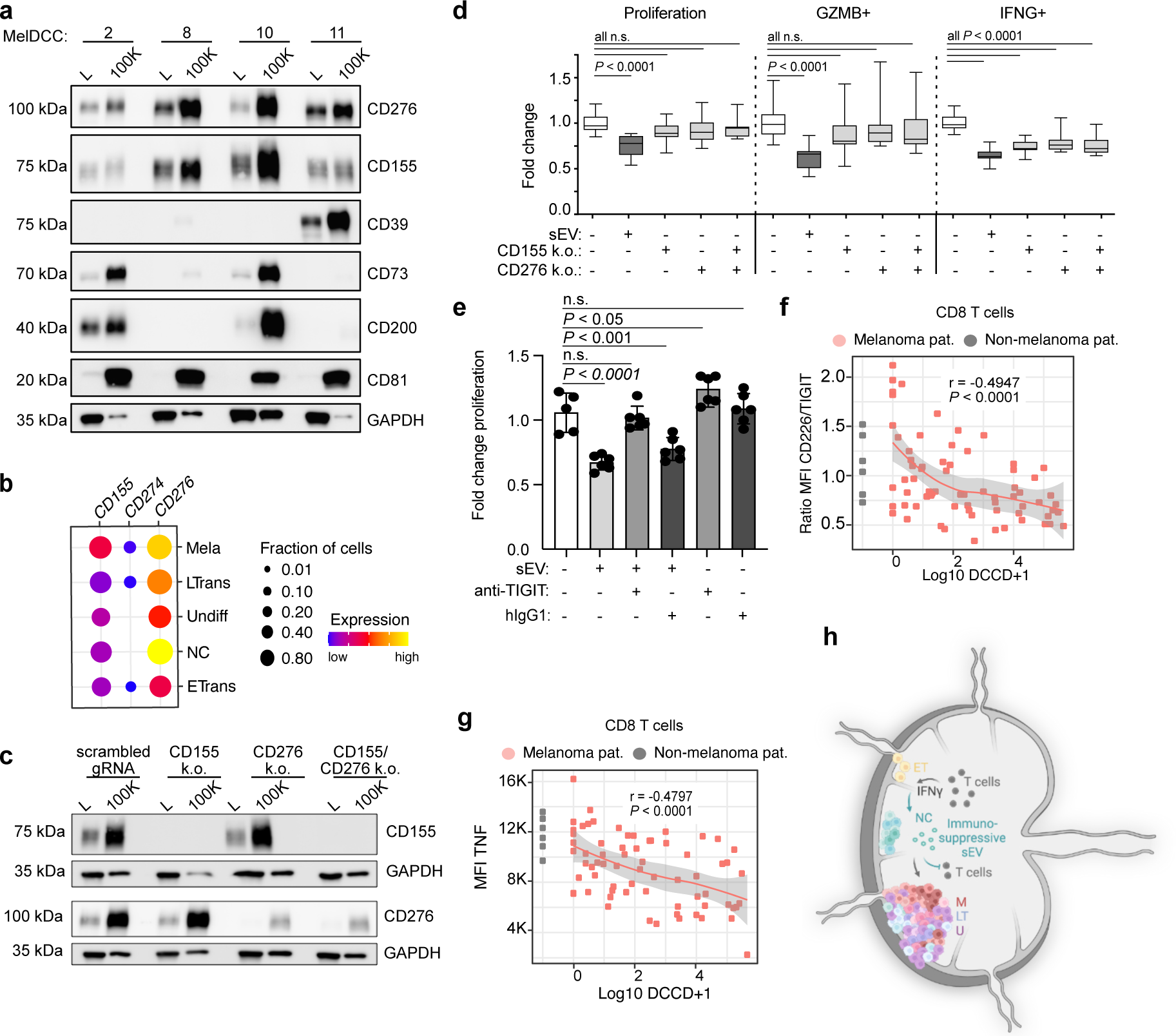
Analysis of sEV-associated immune checkpoint ligands. **a,** Western blot analysis for expression of immune checkpoint ligands in MelDCC lines (L) and their respective 100K pellets. **b,** Expression of *CD155*, *CD276* and *CD274* (PD-L1) in scRNA-seq data of DCC of melanoma patients (n = 164). **c,** Western blot analysis for CD155 and CD276 expression in MelDCC 10a controls or CD155/CD276 single or double knock-outs and respective 100K pellets. **d,** Flow cytometric analysis of CD8 T cell proliferation and percentage of IFNG^+^ and GZMB^+^ CD8 T cells at day 4 after anti-CD3/CD28 stimulation and addition of PBS or sEV of MelDCC 10a (wt EV, n = 23) and MelDCC 10a with CD155/CD276 single (CD155 k.o. n = 23, CD276 k.o., n = 12) or double knock-out (CD155/CD276 k.o., n = 11) 18 h before anti-CD3/CD28 stimulation. Shown is the pooled analysis of four experiments. **e,** Flow cytometric analysis of CD8 T cell proliferation at day 4 after anti-CD3/CD28 stimulation and addition of PBS, MelDCC 10a-sEV in the presence or absence of 10 µg anti-human TIGIT antibody or human IgG1 isotype control. **f, g,** Flow cytometric analysis of CD226/TIGIT and TNF expression in CD8 T cells (f) and PD1^+^TIM3^+/-^CD8 T cells (g), respectively, from lymph nodes of melanoma patients (n = 69) and non-melanoma patients (n = 6). Red and grey symbols indicate detected values of individual lymph nodes from melanoma and non-melanoma patients, respectively. The grey area indicates the standard error of the smoothed curve (red line). **h,** Schematic illustration of metastatic colony formation in human melanoma. Melanoma DCC switch phenotype during metastatic colonization. Early on, NC-like cells suppress CD8 T cells via immunosuppressive small extracellular vesicles (sEV). NC = neural-crest like, M = melanocytic, U = undifferentiated, ET and LT = early and late transitory phenotype, respectively; Created with BioRender.com. *P* values in **d, e** one-way ANOVA with Dunnett’s post hoc multiple comparison test, **f,** Pearson’s correlation. Boxes mark the median, lower-quartile, and upper-quartile, and lines connecting the extremes.

To test the effect of DCC-derived sEV on CD8 T cell function, we purified sEV by differential ultracentrifugation ^29^ from culture supernatant of several MelDCC lines. The pellet obtained at 100,000g (100K pellet) contained highly purified sEV, as WB revealed strong signals for the sEV markers TSG101 and CD81 and loss of GRP94, a marker for larger EV and absence of non-vesicle or exomere markers (Fig. 5c, ED Fig. 6i). Transmission electron microscopy and NTA confirmed the isolation of intact sEV at the expected size (median size 108-120 nm), production of 5.2-9.5×10^9^ sEV per million cells within 48 h, i.e. 1.8-3.2 sEV/cell/min, and presence of proteins in their membrane (Fig. 5d, ED Fig. 6c, d). To analyse the effect of sEV on CD8 T cell function, we added sEV of six MelDCC lines to healthy donor-derived polyclonal, CFSE-labeled CD8 T cells (Fig. 5e; ED Fig. 6e-g) or MART-1_27L26-35_ specific human CD8 T cells expanded from a HLA-A2 02:10 healthy donor (Fig. 5f), either 18 h before or after anti-CD3/CD28 stimulation. Small EV were added in a ratio of 10:1 or 50:1 (ED Fig. 6f), that is sEV produced by 10 and 50 MelDCC per CD8 T cell. Flow cytometric analysis at day 4 showed that sEV significantly suppressed proliferation and differentiation into IFNG/GZMB-producing cells of polyclonal and antigen-specific CD8 T cells at both time points (*P* < 0.0001 Dunnett’s post-hoc analysis Fig. 5e, f; ED Fig. 6e-g) with effector cytokine production being affected at lower sEV concentrations than proliferation (ED Fig. 6f). Moreover, sEV significantly reduced the ability of MART-1_27L26-35_ specific human CD8 T cells to eliminate antigen-loaded T2 target cells (Fig. 5g, ED Fig. 6h). Removal of proteins from the 100K pellet using size exclusion chromatography confirmed that sEV were responsible for the suppressive effect, and not any co-purified proteins or exomeres (ED Fig. 6i, j). We finally confirmed the T cell suppressive effect of sEV by adding sEV to fresh, *ex vivo*-derived human precision cut lung slices (PCLS) (n = 4 patients). Also in this setting, melanoma DCC-derived sEV significantly reduced the frequency of Ki67^+^ CD8 T cells in a dose-dependent manner (Fig. 5h).

### Identification of the immunosuppressive sEV cargo

Inhibition of CD8 T cell proliferation and effector cytokine production pointed towards the presence of inhibitory immune checkpoint ligands (ICL) in sEV as reported for CD274 (PD-L1) ^30^. We therefore mined the transcriptome data and mass spectrometry proteomics results of MelDCC lines and sEV, respectively (ED Fig. 7a, Supplementary Table 4) and tested candidate ICL by western blot (WB) analysis. Whereas CD39, CD73 and CD200 showed cell line specific expression patterns, CD155 and CD276 were detected in all MelDCC lines and sEV tested (Fig. 6a, ED Fig. 7b). Interestingly, CD274 was not detected on MelDCC lines and their sEV (ED Fig. 7c). CD274 expression could be induced by exogenous IFNG. This was, however, cell line dependent and downregulation of CD274 after 14-28 days of IFNG treatment was also observed (ED Fig. 7d). In contrast, CD155 and CD276 were expressed constitutively on all MelDCC lines and their expression was independent of exogenous IFNG stimulation (ED Fig. 7a, d). *CD155* and *CD276* were also significantly more frequently expressed than *CD274* in DCC from sentinel lymph nodes (*P* < 0.0001, Fisher’s exact test, Fig. 6b), in melanoma cell lines from primary tumour and metastasis (cancer cell encyclopedia^26^; ^31^), human epidermal melanocytes ^25^ and mouse melanoma models ^23^ (ED Fig. 7e, f). We confirmed by size-exclusion chromatography that CD155 and CD276 were sEV-associated (ED Fig. 7g) and used CRISPR/Cas technology to knock them out in MelDCC 10a (Fig. 6c, ED Fig. 7h). The single knock out of CD155 or CD276 was sufficient to significantly reduce the suppressive effect of MelDCC 10a-derived sEV on T cell proliferation, with no additive effect of the double knock-out (all *P* < 0.0001 Dunnett’s post hoc analysis, Fig. 6d). Of note, the knock-out of CD155 and/or CD276 increased the percentage of GZMB^+^ cells but had little effect on the percentage of IFNG^+^ cells indicating involvement of additional suppressive molecules or different susceptibilities to ICL inhibition among IFNG/GZMB CD8 T cell subpopulations (Fig. 6d). Next, we investigated the CD155 and CD276 topology in sEV by subjecting sEV to a mild proteinase K (PK) treatment followed by WB, using antibodies targeting the extracellular domains of CD155 and CD276. We used GAPDH and CD81, a mainly luminal protein and a common membrane-associated sEV-marker, respectively, as controls. As expected, GAPDH was protected from PK degradation, while CD81, CD155 and CD276 were sensitive to PK ^29^(ED Fig. 7i), confirming their surface-topology.

### The immunosuppressive effect correlates with DCCD

We next analysed the expression of the CD155 and CD276 receptors on healthy donor-derived CD8 T cells. We could not confirm the expression of the putative CD276 receptor IL20RA^32^ (ED Fig. 7j) arguing against involvement of intra- or extracellularly located IL20RA in CD276 mediated suppression of CD8 T cells by sEV. In contrast, 12 % and 64 % of CD8 T cells expressed the T cell immunoreceptor with Ig and ITIM domains (TIGIT) and CD226, respectively, two known receptors for CD155 on CD8 T cells (ED Fig. 7k). When we added an anti-TIGIT antibody to cultures of stimulated CD8 T cells with sEV, the suppressive effect of sEV on CD8 T cell proliferation was completely neutralized as compared to the isotype control (Fig. 6e). As TIGIT competes with the co-stimulatory receptor CD226 for binding to CD155 and binds to CD155 with higher affinity than CD226 ^33^, we tested 75 sentinel and regional lymph nodes from melanoma patients by flow cytometry for their TIGIT and CD226 expression. In line with the reported ubiquitination and proteasomal degradation of CD226 in CD8 T cells by tumour-derived CD155 ^34^, the ratio of CD226/TIGIT expression on CD8 T cells decreased with increasing DCCD (*P* < 0.0001 Pearson’s correlation, Fig. 6f). The decrease in the CD226/TIGIT ratio of CD8 T cells was mainly caused by a decrease in CD226 expression (ED Fig. 7l), most pronounced at the pre-colonizing stage (DCCD < 100, Fig. 6f) and culminated at high DCCD in PD1^+^Tim3^+^ T cells that had completely lost CD226 expression (ED Fig. 7m). Moreover, the decrease in the ratio of CD226/TIGIT expression was paralleled by a steady decline in the ability of CD8 T cells to produce TNF (*P* < 0.0001 Pearson’s correlation, Fig. 6g). These findings, together with the IFNG signature of NC-like cells and their increased secretion of immunosuppressive sEV suggest that selection of DCC by and adaptation to immune selective forces is operative in patients already at the pre-colonizing stage and is associated with the NC phenotype.

## Discussion

Here we dissected the earliest steps of metastatic colony formation in human melanoma by analysing treatment-naïve disseminated melanoma cells in SLN of melanoma patients. Our findings suggest that melanoma cells enter the lymph node of patients expressing mostly the phenotype of early-transitory melanoma ^21^ and acquire a NC-like phenotype at the start of colony formation as a result of IFNG exposure. This finding is in line with data from a recent mouse model showing that melanoma cells arrive at the lymph node with a mesenchymal phenotype, devoid of a NC-like signature, but which changes when cells invade from the subcapsular sinus into the cortical region ^23^. Apparently, the induction of the NC-like phenotype results from T cell attacks and surviving melanoma cells activate production of immunosuppressive sEV that enables them to progress to colony formation via proliferative programs of either the undifferentiated, melanocytic or late-transitory subtype (Fig. 6h). It is tempting to speculate that immunosuppression-enabled growth of first colonies leads to a dilution of IFNG which further boosts phenotype switching from NC-like to one of the others. Thus, we found a highly dynamic and microenvironmentally-induced plasticity of metastasis-founding melanoma cells. The high plasticity of melanoma cells has been observed under various conditions repeatedly ^35–37^, but its dynamic nature in patients at the earliest steps of metastasis formation at the secondary site was not known so far.

To uncover it, we started a scrupulous search for candidate metastasis founder cells (MFC). Twenty years ago, we had begun to compare gp100, S100, Melan A and MCSP for their ability to detect early DCC and their impact on survival of patients and found that gp100 generated the most accurate prognostic information as single marker ^11,12^. Lack of specificity made MCSP unsuitable as single marker, but addition of melanoma-associated transcripts (either *PMEL*, *MLANA* or *DCT*), defining MCSP^+^MT^+^ cells, outperformed the gp100 assay regarding clinical impact. In fact, even in the best prognostic group of gp100-negative patients, detection of single MCSP^+^MT^+^ cells puts the patients at high risk to die from melanoma, indicating that we identified a marker combination for metastasis founder cells. It is notable that MCSP expression peaks in human development at fetal week 18 and is preserved over all subtypes in human and murine melanoma ^23,25^. Which of the multiple functions of MCSP identified so far equips the cells with MFC potential remains to be elucidated ^13^.

Having identified MFC, we could quantitatively track them over metastasis formation via the DCCD that is highly correlated with microscopic presentation of lymph node invading melanoma cells from single cells over micro- to macroscopic colony formation ^2,11^. In addition to the acquisition of genetic alterations ^2^, we now found that melanoma phenotypic subtypes and the immune response significantly correlate with the DCCD, further emphasizing the dynamic interdependencies at early colonization. The predominant phenotype at the transition from single invading DCC to micrometastasis was characterized by a NC-like gene expression program that also reflected encounter with immune cells, such as an IFNG signature, and showed activation of the exosomal pathway. We provide evidence that at least CD8 T cells are targeted by the sEV cargo and identified CD155 and CD276, but not CD274 (PD-L1), as sEV-associated immunosuppressors. The relevance of CD155-mediated suppression for early metastatic colonization was demonstrated by the fact that the decrease in the ratio of CD226/TIGIT expression on CD8 T cells (as a marker of suppression) is most pronounced at the pre-colonizing stage and preceded the accumulation of exhausted CD8 T cells.

In summary, we have identified the strongest candidate for metastasis founder cells in melanoma and their early immune interactions, providing a foundation for advancing melanoma immunotherapy. Targeting MCSP could be an optimal strategy for eliminating metastasis founder cells. First, it has been found in our study and by others to be expressed in all phenotypes forming nascent metastases and therefore likely to be expressed by MFC in organs other than lymph node. Second, by-passing immunosuppression by MCSP-directed CAR-T cells may be particularly promising. Additionally, our study underscores the potential significance of CD155 and CD276 as superior or supplementary targets compared to PD-L1/PD-1 in the context of adjuvant immune checkpoint blockade.

## Online Methods

### Patient material

Human disseminated cancer cells were obtained from sentinel or regional lymph nodes of melanoma patients. Skin draining control lymph nodes were obtained from non-melanoma patients. Written informed consent of melanoma and control patients was obtained and the ethics committee of the University of Regensburg (ethics vote 07-079 and 18-948-101) approved lymph node sampling and analysis of isolated cells. Human peripheral blood mononuclear cells were obtained from a healthy donor (ethics vote 18-948-101) and human tumour-free lung samples from patients with lung cancer (ethics vote 2701-2015, ethic committee Medical School Hannover).

### Cell lines

MelDCC lines were established from DCC-derived xenografts ^2^ or DCC directly propagated in vitro (manuscript in preparation). Their patient origin was verified by STR analysis (Cell-ID™, Promega), their melanoma origin by a human pathologist and their aberrant genotype by CGH. MelDCC lines, NCI-H1975 and HeLa cells were propagated in RPMI 1640 medium supplemented with 10 % fetal bovine serum, 2 mM L-glutamine and 1 % penicillin/streptomycin (P/S) (all Pan-Biotech). Adult human epidermal melanocytes (Lonza) were cultured in MGM-4 melanocyte growth media including supplements (CaCl_2_, PMA, hFGF, Insulin, BPE, Hydrocortisone, FBS, GA-1000) (all Lonza) and 1 % P/S (Pan-Biotech) and used only at passage 1 for scRNA-seq. All cell lines were kept at 37 °C and 5 % CO_2_ in a fully humidified incubator and negatively tested for mycoplasma by PCR.

### Lymph node disaggregation and immunocytology

Quantitative immunocytology was performed as described ^2,11,12^ after SLN biopsy or removal of regional lymph nodes using unfixed SLN tissue. Immunocytological staining was carried out using primary antibodies against gp100 (clone HMB45, Agilent) or MCSP (melanoma chondroitin sulphate proteoglycan, clone 9.2.27, BD Biosciences). A lymph node was defined as gp100- or MCSP-positive if it contained at least one gp100- or MCSP-positive cell. The number of positive cells per million lymphocytes (DCCD, disseminated cancer cell density) was recorded as DCCD_gp100_ or DCCD_MCSP_. MCSP+ cells were isolated with a micromanipulator (Eppendorf PatchMan NP2, Eppendorf) and cells were subjected to whole transcriptome amplification (WTA) to isolate RNA and whole genome amplification (WGA) for isolation of genomic DNA. Images of MCSP^+^ cells were acquired on an Axiovert 200M (Zeiss) or IX81 (Olympus) microscope.

Combined immunofluorescent staining of SLN cell suspensions on adhesion slides for MCSP, MelanA and CD74 was carried out as follows: Unspecific binding of antibodies was blocked with 300 µl/spot of Human TruStain FcX (Biolegend, 1:20) in 5 % BSA/PBS for 30 min at RT. Blocking buffer was discarded and slides were stained with an unconjugated anti-human MCSP antibody (clone LHM2, Abcam,1 mg/ml, 1:176 dilution) in 5 % BSA/PBS with 150 µl/spot for 60 min at RT. Slides were washed 3 times for 3 min with PBS before a goat-anti mouse IgG1 AF546 antibody (Invitrogen, 2 mg/ml, 1:150 dilution) in 5 % BSA/PBS with 150 µl/spot was applied for 30 min at RT. Slides were washed 3 times for 3 min with PBS and free binding sites of the goat-anti mouse IgG1 AF546 antibody were blocked with 300 µl/spot of 5 % mouse serum (DAKO) in PBS. Slide were then stained with an anti-human MelanA biotinylated antibody (clone A-103, Invitrogen, 0.1 mg/ml, 1:100 dilution) and anti-human CD74 APC antibody (clone LN2, Biolegend, 0.15 mg/ml, 1:20 dilution) in 5 % BSA/PBS with 150 µl/spot for 60 min at RT. Slides were washed 3 times for 3 min with PBS before staining with Streptavidin-AF488 (Invitrogen, 2 mg/ml, 1:250 dilution) in 5 % BSA/PBS with 150 µl/spot for 30 min at RT. Slides were washed 3 times for 3 min with PBS and stained with DAPI (Roche, 5 mg/ml, 1:25.000 dilution) in PBS with 150 µl/spot for 10 min at RT. Slides were washed 2 times for 2 min with PBS and fixed with 150 µl/spot 1% formaldehyde in PBS for 5 min at RT. Slides were washed 3 times for 3 min with PBS and stored at 4°C until image analysis. Images were acquired at a IX81 (Olympus) microscope with identical settings for all samples, fluorescence intensity quantification of MelanA and CD74 was performed with ImageJ/Fiji.

### Patient-Survival analysis

Kaplan-Meier survival curves were created with survfit function in survival_3.3-1^38^. For univariable proportional hazard model, coxph function from survival_3.3-1 was applied to each known patient related feature including age, gender, MT status, log10(DCCD_MCSP_), log10(DCCD_gp100_), N status, thickness and ulceration. Further, features were selected with selectCox function from pec_2022.05.04 for multivariable proportional hazard model. Only the selected features are included in the final multivariable proportional models, which are visualized with ggforest function from survminer_0.4.9. For cox regression models, proportional assumption is checked with cox.zph function from survival_3.3.1.

### Whole genome amplification and analysis of copy number alterations

Single-cell genomic DNA was collected during WTA procedure by precipitation and subjected to whole genome amplification (WGA), using the previously described ^15,39^ or the commercially available Ampli1^TM^ WGA Kit (Ampli1^TM^ WGA, Menarini Silicon Biosystems). CNA analysis was performed using the Ampli^TM^ LowPass kit (Menarini Silicon Biosystems) according to the manufacturer’s instructions. The Ampli1^TM^ LowPass libraries were sequenced in single-end mode on a MiSeq device using the MiSeq Reagent Kit v3 (150 cycles) (Illumina) or on the NovaSeq6000 (Cegat). Genomic coordinates analysed with the LowPass bioinformatics analysis pipeline (Menarini Silicon Biosystems) or HIENA (Fraunhofer ITEM) were submitted to the Progenetix (v4.0, 2022) user data tool ^40^ to generate cumulative frequency plots.

### Whole transcriptome amplification (WTA)

Isolation of mRNA from a single cell, reverse transcription, and global amplification of the first strand cDNA was carried out as previously described ^14,15^. The quality of WTA products was assessed by expression analysis of three housekeeping genes as previously described ^4^.

### Marker expression analysis in single cells

Endpoint PCR for specific transcripts was carried out on all WTA products as previously described ^41^. Primers used were:

*PMEL* (gp100) 5’-TCCAAAGTCCCAGGTGTAGG-3’ and 3 ’-CCTCTTGCTCATTCCAGCTC-5’, *DCT* (dopachrome tautomerase) 5 ’-CCAGCTGGGAAACTGTCTGT-3’ and 3 ’-AACCCTTCCAAAGCATTCCT-5’, *MLANA* (melan A) 5 ’-ATAAGCAGGTGGAGCATTGG-3’ and 3 ’-GCTCATCGGCTGTTGGTATT-5’ and *PTPRC* (CD45) 5’-TTAGGGACACGGCTGACTTC-3’ and 3’-ATAAACACTGTCCCGTTTCG-5’.

### NGS mRNA library preparation and sequencing

Single cell mRNA-seq: NGS mRNA library preparation of MCSP^+^ cells from melanoma and non-melanoma patients and human epidermal melanocytes (Lonza) was performed as previously described ^4^. Resulting libraries were quantified with the MiSeq System using MiSeq Reagent Kit v2 (50-cycles) (Illumina^®^), pooled in equal molar ratios and sequenced on Illumina NovaSeq6000 platforms. 1-2 million MelDCC cells were harvested using the RNeasy mini kit (Qiagen) according to the manufacturer’s instructions. RNA-seq libraries were generated using the TruSeq Stranded mRNA Library Prep Kit (Illumina) according to the manufacturer’s instructions. Libraries were sequenced single-end (SE-82-10) on a NextSeq550 at the NGS Core Unit of the Leibniz-Institute for Immunotherapy, Regensburg and University Medical Center Regensburg, Germany).

### scRNA-seq data analysis

FastQC_0.11.5 ^42^ and MultiQC_1.12 ^43^ were used to evaluate sequence quality of the samples. After trimming with BBDuk ^44^ reads were mapped to human reference genome GRCh38 with gene annotation GRCh38.96 using STAR_2.6.1c ^45^. Expected gene counts were calculated with RSEM_1.3.1 ^46^. Cells with less than 50,000 counts, mitochondrial gene counts of more than 70 % or less than 1,000 expressed genes were excluded, and only genes expressed in at least 3 cells were kept. The Seurat_4.1.0 ^47^ pipeline was used to process the expression data of single cells: The NormalizeData and FindVariableFeatures functions were used to normalize expression and find the top 2,000 most variable genes. Considering the relative low number of cells, the number of dimensions resulting from RunPCA was selected carefully according to Elbowplot and JackStrawplot. Further testing of other dimensions (5, 10, 15, 20, 25, 30) revealed that 20 was the minimal number of dimensions needed to separate melanocytes from DCC. RunUMAP was then run on the top 20 components of PCA with parameters umap.method = “umap-learn”, n.epochs = 1000, min.dist 0.1 and n.neighbors = 15. Clusters were calculated by FindNeighbors with reduction = “umap”, k.param=10, dims = 1:2 and FindClusters with resolution = 0.2 based on UMAP.

To test the robustness and consistency of clusters, the clusterMany function in clusterExperiment (version 2.12.0) ^48^ was used with parameters clusterFunction = c (“pam”, “clara”, “kmeans”, “spectral”, “hierarchicalK”, “tight”), ks = 3:10, minSizes = 3, isCount = F, reduceMethod = c (“PCA”), nReducedDims = c (5, 10, 15, 20, 25, 30) to get 288 individual clusterings for DCC. The proportion of times that pairs of cells clustered together, i.e. the consensus, was calculated. For each Seurat cluster, the consensus of cell pairs from the same cluster and across different clusters was checked. Bluster_1.2.1 was used to further confirm the modularity of clusters based on both UMAP and PCA results. FindAllMarkers was used to detect cluster specific markers using “MAST” ^49^. Upregulated genes with *P* < 0.01 were used as cluster markers and input for Enrichr.

To infer subtypes of melanoma DCC, gene expression signature scores were calculated with AUCell ^50^ with aucMaxRank = 0.3, and rescaled to range [0,1] with rescale from R package scales_1.2.0. To infer pseudotime and cell trajectory, slingshot_2.0.0 ^28^ was run with UMAP dimension reduction and Seurat clusters. The cluster with the lowest DCCD level (cluster 0) was set as start cluster. To confirm the trajectories inferred by slingshot, an Elastic structure ^51^ was constructed using the function computeElasticPrincipalTree from ElPiGraph.R(version 1.0.0) and UMAP with parameter NumNodes = 6, followed by the function GetSubGraph with parameter Structure = ‘end2end’. NumNodes = 6 is the minimal number that can bind all Seurat clusters in one tree. The node with lowest DCCD level was set to be the tree root. For each lineage retrieved, pseudotime was calculated with the function getPseudotime.

For visualization, signature score lines were plotted with the geom_smooth function with method=loess and span = 1 from ggplot2_3.3.6. To check how signatures changed with pseudotime in each lineage, gam function from gam_1.20.1 ^52^ was used to fit the generalized additive model with loess smooth. ANOVA *P* values for the parametric effect and nonparametric effect were extracted and the minimum of the two was used to test if signature scores changed with pseudotime in each lineage for both slingshot and ElPiGraph.

scWGCNA_0.0.0.9000 ^41^ and WGCNA_1.70-3 ^42^ were used to generate the weighted correlation network on the top 1,000 most variant genes. Meta cell for each cluster were constructed with the function construct_metacells with parameters k = 5, reduction = ‘pca’. Thereafter, weighted gene co-expression network was constructed with the blockwiseConsensusModules function from WGCNA_1.70-3 ^53^ with parameters consensusQuantile = 0.3, power = 10, networkType = signed, mergeCutHeight = 0.2, and minModuleSiz e= 50. Power = 10 was chosen according to the result of function pickSoftThereshold. The respective gene modules were obtained accordingly.

To check the preservation of the modules, the function modulePreservation from WGCNA_1.70-3 was used with resampling of all samples with replacement and with the downsampled half of the samples without replacement. Both results showed that detected modules were conserved, whereas random modules and the genes that could not be included in a particular module showed low conservation. Only modules with an ANOVA *P* value < 0.001 between clusters were processed for pathway enrichment analysis Enrichr ^54^ based on Gene Ontology 2015 and MSIGDB_HALLMARK 2021 ^55^ databases.

CONICsmat 0.0.0.1 ^56^ was used to infer copy number alterations from the gene expression of the chromosome arm levels. In addition to cells already contained in the melanocyte cluster and lymph node cluster, 200 normal melanocyte cells ^57^ were included in the analysis as control cells. Cells were separated into two clusters based on posterior expression using the plotHistogram function with the “ward” clustering method. Chromosome arms with a BIC (Bayesian information criterion) difference larger than 70 and adjusted *P* value < 0.01 were kept as candidate areas with CNA. Posterior probabilities of CNA were binarized on specific chromosome arms using the binarizeMatrix function with parameter threshold = 0.9.

SCENTIC 1.2.4 was used to infer transcript factor regulatory networks with GENIE3 and the motif database from: https://resources.aertslab.org/cistarget/databases/old/homo_sapiens/hg19/refseq_r45/ mc9nr/gene_based/hg19-tss-centered-10kb-7species.mc9nr.feather. Only regulatons with highest score > 0.3, and in at least 5 cells that scored larger than 0.1, were kept. Differentially activated transcription factors for each cluster were identified using pair-wise-t test and filtered with largest BH adjust p value < 0.05.

Skin and lymph node tissue cells with cell type annotation were downloaded from the human cell atlas ^58^. Cell types with less than 50 cells were removed to reduce the noise in the cell type annotation. To integrate data sets, integration anchors were computed following SCTransform normalization, followed by IntegrateData. The integrated data was transformed into PCA with RunPCA and using RunUMAP with parameter reduction = “pca”, dims = 1:30.

### Bulk RNA-seq data analysis of MelDCC

Gene counts were calculated using the same methods as for scRNA-seq. Gene expression was normalized with log counts per million. GSVA ^59^ with the marker gene set for each single cell melanoma DCC cluster (cluster 0 to cluster 4) was used to get the DCC cluster signature scores for every MelDCC-line.

### Isolation of sEV by differential ultracentrifugation

2×10^6^ MelDCC were seeded in T175 cell culture flasks and cultured in 25 ml medium (RPMI 1640, 10 % fetal bovine serum, 2 mM L-glutamine, 1 % P/S, all Pan-Biotech). After 3 days medium was removed, cells were washed with PBS and 30 ml of FBS-free medium (RPMI 1640, 2 mM L-glutamine, 1 % P/S) was added. The conditioned medium was collected after 48 h and subjected to differential ultracentrifugation as previously described ^29^. Briefly, conditioned medium was centrifuged at 300g for 10 min at 4 °C to remove dead cells and debris. The supernatant was centrifuged at 2,000g for 20 min at 4 °C (2K pellet) and then centrifuged at 10,000g (10K pellet) for 40 min at 4 °C. Finally, the supernatant was centrifuged at 100,000g (100K pellet) for 90 min at 4 °C. All pellets were pooled and washed in 35-50 ml PBS and recentrifuged at the same speed they were initially harvested before resuspension in 1 µl PBS/1×10^6^ EV-producing cells at the time of supernatant-harvesting.

### Size exclusion chromatography

100K pellets obtained by differential ultracentrifugation were pooled before the last washing step and loaded onto a SEC Column (Plastic XXL Column filled with 45 ml 2 % BCL Agarose Bead Standard (50-150 µm) and a Plastic XXL Column Frit on top, all Agarose Bead Technologies). The column was attached to a quartz flow cell inserted into a Zetasizer Nano ZS (Malvern Panalytical) allowing particle detection in real time and thus determination of the boundary between the vesicle- and protein-fraction. The vesicle fraction was concentrated by centrifugation at 100,000g for 90 min at 4 °C. The protein fraction was concentrated by centrifugal ultrafiltration, first with a Macrosep Advance Centrifugal Device with Omega Membrane 3K (Pall) at 3,200g for 30 min at 4 °C and then with a Vivaspin 500, 5,000 MWCO PES (Sartorius) to a low volume suited for further use. The vesicle and protein fraction were characterized by WB and used in CD8 T cell assays.

### Label-Free Mass Spectrometry-based Proteomics

100K EV of MelDCC 10a in duplicates were resuspended in 50 µl GASP-buffer (50 mM Tris-HCl, pH 8.8, 6 M urea, 1.5 M thiourea, 4 % SDS) and sonicated for 15 Cycles 60/30 with the BioRuptor pico (Diagenode). Cell debris was separated by centrifugation 20,000g for 15 min and at 4 °C and proteins were quantified with the SERVA Purple Protein Quantification Assay (SERVA Electrophoresis GmbH) according to the manufacturer’s instructions. Sample preparation for LC-MS was done according to the GASP protocol ^60^ with slight modifications ^61^. Two micrograms of the resulting peptide mixtures were spiked with 100 fmol of the retention time standard RePLiCal (Polyquant GmbH) and analysed on the Eksigent ekspert™ nanoLC 400 system coupled to a TripleTOF 5600^+^ mass spectrometer (SCIEX). Samples were loaded onto a YMC-Triart C18 trap column (3 μm particle size, 0.5 cm length; YMC America, Inc.) at a flow-rate of 10 µL·min^−1^ for 5 min (isocratic conditions A: 0.1 % formic acid, 0.1 % ACN). Peptides were then separated on a reverse-phase column (YMC-Triart C18, 1.9 µm particle size, 120 Å, flow-rate of 5 µL·min^−1^, 40 °C) using a 94-min binary acetonitrile gradient (3–40 % B in 87 min, 40–45 % B in 7 min). Duplicate samples were run twice.

The sequential window acquisition of all theoretical fragment-ion spectra (SWATH) runs was accomplished using a 50 ms full-MS scan from 400–1,000 *m/z* and 60 subsequent SWATH windows of variable size for 35 ms each (mass range, 230–1,500 *m/z*). The respective libraries were generated from pooled samples measured in data-dependent acquisitions mode (DDA) using the TOP20 method with a full-MS scan for 250 ms and MS/MS scans for 50 ms each. MS/MS spectra from the DDA runs were searched against the respective UniProt database (swissprot-human 12-2021) using ProteinPilot 5.0 and imported in PeakView 2.1 using the SWATH MicroApp 2.0 allowing six peptides per protein and six transitions per peptide. Raw values were normalized to total intensity.

### Immuno-(western) blotting

WB was performed as previously described (Werner-Klein 2020) with minor modifications: Protein (cell lysates) or sEV were denatured by incubation for 5 min at 95 °C in the presence of 1X Leammli Buffer (Bio-Rad) containing 10 % 2-Mercaptoethanol (Sigma-Aldrich) and loaded on 10 % or 4-20 % Mini-PROTEAN^®^ TGX Stain-Free™ Protein Gels (Bio-Rad). Incubation of blotted PVDF membranes was performed over night with antibodies directed to: CD81 (clone B-11, 0.2 mg/ml, 1:10,000 dilution), GAPDH (clone 6C5, 0.1 mg/ml, 1:1,000 dilution), IL20Rα (clone EE09, 0.1 mg/ml, 1:500 dilution) all Santa Cruz Biotechnology, Calnexin (clone 37/Calnexin, 0.25 mg/ml, 1:1000 dilution), HSP70 (clone 7/Hsp70, 0.25 mg/ml, 1:1,000 dilution), TSG101 (clone 51/TSG101, 0.25 mg/ml, 1:1,000 dilution) both BD Biosciences, CD39 (clone EPR20627, 0.613 mg/ml, 1:1,000 dilution), CD73 (clone EPR6114, 2.235 mg/ml 1:1,000 dilution), CD200 (clone EPR22412-229, 0.507 mg/ml, 1:1,000 dilution), CD276 (clone EPR20115, 0.516 mg/ml, 1:5,000 dilution), PD-L1 (clone EPR19759, 0.443 mg/ml, 1:1,000 dilution), CD155 (clone EPR17302, 0.151 mg/ml, 1:2,000 dilution) all Abcam, GRP94 (clone B-11, 1 mg/ml, 1:1,000 dilution) Enzo Life Sciences, Albumin (clone J32-10, 1 mg/ml, 1:1,000 dilution) Invitrogen, ACLY (polyclonal rabbit, 0.037 mg/ml, 1:1,000 dilution), Fibronectin (clone E5H6X, 0.1 mg/ml, 1:1,000 dilution), and Histone H2A (polyclonal rabbit, 0.01 mg/ml, 1:1,000 dilution) all from Cell Signaling. After washing, the blots were incubated with HRP-conjugated anti-mouse, anti-rat or anti-rabbit IgG secondary antibody (all Sigma Aldrich) at a dilution of 1:10,000 for 2 h at RT. Protein bands were visualized using SuperSignal^TM^ West Pico PLUS Chemiluminescent Substrate (Thermo Fisher Scientific). Chemiluminescence was recorded by a ChemiDoc^TM^ MP Imaging System (Bio-Rad) and analysed with Image Lab^TM^ (version 6.1, Bio-Rad).

### NTA measurement of sEV

For detection of CD81-positive sEV, 1 µl 100K pellet and 1 µl anti-human CD81 PE/Dazzle 594 (clone 5A6, Biolegend) were incubated for 30 min at RT in PBS (Gibco) in a total volume of 10 µl and then diluted 1:10,000 in PBS in a final volume of 10 ml. For enumeration of sEV in the culture-supernatant of MelDCC lines, cells were incubated in EV-production medium, i.e. phenolred and serum-free RPMI (Gibco) supplemented with 2 mM L-glutamine and 1 % (P/S) (all Pan-Biotech). 48 h after exchange to EV-production medium, the culture supernatant was harvested, centrifuged at 300g for 10 min and filtered (0.22 µm). sEV were stained with 1 µl of a 1:10 dilution of CellMaskGreen Plasma Membrane Stain (Thermo Fisher Scientific) in PBS in a total volume of 100 µl for 1 h at RT and then diluted 1:25 to 30 in PBS in a final volume of 1 ml. For EV analysis, the manufacturer’s default software settings were selected accordingly: For each measurement, one cycle was performed by scanning 11 cell positions and capturing 30 frames per position. In the scatter mode a sensitivity of 80 and a trace length of 15 was chosen, whereas for the measurement of CD81- or CMG-stained particles with the 500 nm LP fluorescence filter a sensitivity of 96 and a trace length of 7 or 10 was used. Data analysis was performed with built-in ZetaView Software version 8.05.14 SP7 and FlowJo version 10.8.1.

### Transmission electron microscopy (TEM) imaging of EV

For negative staining, EV samples were isolated from MelDCC line cells by differential ultracentrifugation and diluted 1:10 using HEPES buffered saline (pH 7.4) and 4 µL of this suspension was applied onto hydrophilized, carbon-coated grids (400 mesh, Cu; Plano) for 30 s, blotted with filter paper and stained with a 1 % Uranyl acetate solution for 30 s, before final blotting and air-drying. Undiluted EV samples were also prepared for freeze-etching as described earlier ^62^ including unidirectional Platinum/carbon shadowing (angle 45°; 1.5 nm) and carbon coating (angle: 90°; 15 nm). Screening and imaging of samples were done using a 200 kV field-emitter transmission electron microscope (JEM-2100F; JEOL GmbH) equipped with a 16 megapixel CMOS camera (F416; TVIPS GmbH). Full control of the microscope and imaging parameters was provided using the SerialEM software package ^63^. Images were recorded at various magnifications (2,000 x up to 40,000 x), resulting in pixel sizes of about 5.5 nm down to 0.28 nm.

### Topology assessment of EV-associated immune checkpoint ligands

sEV of 2.5×10^6^ or 0.5×10^6^ MelDCC/western blot lane for the analysis of CD155 or CD276 and CD81 topology were incubated for 1 or 2 h at 37 °C in 50mM Tris-HCl pH 8.0 with 5 mM CaCl_2_ (Sigma-Aldrich) in the presence or absence of 0.125 mg/ml Proteinase K (Promega or Roche) with or without 1 % Triton X-100 (Sigma-Aldrich). The proteinase activity was inhibited by incubation with by 1 mM phenylmethylsulfonyl (Sigma-Aldrich) for 10 minutes at RT before WB against CD155, CD276, CD81 and GAPDH was performed.

### CRIPSR/Cas9 genetic engineering

Knock-down of CD155 and CD276 in MelDCC10 was performed with 1.5 μM Cas9 protein (Alt-R® S.p. HiFi Cas9 Nuclease V3), 1.8 μM tracrRNA (Alt-R CRISPR-Cas9 tracrRNA), 1.8 μM CD276 AD crRNA (Hs.Cas9.CD276.1.AD) or PVR AB crRNA (Hs.Cas9.PVR.1.AB), or control crRNA, 1.8 μM Alt-R Cas9 electroporation enhancer (all IDT) and 100,000 cells per transfection. Transfection was performed using the NEON transfection instrument (Thermo Fisher, settings: 1600 V, 10 ms pulse width, 3 pulses).

### Induction of dedifferentiation in MelDCC lines

Lyophilized IFNG (Peprotech) was dissolved at a concentration of 200.000 U/ml in 0.1 % bovine serum albumin (BSA) in PBS. 1×10^6^ MelDCC were seeded in T25 flasks one day before 500 U IFNG (Peprotech) was added to the culture medium. IFNG-containing medium was exchanged every 2-3 days for a total of 28 days. On day 28, the medium was replaced with EV-production medium, i.e. phenolred and serum-free RPMI (Gibco) supplemented with 2 mM L-glutamine and 1 % P/S (all Pan-Biotech). Control cells were plated with 400,000 cells/T25 flask and medium was exchanged to EV-production medium after 48 h. After another 48 h, the culture supernatant of IFNG-treated and control cells was harvested, centrifuged at 300g for 10 min and then passed over a 0.22 µm filter before NTA was performed. In addition, flow cytometric analysis of seeded MelDCC (d0) and at day 28 and day 30 of IFNG-treatment was performed and the cell number of IFNG-treated/control cells at time of supernatant harvest recorded.

### CD8 T cell isolation

Peripheral blood mononuclear cells were prepared by a density gradient centrifugation (60 % Percoll solution, GE Healthcare) and cryoconserved in liquid nitrogen. Mononuclear cells were thawed in the presence of 100 µg/ml DNAse I (Roche) and rested overnight with 2×10^6^ cells/well in a 96-well plate in 200 µl RPMI 1640 medium supplemented with 10 % fetal bovine serum, 2 mM L-glutamine, 1 % P/S (all Pan-Biotech), 100 mM HEPES (Sigma Aldrich), and 0.1 % 2-Mercaptoethanol (Thermo Fisher). The next day, cells were pooled and CD8 T cells were isolated using the MojoSort Human CD8 T Cell Isolation Kit (Biolegend) according to the manufacturer’s instruction with the modification that LS columns (Miltenyi Biotec) and MACS-buffer (PBS (Thermo Fisher), 0.5 % BSA (Roche), 2 mM EDTA (Thermo Fisher) were used.

### Testing of CD8 T cell inhibitory function of sEV

Polyclonal or MART-1_27L26-35_ specific CD8 T cells were labeled with 2 µM CFSE (eBioscience) for 10 min at 37 °C in PBS/1 % FBS. The reaction was stopped with 10 % FBS containing medium. 1×10^5^ CFSE-labeled CD8 T cells/well were stimulated on 96 well plates coated with 2 µg/ml anti-CD3 (OKT3) and 2 µg/ml CD28 (CD28.2) antibodies (both Biolegend) in 200 µl PBS and overnight at 4 °C. sEV (wildtype or with CD155 and/or CD276 knock-out) were added either 18 h before or after anti-CD3/CD28 stimulation in a total volume of 200 µl and a ratio of 50:1, if not indicated otherwise, that is sEV produced by 50 MelDCC line cells/CD8 T cell. To block CD155/TIGIT interaction, anti-TIGIT Neutralizing Antibody (human monoclonal, BPS Bioscience, catalogue number 71340) or isotype control (QA16A12, Biolegend) was added at a final concentration of 10 µg/ml. At day 4, cultures were re-stimulated for 4 h with 1x Cell Stimulation Cocktail that included protein transport inhibitors (eBioscience). Proliferation as function of CFSE-dilution and production of the effector cytokines INFG and GZMB were assessed by flow cytometry.

### In vitro cytotoxicity assay

This assay is based on the specific *in vitro* elimination of MART-1_27L26-35_ loaded T2 cells labeled with CFSE (ebioscience) over unloaded T2 cells, which were labeled with the CellTrace Violet Cell Proliferation Kit (ebioscience). To generate antigen-specific CD8 T cells, CD8 T cells of a HLA A02:01 healthy donor were expanded using artificial APC loaded with MART-1_27L26-35_ as previously described ^64^. To test the MART-1_27L26-35_ specific cytotoxic activity, MART-1_27L26-35_ specific CD8 T cells were exposed to PBS or sEV at a ratio of 50:1, i.e. sEV isolated from 50 MelDCC 10a cells per CD8 T cell, prior to stimulation on anti-CD3/CD28 coated plates (2 µg/ml). At day 4, CD8 T cells were harvested and added in a 1:1 ratio to 25,000 target cells on a round-bottom 96 well plate. Target cells were loaded with 10 µg/ml MART-1_27L26-35_ peptide for 1 h at 37 °C and 5 % CO_2_. To remove unbound peptide, the cells were washed two times with PBS. Peptide-loaded cells were labeled with 2 µM CFSE whereas unloaded cells were labeled with 2 µM CellTrace Violet. Equal numbers of CFSE and CellTrace Violet labeled T2 cells were mixed and CD8 T cells were added. After 24 h, target cells were analysed by flow cytometry. The percentage of MART-1_27L26-35_-specific cytotoxic lysis was calculated as follows: percent-specific cytotoxic lysis = (1 - (r with T cells/r without T cells) × 100); r = (% CFSE^+^/% CellTrace Violet^+^).

### Precision Cut Lung Slices

Precision cut lung slices were generated from tumour distant tissue after tumour resections as described previously ^65^. In short, lung lobes were filled with 4 % low melting agarose, punched into 8 mm cores and sliced into 300 µm thick slices using a Krumdiek Tissue Slicer (Alamaba R&D). PCLS were cultured in 48 well plates with one PCLS/well in 250 µL DMEM F/12 and treated with anti-CD3/CD28 beads 18 hours prior to addition of sEV or 5 µM pimecrolimus. Flow cytometric analysis was performed 6 days post treatment. For Flowcytometric analysis 6 PCLS per condition were dissociated pre- and post-treatment with sEV.

### Flow cytometry

Single cell suspensions of lymph nodes or PCLS, CD8/EV co-cultures or MelDCC-lines were incubated for 5 min at 4 °C with PBS/10 % AB-serum (Bio-Rad) or human TruStain FcX receptor blocking solution (Biolegend) to reduce non-specific antibody binding, stained with fluorescence-labeled antibodies for 30 min at 4 °C and washed once with PBS/2 % FBS/0.01 % NaN_3_. For intracellular cytokine staining, cells were fixed for 20 min at RT with FluoroFix (Biolegend) and permeabilized with Intracellular Staining Permeabilization Wash Buffer (Biolegend) according to the manufacturer’s instructions. Intranuclear staining was conducted with the Foxp3/Transcription Factor Staining Buffer Set (Thermo Fisher) according to the manufacturer’s instructions. Cells were stained for 30 min using the following antibodies: anti-human CD45 FITC, AF488 or PerCP (HI30), anti-human CD3 AF700 (SK7 or UCHT1), anti-human CD3 PerCP/Cyanine5.5 (SK7 or HIT3a), anti-human CD4 BV650 (RPA-T4) or AF700 (SK3), anti-human CD8 AF700 (HIT8α) or BV510 and BV 421(RPA-T8), anti-human CD45RA BV421 (HI100), anti-human CD127 PE-Dazzle (A019D5), anti-human CD197 PE/Cy7 (G043H7), anti-human Ki-67 APC (Ki-67), anti-human IFNG PE (4S.B3), anti-human GZMB AF647 (GB11), anti-human CD226 PE/Cy7 (DNAM-1), anti-human TIGIT BV421 (VSTM3), anti-human TIM-3 (CD366) BV510, anti-human PD-1 BV711 or BV 421 (EH12.2H7), anti-human CD155 PE or PE/Cy 7 (SKII.4), anti-human CD271 (ME20.4) PE, anti-human CD271 (PD-L1; 29E.2A3), BV711 anti-human CD276 (B7-H3) APC (MIH42) all from Biolegend; anti-human CD8 BUV805 (HIT8α), anti-human TIM-3 (CD366) BUV615 (7D3), anti-human TNF BUV395 (Mab11) all from BD Biosciences; anti-human CD276 PerCP-eFluor710 (7-517, Thermo Fisher Scientific), anti-human MelanA AF647 (A103, Santa Cruz) and anti-MCSP (9.2.27, BD Biosciences). Unspecific binding was controlled by staining with the following Isotype controls: mouse IgG1, κ BUV395 (clone X40) and mouse IgG1, κ BUV615 (clone X40) from BD Biosciences; mouse IgG1, κ PE, BV711, PE/Cy7, BV421, APC, AF647 (clone MOPC-21), mouse IgG2b, κ BV711 (MPC11) and mouse IgG2a, κ PE (MOP-173) all from Biolegend; mouse IgG1, κ PerCP-eFluor710 (P3.6.2.8.1; Thermo Fisher Scientific). Fixable Viability Dye eFluor 780 (ebioscience) or Zombie NIR Fixable Viability Kit (Biolegend) was used for live/dead cell discrimination. Cells were analysed on a LSR II, FACSCelesta^TM^, FACSymphony™ A5 SORP or Cytoflex (Beckman Coulter) machine and data was analysed with FloJo 10.8.1 (Tree Star). Sorting of CD155/CD276 MelDCC lines after CD155/CD276 CRIPSR/Cas9 knock-out was performed with a FACSAria™ IIu cell sorter (BD Bioscience).

### Statistical analysis

Statistical analysis was performed using the GraphPad Prism 9.3.1 software (GraphPad Software, Inc.) and R (version 4.1.0). Differences in mean values between groups were analysed by Student’s *t*-test or one-way ANOVA followed by post-hoc statistical testing, where appropriate. Univariable and multivariable analysis and survival analyses were performed using Cox-regression and Mantel-Cox test with a log-rank test. All tests were realized two-sided. A *P* value of less than 0.05 was considered statistically significant. DCCD+1 was log transformed with X=Log10(X).

## Data and material availability

The RNA sequencing data are deposited at the European Genome-Phenome Archive (EGA) database under accession number EGAS00001006702. The availability of patient-derived material and raw sequencing data is restricted based on individual patient consent and the General Data Protection Regulation (GDPR). All the other data supporting the findings of this study are available within the article and its Extended Data and source files and from the corresponding author upon reasonable request. A reporting summary for this article is available as a Supplementary Information file.

## Acknowledgements

We thank Sandra Grunewald, Anthea Povall, Isabell Blochberger, Irene Nebeja, Michaela Becker and Stefanie Güldener for excellent technical assistance. We are grateful to Mathias Oelke and Jonathan Schneck for providing aAPC. This work was supported by grants to CAK from the Deutsche Krebshilfe (Nr. 70113472 and 70112463 to CAK and LG), the Deutsche Forschungsgemeinschaft (DFG; KL 1233/14-1) and the Wilhelm Sander Stiftung (2020.129.1). The work of MW-K has been funded by the DFG (WE 4632-4/1, WE 4632/5-1/FOR2127 and B09/TRR 305) and Wilhelm Sander Stiftung (2015.016.1). The BD FACSymphony^TM^ was supported by the DFG (INST 89/518-1 FUGG).

## Authors’ Contributions

**Conception and design:** C. A. Klein and M. Werner-Klein

**Development of methodology:** S. M. Guetter, C. König, H. Koerkel-Qu, M. Werner-Klein, J. Warfsmann, R. Rachel, A. Braun

**Acquisition of data:** S. M. Guetter, C. König, H. Koerkel-Qu, A. Markiewicz, M. Katzer, M. Berneburg, S. Scheitler, P. Renner, B. Cucuruz, L. Guttenberger, V. Naimer, K. Weidele, S. Treitschke, C. Werno, H. Jaser, T. Bargmann, F. Weber, R. Rachel, F. Baumann, L. Schmidleithner, K. Schambeck, P. Mohammadi, A. Ulmer, S. Haferkamp, M. Werner-Klein,

**Analysis and interpretation of data:** S. M. Guetter, C. König, H. Koerkel-Qu, M. Werner-Klein, C. A. Klein

**Writing of the manuscript:** M. Werner-Klein, C. König, S. M. Guetter, H. Koerkel-Qu and C. A. Klein

**Review and/or revision of the manuscript:** all authors

**Competing interest:** The authors declare no potential conflicts of interest.

## Notes

### Competing Interest Statement

The authors have declared no competing interest.

