## Extended Data Fig. 1-7 for "Metastasis founder cells activate immunosuppression early in human melanoma metastatic colonization"

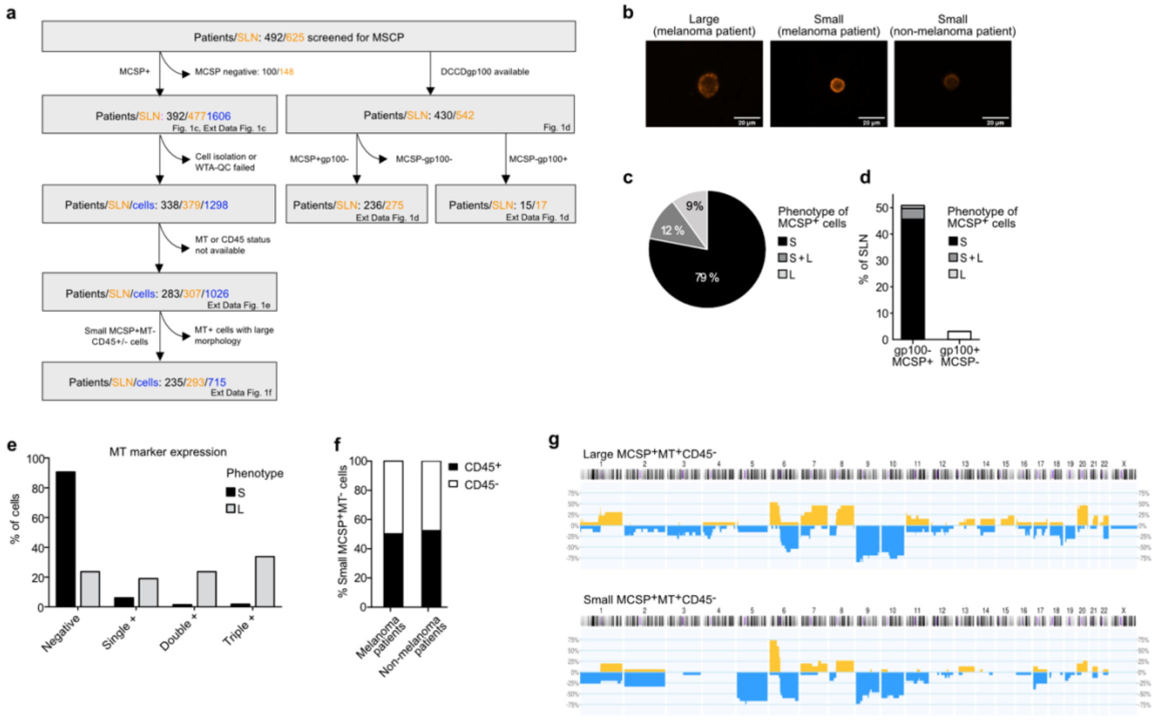


**Extended Data Figure 1: Expression of melanoma markers and CD45 in MCSP+ cells.**

**a,** Schematic overview of analyzed patient samples, exclusion criteria and reference individual figure panels. Number of patient numbers, SLN and cells are depicted in black, orange and blue, respectively. **b,** Staining intensity of MCSP$^+$ cells in SLN and LN of melanoma and non-melanoma patients. Scale bar as indicated. See also Fig. 1b for merged images of fluorescence and bright field channels of MCSP$^+$ cells in SLN of melanoma patients. **c**, Percentage of MCSP$^+$ SLN of melanoma patients with MCSP$^+$ cells separated according to their phenotype (diameter) into small (S), small and large (S+L), large (L) cells. SLN numbers see panel a and main text. **d**, Percentage of gp100$^-$MCSP$^+$ (n = 275) and gp100$^+$MCSP$^-$ SLN (n = 17) among SLN of melanoma patients with staining-results for both gp100 and MCSP (n = 542). gp100$^-$MCSP$^+$ SLN are annotated according to the observed phenotype (diameter) of detected MCSP$^+$ cells. **e**, Percentage of MCSP$^+$ small (n = 789) and large (n = 237) cells negative or positive for melanoma marker expression (*gp100*, *DCT* and *MLANA*) either alone (single$^+$) or in combination (double$^+$ (any two markers) or triple$^+$). **f**, Percentage of MCSP$^+$MT$^-$ cells with or without *CD45* expression isolated from melanoma-patients (MT$^-$ small cells, n = 715) and non-melanoma patients (MT$^-$ small cells, n = 61). **g**, Cumulative frequency plots of genomic aberrations in

$MCSP^+MT^+CD45^-$ large (n = 13) and small (n = 15) cells with genomic gains and losses in orange and blue, respectively.

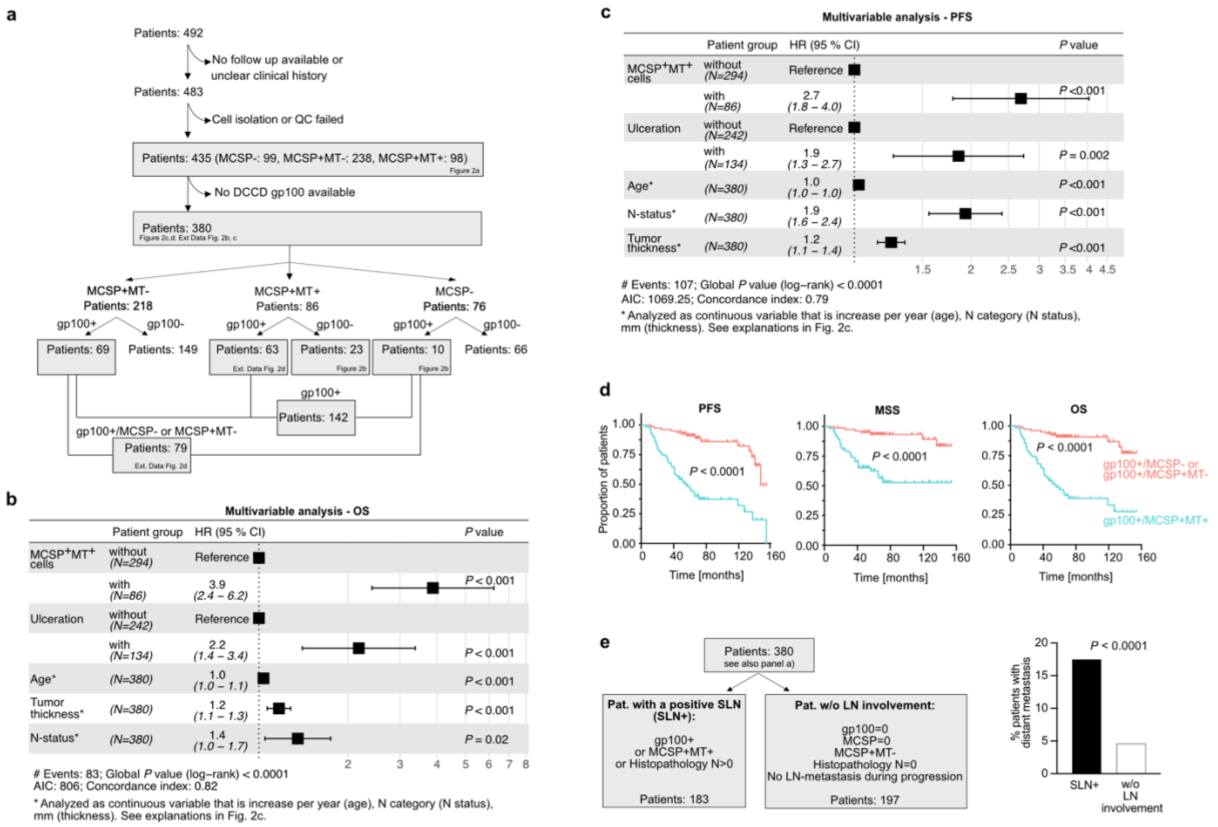


**Extended Data Figure 2: MCSP+ DCC impact on progression and patient survival.**

**a,** Schematic overview of analyzed patient samples and exclusion criteria. Note reference for individual figure panels. **b, c,** Multivariable Cox regression analysis for OS (**b**) and PFS (**c**) (n = 380) comprising the most informative, backward selected features. Ulceration group a and b denotes patients without and with ulceration, and MCSP$^{+}$MT$^{+}$ group 0 and 1 categorizes patients without and with MCSP$^{+}$MT$^{+}$ cells. Group 0 MCSP$^{+}$MT$^{+}$ patients and patients without ulceration were used as reference for defining the hazard ratio (HR). **d,** Kaplan Meier curves of PFS, MSS and OS of patients with gp100$^{+}$ cells stratified according to whether MCSP$^{+}$ DCC (n=63) were co-detected in the SLN or not (n=79). **e,** Impact of lymph node involvement on distant metastasis. SLN$^{+}$: patients with gp100$^{+}$ and/or MCSP$^{+}$MT$^{+}$ cells in the SLN or with a positive SLN by histopathology. LN$^{-}$: patients lacking gp100$^{+}$ or MCSP$^{+}$MT$^{+}$ cells in SLN, a negative SLN by histopathology and without evidence of LN-involvement at any time during the disease. *P* values in **b, c,** Wald test. **d,** Fisher's exact test. **e,** Log-rank test. See Supplementary Table 1 for baseline characteristics of study cohort.

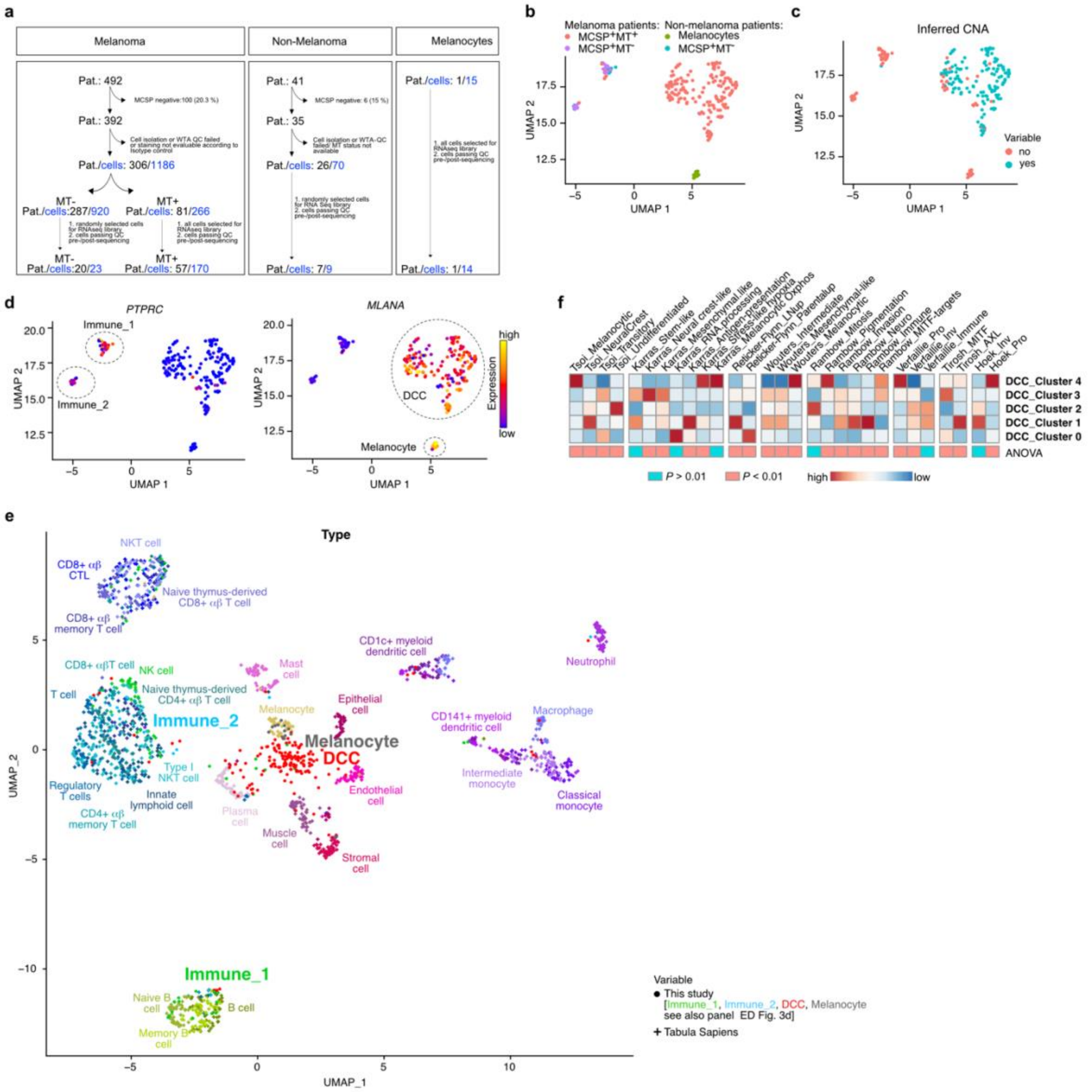

**Extended Data Figure 3: Molecular subtypes of melanoma DCC.**

**a,** Schematic overview of analyzed patient samples and exclusion criteria. The number of patients and cells are depicted in black and blue, respectively. Patient numbers are depicted in black, cell numbers in blue. **b-d,** UMAP of scRNA-seq data of lymph node-derived small and large MCSP$^{+}$MT$^{+}$ cells (n = 170) and MCSP$^{+}$MT$^{-}$ cells (n = 23) from melanoma patients, lymph node-derived MCSP$^{+}$ cells from non-melanoma patients (n = 9) and cultured human melanocytes (n = 14) with Seurat. Cells are annotated by their patient origin (melanoma, non-melanoma patient) and cell type (melanocyte, MCSP/MT-status) (b), gene expression inferred CNA (c) or expression of marker genes for immune cells (*PTPRC, CD45*) and cells of melanocytic origin (*MLANA*) (d). **e,** scRNA-seq data of Immune_1, Immune_2, DCC, melanocytes (see panel d) integrated into skin and lymph node of the Human cell atlas (Tabula sapiens).

Each cell type of the human cell atlas (n = 33) was downsampled to contain 50 cells. Each cell is a point and colored by its assigned cluster. **f,** AUCell scores of published melanoma subtype marker gene sets averaged and replotted over DCC clusters.

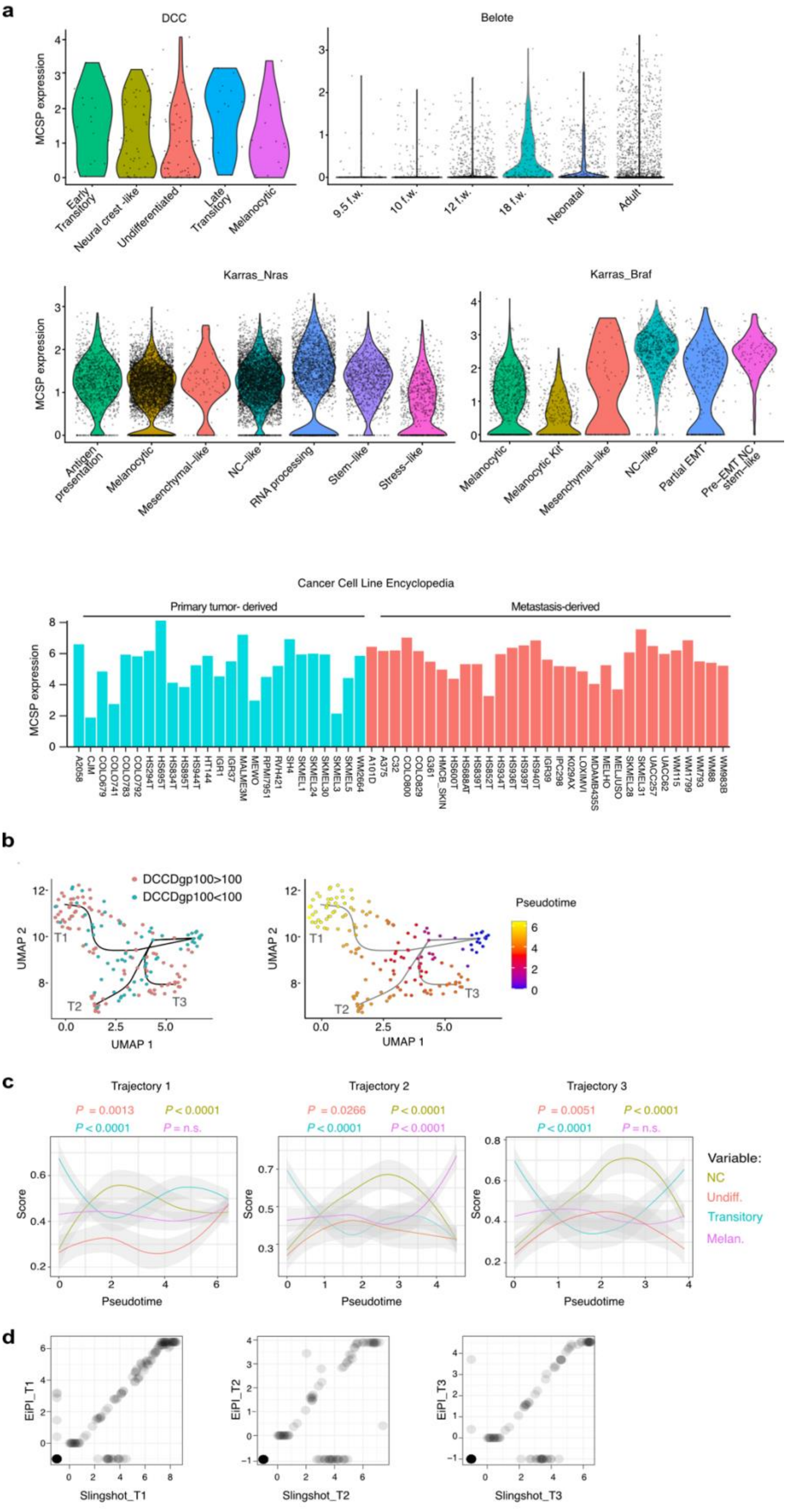
a
DCC
MCSP expression
Early Transitory
Neural crest -like
Undifferentiated
Late Transitory
Melanocytic
Belote
9.5 f.w.
10 f.w.
12 f.w.
18 f.w.
Neonatal
Adult
Karras_Nras
Antigen presentation
Melanocytic
Mesenchymal-like
NC-like
RNA processing
Stem-like
Stress-like
Karras_Braf
Melanocytic
Melanocytic Kit
Mesenchymal-like
NC-like
Partial EMT
Pre-EMT NC stem-like
Cancer Cell Line Encyclopedia
Primary tumor- derived
Metastasis-derived
A2058 CJM COLO679 COLO741 COLO783 COLO792 HS294T HS695T HS834T HS895T HS944T HT144 IGR1 IGR37 MALME3M MEWO RPMI7951 RVH421 SH4 SKMEL1 SKMEL24 SKMEL30 SKMEL3 SKMEL5 WM2664
A101D A375 C32 COLO800 COLO829 G361 HMCB_SKIN HS600T HS688AT HS839T HS852T HS934T HS936T HS939T HS940T IGR39 IPC298 K029AX LOXIMVI MDAMB435S MELHO MELJUSO SKMEL28 SKMEL31 UACC257 UACC62 WM115 WM1799 WM793 WM88 WM983B
b
DCCDgp100>100
DCCDgp100<100
UMAP 2
UMAP 1
T1
T2
T3
Pseudotime
c
Trajectory 1
P = 0.0013
P < 0.0001
P < 0.0001
P = n.s.
Trajectory 2
P = 0.0266
P < 0.0001
P < 0.0001
P < 0.0001
Trajectory 3
P = 0.0051
P < 0.0001
P < 0.0001
P = n.s.
Score
Pseudotime
Variable:
NC
Undiff.
Transitory
Melan.
d
EiPI_T1
Slingshot_T1
EiPI_T2
Slingshot_T2
EiPI_T3
Slingshot_T3

**Extended Data Figure 4: MCSP expression and progression of melanoma subtypes during metastatic lymph node colonization.**

**a,** MCSP expression in melanoma DCC-clusters and published scRNA-seq data-sets [1,2] or in melanoma cell lines of the Cancer Cell Line Encyclopedia [3]. **b,** Inferred trajectories (T1, T2, T3) with ElPiGraph. Left: each cell is colored according to its DCCD (DCCD < 100 blue, DCCD > 100 red). Right: each cell is colored according to its pseudotime. **c,** Melanocytic, neural-crest-like, transitory and undifferentiated signature scores [4] with AUCell along pseudotime (EiPl) of trajectory 1-3**. d,** Inferred pseudotime using Slingshot and ElPiGraph for trajectory 1-3.

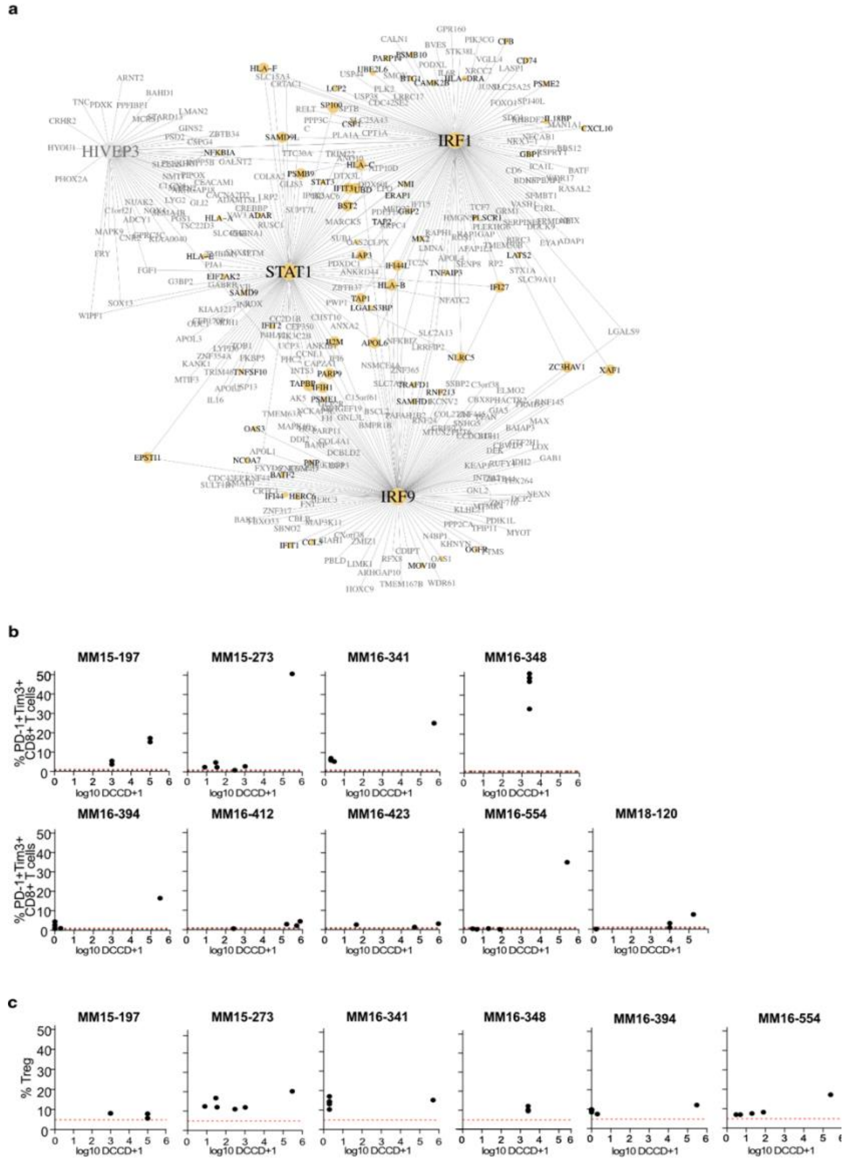


**Extended Data Figure 5: DCC and immune cell communication.**

**a,** Transcription factor regulatory network highly activated in cluster 1 DCC. Black letters and yellow circles indicate genes involved in interferon alpha/gamma signaling pathways. **b, c** Percentage of PD1$^+$TIM3$^+$ CD8 T cells (b) and CD4$^+$CD25$^+$CD127$^-$ regulatory T cells (c) versus the DCCD of lymph nodes originating from the same regional bed of melanoma patients (n = 6 - 9).

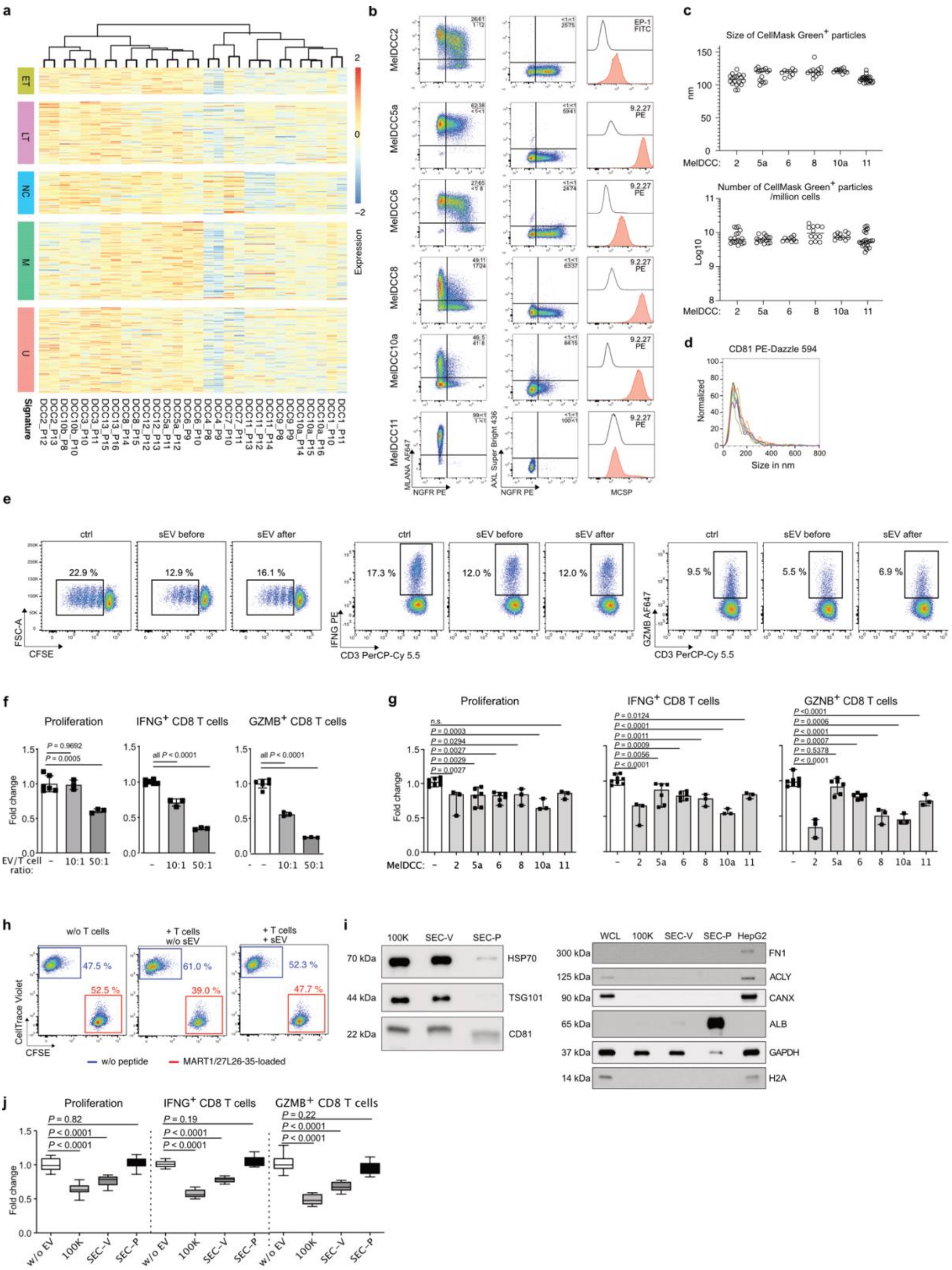


**Extended Data Figure 6: Impact of sEV on CD8 T cell function.**

**a,** Expression of DCC marker genes for the early and late transitional (ET and LT), Neural crest-like (NC), undifferentiated (U) and melanocytic (M) phenotype in MelDCC1-13 DCC. Each cell line was analyzed by bulk RNAseq and in duplicates or triplicates of consecutive passages. **b,** Flowcytometric analysis of MelDCC 2, 5a, 6, 8, 10a, 11 for NGFR (neural-crest marker), MelanA (melanocytic marker), AXL (invasiveness marker) and MCSP expression. **c,** Nanoparticle tracking analysis based enumeration of sEV secreted per million cells of indicated MelDCC lines within 48 hours. **d,** Diameter of CD81 positive sEV of MelDCC 10a (n = 7) as determined by NTA. **e-g,** Flow cytometric analysis of proliferation and IFNG and GZMB production of CD8 T cells 4 days after anti-CD3/CD28 stimulation and exposed to PBS (n = 6) or sEV (MelDCC 10a (e, f); MelDCC 2, 5a, 6, 8, 10a, 11 (g); n = 3-6). sEV were added 18 h before anti-CD3/CD28 stimulation in a 50:1 ratio, if not indicated otherwise, i.e. sEV produced by 50 MelDCC cells per CD8 T cell. **h,** Flow cytometric analysis of the antigen-specific cytotoxicity assay (see Fig. 5g) with $MART1_{27L26\text{-}35}$-loaded CFSE-labeled T2 cells and non-loaded, CellTrace Violet T2 cells as non-target reference population. **i,** Western blot analysis of a 100K EV-preparation or a 100K EV-preparation separated by size exclusion chromatography into a vesicle (SEC-V) and protein (SEC-P) fraction. Antibodies considered as markers for small (CD81, TSG101) and large EV (GRP94), pan-EV marker (HSP70) or non-vesicular contaminants (FN1, ACLY, CANX, ALB, H2A) were used. This albumin signal in the SEC-P fraction is attributable to a washing step absent in the size-exclusion chromatography SEC-workflow as compared to EV preparation via ultracentrifugation **j,** Flowcytometric analysis of proliferation and IFNG and GZMB production of $MART1_{27L26\text{-}35}$-specific CD8 T cells. CD8 T cells were exposed to PBS (n = 6) or sEV of MelDCC 10a (n = 6) 18 h prior to anti-CD3/CD28 stimulation. *P* values in **f, g, j** one-way ANOVA with Dunnett's post hoc multiple comparison test.

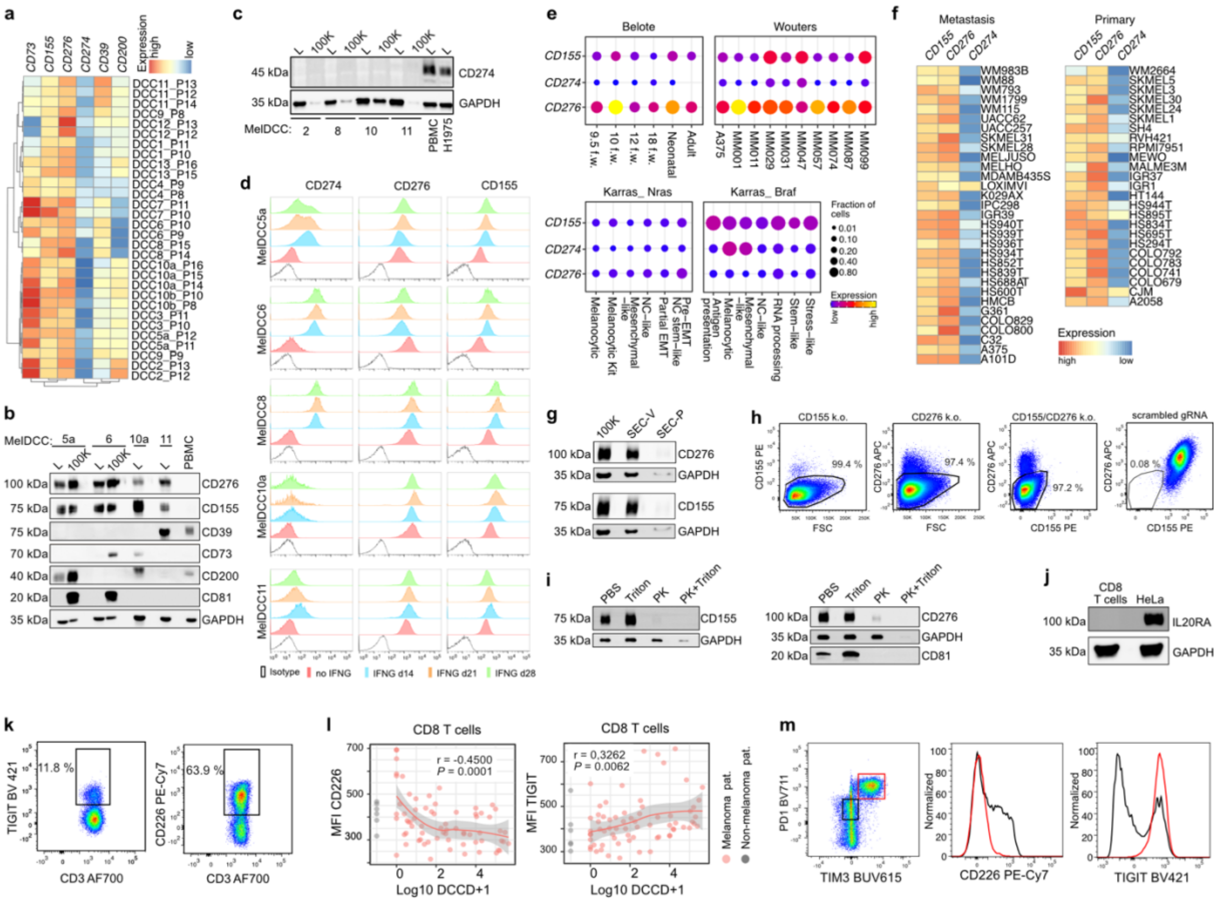


**Extended Data Figure 7: Analysis of sEV-associated immune checkpoint ligands.**

**a,** Expression of CD155, CD274 and CD276 in MelDCC 1-13. Each cell line was analyzed by bulk RNAseq and in duplicates or triplicates of consecutive passages. **b, c,** Western blot analysis for expression of immune checkpoint ligands in MelDCC lines (L) and their respective 100K pellets. **d,** Flowcytometric analysis of CD155, CD274 and CD276 expression in MelDCC lines cultured in the absence (red) or presence of 500 U IFNG for 14 (blue), 21 (orange) or 28 (green) days. As isotype controls did not differ between the time points, only the isotype control of untreated cells at d0 is shown. **d, e,** Expression of *CD155*, *CD276* and *CD274* (PD-L1) in publicly available scRNA-seq data of melanoma [1,2,5] or bulkRNAseq of cell lines of the cancer cell line encyclopedia[3]. **g,** Western blot analysis for the presence of CD155 and CD276 in the 100K EV-preparation separated by size exclusion chromatography into a vesicle (SEC-V) and protein (SEV-P) fraction. **h,** Flow cytometric analysis for CD155 and CD276 expression in MelDCC 10a CD155/CD276 single or double knock-out and controls. **i,** Western blot analysis of sEV treated with PBS, Triton X-100, Proteinase K or Proteinase K plus Triton X-100 for CD81, CD155, CD276 and GADPH. **j,** Western blot analysis for IL20RA expression in CD8

T cells from a healthy donor and HeLa cells as positive control. **k-m,** Flowcytometric analysis of TIGIT and CD226 expression in polyclonal CD8 T cells from peripheral blood of a healthy donor (k) and lymph nodes of melanoma patients (n = 69) and non-melanoma patients (n = 6) (l, m,). Red and grey symbols in l indicate detected values of individual lymph nodes from melanoma and non-melanoma patients, respectively. The grey area indicates the standard error of the smoothed curve (red line). *P* values in l according to Pearson's correlation.

### References Extended Data


1 Karras, P. *et al.* A cellular hierarchy in melanoma uncouples growth and metastasis. *Nature* **610**, 190-198 (2022). https://doi.org:10.1038/s41586-022-05242-7

2 Belote, R. L. *et al.* Human melanocyte development and melanoma dedifferentiation at single-cell resolution. *Nat Cell Biol* **23**, 1035-1047 (2021). https://doi.org:10.1038/s41556-021-00740-8

3 Barretina, J. *et al.* The Cancer Cell Line Encyclopedia enables predictive modelling of anticancer drug sensitivity. *Nature* **483**, 603-607 (2012). https://doi.org:10.1038/nature11003

4 Tsoi, J. *et al.* Multi-stage Differentiation Defines Melanoma Subtypes with Differential Vulnerability to Drug-Induced Iron-Dependent Oxidative Stress. *Cancer Cell* **33**, 890-904 e895 (2018). https://doi.org:10.1016/j.ccell.2018.03.017

5 Wouters, J. *et al.* Robust gene expression programs underlie recurrent cell states and phenotype switching in melanoma. *Nat Cell Biol* **22**, 986-998 (2020). https://doi.org:10.1038/s41556-020-0547-3
