## Supplementary Table 1 for "Metastasis founder cells activate immunosuppression early in human melanoma metastatic colonization"

| Characteristic | Number of patients | Percentage [%] | Median | Range | IR |
|---|---|---|---|---|---|
| **Patients** | **435** | | | | |
| **Gender** | | | | | |
| Female | 192 | 44.1 | | | |
| Male | 243 | 55.9 | | | |
| **Age (years)** | | | 56 | 15-85 | |
| **Breslow´s thickness (mm)** | | | 2 | 0.2-15.0 | 1-3 |
| **Ulceration** | | | | | |
| No | 278 | 63.9 | | | |
| Yes | 153 | 35.2 | | | |
| Not specified | 4 | 0.9 | | | |
| **Localization** | | | | | |
| Extremities | 252 | 57.9 | | | |
| Trunk or head | 183 | 42.1 | | | |
| **Nodal status histopathology** | | | | | |
| Negative | 342 | 78.6 | | | |
| Positive | 93 | 21.4 | | | |
| **Clinical stage** | | | | | |
| IA | 49 | 11.3 | | | |
| IB | 168 | 38.6 | | | |
| II | 1 | 0.2 | | | |
| IIA | 73 | 16.8 | | | |
| IIB | 40 | 9.2 | | | |
| IIC | 12 | 2.8 | | | |
| III | 4 | 0.9 | | | |
| IIIA | 32 | 7.4 | | | |
| IIIB | 38 | 8.7 | | | |
| IIIC | 17 | 3.9 | | | |
| IIID | 1 | 0.2 | | | |
| **DCCD [a] (MCSP)** | | | 5 | 0-400.000 | 1-12 |
| **DCCD [b] (gp100)** | | | 0 | 0-500.000 | 0-1 |
| **Survival** | | | | | |
| Deceased | 98 | 22.5 | | | |
| Alive | 337 | 77.5 | | | |
| **Adjuvant therapy** | | | | | |
| Yes[c] | 54 | 12.4 | | | |
| No/not specified | 381 | 87.6 | | | |

IR, interquartile range.

[a] Disseminated cancer cell density in the sentinel node; number of MCSP positive cells per million isolated cells, if more than one node per patient was positive, the node with the highest cancer cell density was taken.

[b] Disseminated cancer cell density in the sentinel node; number of gp100 positive cells per million isolated cells, if more than one node per patient was positive, the node with the highest cancer cell density was taken.
