## Supplementary Table 2 for "Metastasis founder cells activate immunosuppression early in human melanoma metastatic colonization"

**Fig. 3b**

| **Cluster comparisons** | **Melanocytic_adjusted p value** | **NeuralCrest_adjusted p value** | **Transitory_adjusted p value** | **Undifferentiated_adjusted p value** |
|---|---|---|---|---|
| 0 - 1 | 0.78 | 5.00E-06 | 1.60E-05 | 0.37 |
| 0 - 2 | 0.78 | 0.58 | 0.9 | 1 |
| 0 - 3 | 0.78 | 0.014 | 0.00099 | 0.0028 |
| 0 - 4 | 3.10E-06 | 0.79 | 4.90E-11 | 1 |
| 1 - 2 | 0.78 | 0.00031 | 0.12 | 0.27 |
| 1 - 3 | 0.2 | 0.0068 | 0.003 | 1 |
| 1 - 4 | 1.30E-05 | 0.00025 | 1.40E-07 | 1 |
| 2 - 3 | 0.66 | 0.79 | 0.024 | 0.26 |
| 2 - 4 | 6.80E-06 | 0.1 | 4.90E-11 | 0.092 |
| 3 - 4 | 7.90E-07 | 0.79 | 2.30E-06 | 1 |

**Post-hoc analysis: pairwise Student's t-test**

**Fig. 3c**

| **Cluster comparisons** | **DCCDgp100_adjusted p value** |
|---|---|
| 0 - 1 | 0.956368 |
| 0 - 2 | 0.048148 |
| 0 - 3 | 0.000062 |
| 0 - 4 | 0.000693 |
| 1 - 2 | 0.029792 |
| 1 - 3 | 0.000013 |
| 1 - 4 | 0.000267 |
| 2 - 3 | 0.005775 |
| 2 - 4 | 0.048148 |
| 3 - 4 | 0.956368 |

**Post-hoc analysis: pairwise Student's t-test**

**Fig. 3d**

| | Early Transitory | NCSC | Undifferentiated | Late Transitory |
|---|---|---|---|---|
| **NCSC** | 0.08417 | - | - | - |
| **Undifferentiated** | 0.00017 | 0.00516 | - | - |
| **Late Transitory** | 3.50E-06 | 0.00011 | 0.01627 | - |
| **Melanocytic** | 0.00103 | 0.01867 | 0.53596 | 0.22414 |

**Fisher's exact test**

**Fig. 4b**

| Group1 | Group2 | MEyellow_adjusted p value | MEturquoise_adjusted p value | MEblue_adjusted p value | MEbrown_adju |
|---|---|---|---|---|---|
| Early Trans | NCSC | 2.80E-06 | 2.70E-09 | 8.80E-06 | 2.10E-10 |
| Early Trans | Undifferent. | 0.17 | 6.80E-11 | 0.004 | 4.00E-08 |
| Early Trans | Late Trans | 0.17 | 3.00E-06 | 0.81 | 1.60E-05 |
| Early Trans | Mela | 2.10E-06 | 2.70E-10 | 0.38 | 0.98 |
| NCSC | Undifferent. | 9.40E-23 | 7.70E-17 | 2.40E-10 | 4.70E-15 |
| NCSC | Late Trans | 2.60E-05 | 1.70E-12 | 2.20E-06 | 8.80E-06 |
| NCSC | Mela | 8.10E-06 | 1.10E-06 | 1.30E-07 | 3.60E-10 |
| Undifferent. | Late Trans | 0.0018 | 4.00E-19 | 6.40E-10 | 6.80E-11 |
| Undifferent. | Mela | 2.10E-06 | 0.00064 | 0.01 | 1.20E-05 |
| Late Trans | Mela | 2.80E-06 | 3.40E-15 | 0.09 | 5.90E-05 |

**Post-hoc analysis: pairwise Student's t-test**
