## Supplementary Table 3 for "Metastasis founder cells activate immunosuppression early in human melanoma metastatic colonization"

| | p_val | avg_log2FC | pct.1 | pct.2 | p_val_adj | gene | seurat_clusters |
|---|---|---|---|---|---|---|---|
| GPX3.1 | 6.47E-08 | 2.94710526 | 0.786 | 0.331 | 0.001412156 | GPX3 | 2 |
| LYSMD1 | 0.000134865 | 2.811222726 | 0.679 | 0.271 | 1 | LYSMD1 | 2 |
| MGP | 5.02E-10 | 2.686578971 | 0.929 | 0.391 | 1.10E-05 | MGP | 2 |
| RNASE1.1 | 5.37E-10 | 2.553831939 | 0.929 | 0.579 | 1.17E-05 | RNASE1 | 2 |
| ITIH5 | 1.32E-07 | 2.437791254 | 0.714 | 0.233 | 0.002870231 | ITIH5 | 2 |
| FGL2.1 | 4.38E-07 | 2.418932705 | 0.679 | 0.218 | 0.009567076 | FGL2 | 2 |
| SPARCL1.1 | 1.36E-12 | 2.276536616 | 0.964 | 0.459 | 2.96E-08 | SPARCL1 | 2 |
| TTR.1 | 0.000213378 | 2.262979312 | 0.571 | 0.195 | 1 | TTR | 2 |
| SPP1 | 1.07E-07 | 2.183424711 | 0.893 | 0.549 | 0.002345022 | SPP1 | 2 |
| MT2A.1 | 3.70E-07 | 2.05385609 | 0.929 | 0.782 | 0.00807232 | MT2A | 2 |
| COL1A2 | 0.000260264 | 2.025725448 | 0.75 | 0.474 | 1 | COL1A2 | 2 |
| NDRG1 | 3.84E-08 | 2.017094573 | 1 | 0.827 | 0.000838225 | NDRG1 | 2 |
| KLHL29 | 2.29E-05 | 1.69797565 | 0.857 | 0.406 | 0.499657303 | KLHL29 | 2 |
| ANGPTL7 | 0.001115804 | 1.69042802 | 0.464 | 0.173 | 1 | ANGPTL7 | 2 |
| ANGPT2 | 6.15E-09 | 1.586963385 | 0.857 | 0.353 | 0.000134193 | ANGPT2 | 2 |
| FCER1G | 0.001062815 | 1.585715553 | 0.464 | 0.173 | 1 | FCER1G | 2 |
| A2M.1 | 6.41E-05 | 1.583127499 | 1 | 0.88 | 1 | A2M | 2 |
| DAGLB | 2.33E-05 | 1.530021274 | 0.821 | 0.368 | 0.508178891 | DAGLB | 2 |
| ALDH1A3.1 | 0.005617622 | 1.438228742 | 0.75 | 0.444 | 1 | ALDH1A3 | 2 |
| ESRRA.1 | 0.000474127 | 1.426865744 | 0.75 | 0.353 | 1 | ESRRA | 2 |
| FGFR1 | 0.007394514 | 1.423086173 | 0.679 | 0.511 | 1 | FGFR1 | 2 |
| FGFBP2 | 4.41E-06 | 1.368004059 | 0.571 | 0.158 | 0.096338498 | FGFBP2 | 2 |
| CYHR1 | 6.93E-05 | 1.354169615 | 0.821 | 0.429 | 1 | CYHR1 | 2 |
| ZNF316.1 | 0.000125373 | 1.331940754 | 0.929 | 0.662 | 1 | ZNF316 | 2 |
| ZMAT2.1 | 5.38E-06 | 1.312928462 | 0.929 | 0.669 | 0.117487306 | ZMAT2 | 2 |
| ALDH1A1 | 4.24E-05 | 1.291275124 | 0.857 | 0.534 | 0.924752786 | ALDH1A1 | 2 |
| KCNJ10.1 | 3.48E-07 | 1.28742182 | 0.75 | 0.233 | 0.007602337 | KCNJ10 | 2 |
| CFH.1 | 4.81E-08 | 1.283280768 | 0.75 | 0.195 | 0.001050262 | CFH | 2 |
| SPRY4 | 6.10E-07 | 1.23680559 | 0.929 | 0.782 | 0.0133119 | SPRY4 | 2 |
| SOD2.1 | 0.006571084 | 1.233823077 | 0.964 | 0.752 | 1 | SOD2 | 2 |
| C1S | 0.001135413 | 1.219466827 | 0.714 | 0.338 | 1 | C1S | 2 |
| NELL1 | 0.001365555 | 1.215019077 | 0.393 | 0.256 | 1 | NELL1 | 2 |
| EFEMP2.1 | 4.53E-06 | 1.206331176 | 0.75 | 0.436 | 0.098887205 | EFEMP2 | 2 |
| SERINC5 | 7.10E-07 | 1.196716728 | 0.786 | 0.632 | 0.015501528 | SERINC5 | 2 |
| APOD.1 | 0.002410185 | 1.187251363 | 1 | 0.955 | 1 | APOD | 2 |
| IRS2 | 0.001200599 | 1.168102228 | 0.857 | 0.579 | 1 | IRS2 | 2 |
| TIMP1.1 | 0.000177947 | 1.146875493 | 0.964 | 0.677 | 1 | TIMP1 | 2 |
| AKAP12 | 4.18E-05 | 1.140249092 | 0.964 | 0.714 | 0.912244732 | AKAP12 | 2 |
| TSPAN7.1 | 2.87E-06 | 1.137014174 | 0.893 | 0.549 | 0.062681069 | TSPAN7 | 2 |
| FMOD | 3.88E-06 | 1.107834306 | 0.536 | 0.113 | 0.084652995 | FMOD | 2 |
| SPARC.1 | 0.003434318 | 1.095014378 | 1 | 0.932 | 1 | SPARC | 2 |
| AEBP1 | 9.91E-05 | 1.077324495 | 1 | 0.955 | 1 | AEBP1 | 2 |
| ARHGEF34P | 4.97E-09 | 1.075317132 | 0.786 | 0.18 | 0.000108505 | ARHGEF34P | 2 |
| PLEKHB1 | 1.90E-05 | 1.07399512 | 0.857 | 0.564 | 0.415609998 | PLEKHB1 | 2 |
| ABHD2.1 | 0.001622455 | 1.069318744 | 1 | 0.82 | 1 | ABHD2 | 2 |
| EMILIN1 | 1.31E-06 | 1.058722616 | 0.786 | 0.323 | 0.02855659 | EMILIN1 | 2 |
| DKK3 | 0.009859643 | 1.052635791 | 0.464 | 0.188 | 1 | DKK3 | 2 |
| GPR37 | 2.51E-06 | 1.035141457 | 0.714 | 0.308 | 0.054857916 | GPR37 | 2 |
| FCGR2C | 9.04E-05 | 1.032516423 | 0.607 | 0.218 | 1 | FCGR2C | 2 |
| SERPINE2 | 1.21E-06 | 1.023197945 | 1 | 0.977 | 0.026351875 | SERPINE2 | 2 |
| MATN2.1 | 0.000118104 | 0.997804503 | 0.821 | 0.406 | 1 | MATN2 | 2 |
| ERGIC1 | 0.001558703 | 0.996781501 | 0.964 | 0.842 | 1 | ERGIC1 | 2 |
| SYNM.1 | 0.000148706 | 0.996321654 | 0.893 | 0.579 | 1 | SYNM | 2 |
| GJB2 | 1.23E-05 | 0.990276886 | 0.5 | 0.113 | 0.268642935 | GJB2 | 2 |
| SDC2 | 8.27E-06 | 0.967836198 | 0.643 | 0.233 | 0.180565196 | SDC2 | 2 |
| SWAP70 | 0.000560745 | 0.965683499 | 1 | 0.827 | 1 | SWAP70 | 2 |
| BAALC | 1.13E-10 | 0.960173608 | 0.893 | 0.316 | 2.46E-06 | BAALC | 2 |
| POSTN.1 | 2.61E-06 | 0.944820735 | 0.964 | 0.541 | 0.057024245 | POSTN | 2 |
| LRATD2 | 2.09E-05 | 0.944334009 | 0.714 | 0.278 | 0.457129411 | LRATD2 | 2 |
| CD59 | 7.46E-05 | 0.939134556 | 1 | 0.977 | 1 | CD59 | 2 |
| NBL1 | 0.000700566 | 0.937437704 | 0.714 | 0.376 | 1 | NBL1 | 2 |
| CTDNEP1.1 | 0.002077275 | 0.928306995 | 0.786 | 0.466 | 1 | CTDNEP1 | 2 |
| TNFRSF21 | 2.23E-07 | 0.907753253 | 0.929 | 0.414 | 0.004861715 | TNFRSF21 | 2 |
| HERC2P2.1 | 0.004638759 | 0.90573731 | 0.893 | 0.707 | 1 | HERC2P2 | 2 |
| AIF1L | 0.005618354 | 0.89871215 | 0.786 | 0.534 | 1 | AIF1L | 2 |
| BIRC2 | 0.000798377 | 0.891128312 | 0.714 | 0.338 | 1 | BIRC2 | 2 |
| LHFPL2 | 0.000606102 | 0.890634111 | 0.964 | 0.752 | 1 | LHFPL2 | 2 |
| PDE3B.1 | 2.38E-07 | 0.877701525 | 0.964 | 0.481 | 0.005193543 | PDE3B | 2 |
| WFS1.1 | 0.007375863 | 0.87615821 | 0.643 | 0.421 | 1 | WFS1 | 2 |
| PTN | 0.00047746 | 0.873391681 | 0.321 | 0.06 | 1 | PTN | 2 |
| VAPB.1 | 0.001841743 | 0.871236655 | 0.929 | 0.677 | 1 | VAPB | 2 |
| MVP.1 | 7.06E-05 | 0.869793462 | 0.964 | 0.639 | 1 | MVP | 2 |
| ST6GAL1 | 0.000362116 | 0.857422352 | 1 | 0.759 | 1 | ST6GAL1 | 2 |
| STAT3.1 | 0.001907531 | 0.835608378 | 0.964 | 0.805 | 1 | STAT3 | 2 |
| STAT5B | 0.002368112 | 0.824636095 | 0.786 | 0.444 | 1 | STAT5B | 2 |
| MGST1 | 0.001960899 | 0.821794813 | 0.464 | 0.158 | 1 | MGST1 | 2 |
| TSPAN14 | 0.000807649 | 0.818660401 | 0.964 | 0.857 | 1 | TSPAN14 | 2 |
| HKDC1.1 | 5.23E-08 | 0.818359974 | 0.786 | 0.211 | 0.001142074 | HKDC1 | 2 |
| OBSL1 | 1.78E-05 | 0.817979968 | 0.857 | 0.429 | 0.388237595 | OBSL1 | 2 |
| EIF3M.1 | 0.00315558 | 0.817277274 | 1 | 0.782 | 1 | EIF3M | 2 |
| LOXL4 | 0.008927279 | 0.816661888 | 0.857 | 0.602 | 1 | LOXL4 | 2 |
| GBP4.1 | 5.81E-05 | 0.807742702 | 0.786 | 0.338 | 1 | GBP4 | 2 |
| ANKS1A | 0.003037626 | 0.806833557 | 1 | 0.85 | 1 | ANKS1A | 2 |
| HSPA7 | 0.000266126 | 0.806785751 | 0.643 | 0.256 | 1 | HSPA7 | 2 |
| SELENOP | 5.89E-08 | 0.799260192 | 0.571 | 0.09 | 0.001286148 | SELENOP | 2 |
| ARHGEF5 | 1.95E-05 | 0.798129756 | 0.714 | 0.293 | 0.424469616 | ARHGEF5 | 2 |
| CREB3.1 | 0.000903691 | 0.795023011 | 0.571 | 0.233 | 1 | CREB3 | 2 |
| NOMO1.1 | 0.000790529 | 0.794487644 | 1 | 0.782 | 1 | NOMO1 | 2 |
| GPNMB.1 | 0.000962239 | 0.793116307 | 1 | 0.985 | 1 | GPNMB | 2 |
| FABP7 | 6.62E-05 | 0.790509436 | 0.607 | 0.188 | 1 | FABP7 | 2 |
| EIF4H.1 | 0.000464585 | 0.78751115 | 1 | 0.917 | 1 | EIF4H | 2 |
| ATP1B1.1 | 1.39E-07 | 0.786039416 | 0.929 | 0.444 | 0.003031603 | ATP1B1 | 2 |
| CSPG4.1 | 0.000202359 | 0.781404332 | 1 | 0.917 | 1 | CSPG4 | 2 |
| PARP10 | 0.007304736 | 0.778975821 | 0.714 | 0.459 | 1 | PARP10 | 2 |
| POM121.1 | 0.001661508 | 0.773916906 | 1 | 0.797 | 1 | POM121 | 2 |

| | | | | | | | |
|---|---|---|---|---|---|---|---|
| SEMA3D | 5.27E-06 | 0.771565093 | 0.607 | 0.195 | 0.114914498 | SEMA3D | 2 |
| MGAT5 | 0.000400104 | 0.770455959 | 0.964 | 0.759 | 1 | MGAT5 | 2 |
| MX1 | 0.001791305 | 0.763204433 | 0.607 | 0.361 | 1 | MX1 | 2 |
| C7orf50 | 0.000683964 | 0.761235038 | 0.607 | 0.271 | 1 | C7orf50 | 2 |
| CC2D1A | 4.99E-06 | 0.760775841 | 0.929 | 0.496 | 0.108973465 | CC2D1A | 2 |
| DCTN6.1 | 0.00111862 | 0.756290448 | 0.786 | 0.414 | 1 | DCTN6 | 2 |
| TFCP2L1 | 0.000419951 | 0.751588848 | 0.393 | 0.135 | 1 | TFCP2L1 | 2 |
| STRN4 | 0.000121466 | 0.745143122 | 0.893 | 0.534 | 1 | STRN4 | 2 |
| CAV1.1 | 0.000930847 | 0.744301328 | 1 | 0.805 | 1 | CAV1 | 2 |
| ADGRG1.1 | 0.007041895 | 0.743283725 | 1 | 0.917 | 1 | ADGRG1 | 2 |
| SPON2.1 | 6.21E-06 | 0.741902276 | 0.714 | 0.226 | 0.135476023 | SPON2 | 2 |
| F5 | 1.65E-08 | 0.728920035 | 0.607 | 0.09 | 0.000360068 | F5 | 2 |
| SEMA5A | 0.002147793 | 0.728791156 | 0.893 | 0.662 | 1 | SEMA5A | 2 |
| GPM6B.1 | 0.000341852 | 0.728245929 | 1 | 0.827 | 1 | GPM6B | 2 |
| CYFIP2 | 1.04E-05 | 0.726546271 | 0.75 | 0.338 | 0.227779522 | CYFIP2 | 2 |
| ABCA6 | 3.90E-09 | 0.726476194 | 0.857 | 0.301 | 8.52E-05 | ABCA6 | 2 |
| LRP2 | 0.000741738 | 0.725438307 | 0.571 | 0.226 | 1 | LRP2 | 2 |
| SEM1.1 | 0.000964362 | 0.724571363 | 0.857 | 0.632 | 1 | SEM1 | 2 |
| DAB2IP | 0.000375778 | 0.7226582 | 0.821 | 0.451 | 1 | DAB2IP | 2 |
| CPSF1 | 0.008099178 | 0.72239805 | 0.821 | 0.729 | 1 | CPSF1 | 2 |
| TBX2 | 0.006179982 | 0.719682257 | 0.786 | 0.504 | 1 | TBX2 | 2 |
| NUMA1.1 | 0.004956244 | 0.716190636 | 1 | 0.94 | 1 | NUMA1 | 2 |
| SORL1 | 0.000147313 | 0.711298676 | 0.964 | 0.677 | 1 | SORL1 | 2 |
| DUSP6.1 | 0.001973657 | 0.708588144 | 0.964 | 0.767 | 1 | DUSP6 | 2 |
| CCHCR1 | 0.007450806 | 0.70733078 | 0.393 | 0.338 | 1 | CCHCR1 | 2 |
| G6PD | 0.000256603 | 0.706617365 | 0.786 | 0.451 | 1 | G6PD | 2 |
| GPAT4.1 | 0.007563728 | 0.705045653 | 0.929 | 0.82 | 1 | GPAT4 | 2 |
| TSR2.1 | 0.008042544 | 0.694744483 | 0.714 | 0.526 | 1 | TSR2 | 2 |
| NAGLU | 0.003158192 | 0.69358989 | 0.786 | 0.451 | 1 | NAGLU | 2 |
| LPAR1 | 1.43E-09 | 0.69260682 | 0.643 | 0.083 | 3.11E-05 | LPAR1 | 2 |
| FKBP10.1 | 0.006632377 | 0.687269486 | 0.964 | 0.835 | 1 | FKBP10 | 2 |
| CNST | 0.006879561 | 0.679543776 | 0.821 | 0.534 | 1 | CNST | 2 |
| TRAF4 | 0.000444754 | 0.675683121 | 0.857 | 0.489 | 1 | TRAF4 | 2 |
| HLA-C.1 | 0.006656691 | 0.671937947 | 0.964 | 0.759 | 1 | HLA-C | 2 |
| PRXL2A | 0.002562341 | 0.670264925 | 0.857 | 0.624 | 1 | PRXL2A | 2 |
| MET | 0.002080763 | 0.668362955 | 0.786 | 0.481 | 1 | MET | 2 |
| JADE1 | 0.002044987 | 0.668226191 | 0.821 | 0.602 | 1 | JADE1 | 2 |
| CD320 | 0.007094644 | 0.665521659 | 0.393 | 0.135 | 1 | CD320 | 2 |
| ING5 | 0.007056055 | 0.664920032 | 0.714 | 0.519 | 1 | ING5 | 2 |
| RPL23 | 0.005553009 | 0.66342105 | 1 | 0.955 | 1 | RPL23 | 2 |
| LONP2 | 0.003776863 | 0.65954389 | 1 | 0.962 | 1 | LONP2 | 2 |
| CFI | 3.61E-05 | 0.656472136 | 0.5 | 0.12 | 0.788608932 | CFI | 2 |
| CRYAB | 0.001351158 | 0.65325944 | 0.821 | 0.594 | 1 | CRYAB | 2 |
| DIRAS1 | 0.000725313 | 0.651957084 | 0.643 | 0.263 | 1 | DIRAS1 | 2 |
| COQ9.1 | 0.00011447 | 0.651868434 | 0.714 | 0.338 | 1 | COQ9 | 2 |
| SEC11C | 0.000175791 | 0.649725333 | 0.964 | 0.677 | 1 | SEC11C | 2 |
| RNF157 | 8.54E-05 | 0.649584308 | 0.786 | 0.383 | 1 | RNF157 | 2 |
| CORO2B | 3.73E-05 | 0.642609625 | 0.929 | 0.526 | 0.813441013 | CORO2B | 2 |
| KIAA1614 | 0.000170888 | 0.641554902 | 0.75 | 0.338 | 1 | KIAA1614 | 2 |
| ARHGEF35-AS1 | 0.000449816 | 0.63795071 | 0.714 | 0.376 | 1 | ARHGEF35-AS1 | 2 |
| ZNF282 | 0.000356162 | 0.635733708 | 0.679 | 0.301 | 1 | ZNF282 | 2 |
| APOE.1 | 0.003836744 | 0.634866702 | 1 | 0.805 | 1 | APOE | 2 |
| STK10 | 0.000121281 | 0.633659004 | 1 | 0.85 | 1 | STK10 | 2 |
| EXT2.1 | 0.002269487 | 0.63286203 | 0.857 | 0.549 | 1 | EXT2 | 2 |
| ABCA8 | 1.01E-06 | 0.632360906 | 0.75 | 0.226 | 0.021985434 | ABCA8 | 2 |
| BCAP29 | 0.000150178 | 0.631978367 | 0.893 | 0.496 | 1 | BCAP29 | 2 |
| AKAP6.1 | 1.96E-06 | 0.626557532 | 0.857 | 0.376 | 0.04284807 | AKAP6 | 2 |
| SKP1.1 | 0.001511982 | 0.62190191 | 1 | 0.955 | 1 | SKP1 | 2 |
| VWA2 | 0.001570239 | 0.621155941 | 0.429 | 0.128 | 1 | VWA2 | 2 |
| COL4A2 | 0.000346849 | 0.617251743 | 0.821 | 0.436 | 1 | COL4A2 | 2 |
| SLC1A4 | 0.005940565 | 0.616471648 | 0.75 | 0.459 | 1 | SLC1A4 | 2 |
| FAM95B1 | 6.80E-06 | 0.615984204 | 0.857 | 0.368 | 0.14835896 | FAM95B1 | 2 |
| TMED4.1 | 0.000422267 | 0.614597873 | 0.857 | 0.639 | 1 | TMED4 | 2 |
| DDR2 | 0.000360976 | 0.613581889 | 1 | 0.737 | 1 | DDR2 | 2 |
| MARCKSL1.1 | 0.007632763 | 0.612650687 | 0.964 | 0.759 | 1 | MARCKSL1 | 2 |
| SEMA4B | 6.05E-06 | 0.611184919 | 0.714 | 0.233 | 0.132098345 | SEMA4B | 2 |
| SEMA3E | 2.14E-08 | 0.610367934 | 0.5 | 0.045 | 0.00046641 | SEMA3E | 2 |
| PTPRS.1 | 0.002907144 | 0.607432611 | 0.964 | 0.767 | 1 | PTPRS | 2 |
| DCHS1 | 7.32E-06 | 0.607080289 | 0.643 | 0.188 | 0.159851356 | DCHS1 | 2 |
| RPS6KA3 | 3.51E-06 | 0.606396955 | 0.964 | 0.602 | 0.076619722 | RPS6KA3 | 2 |
| TNFRSF1A | 1.91E-05 | 0.602718171 | 0.857 | 0.436 | 0.415938704 | TNFRSF1A | 2 |
| PEPD.1 | 0.005104433 | 0.602517623 | 0.857 | 0.564 | 1 | PEPD | 2 |
| IGFBP4.1 | 0.000318259 | 0.601744383 | 0.821 | 0.459 | 1 | IGFBP4 | 2 |
| NOB1.1 | 0.000733582 | 0.598051764 | 0.821 | 0.444 | 1 | NOB1 | 2 |
| GDF11 | 0.000829249 | 0.596889701 | 0.929 | 0.602 | 1 | GDF11 | 2 |
| NPEPL1.1 | 7.65E-05 | 0.594544957 | 0.893 | 0.481 | 1 | NPEPL1 | 2 |
| WDR83OS.1 | 0.002803786 | 0.593988568 | 0.786 | 0.459 | 1 | WDR83OS | 2 |
| HPGD | 0.000776206 | 0.5939465 | 0.643 | 0.278 | 1 | HPGD | 2 |
| CEROX1 | 3.56E-06 | 0.593933592 | 0.679 | 0.195 | 0.077711418 | CEROX1 | 2 |
| ETV5 | 0.001007642 | 0.593767772 | 1 | 0.887 | 1 | ETV5 | 2 |
| DENND2B.1 | 0.000757002 | 0.591783254 | 0.964 | 0.782 | 1 | DENND2B | 2 |
| PTPRG.1 | 6.46E-05 | 0.590891053 | 0.964 | 0.624 | 1 | PTPRG | 2 |
| FRAS1 | 0.00328121 | 0.590344294 | 0.571 | 0.256 | 1 | FRAS1 | 2 |
| RGS2 | 0.000390694 | 0.585775132 | 0.464 | 0.12 | 1 | RGS2 | 2 |
| SASH1 | 0.008654752 | 0.58127898 | 0.893 | 0.722 | 1 | SASH1 | 2 |
| TMEM98.1 | 6.79E-05 | 0.580836386 | 0.893 | 0.511 | 1 | TMEM98 | 2 |
| FAM160B2.1 | 0.003524502 | 0.580543142 | 0.821 | 0.549 | 1 | FAM160B2 | 2 |
| RNF114 | 0.000436222 | 0.578930254 | 0.964 | 0.707 | 1 | RNF114 | 2 |
| PPP2R2B | 7.34E-05 | 0.577789945 | 0.464 | 0.098 | 1 | PPP2R2B | 2 |
| MCHR1 | 0.001163058 | 0.577680293 | 0.393 | 0.098 | 1 | MCHR1 | 2 |
| SNED1 | 0.000564878 | 0.577428148 | 0.643 | 0.301 | 1 | SNED1 | 2 |
| SPTBN1.1 | 0.004328208 | 0.575071604 | 1 | 0.977 | 1 | SPTBN1 | 2 |
| CAPN5 | 0.000544138 | 0.572742116 | 0.75 | 0.406 | 1 | CAPN5 | 2 |
| CDH19 | 6.44E-05 | 0.569721268 | 1 | 0.714 | 1 | CDH19 | 2 |
| PCDHGC3.1 | 0.002329667 | 0.567586731 | 1 | 0.842 | 1 | PCDHGC3 | 2 |
| SMARCD2 | 0.00012797 | 0.567186738 | 0.964 | 0.624 | 1 | SMARCD2 | 2 |

| | | | | | | | |
|---|---|---|---|---|---|---|---|
| BAG6.1 | 0.007619186 | 0.56371373 | 1 | 0.88 | 1 | BAG6 | 2 |
| TGFBRAP1 | 0.008146754 | 0.56125612 | 0.857 | 0.594 | 1 | TGFBRAP1 | 2 |
| CDKN2B | 0.00287979 | 0.56090756 | 0.25 | 0.286 | 1 | CDKN2B | 2 |
| PTPMT1 | 0.0009565 | 0.558109195 | 0.607 | 0.241 | 1 | PTPMT1 | 2 |
| DROSHA | 0.000321938 | 0.556390323 | 0.893 | 0.624 | 1 | DROSHA | 2 |
| TCN2 | 0.007616005 | 0.555802936 | 0.429 | 0.18 | 1 | TCN2 | 2 |
| ENPP2 | 1.00E-04 | 0.552978753 | 0.964 | 0.639 | 1 | ENPP2 | 2 |
| AC020907.1 | 0.000103012 | 0.552231966 | 0.714 | 0.286 | 1 | AC020907.1 | 2 |
| SSH3 | 0.002085359 | 0.55037598 | 0.536 | 0.203 | 1 | SSH3 | 2 |
| SIRT7 | 0.001624328 | 0.549529474 | 0.536 | 0.203 | 1 | SIRT7 | 2 |
| HPN | 1.31E-05 | 0.547569953 | 0.571 | 0.15 | 0.286774786 | HPN | 2 |
| RCN1.1 | 0.000961898 | 0.544197103 | 1 | 0.767 | 1 | RCN1 | 2 |
| ANKHD1.1 | 0.000731593 | 0.541846964 | 1 | 0.805 | 1 | ANKHD1 | 2 |
| RBM14 | 0.005782266 | 0.541809862 | 0.893 | 0.617 | 1 | RBM14 | 2 |
| MINK1.1 | 0.00013414 | 0.539545067 | 1 | 0.729 | 1 | MINK1 | 2 |
| ATP1B2 | 2.42E-06 | 0.532562187 | 0.5 | 0.083 | 0.052822756 | ATP1B2 | 2 |
| BOC.1 | 5.21E-08 | 0.528037263 | 0.786 | 0.226 | 0.001138012 | BOC | 2 |
| PIR | 0.008674347 | 0.527593159 | 0.893 | 0.662 | 1 | PIR | 2 |
| SESN3 | 0.002923902 | 0.524259259 | 0.893 | 0.617 | 1 | SESN3 | 2 |
| ITM2C.1 | 0.0024001 | 0.523046353 | 0.929 | 0.669 | 1 | ITM2C | 2 |
| PTPRZ1 | 0.000349596 | 0.520181124 | 0.964 | 0.662 | 1 | PTPRZ1 | 2 |
| DNAH9 | 9.55E-05 | 0.519914811 | 0.429 | 0.083 | 1 | DNAH9 | 2 |
| GSN-AS1 | 0.000514049 | 0.519585286 | 0.821 | 0.466 | 1 | GSN-AS1 | 2 |
| TRPM4 | 0.001528606 | 0.518590523 | 0.571 | 0.226 | 1 | TRPM4 | 2 |
| STMN3 | 0.001435167 | 0.518331698 | 0.571 | 0.218 | 1 | STMN3 | 2 |
| CLK1.1 | 0.005267475 | 0.516051814 | 1 | 0.835 | 1 | CLK1 | 2 |
| CEP170B | 0.00334781 | 0.512838032 | 0.821 | 0.526 | 1 | CEP170B | 2 |
| GDNF | 0.00344724 | 0.509414849 | 0.25 | 0.045 | 1 | GDNF | 2 |
| SBSN | 0.007355359 | 0.509265911 | 0.143 | 0.008 | 1 | SBSN | 2 |
| MAP2 | 0.00014175 | 0.508925391 | 0.893 | 0.541 | 1 | MAP2 | 2 |
| TRAM2.1 | 0.002964334 | 0.504074108 | 0.929 | 0.632 | 1 | TRAM2 | 2 |
| FBN1 | 0.002877443 | 0.502638731 | 0.643 | 0.338 | 1 | FBN1 | 2 |
| EYA1 | 0.000634402 | 0.500309618 | 0.464 | 0.135 | 1 | EYA1 | 2 |
| BCAP31.1 | 0.009930803 | 0.49978854 | 0.893 | 0.692 | 1 | BCAP31 | 2 |
| MT1E | 4.74E-05 | 0.499684798 | 0.643 | 0.301 | 1 | MT1E | 2 |
| FAXDC2 | 0.006994537 | 0.499403687 | 0.857 | 0.571 | 1 | FAXDC2 | 2 |
| DBN1 | 0.004371077 | 0.498731394 | 0.821 | 0.519 | 1 | DBN1 | 2 |
| KCTD20.1 | 0.002552996 | 0.498529912 | 0.964 | 0.707 | 1 | KCTD20 | 2 |
| SLC35E2B.1 | 0.001307146 | 0.497782888 | 0.964 | 0.677 | 1 | SLC35E2B | 2 |
| NDRG2.1 | 7.27E-05 | 0.492046062 | 0.821 | 0.383 | 1 | NDRG2 | 2 |
| TMEM204.1 | 0.003651131 | 0.490684082 | 0.571 | 0.241 | 1 | TMEM204 | 2 |
| CCDC80 | 0.005248782 | 0.490160735 | 0.786 | 0.474 | 1 | CCDC80 | 2 |
| GNG7 | 0.002982753 | 0.490090838 | 0.857 | 0.534 | 1 | GNG7 | 2 |
| CREB5 | 3.65E-05 | 0.489923197 | 0.786 | 0.338 | 0.79652576 | CREB5 | 2 |
| AHI1 | 0.001802095 | 0.487974079 | 0.75 | 0.391 | 1 | AHI1 | 2 |
| OPN3 | 7.93E-06 | 0.483350891 | 0.857 | 0.451 | 0.17299175 | OPN3 | 2 |
| GJB6 | 0.000114481 | 0.482155879 | 0.357 | 0.06 | 1 | GJB6 | 2 |
| IFIT3 | 0.006265382 | 0.481696976 | 0.714 | 0.398 | 1 | IFIT3 | 2 |
| AHR | 3.74E-05 | 0.478812247 | 0.893 | 0.496 | 0.816250066 | AHR | 2 |
| CAPZB | 0.000902683 | 0.474471243 | 0.964 | 0.714 | 1 | CAPZB | 2 |
| FAM167B | 0.003283521 | 0.47364751 | 0.429 | 0.135 | 1 | FAM167B | 2 |
| DMD | 0.000788717 | 0.472232541 | 0.714 | 0.361 | 1 | DMD | 2 |
| MZT2A | 0.002576295 | 0.470307471 | 0.286 | 0.06 | 1 | MZT2A | 2 |
| SRGAP1 | 0.002931476 | 0.469404675 | 1 | 0.82 | 1 | SRGAP1 | 2 |
| ZCRB1 | 0.003830791 | 0.467534598 | 0.893 | 0.602 | 1 | ZCRB1 | 2 |
| SPTBN2 | 0.00075574 | 0.467469996 | 0.643 | 0.278 | 1 | SPTBN2 | 2 |
| MTSS2.1 | 0.000363472 | 0.466192445 | 0.893 | 0.526 | 1 | MTSS2 | 2 |
| KLHL26 | 0.004048497 | 0.46525129 | 0.321 | 0.083 | 1 | KLHL26 | 2 |
| TARS1.1 | 0.005419933 | 0.465075476 | 0.857 | 0.594 | 1 | TARS1 | 2 |
| UBE2Z | 0.001499928 | 0.463886484 | 1 | 0.774 | 1 | UBE2Z | 2 |
| CWC15.1 | 0.000326139 | 0.462983063 | 0.964 | 0.632 | 1 | CWC15 | 2 |
| AZGP1 | 1.04E-07 | 0.461064962 | 0.893 | 0.331 | 0.002267337 | AZGP1 | 2 |
| YWHAE | 0.002092529 | 0.460883363 | 1 | 0.94 | 1 | YWHAE | 2 |
| SHROOM4.1 | 0.001576254 | 0.45973503 | 0.714 | 0.346 | 1 | SHROOM4 | 2 |
| BMPR2.1 | 0.001135879 | 0.459528676 | 0.964 | 0.699 | 1 | BMPR2 | 2 |
| TIMM50 | 0.004122218 | 0.459460053 | 0.857 | 0.541 | 1 | TIMM50 | 2 |
| CYB5B.1 | 0.00351723 | 0.455714565 | 0.893 | 0.654 | 1 | CYB5B | 2 |
| TMBIM4.1 | 0.000399904 | 0.4533379 | 0.857 | 0.526 | 1 | TMBIM4 | 2 |
| SCN1B | 2.52E-06 | 0.452626339 | 0.714 | 0.211 | 0.054997858 | SCN1B | 2 |
| FNIP2 | 0.002999572 | 0.451489372 | 0.893 | 0.586 | 1 | FNIP2 | 2 |
| PRICKLE2.1 | 0.000813908 | 0.447334657 | 0.75 | 0.376 | 1 | PRICKLE2 | 2 |
| TTC19.1 | 0.000140804 | 0.445955438 | 0.857 | 0.504 | 1 | TTC19 | 2 |
| PDGFD | 2.48E-06 | 0.445846771 | 0.571 | 0.113 | 0.054189357 | PDGFD | 2 |
| CTSK | 0.009138041 | 0.444805998 | 0.929 | 0.737 | 1 | CTSK | 2 |
| MAPRE2.1 | 0.000948817 | 0.444621797 | 0.857 | 0.504 | 1 | MAPRE2 | 2 |
| THUMPD3.1 | 0.002954021 | 0.443573455 | 0.857 | 0.526 | 1 | THUMPD3 | 2 |
| TRIB1.1 | 0.006048726 | 0.443125733 | 0.964 | 0.729 | 1 | TRIB1 | 2 |
| MADD.1 | 0.003291988 | 0.442491338 | 0.964 | 0.699 | 1 | MADD | 2 |
| SLC39A11.1 | 0.000526505 | 0.440924832 | 0.679 | 0.286 | 1 | SLC39A11 | 2 |
| ITM2B | 0.004364612 | 0.437353464 | 1 | 0.857 | 1 | ITM2B | 2 |
| ECHDC2 | 5.27E-07 | 0.435765488 | 0.964 | 0.474 | 0.011499084 | ECHDC2 | 2 |
| ABCA9 | 1.71E-09 | 0.435214199 | 0.679 | 0.105 | 3.74E-05 | ABCA9 | 2 |
| SETX | 0.001857072 | 0.434425159 | 1 | 0.782 | 1 | SETX | 2 |
| AP2B1.1 | 0.006412921 | 0.433407977 | 0.929 | 0.699 | 1 | AP2B1 | 2 |
| LINC01697 | 8.47E-06 | 0.432514912 | 0.464 | 0.075 | 0.184903556 | LINC01697 | 2 |
| PTDSS1 | 0.002688241 | 0.431348533 | 0.964 | 0.722 | 1 | PTDSS1 | 2 |
| FCHSD1 | 0.000384941 | 0.430514531 | 0.75 | 0.346 | 1 | FCHSD1 | 2 |
| SUMF2 | 0.001306407 | 0.429856642 | 1 | 0.782 | 1 | SUMF2 | 2 |
| DLGAP1 | 8.01E-07 | 0.429431121 | 0.714 | 0.195 | 0.017474971 | DLGAP1 | 2 |
| MTUS1 | 6.96E-05 | 0.427593246 | 1 | 0.684 | 1 | MTUS1 | 2 |
| AFAP1 | 0.000653401 | 0.426461031 | 0.786 | 0.414 | 1 | AFAP1 | 2 |
| PCDHGB7 | 0.001110701 | 0.424466457 | 0.464 | 0.143 | 1 | PCDHGB7 | 2 |
| NEURL4.1 | 0.000458481 | 0.423230088 | 0.821 | 0.459 | 1 | NEURL4 | 2 |
| LOXL3 | 0.004481621 | 0.420949255 | 0.536 | 0.218 | 1 | LOXL3 | 2 |
| TOP1MT | 0.001740909 | 0.420636518 | 0.786 | 0.459 | 1 | TOP1MT | 2 |
| ENSA.1 | 0.000231859 | 0.419859382 | 0.964 | 0.632 | 1 | ENSA | 2 |

| | | | | | | | |
|---|---|---|---|---|---|---|---|
| RNF11 | 0.00079791 | 0.419462316 | 0.75 | 0.429 | 1 | RNF11 | 2 |
| FAM234A.1 | 0.000940636 | 0.419401585 | 0.929 | 0.594 | 1 | FAM234A | 2 |
| SSH2 | 0.008734353 | 0.41809607 | 0.786 | 0.534 | 1 | SSH2 | 2 |
| WHRN | 0.000632497 | 0.416962003 | 0.714 | 0.323 | 1 | WHRN | 2 |
| SCAMP4 | 0.007448511 | 0.414053913 | 0.821 | 0.526 | 1 | SCAMP4 | 2 |
| ZBTB20 | 8.11E-05 | 0.412856432 | 0.964 | 0.609 | 1 | ZBTB20 | 2 |
| SNF8 | 0.006282252 | 0.410158733 | 0.786 | 0.474 | 1 | SNF8 | 2 |
| CD55 | 0.001339762 | 0.409820594 | 0.857 | 0.511 | 1 | CD55 | 2 |
| CPNE5 | 0.000937387 | 0.409609429 | 0.464 | 0.135 | 1 | CPNE5 | 2 |
| TSPYL2 | 0.002963764 | 0.40842015 | 1 | 0.797 | 1 | TSPYL2 | 2 |
| PLEKHB2.1 | 0.00569882 | 0.406038562 | 0.929 | 0.707 | 1 | PLEKHB2 | 2 |
| MYEF2.1 | 0.003374795 | 0.405534586 | 0.964 | 0.729 | 1 | MYEF2 | 2 |
| STK32A | 0.000229177 | 0.404996539 | 0.964 | 0.684 | 1 | STK32A | 2 |
| MPZL2 | 0.000173796 | 0.402972989 | 0.5 | 0.128 | 1 | MPZL2 | 2 |
| GTF2A2 | 0.005762607 | 0.400424195 | 0.643 | 0.316 | 1 | GTF2A2 | 2 |
| ATP6V1A | 0.001703451 | 0.400392299 | 0.893 | 0.586 | 1 | ATP6V1A | 2 |
| S100B | 0.001431901 | 0.400057635 | 1 | 0.789 | 1 | S100B | 2 |
| CXXC5.1 | 0.001567768 | 0.396668952 | 0.75 | 0.414 | 1 | CXXC5 | 2 |
| PIP4K2B.1 | 0.00400931 | 0.396072805 | 1 | 0.797 | 1 | PIP4K2B | 2 |
| AHCYL2 | 0.002087606 | 0.395831824 | 0.893 | 0.586 | 1 | AHCYL2 | 2 |
| HSD17B14 | 0.001516984 | 0.394341463 | 0.429 | 0.135 | 1 | HSD17B14 | 2 |
| CCT4.1 | 0.002665175 | 0.393048573 | 0.893 | 0.579 | 1 | CCT4 | 2 |
| GRIP1 | 0.0003263 | 0.392630256 | 0.5 | 0.15 | 1 | GRIP1 | 2 |
| SPOUT1 | 0.009497665 | 0.392232791 | 0.607 | 0.368 | 1 | SPOUT1 | 2 |
| TOLLIP | 0.001262094 | 0.390674351 | 0.893 | 0.556 | 1 | TOLLIP | 2 |
| MEGF9 | 0.002716956 | 0.390336446 | 0.786 | 0.444 | 1 | MEGF9 | 2 |
| ARRDC3.1 | 0.000299472 | 0.388442576 | 0.929 | 0.564 | 1 | ARRDC3 | 2 |
| TTC1.1 | 3.99E-05 | 0.387641414 | 1 | 0.662 | 0.86968851 | TTC1 | 2 |
| INTS9 | 6.24E-07 | 0.385950871 | 0.857 | 0.331 | 0.013621736 | INTS9 | 2 |
| UBE2L6.1 | 0.001697455 | 0.38541829 | 0.821 | 0.466 | 1 | UBE2L6 | 2 |
| SATB1 | 1.89E-06 | 0.385144807 | 0.857 | 0.368 | 0.041190741 | SATB1 | 2 |
| TBCA | 0.009689312 | 0.383869551 | 0.964 | 0.797 | 1 | TBCA | 2 |
| ST3GAL1 | 0.003476774 | 0.381307533 | 0.893 | 0.586 | 1 | ST3GAL1 | 2 |
| SGCE.1 | 0.000850462 | 0.380569719 | 0.893 | 0.564 | 1 | SGCE | 2 |
| PTMS.1 | 0.00370906 | 0.380154083 | 0.821 | 0.489 | 1 | PTMS | 2 |
| TMPRSS9 | 0.000356666 | 0.379848403 | 0.429 | 0.113 | 1 | TMPRSS9 | 2 |
| AP006287.1 | 0.000277747 | 0.379145032 | 0.143 | 0 | 1 | AP006287.1 | 2 |
| KRTCAP2 | 0.000225901 | 0.379035226 | 0.929 | 0.556 | 1 | KRTCAP2 | 2 |
| OPLAH | 0.005609638 | 0.377770277 | 0.393 | 0.12 | 1 | OPLAH | 2 |
| CHL1 | 0.000318239 | 0.377753531 | 1 | 0.744 | 1 | CHL1 | 2 |
| MAF1 | 0.005902831 | 0.375293755 | 0.643 | 0.323 | 1 | MAF1 | 2 |
| CADM1 | 0.000450872 | 0.37356565 | 0.75 | 0.353 | 1 | CADM1 | 2 |
| SLC9A1 | 0.002027694 | 0.373155681 | 0.857 | 0.519 | 1 | SLC9A1 | 2 |
| EPDR1 | 1.75E-06 | 0.37291649 | 0.786 | 0.278 | 0.038259651 | EPDR1 | 2 |
| SP110.1 | 0.007709661 | 0.371874564 | 0.714 | 0.444 | 1 | SP110 | 2 |
| SLC25A12 | 0.001446031 | 0.37173617 | 0.643 | 0.278 | 1 | SLC25A12 | 2 |
| SEZ6L2 | 0.006615887 | 0.370084233 | 0.571 | 0.256 | 1 | SEZ6L2 | 2 |
| PLEKHA4.1 | 0.001412664 | 0.368584463 | 0.929 | 0.617 | 1 | PLEKHA4 | 2 |
| PCDHGA12.1 | 0.001302643 | 0.368196948 | 0.714 | 0.368 | 1 | PCDHGA12 | 2 |
| MIA.1 | 0.006807621 | 0.367054609 | 0.857 | 0.564 | 1 | MIA | 2 |
| CSRP1 | 0.005887644 | 0.36545798 | 1 | 0.88 | 1 | CSRP1 | 2 |
| ATP6V1B2 | 0.00462995 | 0.365411728 | 0.964 | 0.789 | 1 | ATP6V1B2 | 2 |
| HFE.1 | 8.98E-05 | 0.365126149 | 0.714 | 0.278 | 1 | HFE | 2 |
| PDGFA | 1.53E-06 | 0.364969325 | 0.714 | 0.203 | 0.033315607 | PDGFA | 2 |
| INPP5F | 0.001817414 | 0.364595558 | 0.964 | 0.692 | 1 | INPP5F | 2 |
| PIGS | 2.53E-05 | 0.363929656 | 0.893 | 0.444 | 0.552339796 | PIGS | 2 |
| TES | 0.004976203 | 0.363292647 | 0.821 | 0.534 | 1 | TES | 2 |
| CHST11 | 0.003728384 | 0.362441055 | 0.929 | 0.699 | 1 | CHST11 | 2 |
| AC092068.2 | 0.000298949 | 0.357550377 | 0.286 | 0.03 | 1 | AC092068.2 | 2 |
| ACSL3 | 0.009814141 | 0.35714912 | 1 | 0.88 | 1 | ACSL3 | 2 |
| SEPTIN7 | 0.008202885 | 0.356693391 | 1 | 0.85 | 1 | SEPTIN7 | 2 |
| MAOB | 0.000913093 | 0.353767836 | 0.429 | 0.12 | 1 | MAOB | 2 |
| TMEM59.1 | 0.00528461 | 0.353702776 | 0.964 | 0.722 | 1 | TMEM59 | 2 |
| CDK18.1 | 0.001070977 | 0.352527317 | 0.679 | 0.323 | 1 | CDK18 | 2 |
| AL139241.1 | 4.81E-06 | 0.351931613 | 0.607 | 0.143 | 0.104915957 | AL139241.1 | 2 |
| AAMP | 0.003233447 | 0.350840521 | 0.893 | 0.579 | 1 | AAMP | 2 |
| VPS41 | 0.009366385 | 0.350812028 | 0.964 | 0.767 | 1 | VPS41 | 2 |
| GPR107 | 0.008416552 | 0.350654787 | 0.929 | 0.677 | 1 | GPR107 | 2 |
| SERTAD2 | 0.000182735 | 0.349815433 | 0.929 | 0.556 | 1 | SERTAD2 | 2 |
| THBD | 0.001581601 | 0.347827651 | 0.357 | 0.09 | 1 | THBD | 2 |
| BCAR1 | 0.004349471 | 0.347363564 | 0.714 | 0.376 | 1 | BCAR1 | 2 |
| PPP1R12B | 0.004508044 | 0.345917445 | 0.857 | 0.541 | 1 | PPP1R12B | 2 |
| NFATC2 | 0.001327437 | 0.345007693 | 0.857 | 0.504 | 1 | NFATC2 | 2 |
| SLC27A1 | 0.009415284 | 0.344179974 | 0.929 | 0.707 | 1 | SLC27A1 | 2 |
| USP9X.1 | 0.001374566 | 0.344076735 | 1 | 0.767 | 1 | USP9X | 2 |
| SLC39A3 | 0.009070267 | 0.342714078 | 0.214 | 0.038 | 1 | SLC39A3 | 2 |
| BAHD1.1 | 0.001122912 | 0.341296986 | 0.643 | 0.271 | 1 | BAHD1 | 2 |
| SYMPK.1 | 0.00209448 | 0.339066157 | 0.964 | 0.707 | 1 | SYMPK | 2 |
| NCDN | 4.75E-05 | 0.33889764 | 0.821 | 0.391 | 1 | NCDN | 2 |
| MYDGF | 0.008425301 | 0.33836369 | 0.679 | 0.398 | 1 | MYDGF | 2 |
| MOXD1 | 0.000141068 | 0.338219596 | 0.5 | 0.135 | 1 | MOXD1 | 2 |
| GLCE.1 | 0.002175732 | 0.337803892 | 0.75 | 0.398 | 1 | GLCE | 2 |
| ZNF507 | 0.000778854 | 0.335398197 | 0.786 | 0.406 | 1 | ZNF507 | 2 |
| SLC5A3.1 | 0.002299689 | 0.335282178 | 0.857 | 0.526 | 1 | SLC5A3 | 2 |
| GAREM1 | 1.82E-05 | 0.334387603 | 0.714 | 0.256 | 0.397593708 | GAREM1 | 2 |
| FAM174B | 0.001372585 | 0.334349135 | 0.571 | 0.226 | 1 | FAM174B | 2 |
| SUN2 | 0.001208647 | 0.333419258 | 0.857 | 0.526 | 1 | SUN2 | 2 |
| CLPTM1.1 | 0.00325579 | 0.333386771 | 1 | 0.782 | 1 | CLPTM1 | 2 |
| TNS3 | 0.005775653 | 0.332979835 | 1 | 0.805 | 1 | TNS3 | 2 |
| MFSD6 | 0.000201564 | 0.331676714 | 0.786 | 0.391 | 1 | MFSD6 | 2 |
| LDLRAD3.1 | 0.005938184 | 0.329400411 | 0.893 | 0.617 | 1 | LDLRAD3 | 2 |
| ZNF7 | 0.008043734 | 0.329032897 | 0.786 | 0.474 | 1 | ZNF7 | 2 |
| ITFG1 | 0.000913576 | 0.327734748 | 0.857 | 0.496 | 1 | ITFG1 | 2 |
| MYO6 | 0.005108011 | 0.326692311 | 0.5 | 0.203 | 1 | MYO6 | 2 |
| MPDU1 | 0.00648365 | 0.326644473 | 0.75 | 0.444 | 1 | MPDU1 | 2 |
| STXBP4 | 0.000266786 | 0.325829423 | 0.536 | 0.165 | 1 | STXBP4 | 2 |

| | | | | | | | |
|---|---|---|---|---|---|---|---|
| ASB9 | 0.004933576 | 0.324490511 | 0.679 | 0.383 | 1 | ASB9 | 2 |
| DLC1 | 0.002651288 | 0.323310878 | 0.964 | 0.774 | 1 | DLC1 | 2 |
| PPP1R9A.1 | 0.000465532 | 0.320508416 | 0.821 | 0.451 | 1 | PPP1R9A | 2 |
| KDELR2 | 0.009868565 | 0.318769647 | 0.929 | 0.684 | 1 | KDELR2 | 2 |
| NDUFA10.1 | 0.000218442 | 0.318386759 | 0.964 | 0.617 | 1 | NDUFA10 | 2 |
| PDK4 | 8.02E-07 | 0.317589875 | 0.607 | 0.12 | 0.017503301 | PDK4 | 2 |
| CICP14 | 0.000384343 | 0.315853509 | 0.75 | 0.368 | 1 | CICP14 | 2 |
| ALKBH5 | 0.006711003 | 0.315760406 | 0.786 | 0.474 | 1 | ALKBH5 | 2 |
| SLC39A14.1 | 0.001782298 | 0.314755481 | 0.786 | 0.444 | 1 | SLC39A14 | 2 |
| GDF15 | 0.000355912 | 0.31311191 | 0.714 | 0.308 | 1 | GDF15 | 2 |
| HEXIM1.1 | 0.004797578 | 0.312355813 | 0.929 | 0.662 | 1 | HEXIM1 | 2 |
| LYNX1 | 2.71E-06 | 0.312128572 | 0.464 | 0.06 | 0.059226657 | LYNX1 | 2 |
| FAF2.1 | 0.002051194 | 0.311736706 | 0.893 | 0.564 | 1 | FAF2 | 2 |
| GRIA4 | 6.40E-05 | 0.311056846 | 0.321 | 0.03 | 1 | GRIA4 | 2 |
| STING1 | 0.001502345 | 0.309300731 | 0.786 | 0.421 | 1 | STING1 | 2 |
| INPP1.1 | 0.001358241 | 0.308559216 | 0.821 | 0.459 | 1 | INPP1 | 2 |
| MINDY1 | 4.23E-05 | 0.308530669 | 0.786 | 0.331 | 0.922944573 | MINDY1 | 2 |
| ADGRL1 | 0.004412817 | 0.308124483 | 0.75 | 0.421 | 1 | ADGRL1 | 2 |
| MKRN1 | 0.004967795 | 0.305174794 | 0.893 | 0.609 | 1 | MKRN1 | 2 |
| KDM5C.1 | 0.001871837 | 0.304423781 | 1 | 0.797 | 1 | KDM5C | 2 |
| RHOJ | 0.004871075 | 0.301401102 | 0.857 | 0.556 | 1 | RHOJ | 2 |
| EMG1 | 0.000232903 | 0.299936722 | 0.679 | 0.293 | 1 | EMG1 | 2 |
| DHRS11 | 0.002005818 | 0.299285496 | 0.607 | 0.256 | 1 | DHRS11 | 2 |
| PLXND1 | 0.003935867 | 0.296833375 | 0.893 | 0.624 | 1 | PLXND1 | 2 |
| PROS1.1 | 6.81E-06 | 0.295855834 | 0.821 | 0.331 | 0.148656458 | PROS1 | 2 |
| PMS2 | 0.006670273 | 0.295434049 | 0.786 | 0.466 | 1 | PMS2 | 2 |
| FREM1 | 0.007492574 | 0.295341128 | 0.429 | 0.165 | 1 | FREM1 | 2 |
| CAVIN1 | 3.03E-05 | 0.295193246 | 0.893 | 0.451 | 0.661079338 | CAVIN1 | 2 |
| SRPRA.1 | 0.001812493 | 0.29512537 | 1 | 0.789 | 1 | SRPRA | 2 |
| PCBP4.1 | 0.007782964 | 0.29401109 | 0.643 | 0.331 | 1 | PCBP4 | 2 |
| PLEKHA2 | 0.007525231 | 0.29398432 | 0.964 | 0.797 | 1 | PLEKHA2 | 2 |
| CHRDL1 | 7.75E-08 | 0.293853539 | 0.429 | 0.023 | 0.001691369 | CHRDL1 | 2 |
| MED10 | 0.000517211 | 0.292258675 | 0.571 | 0.195 | 1 | MED10 | 2 |
| TFIP11.1 | 0.001734044 | 0.288635205 | 0.75 | 0.391 | 1 | TFIP11 | 2 |
| HOGA1 | 0.005674222 | 0.288012331 | 0.393 | 0.12 | 1 | HOGA1 | 2 |
| LAMB1.1 | 0.007442221 | 0.285825314 | 0.964 | 0.737 | 1 | LAMB1 | 2 |
| CAPN11 | 0.002743275 | 0.284385333 | 0.214 | 0.023 | 1 | CAPN11 | 2 |
| FNDC3A | 0.001119201 | 0.284360778 | 1 | 0.774 | 1 | FNDC3A | 2 |
| EI24.1 | 0.00912593 | 0.284159274 | 0.964 | 0.767 | 1 | EI24 | 2 |
| SIPA1L2 | 0.000206619 | 0.282168994 | 0.893 | 0.496 | 1 | SIPA1L2 | 2 |
| PAFAH1B2 | 0.00721408 | 0.281376522 | 0.857 | 0.571 | 1 | PAFAH1B2 | 2 |
| SORBS2 | 0.001501591 | 0.280885799 | 0.714 | 0.346 | 1 | SORBS2 | 2 |
| C2CD2.1 | 0.000411068 | 0.28019812 | 0.679 | 0.278 | 1 | C2CD2 | 2 |
| CSPG5 | 0.000474283 | 0.2778917 | 0.429 | 0.105 | 1 | CSPG5 | 2 |
| CDH1 | 0.001265857 | 0.276096491 | 1 | 0.759 | 1 | CDH1 | 2 |
| BNIP1 | 1.09E-05 | 0.273919321 | 0.393 | 0.045 | 0.237283839 | BNIP1 | 2 |
| LGI4.1 | 8.29E-05 | 0.272940506 | 0.929 | 0.526 | 1 | LGI4 | 2 |
| CHPT1 | 1.13E-05 | 0.272181704 | 0.821 | 0.346 | 0.246577279 | CHPT1 | 2 |
| RXRG | 0.002426696 | 0.271444395 | 0.679 | 0.323 | 1 | RXRG | 2 |
| OR2A9P | 6.35E-08 | 0.270374415 | 0.714 | 0.158 | 0.001386432 | OR2A9P | 2 |
| TMEM178B | 0.009061452 | 0.270295435 | 0.536 | 0.241 | 1 | TMEM178B | 2 |
| FHAD1 | 0.007044858 | 0.267289059 | 0.214 | 0.03 | 1 | FHAD1 | 2 |
| UBQLN2 | 0.003345476 | 0.267145249 | 0.714 | 0.368 | 1 | UBQLN2 | 2 |
| MX2 | 0.008336312 | 0.266549677 | 0.714 | 0.414 | 1 | MX2 | 2 |
| NUP205 | 0.001327411 | 0.265424674 | 0.964 | 0.669 | 1 | NUP205 | 2 |
| SLC9A3 | 0.000336926 | 0.264795737 | 0.571 | 0.203 | 1 | SLC9A3 | 2 |
| BTN3A3.1 | 0.002334956 | 0.264747822 | 0.857 | 0.519 | 1 | BTN3A3 | 2 |
| AHCTF1 | 0.009640621 | 0.264661321 | 0.786 | 0.496 | 1 | AHCTF1 | 2 |
| RTN4RL1 | 0.000111778 | 0.263859822 | 0.464 | 0.113 | 1 | RTN4RL1 | 2 |
| ARHGEF37 | 0.000208056 | 0.263848922 | 0.643 | 0.233 | 1 | ARHGEF37 | 2 |
| AC138409.1 | 0.001324843 | 0.26322572 | 0.786 | 0.421 | 1 | AC138409.1 | 2 |
| PRKRIP1 | 0.007733581 | 0.263141372 | 0.821 | 0.519 | 1 | PRKRIP1 | 2 |
| GNG11 | 3.22E-05 | 0.262400144 | 0.679 | 0.226 | 0.702701412 | GNG11 | 2 |
| FAM20A.1 | 0.00102569 | 0.262230793 | 0.429 | 0.113 | 1 | FAM20A | 2 |
| STARD7 | 0.004957398 | 0.261787298 | 0.893 | 0.609 | 1 | STARD7 | 2 |
| PLRG1 | 0.001383951 | 0.260014709 | 0.75 | 0.383 | 1 | PLRG1 | 2 |
| IL27RA | 0.002287979 | 0.259160275 | 0.464 | 0.165 | 1 | IL27RA | 2 |
| RNF216 | 0.002457986 | 0.258793849 | 0.929 | 0.639 | 1 | RNF216 | 2 |
| HIBCH | 0.001421551 | 0.258468983 | 0.964 | 0.677 | 1 | HIBCH | 2 |
| LRRTM4 | 0.006792071 | 0.257122325 | 0.464 | 0.173 | 1 | LRRTM4 | 2 |
| DNHD1.1 | 4.29E-05 | 0.256953789 | 0.786 | 0.331 | 0.93591187 | DNHD1 | 2 |
| C1QTNF1 | 6.32E-05 | 0.256849334 | 0.5 | 0.113 | 1 | C1QTNF1 | 2 |
| MYBBP1A | 0.007912281 | 0.256441307 | 0.857 | 0.564 | 1 | MYBBP1A | 2 |
| OR2A20P | 8.02E-07 | 0.256358109 | 0.643 | 0.143 | 0.017497797 | OR2A20P | 2 |
| MIEF2 | 0.008532017 | 0.256290832 | 0.321 | 0.09 | 1 | MIEF2 | 2 |
| TUSC3.1 | 1.08E-05 | 0.255377618 | 0.893 | 0.429 | 0.235731957 | TUSC3 | 2 |
| GARRE1 | 0.005791015 | 0.255017883 | 0.821 | 0.504 | 1 | GARRE1 | 2 |
| AC091390.4 | 0.000889167 | 0.254885476 | 0.571 | 0.211 | 1 | AC091390.4 | 2 |
| CREB3L2.1 | 0.008400084 | 0.254714484 | 0.929 | 0.669 | 1 | CREB3L2 | 2 |
| MRPL15 | 0.004801846 | 0.253820494 | 0.607 | 0.286 | 1 | MRPL15 | 2 |
| ZHX1.1 | 0.000193109 | 0.253255597 | 0.893 | 0.496 | 1 | ZHX1 | 2 |
| VPS4A.1 | 0.000297733 | 0.253052797 | 0.929 | 0.564 | 1 | VPS4A | 2 |
| ADSS2 | 0.007926779 | 0.25164128 | 0.929 | 0.669 | 1 | ADSS2 | 2 |
| FURIN.1 | 0.001554033 | 0.250698436 | 0.786 | 0.421 | 1 | FURIN | 2 |
