## Supplementary Table 4 for "Metastasis founder cells activate immunosuppression early in human melanoma metastatic colonization"

| Protein | MelDCC_10 100K_01a | MelDCC_10 100K_01b | MelDCC_10 100K_02a | MelDCC_10 100K_02b | Protein | MW | SD | RSD | Entry | Entry Name | Entry name | Protein names | Gene names |
|---|---|---|---|---|---|---|---|---|---|---|---|---|---|
| sp\|Q15149\|PLEC_HUMAN | 480,513 | 407,895 | 810,309 | 769,772 | sp\|Q15149\|PLEC_HUMAN | 498,208 | 198,295 | 40% | Q15149 | PLEC_HUMAN | PLEC_HUMAN | Plectin (PCN) (PLTN) (Hemidesmosomal protein 1) (HD1) (Plectin-1) | PLEC PLEC1 |
| sp\|Q14204\|DYHC1_HUMAN | 758,780 | 939,313 | 1,167,799 | 978,511 | sp\|Q14204\|DYHC1_HUMAN | 795,248 | 214,343 | 27% | Q14204 | DYHC1_HUMAN | DYHC1_HUMAN | Cytoplasmic dynein 1 heavy chain 1 (Cytoplasmic dynein heavy chain 1) (Dynein heavy chain, cytosolic) | DYNC1H1 DHC1 DNCH1 DNCL DNECL DYHC KIAA0325 |
| sp\|P49327\|FAS_HUMAN | 2,457,634 | 2,667,144 | 2,663,888 | 3,522,243 | sp\|P49327\|FAS_HUMAN | 2,246,221 | 742,810 | 33% | P49327 | FAS_HUMAN | FAS_HUMAN | Fatty acid synthase (EC 2.3.1.85) (Type I fatty acid synthase) [Includes: [Acyl-carrier-protein] S-acetyltransferase (EC 2.3.1.38); [Acyl-carrier-protein] S-malonyltransferase (EC 2.3.1.39); 3-oxoacyl-[acyl-carrier-protein] synthase (EC 2.3.1.41); 3-oxoacyl-[acyl-carrier-protein] reductase (EC 1.1.1.100); 3-hydroxyacyl-[acyl-carrier-protein] dehydratase (EC 4.2.1.59); Enoyl-[acyl-carrier-protein] reductase (EC 1.3.1.39); Acyl-[acyl-carrier-protein] hydrolase (EC 3.1.2.14)] | FASN FAS |
| sp\|P35579\|MYH9_HUMAN | 1,678,226 | 1,909,809 | 1,321,321 | 1,785,295 | sp\|P35579\|MYH9_HUMAN | 1,402,067 | 339,535 | 24% | P35579 | MYH9_HUMAN | MYH9_HUMAN | Myosin-9 (Cellular myosin heavy chain, type A) (Myosin heavy chain 9) (Myosin heavy chain, non-muscle IIa) (Non-muscle myosin heavy chain A) (NMMHC-A) (Non-muscle myosin heavy chain IIa) (NMMHC II-a) (NMMHC-IIA) | MYH9 |
| sp\|P24821\|TENA_HUMAN | 6,168,101 | 6,292,763 | 6,198,642 | 6,866,582 | sp\|P24821\|TENA_HUMAN | 6,541,956 | 724,917 | 11% | P24821 | TENA_HUMAN | TENA_HUMAN | Tenascin (TN) (Cytotactin) (GMEM) (GP 150-225) (Glioma-associated-extracellular matrix antigen) (Hexabrachion) (JI) (Myotendinous antigen) (Neuronectin) (Tenascin-C) (TN-C) | TNC HXB |
| sp\|Q00610\|CLH1_HUMAN | 2,458,702 | 2,380,338 | 3,135,943 | 3,482,754 | sp\|Q00610\|CLH1_HUMAN | 2,608,488 | 589,359 | 23% | Q00610 | CLH1_HUMAN | CLH1_HUMAN | Clathrin heavy chain 1 (Clathrin heavy chain on chromosome 17) (CLH-17) | CLTC CLH17 CLTCL2 KIAA0034 |
| sp\|P98160\|PGBM_HUMAN | 3,040,182 | 2,217,206 | 4,925,559 | 3,582,379 | sp\|P98160\|PGBM_HUMAN | 3,204,053 | 875,940 | 27% | P98160 | PGBM_HUMAN | PGBM_HUMAN | Basement membrane-specific heparan sulfate proteoglycan core protein (HSPG) (Perlecan) (PLC) [Cleaved into: Endorepellin; LG3 peptide] | HSPG2 |
| sp\|Q6UVK1\|CSPG4_HUMAN | 6,792,458 | 6,268,413 | 10,903,839 | 10,329,746 | sp\|Q6UVK1\|CSPG4_HUMAN | 7,863,292 | 2,023,857 | 26% | Q6UVK1 | CSPG4_HUMAN | CSPG4_HUMAN | Chondroitin sulfate proteoglycan 4 (Chondroitin sulfate proteoglycan NG2) (Melanoma chondroitin sulfate proteoglycan) (Melanoma-associated chondroitin sulfate proteoglycan) | CSPG4 MCSP |
| sp\|O43707\|ACTN4_HUMAN | 1,184,508 | 1,919,656 | 665,711 | 720,523 | sp\|O43707\|ACTN4_HUMAN | 932,689 | 522,022 | 56% | O43707 | ACTN4_HUMAN | ACTN4_HUMAN | Alpha-actinin-4 (Non-muscle alpha-actinin 4) | ACTN4 |
| sp\|P21333\|FLNA_HUMAN | 578,113 | 831,050 | 598,207 | 789,196 | sp\|P21333\|FLNA_HUMAN | 834,020 | 189,929 | 23% | P21333 | FLNA_HUMAN | FLNA_HUMAN | Filamin-A (FLN-A) (Actin-binding protein 280) (ABP-280) (Alpha-filamin) (Endothelial actin-binding protein) (Filamin-1) (Non-muscle filamin) | FLNA FLN FLN1 |
| sp\|P46940\|IQGA1_HUMAN | 1,573,325 | 1,980,967 | 2,549,295 | 2,725,098 | sp\|P46940\|IQGA1_HUMAN | 1,916,835 | 519,627 | 27% | P46940 | IQGA1_HUMAN | IQGA1_HUMAN | Ras GTPase-activating-like protein IQGAP1 (p195) | IQGAP1 KIAA0051 |
| sp\|P08238\|HS90B_HUMAN | 4,963,524 | 5,313,088 | 3,906,588 | 4,223,299 | sp\|P08238\|HS90B_HUMAN | 3,816,806 | 973,716 | 26% | P08238 | HS90B_HUMAN | HS90B_HUMAN | Heat shock protein HSP 90-beta (HSP 90) (Heat shock 84 kDa) (HSP 84) (HSP84) | HSP90AB1 HSP90B HSPC2 HSPCB |

| | | | | | | | | | | | | | |
|---|---|---|---|---|---|---|---|---|---|---|---|---|---|
| sp\|P08670\|VIME_HUMAN | 2,797,422 | 2,977,716 | 6,735,817 | 7,904,613 | sp\|P08670\|VIME_HUMAN | 3,908,352 | 2,188,403 | 56% | P08670 | VIME_HUMAN | VIME_HUMAN | Vimentin | VIM |
| sp\|P42704\|LPPRC_HUMAN | 32,728 | 37,198 | 59,746 | 23,003 | sp\|P42704\|LPPRC_HUMAN | 31,076 | 13,036 | 42% | P42704 | LPPRC_HUMAN | LPPRC_HUMAN | Leucine-rich PPR motif-containing protein, mitochondrial (130 kDa leucine-rich protein) (LRP 130) (GP130) | LRPPRC LRP130 |
| sp\|Q9Y490\|TLN1_HUMAN | 525,241 | 724,484 | 502,530 | 567,395 | sp\|Q9Y490\|TLN1_HUMAN | 783,456 | 279,622 | 36% | Q9Y490 | TLN1_HUMAN | TLN1_HUMAN | Talin-1 | TLN1 KIAA1027 TLN |
| sp\|P05023\|AT1A1_HUMAN | 10,695,496 | 10,611,811 | 13,736,968 | 14,052,902 | sp\|P05023\|AT1A1_HUMAN | 11,003,190 | 2,049,372 | 19% | P05023 | AT1A1_HUMAN | AT1A1_HUMAN | Sodium/potassium-transporting ATPase subunit alpha-1 (Na(+)/K(+) ATPase alpha-1 subunit) (EC 7.2.2.13) (Sodium pump subunit alpha-1) | ATP1A1 |
| sp\|P11216\|PYGB_HUMAN | 2,073,784 | 1,809,059 | 2,711,637 | 2,838,021 | sp\|P11216\|PYGB_HUMAN | 2,037,402 | 480,866 | 24% | P11216 | PYGB_HUMAN | PYGB_HUMAN | Glycogen phosphorylase, brain form (EC 2.4.1.1) | PYGB |
| sp\|P26038\|MOES_HUMAN | 5,599,948 | 6,791,514 | 5,227,243 | 6,712,522 | sp\|P26038\|MOES_HUMAN | 5,867,057 | 958,942 | 16% | P26038 | MOES_HUMAN | MOES_HUMAN | Moesin (Membrane-organizing extension spike protein) | MSN |
| sp\|O00159\|MYO1C_HUMAN | 5,319,245 | 5,432,543 | 6,663,051 | 7,146,738 | sp\|O00159\|MYO1C_HUMAN | 5,535,138 | 1,023,757 | 18% | O00159 | MYO1C_HUMAN | MYO1C_HUMAN | Unconventional myosin-Ic (Myosin I beta) (MMI-beta) (MMIb) | MYO1C |
| sp\|P13639\|EF2_HUMAN | 2,642,758 | 3,192,620 | 3,117,280 | 3,496,727 | sp\|P13639\|EF2_HUMAN | 2,901,420 | 343,372 | 12% | P13639 | EF2_HUMAN | EF2_HUMAN | Elongation factor 2 (EF-2) | EEF2 EF2 |
| sp\|Q6YHK3\|CD109_HUMAN | 3,095,479 | 3,338,712 | 2,258,879 | 2,768,530 | sp\|Q6YHK3\|CD109_HUMAN | 2,408,283 | 677,542 | 28% | Q6YHK3 | CD109_HUMAN | CD109_HUMAN | CD109 antigen (150 kDa TGF-beta-1-binding protein) (C3 and PZP-like alpha-2-macroglobulin domain-containing protein 7) (Platelet-specific Gov antigen) (p180) (r150) (CD antigen CD109) | CD109 CPAMD7 |
| sp\|P06756\|ITAV_HUMAN | 3,221,666 | 3,452,075 | 4,140,008 | 4,327,933 | sp\|P06756\|ITAV_HUMAN | 3,215,445 | 793,278 | 25% | P06756 | ITAV_HUMAN | ITAV_HUMAN | Integrin alpha-V (Vitronectin receptor) (Vitronectin receptor subunit alpha) (CD antigen CD51) [Cleaved into: Integrin alpha-V heavy chain; Integrin alpha-V light chain] | ITGAV MSK8 VNRA VTNR |
| sp\|Q8WUM4\|PDC6I_HUMAN | 4,379,956 | 4,942,476 | 2,314,714 | 2,377,555 | sp\|Q8WUM4\|PDC6I_HUMAN | 2,851,956 | 1,273,454 | 45% | Q8WUM4 | PDC6I_HUMAN | PDC6I_HUMAN | Programmed cell death 6-interacting protein (PDCD6-interacting protein) (ALG-2-interacting protein 1) (ALG-2-interacting protein X) (Hp95) | PDCD6IP AIP1 ALIX KIAA1375 |
| sp\|P04264\|K2C1_HUMAN | 35,847,265 | 15,929,015 | 24,210,948 | 16,218,337 | sp\|P04264\|K2C1_HUMAN | 26,967,391 | 9,641,768 | 36% | P04264 | K2C1_HUMAN | K2C1_HUMAN | Keratin, type II cytoskeletal 1 (67 kDa cytokeratin) (Cytokeratin-1) (CK-1) (Hair alpha protein) (Keratin-1) (K1) (Type-II keratin Kb1) | KRT1 KRTA |
| sp\|P14618\|KPYM_HUMAN | 21,706,801 | 20,693,657 | 28,401,114 | 18,756,570 | sp\|P14618\|KPYM_HUMAN | 19,223,839 | 4,966,060 | 26% | P14618 | KPYM_HUMAN | KPYM_HUMAN | Pyruvate kinase PKM (EC 2.7.1.40) (Cytosolic thyroid hormone-binding protein) (CTHBP) (Opa-interacting protein 3) (OIP-3) (Pyruvate kinase 2/3) (Pyruvate kinase muscle isozyme) (Threonine-protein kinase PKM2) (EC 2.7.11.1) (Thyroid hormone-binding protein 1) (THBP1) (Tumor M2-PK) (Tyrosine-protein kinase PKM2) (EC 2.7.10.2) (p58) | PKM OIP3 PK2 PK3 PKM2 |
| sp\|P11142\|HSP7C_HUMAN | 10,545,110 | 11,642,220 | 7,648,001 | 8,786,733 | sp\|P11142\|HSP7C_HUMAN | 7,745,745 | 2,584,319 | 33% | P11142 | HSP7C_HUMAN | HSP7C_HUMAN | Heat shock cognate 71 kDa protein (EC 3.6.4.10) (Heat shock 70 kDa protein 8) (Lipopolysaccharide-associated protein 1) (LAP-1) (LPS-associated protein 1) | HSPA8 HSC70 HSP73 HSPA10 |

| | | | | | | | | | | | | | |
|---|---|---|---|---|---|---|---|---|---|---|---|---|---|
| sp\|P10809\|CH60_HUMAN | 524,304 | 661,427 | 290,834 | 366,842 | sp\|P10809\|CH60_HUMAN | 443,364 | 206,684 | 47% | P10809 | CH60_HUMAN | CH60_HUMAN | 60 kDa heat shock protein, mitochondrial (EC 5.6.1.7) (60 kDa chaperonin) (Chaperonin 60) (CPN60) (Heat shock protein 60) (HSP-60) (Hsp60) (HuCHA60) (Mitochondrial matrix protein P1) (P60 lymphocyte protein) | HSPD1 HSP60 |
| sp\|Q7KZF4\|SND1_HUMAN | 617,784 | 762,017 | 1,113,910 | 1,240,968 | sp\|Q7KZF4\|SND1_HUMAN | 770,775 | 276,835 | 36% | Q7KZF4 | SND1_HUMAN | SND1_HUMAN | Staphylococcal nuclease domain-containing protein 1 (EC 3.1.31.1) (100 kDa coactivator) (EBNA2 coactivator p100) (Tudor domain-containing protein 11) (p100 co-activator) | SND1 TDRD11 |
| sp\|P13667\|PDIA4_HUMAN | 482,932 | 603,632 | 505,747 | 466,798 | sp\|P13667\|PDIA4_HUMAN | 394,782 | 137,029 | 35% | P13667 | PDIA4_HUMAN | PDIA4_HUMAN | Protein disulfide-isomerase A4 (EC 5.3.4.1) (Endoplasmic reticulum resident protein 70) (ER protein 70) (ERp70) (Endoplasmic reticulum resident protein 72) (ER protein 72) (ERp-72) (ERp72) | PDIA4 ERP70 ERP72 |
| sp\|P55072\|TERA_HUMAN | 1,618,175 | 1,835,000 | 1,448,645 | 1,601,646 | sp\|P55072\|TERA_HUMAN | 1,345,015 | 321,079 | 24% | P55072 | TERA_HUMAN | TERA_HUMAN | Transitional endoplasmic reticulum ATPase (TER ATPase) (EC 3.6.4.6) (15S Mg(2+)-ATPase p97 subunit) (Valosin-containing protein) (VCP) | VCP |
| sp\|P14625\|ENPL_HUMAN | 712,358 | 791,892 | 566,949 | 672,409 | sp\|P14625\|ENPL_HUMAN | 541,409 | 168,467 | 31% | P14625 | ENPL_HUMAN | ENPL_HUMAN | Endoplasmin (94 kDa glucose-regulated protein) (GRP-94) (Heat shock protein 90 kDa beta member 1) (Tumor rejection antigen 1) (gp96 homolog) | HSP90B1 GRP94 TRA1 |
| sp\|P30101\|PDIA3_HUMAN | 759,039 | 915,989 | 1,144,910 | 1,031,777 | sp\|P30101\|PDIA3_HUMAN | 759,010 | 251,005 | 33% | P30101 | PDIA3_HUMAN | PDIA3_HUMAN | Protein disulfide-isomerase A3 (EC 5.3.4.1) (58 kDa glucose-regulated protein) (58 kDa microsomal protein) (p58) (Disulfide isomerase ER-60) (Endoplasmic reticulum resident protein 57) (ER protein 57) (ERp57) (Endoplasmic reticulum resident protein 60) (ER protein 60) (ERp60) | PDIA3 ERP57 ERP60 GRP58 |
| sp\|P07237\|PDIA1_HUMAN | 766,701 | 873,704 | 223,150 | 505,941 | sp\|P07237\|PDIA1_HUMAN | 478,717 | 230,875 | 48% | P07237 | PDIA1_HUMAN | PDIA1_HUMAN | Protein disulfide-isomerase (PDI) (EC 5.3.4.1) (Cellular thyroid hormone-binding protein) (Prolyl 4-hydroxylase subunit beta) (p55) | P4HB ERBA2L PDI PDIA1 PO4DB |
| sp\|P35527\|K1C9_HUMAN | 18,227,244 | 6,619,674 | 10,321,019 | 5,923,517 | sp\|P35527\|K1C9_HUMAN | 10,984,694 | 4,479,532 | 41% | P35527 | K1C9_HUMAN | K1C9_HUMAN | Keratin, type I cytoskeletal 9 (Cytokeratin-9) (CK-9) (Keratin-9) (K9) | KRT9 |
| sp\|P11021\|GRP78_HUMAN | 965,301 | 1,032,789 | 879,926 | 857,433 | sp\|P11021\|GRP78_HUMAN | 707,650 | 257,229 | 36% | P11021 | GRP78_HUMAN | BIP_HUMAN | Endoplasmic reticulum chaperone BiP (EC 3.6.4.10) (78 kDa glucose-regulated protein) (GRP-78) (Binding-immunoglobulin protein) (BiP) (Heat shock protein 70 family protein 5) (HSP70 family protein 5) (Heat shock protein family A member 5) (Immunoglobulin heavy chain-binding protein) | HSPA5 GRP78 |
| sp\|Q01082\|SPTB2_HUMAN | 232,763 | 359,938 | 527,856 | 566,917 | sp\|Q01082\|SPTB2_HUMAN | 402,131 | 127,135 | 32% | Q01082 | SPTB2_HUMAN | SPTB2_HUMAN | Spectrin beta chain, non-erythrocytic 1 (Beta-II spectrin) (Fodrin beta chain) (Spectrin, non-erythroid beta chain 1) | SPTBN1 SPTB2 |
| sp\|Q14697\|GANAB_HUMAN | 659,158 | 543,883 | 694,847 | 1,014,102 | sp\|Q14697\|GANAB_HUMAN | 607,940 | 202,793 | 33% | Q14697 | GANAB_HUMAN | GANAB_HUMAN | Neutral alpha-glucosidase AB (EC 3.2.1.207) (Alpha-glucosidase 2) (Glucosidase II subunit alpha) | GANAB G2AN KIAA0088 |

| | | | | | | | | | | | | | |
|---|---|---|---|---|---|---|---|---|---|---|---|---|---|
| sp\|P06733\|ENOA_HUMAN | 4,289,638 | 5,611,789 | 2,488,801 | 3,088,973 | sp\|P06733\|ENOA_HUMAN | 3,464,979 | 1,277,973 | 37% | P06733 | ENOA_HUMAN | ENOA_HUMAN | Alpha-enolase (EC 4.2.1.11) (2-phospho-D-glycerate hydro-lyase) (C-myc promoter-binding protein) (Enolase 1) (MBP-1) (MPB-1) (Non-neural enolase) (NNE) (Phosphopyruvate hydratase) (Plasminogen-binding protein) | ENO1 ENO1L1 MBPB1 MPB1 |
| sp\|P40939\|ECHA_HUMAN | 235,561 | 295,724 | 429,378 | 244,973 | sp\|P40939\|ECHA_HUMAN | 235,736 | 93,185 | 40% | P40939 | ECHA_HUMAN | ECHA_HUMAN | Trifunctional enzyme subunit alpha, mitochondrial (78 kDa gastrin-binding protein) (Monolysocardiolipin acyltransferase) (EC 2.3.1.-) (TP-alpha) [Includes: Long-chain enoyl-CoA hydratase (EC 4.2.1.17); Long chain 3-hydroxyacyl-CoA dehydrogenase (EC 1.1.1.211)] | HADHA HADH |
| sp\|Q14764\|MVP_HUMAN | 3,623,068 | 2,503,480 | 4,489,290 | 3,129,073 | sp\|Q14764\|MVP_HUMAN | 2,544,765 | 1,124,447 | 44% | Q14764 | MVP_HUMAN | MVP_HUMAN | Major vault protein (MVP) (Lung resistance-related protein) | MVP LRP |
| sp\|P68371\|TBB4B_HUMAN | 3,018,758 | 2,778,157 | 4,118,949 | 3,864,613 | sp\|P68371\|TBB4B_HUMAN | 2,451,626 | 1,240,200 | 51% | P68371 | TBB4B_HUMAN | TBB4B_HUMAN | Tubulin beta-4B chain (Tubulin beta-2 chain) (Tubulin beta-2C chain) | TUBB4B TUBB2C |
| sp\|P19367\|HXK1_HUMAN | 68,553 | 108,952 | 57,132 | 126,238 | sp\|P19367\|HXK1_HUMAN | 79,569 | 27,574 | 35% | P19367 | HXK1_HUMAN | HXK1_HUMAN | Hexokinase-1 (EC 2.7.1.1) (Brain form hexokinase) (Hexokinase type I) (HK I) (Hexokinase-A) | HK1 |
| sp\|P35908\|K22E_HUMAN | 8,819,277 | 4,060,212 | 6,906,713 | 5,424,490 | sp\|P35908\|K22E_HUMAN | 6,978,860 | 2,493,174 | 36% | P35908 | K22E_HUMAN | K22E_HUMAN | Keratin, type II cytoskeletal 2 epidermal (Cytokeratin-2e) (CK-2e) (Epithelial keratin-2e) (Keratin-2 epidermis) (Keratin-2e) (K2e) (Type-II keratin Kb2) | KRT2 KRT2A KRT2E |
| sp\|P25705\|ATPA_HUMAN | 1,238,658 | 1,284,990 | 1,572,922 | 1,593,184 | sp\|P25705\|ATPA_HUMAN | 1,025,169 | 454,251 | 44% | P25705 | ATPA_HUMAN | ATPA_HUMAN | ATP synthase subunit alpha, mitochondrial (ATP synthase F1 subunit alpha) | ATP5F1A ATP5A ATP5A1 ATP5AL2 ATPM |
| sp\|P50990\|TCPQ_HUMAN | 1,569,303 | 1,451,047 | 1,720,825 | 1,720,284 | sp\|P50990\|TCPQ_HUMAN | 1,335,060 | 326,574 | 24% | P50990 | TCPQ_HUMAN | TCPQ_HUMAN | T-complex protein 1 subunit theta (TCP-1-theta) (CCT-theta) (Chaperonin containing T-complex polypeptide 1 subunit 8) (Renal carcinoma antigen NY-REN-15) | CCT8 C21orf112 CCTQ KIAA0002 |
| sp\|P13645\|K1C10_HUMAN | 12,850,222 | 6,494,894 | 11,284,578 | 8,982,457 | sp\|P13645\|K1C10_HUMAN | 12,009,977 | 5,211,506 | 43% | P13645 | K1C10_HUMAN | K1C10_HUMAN | Keratin, type I cytoskeletal 10 (Cytokeratin-10) (CK-10) (Keratin-10) (K10) | KRT10 KPP |
| sp\|P07355\|ANXA2_HUMAN | 15,025,974 | 14,944,716 | 19,853,347 | 17,885,489 | sp\|P07355\|ANXA2_HUMAN | 14,129,992 | 3,581,202 | 25% | P07355 | ANXA2_HUMAN | ANXA2_HUMAN | Annexin A2 (Annexin II) (Annexin-2) (Calpactin I heavy chain) (Calpactin-1 heavy chain) (Chromobindin-8) (Lipocortin II) (Placental anticoagulant protein IV) (PAP-IV) (Protein I) (p36) | ANXA2 ANX2 ANX2L4 CAL1H LPC2D |
| sp\|P08582\|TRFM_HUMAN | 4,137,812 | 4,228,278 | 3,051,837 | 5,572,531 | sp\|P08582\|TRFM_HUMAN | 3,758,506 | 886,299 | 24% | P08582 | TRFM_HUMAN | TRFM_HUMAN | Melanotransferrin (Melanoma-associated antigen p97) (CD antigen CD228) | MELTF MAP97 MFI2 |
| sp\|P16112\|PGCA_HUMAN | 8,247,780 | 7,471,044 | 12,534,396 | 11,274,018 | sp\|P16112\|PGCA_HUMAN | 10,884,986 | 2,205,221 | 20% | P16112 | PGCA_HUMAN | PGCA_HUMAN | Aggrecan core protein (Cartilage-specific proteoglycan core protein) (CSPCP) (Chondroitin sulfate proteoglycan core protein 1) (Chondroitin sulfate proteoglycan 1) [Cleaved into: Aggrecan core protein 2] | ACAN AGC1 CSPG1 MSK16 |

| | | | | | | | | | | | | | |
|---|---|---|---|---|---|---|---|---|---|---|---|---|---|
| sp\|P35580\|MYH10_HUMAN | 334,622 | 379,343 | 628,112 | 638,722 | sp\|P35580\|MYH10_HUMAN | 434,779 | 138,752 | 32% | P35580 | MYH10_HUMAN | MYH10_HUMAN | Myosin-10 (Cellular myosin heavy chain, type B) (Myosin heavy chain 10) (Myosin heavy chain, non-muscle IIb) (Non-muscle myosin heavy chain B) (NMMHC-B) (Non-muscle myosin heavy chain IIb) (NMMHC II-b) (NMMHC-IIB) | MYH10 |
| sp\|Q08431\|MFGM_HUMAN | 12,710,409 | 19,700,464 | 16,663,246 | 16,356,582 | sp\|Q08431\|MFGM_HUMAN | 17,454,060 | 3,064,781 | 18% | Q08431 | MFGM_HUMAN | MFGM_HUMAN | Lactadherin (Breast epithelial antigen BA46) (HMFG) (MFGM) (Milk fat globule-EGF factor 8) (MFG-E8) (SED1) [Cleaved into: Lactadherin short form; Medin] | MFGE8 |
| sp\|P12814\|ACTN1_HUMAN | 524,041 | 747,586 | 325,637 | 534,132 | sp\|P12814\|ACTN1_HUMAN | 495,525 | 140,501 | 28% | P12814 | ACTN1_HUMAN | ACTN1_HUMAN | Alpha-actinin-1 (Alpha-actinin cytoskeletal isoform) (F-actin cross-linking protein) (Non-muscle alpha-actinin-1) | ACTN1 |
| sp\|P08195\|4F2_HUMAN | 7,384,342 | 10,064,077 | 7,605,046 | 14,281,141 | sp\|P08195\|4F2_HUMAN | 10,795,350 | 4,285,597 | 40% | P08195 | 4F2_HUMAN | 4F2_HUMAN | 4F2 cell-surface antigen heavy chain (4F2hc) (4F2 heavy chain antigen) (Lymphocyte activation antigen 4F2 large subunit) (Solute carrier family 3 member 2) (CD antigen CD98) | SLC3A2 MDU1 |
| sp\|P02786\|TFR1_HUMAN | 1,909,109 | 1,825,647 | 2,481,562 | 2,236,292 | sp\|P02786\|TFR1_HUMAN | 1,802,066 | 453,459 | 25% | P02786 | TFR1_HUMAN | TFR1_HUMAN | Transferrin receptor protein 1 (TR) (TfR) (TfR1) (Trfr) (T9) (p90) (CD antigen CD71) [Cleaved into: Transferrin receptor protein 1, serum form (sTfR)] | TFRC |
| sp\|P06576\|ATPB_HUMAN | 1,047,330 | 980,823 | 1,223,862 | 1,239,309 | sp\|P06576\|ATPB_HUMAN | 828,216 | 336,735 | 41% | P06576 | ATPB_HUMAN | ATPB_HUMAN | ATP synthase subunit beta, mitochondrial (EC 7.1.2.2) (ATP synthase F1 subunit beta) | ATP5F1B ATP5B ATPMB ATPSB |
| sp\|P38646\|GRP75_HUMAN | 2,099,316 | 2,494,436 | 1,528,453 | 1,755,880 | sp\|P38646\|GRP75_HUMAN | 1,591,583 | 516,528 | 32% | P38646 | GRP75_HUMAN | GRP75_HUMAN | Stress-70 protein, mitochondrial (75 kDa glucose-regulated protein) (GRP-75) (Heat shock 70 kDa protein 9) (Mortalin) (MOT) (Peptide-binding protein 74) (PBP74) | HSPA9 GRP75 HSPA9B mt-HSP70 |
| sp\|Q9P265\|DIP2B_HUMAN | 987,061 | 1,093,876 | 892,699 | 1,104,504 | sp\|Q9P265\|DIP2B_HUMAN | 888,951 | 173,751 | 20% | Q9P265 | DIP2B_HUMAN | DIP2B_HUMAN | Disco-interacting protein 2 homolog B (DIP2 homolog B) | DIP2B KIAA1463 |
| sp\|P08758\|ANXA5_HUMAN | 15,594,804 | 13,898,367 | 20,084,025 | 16,038,676 | sp\|P08758\|ANXA5_HUMAN | 13,702,606 | 3,434,206 | 25% | P08758 | ANXA5_HUMAN | ANXA5_HUMAN | Annexin A5 (Anchorin CII) (Annexin V) (Annexin-5) (Calphobindin I) (CPB-I) (Endonexin II) (Lipocortin V) (Placental anticoagulant protein 4) (PP4) (Placental anticoagulant protein I) (PAP-I) (Thromboplastin inhibitor) (Vascular anticoagulant-alpha) (VAC-alpha) | ANXA5 ANX5 ENX2 PP4 |
| sp\|Q9BQE3\|TBA1C_HUMAN | 145,720 | 206,005 | 18,055 | 15,396 | sp\|Q9BQE3\|TBA1C_HUMAN | 69,237 | 71,816 | 104% | Q9BQE3 | TBA1C_HUMAN | TBA1C_HUMAN | Tubulin alpha-1C chain (Alpha-tubulin 6) (Tubulin alpha-6 chain) [Cleaved into: Detyrosinated tubulin alpha-1C chain] | TUBA1C TUBA6 |
| sp\|P23229\|ITA6_HUMAN | 2,541,270 | 2,525,534 | 3,445,823 | 3,710,889 | sp\|P23229\|ITA6_HUMAN | 3,236,662 | 700,530 | 22% | P23229 | ITA6_HUMAN | ITA6_HUMAN | Integrin alpha-6 (CD49 antigen-like family member F) (VLA-6) (CD antigen CD49f) [Cleaved into: Integrin alpha-6 heavy chain; Integrin alpha-6 light chain; Processed integrin alpha-6 (Alpha6p)] | ITGA6 |
| sp\|P49368\|TCPG_HUMAN | 1,264,226 | 1,206,969 | 1,383,098 | 1,402,091 | sp\|P49368\|TCPG_HUMAN | 1,044,137 | 318,303 | 30% | P49368 | TCPG_HUMAN | TCPG_HUMAN | T-complex protein 1 subunit gamma (TCP-1-gamma) (CCT-gamma) (hTRiC5) | CCT3 CCTG TRIC5 |

| | | | | | | | | | | | | | |
|---|---|---|---|---|---|---|---|---|---|---|---|---|---|
| sp\|P21589\|5NTD_HUMAN | 6,974,094 | 7,496,042 | 4,209,883 | 4,902,599 | sp\|P21589\|5NTD_HUMAN | 4,267,661 | 2,070,577 | 49% | P21589 | 5NTD_HUMAN | 5NTD_HUMAN | 5'-nucleotidase (5'-NT) (EC 3.1.3.5) (Ecto-5'-nucleotidase) (CD antigen CD73) | NT5E NT5 NTE |
| sp\|P07900\|HS90A_HUMAN | 956,832 | 1,088,889 | 685,152 | 814,623 | sp\|P07900\|HS90A_HUMAN | 819,987 | 150,796 | 18% | P07900 | HS90A_HUMAN | HS90A_HUMAN | Heat shock protein HSP 90-alpha (EC 3.6.4.10) (Heat shock 86 kDa) (HSP 86) (HSP86) (Lipopolysaccharide-associated protein 2) (LAP-2) (LPS-associated protein 2) (Renal carcinoma antigen NY-REN-38) | HSP90AA1 HSP90A HSPC1 HSPCA |
| sp\|P48681\|NEST_HUMAN | 182,827 | 231,151 | 550,019 | 477,197 | sp\|P48681\|NEST_HUMAN | 342,269 | 202,235 | 59% | P48681 | NEST_HUMAN | NEST_HUMAN | Nestin | NES Nbla00170 |
| sp\|P53396\|ACLY_HUMAN | 382,371 | 461,475 | 337,229 | 357,101 | sp\|P53396\|ACLY_HUMAN | 348,745 | 55,867 | 16% | P53396 | ACLY_HUMAN | ACLY_HUMAN | ATP-citrate synthase (EC 2.3.3.8) (ATP-citrate (pro-S-)-lyase) (ACL) (Citrate cleavage enzyme) | ACLY |
| sp\|P00558\|PGK1_HUMAN | 1,836,628 | 2,278,955 | 1,219,626 | 1,470,489 | sp\|P00558\|PGK1_HUMAN | 1,482,105 | 473,036 | 32% | P00558 | PGK1_HUMAN | PGK1_HUMAN | Phosphoglycerate kinase 1 (EC 2.7.2.3) (Cell migration-inducing gene 10 protein) (Primer recognition protein 2) (PRP 2) | PGK1 PGKA MIG10 OK/SW-cl.110 |
| sp\|Q13813\|SPTN1_HUMAN | 171,955 | 225,388 | 283,297 | 464,603 | sp\|Q13813\|SPTN1_HUMAN | 242,645 | 108,715 | 45% | Q13813 | SPTN1_HUMAN | SPTN1_HUMAN | Spectrin alpha chain, non-erythrocytic 1 (Alpha-II spectrin) (Fodrin alpha chain) (Spectrin, non-erythroid alpha subunit) | SPTAN1 NEAS SPTA2 |
| sp\|P43121\|MUC18_HUMAN | 8,159,485 | 10,078,190 | 8,336,785 | 9,708,649 | sp\|P43121\|MUC18_HUMAN | 7,478,118 | 1,842,123 | 25% | P43121 | MUC18_HUMAN | MUC18_HUMAN | Cell surface glycoprotein MUC18 (Cell surface glycoprotein P1H12) (Melanoma cell adhesion molecule) (Melanoma-associated antigen A32) (Melanoma-associated antigen MUC18) (S-endo 1 endothelial-associated antigen) (CD antigen CD146) | MCAM MUC18 |
| sp\|P47895\|AL1A3_HUMAN | 1,702,077 | 1,836,285 | 1,723,247 | 1,868,001 | sp\|P47895\|AL1A3_HUMAN | 1,516,930 | 304,426 | 20% | P47895 | AL1A3_HUMAN | AL1A3_HUMAN | Aldehyde dehydrogenase family 1 member A3 (EC 1.2.1.36) (Aldehyde dehydrogenase 6) (Retinaldehyde dehydrogenase 3) (RALDH-3) (RalDH3) | ALDH1A3 ALDH6 |
| sp\|P27824\|CALX_HUMAN | 396,399 | 390,863 | 257,616 | 263,606 | sp\|P27824\|CALX_HUMAN | 249,275 | 99,380 | 40% | P27824 | CALX_HUMAN | CALX_HUMAN | Calnexin (IP90) (Major histocompatibility complex class I antigen-binding protein p88) (p90) | CANX |
| sp\|P78371\|TCPB_HUMAN | 748,433 | 746,849 | 878,190 | 886,539 | sp\|P78371\|TCPB_HUMAN | 716,625 | 132,221 | 18% | P78371 | TCPB_HUMAN | TCPB_HUMAN | T-complex protein 1 subunit beta (TCP-1-beta) (CCT-beta) | CCT2 99D8.1 CCTB |
| sp\|P02545\|LMNA_HUMAN | 233,294 | 214,145 | 520,733 | 642,867 | sp\|P02545\|LMNA_HUMAN | 295,442 | 184,723 | 63% | P02545 | LMNA_HUMAN | LMNA_HUMAN | Prelamin-A/C [Cleaved into: Lamin-A/C (70 kDa lamin) (Renal carcinoma antigen NY-REN-32)] | LMNA LMN1 |
| sp\|O95573\|ACSL3_HUMAN | 444,620 | 585,524 | 601,852 | 924,443 | sp\|O95573\|ACSL3_HUMAN | 470,298 | 225,913 | 48% | O95573 | ACSL3_HUMAN | ACSL3_HUMAN | Fatty acid CoA ligase Acsl3 (Arachidonate--CoA ligase) (EC 6.2.1.15) (Long-chain acyl-CoA synthetase 3) (LACS 3) (Long-chain-fatty-acid--CoA ligase 3) (EC 6.2.1.3) (Medium-chain acyl-CoA ligase Acsl3) (EC 6.2.1.2) | ACSL3 ACS3 FACL3 LACS3 |
| sp\|P63261\|ACTG_HUMAN | 15,214,299 | 18,448,286 | 21,160,669 | 25,442,087 | sp\|P63261\|ACTG_HUMAN | 18,450,575 | 3,631,227 | 20% | P63261 | ACTG_HUMAN | ACTG_HUMAN | Actin, cytoplasmic 2 (Gamma-actin) [Cleaved into: Actin, cytoplasmic 2, N-terminally processed] | ACTG1 ACTG |
| sp\|Q07065\|CKAP4_HUMAN | 196,549 | 122,385 | 241,076 | 186,358 | sp\|Q07065\|CKAP4_HUMAN | 166,022 | 59,927 | 36% | Q07065 | CKAP4_HUMAN | CKAP4_HUMAN | Cytoskeleton-associated protein 4 (63-kDa cytoskeleton-linking membrane protein) (Climp-63) (p63) | CKAP4 |

| | | | | | | | | | | | | | |
|---|---|---|---|---|---|---|---|---|---|---|---|---|---|
| sp\|P22314\|UBA1_HUMAN | 933,832 | 960,343 | 1,526,268 | 1,212,964 | sp\|P22314\|UBA1_HUMAN | 1,042,286 | 258,491 | 25% | P22314 | UBA1_HUMAN | UBA1_HUMAN | Ubiquitin-like modifier-activating enzyme 1 (EC 6.2.1.45) (Protein A1S9) (Ubiquitin-activating enzyme E1) | UBA1 A1S9T UBE1 |
| sp\|Q9UQ80\|PA2G4_HUMAN | 579,509 | 834,172 | 854,728 | 940,262 | sp\|Q9UQ80\|PA2G4_HUMAN | 724,643 | 138,121 | 19% | Q9UQ80 | PA2G4_HUMAN | PA2G4_HUMAN | Proliferation-associated protein 2G4 (Cell cycle protein p38-2G4 homolog) (hG4-1) (ErbB3-binding protein 1) | PA2G4 EBP1 |
| sp\|P04075\|ALDOA_HUMAN | 2,047,212 | 3,051,162 | 1,419,141 | 2,134,905 | sp\|P04075\|ALDOA_HUMAN | 1,996,787 | 546,572 | 27% | P04075 | ALDOA_HUMAN | ALDOA_HUMAN | Fructose-bisphosphate aldolase A (EC 4.1.2.13) (Lung cancer antigen NY-LU-1) (Muscle-type aldolase) | ALDOA ALDA |
| sp\|O75083\|WDR1_HUMAN | 1,737,494 | 2,062,531 | 1,914,773 | 2,277,416 | sp\|O75083\|WDR1_HUMAN | 1,884,220 | 196,491 | 10% | O75083 | WDR1_HUMAN | WDR1_HUMAN | WD repeat-containing protein 1 (Actin-interacting protein 1) (AIP1) (NORI-1) | WDR1 |
| sp\|P31939\|PUR9_HUMAN | 714,956 | 741,342 | 602,182 | 551,779 | sp\|P31939\|PUR9_HUMAN | 611,078 | 92,435 | 15% | P31939 | PUR9_HUMAN | PUR9_HUMAN | Bifunctional purine biosynthesis protein ATIC (AICAR transformylase/inosine monophosphate cyclohydrolase) (ATIC) [Cleaved into: Bifunctional purine biosynthesis protein ATIC, N-terminally processed] [Includes: Phosphoribosylaminoimidazolecarboxamide formyltransferase (EC 2.1.2.3) (5-aminoimidazole-4-carboxamide ribonucleotide formyltransferase) (AICAR formyltransferase) (AICAR transformylase); Inosine 5'-monophosphate cyclohydrolase (IMP cyclohydrolase) (EC 3.5.4.10) (IMP synthase) (Inosinicase)] | ATIC PURH OK/SW-cl.86 |
| sp\|P02649\|APOE_HUMAN | 1,285,886 | 1,300,817 | 2,339,056 | 1,862,002 | sp\|P02649\|APOE_HUMAN | 6,928,822 | 5,848,850 | 84% | P02649 | APOE_HUMAN | APOE_HUMAN | Apolipoprotein E (Apo-E) | APOE |
| sp\|P05106\|ITB3_HUMAN | 1,576,343 | 1,573,193 | 1,060,082 | 1,165,011 | sp\|P05106\|ITB3_HUMAN | 1,160,585 | 321,690 | 28% | P05106 | ITB3_HUMAN | ITB3_HUMAN | Integrin beta-3 (Platelet membrane glycoprotein IIIa) (GPIIIa) (CD antigen CD61) | ITGB3 GP3A |
| sp\|P10915\|HPLN1_HUMAN | 22,643,568 | 18,310,706 | 28,662,897 | 24,085,908 | sp\|P10915\|HPLN1_HUMAN | 20,902,160 | 4,923,035 | 24% | P10915 | HPLN1_HUMAN | HPLN1_HUMAN | Hyaluronan and proteoglycan link protein 1 (Cartilage-linking protein 1) (Cartilage-link protein) (Proteoglycan link protein) | HAPLN1 CRTL1 |
| sp\|P02751\|FINC_HUMAN | 537,658 | 646,333 | 533,883 | 718,008 | sp\|P02751\|FINC_HUMAN | 1,510,569 | 1,151,694 | 76% | P02751 | FINC_HUMAN | FINC_HUMAN | Fibronectin (FN) (Cold-insoluble globulin) (CIG) [Cleaved into: Anastellin; Ugl-Y1; Ugl-Y2; Ugl-Y3] | FN1 FN |
| sp\|P22102\|PUR2_HUMAN | 468,818 | 439,351 | 426,894 | 882,012 | sp\|P22102\|PUR2_HUMAN | 554,395 | 176,291 | 32% | P22102 | PUR2_HUMAN | PUR2_HUMAN | Trifunctional purine biosynthetic protein adenosine-3 [Includes: Phosphoribosylamine--glycine ligase (EC 6.3.4.13) (Glycinamide ribonucleotide synthetase) (GARS) (Phosphoribosylglycinamide synthetase); Phosphoribosylformylglycinamidine cyclo-ligase (EC 6.3.3.1) (AIR synthase) (AIRS) (Phosphoribosyl-aminoimidazole synthetase); Phosphoribosylglycinamide formyltransferase (EC 2.1.2.2) (5'-phosphoribosylglycinamide transformylase) (GAR transformylase) (GART)] | GART PGFT PRGS |

| | | | | | | | | | | | | | |
|---|---|---|---|---|---|---|---|---|---|---|---|---|---|
| sp\|P06396\|GELS_HUMAN | 965,095 | 1,199,683 | 687,417 | 1,151,869 | sp\|P06396\|GELS_HUMAN | 1,176,763 | 392,756 | 33% | P06396 | GELS_HUMAN | GELS_HUMAN | Gelsolin (AGEL) (Actin-depolymerizing factor) (ADF) (Brevin) | GSN |
| sp\|Q9P2B2\|FPRP_HUMAN | 2,059,612 | 2,039,486 | 2,490,741 | 2,285,976 | sp\|Q9P2B2\|FPRP_HUMAN | 1,899,893 | 487,869 | 26% | Q9P2B2 | FPRP_HUMAN | FPRP_HUMAN | Prostaglandin F2 receptor negative regulator (CD9 partner 1) (CD9P-1) (Glu-Trp-Ile EWI motif-containing protein F) (EWI-F) (Prostaglandin F2-alpha receptor regulatory protein) (Prostaglandin F2-alpha receptor-associated protein) (CD antigen CD315) | PTGFRN CD9P1 EWIF FPRP KIAA1436 |
| sp\|P26641\|EF1G_HUMAN | 1,614,959 | 2,566,597 | 1,699,211 | 2,377,493 | sp\|P26641\|EF1G_HUMAN | 1,502,327 | 750,326 | 50% | P26641 | EF1G_HUMAN | EF1G_HUMAN | Elongation factor 1-gamma (EF-1-gamma) (eEF-1B gamma) | EEF1G EF1G PRO1608 |
| sp\|P48643\|TCPE_HUMAN | 393,569 | 394,080 | 502,801 | 537,572 | sp\|P48643\|TCPE_HUMAN | 392,026 | 118,800 | 30% | P48643 | TCPE_HUMAN | TCPE_HUMAN | T-complex protein 1 subunit epsilon (TCP-1-epsilon) (CCT-epsilon) | CCT5 CCTE KIAA0098 |
| sp\|P07996\|TSP1_HUMAN | 201,224 | 270,221 | 212,399 | 403,450 | sp\|P07996\|TSP1_HUMAN | 1,956,815 | 1,901,197 | 97% | P07996 | TSP1_HUMAN | TSP1_HUMAN | Thrombospondin-1 (Glycoprotein G) | THBS1 TSP TSP1 |
| sp\|Q01518\|CAP1_HUMAN | 1,069,805 | 1,811,144 | 466,883 | 665,254 | sp\|Q01518\|CAP1_HUMAN | 961,416 | 450,144 | 47% | Q01518 | CAP1_HUMAN | CAP1_HUMAN | Adenylyl cyclase-associated protein 1 (CAP 1) | CAP1 CAP |
| sp\|P30153\|2AAA_HUMAN | 619,446 | 653,016 | 757,859 | 814,555 | sp\|P30153\|2AAA_HUMAN | 702,822 | 80,555 | 11% | P30153 | 2AAA_HUMAN | 2AAA_HUMAN | Serine/threonine-protein phosphatase 2A 65 kDa regulatory subunit A alpha isoform (Medium tumor antigen-associated 61 kDa protein) (PP2A subunit A isoform PR65-alpha) (PP2A subunit A isoform R1-alpha) | PPP2R1A |
| sp\|P62258\|1433E_HUMAN | 2,033,536 | 2,934,582 | 1,924,033 | 2,473,160 | sp\|P62258\|1433E_HUMAN | 2,356,056 | 415,108 | 18% | P62258 | 1433E_HUMAN | 1433E_HUMAN | 14-3-3 protein epsilon (14-3-3E) | YWHAE |
| sp\|Q99832\|TCPH_HUMAN | 627,212 | 648,576 | 745,772 | 746,587 | sp\|Q99832\|TCPH_HUMAN | 554,533 | 160,717 | 29% | Q99832 | TCPH_HUMAN | TCPH_HUMAN | T-complex protein 1 subunit eta (TCP-1-eta) (CCT-eta) (HIV-1 Nef-interacting protein) [Cleaved into: T-complex protein 1 subunit eta, N-terminally processed] | CCT7 CCTH NIP7-1 |
| sp\|P17987\|TCPA_HUMAN | 1,252,069 | 1,073,701 | 1,416,367 | 1,312,310 | sp\|P17987\|TCPA_HUMAN | 1,007,563 | 326,782 | 32% | P17987 | TCPA_HUMAN | TCPA_HUMAN | T-complex protein 1 subunit alpha (TCP-1-alpha) (CCT-alpha) | TCP1 CCT1 CCTA |
| sp\|P09543\|CN37_HUMAN | 1,816,821 | 1,872,515 | 1,470,975 | 1,327,003 | sp\|P09543\|CN37_HUMAN | 1,536,460 | 306,173 | 20% | P09543 | CN37_HUMAN | CN37_HUMAN | 2',3'-cyclic-nucleotide 3'-phosphodiesterase (CNP) (CNPase) (EC 3.1.4.37) | CNP |
| sp\|Q9HBA0\|TRPV4_HUMAN | 1,429,858 | 1,792,485 | 2,508,650 | 2,325,586 | sp\|Q9HBA0\|TRPV4_HUMAN | 2,084,880 | 352,689 | 17% | Q9HBA0 | TRPV4_HUMAN | TRPV4_HUMAN | Transient receptor potential cation channel subfamily V member 4 (TrpV4) (Osm-9-like TRP channel 4) (OTRPC4) (Transient receptor potential protein 12) (TRP12) (Vanilloid receptor-like channel 2) (Vanilloid receptor-like protein 2) (VRL-2) (Vanilloid receptor-related osmotically-activated channel) (VR-OAC) | TRPV4 VRL2 VROAC |
| sp\|P00338\|LDHA_HUMAN | 5,301,013 | 6,324,727 | 2,920,294 | 3,575,541 | sp\|P00338\|LDHA_HUMAN | 4,301,469 | 1,364,264 | 32% | P00338 | LDHA_HUMAN | LDHA_HUMAN | L-lactate dehydrogenase A chain (LDH-A) (EC 1.1.1.27) (Cell proliferation-inducing gene 19 protein) (LDH muscle subunit) (LDH-M) (Renal carcinoma antigen NY-REN-59) | LDHA PIG19 |
| sp\|P50395\|GDIB_HUMAN | 835,462 | 1,009,641 | 446,921 | 428,950 | sp\|P50395\|GDIB_HUMAN | 627,083 | 235,031 | 37% | P50395 | GDIB_HUMAN | GDIB_HUMAN | Rab GDP dissociation inhibitor beta (Rab GDI beta) (Guanosine diphosphate dissociation inhibitor 2) (GDI-2) | GDI2 RABGDIB |
| sp\|P49588\|SYAC_HUMAN | 459,117 | 500,865 | 542,758 | 543,452 | sp\|P49588\|SYAC_HUMAN | 469,665 | 67,656 | 14% | P49588 | SYAC_HUMAN | SYAC_HUMAN | Alanine--tRNA ligase, cytoplasmic (EC 6.1.1.7) (Alanyl-tRNA synthetase) (AlaRS) (Renal carcinoma antigen NY-REN-42) | AARS1 AARS |

| | | | | | | | | | | | | | |
|---|---|---|---|---|---|---|---|---|---|---|---|---|---|
| sp\|P04406\|G3P_HUMAN | 12,137,948 | 12,902,235 | 8,643,517 | 9,087,032 | sp\|P04406\|G3P_HUMAN | 9,183,457 | 2,583,846 | 28% | P04406 | G3P_HUMAN | G3P_HUMAN | Glyceraldehyde-3-phosphate dehydrogenase (GAPDH) (EC 1.2.1.12) (Peptidyl-cysteine S-nitrosylase GAPDH) (EC 2.6.99.-) | GAPDH GAPD CDABP0047 OK/SW-cl.12 |
| sp\|Q08380\|LG3BP_HUMAN | 10,479,448 | 8,404,663 | 10,935,255 | 8,690,342 | sp\|Q08380\|LG3BP_HUMAN | 7,731,321 | 2,309,095 | 30% | Q08380 | LG3BP_HUMAN | LG3BP_HUMAN | Galectin-3-binding protein (Basement membrane autoantigen p105) (Lectin galactoside-binding soluble 3-binding protein) (Mac-2-binding protein) (MAC2BP) (Mac-2 BP) (Tumor-associated antigen 90K) | LGALS3BP M2BP |
| sp\|Q14152\|EIF3A_HUMAN | 412,041 | 413,495 | 480,065 | 550,761 | sp\|Q14152\|EIF3A_HUMAN | 504,776 | 132,127 | 26% | Q14152 | EIF3A_HUMAN | EIF3A_HUMAN | Eukaryotic translation initiation factor 3 subunit A (eIF3a) (Eukaryotic translation initiation factor 3 subunit 10) (eIF-3-theta) (eIF3 p167) (eIF3 p180) (eIF3 p185) | EIF3A EIF3S10 KIAA0139 |
| sp\|P22234\|PUR6_HUMAN | 820,475 | 925,877 | 420,984 | 666,866 | sp\|P22234\|PUR6_HUMAN | 618,907 | 200,528 | 32% | P22234 | PUR6_HUMAN | PUR6_HUMAN | Bifunctional phosphoribosylaminoimidazole carboxylase/phosphoribosylaminoimidazole succinocarboxamide synthetase (PAICS) [Includes: Phosphoribosylaminoimidazole carboxylase (EC 4.1.1.21) (AIR carboxylase) (AIRC); Phosphoribosylaminoimidazole succinocarboxamide synthetase (EC 6.3.2.6) (SAICAR synthetase)] | PAICS ADE2 AIRC PAIS |
| sp\|Q7L576\|CYFP1_HUMAN | 584,548 | 556,813 | 736,589 | 536,844 | sp\|Q7L576\|CYFP1_HUMAN | 582,539 | 69,550 | 12% | Q7L576 | CYFP1_HUMAN | CYFP1_HUMAN | Cytoplasmic FMR1-interacting protein 1 (Specifically Rac1-associated protein 1) (Sra-1) (p140sra-1) | CYFIP1 KIAA0068 |
| sp\|P60842\|IF4A1_HUMAN | 441,306 | 682,279 | 348,738 | 365,815 | sp\|P60842\|IF4A1_HUMAN | 422,508 | 141,560 | 34% | P60842 | IF4A1_HUMAN | IF4A1_HUMAN | Eukaryotic initiation factor 4A-I (eIF-4A-I) (eIF4A-I) (EC 3.6.4.13) (ATP-dependent RNA helicase eIF4A-1) | EIF4A1 DDX2A EIF4A |
| sp\|P05556\|ITB1_HUMAN | 4,013,283 | 4,870,658 | 2,446,140 | 2,664,963 | sp\|P05556\|ITB1_HUMAN | 2,999,410 | 1,146,864 | 38% | P05556 | ITB1_HUMAN | ITB1_HUMAN | Integrin beta-1 (Fibronectin receptor subunit beta) (Glycoprotein IIa) (GPIIA) (VLA-4 subunit beta) (CD antigen CD29) | ITGB1 FNRB MDF2 MSK12 |
| sp\|P11586\|C1TC_HUMAN | 524,441 | 489,057 | 283,200 | 426,071 | sp\|P11586\|C1TC_HUMAN | 393,086 | 92,124 | 23% | P11586 | C1TC_HUMAN | C1TC_HUMAN | C-1-tetrahydrofolate synthase, cytoplasmic (C1-THF synthase) (Epididymis secretory sperm binding protein) [Cleaved into: C-1-tetrahydrofolate synthase, cytoplasmic, N-terminally processed] [Includes: Methylenetetrahydrofolate dehydrogenase (EC 1.5.1.5); Methenyltetrahydrofolate cyclohydrolase (EC 3.5.4.9); Formyltetrahydrofolate synthetase (EC 6.3.4.3)] | MTHFD1 MTHFC MTHFD |
| sp\|P04843\|RPN1_HUMAN | 288,537 | 249,200 | 232,481 | 227,146 | sp\|P04843\|RPN1_HUMAN | 244,567 | 63,010 | 26% | P04843 | RPN1_HUMAN | RPN1_HUMAN | Dolichyl-diphosphooligosaccharide--protein glycosyltransferase subunit 1 (Dolichyl-diphosphooligosaccharide--protein glycosyltransferase 67 kDa subunit) (Ribophorin I) (RPN-I) (Ribophorin-1) | RPN1 |
| sp\|P49411\|EFTU_HUMAN | 503,465 | 476,152 | 452,528 | 553,782 | sp\|P49411\|EFTU_HUMAN | 376,957 | 132,177 | 35% | P49411 | EFTU_HUMAN | EFTU_HUMAN | Elongation factor Tu, mitochondrial (EF-Tu) (P43) | TUFM |
| sp\|Q16363\|LAMA4_HUMAN | 1,726,308 | 1,634,716 | 1,669,715 | 1,703,250 | sp\|Q16363\|LAMA4_HUMAN | 1,308,349 | 424,187 | 32% | Q16363 | LAMA4_HUMAN | LAMA4_HUMAN | Laminin subunit alpha-4 (Laminin-14 subunit alpha) (Laminin-8 subunit alpha) (Laminin-9 subunit alpha) | LAMA4 |

| | | | | | | | | | | | | | |
|---|---|---|---|---|---|---|---|---|---|---|---|---|---|
| sp\|Q969P0\|IGSF8_HUMAN | 11,916,824 | 10,157,355 | 18,598,686 | 17,582,870 | sp\|Q969P0\|IGSF8_HUMAN | 13,167,277 | 3,954,975 | 30% | Q969P0 | IGSF8_HUMAN | IGSF8_HUMAN | Immunoglobulin superfamily member 8 (IgSF8) (CD81 partner 3) (Glu-Trp-Ile EWI motif-containing protein 2) (EWI-2) (Keratinocytes-associated transmembrane protein 4) (KCT-4) (LIR-D1) (Prostaglandin regulatory-like protein) (PGRL) (CD antigen CD316) | IGSF8 CD81P3 EWI2 KCT4 |
| sp\|Q9Y4L1\|HYOU1_HUMAN | 165,200 | 79,390 | 170,182 | 96,437 | sp\|Q9Y4L1\|HYOU1_HUMAN | 145,036 | 41,423 | 29% | Q9Y4L1 | HYOU1_HUMAN | HYOU1_HUMAN | Hypoxia up-regulated protein 1 (150 kDa oxygen-regulated protein) (ORP-150) (170 kDa glucose-regulated protein) (GRP-170) | HYOU1 GRP170 ORP150 |
| sp\|P14543\|NID1_HUMAN | 1,534,851 | 1,288,867 | 2,461,999 | 2,208,690 | sp\|P14543\|NID1_HUMAN | 1,776,174 | 408,797 | 23% | P14543 | NID1_HUMAN | NID1_HUMAN | Nidogen-1 (NID-1) (Entactin) | NID1 NID |
| sp\|P12268\|IMDH2_HUMAN | 993,907 | 1,051,039 | 1,109,522 | 1,172,229 | sp\|P12268\|IMDH2_HUMAN | 859,676 | 246,094 | 29% | P12268 | IMDH2_HUMAN | IMDH2_HUMAN | Inosine-5'-monophosphate dehydrogenase 2 (IMP dehydrogenase 2) (IMPD 2) (IMPDH 2) (EC 1.1.1.205) (Inosine-5'-monophosphate dehydrogenase type II) (IMP dehydrogenase II) (IMPDH-II) | IMPDH2 IMPD2 |
| sp\|P68104\|EF1A1_HUMAN | 5,038,785 | 5,964,952 | 2,077,709 | 3,035,879 | sp\|P68104\|EF1A1_HUMAN | 3,391,604 | 1,621,911 | 48% | P68104 | EF1A1_HUMAN | EF1A1_HUMAN | Elongation factor 1-alpha 1 (EF-1-alpha-1) (Elongation factor Tu) (EF-Tu) (Eukaryotic elongation factor 1 A-1) (eEF1A-1) (Leukocyte receptor cluster member 7) | EEF1A1 EEF1A EF1A LENG7 |
| sp\|P00367\|DHE3_HUMAN | 158,495 | 211,413 | 146,388 | 175,064 | sp\|P00367\|DHE3_HUMAN | 142,407 | 39,915 | 28% | P00367 | DHE3_HUMAN | DHE3_HUMAN | Glutamate dehydrogenase 1, mitochondrial (GDH 1) (EC 1.4.1.3) | GLUD1 GLUD |
| sp\|P50991\|TCPD_HUMAN | 1,044,427 | 1,022,811 | 1,118,392 | 1,055,425 | sp\|P50991\|TCPD_HUMAN | 925,112 | 177,299 | 19% | P50991 | TCPD_HUMAN | TCPD_HUMAN | T-complex protein 1 subunit delta (TCP-1-delta) (CCT-delta) (Stimulator of TAR RNA-binding) | CCT4 CCTD SRB |
| sp\|P52789\|HXK2_HUMAN | 204,507 | 238,622 | 243,568 | 261,618 | sp\|P52789\|HXK2_HUMAN | 280,770 | 116,548 | 42% | P52789 | HXK2_HUMAN | HXK2_HUMAN | Hexokinase-2 (EC 2.7.1.1) (Hexokinase type II) (HK II) (Hexokinase-B) (Muscle form hexokinase) | HK2 |
| sp\|P05141\|ADT2_HUMAN | 350,447 | 462,122 | 235,085 | 182,384 | sp\|P05141\|ADT2_HUMAN | 256,281 | 105,031 | 41% | P05141 | ADT2_HUMAN | ADT2_HUMAN | ADP/ATP translocase 2 (ADP,ATP carrier protein 2) (ADP,ATP carrier protein, fibroblast isoform) (Adenine nucleotide translocator 2) (ANT 2) (Solute carrier family 25 member 5) [Cleaved into: ADP/ATP translocase 2, N-terminally processed] | SLC25A5 AAC2 ANT2 |
| sp\|Q9Y4F1\|FARP1_HUMAN | 587,231 | 830,042 | 928,464 | 1,104,700 | sp\|Q9Y4F1\|FARP1_HUMAN | 837,612 | 160,225 | 19% | Q9Y4F1 | FARP1_HUMAN | FARP1_HUMAN | FERM, ARHGEF and pleckstrin domain-containing protein 1 (Chondrocyte-derived ezrin-like protein) (FERM, RhoGEF and pleckstrin domain-containing protein 1) (Pleckstrin homology domain-containing family C member 2) (PH domain-containing family C member 2) | FARP1 CDEP PLEKHC2 |
| sp\|P50995\|ANX11_HUMAN | 3,494,833 | 1,978,204 | 4,156,346 | 3,366,508 | sp\|P50995\|ANX11_HUMAN | 2,910,457 | 780,654 | 27% | P50995 | ANX11_HUMAN | ANX11_HUMAN | Annexin A11 (56 kDa autoantigen) (Annexin XI) (Annexin-11) (Calcyclin-associated annexin 50) (CAP-50) | ANXA11 ANX11 |
| sp\|P11940\|PABP1_HUMAN | 435,504 | 507,463 | 218,671 | 298,899 | sp\|P11940\|PABP1_HUMAN | 330,587 | 106,656 | 32% | P11940 | PABP1_HUMAN | PABP1_HUMAN | Polyadenylate-binding protein 1 (PABP-1) (Poly(A)-binding protein 1) | PABPC1 PAB1 PABP PABP1 PABPC2 |

| | | | | | | | | | | | | | |
|---|---|---|---|---|---|---|---|---|---|---|---|---|---|
| sp\|Q14974\|IMB1_HUMAN | 457,360 | 605,864 | 615,699 | 734,978 | sp\|Q14974\|IMB1_HUMAN | 575,561 | 91,598 | 16% | Q14974 | IMB1_HUMAN | IMB1_HUMAN | Importin subunit beta-1 (Importin-90) (Karyopherin subunit beta-1) (Nuclear factor p97) (Pore targeting complex 97 kDa subunit) (PTAC97) | KPNB1 NTF97 |
| sp\|Q12931\|TRAP1_HUMAN | 96,029 | 86,195 | 162,989 | 154,410 | sp\|Q12931\|TRAP1_HUMAN | 117,238 | 44,122 | 38% | Q12931 | TRAP1_HUMAN | TRAP1_HUMAN | Heat shock protein 75 kDa, mitochondrial (HSP 75) (TNFR-associated protein 1) (Tumor necrosis factor type 1 receptor-associated protein) (TRAP-1) | TRAP1 HSP75 |
| sp\|O15061\|SYNEM_HUMAN | 62,252 | 85,543 | 210,570 | 115,865 | sp\|O15061\|SYNEM_HUMAN | 122,010 | 61,605 | 50% | O15061 | SYNEM_HUMAN | SYNEM_HUMAN | Synemin (Desmuslin) | SYNM DMN KIAA0353 SYN |
| sp\|P43304\|GPDM_HUMAN | 97,556 | 116,127 | 141,902 | 123,908 | sp\|P43304\|GPDM_HUMAN | 112,051 | 31,365 | 28% | P43304 | GPDM_HUMAN | GPDM_HUMAN | Glycerol-3-phosphate dehydrogenase, mitochondrial (GPD-M) (GPDH-M) (EC 1.1.5.3) (mitohondrial glycerophosphate dehydrogenase gene) (mGDH) (mtGPD) | GPD2 |
| sp\|O75874\|IDHC_HUMAN | 848,324 | 736,500 | 545,132 | 643,919 | sp\|O75874\|IDHC_HUMAN | 599,672 | 160,946 | 27% | O75874 | IDHC_HUMAN | IDHC_HUMAN | Isocitrate dehydrogenase [NADP] cytoplasmic (IDH) (IDH1) (EC 1.1.1.42) (Cytosolic NADP-isocitrate dehydrogenase) (IDPc) (NADP(+)-specific ICDH) (Oxalosuccinate decarboxylase) | IDH1 PICD |
| sp\|P40227\|TCPZ_HUMAN | 424,304 | 440,815 | 269,186 | 298,697 | sp\|P40227\|TCPZ_HUMAN | 334,004 | 86,576 | 26% | P40227 | TCPZ_HUMAN | TCPZ_HUMAN | T-complex protein 1 subunit zeta (TCP-1-zeta) (Acute morphine dependence-related protein 2) (CCT-zeta-1) (HTR3) (Tcp20) | CCT6A CCT6 CCTZ |
| sp\|Q9Y2E4\|DIP2C_HUMAN | 527,524 | 607,759 | 540,794 | 661,737 | sp\|Q9Y2E4\|DIP2C_HUMAN | 604,724 | 100,137 | 17% | Q9Y2E4 | DIP2C_HUMAN | DIP2C_HUMAN | Disco-interacting protein 2 homolog C (DIP2 homolog C) | DIP2C KIAA0934 |
| sp\|P40926\|MDHM_HUMAN | 564,425 | 937,754 | 345,358 | 569,148 | sp\|P40926\|MDHM_HUMAN | 519,980 | 189,359 | 36% | P40926 | MDHM_HUMAN | MDHM_HUMAN | Malate dehydrogenase, mitochondrial (EC 1.1.1.37) | MDH2 |
| sp\|O60701\|UGDH_HUMAN | 458,597 | 405,927 | 692,361 | 574,025 | sp\|O60701\|UGDH_HUMAN | 461,380 | 124,097 | 27% | O60701 | UGDH_HUMAN | UGDH_HUMAN | UDP-glucose 6-dehydrogenase (UDP-Glc dehydrogenase) (UDP-GlcDH) (UDPGDH) (EC 1.1.1.22) | UGDH |
| sp\|P23526\|SAHH_HUMAN | 1,432,874 | 1,606,024 | 1,490,529 | 1,604,701 | sp\|P23526\|SAHH_HUMAN | 1,716,524 | 228,974 | 13% | P23526 | SAHH_HUMAN | SAHH_HUMAN | Adenosylhomocysteinase (AdoHcyase) (EC 3.3.1.1) (S-adenosyl-L-homocysteine hydrolase) | AHCY SAHH |
| sp\|Q9UJS0\|CMC2_HUMAN | 97,298 | 114,085 | 118,276 | 105,219 | sp\|Q9UJS0\|CMC2_HUMAN | 89,323 | 26,264 | 29% | Q9UJS0 | CMC2_HUMAN | CMC2_HUMAN | Calcium-binding mitochondrial carrier protein Aralar2 (Citrin) (Mitochondrial aspartate glutamate carrier 2) (Solute carrier family 25 member 13) | SLC25A13 ARALAR2 |
| sp\|P04899\|GNAI2_HUMAN | 2,691,214 | 2,937,093 | 3,591,818 | 3,400,440 | sp\|P04899\|GNAI2_HUMAN | 3,172,363 | 417,870 | 13% | P04899 | GNAI2_HUMAN | GNAI2_HUMAN | Guanine nucleotide-binding protein G(i) subunit alpha-2 (Adenylate cyclase-inhibiting G alpha protein) | GNAI2 GNAI2B |
| sp\|P0DMV9\|HS71B_HUMAN | 1,074,479 | 1,353,366 | 861,065 | 1,184,664 | sp\|P0DMV9\|HS71B_HUMAN | 1,204,327 | 174,335 | 14% | P0DMV9 | HS71B_HUMAN | HS71B_HUMAN | Heat shock 70 kDa protein 1B (Heat shock 70 kDa protein 2) (HSP70-2) (HSP70.2) | HSPA1B HSP72 |
| sp\|P50454\|SERPH_HUMAN | 332,035 | 285,810 | 472,489 | 339,779 | sp\|P50454\|SERPH_HUMAN | 289,055 | 91,940 | 32% | P50454 | SERPH_HUMAN | SERPH_HUMAN | Serpin H1 (47 kDa heat shock protein) (Arsenic-transactivated protein 3) (AsTP3) (Cell proliferation-inducing gene 14 protein) (Collagen-binding protein) (Colligin) (Rheumatoid arthritis-related antigen RA-A47) | SERPINH1 CBP1 CBP2 HSP47 SERPINH2 PIG14 |

| | | | | | | | | | | | | | |
|---|---|---|---|---|---|---|---|---|---|---|---|---|---|
| sp\|P34897\|GLYM_HUMAN | 449,624 | 440,398 | 767,015 | 628,586 | sp\|P34897\|GLYM_HUMAN | 468,261 | 174,942 | 37% | P34897 | GLYM_HUMAN | GLYM_HUMAN | Serine hydroxymethyltransferase, mitochondrial (SHMT) (EC 2.1.2.1) (Glycine hydroxymethyltransferase) (Serine methylase) | SHMT2 |
| sp\|P31948\|STIP1_HUMAN | 527,271 | 645,328 | 722,744 | 894,095 | sp\|P31948\|STIP1_HUMAN | 610,685 | 140,246 | 23% | P31948 | STIP1_HUMAN | STIP1_HUMAN | Stress-induced-phosphoprotein 1 (STI1) (Hsc70/Hsp90-organizing protein) (Hop) (Renal carcinoma antigen NY-REN-11) (Transformation-sensitive protein IEF SSP 3521) | STIP1 |
| sp\|P26640\|SYVC_HUMAN | 242,200 | 259,179 | 396,420 | 159,919 | sp\|P26640\|SYVC_HUMAN | 243,868 | 78,384 | 32% | P26640 | SYVC_HUMAN | SYVC_HUMAN | Valine--tRNA ligase (EC 6.1.1.9) (Protein G7a) (Valyl-tRNA synthetase) (ValRS) | VARS1 G7A VARS VARS2 |
| sp\|P14868\|SYDC_HUMAN | 624,965 | 523,435 | 584,642 | 617,962 | sp\|P14868\|SYDC_HUMAN | 472,435 | 144,901 | 31% | P14868 | SYDC_HUMAN | SYDC_HUMAN | Aspartate--tRNA ligase, cytoplasmic (EC 6.1.1.12) (Aspartyl-tRNA synthetase) (AspRS) (Cell proliferation-inducing gene 40 protein) | DARS1 DARS PIG40 |
| sp\|Q13200\|PSMD2_HUMAN | 437,911 | 379,156 | 509,528 | 357,241 | sp\|Q13200\|PSMD2_HUMAN | 395,357 | 98,825 | 25% | Q13200 | PSMD2_HUMAN | PSMD2_HUMAN | 26S proteasome non-ATPase regulatory subunit 2 (26S proteasome regulatory subunit RPN1) (26S proteasome regulatory subunit S2) (26S proteasome subunit p97) (Protein 55.11) (Tumor necrosis factor type 1 receptor-associated protein 2) | PSMD2 TRAP2 |
| sp\|P05534\|1A24_HUMAN | 3,649,052 | 3,300,206 | 4,265,879 | 4,221,038 | sp\|P05534\|1A24_HUMAN | 3,211,027 | 845,825 | 26% | P05534 | 1A24_HUMAN | HLAA_HUMAN | HLA class I histocompatibility antigen, A alpha chain (Human leukocyte antigen A) (HLA-A) | HLA-A HLAA |
| sp\|P55084\|ECHB_HUMAN | 569,340 | 183,205 | 336,247 | 209,003 | sp\|P55084\|ECHB_HUMAN | 264,092 | 208,934 | 79% | P55084 | ECHB_HUMAN | ECHB_HUMAN | Trifunctional enzyme subunit beta, mitochondrial (TP-beta) [Includes: 3-ketoacyl-CoA thiolase (EC 2.3.1.155) (EC 2.3.1.16) (Acetyl-CoA acyltransferase) (Beta-ketothiolase)] | HADHB MSTP029 |
| sp\|P27797\|CALR_HUMAN | 465,994 | 544,932 | 320,767 | 347,130 | sp\|P27797\|CALR_HUMAN | 314,139 | 138,438 | 44% | P27797 | CALR_HUMAN | CALR_HUMAN | Calreticulin (CRP55) (Calregulin) (Endoplasmic reticulum resident protein 60) (ERp60) (HACBP) (grp60) | CALR CRTC |
| sp\|P00505\|AATM_HUMAN | 416,439 | 501,811 | 647,881 | 834,844 | sp\|P00505\|AATM_HUMAN | 496,942 | 190,467 | 38% | P00505 | AATM_HUMAN | AATM_HUMAN | Aspartate aminotransferase, mitochondrial (mAspAT) (EC 2.6.1.1) (EC 2.6.1.7) (Fatty acid-binding protein) (FABP-1) (Glutamate oxaloacetate transaminase 2) (Kynurenine aminotransferase 4) (Kynurenine aminotransferase IV) (Kynurenine--oxoglutarate transaminase 4) (Kynurenine--oxoglutarate transaminase IV) (Plasma membrane-associated fatty acid-binding protein) (FABPpm) (Transaminase A) | GOT2 KYAT4 |
| sp\|O95782\|AP2A1_HUMAN | 749,874 | 810,254 | 864,987 | 857,997 | sp\|O95782\|AP2A1_HUMAN | 774,309 | 98,489 | 13% | O95782 | AP2A1_HUMAN | AP2A1_HUMAN | AP-2 complex subunit alpha-1 (100 kDa coated vesicle protein A) (Adaptor protein complex AP-2 subunit alpha-1) (Adaptor-related protein complex 2 subunit alpha-1) (Alpha-adaptin A) (Alpha1-adaptin) (Clathrin assembly protein complex 2 alpha-A large chain) (Plasma membrane adaptor HA2/AP2 adaptin alpha A subunit) | AP2A1 ADTAA CLAPA1 |

| | | | | | | | | | | | | | |
|---|---|---|---|---|---|---|---|---|---|---|---|---|---|
| sp\|P36578\|RL4_HUMAN | 993,122 | 1,264,199 | 1,076,698 | 962,498 | sp\|P36578\|RL4_HUMAN | 735,730 | 379,595 | 52% | P36578 | RL4_HUMAN | RL4_HUMAN | 60S ribosomal protein L4 (60S ribosomal protein L1) (Large ribosomal subunit protein uL4) | RPL4 RPL1 |
| sp\|Q06830\|PRDX1_HUMAN | 2,431,838 | 2,786,812 | 1,872,300 | 2,120,495 | sp\|Q06830\|PRDX1_HUMAN | 1,963,058 | 512,650 | 26% | Q06830 | PRDX1_HUMAN | PRDX1_HUMAN | Peroxiredoxin-1 (EC 1.11.1.24) (Natural killer cell-enhancing factor A) (NKEF-A) (Proliferation-associated gene protein) (PAG) (Thioredoxin peroxidase 2) (Thioredoxin-dependent peroxide reductase 2) (Thioredoxin-dependent peroxiredoxin 1) | PRDX1 PAGA PAGB TDPX2 |
| sp\|P61247\|RS3A_HUMAN | 2,262,537 | 2,313,940 | 1,877,307 | 1,579,337 | sp\|P61247\|RS3A_HUMAN | 1,415,539 | 684,144 | 48% | P61247 | RS3A_HUMAN | RS3A_HUMAN | 40S ribosomal protein S3a (Small ribosomal subunit protein eS1) (v-fos transformation effector protein) (Fte-1) | RPS3A FTE1 MFTL |
| sp\|P15311\|EZRI_HUMAN | 714,338 | 1,135,579 | 507,072 | 731,400 | sp\|P15311\|EZRI_HUMAN | 759,051 | 206,667 | 27% | P15311 | EZRI_HUMAN | EZRI_HUMAN | Ezrin (Cytovillin) (Villin-2) (p81) | EZR VIL2 |
| sp\|P13647\|K2C5_HUMAN | 1,747,451 | 924,854 | 2,056,512 | 1,635,196 | sp\|P13647\|K2C5_HUMAN | 2,155,530 | 880,662 | 41% | P13647 | K2C5_HUMAN | K2C5_HUMAN | Keratin, type II cytoskeletal 5 (58 kDa cytokeratin) (Cytokeratin-5) (CK-5) (Keratin-5) (K5) (Type-II keratin Kb5) | KRT5 |
| sp\|P34932\|HSP74_HUMAN | 351,621 | 441,120 | 399,640 | 475,202 | sp\|P34932\|HSP74_HUMAN | 393,796 | 56,902 | 14% | P34932 | HSP74_HUMAN | HSP74_HUMAN | Heat shock 70 kDa protein 4 (HSP70RY) (Heat shock 70-related protein APG-2) | HSPA4 APG2 |
| sp\|P54577\|SYYC_HUMAN | 253,375 | 278,683 | 224,021 | 211,557 | sp\|P54577\|SYYC_HUMAN | 241,316 | 33,935 | 14% | P54577 | SYYC_HUMAN | SYYC_HUMAN | Tyrosine--tRNA ligase, cytoplasmic (EC 6.1.1.1) (Tyrosyl-tRNA synthetase) (TyrRS) [Cleaved into: Tyrosine--tRNA ligase, cytoplasmic, N-terminally processed] | YARS1 YARS |
| sp\|P09525\|ANXA4_HUMAN | 3,999,889 | 2,661,992 | 6,057,969 | 4,232,876 | sp\|P09525\|ANXA4_HUMAN | 3,368,110 | 1,389,262 | 41% | P09525 | ANXA4_HUMAN | ANXA4_HUMAN | Annexin A4 (35-beta calcimedin) (Annexin IV) (Annexin-4) (Carbohydrate-binding protein p33/p41) (Chromobindin-4) (Endonexin I) (Lipocortin IV) (P32.5) (PP4-X) (Placental anticoagulant protein II) (PAP-II) (Protein II) | ANXA4 ANX4 |
| sp\|P07195\|LDHB_HUMAN | 2,986,465 | 4,092,369 | 1,774,472 | 2,375,035 | sp\|P07195\|LDHB_HUMAN | 2,829,586 | 923,897 | 33% | P07195 | LDHB_HUMAN | LDHB_HUMAN | L-lactate dehydrogenase B chain (LDH-B) (EC 1.1.1.27) (LDH heart subunit) (LDH-H) (Renal carcinoma antigen NY-REN-46) | LDHB |
| sp\|P21796\|VDAC1_HUMAN | 370,231 | 418,060 | 192,489 | 246,255 | sp\|P21796\|VDAC1_HUMAN | 255,308 | 89,390 | 35% | P21796 | VDAC1_HUMAN | VDAC1_HUMAN | Voltage-dependent anion-selective channel protein 1 (VDAC-1) (hVDAC1) (Outer mitochondrial membrane protein porin 1) (Plasmalemmal porin) (Porin 31HL) (Porin 31HM) | VDAC1 VDAC |
| sp\|Q99623\|PHB2_HUMAN | 412,843 | 555,706 | 521,065 | 987,530 | sp\|Q99623\|PHB2_HUMAN | 472,010 | 264,166 | 56% | Q99623 | PHB2_HUMAN | PHB2_HUMAN | Prohibitin-2 (B-cell receptor-associated protein BAP37) (D-prohibitin) (Repressor of estrogen receptor activity) | PHB2 BAP REA |
| sp\|P06744\|G6PI_HUMAN | 1,044,223 | 1,325,771 | 354,035 | 453,626 | sp\|P06744\|G6PI_HUMAN | 709,276 | 372,605 | 53% | P06744 | G6PI_HUMAN | G6PI_HUMAN | Glucose-6-phosphate isomerase (GPI) (EC 5.3.1.9) (Autocrine motility factor) (AMF) (Neuroleukin) (NLK) (Phosphoglucose isomerase) (PGI) (Phosphohexose isomerase) (PHI) (Sperm antigen 36) (SA-36) | GPI |
| sp\|P13489\|RINI_HUMAN | 1,029,875 | 1,052,966 | 1,358,156 | 1,303,054 | sp\|P13489\|RINI_HUMAN | 914,623 | 311,665 | 34% | P13489 | RINI_HUMAN | RINI_HUMAN | Ribonuclease inhibitor (Placental ribonuclease inhibitor) (Placental RNase inhibitor) (Ribonuclease/angiogenin inhibitor 1) (RAI) | RNH1 PRI RNH |

| | | | | | | | | | | | | | |
|---|---|---|---|---|---|---|---|---|---|---|---|---|---|
| sp\|P23528\|COF1_HUMAN | 3,964,186 | 5,002,649 | 2,104,992 | 2,362,480 | sp\|P23528\|COF1_HUMAN | 2,677,299 | 1,338,349 | 50% | P23528 | COF1_HUMAN | COF1_HUMAN | Cofilin-1 (18 kDa phosphoprotein) (p18) (Cofilin, non-muscle isoform) | CFL1 CFL |
| sp\|P63244\|RACK1_HUMAN | 1,381,559 | 1,455,274 | 1,279,333 | 1,232,763 | sp\|P63244\|RACK1_HUMAN | 998,371 | 416,374 | 42% | P63244 | RACK1_HUMAN | RACK1_HUMAN | Receptor of activated protein C kinase 1 (Cell proliferation-inducing gene 21 protein) (Guanine nucleotide-binding protein subunit beta-2-like 1) (Guanine nucleotide-binding protein subunit beta-like protein 12.3) (Human lung cancer oncogene 7 protein) (HLC-7) (Receptor for activated C kinase) (Small ribosomal subunit protein RACK1) [Cleaved into: Receptor of activated protein C kinase 1, N-terminally processed (Guanine nucleotide-binding protein subunit beta-2-like 1, N-terminally processed)] | RACK1 GNB2L1 HLC7 PIG21 |
| sp\|P35232\|PHB_HUMAN | 357,100 | 408,954 | 442,358 | 572,645 | sp\|P35232\|PHB_HUMAN | 349,272 | 123,868 | 35% | P35232 | PHB_HUMAN | PHB1_HUMAN | Prohibitin 1 | PHB1 PHB |
| sp\|P04083\|ANXA1_HUMAN | 1,361,268 | 1,253,538 | 1,675,057 | 1,331,630 | sp\|P04083\|ANXA1_HUMAN | 1,214,912 | 245,750 | 20% | P04083 | ANXA1_HUMAN | ANXA1_HUMAN | Annexin A1 (Annexin I) (Annexin-1) (Calpactin II) (Calpactin-2) (Chromobindin-9) (Lipocortin I) (Phospholipase A2 inhibitory protein) (p35) [Cleaved into: Annexin Ac2-26] | ANXA1 ANX1 LPC1 |
| sp\|P33176\|KINH_HUMAN | 538,752 | 570,195 | 441,182 | 502,279 | sp\|P33176\|KINH_HUMAN | 436,040 | 106,952 | 25% | P33176 | KINH_HUMAN | KINH_HUMAN | Kinesin-1 heavy chain (Conventional kinesin heavy chain) (Ubiquitous kinesin heavy chain) (UKHC) | KIF5B KNS KNS1 |
| sp\|P62701\|RS4X_HUMAN | 2,573,061 | 2,232,718 | 1,705,110 | 2,130,389 | sp\|P62701\|RS4X_HUMAN | 1,502,587 | 749,832 | 50% | P62701 | RS4X_HUMAN | RS4X_HUMAN | 40S ribosomal protein S4, X isoform (SCR10) (Single copy abundant mRNA protein) (Small ribosomal subunit protein eS4) | RPS4X CCG2 RPS4 SCAR |
| sp\|P23284\|PPIB_HUMAN | 1,524,904 | 2,434,753 | 1,849,263 | 1,797,708 | sp\|P23284\|PPIB_HUMAN | 1,404,086 | 599,017 | 43% | P23284 | PPIB_HUMAN | PPIB_HUMAN | Peptidyl-prolyl cis-trans isomerase B (PPIase B) (EC 5.2.1.8) (CYP-S1) (Cyclophilin B) (Rotamase B) (S-cyclophilin) (SCYLP) | PPIB CYPB |
| sp\|Q9NPH3\|IL1AP_HUMAN | 1,138,357 | 1,239,943 | 1,026,188 | 1,443,005 | sp\|Q9NPH3\|IL1AP_HUMAN | 1,087,590 | 251,400 | 23% | Q9NPH3 | IL1AP_HUMAN | IL1AP_HUMAN | Interleukin-1 receptor accessory protein (IL-1 receptor accessory protein) (IL-1RAcP) (EC 3.2.2.6) (Interleukin-1 receptor 3) (IL-1R-3) (IL-1R3) | IL1RAP C3orf13 IL1R3 |
| sp\|P63104\|1433Z_HUMAN | 2,893,041 | 3,891,198 | 1,169,064 | 1,800,139 | sp\|P63104\|1433Z_HUMAN | 2,280,447 | 1,071,655 | 47% | P63104 | 1433Z_HUMAN | 1433Z_HUMAN | 14-3-3 protein zeta/delta (Protein kinase C inhibitor protein 1) (KCIP-1) | YWHAZ |
| sp\|Q15084\|PDIA6_HUMAN | 800,394 | 1,037,079 | 736,945 | 1,199,833 | sp\|Q15084\|PDIA6_HUMAN | 759,033 | 246,394 | 32% | Q15084 | PDIA6_HUMAN | PDIA6_HUMAN | Protein disulfide-isomerase A6 (EC 5.3.4.1) (Endoplasmic reticulum protein 5) (ER protein 5) (ERp5) (Protein disulfide isomerase P5) (Thioredoxin domain-containing protein 7) | PDIA6 ERP5 P5 TXNDC7 |
| sp\|P16615\|AT2A2_HUMAN | 113,552 | 94,405 | 109,095 | 79,469 | sp\|P16615\|AT2A2_HUMAN | 103,768 | 40,430 | 39% | P16615 | AT2A2_HUMAN | AT2A2_HUMAN | Sarcoplasmic/endoplasmic reticulum calcium ATPase 2 (SERCA2) (SR Ca(2+)-ATPase 2) (EC 7.2.2.10) (Calcium pump 2) (Calcium-transporting ATPase sarcoplasmic reticulum type, slow twitch skeletal muscle isoform) (Endoplasmic reticulum class 1/2 Ca(2+) ATPase) | ATP2A2 ATP2B |

| | | | | | | | | | | | | | |
|---|---|---|---|---|---|---|---|---|---|---|---|---|---|
| sp\|P01024\|CO3_HUMAN | 480,098 | 333,767 | 332,795 | 167,684 | sp\|P01024\|CO3_HUMAN | 4,862,901 | 5,080,550 | 104% | P01024 | CO3_HUMAN | CO3_HUMAN | Complement C3 (C3 and PZP-like alpha-2-macroglobulin domain-containing protein 1) [Cleaved into: Complement C3 beta chain; C3-beta-c (C3bc); Complement C3 alpha chain; C3a anaphylatoxin; Acylation stimulating protein (ASP) (C3adesArg); Complement C3b alpha' chain; Complement C3c alpha' chain fragment 1; Complement C3dg fragment; Complement C3g fragment; Complement C3d fragment; Complement C3f fragment; Complement C3c alpha' chain fragment 2] | C3 CPAMD1 |
| sp\|O00154\|BACH_HUMAN | 1,614,435 | 1,666,363 | 1,209,504 | 1,283,288 | sp\|O00154\|BACH_HUMAN | 1,133,014 | 386,024 | 34% | O00154 | BACH_HUMAN | BACH_HUMAN | Cytosolic acyl coenzyme A thioester hydrolase (EC 3.1.2.2) (Acyl-CoA thioesterase 7) (Brain acyl-CoA hydrolase) (BACH) (hBACH) (CTE-IIa) (CTE-II) (Long chain acyl-CoA thioester hydrolase) | ACOT7 BACH |
| sp\|P23396\|RS3_HUMAN | 2,524,745 | 2,545,371 | 2,896,348 | 2,780,858 | sp\|P23396\|RS3_HUMAN | 1,884,753 | 886,959 | 47% | P23396 | RS3_HUMAN | RS3_HUMAN | 40S ribosomal protein S3 (EC 4.2.99.18) (Small ribosomal subunit protein uS3) | RPS3 OK/SW-cl.26 |
| sp\|Q99536\|VAT1_HUMAN | 1,391,699 | 2,079,884 | 1,035,227 | 1,334,044 | sp\|Q99536\|VAT1_HUMAN | 1,266,401 | 423,783 | 33% | Q99536 | VAT1_HUMAN | VAT1_HUMAN | Synaptic vesicle membrane protein VAT-1 homolog (EC 1.-.-.-) | VAT1 |
| sp\|O75131\|CPNE3_HUMAN | 888,458 | 744,143 | 1,024,768 | 626,506 | sp\|O75131\|CPNE3_HUMAN | 719,573 | 204,424 | 28% | O75131 | CPNE3_HUMAN | CPNE3_HUMAN | Copine-3 (Copine III) | CPNE3 CPN3 KIAA0636 |
| sp\|P46781\|RS9_HUMAN | 3,994,671 | 3,644,363 | 4,065,387 | 3,252,936 | sp\|P46781\|RS9_HUMAN | 2,580,376 | 1,271,804 | 49% | P46781 | RS9_HUMAN | RS9_HUMAN | 40S ribosomal protein S9 (Small ribosomal subunit protein uS4) | RPS9 |
| sp\|P21281\|VATB2_HUMAN | 323,382 | 295,694 | 465,489 | 532,346 | sp\|P21281\|VATB2_HUMAN | 383,548 | 97,256 | 25% | P21281 | VATB2_HUMAN | VATB2_HUMAN | V-type proton ATPase subunit B, brain isoform (V-ATPase subunit B 2) (Endomembrane proton pump 58 kDa subunit) (HO57) (Vacuolar proton pump subunit B 2) | ATP6V1B2 ATP6B2 VPP3 |
| sp\|O00571\|DDX3X_HUMAN | 704,784 | 659,146 | 835,320 | 765,091 | sp\|O00571\|DDX3X_HUMAN | 580,747 | 182,315 | 31% | O00571 | DDX3X_HUMAN | DDX3X_HUMAN | ATP-dependent RNA helicase DDX3X (EC 3.6.4.13) (CAP-Rf) (DEAD box protein 3, X-chromosomal) (DEAD box, X isoform) (DBX) (Helicase-like protein 2) (HLP2) | DDX3X DBX DDX3 |
| sp\|P18669\|PGAM1_HUMAN | 887,520 | 1,228,147 | 619,237 | 944,927 | sp\|P18669\|PGAM1_HUMAN | 873,905 | 218,796 | 25% | P18669 | PGAM1_HUMAN | PGAM1_HUMAN | Phosphoglycerate mutase 1 (EC 5.4.2.11) (EC 5.4.2.4) (BPG-dependent PGAM 1) (Phosphoglycerate mutase isozyme B) (PGAM-B) | PGAM1 PGAMA CDABP0006 |
| sp\|P39023\|RL3_HUMAN | 922,435 | 1,043,950 | 959,569 | 876,006 | sp\|P39023\|RL3_HUMAN | 724,074 | 248,112 | 34% | P39023 | RL3_HUMAN | RL3_HUMAN | 60S ribosomal protein L3 (HIV-1 TAR RNA-binding protein B) (TARBP-B) (Large ribosomal subunit protein uL3) | RPL3 OK/SW-cl.32 |
| sp\|Q96KP4\|CNDP2_HUMAN | 497,116 | 672,950 | 351,412 | 440,209 | sp\|Q96KP4\|CNDP2_HUMAN | 483,548 | 126,881 | 26% | Q96KP4 | CNDP2_HUMAN | CNDP2_HUMAN | Cytosolic non-specific dipeptidase (EC 3.4.13.18) (CNDP dipeptidase 2) (Glutamate carboxypeptidase-like protein 1) (Peptidase A) (Threonyl dipeptidase) | CNDP2 CN2 CPGL HEL-S-13 PEPA |

| | | | | | | | | | | | | | |
|---|---|---|---|---|---|---|---|---|---|---|---|---|---|
| sp\|O60506\|HNRPQ_HUMAN | 679,940 | 510,572 | 306,765 | 562,076 | sp\|O60506\|HNRPQ_HUMAN | 478,006 | 144,856 | 30% | O60506 | HNRPQ_HUMAN | HNRPQ_HUMAN | Heterogeneous nuclear ribonucleoprotein Q (hnRNP Q) (Glycine- and tyrosine-rich RNA-binding protein) (GRY-RBP) (NS1-associated protein 1) (Synaptotagmin-binding, cytoplasmic RNA-interacting protein) | SYNCRIP HNRPQ NSAP1 |
| sp\|P18124\|RL7_HUMAN | 1,976,762 | 1,778,558 | 1,774,438 | 1,664,724 | sp\|P18124\|RL7_HUMAN | 1,285,652 | 556,459 | 43% | P18124 | RL7_HUMAN | RL7_HUMAN | 60S ribosomal protein L7 (Large ribosomal subunit protein uL30) | RPL7 |
| sp\|P61158\|ARP3_HUMAN | 1,187,705 | 1,485,795 | 956,887 | 1,168,641 | sp\|P61158\|ARP3_HUMAN | 1,021,764 | 242,087 | 24% | P61158 | ARP3_HUMAN | ARP3_HUMAN | Actin-related protein 3 (Actin-like protein 3) | ACTR3 ARP3 |
| sp\|P51149\|RAB7A_HUMAN | 3,058,421 | 3,144,405 | 2,374,998 | 2,738,293 | sp\|P51149\|RAB7A_HUMAN | 2,485,844 | 459,383 | 18% | P51149 | RAB7A_HUMAN | RAB7A_HUMAN | Ras-related protein Rab-7a (EC 3.6.5.2) | RAB7A RAB7 |
| sp\|P11498\|PYC_HUMAN | 24,347 | 28,618 | 51,607 | 33,901 | sp\|P11498\|PYC_HUMAN | 33,721 | 9,546 | 28% | P11498 | PYC_HUMAN | PYC_HUMAN | Pyruvate carboxylase, mitochondrial (EC 6.4.1.1) (Pyruvic carboxylase) (PCB) | PC |
| sp\|O43776\|SYNC_HUMAN | 549,768 | 604,552 | 467,214 | 870,571 | sp\|O43776\|SYNC_HUMAN | 585,771 | 123,491 | 21% | O43776 | SYNC_HUMAN | SYNC_HUMAN | Asparagine--tRNA ligase, cytoplasmic (EC 6.1.1.22) (Asparaginyl-tRNA synthetase) (AsnRS) (Asparaginyl-tRNA synthetase 1) | NARS1 NARS NRS |
| sp\|Q9ULV4\|COR1C_HUMAN | 685,141 | 893,456 | 721,579 | 1,151,787 | sp\|Q9ULV4\|COR1C_HUMAN | 880,288 | 157,542 | 18% | Q9ULV4 | COR1C_HUMAN | COR1C_HUMAN | Coronin-1C (Coronin-3) (hCRNN4) | CORO1C CRN2 CRNN4 |
| sp\|P55268\|LAMB2_HUMAN | 1,900,113 | 1,237,535 | 2,883,323 | 2,610,944 | sp\|P55268\|LAMB2_HUMAN | 1,713,206 | 711,290 | 42% | P55268 | LAMB2_HUMAN | LAMB2_HUMAN | Laminin subunit beta-2 (Laminin B1s chain) (Laminin-11 subunit beta) (Laminin-14 subunit beta) (Laminin-15 subunit beta) (Laminin-3 subunit beta) (Laminin-4 subunit beta) (Laminin-7 subunit beta) (Laminin-9 subunit beta) (S-laminin subunit beta) (S-LAM beta) | LAMB2 LAMS |
| sp\|P60174\|TPIS_HUMAN | 1,296,572 | 1,805,972 | 740,542 | 869,699 | sp\|P60174\|TPIS_HUMAN | 1,003,176 | 497,415 | 50% | P60174 | TPIS_HUMAN | TPIS_HUMAN | Triosephosphate isomerase (TIM) (EC 5.3.1.1) (Methylglyoxal synthase) (EC 4.2.3.3) (Triose-phosphate isomerase) | TPI1 TPI |
| sp\|Q8WWI5\|CTL1_HUMAN | 1,070,719 | 1,443,001 | 1,214,579 | 1,532,872 | sp\|Q8WWI5\|CTL1_HUMAN | 1,143,024 | 244,954 | 21% | Q8WWI5 | CTL1_HUMAN | CTL1_HUMAN | Choline transporter-like protein 1 (CDw92) (Solute carrier family 44 member 1) (CD antigen CD92) | SLC44A1 CD92 CDW92 CTL1 |
| sp\|Q01813\|PFKAP_HUMAN | 720,713 | 709,685 | 312,796 | 520,004 | sp\|Q01813\|PFKAP_HUMAN | 509,609 | 146,682 | 29% | Q01813 | PFKAP_HUMAN | PFKAP_HUMAN | ATP-dependent 6-phosphofructokinase, platelet type (ATP-PFK) (PFK-P) (EC 2.7.1.11) (6-phosphofructokinase type C) (Phosphofructo-1-kinase isozyme C) (PFK-C) (Phosphohexokinase) | PFKP PFKF |
| sp\|P41250\|GARS_HUMAN | 216,589 | 347,184 | 189,843 | 223,283 | sp\|P41250\|GARS_HUMAN | 306,622 | 113,149 | 37% | P41250 | GARS_HUMAN | GARS_HUMAN | Glycine--tRNA ligase (EC 6.1.1.14) (Diadenosine tetraphosphate synthetase) (Ap4A synthetase) (EC 2.7.7.-) (Glycyl-tRNA synthetase) (GlyRS) (Glycyl-tRNA synthetase 1) | GARS1 GARS |
| sp\|Q12965\|MYO1E_HUMAN | 695,926 | 819,875 | 523,693 | 549,643 | sp\|Q12965\|MYO1E_HUMAN | 585,684 | 145,203 | 25% | Q12965 | MYO1E_HUMAN | MYO1E_HUMAN | Unconventional myosin-Ie (Myosin-Ic) (Unconventional myosin 1E) | MYO1E MYO1C |
| sp\|P07384\|CAN1_HUMAN | 918,709 | 1,093,615 | 945,934 | 1,274,244 | sp\|P07384\|CAN1_HUMAN | 886,537 | 214,495 | 24% | P07384 | CAN1_HUMAN | CAN1_HUMAN | Calpain-1 catalytic subunit (EC 3.4.22.52) (Calcium-activated neutral proteinase 1) (CANP 1) (Calpain mu-type) (Calpain-1 large subunit) (Cell proliferation-inducing gene 30 protein) (Micromolar-calpain) (muCANP) | CAPN1 CANPL1 PIG30 |
| sp\|P13797\|PLST_HUMAN | 392,719 | 573,305 | 429,533 | 511,906 | sp\|P13797\|PLST_HUMAN | 486,350 | 57,251 | 12% | P13797 | PLST_HUMAN | PLST_HUMAN | Plastin-3 (T-plastin) | PLS3 |

| | | | | | | | | | | | | | |
|---|---|---|---|---|---|---|---|---|---|---|---|---|---|
| sp\|Q96QK1\|VPS35_HUMAN | 383,849 | 466,379 | 636,475 | 693,310 | sp\|Q96QK1\|VPS35_HUMAN | 477,232 | 135,459 | 28% | Q96QK1 | VPS35_HUMAN | VPS35_HUMAN | Vacuolar protein sorting-associated protein 35 (hVPS35) (Maternal-embryonic 3) (Vesicle protein sorting 35) | VPS35 MEM3 TCCCTA00141 |
| sp\|Q02790\|FKBP4_HUMAN | 204,788 | 248,256 | 202,875 | 251,819 | sp\|Q02790\|FKBP4_HUMAN | 197,372 | 48,924 | 25% | Q02790 | FKBP4_HUMAN | FKBP4_HUMAN | Peptidyl-prolyl cis-trans isomerase FKBP4 (PPIase FKBP4) (EC 5.2.1.8) (51 kDa FK506-binding protein) (FKBP51) (52 kDa FK506-binding protein) (52 kDa FKBP) (FKBP-52) (59 kDa immunophilin) (p59) (FK506-binding protein 4) (FKBP-4) (FKBP59) (HSP-binding immunophilin) (HBI) (Immunophilin FKBP52) (Rotamase) [Cleaved into: Peptidyl-prolyl cis-trans isomerase FKBP4, N-terminally processed] | FKBP4 FKBP52 |
| sp\|Q99798\|ACON_HUMAN | 126,395 | 107,881 | 184,021 | 185,835 | sp\|Q99798\|ACON_HUMAN | 156,196 | 68,187 | 44% | Q99798 | ACON_HUMAN | ACON_HUMAN | Aconitate hydratase, mitochondrial (Aconitase) (EC 4.2.1.3) (Citrate hydro-lyase) | ACO2 |
| sp\|Q06210\|GFPT1_HUMAN | 528,722 | 546,344 | 529,269 | 709,946 | sp\|Q06210\|GFPT1_HUMAN | 521,454 | 98,671 | 19% | Q06210 | GFPT1_HUMAN | GFPT1_HUMAN | Glutamine--fructose-6-phosphate aminotransferase [isomerizing] 1 (EC 2.6.1.16) (D-fructose-6-phosphate amidotransferase 1) (Glutamine:fructose-6-phosphate amidotransferase 1) (GFAT 1) (GFAT1) (Hexosephosphate aminotransferase 1) | GFPT1 GFAT GFPT |
| sp\|P62937\|PPIA_HUMAN | 5,765,489 | 7,101,078 | 5,239,851 | 5,386,835 | sp\|P62937\|PPIA_HUMAN | 5,050,365 | 1,084,248 | 21% | P62937 | PPIA_HUMAN | PPIA_HUMAN | Peptidyl-prolyl cis-trans isomerase A (PPIase A) (EC 5.2.1.8) (Cyclophilin A) (Cyclosporin A-binding protein) (Rotamase A) [Cleaved into: Peptidyl-prolyl cis-trans isomerase A, N-terminally processed] | PPIA CYPA |
| sp\|O00299\|CLIC1_HUMAN | 3,295,524 | 4,327,193 | 4,803,516 | 5,386,507 | sp\|O00299\|CLIC1_HUMAN | 4,130,815 | 753,663 | 18% | O00299 | CLIC1_HUMAN | CLIC1_HUMAN | Chloride intracellular channel protein 1 (Chloride channel ABP) (Nuclear chloride ion channel 27) (NCC27) (Regulatory nuclear chloride ion channel protein) (hRNCC) | CLIC1 G6 NCC27 |
| sp\|P38606\|VATA_HUMAN | 395,648 | 387,557 | 456,150 | 761,702 | sp\|P38606\|VATA_HUMAN | 503,869 | 154,434 | 31% | P38606 | VATA_HUMAN | VATA_HUMAN | V-type proton ATPase catalytic subunit A (V-ATPase subunit A) (EC 7.1.2.2) (V-ATPase 69 kDa subunit) (Vacuolar ATPase isoform VA68) (Vacuolar proton pump subunit alpha) | ATP6V1A ATP6A1 ATP6V1A1 VPP2 |
| sp\|P54886\|P5CS_HUMAN | 70,589 | 106,728 | 105,773 | 100,949 | sp\|P54886\|P5CS_HUMAN | 94,932 | 31,132 | 33% | P54886 | P5CS_HUMAN | P5CS_HUMAN | Delta-1-pyrroline-5-carboxylate synthase (P5CS) (Aldehyde dehydrogenase family 18 member A1) [Includes: Glutamate 5-kinase (GK) (EC 2.7.2.11) (Gamma-glutamyl kinase); Gamma-glutamyl phosphate reductase (GPR) (EC 1.2.1.41) (Glutamate-5-semialdehyde dehydrogenase) (Glutamyl-gamma-semialdehyde dehydrogenase)] | ALDH18A1 GSAS P5CS PYCS |

| | | | | | | | | | | | | | |
|---|---|---|---|---|---|---|---|---|---|---|---|---|---|
| sp\|Q9Y6N5\|SQOR_HUMAN | 268,323 | 150,680 | 135,551 | 155,084 | sp\|Q9Y6N5\|SQOR_HUMAN | 167,022 | 50,612 | 30% | Q9Y6N5 | SQOR_HUMAN | SQOR_HUMAN | Sulfide:quinone oxidoreductase, mitochondrial (SQOR) (EC 1.8.5.8) (Sulfide dehydrogenase-like) (Sulfide quinone oxidoreductase) | SQOR SQRDL CGI-44 |
| sp\|O60568\|PLOD3_HUMAN | 375,623 | 320,957 | 501,652 | 322,456 | sp\|O60568\|PLOD3_HUMAN | 355,907 | 103,569 | 29% | O60568 | PLOD3_HUMAN | PLOD3_HUMAN | Multifunctional procollagen lysine hydroxylase and glycosyltransferase LH3 [Includes: Procollagen-lysine,2-oxoglutarate 5-dioxygenase 3 (EC 1.14.11.4) (Lysyl hydroxylase 3) (LH3); Procollagen glycosyltransferase (EC 2.4.1.50) (EC 2.4.1.66) (Galactosylhydroxylysine-glucosyltransferase) (Procollagen galactosyltransferase) (Procollagen glucosyltransferase)] | PLOD3 |
| sp\|P13612\|ITA4_HUMAN | 1,144,015 | 1,040,987 | 1,685,868 | 1,658,986 | sp\|P13612\|ITA4_HUMAN | 1,331,122 | 307,641 | 23% | P13612 | ITA4_HUMAN | ITA4_HUMAN | Integrin alpha-4 (CD49 antigen-like family member D) (Integrin alpha-IV) (VLA-4 subunit alpha) (CD antigen CD49d) | ITGA4 CD49D |
| sp\|P43007\|SATT_HUMAN | 4,601,147 | 9,288,766 | 3,977,564 | 6,680,607 | sp\|P43007\|SATT_HUMAN | 6,497,620 | 2,171,427 | 33% | P43007 | SATT_HUMAN | SATT_HUMAN | Neutral amino acid transporter A (Alanine/serine/cysteine/threonine transporter 1) (ASCT-1) (Solute carrier family 1 member 4) | SLC1A4 ASCT1 SATT |
| sp\|Q9NVI7\|ATD3A_HUMAN | 73,629 | 54,998 | 84,798 | 72,648 | sp\|Q9NVI7\|ATD3A_HUMAN | 61,416 | 23,397 | 38% | Q9NVI7 | ATD3A_HUMAN | ATD3A_HUMAN | ATPase family AAA domain-containing protein 3A | ATAD3A |
| sp\|P61586\|RHOA_HUMAN | 177,077 | 302,666 | 171,484 | 154,997 | sp\|P61586\|RHOA_HUMAN | 184,232 | 73,525 | 40% | P61586 | RHOA_HUMAN | RHOA_HUMAN | Transforming protein RhoA (EC 3.6.5.2) (Rho cDNA clone 12) (h12) | RHOA ARH12 ARHA RHO12 |
| sp\|Q9Y262\|EIF3L_HUMAN | 265,725 | 340,689 | 411,588 | 524,839 | sp\|Q9Y262\|EIF3L_HUMAN | 317,604 | 107,401 | 34% | Q9Y262 | EIF3L_HUMAN | EIF3L_HUMAN | Eukaryotic translation initiation factor 3 subunit L (eIF3l) (Eukaryotic translation initiation factor 3 subunit 6-interacting protein) (Eukaryotic translation initiation factor 3 subunit E-interacting protein) | EIF3L EIF3EIP EIF3S6IP HSPC021 HSPC025 MSTP005 |
| sp\|P61224\|RAP1B_HUMAN | 771,859 | 857,557 | 199,520 | 242,510 | sp\|P61224\|RAP1B_HUMAN | 509,426 | 332,858 | 65% | P61224 | RAP1B_HUMAN | RAP1B_HUMAN | Ras-related protein Rap-1b (EC 3.6.5.2) (GTP-binding protein smg p21B) | RAP1B OK/SW-cl.11 |
| sp\|Q5JWF2\|GNAS1_HUMAN | 1,050,827 | 1,268,000 | 865,243 | 1,039,791 | sp\|Q5JWF2\|GNAS1_HUMAN | 912,431 | 202,834 | 22% | Q5JWF2 | GNAS1_HUMAN | GNAS1_HUMAN | Guanine nucleotide-binding protein G(s) subunit alpha isoforms XLas (Adenylate cyclase-stimulating G alpha protein) (Extra large alphas protein) (XLalphas) | GNAS GNAS1 |
| sp\|Q92598\|HS105_HUMAN | 337,601 | 336,623 | 313,615 | 521,084 | sp\|Q92598\|HS105_HUMAN | 365,961 | 80,180 | 22% | Q92598 | HS105_HUMAN | HS105_HUMAN | Heat shock protein 105 kDa (Antigen NY-CO-25) (Heat shock 110 kDa protein) | HSPH1 HSP105 HSP110 KIAA0201 |
| sp\|P48147\|PPCE_HUMAN | 407,759 | 423,415 | 473,144 | 573,176 | sp\|P48147\|PPCE_HUMAN | 395,216 | 103,200 | 26% | P48147 | PPCE_HUMAN | PPCE_HUMAN | Prolyl endopeptidase (PE) (EC 3.4.21.26) (Post-proline cleaving enzyme) | PREP PEP |
| sp\|P15880\|RS2_HUMAN | 1,768,612 | 1,627,703 | 1,262,250 | 1,021,857 | sp\|P15880\|RS2_HUMAN | 943,041 | 562,951 | 60% | P15880 | RS2_HUMAN | RS2_HUMAN | 40S ribosomal protein S2 (40S ribosomal protein S4) (Protein LLRep3) (Small ribosomal subunit protein uS5) | RPS2 RPS4 |
| sp\|Q9UHD8\|SEPT9_HUMAN | 478,207 | 649,617 | 237,395 | 242,063 | sp\|Q9UHD8\|SEPT9_HUMAN | 357,556 | 166,438 | 47% | Q9UHD8 | SEPT9_HUMAN | SEPT9_HUMAN | Septin-9 (MLL septin-like fusion protein MSF-A) (MLL septin-like fusion protein) (Ovarian/Breast septin) (Ov/Br septin) (Septin D1) | SEPTIN9 KIAA0991 MSF SEPT9 |

| | | | | | | | | | | | | | |
|---|---|---|---|---|---|---|---|---|---|---|---|---|---|
| sp\|P13611\|CSPG2_HUMAN | 1,356,667 | 1,459,646 | 1,463,288 | 1,777,497 | sp\|P13611\|CSPG2_HUMAN | 1,525,966 | 253,087 | 17% | P13611 | CSPG2_HUMAN | CSPG2_HUMAN | Versican core protein (Chondroitin sulfate proteoglycan core protein 2) (Chondroitin sulfate proteoglycan 2) (Glial hyaluronate-binding protein) (GHAP) (Large fibroblast proteoglycan) (PG-M) | VCAN CSPG2 |
| sp\|Q15758\|AAAT_HUMAN | 1,065,828 | 2,779,764 | 1,964,400 | 3,251,274 | sp\|Q15758\|AAAT_HUMAN | 2,719,582 | 800,282 | 29% | Q15758 | AAAT_HUMAN | AAAT_HUMAN | Neutral amino acid transporter B(0) (ATB(0)) (Baboon M7 virus receptor) (RD114/simian type D retrovirus receptor) (Sodium-dependent neutral amino acid transporter type 2) (Solute carrier family 1 member 5) | SLC1A5 ASCT2 M7V1 RDR RDRC |
| sp\|P23368\|MAOM_HUMAN | 42,286 | 43,819 | 36,575 | 51,112 | sp\|P23368\|MAOM_HUMAN | 49,039 | 14,003 | 29% | P23368 | MAOM_HUMAN | MAOM_HUMAN | NAD-dependent malic enzyme, mitochondrial (NAD-ME) (EC 1.1.1.38) (Malic enzyme 2) | ME2 |
| sp\|Q15365\|PCBP1_HUMAN | 1,040,918 | 1,375,813 | 1,024,869 | 1,047,678 | sp\|Q15365\|PCBP1_HUMAN | 925,498 | 250,520 | 27% | Q15365 | PCBP1_HUMAN | PCBP1_HUMAN | Poly(rC)-binding protein 1 (Alpha-CP1) (Heterogeneous nuclear ribonucleoprotein E1) (hnRNP E1) (Nucleic acid-binding protein SUB2.3) | PCBP1 |
| sp\|P33897\|ABCD1_HUMAN | 94,066 | 117,399 | 86,091 | 115,089 | sp\|P33897\|ABCD1_HUMAN | 91,088 | 23,871 | 26% | P33897 | ABCD1_HUMAN | ABCD1_HUMAN | ATP-binding cassette sub-family D member 1 (EC 3.1.2.-) (EC 7.6.2.-) (Adrenoleukodystrophy protein) (ALDP) | ABCD1 ALD |
| sp\|P05198\|IF2A_HUMAN | 420,893 | 458,245 | 545,089 | 605,352 | sp\|P05198\|IF2A_HUMAN | 589,364 | 208,820 | 35% | P05198 | IF2A_HUMAN | IF2A_HUMAN | Eukaryotic translation initiation factor 2 subunit 1 (Eukaryotic translation initiation factor 2 subunit alpha) (eIF-2-alpha) (eIF-2A) (eIF-2alpha) | EIF2S1 EIF2A |
| sp\|P60981\|DEST_HUMAN | 1,974,185 | 2,518,321 | 1,138,901 | 1,477,707 | sp\|P60981\|DEST_HUMAN | 1,492,763 | 569,488 | 38% | P60981 | DEST_HUMAN | DEST_HUMAN | Destrin (Actin-depolymerizing factor) (ADF) | DSTN ACTDP DSN |
| sp\|P45880\|VDAC2_HUMAN | 542,488 | 641,906 | 295,319 | 338,115 | sp\|P45880\|VDAC2_HUMAN | 324,226 | 179,610 | 55% | P45880 | VDAC2_HUMAN | VDAC2_HUMAN | Voltage-dependent anion-selective channel protein 2 (VDAC-2) (hVDAC2) (Outer mitochondrial membrane protein porin 2) | VDAC2 |
| sp\|P62879\|GBB2_HUMAN | 1,227,010 | 1,474,277 | 788,259 | 1,251,151 | sp\|P62879\|GBB2_HUMAN | 1,116,895 | 215,176 | 19% | P62879 | GBB2_HUMAN | GBB2_HUMAN | Guanine nucleotide-binding protein G(I)/G(S)/G(T) subunit beta-2 (G protein subunit beta-2) (Transducin beta chain 2) | GNB2 |
| sp\|P05388\|RLA0_HUMAN | 1,295,230 | 1,638,536 | 597,490 | 728,611 | sp\|P05388\|RLA0_HUMAN | 772,842 | 487,510 | 63% | P05388 | RLA0_HUMAN | RLA0_HUMAN | 60S acidic ribosomal protein P0 (60S ribosomal protein L10E) (Large ribosomal subunit protein uL10) | RPLP0 |
| sp\|P60228\|EIF3E_HUMAN | 291,130 | 270,767 | 275,279 | 304,720 | sp\|P60228\|EIF3E_HUMAN | 238,221 | 53,755 | 23% | P60228 | EIF3E_HUMAN | EIF3E_HUMAN | Eukaryotic translation initiation factor 3 subunit E (eIF3e) (Eukaryotic translation initiation factor 3 subunit 6) (Viral integration site protein INT-6 homolog) (eIF-3 p48) | EIF3E EIF3S6 INT6 |
| sp\|P38117\|ETFB_HUMAN | 475,389 | 438,916 | 699,301 | 330,246 | sp\|P38117\|ETFB_HUMAN | 425,581 | 130,757 | 31% | P38117 | ETFB_HUMAN | ETFB_HUMAN | Electron transfer flavoprotein subunit beta (Beta-ETF) | ETFB FP585 |
| sp\|O00410\|IPO5_HUMAN | 184,369 | 186,113 | 162,099 | 189,140 | sp\|O00410\|IPO5_HUMAN | 199,846 | 30,942 | 15% | O00410 | IPO5_HUMAN | IPO5_HUMAN | Importin-5 (Imp5) (Importin subunit beta-3) (Karyopherin beta-3) (Ran-binding protein 5) (RanBP5) | IPO5 KPNB3 RANBP5 |
| sp\|P11413\|G6PD_HUMAN | 495,810 | 656,241 | 694,581 | 845,325 | sp\|P11413\|G6PD_HUMAN | 682,495 | 129,837 | 19% | P11413 | G6PD_HUMAN | G6PD_HUMAN | Glucose-6-phosphate 1-dehydrogenase (G6PD) (EC 1.1.1.49) | G6PD |
| sp\|O43852\|CALU_HUMAN | 227,497 | 163,835 | 442,003 | 430,938 | sp\|O43852\|CALU_HUMAN | 240,970 | 132,742 | 55% | O43852 | CALU_HUMAN | CALU_HUMAN | Calumenin (Crocalbin) (IEF SSP 9302) | CALU |

| | | | | | | | | | | | | | |
|---|---|---|---|---|---|---|---|---|---|---|---|---|---|
| sp\|Q10567\|AP1B1_HUMAN | 131,695 | 194,893 | 136,976 | 125,477 | sp\|Q10567\|AP1B1_HUMAN | 133,128 | 31,888 | 24% | Q10567 | AP1B1_HUMAN | AP1B1_HUMAN | AP-1 complex subunit beta-1 (Adaptor protein complex AP-1 subunit beta-1) (Adaptor-related protein complex 1 subunit beta-1) (Beta-1-adaptin) (Beta-adaptin 1) (Clathrin assembly protein complex 1 beta large chain) (Golgi adaptor HA1/AP1 adaptin beta subunit) | AP1B1 ADTB1 BAM22 CLAPB2 |
| sp\|P11047\|LAMC1_HUMAN | 2,024,182 | 2,007,612 | 2,436,959 | 2,342,653 | sp\|P11047\|LAMC1_HUMAN | 1,834,982 | 425,700 | 23% | P11047 | LAMC1_HUMAN | LAMC1_HUMAN | Laminin subunit gamma-1 (Laminin B2 chain) (Laminin-1 subunit gamma) (Laminin-10 subunit gamma) (Laminin-11 subunit gamma) (Laminin-2 subunit gamma) (Laminin-3 subunit gamma) (Laminin-4 subunit gamma) (Laminin-6 subunit gamma) (Laminin-7 subunit gamma) (Laminin-8 subunit gamma) (Laminin-9 subunit gamma) (S-laminin subunit gamma) (S-LAM gamma) | LAMC1 LAMB2 |
| sp\|Q9HDC9\|APMAP_HUMAN | 110,284 | 106,875 | 160,418 | 136,625 | sp\|Q9HDC9\|APMAP_HUMAN | 100,928 | 38,802 | 38% | Q9HDC9 | APMAP_HUMAN | APMAP_HUMAN | Adipocyte plasma membrane-associated protein (Protein BSCv) | APMAP C20orf3 UNQ1869/PRO4305 |
| sp\|P46777\|RL5_HUMAN | 1,036,293 | 862,187 | 750,427 | 713,290 | sp\|P46777\|RL5_HUMAN | 582,442 | 294,383 | 51% | P46777 | RL5_HUMAN | RL5_HUMAN | 60S ribosomal protein L5 (Large ribosomal subunit protein uL18) | RPL5 MSTP030 |
| sp\|Q7L1Q6\|BZW1_HUMAN | 452,849 | 429,458 | 142,635 | 286,446 | sp\|Q7L1Q6\|BZW1_HUMAN | 352,883 | 118,881 | 34% | Q7L1Q6 | BZW1_HUMAN | 5MP2_HUMAN | eIF5-mimic protein 2 (Basic leucine zipper and W2 domain-containing protein 1) (Protein Orf) | BZW1 5MP2 BZAP45 KIAA0005 |
| sp\|P61163\|ACTZ_HUMAN | 228,138 | 257,873 | 218,044 | 335,484 | sp\|P61163\|ACTZ_HUMAN | 233,863 | 48,616 | 21% | P61163 | ACTZ_HUMAN | ACTZ_HUMAN | Alpha-centractin (Centractin) (ARP1) (Actin-RPV) (Centrosome-associated actin homolog) | ACTR1A CTRN1 |
| sp\|Q8NBS9\|TXND5_HUMAN | 166,016 | 213,911 | 151,749 | 312,965 | sp\|Q8NBS9\|TXND5_HUMAN | 182,121 | 79,702 | 44% | Q8NBS9 | TXND5_HUMAN | TXND5_HUMAN | Thioredoxin domain-containing protein 5 (EC 1.8.4.-) (EC 5.3.4.1) (Endoplasmic reticulum resident protein 46) (ER protein 46) (ERp46) (Thioredoxin-like protein p46) | TXNDC5 TLP46 UNQ364/PRO700 |
| sp\|P41091\|IF2G_HUMAN | 417,254 | 426,107 | 325,864 | 341,576 | sp\|P41091\|IF2G_HUMAN | 356,941 | 49,095 | 14% | P41091 | IF2G_HUMAN | IF2G_HUMAN | Eukaryotic translation initiation factor 2 subunit 3 (EC 3.6.5.3) (Eukaryotic translation initiation factor 2 subunit gamma X) (eIF-2-gamma X) (eIF-2gX) | EIF2S3 EIF2G |
| sp\|P61978\|HNRPK_HUMAN | 560,815 | 689,372 | 751,233 | 602,982 | sp\|P61978\|HNRPK_HUMAN | 451,285 | 221,913 | 49% | P61978 | HNRPK_HUMAN | HNRPK_HUMAN | Heterogeneous nuclear ribonucleoprotein K (hnRNP K) (Transformation up-regulated nuclear protein) (TUNP) | HNRNPK HNRPK |
| sp\|P62714\|PP2AB_HUMAN | 350,503 | 514,815 | 351,419 | 402,378 | sp\|P62714\|PP2AB_HUMAN | 378,310 | 99,620 | 26% | P62714 | PP2AB_HUMAN | PP2AB_HUMAN | Serine/threonine-protein phosphatase 2A catalytic subunit beta isoform (PP2A-beta) (EC 3.1.3.16) | PPP2CB |
| sp\|P28838\|AMPL_HUMAN | 335,591 | 372,327 | 292,510 | 218,207 | sp\|P28838\|AMPL_HUMAN | 281,507 | 57,714 | 21% | P28838 | AMPL_HUMAN | AMPL_HUMAN | Cytosol aminopeptidase (EC 3.4.11.1) (Cysteinylglycine-S-conjugate dipeptidase) (EC 3.4.13.23) (Leucine aminopeptidase 3) (LAP-3) (Leucyl aminopeptidase) (Peptidase S) (Proline aminopeptidase) (EC 3.4.11.5) (Prolyl aminopeptidase) | LAP3 LAPEP PEPS |

| | | | | | | | | | | | | | |
|---|---|---|---|---|---|---|---|---|---|---|---|---|---|
| sp\|Q96TA1\|NIBL1_HUMAN | 612,650 | 722,278 | 335,665 | 946,441 | sp\|Q96TA1\|NIBL1_HUMAN | 703,473 | 192,783 | 27% | Q96TA1 | NIBL1_HUMAN | NIBA2_HUMAN | Protein Niban 2 (Meg-3) (Melanoma invasion by ERK) (MINERVA) (Niban-like protein 1) (Protein FAM129B) | NIBAN2 C9orf88 FAM129B |
| sp\|Q00325\|MPCP_HUMAN | 273,689 | 327,033 | 246,516 | 102,904 | sp\|Q00325\|MPCP_HUMAN | 203,901 | 73,541 | 36% | Q00325 | MPCP_HUMAN | MPCP_HUMAN | Phosphate carrier protein, mitochondrial (Phosphate transport protein) (PTP) (Solute carrier family 25 member 3) | SLC25A3 PHC OK/SW-cl.48 |
| sp\|O95373\|IPO7_HUMAN | 224,968 | 239,959 | 121,864 | 271,480 | sp\|O95373\|IPO7_HUMAN | 222,725 | 47,756 | 21% | O95373 | IPO7_HUMAN | IPO7_HUMAN | Importin-7 (Imp7) (Ran-binding protein 7) (RanBP7) | IPO7 RANBP7 |
| sp\|P29401\|TKT_HUMAN | 806,373 | 1,012,480 | 763,485 | 810,579 | sp\|P29401\|TKT_HUMAN | 741,386 | 175,712 | 24% | P29401 | TKT_HUMAN | TKT_HUMAN | Transketolase (TK) (EC 2.2.1.1) | TKT |
| sp\|Q9H223\|EHD4_HUMAN | 921,227 | 1,160,724 | 1,107,315 | 1,209,002 | sp\|Q9H223\|EHD4_HUMAN | 1,045,401 | 219,816 | 21% | Q9H223 | EHD4_HUMAN | EHD4_HUMAN | EH domain-containing protein 4 (Hepatocellular carcinoma-associated protein 10/11) (PAST homolog 4) | EHD4 HCA10 HCA11 PAST4 FKSG7 |
| sp\|P07339\|CATD_HUMAN | 1,079,625 | 937,934 | 848,777 | 679,520 | sp\|P07339\|CATD_HUMAN | 674,349 | 262,590 | 39% | P07339 | CATD_HUMAN | CATD_HUMAN | Cathepsin D (EC 3.4.23.5) [Cleaved into: Cathepsin D light chain; Cathepsin D heavy chain] | CTSD CPSD |
| sp\|P09211\|GSTP1_HUMAN | 1,420,666 | 2,355,739 | 447,630 | 688,411 | sp\|P09211\|GSTP1_HUMAN | 1,014,154 | 739,658 | 73% | P09211 | GSTP1_HUMAN | GSTP1_HUMAN | Glutathione S-transferase P (EC 2.5.1.18) (GST class-pi) (GSTP1-1) | GSTP1 FAEES3 GST3 |
| sp\|P61981\|1433G_HUMAN | 853,581 | 1,104,864 | 732,541 | 865,283 | sp\|P61981\|1433G_HUMAN | 847,609 | 179,656 | 21% | P61981 | 1433G_HUMAN | 1433G_HUMAN | 14-3-3 protein gamma (Protein kinase C inhibitor protein 1) (KCIP-1) [Cleaved into: 14-3-3 protein gamma, N-terminally processed] | YWHAG |
| sp\|P07737\|PROF1_HUMAN | 2,808,049 | 4,239,062 | 1,916,044 | 1,883,991 | sp\|P07737\|PROF1_HUMAN | 2,615,491 | 1,150,883 | 44% | P07737 | PROF1_HUMAN | PROF1_HUMAN | Profilin-1 (Epididymis tissue protein Li 184a) (Profilin I) | PFN1 |
| sp\|P62820\|RAB1A_HUMAN | 387,754 | 526,608 | 184,850 | 252,857 | sp\|P62820\|RAB1A_HUMAN | 299,125 | 112,296 | 38% | P62820 | RAB1A_HUMAN | RAB1A_HUMAN | Ras-related protein Rab-1A (EC 3.6.5.2) (YPT1-related protein) | RAB1A RAB1 |
| sp\|Q99714\|HCD2_HUMAN | 267,846 | 293,524 | 243,760 | 268,460 | sp\|Q99714\|HCD2_HUMAN | 226,420 | 51,758 | 23% | Q99714 | HCD2_HUMAN | HCD2_HUMAN | 3-hydroxyacyl-CoA dehydrogenase type-2 (EC 1.1.1.35) (17-beta-estradiol 17-dehydrogenase) (EC 1.1.1.62) (2-methyl-3-hydroxybutyryl-CoA dehydrogenase) (MHBD) (3-alpha-(17-beta)-hydroxysteroid dehydrogenase (NAD(+))) (EC 1.1.1.239) (3-hydroxy-2-methylbutyryl-CoA dehydrogenase) (EC 1.1.1.178) (3-hydroxyacyl-CoA dehydrogenase type II) (3alpha(or 20beta)-hydroxysteroid dehydrogenase) (EC 1.1.1.53) (7-alpha-hydroxysteroid dehydrogenase) (EC 1.1.1.159) (Endoplasmic reticulum-associated amyloid beta-peptide-binding protein) (Mitochondrial ribonuclease P protein 2) (Mitochondrial RNase P protein 2) (Short chain | HSD17B10 ERAB HADH2 MRPP2 SCHAD SDR5C1 XH98G2 |
| sp\|P43034\|LIS1_HUMAN | 275,331 | 439,794 | 288,555 | 255,085 | sp\|P43034\|LIS1_HUMAN | 325,227 | 107,779 | 33% | P43034 | LIS1_HUMAN | LIS1_HUMAN | Platelet-activating factor acetylhydrolase IB subunit beta (Lissencephaly-1 protein) (LIS-1) (PAF acetylhydrolase 45 kDa subunit) (PAF-AH 45 kDa subunit) (PAF-AH alpha) (PAFAH alpha) | PAFAH1B1 LIS1 MDCR MDS PAFAHA |
| sp\|Q9UHN6\|TMEM2_HUMAN | 673,515 | 642,837 | 805,046 | 753,969 | sp\|Q9UHN6\|TMEM2_HUMAN | 735,918 | 218,630 | 30% | Q9UHN6 | TMEM2_HUMAN | CEIP2_HUMAN | Cell surface hyaluronidase (EC 3.2.1.35) (Cell migration-inducing hyaluronidase 2) (Transmembrane protein 2) | CEMIP2 KIAA1412 TMEM2 |
| sp\|Q16181\|SEPT7_HUMAN | 226,397 | 287,373 | 266,524 | 205,219 | sp\|Q16181\|SEPT7_HUMAN | 235,367 | 49,255 | 21% | Q16181 | SEPT7_HUMAN | SEPT7_HUMAN | Septin-7 (CDC10 protein homolog) | SEPTIN7 CDC10 SEPT7 |

| | | | | | | | | | | | | | |
|---|---|---|---|---|---|---|---|---|---|---|---|---|---|
| sp\|P27105\|STOM_HUMAN | 3,162,149 | 3,660,329 | 2,412,933 | 2,745,080 | sp\|P27105\|STOM_HUMAN | 2,332,367 | 851,335 | 37% | P27105 | STOM_HUMAN | STOM_HUMAN | Stomatin (Erythrocyte band 7 integral membrane protein) (Erythrocyte membrane protein band 7.2) (Protein 7.2b) | STOM BND7 EPB72 |
| sp\|P22626\|ROA2_HUMAN | 593,790 | 639,419 | 500,275 | 701,226 | sp\|P22626\|ROA2_HUMAN | 414,955 | 219,615 | 53% | P22626 | ROA2_HUMAN | ROA2_HUMAN | Heterogeneous nuclear ribonucleoproteins A2/B1 (hnRNP A2/B1) | HNRNPA2B1 HNRPA2B1 |
| sp\|Q02878\|RL6_HUMAN | 826,030 | 872,657 | 522,494 | 534,280 | sp\|Q02878\|RL6_HUMAN | 568,297 | 260,882 | 46% | Q02878 | RL6_HUMAN | RL6_HUMAN | 60S ribosomal protein L6 (Large ribosomal subunit protein eL6) (Neoplasm-related protein C140) (Tax-responsive enhancer element-binding protein 107) (TaxREB107) | RPL6 TXREB1 |
| sp\|P47897\|SYQ_HUMAN | 289,963 | 256,557 | 420,049 | 528,565 | sp\|P47897\|SYQ_HUMAN | 322,401 | 107,839 | 33% | P47897 | SYQ_HUMAN | SYQ_HUMAN | Glutamine--tRNA ligase (EC 6.1.1.18) (Glutaminyl-tRNA synthetase) (GlnRS) | QARS1 QARS |
| sp\|Q9Y265\|RUVB1_HUMAN | 518,200 | 470,161 | 428,949 | 524,604 | sp\|Q9Y265\|RUVB1_HUMAN | 351,706 | 154,048 | 44% | Q9Y265 | RUVB1_HUMAN | RUVB1_HUMAN | RuvB-like 1 (EC 3.6.4.12) (49 kDa TATA box-binding protein-interacting protein) (49 kDa TBP-interacting protein) (54 kDa erythrocyte cytosolic protein) (ECP-54) (INO80 complex subunit H) (Nuclear matrix protein 238) (NMP 238) (Pontin 52) (TIP49a) (TIP60-associated protein 54-alpha) (TAP54-alpha) | RUVBL1 INO80H NMP238 TIP49 TIP49A |
| sp\|P23634\|AT2B4_HUMAN | 467,474 | 520,318 | 378,860 | 458,479 | sp\|P23634\|AT2B4_HUMAN | 430,630 | 49,890 | 12% | P23634 | AT2B4_HUMAN | AT2B4_HUMAN | Plasma membrane calcium-transporting ATPase 4 (PMCA4) (EC 7.2.2.10) (Matrix-remodeling-associated protein 1) (Plasma membrane calcium ATPase isoform 4) (Plasma membrane calcium pump isoform 4) | ATP2B4 ATP2B2 MXRA1 |
| sp\|Q15019\|SEPT2_HUMAN | 405,371 | 503,851 | 309,442 | 555,217 | sp\|Q15019\|SEPT2_HUMAN | 449,780 | 90,384 | 20% | Q15019 | SEPT2_HUMAN | SEPT2_HUMAN | Septin-2 (Neural precursor cell expressed developmentally down-regulated protein 5) (NEDD-5) | SEPTIN2 DIFF6 KIAA0158 NEDD5 SEPT2 |
| sp\|P48735\|IDHP_HUMAN | 149,504 | 182,307 | 113,493 | 115,910 | sp\|P48735\|IDHP_HUMAN | 129,412 | 33,160 | 26% | P48735 | IDHP_HUMAN | IDHP_HUMAN | Isocitrate dehydrogenase [NADP], mitochondrial (IDH) (EC 1.1.1.42) (ICD-M) (IDP) (NADP(+)-specific ICDH) (Oxalosuccinate decarboxylase) | IDH2 |
| sp\|P07942\|LAMB1_HUMAN | 817,198 | 835,458 | 1,323,974 | 845,186 | sp\|P07942\|LAMB1_HUMAN | 928,033 | 285,209 | 31% | P07942 | LAMB1_HUMAN | LAMB1_HUMAN | Laminin subunit beta-1 (Laminin B1 chain) (Laminin-1 subunit beta) (Laminin-10 subunit beta) (Laminin-12 subunit beta) (Laminin-2 subunit beta) (Laminin-6 subunit beta) (Laminin-8 subunit beta) | LAMB1 |
| sp\|Q15907\|RB11B_HUMAN | 3,350,561 | 4,251,580 | 2,899,584 | 4,211,860 | sp\|Q15907\|RB11B_HUMAN | 3,363,993 | 630,055 | 19% | Q15907 | RB11B_HUMAN | RB11B_HUMAN | Ras-related protein Rab-11B (EC 3.6.5.2) (GTP-binding protein YPT3) | RAB11B YPT3 |
| sp\|P28482\|MK01_HUMAN | 534,250 | 406,948 | 369,778 | 437,491 | sp\|P28482\|MK01_HUMAN | 389,007 | 84,690 | 22% | P28482 | MK01_HUMAN | MK01_HUMAN | Mitogen-activated protein kinase 1 (MAP kinase 1) (MAPK 1) (EC 2.7.11.24) (ERT1) (Extracellular signal-regulated kinase 2) (ERK-2) (MAP kinase isoform p42) (p42-MAPK) (Mitogen-activated protein kinase 2) (MAP kinase 2) (MAPK 2) | MAPK1 ERK2 PRKM1 PRKM2 |
| sp\|Q13596\|SNX1_HUMAN | 188,993 | 252,709 | 140,068 | 280,972 | sp\|Q13596\|SNX1_HUMAN | 197,636 | 53,524 | 27% | Q13596 | SNX1_HUMAN | SNX1_HUMAN | Sorting nexin-1 | SNX1 |

| | | | | | | | | | | | | | |
|---|---|---|---|---|---|---|---|---|---|---|---|---|---|
| sp\|Q9NYU2\|UGGG1_HUMAN | 33,940 | 36,407 | 16,549 | 68,076 | sp\|Q9NYU2\|UGGG1_HUMAN | 29,881 | 18,896 | 63% | Q9NYU2 | UGGG1_HUMAN | UGGG1_HUMAN | UDP-glucose:glycoprotein glucosyltransferase 1 (UGT1) (hUGT1) (EC 2.4.1.-) (UDP--Glc:glycoprotein glucosyltransferase) (UDP-glucose ceramide glucosyltransferase-like 1) | UGGT1 GT UGCGL1 UGGT UGT1 UGTR |
| sp\|Q16891\|MIC60_HUMAN | 64,121 | 171,819 | 256,765 | 192,307 | sp\|Q16891\|MIC60_HUMAN | 150,350 | 67,448 | 45% | Q16891 | MIC60_HUMAN | MIC60_HUMAN | MICOS complex subunit MIC60 (Cell proliferation-inducing gene 4/52 protein) (Mitochondrial inner membrane protein) (Mitofilin) (p87/89) | IMMT HMP MIC60 MINOS2 PIG4 PIG52 |
| sp\|P54136\|SYRC_HUMAN | 267,431 | 291,627 | 307,707 | 334,831 | sp\|P54136\|SYRC_HUMAN | 267,394 | 51,288 | 19% | P54136 | SYRC_HUMAN | SYRC_HUMAN | Arginine--tRNA ligase, cytoplasmic (EC 6.1.1.19) (Arginyl-tRNA synthetase) (ArgRS) | RARS1 RARS |
| sp\|P22392\|NDKB_HUMAN | 83,872 | 313,931 | 37,947 | 26,261 | sp\|P22392\|NDKB_HUMAN | 124,262 | 117,307 | 94% | P22392 | NDKB_HUMAN | NDKB_HUMAN | Nucleoside diphosphate kinase B (NDK B) (NDP kinase B) (EC 2.7.4.6) (C-myc purine-binding transcription factor PUF) (Histidine protein kinase NDKB) (EC 2.7.13.3) (nm23-H2) | NME2 NM23B |
| sp\|P04181\|OAT_HUMAN | 93,786 | 143,139 | 76,232 | 132,000 | sp\|P04181\|OAT_HUMAN | 96,220 | 27,007 | 28% | P04181 | OAT_HUMAN | OAT_HUMAN | Ornithine aminotransferase, mitochondrial (EC 2.6.1.13) (Ornithine delta-aminotransferase) (Ornithine--oxo-acid aminotransferase) [Cleaved into: Ornithine aminotransferase, hepatic form; Ornithine aminotransferase, renal form] | OAT |
| sp\|P51148\|RAB5C_HUMAN | 2,239,680 | 2,285,272 | 2,098,251 | 2,484,757 | sp\|P51148\|RAB5C_HUMAN | 1,996,626 | 333,032 | 17% | P51148 | RAB5C_HUMAN | RAB5C_HUMAN | Ras-related protein Rab-5C (EC 3.6.5.2) (L1880) (RAB5L) | RAB5C RABL |
| sp\|P08865\|RSSA_HUMAN | 1,131,437 | 1,235,119 | 1,230,186 | 1,389,947 | sp\|P08865\|RSSA_HUMAN | 1,048,638 | 231,488 | 22% | P08865 | RSSA_HUMAN | RSSA_HUMAN | 40S ribosomal protein SA (37 kDa laminin receptor precursor) (37LRP) (37/67 kDa laminin receptor) (LRP/LR) (67 kDa laminin receptor) (67LR) (Colon carcinoma laminin-binding protein) (Laminin receptor 1) (LamR) (Laminin-binding protein precursor p40) (LBP/p40) (Multidrug resistance-associated protein MGr1-Ag) (NEM/1CHD4) (Small ribosomal subunit protein uS2) | RPSA LAMBR LAMR1 |
| sp\|P17931\|LEG3_HUMAN | 2,431,591 | 3,409,252 | 2,353,706 | 2,606,239 | sp\|P17931\|LEG3_HUMAN | 2,306,732 | 611,174 | 26% | P17931 | LEG3_HUMAN | LEG3_HUMAN | Galectin-3 (Gal-3) (35 kDa lectin) (Carbohydrate-binding protein 35) (CBP 35) (Galactose-specific lectin 3) (Galactoside-binding protein) (GALBP) (IgE-binding protein) (L-31) (Laminin-binding protein) (Lectin L-29) (Mac-2 antigen) | LGALS3 MAC2 |
| sp\|P62333\|PRS10_HUMAN | 456,752 | 471,296 | 346,155 | 524,814 | sp\|P62333\|PRS10_HUMAN | 393,795 | 79,035 | 20% | P62333 | PRS10_HUMAN | PRS10_HUMAN | 26S proteasome regulatory subunit 10B (26S proteasome AAA-ATPase subunit RPT4) (Proteasome 26S subunit ATPase 6) (Proteasome subunit p42) | PSMC6 SUG2 |
| sp\|P52209\|6PGD_HUMAN | 299,369 | 342,187 | 221,351 | 211,638 | sp\|P52209\|6PGD_HUMAN | 295,258 | 89,963 | 30% | P52209 | 6PGD_HUMAN | 6PGD_HUMAN | 6-phosphogluconate dehydrogenase, decarboxylating (EC 1.1.1.44) | PGD PGDH |

| | | | | | | | | | | | | | |
|---|---|---|---|---|---|---|---|---|---|---|---|---|---|
| sp\|P16070\|CD44_HUMAN | 12,011,221 | 13,241,699 | 13,587,051 | 16,130,152 | sp\|P16070\|CD44_HUMAN | 13,580,307 | 1,488,034 | 11% | P16070 | CD44_HUMAN | CD44_HUMAN | CD44 antigen (CDw44) (Epican) (Extracellular matrix receptor III) (ECMR-III) (GP90 lymphocyte homing/adhesion receptor) (HUTCH-I) (Heparan sulfate proteoglycan) (Hermes antigen) (Hyaluronate receptor) (Phagocytic glycoprotein 1) (PGP-1) (Phagocytic glycoprotein I) (PGP-I) (CD antigen CD44) | CD44 LHR MDU2 MDU3 MIC4 |
| sp\|P28331\|NDUS1_HUMAN | 44,231 | 47,500 | 109,689 | 95,188 | sp\|P28331\|NDUS1_HUMAN | 98,457 | 55,874 | 57% | P28331 | NDUS1_HUMAN | NDUS1_HUMAN | NADH-ubiquinone oxidoreductase 75 kDa subunit, mitochondrial (EC 7.1.1.2) (Complex I-75kD) (CI-75kD) | NDUFS1 |
| sp\|P62136\|PP1A_HUMAN | 643,286 | 770,077 | 312,528 | 432,014 | sp\|P62136\|PP1A_HUMAN | 460,337 | 173,478 | 38% | P62136 | PP1A_HUMAN | PP1A_HUMAN | Serine/threonine-protein phosphatase PP1-alpha catalytic subunit (PP-1A) (EC 3.1.3.16) | PPP1CA PPP1A |
| sp\|P47756\|CAPZB_HUMAN | 801,133 | 905,080 | 582,526 | 708,050 | sp\|P47756\|CAPZB_HUMAN | 638,635 | 195,247 | 31% | P47756 | CAPZB_HUMAN | CAPZB_HUMAN | F-actin-capping protein subunit beta (CapZ beta) | CAPZB |
| sp\|Q9H9B4\|SFXN1_HUMAN | 51,938 | 63,432 | 87,356 | 96,225 | sp\|Q9H9B4\|SFXN1_HUMAN | 124,637 | 59,435 | 48% | Q9H9B4 | SFXN1_HUMAN | SFXN1_HUMAN | Sideroflexin-1 | SFXN1 |
| sp\|P35241\|RADI_HUMAN | 432,022 | 646,090 | 178,883 | 274,982 | sp\|P35241\|RADI_HUMAN | 418,937 | 202,090 | 48% | P35241 | RADI_HUMAN | RADI_HUMAN | Radixin | RDX |
| sp\|O95831\|AIFM1_HUMAN | 66,435 | 80,356 | 81,007 | 97,056 | sp\|O95831\|AIFM1_HUMAN | 87,731 | 29,126 | 33% | O95831 | AIFM1_HUMAN | AIFM1_HUMAN | Apoptosis-inducing factor 1, mitochondrial (EC 1.6.99.-) (Programmed cell death protein 8) | AIFM1 AIF PDCD8 |
| sp\|Q8NBQ5\|DHB11_HUMAN | 195,255 | 202,584 | 225,270 | 170,922 | sp\|Q8NBQ5\|DHB11_HUMAN | 169,667 | 39,204 | 23% | Q8NBQ5 | DHB11_HUMAN | DHB11_HUMAN | Estradiol 17-beta-dehydrogenase 11 (EC 1.1.1.62) (17-beta-hydroxysteroid dehydrogenase 11) (17-beta-HSD 11) (17bHSD11) (17betaHSD11) (17-beta-hydroxysteroid dehydrogenase XI) (17-beta-HSD XI) (17betaHSDXI) (Cutaneous T-cell lymphoma-associated antigen HD-CL-03) (CTCL-associated antigen HD-CL-03) (Dehydrogenase/reductase SDR family member 8) (Retinal short-chain dehydrogenase/reductase 2) (retSDR2) (Short chain dehydrogenase/reductase family 16C member 2) | HSD17B11 DHRS8 PAN1B SDR16C2 PSEC0029 UNQ207/PRO233 |
| sp\|O00303\|EIF3F_HUMAN | 315,608 | 346,570 | 277,642 | 153,025 | sp\|O00303\|EIF3F_HUMAN | 239,894 | 67,882 | 28% | O00303 | EIF3F_HUMAN | EIF3F_HUMAN | Eukaryotic translation initiation factor 3 subunit F (eIF3f) (Deubiquitinating enzyme eIF3f) (EC 3.4.19.12) (Eukaryotic translation initiation factor 3 subunit 5) (eIF-3-epsilon) (eIF3 p47) | EIF3F EIF3S5 |
| sp\|Q9Y6M1\|IF2B2_HUMAN | 209,608 | 288,911 | 76,849 | 385,712 | sp\|Q9Y6M1\|IF2B2_HUMAN | 194,837 | 103,996 | 53% | Q9Y6M1 | IF2B2_HUMAN | IF2B2_HUMAN | Insulin-like growth factor 2 mRNA-binding protein 2 (IGF2 mRNA-binding protein 2) (IMP-2) (Hepatocellular carcinoma autoantigen p62) (IGF-II mRNA-binding protein 2) (VICKZ family member 2) | IGF2BP2 IMP2 VICKZ2 |
| sp\|P17655\|CAN2_HUMAN | 226,625 | 353,040 | 138,935 | 321,325 | sp\|P17655\|CAN2_HUMAN | 255,213 | 80,419 | 32% | P17655 | CAN2_HUMAN | CAN2_HUMAN | Calpain-2 catalytic subunit (EC 3.4.22.53) (Calcium-activated neutral proteinase 2) (CANP 2) (Calpain M-type) (Calpain large polypeptide L2) (Calpain-2 large subunit) (Millimolar-calpain) (M-calpain) | CAPN2 CANPL2 |

| | | | | | | | | | | | | | |
|---|---|---|---|---|---|---|---|---|---|---|---|---|---|
| sp\|P06865\|HEXA_HUMAN | 283,969 | 286,694 | 268,040 | 143,685 | sp\|P06865\|HEXA_HUMAN | 209,328 | 61,073 | 29% | P06865 | HEXA_HUMAN | HEXA_HUMAN | Beta-hexosaminidase subunit alpha (EC 3.2.1.52) (Beta-N-acetylhexosaminidase subunit alpha) (Hexosaminidase subunit A) (N-acetyl-beta-glucosaminidase subunit alpha) | HEXA |
| sp\|P17812\|PYRG1_HUMAN | 213,788 | 215,407 | 176,759 | 188,325 | sp\|P17812\|PYRG1_HUMAN | 161,759 | 42,646 | 26% | P17812 | PYRG1_HUMAN | PYRG1_HUMAN | CTP synthase 1 (EC 6.3.4.2) (CTP synthetase 1) (UTP--ammonia ligase 1) | CTPS1 CTPS |
| sp\|Q99541\|PLIN2_HUMAN | 124,127 | 183,404 | 151,406 | 149,294 | sp\|Q99541\|PLIN2_HUMAN | 191,160 | 90,497 | 47% | Q99541 | PLIN2_HUMAN | PLIN2_HUMAN | Perilipin-2 (Adipophilin) (Adipose differentiation-related protein) (ADRP) | PLIN2 ADFP |
| sp\|P08133\|ANXA6_HUMAN | 1,438,377 | 1,078,867 | 1,588,519 | 1,322,552 | sp\|P08133\|ANXA6_HUMAN | 1,058,587 | 353,418 | 33% | P08133 | ANXA6_HUMAN | ANXA6_HUMAN | Annexin A6 (67 kDa calelectrin) (Annexin VI) (Annexin-6) (Calphobindin-II) (CPB-II) (Chromobindin-20) (Lipocortin VI) (Protein III) (p68) (p70) | ANXA6 ANX6 |
| sp\|P12004\|PCNA_HUMAN | 808,296 | 817,466 | 340,601 | 414,788 | sp\|P12004\|PCNA_HUMAN | 480,515 | 220,647 | 46% | P12004 | PCNA_HUMAN | PCNA_HUMAN | Proliferating cell nuclear antigen (PCNA) (Cyclin) | PCNA |
| sp\|P61106\|RAB14_HUMAN | 786,805 | 967,076 | 814,372 | 764,103 | sp\|P61106\|RAB14_HUMAN | 730,701 | 131,600 | 18% | P61106 | RAB14_HUMAN | RAB14_HUMAN | Ras-related protein Rab-14 | RAB14 |
| sp\|Q9Y266\|NUDC_HUMAN | 139,029 | 221,514 | 124,436 | 197,816 | sp\|Q9Y266\|NUDC_HUMAN | 167,916 | 44,702 | 27% | Q9Y266 | NUDC_HUMAN | NUDC_HUMAN | Nuclear migration protein nudC (Nuclear distribution protein C homolog) | NUDC |
| sp\|P41252\|SYIC_HUMAN | 224,916 | 378,361 | 159,500 | 487,779 | sp\|P41252\|SYIC_HUMAN | 267,338 | 111,386 | 42% | P41252 | SYIC_HUMAN | SYIC_HUMAN | Isoleucine--tRNA ligase, cytoplasmic (EC 6.1.1.5) (Isoleucyl-tRNA synthetase) (IRS) (IleRS) | IARS1 IARS |
| sp\|P04792\|HSPB1_HUMAN | 911,084 | 936,185 | 908,671 | 821,615 | sp\|P04792\|HSPB1_HUMAN | 745,834 | 203,892 | 27% | P04792 | HSPB1_HUMAN | HSPB1_HUMAN | Heat shock protein beta-1 (HspB1) (28 kDa heat shock protein) (Estrogen-regulated 24 kDa protein) (Heat shock 27 kDa protein) (HSP 27) (Stress-responsive protein 27) (SRP27) | HSPB1 HSP27 HSP28 |
| sp\|P40121\|CAPG_HUMAN | 720,068 | 660,558 | 858,348 | 1,292,200 | sp\|P40121\|CAPG_HUMAN | 915,337 | 232,394 | 25% | P40121 | CAPG_HUMAN | CAPG_HUMAN | Macrophage-capping protein (Actin regulatory protein CAP-G) | CAPG AFCP MCP |
| sp\|P04632\|CPNS1_HUMAN | 473,251 | 576,171 | 491,424 | 503,697 | sp\|P04632\|CPNS1_HUMAN | 460,100 | 106,423 | 23% | P04632 | CPNS1_HUMAN | CPNS1_HUMAN | Calpain small subunit 1 (CSS1) (Calcium-activated neutral proteinase small subunit) (CANP small subunit) (Calcium-dependent protease small subunit) (CDPS) (Calcium-dependent protease small subunit 1) (Calpain regulatory subunit) | CAPNS1 CAPN4 CAPNS |
| sp\|Q03405\|UPAR_HUMAN | 3,189,519 | 2,896,699 | 3,348,811 | 3,132,646 | sp\|Q03405\|UPAR_HUMAN | 3,136,718 | 844,675 | 27% | Q03405 | UPAR_HUMAN | UPAR_HUMAN | Urokinase plasminogen activator surface receptor (U-PAR) (uPAR) (Monocyte activation antigen Mo3) (CD antigen CD87) | PLAUR MO3 UPAR |
| sp\|Q02543\|RL18A_HUMAN | 1,719,202 | 1,432,928 | 1,577,112 | 1,684,880 | sp\|Q02543\|RL18A_HUMAN | 1,162,156 | 482,020 | 41% | Q02543 | RL18A_HUMAN | RL18A_HUMAN | 60S ribosomal protein L18a (Large ribosomal subunit protein eL20) | RPL18A |
| sp\|P54709\|AT1B3_HUMAN | 3,864,904 | 4,846,595 | 2,901,196 | 3,508,148 | sp\|P54709\|AT1B3_HUMAN | 3,315,565 | 780,712 | 24% | P54709 | AT1B3_HUMAN | AT1B3_HUMAN | Sodium/potassium-transporting ATPase subunit beta-3 (Sodium/potassium-dependent ATPase subunit beta-3) (ATPB-3) (CD antigen CD298) | ATP1B3 |
| sp\|P19823\|ITIH2_HUMAN | 394,680 | 448,286 | 429,083 | 352,558 | sp\|P19823\|ITIH2_HUMAN | 3,008,356 | 2,846,615 | 95% | P19823 | ITIH2_HUMAN | ITIH2_HUMAN | Inter-alpha-trypsin inhibitor heavy chain H2 (ITI heavy chain H2) (ITI-HC2) (Inter-alpha-inhibitor heavy chain 2) (Inter-alpha-trypsin inhibitor complex component II) (Serum-derived hyaluronan-associated protein) (SHAP) | ITIH2 IGHEP2 |

| | | | | | | | | | | | | | |
|---|---|---|---|---|---|---|---|---|---|---|---|---|---|
| sp\|Q9NSD9\|SYFB_HUMAN | 137,405 | 175,656 | 169,824 | 174,566 | sp\|Q9NSD9\|SYFB_HUMAN | 164,368 | 42,404 | 26% | Q9NSD9 | SYFB_HUMAN | SYFB_HUMAN | Phenylalanine--tRNA ligase beta subunit (EC 6.1.1.20) (Phenylalanyl-tRNA synthetase beta subunit) (PheRS) | FARSB FARSLB FRSB HSPC173 |
| sp\|Q13418\|ILK_HUMAN | 437,420 | 454,422 | 274,275 | 338,613 | sp\|Q13418\|ILK_HUMAN | 449,695 | 115,891 | 26% | Q13418 | ILK_HUMAN | ILK_HUMAN | Integrin-linked protein kinase (EC 2.7.11.1) (59 kDa serine/threonine-protein kinase) (Beta-integrin-linked kinase) (ILK-1) (ILK-2) (p59ILK) | ILK ILK1 ILK2 |
| sp\|P78417\|GSTO1_HUMAN | 927,773 | 1,518,579 | 519,422 | 893,555 | sp\|P78417\|GSTO1_HUMAN | 852,770 | 356,295 | 42% | P78417 | GSTO1_HUMAN | GSTO1_HUMAN | Glutathione S-transferase omega-1 (GSTO-1) (EC 2.5.1.18) (Glutathione S-transferase omega 1-1) (GSTO 1-1) (Glutathione-dependent dehydroascorbate reductase) (EC 1.8.5.1) (Monomethylarsonic acid reductase) (MMA(V) reductase) (EC 1.20.4.2) (S-(Phenacyl)glutathione reductase) (SPG-R) | GSTO1 GSTTLP28 |
| sp\|P63000\|RAC1_HUMAN | 1,978,891 | 2,117,890 | 1,420,350 | 1,665,662 | sp\|P63000\|RAC1_HUMAN | 1,522,942 | 493,601 | 32% | P63000 | RAC1_HUMAN | RAC1_HUMAN | Ras-related C3 botulinum toxin substrate 1 (EC 3.6.5.2) (Cell migration-inducing gene 5 protein) (Ras-like protein TC25) (p21-Rac1) | RAC1 TC25 MIG5 |
| sp\|P51659\|DHB4_HUMAN | 2,464,068 | 2,732,054 | 3,596,077 | 4,323,373 | sp\|P51659\|DHB4_HUMAN | 2,982,089 | 834,560 | 28% | P51659 | DHB4_HUMAN | DHB4_HUMAN | Peroxisomal multifunctional enzyme type 2 (MFE-2) (17-beta-hydroxysteroid dehydrogenase 4) (17-beta-HSD 4) (D-bifunctional protein) (DBP) (Multifunctional protein 2) (MFP-2) (Short chain dehydrogenase/reductase family 8C member 1) [Cleaved into: (3R)-hydroxyacyl-CoA dehydrogenase (EC 1.1.1.n12); Enoyl-CoA hydratase 2 (EC 4.2.1.107) (EC 4.2.1.119) (3-alpha,7-alpha,12-alpha-trihydroxy-5-beta-cholest-24-enoyl-CoA hydratase)] | HSD17B4 EDH17B4 SDR8C1 |
| sp\|P04040\|CATA_HUMAN | 87,772 | 81,000 | 62,725 | 67,231 | sp\|P04040\|CATA_HUMAN | 99,147 | 54,292 | 55% | P04040 | CATA_HUMAN | CATA_HUMAN | Catalase (EC 1.11.1.6) | CAT |
| sp\|O15144\|ARPC2_HUMAN | 1,067,455 | 1,268,887 | 994,965 | 1,332,374 | sp\|O15144\|ARPC2_HUMAN | 1,090,984 | 144,245 | 13% | O15144 | ARPC2_HUMAN | ARPC2_HUMAN | Actin-related protein 2/3 complex subunit 2 (Arp2/3 complex 34 kDa subunit) (p34-ARC) | ARPC2 ARC34 PRO2446 |
| sp\|Q01469\|FABP5_HUMAN | 1,358,153 | 1,485,923 | 876,337 | 1,009,977 | sp\|Q01469\|FABP5_HUMAN | 1,157,031 | 392,076 | 34% | Q01469 | FABP5_HUMAN | FABP5_HUMAN | Fatty acid-binding protein 5 (Epidermal-type fatty acid-binding protein) (E-FABP) (Fatty acid-binding protein, epidermal) (Psoriasis-associated fatty acid-binding protein homolog) (PA-FABP) | FABP5 |
| sp\|P62241\|RS8_HUMAN | 1,949,003 | 1,616,916 | 1,541,707 | 1,140,070 | sp\|P62241\|RS8_HUMAN | 1,061,834 | 579,565 | 55% | P62241 | RS8_HUMAN | RS8_HUMAN | 40S ribosomal protein S8 (Small ribosomal subunit protein eS8) | RPS8 OK/SW-cl.83 |
| sp\|Q9Y696\|CLIC4_HUMAN | 651,826 | 1,017,215 | 675,381 | 707,089 | sp\|Q9Y696\|CLIC4_HUMAN | 793,147 | 150,979 | 19% | Q9Y696 | CLIC4_HUMAN | CLIC4_HUMAN | Chloride intracellular channel protein 4 (Intracellular chloride ion channel protein p64H1) | CLIC4 |
| sp\|P52907\|CAZA1_HUMAN | 296,791 | 574,464 | 567,948 | 393,499 | sp\|P52907\|CAZA1_HUMAN | 430,399 | 153,409 | 36% | P52907 | CAZA1_HUMAN | CAZA1_HUMAN | F-actin-capping protein subunit alpha-1 (CapZ alpha-1) | CAPZA1 |
| sp\|P26373\|RL13_HUMAN | 1,912,541 | 1,737,472 | 2,702,342 | 2,437,920 | sp\|P26373\|RL13_HUMAN | 1,696,610 | 645,704 | 38% | P26373 | RL13_HUMAN | RL13_HUMAN | 60S ribosomal protein L13 (Breast basic conserved protein 1) (Large ribosomal subunit protein eL13) | RPL13 BBC1 OK/SW-cl.46 |

| | | | | | | | | | | | | | |
|---|---|---|---|---|---|---|---|---|---|---|---|---|---|
| sp\|P01023\|A2MG_HUMAN | 412,555 | 272,181 | 369,751 | 102,199 | sp\|P01023\|A2MG_HUMAN | 7,807,735 | 8,145,489 | 104% | P01023 | A2MG_HUMAN | A2MG_HUMAN | Alpha-2-macroglobulin (Alpha-2-M) (C3 and PZP-like alpha-2-macroglobulin domain-containing protein 5) | A2M CPAMD5 FWP007 |
| sp\|Q9Y6C9\|MTCH2_HUMAN | 117,477 | 123,944 | 367,728 | 384,747 | sp\|Q9Y6C9\|MTCH2_HUMAN | 195,831 | 133,526 | 68% | Q9Y6C9 | MTCH2_HUMAN | MTCH2_HUMAN | Mitochondrial carrier homolog 2 (Met-induced mitochondrial protein) | MTCH2 MIMP HSPC032 |
| sp\|Q96CW1\|AP2M1_HUMAN | 935,068 | 924,701 | 955,482 | 980,497 | sp\|Q96CW1\|AP2M1_HUMAN | 821,000 | 139,082 | 17% | Q96CW1 | AP2M1_HUMAN | AP2M1_HUMAN | AP-2 complex subunit mu (AP-2 mu chain) (Adaptin-mu2) (Adaptor protein complex AP-2 subunit mu) (Adaptor-related protein complex 2 subunit mu) (Clathrin assembly protein complex 2 mu medium chain) (Clathrin coat assembly protein AP50) (Clathrin coat-associated protein AP50) (HA2 50 kDa subunit) (Plasma membrane adaptor AP-2 50 kDa protein) | AP2M1 CLAPM1 KIAA0109 |
| sp\|Q13011\|ECH1_HUMAN | 159,516 | 257,676 | 225,265 | 272,695 | sp\|Q13011\|ECH1_HUMAN | 201,528 | 59,052 | 29% | Q13011 | ECH1_HUMAN | ECH1_HUMAN | Delta(3,5)-Delta(2,4)-dienoyl-CoA isomerase, mitochondrial (EC 5.3.3.-) | ECH1 |
| sp\|P54920\|SNAA_HUMAN | 497,426 | 736,312 | 398,661 | 631,048 | sp\|P54920\|SNAA_HUMAN | 520,908 | 153,338 | 29% | P54920 | SNAA_HUMAN | SNAA_HUMAN | Alpha-soluble NSF attachment protein (SNAP-alpha) (N-ethylmaleimide-sensitive factor attachment protein alpha) | NAPA SNAPA |
| sp\|Q13162\|PRDX4_HUMAN | 179,279 | 213,528 | 199,879 | 238,179 | sp\|Q13162\|PRDX4_HUMAN | 217,318 | 73,503 | 34% | Q13162 | PRDX4_HUMAN | PRDX4_HUMAN | Peroxiredoxin-4 (EC 1.11.1.24) (Antioxidant enzyme AOE372) (AOE37-2) (Peroxiredoxin IV) (Prx-IV) (Thioredoxin peroxidase AO372) (Thioredoxin-dependent peroxide reductase A0372) (Thioredoxin-dependent peroxiredoxin 4) | PRDX4 |
| sp\|P35625\|TIMP3_HUMAN | 3,370,352 | 2,904,198 | 2,808,337 | 2,324,436 | sp\|P35625\|TIMP3_HUMAN | 2,950,431 | 466,768 | 16% | P35625 | TIMP3_HUMAN | TIMP3_HUMAN | Metalloproteinase inhibitor 3 (Protein MIG-5) (Tissue inhibitor of metalloproteinases 3) (TIMP-3) | TIMP3 |
| sp\|P18085\|ARF4_HUMAN | 549,213 | 579,068 | 532,642 | 606,483 | sp\|P18085\|ARF4_HUMAN | 527,031 | 72,012 | 14% | P18085 | ARF4_HUMAN | ARF4_HUMAN | ADP-ribosylation factor 4 | ARF4 ARF2 |
| sp\|Q99613\|EIF3C_HUMAN | 210,001 | 467,580 | 464,379 | 491,311 | sp\|Q99613\|EIF3C_HUMAN | 365,773 | 99,742 | 27% | Q99613 | EIF3C_HUMAN | EIF3C_HUMAN | Eukaryotic translation initiation factor 3 subunit C (eIF3c) (Eukaryotic translation initiation factor 3 subunit 8) (eIF3 p110) | EIF3C EIF3S8 |
| sp\|Q9Y230\|RUVB2_HUMAN | 671,169 | 631,954 | 982,516 | 981,025 | sp\|Q9Y230\|RUVB2_HUMAN | 626,479 | 250,627 | 40% | Q9Y230 | RUVB2_HUMAN | RUVB2_HUMAN | RuvB-like 2 (EC 3.6.4.12) (48 kDa TATA box-binding protein-interacting protein) (48 kDa TBP-interacting protein) (51 kDa erythrocyte cytosolic protein) (ECP-51) (INO80 complex subunit J) (Repressing pontin 52) (Reptin 52) (TIP49b) (TIP60-associated protein 54-beta) (TAP54-beta) | RUVBL2 INO80J TIP48 TIP49B CGI-46 |
| sp\|P62249\|RS16_HUMAN | 2,600,548 | 2,370,792 | 2,568,401 | 2,120,420 | sp\|P62249\|RS16_HUMAN | 1,663,188 | 820,143 | 49% | P62249 | RS16_HUMAN | RS16_HUMAN | 40S ribosomal protein S16 (Small ribosomal subunit protein uS9) | RPS16 |
| sp\|P61313\|RL15_HUMAN | 776,740 | 658,990 | 1,276,805 | 913,188 | sp\|P61313\|RL15_HUMAN | 593,421 | 382,099 | 64% | P61313 | RL15_HUMAN | RL15_HUMAN | 60S ribosomal protein L15 (Large ribosomal subunit protein eL15) | RPL15 EC45 TCBAP0781 |
| sp\|P52292\|IMA1_HUMAN | 132,486 | 217,190 | 68,198 | 174,007 | sp\|P52292\|IMA1_HUMAN | 151,345 | 47,103 | 31% | P52292 | IMA1_HUMAN | IMA1_HUMAN | Importin subunit alpha-1 (Karyopherin subunit alpha-2) (RAG cohort protein 1) (SRP1-alpha) | KPNA2 RCH1 SRP1 |

| | | | | | | | | | | | | | |
|---|---|---|---|---|---|---|---|---|---|---|---|---|---|
| sp\|P62244\|RS15A_HUMAN | 1,410,813 | 1,756,233 | 1,214,335 | 1,404,446 | sp\|P62244\|RS15A_HUMAN | 1,094,149 | 418,736 | 38% | P62244 | RS15A_HUMAN | RS15A_HUMAN | 40S ribosomal protein S15a (Small ribosomal subunit protein uS8) | RPS15A OK/SW-cl.82 |
| sp\|P26639\|SYTC_HUMAN | 208,443 | 277,502 | 257,356 | 364,057 | sp\|P26639\|SYTC_HUMAN | 279,389 | 44,516 | 16% | P26639 | SYTC_HUMAN | SYTC_HUMAN | Threonine--tRNA ligase 1, cytoplasmic (EC 6.1.1.3) (Threonyl-tRNA synthetase) (ThrRS) (Threonyl-tRNA synthetase 1) | TARS1 TARS |
| sp\|P30040\|ERP29_HUMAN | 107,744 | 84,050 | 77,752 | 90,185 | sp\|P30040\|ERP29_HUMAN | 79,843 | 16,949 | 21% | P30040 | ERP29_HUMAN | ERP29_HUMAN | Endoplasmic reticulum resident protein 29 (ERp29) (Endoplasmic reticulum resident protein 28) (ERp28) (Endoplasmic reticulum resident protein 31) (ERp31) | ERP29 C12orf8 ERP28 |
| sp\|P16152\|CBR1_HUMAN | 528,803 | 621,956 | 709,727 | 844,907 | sp\|P16152\|CBR1_HUMAN | 623,966 | 115,777 | 19% | P16152 | CBR1_HUMAN | CBR1_HUMAN | Carbonyl reductase [NADPH] 1 (EC 1.1.1.184) (15-hydroxyprostaglandin dehydrogenase [NADP(+)]) (EC 1.1.1.196) (EC 1.1.1.197) (20-beta-hydroxysteroid dehydrogenase) (Alcohol dehydrogenase [NAD(P)+] CBR1) (EC 1.1.1.71) (NADPH-dependent carbonyl reductase 1) (Prostaglandin 9-ketoreductase) (PG-9-KR) (Prostaglandin-E(2) 9-reductase) (EC 1.1.1.189) (Short chain dehydrogenase/reductase family 21C member 1) | CBR1 CBR CRN SDR21C1 |
| sp\|P25786\|PSA1_HUMAN | 503,402 | 529,623 | 611,962 | 594,793 | sp\|P25786\|PSA1_HUMAN | 529,464 | 53,053 | 10% | P25786 | PSA1_HUMAN | PSA1_HUMAN | Proteasome subunit alpha type-1 (30 kDa prosomal protein) (PROS-30) (Macropain subunit C2) (Multicatalytic endopeptidase complex subunit C2) (Proteasome component C2) (Proteasome nu chain) | PSMA1 HC2 NU PROS30 PSC2 |
| sp\|P08754\|GNAI3_HUMAN | 1,480,690 | 1,622,538 | 2,102,006 | 2,321,936 | sp\|P08754\|GNAI3_HUMAN | 1,779,547 | 383,885 | 22% | P08754 | GNAI3_HUMAN | GNAI3_HUMAN | Guanine nucleotide-binding protein G(i) subunit alpha-3 (G(i) alpha-3) | GNAI3 |
| sp\|P39656\|OST48_HUMAN | 184,209 | 201,655 | 247,725 | 263,840 | sp\|P39656\|OST48_HUMAN | 161,430 | 75,056 | 46% | P39656 | OST48_HUMAN | OST48_HUMAN | Dolichyl-diphosphooligosaccharide--protein glycosyltransferase 48 kDa subunit (DDOST 48 kDa subunit) (Oligosaccharyl transferase 48 kDa subunit) | DDOST KIAA0115 OST48 OK/SW-cl.45 |
| sp\|P40925\|MDHC_HUMAN | 451,380 | 709,947 | 228,746 | 471,628 | sp\|P40925\|MDHC_HUMAN | 433,787 | 154,954 | 36% | P40925 | MDHC_HUMAN | MDHC_HUMAN | Malate dehydrogenase, cytoplasmic (EC 1.1.1.37) (Cytosolic malate dehydrogenase) (Diiodophenylpyruvate reductase) (EC 1.1.1.96) | MDH1 MDHA |
| sp\|O75340\|PDCD6_HUMAN | 1,479,961 | 1,361,563 | 1,449,262 | 1,424,353 | sp\|O75340\|PDCD6_HUMAN | 1,428,432 | 154,398 | 11% | O75340 | PDCD6_HUMAN | PDCD6_HUMAN | Programmed cell death protein 6 (Apoptosis-linked gene 2 protein homolog) (ALG-2) | PDCD6 ALG2 |
| sp\|Q99829\|CPNE1_HUMAN | 810,449 | 890,224 | 705,085 | 792,010 | sp\|Q99829\|CPNE1_HUMAN | 699,082 | 127,668 | 18% | Q99829 | CPNE1_HUMAN | CPNE1_HUMAN | Copine-1 (Chromobindin 17) (Copine I) | CPNE1 CPN1 |
| sp\|Q9BUF5\|TBB6_HUMAN | 531,877 | 633,971 | 578,993 | 714,699 | sp\|Q9BUF5\|TBB6_HUMAN | 496,291 | 150,127 | 30% | Q9BUF5 | TBB6_HUMAN | TBB6_HUMAN | Tubulin beta-6 chain (Tubulin beta class V) | TUBB6 |
| sp\|P30044\|PRDX5_HUMAN | 504,884 | 668,716 | 226,002 | 334,110 | sp\|P30044\|PRDX5_HUMAN | 347,534 | 181,109 | 52% | P30044 | PRDX5_HUMAN | PRDX5_HUMAN | Peroxiredoxin-5, mitochondrial (EC 1.11.1.24) (Alu corepressor 1) (Antioxidant enzyme B166) (AOEB166) (Liver tissue 2D-page spot 71B) (PLP) (Peroxiredoxin V) (Prx-V) (Peroxisomal antioxidant enzyme) (TPx type VI) (Thioredoxin peroxidase PMP20) (Thioredoxin-dependent peroxiredoxin 5) | PRDX5 ACR1 SBBI10 |

| | | | | | | | | | | | | | |
|---|---|---|---|---|---|---|---|---|---|---|---|---|---|
| sp\|Q9H4M9\|EHD1_HUMAN | 611,079 | 833,373 | 648,889 | 581,906 | sp\|Q9H4M9\|EHD1_HUMAN | 602,107 | 121,087 | 20% | Q9H4M9 | EHD1_HUMAN | EHD1_HUMAN | EH domain-containing protein 1 (PAST homolog 1) (hPAST1) (Testilin) | EHD1 PAST PAST1 CDABP0131 |
| sp\|Q13423\|NNTM_HUMAN | 16,910 | 17,305 | 10,413 | 3,964 | sp\|Q13423\|NNTM_HUMAN | 20,214 | 23,699 | 117% | Q13423 | NNTM_HUMAN | NNTM_HUMAN | NAD(P) transhydrogenase, mitochondrial (EC 7.1.1.1) (Nicotinamide nucleotide transhydrogenase) (Pyridine nucleotide transhydrogenase) | NNT |
| sp\|O75534\|CSDE1_HUMAN | 168,196 | 178,347 | 120,521 | 201,845 | sp\|O75534\|CSDE1_HUMAN | 151,994 | 30,354 | 20% | O75534 | CSDE1_HUMAN | CSDE1_HUMAN | Cold shock domain-containing protein E1 (N-ras upstream gene protein) (Protein UNR) | CSDE1 D1S155E KIAA0885 NRU UNR |
| sp\|P50213\|IDH3A_HUMAN | 61,571 | 68,230 | 102,629 | 119,054 | sp\|P50213\|IDH3A_HUMAN | 73,653 | 30,107 | 41% | P50213 | IDH3A_HUMAN | IDH3A_HUMAN | Isocitrate dehydrogenase [NAD] subunit alpha, mitochondrial (EC 1.1.1.41) (Isocitric dehydrogenase subunit alpha) (NAD(+)-specific ICDH subunit alpha) | IDH3A |
| sp\|Q14344\|GNA13_HUMAN | 624,004 | 789,153 | 750,363 | 902,460 | sp\|Q14344\|GNA13_HUMAN | 778,536 | 134,525 | 17% | Q14344 | GNA13_HUMAN | GNA13_HUMAN | Guanine nucleotide-binding protein subunit alpha-13 (G alpha-13) (G-protein subunit alpha-13) | GNA13 |
| sp\|P07814\|SYEP_HUMAN | 83,342 | 199,910 | 145,941 | 96,634 | sp\|P07814\|SYEP_HUMAN | 121,555 | 50,924 | 42% | P07814 | SYEP_HUMAN | SYEP_HUMAN | Bifunctional glutamate/proline--tRNA ligase (Bifunctional aminoacyl-tRNA synthetase) (Cell proliferation-inducing gene 32 protein) (Glutamatyl-prolyl-tRNA synthetase) [Includes: Glutamate--tRNA ligase (EC 6.1.1.17) (Glutamyl-tRNA synthetase) (GluRS); Proline--tRNA ligase (EC 6.1.1.15) (Prolyl-tRNA synthetase)] | EPRS1 EPRS GLNS PARS QARS QPRS PIG32 |
| sp\|P62753\|RS6_HUMAN | 2,046,316 | 1,918,554 | 1,403,938 | 1,156,477 | sp\|P62753\|RS6_HUMAN | 1,077,519 | 664,847 | 62% | P62753 | RS6_HUMAN | RS6_HUMAN | 40S ribosomal protein S6 (Phosphoprotein NP33) (Small ribosomal subunit protein eS6) | RPS6 OK/SW-cl.2 |
| sp\|P02533\|K1C14_HUMAN | 484,850 | 215,175 | 290,865 | 240,142 | sp\|P02533\|K1C14_HUMAN | 398,578 | 167,023 | 42% | P02533 | K1C14_HUMAN | K1C14_HUMAN | Keratin, type I cytoskeletal 14 (Cytokeratin-14) (CK-14) (Keratin-14) (K14) | KRT14 |
| sp\|P00387\|NB5R3_HUMAN | 156,533 | 260,493 | 215,494 | 204,145 | sp\|P00387\|NB5R3_HUMAN | 179,678 | 54,793 | 30% | P00387 | NB5R3_HUMAN | NB5R3_HUMAN | NADH-cytochrome b5 reductase 3 (B5R) (Cytochrome b5 reductase) (EC 1.6.2.2) (Diaphorase-1) [Cleaved into: NADH-cytochrome b5 reductase 3 membrane-bound form; NADH-cytochrome b5 reductase 3 soluble form] | CYB5R3 DIA1 |
| sp\|P35998\|PRS7_HUMAN | 394,066 | 354,199 | 330,222 | 247,489 | sp\|P35998\|PRS7_HUMAN | 296,159 | 59,862 | 20% | P35998 | PRS7_HUMAN | PRS7_HUMAN | 26S proteasome regulatory subunit 7 (26S proteasome AAA-ATPase subunit RPT1) (Proteasome 26S subunit ATPase 2) | PSMC2 MSS1 |
| sp\|Q9Y3I0\|RTCB_HUMAN | 377,266 | 394,371 | 434,139 | 410,500 | sp\|Q9Y3I0\|RTCB_HUMAN | 321,924 | 93,687 | 29% | Q9Y3I0 | RTCB_HUMAN | RTCB_HUMAN | RNA-splicing ligase RtcB homolog (EC 6.5.1.8) (3'-phosphate/5'-hydroxy nucleic acid ligase) | RTCB C22orf28 HSPC117 |

| | | | | | | | | | | | | | |
|---|---|---|---|---|---|---|---|---|---|---|---|---|---|
| sp\|P13010\|XRCC5_HUMAN | 196,313 | 224,156 | 378,953 | 358,121 | sp\|P13010\|XRCC5_HUMAN | 281,098 | 100,535 | 36% | P13010 | XRCC5_HUMAN | XRCC5_HUMAN | X-ray repair cross-complementing protein 5 (EC 3.6.4.-) (86 kDa subunit of Ku antigen) (ATP-dependent DNA helicase 2 subunit 2) (ATP-dependent DNA helicase II 80 kDa subunit) (CTC box-binding factor 85 kDa subunit) (CTC85) (CTCBF) (DNA repair protein XRCC5) (Ku80) (Ku86) (Lupus Ku autoantigen protein p86) (Nuclear factor IV) (Thyroid-lupus autoantigen) (TLAA) (X-ray repair complementing defective repair in Chinese hamster cells 5 (double-strand-break rejoining)) | XRCC5 G22P2 |
| sp\|P0C0L4\|CO4A_HUMAN | 77,882 | 51,320 | 89,977 | 75,041 | sp\|P0C0L4\|CO4A_HUMAN | 247,667 | 190,972 | 77% | P0C0L4 | CO4A_HUMAN | CO4A_HUMAN | Complement C4-A (Acidic complement C4) (C3 and PZP-like alpha-2-macroglobulin domain-containing protein 2) [Cleaved into: Complement C4 beta chain; Complement C4-A alpha chain; C4a anaphylatoxin; C4b-A; C4d-A; Complement C4 gamma chain] | C4A CO4 CPAMD2 |
| sp\|P36542\|ATPG_HUMAN | 201,861 | 229,993 | 136,526 | 138,796 | sp\|P36542\|ATPG_HUMAN | 154,675 | 45,273 | 29% | P36542 | ATPG_HUMAN | ATPG_HUMAN | ATP synthase subunit gamma, mitochondrial (ATP synthase F1 subunit gamma) (F-ATPase gamma subunit) | ATP5F1C ATP5C ATP5C1 ATP5CL1 |
| sp\|P07741\|APT_HUMAN | 815,298 | 907,337 | 1,144,537 | 1,271,482 | sp\|P07741\|APT_HUMAN | 876,731 | 237,027 | 27% | P07741 | APT_HUMAN | APT_HUMAN | Adenine phosphoribosyltransferase (APRT) (EC 2.4.2.7) | APRT |
| sp\|P31930\|QCR1_HUMAN | 105,795 | 64,556 | 157,898 | 157,709 | sp\|P31930\|QCR1_HUMAN | 98,708 | 40,839 | 41% | P31930 | QCR1_HUMAN | QCR1_HUMAN | Cytochrome b-c1 complex subunit 1, mitochondrial (Complex III subunit 1) (Core protein I) (Ubiquinol-cytochrome-c reductase complex core protein 1) | UQCRC1 |
| sp\|Q96AG4\|LRC59_HUMAN | 52,218 | 57,706 | 40,839 | 27,339 | sp\|Q96AG4\|LRC59_HUMAN | 58,105 | 29,001 | 50% | Q96AG4 | LRC59_HUMAN | LRC59_HUMAN | Leucine-rich repeat-containing protein 59 (Ribosome-binding protein p34) (p34) [Cleaved into: Leucine-rich repeat-containing protein 59, N-terminally processed] | LRRC59 PRO1855 |
| sp\|O75390\|CISY_HUMAN | 320,836 | 316,830 | 214,342 | 378,843 | sp\|O75390\|CISY_HUMAN | 235,595 | 93,500 | 40% | O75390 | CISY_HUMAN | CISY_HUMAN | Citrate synthase, mitochondrial (EC 2.3.3.1) (Citrate (Si)-synthase) | CS |
| sp\|O00232\|PSD12_HUMAN | 236,127 | 252,289 | 168,490 | 275,695 | sp\|O00232\|PSD12_HUMAN | 183,453 | 67,305 | 37% | O00232 | PSD12_HUMAN | PSD12_HUMAN | 26S proteasome non-ATPase regulatory subunit 12 (26S proteasome regulatory subunit RPN5) (26S proteasome regulatory subunit p55) | PSMD12 |
| sp\|P48047\|ATPO_HUMAN | 151,522 | 169,565 | 102,418 | 190,223 | sp\|P48047\|ATPO_HUMAN | 151,513 | 44,546 | 29% | P48047 | ATPO_HUMAN | ATPO_HUMAN | ATP synthase subunit O, mitochondrial (ATP synthase peripheral stalk subunit OSCP) (Oligomycin sensitivity conferral protein) (OSCP) | ATP5PO ATP5O ATPO |
| sp\|P60660\|MYL6_HUMAN | 934,893 | 1,054,469 | 989,608 | 993,766 | sp\|P60660\|MYL6_HUMAN | 758,542 | 254,927 | 34% | P60660 | MYL6_HUMAN | MYL6_HUMAN | Myosin light polypeptide 6 (17 kDa myosin light chain) (LC17) (Myosin light chain 3) (MLC-3) (Myosin light chain alkali 3) (Myosin light chain A3) (Smooth muscle and nonmuscle myosin light chain alkali 6) | MYL6 |
| sp\|Q15046\|SYK_HUMAN | 264,712 | 283,143 | 243,839 | 211,491 | sp\|Q15046\|SYK_HUMAN | 214,837 | 56,353 | 26% | Q15046 | SYK_HUMAN | SYK_HUMAN | Lysine--tRNA ligase (EC 2.7.7.-) (EC 6.1.1.6) (Lysyl-tRNA synthetase) (LysRS) | KARS1 KARS KIAA0070 |

| | | | | | | | | | | | | | |
|---|---|---|---|---|---|---|---|---|---|---|---|---|---|
| sp\|P30086\|PEBP1_HUMAN | 700,073 | 901,848 | 319,556 | 516,219 | sp\|P30086\|PEBP1_HUMAN | 574,954 | 220,623 | 38% | P30086 | PEBP1_HUMAN | PEBP1_HUMAN | Phosphatidylethanolamine-binding protein 1 (PEBP-1) (HCNPpp) (Neuropolypeptide h3) (Prostatic-binding protein) (Raf kinase inhibitor protein) (RKIP) [Cleaved into: Hippocampal cholinergic neurostimulating peptide (HCNP)] | PEBP1 PBP PEBP |
| sp\|P29692\|EF1D_HUMAN | 545,706 | 617,646 | 549,847 | 850,129 | sp\|P29692\|EF1D_HUMAN | 555,629 | 136,555 | 25% | P29692 | EF1D_HUMAN | EF1D_HUMAN | Elongation factor 1-delta (EF-1-delta) (Antigen NY-CO-4) | EEF1D EF1D |
| sp\|Q06033\|ITIH3_HUMAN | 109,909 | 74,183 | 92,607 | 69,110 | sp\|Q06033\|ITIH3_HUMAN | 634,077 | 842,811 | 133% | Q06033 | ITIH3_HUMAN | ITIH3_HUMAN | Inter-alpha-trypsin inhibitor heavy chain H3 (ITI heavy chain H3) (ITI-HC3) (Inter-alpha-inhibitor heavy chain 3) (Serum-derived hyaluronan-associated protein) (SHAP) | ITIH3 |
| sp\|O43242\|PSMD3_HUMAN | 675,085 | 768,837 | 431,414 | 103,780 | sp\|O43242\|PSMD3_HUMAN | 356,840 | 263,530 | 74% | O43242 | PSMD3_HUMAN | PSMD3_HUMAN | 26S proteasome non-ATPase regulatory subunit 3 (26S proteasome regulatory subunit RPN3) (26S proteasome regulatory subunit S3) (Proteasome subunit p58) | PSMD3 |
| sp\|O00339\|MATN2_HUMAN | 1,241,772 | 1,213,041 | 1,464,565 | 1,625,965 | sp\|O00339\|MATN2_HUMAN | 1,327,290 | 165,302 | 12% | O00339 | MATN2_HUMAN | MATN2_HUMAN | Matrilin-2 | MATN2 UNQ193/PRO219 |
| sp\|P62263\|RS14_HUMAN | 743,435 | 715,516 | 1,098,005 | 1,026,301 | sp\|P62263\|RS14_HUMAN | 689,784 | 260,648 | 38% | P62263 | RS14_HUMAN | RS14_HUMAN | 40S ribosomal protein S14 (Small ribosomal subunit protein uS11) | RPS14 PRO2640 |
| sp\|P21291\|CSRP1_HUMAN | 510,400 | 597,162 | 260,901 | 197,097 | sp\|P21291\|CSRP1_HUMAN | 373,003 | 181,838 | 49% | P21291 | CSRP1_HUMAN | CSRP1_HUMAN | Cysteine and glycine-rich protein 1 (Cysteine-rich protein 1) (CRP) (CRP1) (Epididymis luminal protein 141) (HEL-141) | CSRP1 CSRP CYRP |
| sp\|P09382\|LEG1_HUMAN | 2,283,236 | 2,380,396 | 1,177,374 | 1,245,317 | sp\|P09382\|LEG1_HUMAN | 1,339,603 | 727,184 | 54% | P09382 | LEG1_HUMAN | LEG1_HUMAN | Galectin-1 (Gal-1) (14 kDa laminin-binding protein) (HLBP14) (14 kDa lectin) (Beta-galactoside-binding lectin L-14-I) (Galaptin) (HBL) (HPL) (Lactose-binding lectin 1) (Lectin galactoside-binding soluble 1) (Putative MAPK-activating protein PM12) (S-Lac lectin 1) | LGALS1 |
| sp\|P09622\|DLDH_HUMAN | 60,280 | 56,978 | 98,352 | 79,254 | sp\|P09622\|DLDH_HUMAN | 104,643 | 57,293 | 55% | P09622 | DLDH_HUMAN | DLDH_HUMAN | Dihydrolipoyl dehydrogenase, mitochondrial (EC 1.8.1.4) (Dihydrolipoamide dehydrogenase) (Glycine cleavage system L protein) | DLD GCSL LAD PHE3 |
| sp\|P27348\|1433T_HUMAN | 984,736 | 1,683,820 | 1,084,538 | 1,364,977 | sp\|P27348\|1433T_HUMAN | 1,383,678 | 316,926 | 23% | P27348 | 1433T_HUMAN | 1433T_HUMAN | 14-3-3 protein theta (14-3-3 protein T-cell) (14-3-3 protein tau) (Protein HS1) | YWHAQ |
| sp\|Q09666\|AHNK_HUMAN | 197,609 | 229,684 | 196,284 | 145,077 | sp\|Q09666\|AHNK_HUMAN | 195,963 | 52,713 | 27% | Q09666 | AHNK_HUMAN | AHNK_HUMAN | Neuroblast differentiation-associated protein AHNAK (Desmoyokin) | AHNAK PM227 |
| sp\|P62081\|RS7_HUMAN | 417,299 | 354,537 | 508,370 | 308,771 | sp\|P62081\|RS7_HUMAN | 302,444 | 146,632 | 48% | P62081 | RS7_HUMAN | RS7_HUMAN | 40S ribosomal protein S7 (Small ribosomal subunit protein eS7) | RPS7 |
| sp\|Q3ZCQ8\|TIM50_HUMAN | 163,348 | 185,183 | 89,921 | 146,588 | sp\|Q3ZCQ8\|TIM50_HUMAN | 114,051 | 46,616 | 41% | Q3ZCQ8 | TIM50_HUMAN | TIM50_HUMAN | Mitochondrial import inner membrane translocase subunit TIM50 | TIMM50 TIM50 PRO1512 |
| sp\|P07093\|GDN_HUMAN | 433,135 | 573,732 | 374,581 | 504,067 | sp\|P07093\|GDN_HUMAN | 492,200 | 116,693 | 24% | P07093 | GDN_HUMAN | GDN_HUMAN | Glia-derived nexin (GDN) (Peptidase inhibitor 7) (PI-7) (Protease nexin 1) (PN-1) (Protease nexin I) (Serpin E2) | SERPINE2 PI7 PN1 |
| sp\|P00734\|THRB_HUMAN | 204,377 | 105,400 | 135,842 | 140,281 | sp\|P00734\|THRB_HUMAN | 2,374,331 | 2,465,373 | 104% | P00734 | THRB_HUMAN | THRB_HUMAN | Prothrombin (EC 3.4.21.5) (Coagulation factor II) [Cleaved into: Activation peptide fragment 1; Activation peptide fragment 2; Thrombin light chain; Thrombin heavy chain] | F2 |

| | | | | | | | | | | | | | |
|---|---|---|---|---|---|---|---|---|---|---|---|---|---|
| sp\|Q9P0V3\|SH3B4_HUMAN | 430,240 | 486,907 | 308,129 | 223,785 | sp\|Q9P0V3\|SH3B4_HUMAN | 364,222 | 85,244 | 23% | Q9P0V3 | SH3B4_HUMAN | SH3B4_HUMAN | SH3 domain-binding protein 4 (EH-binding protein 10) (Transferrin receptor-trafficking protein) | SH3BP4 BOG25 EHB10 TTP |
| sp\|O15143\|ARC1B_HUMAN | 477,235 | 584,545 | 476,954 | 590,117 | sp\|O15143\|ARC1B_HUMAN | 447,428 | 108,019 | 24% | O15143 | ARC1B_HUMAN | ARC1B_HUMAN | Actin-related protein 2/3 complex subunit 1B (Arp2/3 complex 41 kDa subunit) (p41-ARC) | ARPC1B ARC41 |
| sp\|P63241\|IF5A1_HUMAN | 246,000 | 322,429 | 356,455 | 364,281 | sp\|P63241\|IF5A1_HUMAN | 291,676 | 50,511 | 17% | P63241 | IF5A1_HUMAN | IF5A1_HUMAN | Eukaryotic translation initiation factor 5A-1 (eIF-5A-1) (eIF-5A1) (Eukaryotic initiation factor 5A isoform 1) (eIF-5A) (Rev-binding factor) (eIF-4D) | EIF5A |
| sp\|P32969\|RL9_HUMAN | 886,120 | 918,861 | 434,780 | 628,438 | sp\|P32969\|RL9_HUMAN | 552,330 | 238,185 | 43% | P32969 | RL9_HUMAN | RL9_HUMAN | 60S ribosomal protein L9 (Large ribosomal subunit protein uL6) | RPL9 OK/SW-cl.103; RPL9P7; RPL9P8; RPL9P9 |
| sp\|P54652\|HSP72_HUMAN | 838,845 | 1,180,839 | 424,363 | 765,087 | sp\|P54652\|HSP72_HUMAN | 653,319 | 269,308 | 41% | P54652 | HSP72_HUMAN | HSP72_HUMAN | Heat shock-related 70 kDa protein 2 (Heat shock 70 kDa protein 2) | HSPA2 |
| sp\|P61160\|ARP2_HUMAN | 910,088 | 1,216,422 | 626,140 | 727,468 | sp\|P61160\|ARP2_HUMAN | 774,097 | 215,024 | 28% | P61160 | ARP2_HUMAN | ARP2_HUMAN | Actin-related protein 2 (Actin-like protein 2) | ACTR2 ARP2 |
| sp\|P18621\|RL17_HUMAN | 856,481 | 831,626 | 828,740 | 604,203 | sp\|P18621\|RL17_HUMAN | 522,233 | 289,443 | 55% | P18621 | RL17_HUMAN | RL17_HUMAN | 60S ribosomal protein L17 (60S ribosomal protein L23) (Large ribosomal subunit protein uL22) (PD-1) | RPL17 |
| sp\|P62269\|RS18_HUMAN | 1,615,859 | 1,617,179 | 2,148,387 | 2,001,638 | sp\|P62269\|RS18_HUMAN | 1,332,141 | 583,260 | 44% | P62269 | RS18_HUMAN | RS18_HUMAN | 40S ribosomal protein S18 (Ke-3) (Ke3) (Small ribosomal subunit protein uS13) | RPS18 D6S218E |
| sp\|P08237\|PFKAM_HUMAN | 220,471 | 228,072 | 250,235 | 249,040 | sp\|P08237\|PFKAM_HUMAN | 197,422 | 44,722 | 23% | P08237 | PFKAM_HUMAN | PFKAM_HUMAN | ATP-dependent 6-phosphofructokinase, muscle type (ATP-PFK) (PFK-M) (EC 2.7.1.11) (6-phosphofructokinase type A) (Phosphofructo-1-kinase isozyme A) (PFK-A) (Phosphohexokinase) | PFKM PFKX |
| sp\|Q99497\|PARK7_HUMAN | 575,468 | 768,685 | 272,339 | 356,313 | sp\|Q99497\|PARK7_HUMAN | 431,738 | 220,261 | 51% | Q99497 | PARK7_HUMAN | PARK7_HUMAN | Parkinson disease protein 7 (Maillard deglycase) (Oncogene DJ1) (Parkinsonism-associated deglycase) (Protein DJ-1) (DJ-1) (Protein/nucleic acid deglycase DJ-1) (EC 3.1.2.-) (EC 3.5.1.-) (EC 3.5.1.124) | PARK7 |
| sp\|P36871\|PGM1_HUMAN | 283,102 | 495,662 | 425,172 | 313,763 | sp\|P36871\|PGM1_HUMAN | 348,281 | 86,368 | 25% | P36871 | PGM1_HUMAN | PGM1_HUMAN | Phosphoglucomutase-1 (PGM 1) (EC 5.4.2.2) (Glucose phosphomutase 1) | PGM1 |
| sp\|Q14247\|SRC8_HUMAN | 132,217 | 161,453 | 243,192 | 306,161 | sp\|Q14247\|SRC8_HUMAN | 191,886 | 75,973 | 40% | Q14247 | SRC8_HUMAN | SRC8_HUMAN | Src substrate cortactin (Amplaxin) (Oncogene EMS1) | CTTN EMS1 |
| sp\|P62280\|RS11_HUMAN | 378,964 | 319,210 | 305,234 | 298,293 | sp\|P62280\|RS11_HUMAN | 219,582 | 117,340 | 53% | P62280 | RS11_HUMAN | RS11_HUMAN | 40S ribosomal protein S11 (Small ribosomal subunit protein uS17) | RPS11 |
| sp\|P01903\|DRA_HUMAN | 13,250,191 | 15,351,909 | 10,627,093 | 10,967,980 | sp\|P01903\|DRA_HUMAN | 11,159,494 | 2,411,611 | 22% | P01903 | DRA_HUMAN | DRA_HUMAN | HLA class II histocompatibility antigen, DR alpha chain (MHC class II antigen DRA) | HLA-DRA HLA-DRA1 |
| sp\|Q9Y305\|ACOT9_HUMAN | 56,246 | 54,587 | 86,173 | 131,739 | sp\|Q9Y305\|ACOT9_HUMAN | 73,287 | 28,030 | 38% | Q9Y305 | ACOT9_HUMAN | ACOT9_HUMAN | Acyl-coenzyme A thioesterase 9, mitochondrial (Acyl-CoA thioesterase 9) (EC 3.1.2.-) (Acyl-CoA thioester hydrolase 9) | ACOT9 CGI-16 |
| sp\|Q14019\|COTL1_HUMAN | 1,164,493 | 1,398,903 | 828,261 | 1,052,077 | sp\|Q14019\|COTL1_HUMAN | 1,090,140 | 244,048 | 22% | Q14019 | COTL1_HUMAN | COTL1_HUMAN | Coactosin-like protein | COTL1 CLP |
| sp\|P19338\|NUCL_HUMAN | 867,309 | 752,539 | 742,344 | 802,473 | sp\|P19338\|NUCL_HUMAN | 551,052 | 265,210 | 48% | P19338 | NUCL_HUMAN | NUCL_HUMAN | Nucleolin (Protein C23) | NCL |
| sp\|Q9P2J5\|SYLC_HUMAN | 214,714 | 184,894 | 283,015 | 242,527 | sp\|Q9P2J5\|SYLC_HUMAN | 236,264 | 79,097 | 33% | Q9P2J5 | SYLC_HUMAN | SYLC_HUMAN | Leucine--tRNA ligase, cytoplasmic (EC 6.1.1.4) (Leucyl-tRNA synthetase) (LeuRS) | LARS1 KIAA1352 LARS |
| sp\|P20073\|ANXA7_HUMAN | 813,267 | 592,573 | 1,303,774 | 675,514 | sp\|P20073\|ANXA7_HUMAN | 718,592 | 275,363 | 38% | P20073 | ANXA7_HUMAN | ANXA7_HUMAN | Annexin A7 (Annexin VII) (Annexin-7) (Synexin) | ANXA7 ANX7 SNX OK/SW-cl.95 |

| | | | | | | | | | | | | | |
|---|---|---|---|---|---|---|---|---|---|---|---|---|---|
| sp\|Q9Y2J2\|E41L3_HUMAN | 512,764 | 583,639 | 314,060 | 423,698 | sp\|Q9Y2J2\|E41L3_HUMAN | 448,806 | 97,741 | 22% | Q9Y2J2 | E41L3_HUMAN | E41L3_HUMAN | Band 4.1-like protein 3 (4.1B) (Differentially expressed in adenocarcinoma of the lung protein 1) (DAL-1) (Erythrocyte membrane protein band 4.1-like 3) [Cleaved into: Band 4.1-like protein 3, N-terminally processed] | EPB41L3 DAL1 KIAA0987 |
| sp\|O14672\|ADA10_HUMAN | 1,150,726 | 1,264,360 | 1,512,991 | 1,458,052 | sp\|O14672\|ADA10_HUMAN | 1,062,198 | 352,183 | 33% | O14672 | ADA10_HUMAN | ADA10_HUMAN | Disintegrin and metalloproteinase domain-containing protein 10 (ADAM 10) (EC 3.4.24.81) (CDw156) (Kuzbanian protein homolog) (Mammalian disintegrin-metalloprotease) (CD antigen CD156c) | ADAM10 KUZ MADM |
| sp\|Q14108\|SCRB2_HUMAN | 1,050,600 | 923,218 | 1,250,441 | 986,762 | sp\|Q14108\|SCRB2_HUMAN | 809,534 | 284,642 | 35% | Q14108 | SCRB2_HUMAN | SCRB2_HUMAN | Lysosome membrane protein 2 (85 kDa lysosomal membrane sialoglycoprotein) (LGP85) (CD36 antigen-like 2) (Lysosome membrane protein II) (LIMP II) (Scavenger receptor class B member 2) (CD antigen CD36) | SCARB2 CD36L2 LIMP2 LIMPII |
| sp\|O00560\|SDCB1_HUMAN | 5,469,747 | 6,683,643 | 5,688,336 | 6,148,761 | sp\|O00560\|SDCB1_HUMAN | 5,093,653 | 1,119,533 | 22% | O00560 | SDCB1_HUMAN | SDCB1_HUMAN | Syntenin-1 (Melanoma differentiation-associated protein 9) (MDA-9) (Pro-TGF-alpha cytoplasmic domain-interacting protein 18) (TACIP18) (Scaffold protein Pbp1) (Syndecan-binding protein 1) | SDCBP MDA9 SYCL |
| sp\|Q9UNF0\|PACN2_HUMAN | 556,593 | 721,045 | 730,843 | 872,179 | sp\|Q9UNF0\|PACN2_HUMAN | 639,964 | 133,581 | 21% | Q9UNF0 | PACN2_HUMAN | PACN2_HUMAN | Protein kinase C and casein kinase substrate in neurons protein 2 (Syndapin-2) (Syndapin-II) (SdpII) | PACSIN2 |
| sp\|Q9BR76\|COR1B_HUMAN | 402,359 | 433,919 | 574,169 | 714,045 | sp\|Q9BR76\|COR1B_HUMAN | 461,271 | 139,924 | 30% | Q9BR76 | COR1B_HUMAN | COR1B_HUMAN | Coronin-1B (Coronin-2) | CORO1B |
| sp\|Q9UJZ1\|STML2_HUMAN | 69,424 | 80,429 | 69,361 | 90,747 | sp\|Q9UJZ1\|STML2_HUMAN | 93,128 | 21,198 | 23% | Q9UJZ1 | STML2_HUMAN | STML2_HUMAN | Stomatin-like protein 2, mitochondrial (SLP-2) (EPB72-like protein 2) (Paraprotein target 7) (Paratarg-7) | STOML2 SLP2 HSPC108 |
| sp\|Q8NBX0\|SCPDL_HUMAN | 243,367 | 222,269 | 360,481 | 372,130 | sp\|Q8NBX0\|SCPDL_HUMAN | 263,261 | 89,686 | 34% | Q8NBX0 | SCPDL_HUMAN | SCPDL_HUMAN | Saccharopine dehydrogenase-like oxidoreductase (EC 1.-.-.-) | SCCPDH CGI-49 |
| sp\|Q9NZM1\|MYOF_HUMAN | 423,620 | 474,687 | 383,372 | 365,980 | sp\|Q9NZM1\|MYOF_HUMAN | 347,511 | 82,369 | 24% | Q9NZM1 | MYOF_HUMAN | MYOF_HUMAN | Myoferlin (Fer-1-like protein 3) | MYOF FER1L3 KIAA1207 |
| sp\|Q86UX2\|ITIH5_HUMAN | 1,837,635 | 1,246,949 | 2,963,303 | 2,231,678 | sp\|Q86UX2\|ITIH5_HUMAN | 1,796,565 | 679,649 | 38% | Q86UX2 | ITIH5_HUMAN | ITIH5_HUMAN | Inter-alpha-trypsin inhibitor heavy chain H5 (ITI heavy chain H5) (ITI-HC5) (Inter-alpha-inhibitor heavy chain 5) | ITIH5 KIAA1953 PP14776 UNQ311/PRO354 |
| sp\|Q86VP6\|CAND1_HUMAN | 354,929 | 429,508 | 158,063 | 259,994 | sp\|Q86VP6\|CAND1_HUMAN | 296,113 | 100,814 | 34% | Q86VP6 | CAND1_HUMAN | CAND1_HUMAN | Cullin-associated NEDD8-dissociated protein 1 (Cullin-associated and neddylation-dissociated protein 1) (TBP-interacting protein of 120 kDa A) (TBP-interacting protein 120A) (p120 CAND1) | CAND1 KIAA0829 TIP120 TIP120A |
| sp\|P40429\|RL13A_HUMAN | 1,730,871 | 1,435,368 | 1,548,644 | 1,339,292 | sp\|P40429\|RL13A_HUMAN | 1,111,688 | 443,713 | 40% | P40429 | RL13A_HUMAN | RL13A_HUMAN | 60S ribosomal protein L13a (23 kDa highly basic protein) (Large ribosomal subunit protein uL13) | RPL13A |

| | | | | | | | | | | | | | |
|---|---|---|---|---|---|---|---|---|---|---|---|---|---|
| sp\|P23381\|SYWC_HUMAN | 187,478 | 185,466 | 105,425 | 159,123 | sp\|P23381\|SYWC_HUMAN | 148,511 | 37,110 | 25% | P23381 | SYWC_HUMAN | SYWC_HUMAN | Tryptophan--tRNA ligase, cytoplasmic (EC 6.1.1.2) (Interferon-induced protein 53) (IFP53) (Tryptophanyl-tRNA synthetase) (TrpRS) (hWRS) [Cleaved into: T1-TrpRS; T2-TrpRS] | WARS1 IFI53 WARS WRS |
| sp\|P62277\|RS13_HUMAN | 735,937 | 867,370 | 451,096 | 508,830 | sp\|P62277\|RS13_HUMAN | 475,778 | 235,760 | 50% | P62277 | RS13_HUMAN | RS13_HUMAN | 40S ribosomal protein S13 (Small ribosomal subunit protein uS15) | RPS13 |
| sp\|P22570\|ADRO_HUMAN | 54,515 | 14,266 | 192,967 | 73,306 | sp\|P22570\|ADRO_HUMAN | 100,727 | 83,098 | 82% | P22570 | ADRO_HUMAN | ADRO_HUMAN | NADPH:adrenodoxin oxidoreductase, mitochondrial (AR) (Adrenodoxin reductase) (EC 1.18.1.6) (Ferredoxin--NADP(+) reductase) (Ferredoxin reductase) | FDXR ADXR |
| sp\|P53621\|COPA_HUMAN | 423,894 | 365,160 | 364,682 | 405,671 | sp\|P53621\|COPA_HUMAN | 301,363 | 114,089 | 38% | P53621 | COPA_HUMAN | COPA_HUMAN | Coatomer subunit alpha (Alpha-coat protein) (Alpha-COP) (HEP-COP) (HEPCOP) [Cleaved into: Xenin (Xenopsin-related peptide); Proxenin] | COPA |
| sp\|Q01650\|LAT1_HUMAN | 3,330,661 | 4,045,093 | 2,150,778 | 2,530,802 | sp\|Q01650\|LAT1_HUMAN | 2,623,642 | 790,872 | 30% | Q01650 | LAT1_HUMAN | LAT1_HUMAN | Large neutral amino acids transporter small subunit 1 (4F2 light chain) (4F2 LC) (4F2LC) (CD98 light chain) (Integral membrane protein E16) (E16) (L-type amino acid transporter 1) (hLAT1) (Solute carrier family 7 member 5) (y+ system cationic amino acid transporter) | SLC7A5 CD98LC LAT1 MPE16 |
| sp\|P51114\|FXR1_HUMAN | 228,051 | 231,893 | 192,654 | 359,552 | sp\|P51114\|FXR1_HUMAN | 207,039 | 71,938 | 35% | P51114 | FXR1_HUMAN | FXR1_HUMAN | Fragile X mental retardation syndrome-related protein 1 (hFXR1p) | FXR1 |
| sp\|P62906\|RL10A_HUMAN | 900,053 | 1,121,160 | 537,370 | 511,023 | sp\|P62906\|RL10A_HUMAN | 561,320 | 301,292 | 54% | P62906 | RL10A_HUMAN | RL10A_HUMAN | 60S ribosomal protein L10a (CSA-19) (Large ribosomal subunit protein uL1) (Neural precursor cell expressed developmentally down-regulated protein 6) (NEDD-6) | RPL10A NEDD6 |
| sp\|Q9NTK5\|OLA1_HUMAN | 412,131 | 457,408 | 280,290 | 368,477 | sp\|Q9NTK5\|OLA1_HUMAN | 358,924 | 78,771 | 22% | Q9NTK5 | OLA1_HUMAN | OLA1_HUMAN | Obg-like ATPase 1 (DNA damage-regulated overexpressed in cancer 45) (DOC45) (GTP-binding protein 9) | OLA1 GTPBP9 PRO2455 PTD004 |
| sp\|P24539\|AT5F1_HUMAN | 48,050 | 47,886 | 47,832 | 55,216 | sp\|P24539\|AT5F1_HUMAN | 49,329 | 17,187 | 35% | P24539 | AT5F1_HUMAN | AT5F1_HUMAN | ATP synthase F(0) complex subunit B1, mitochondrial (ATP synthase peripheral stalk-membrane subunit b) (ATP synthase proton-transporting mitochondrial F(0) complex subunit B1) (ATP synthase subunit b) (ATPase subunit b) | ATP5PB ATP5F1 |
| sp\|Q9UNM6\|PSD13_HUMAN | 292,904 | 300,180 | 267,602 | 174,554 | sp\|Q9UNM6\|PSD13_HUMAN | 219,105 | 63,884 | 29% | Q9UNM6 | PSD13_HUMAN | PSD13_HUMAN | 26S proteasome non-ATPase regulatory subunit 13 (26S proteasome regulatory subunit RPN9) (26S proteasome regulatory subunit S11) (26S proteasome regulatory subunit p40.5) | PSMD13 |
| sp\|P62805\|H4_HUMAN | 37,032,038 | 15,106,609 | 45,660,190 | 12,220,912 | sp\|P62805\|H4_HUMAN | 20,173,760 | 14,258,264 | 71% | P62805 | H4_HUMAN | H4_HUMAN | Histone H4 | H4C1 H4/A H4FA HIST1H4A; H4C2 H4/I H4FI HIST1H4B; H4C3 H4/G H4FG HIST1H4C; H4C4 H4/B H4FB HIST1H4D; H4C5 H4/J H4FJ HIST1H4E; H4C6 H4/C H4FC HIST1H4F; H4C8 H4/H H4FH HIST1H4H; H4C9 H4/M H4FM HIST1H4I; H4C11 H4/E H4FE HIST1H4J; H4C12 H4/D H4FD HIST1H4K; H4C13 H4/K H4FK HIST1H4L; H4C14 H4/N H4F2 H4FN HIST2H4 HIST2H4A; H4C15 H4/O H4FO HIST2H4B; H4-16 HIST4H4 |

| | | | | | | | | | | | | | |
|---|---|---|---|---|---|---|---|---|---|---|---|---|---|
| sp\|P08559\|ODPA_HUMAN | 100,500 | 118,702 | 233,325 | 227,870 | sp\|P08559\|ODPA_HUMAN | 150,219 | 58,342 | 39% | P08559 | ODPA_HUMAN | ODPA_HUMAN | Pyruvate dehydrogenase E1 component subunit alpha, somatic form, mitochondrial (EC 1.2.4.1) (PDHE1-A type I) | PDHA1 PHE1A |
| sp\|P30048\|PRDX3_HUMAN | 430,751 | 509,453 | 353,617 | 351,041 | sp\|P30048\|PRDX3_HUMAN | 379,808 | 100,313 | 26% | P30048 | PRDX3_HUMAN | PRDX3_HUMAN | Thioredoxin-dependent peroxide reductase, mitochondrial (EC 1.11.1.24) (Antioxidant protein 1) (AOP-1) (HBC189) (Peroxiredoxin III) (Prx-III) (Peroxiredoxin-3) (Protein MER5 homolog) (Thioredoxin-dependent peroxiredoxin 3) | PRDX3 AOP1 |
| sp\|P46782\|RS5_HUMAN | 872,747 | 932,710 | 1,023,557 | 1,080,096 | sp\|P46782\|RS5_HUMAN | 748,191 | 276,364 | 37% | P46782 | RS5_HUMAN | RS5_HUMAN | 40S ribosomal protein S5 (Small ribosomal subunit protein uS7) [Cleaved into: 40S ribosomal protein S5, N-terminally processed] | RPS5 |
| sp\|P61225\|RAP2B_HUMAN | 673,129 | 853,637 | 91,518 | 472,996 | sp\|P61225\|RAP2B_HUMAN | 478,789 | 241,584 | 50% | P61225 | RAP2B_HUMAN | RAP2B_HUMAN | Ras-related protein Rap-2b (EC 3.6.5.2) | RAP2B |
| sp\|P20618\|PSB1_HUMAN | 605,331 | 650,463 | 709,553 | 699,622 | sp\|P20618\|PSB1_HUMAN | 579,482 | 100,737 | 17% | P20618 | PSB1_HUMAN | PSB1_HUMAN | Proteasome subunit beta type-1 (Macropain subunit C5) (Multicatalytic endopeptidase complex subunit C5) (Proteasome component C5) (Proteasome gamma chain) | PSMB1 PSC5 |
| sp\|Q8IWB7\|WDFY1_HUMAN | 193,687 | 235,150 | 100,976 | 73,306 | sp\|Q8IWB7\|WDFY1_HUMAN | 141,410 | 56,900 | 40% | Q8IWB7 | WDFY1_HUMAN | WDFY1_HUMAN | WD repeat and FYVE domain-containing protein 1 (FYVE domain-containing protein localized to endosomes 1) (FENS-1) (Phosphoinositide-binding protein 1) (WD40- and FYVE domain-containing protein 1) (Zinc finger FYVE domain-containing protein 17) | WDFY1 FENS1 KIAA1435 WDF1 ZFYVE17 |
| sp\|P17858\|PFKAL_HUMAN | 267,247 | 305,595 | 243,109 | 275,575 | sp\|P17858\|PFKAL_HUMAN | 245,348 | 43,612 | 18% | P17858 | PFKAL_HUMAN | PFKAL_HUMAN | ATP-dependent 6-phosphofructokinase, liver type (ATP-PFK) (PFK-L) (EC 2.7.1.11) (6-phosphofructokinase type B) (Phosphofructo-1-kinase isozyme B) (PFK-B) (Phosphohexokinase) | PFKL |
| sp\|Q13347\|EIF3I_HUMAN | 151,489 | 151,424 | 124,216 | 171,886 | sp\|Q13347\|EIF3I_HUMAN | 142,171 | 27,380 | 19% | Q13347 | EIF3I_HUMAN | EIF3I_HUMAN | Eukaryotic translation initiation factor 3 subunit I (eIF3i) (Eukaryotic translation initiation factor 3 subunit 2) (TGF-beta receptor-interacting protein 1) (TRIP-1) (eIF-3-beta) (eIF3 p36) | EIF3I EIF3S2 TRIP1 |
| sp\|P31040\|SDHA_HUMAN | 35,908 | 31,974 | 45,200 | 69,992 | sp\|P31040\|SDHA_HUMAN | 36,176 | 16,450 | 45% | P31040 | SDHA_HUMAN | SDHA_HUMAN | Succinate dehydrogenase [ubiquinone] flavoprotein subunit, mitochondrial (EC 1.3.5.1) (Flavoprotein subunit of complex II) (Fp) | SDHA SDH2 SDHF |
| sp\|P37837\|TALDO_HUMAN | 276,085 | 366,233 | 213,117 | 244,391 | sp\|P37837\|TALDO_HUMAN | 256,125 | 65,119 | 25% | P37837 | TALDO_HUMAN | TALDO_HUMAN | Transaldolase (EC 2.2.1.2) | TALDO1 TAL TALDO TALDOR |
| sp\|P24941\|CDK2_HUMAN | 240,112 | 261,893 | 113,710 | 311,268 | sp\|P24941\|CDK2_HUMAN | 183,301 | 78,155 | 43% | P24941 | CDK2_HUMAN | CDK2_HUMAN | Cyclin-dependent kinase 2 (EC 2.7.11.22) (Cell division protein kinase 2) (p33 protein kinase) | CDK2 CDKN2 |
| sp\|P22695\|QCR2_HUMAN | 46,411 | 54,834 | 71,195 | 36,161 | sp\|P22695\|QCR2_HUMAN | 59,785 | 26,846 | 45% | P22695 | QCR2_HUMAN | QCR2_HUMAN | Cytochrome b-c1 complex subunit 2, mitochondrial (Complex III subunit 2) (Core protein II) (Ubiquinol-cytochrome-c reductase complex core protein 2) | UQCRC2 |
| sp\|P62917\|RL8_HUMAN | 1,255,687 | 1,220,211 | 2,613,311 | 2,273,789 | sp\|P62917\|RL8_HUMAN | 1,288,315 | 760,568 | 59% | P62917 | RL8_HUMAN | RL8_HUMAN | 60S ribosomal protein L8 (Large ribosomal subunit protein uL2) | RPL8 |

| | | | | | | | | | | | | | |
|---|---|---|---|---|---|---|---|---|---|---|---|---|---|
| sp\|P06753\|TPM3_HUMAN | 228,721 | 284,508 | 486,319 | 722,643 | sp\|P06753\|TPM3_HUMAN | 448,422 | 168,182 | 38% | P06753 | TPM3_HUMAN | TPM3_HUMAN | Tropomyosin alpha-3 chain (Gamma-tropomyosin) (Tropomyosin-3) (Tropomyosin-5) (hTM5) | TPM3 |
| sp\|P39019\|RS19_HUMAN | 256,869 | 435,673 | 118,025 | 246,427 | sp\|P39019\|RS19_HUMAN | 278,342 | 126,868 | 46% | P39019 | RS19_HUMAN | RS19_HUMAN | 40S ribosomal protein S19 (Small ribosomal subunit protein eS19) | RPS19 |
| sp\|P11177\|ODPB_HUMAN | 59,107 | 53,611 | 50,871 | 21,873 | sp\|P11177\|ODPB_HUMAN | 60,899 | 30,257 | 50% | P11177 | ODPB_HUMAN | ODPB_HUMAN | Pyruvate dehydrogenase E1 component subunit beta, mitochondrial (PDHE1-B) (EC 1.2.4.1) | PDHB PHE1B |
| sp\|P55884\|EIF3B_HUMAN | 390,893 | 438,763 | 191,254 | 427,827 | sp\|P55884\|EIF3B_HUMAN | 343,504 | 123,895 | 36% | P55884 | EIF3B_HUMAN | EIF3B_HUMAN | Eukaryotic translation initiation factor 3 subunit B (eIF3b) (Eukaryotic translation initiation factor 3 subunit 9) (Prt1 homolog) (hPrt1) (eIF-3-eta) (eIF3 p110) (eIF3 p116) | EIF3B EIF3S9 |
| sp\|P04844\|RPN2_HUMAN | 172,854 | 169,322 | 182,385 | 274,190 | sp\|P04844\|RPN2_HUMAN | 190,038 | 55,146 | 29% | P04844 | RPN2_HUMAN | RPN2_HUMAN | Dolichyl-diphosphooligosaccharide--protein glycosyltransferase subunit 2 (Dolichyl-diphosphooligosaccharide--protein glycosyltransferase 63 kDa subunit) (RIBIIR) (Ribophorin II) (RPN-II) (Ribophorin-2) | RPN2 |
| sp\|O60716\|CTND1_HUMAN | 261,261 | 388,576 | 249,389 | 364,534 | sp\|O60716\|CTND1_HUMAN | 387,269 | 126,620 | 33% | O60716 | CTND1_HUMAN | CTND1_HUMAN | Catenin delta-1 (Cadherin-associated Src substrate) (CAS) (p120 catenin) (p120(ctn)) (p120(cas)) | CTNND1 KIAA0384 |
| sp\|P09417\|DHPR_HUMAN | 405,727 | 473,556 | 110,847 | 340,904 | sp\|P09417\|DHPR_HUMAN | 274,020 | 139,842 | 51% | P09417 | DHPR_HUMAN | DHPR_HUMAN | Dihydropteridine reductase (EC 1.5.1.34) (HDHPR) (Quinoid dihydropteridine reductase) (Short chain dehydrogenase/reductase family 33C member 1) | QDPR DHPR SDR33C1 |
| sp\|Q9NQC3\|RTN4_HUMAN | 363,900 | 323,947 | 534,333 | 397,680 | sp\|Q9NQC3\|RTN4_HUMAN | 403,412 | 112,509 | 28% | Q9NQC3 | RTN4_HUMAN | RTN4_HUMAN | Reticulon-4 (Foocen) (Neurite outgrowth inhibitor) (Nogo protein) (Neuroendocrine-specific protein) (NSP) (Neuroendocrine-specific protein C homolog) (RTN-x) (Reticulon-5) | RTN4 KIAA0886 NOGO My043 SP1507 |
| sp\|Q13283\|G3BP1_HUMAN | 33,988 | 77,561 | 65,970 | 116,307 | sp\|Q13283\|G3BP1_HUMAN | 85,472 | 28,179 | 33% | Q13283 | G3BP1_HUMAN | G3BP1_HUMAN | Ras GTPase-activating protein-binding protein 1 (G3BP-1) (EC 3.6.4.12) (EC 3.6.4.13) (ATP-dependent DNA helicase VIII) (hDH VIII) (GAP SH3 domain-binding protein 1) | G3BP1 G3BP |
| sp\|P01111\|RASN_HUMAN | 711,805 | 896,085 | 696,567 | 729,405 | sp\|P01111\|RASN_HUMAN | 711,722 | 85,148 | 12% | P01111 | RASN_HUMAN | RASN_HUMAN | GTPase NRas (EC 3.6.5.2) (Transforming protein N-Ras) | NRAS HRAS1 |
| sp\|P14324\|FPPS_HUMAN | 197,109 | 235,327 | 173,102 | 223,773 | sp\|P14324\|FPPS_HUMAN | 202,663 | 33,244 | 16% | P14324 | FPPS_HUMAN | FPPS_HUMAN | Farnesyl pyrophosphate synthase (FPP synthase) (FPS) (EC 2.5.1.10) ((2E,6E)-farnesyl diphosphate synthase) (Dimethylallyltranstransferase) (EC 2.5.1.1) (Farnesyl diphosphate synthase) (Geranyltranstransferase) | FDPS FPS KIAA1293 |
| sp\|Q96I99\|SUCB2_HUMAN | 81,910 | 83,240 | 47,498 | 113,814 | sp\|Q96I99\|SUCB2_HUMAN | 104,335 | 54,740 | 52% | Q96I99 | SUCB2_HUMAN | SUCB2_HUMAN | Succinate--CoA ligase [GDP-forming] subunit beta, mitochondrial (EC 6.2.1.4) (GTP-specific succinyl-CoA synthetase subunit beta) (G-SCS) (GTPSCS) (Succinyl-CoA synthetase beta-G chain) (SCS-betaG) | SUCLG2 |

| | | | | | | | | | | | | | |
|---|---|---|---|---|---|---|---|---|---|---|---|---|---|
| sp\|Q9UBM7\|DHCR7_HUMAN | 56,782 | 83,579 | 142,359 | 132,017 | sp\|Q9UBM7\|DHCR7_HUMAN | 78,615 | 40,533 | 52% | Q9UBM7 | DHCR7_HUMAN | DHCR7_HUMAN | 7-dehydrocholesterol reductase (7-DHC reductase) (EC 1.3.1.21) (Delta7-sterol reductase) (Sterol Delta(7)-reductase) (Sterol reductase SR-2) | DHCR7 D7SR |
| sp\|P19105\|ML12A_HUMAN | 499,398 | 621,887 | 855,042 | 974,118 | sp\|P19105\|ML12A_HUMAN | 570,401 | 235,858 | 41% | P19105 | ML12A_HUMAN | ML12A_HUMAN | Myosin regulatory light chain 12A (Epididymis secretory protein Li 24) (HEL-S-24) (MLC-2B) (Myosin RLC) (Myosin regulatory light chain 2, nonsarcomeric) (Myosin regulatory light chain MRLC3) | MYL12A MLCB MRLC3 RLC |
| sp\|P25787\|PSA2_HUMAN | 299,132 | 337,258 | 183,681 | 228,884 | sp\|P25787\|PSA2_HUMAN | 245,915 | 51,655 | 21% | P25787 | PSA2_HUMAN | PSA2_HUMAN | Proteasome subunit alpha type-2 (Macropain subunit C3) (Multicatalytic endopeptidase complex subunit C3) (Proteasome component C3) | PSMA2 HC3 PSC3 |
| sp\|O14818\|PSA7_HUMAN | 712,213 | 794,938 | 1,082,898 | 1,224,787 | sp\|O14818\|PSA7_HUMAN | 926,813 | 194,844 | 21% | O14818 | PSA7_HUMAN | PSA7_HUMAN | Proteasome subunit alpha type-7 (Proteasome subunit RC6-1) (Proteasome subunit XAPC7) | PSMA7 HSPC |
| sp\|Q15286\|RAB35_HUMAN | 890,229 | 892,831 | 964,421 | 1,126,271 | sp\|Q15286\|RAB35_HUMAN | 901,186 | 172,512 | 19% | Q15286 | RAB35_HUMAN | RAB35_HUMAN | Ras-related protein Rab-35 (GTP-binding protein RAY) (Ras-related protein Rab-1C) | RAB35 RAB1C RAY |
| sp\|O95819\|M4K4_HUMAN | 564,299 | 585,842 | 600,834 | 1,028,422 | sp\|O95819\|M4K4_HUMAN | 704,937 | 169,659 | 24% | O95819 | M4K4_HUMAN | M4K4_HUMAN | Mitogen-activated protein kinase kinase kinase kinase 4 (EC 2.7.11.1) (HPK/GCK-like kinase HGK) (MAPK/ERK kinase kinase kinase 4) (MEK kinase kinase 4) (MEKKK 4) (Nck-interacting kinase) | MAP4K4 HGK KIAA0687 NIK |
| sp\|P12259\|FA5_HUMAN | 101,218 | 97,651 | 61,925 | 149,178 | sp\|P12259\|FA5_HUMAN | 927,456 | 945,045 | 102% | P12259 | FA5_HUMAN | FA5_HUMAN | Coagulation factor V (Activated protein C cofactor) (Proaccelerin, labile factor) [Cleaved into: Coagulation factor V heavy chain; Coagulation factor V light chain] | F5 |
| sp\|P30050\|RL12_HUMAN | 1,038,768 | 1,313,317 | 581,709 | 810,091 | sp\|P30050\|RL12_HUMAN | 696,547 | 350,813 | 50% | P30050 | RL12_HUMAN | RL12_HUMAN | 60S ribosomal protein L12 (Large ribosomal subunit protein uL11) | RPL12 |
| sp\|P62424\|RL7A_HUMAN | 787,157 | 724,426 | 582,395 | 596,841 | sp\|P62424\|RL7A_HUMAN | 452,582 | 248,703 | 55% | P62424 | RL7A_HUMAN | RL7A_HUMAN | 60S ribosomal protein L7a (Large ribosomal subunit protein eL8) (PLA-X polypeptide) (Surfeit locus protein 3) | RPL7A SURF-3 SURF3 |
| sp\|P04440\|DPB1_HUMAN | 2,100,949 | 2,476,632 | 2,524,191 | 2,411,709 | sp\|P04440\|DPB1_HUMAN | 2,361,469 | 263,076 | 11% | P04440 | DPB1_HUMAN | DPB1_HUMAN | HLA class II histocompatibility antigen, DP beta 1 chain (HLA class II histocompatibility antigen, DP(W4) beta chain) (MHC class II antigen DPB1) | HLA-DPB1 HLA-DP1B |
| sp\|Q08211\|DHX9_HUMAN | 271,363 | 204,631 | 292,698 | 214,386 | sp\|Q08211\|DHX9_HUMAN | 157,106 | 100,020 | 64% | Q08211 | DHX9_HUMAN | DHX9_HUMAN | ATP-dependent RNA helicase A (EC 3.6.4.13) (DEAH box protein 9) (DExH-box helicase 9) (Leukophysin) (LKP) (Nuclear DNA helicase II) (NDH II) (RNA helicase A) | DHX9 DDX9 LKP NDH2 |
| sp\|P68366\|TBA4A_HUMAN | 1,218,518 | 1,307,161 | 359,687 | 929,083 | sp\|P68366\|TBA4A_HUMAN | 852,731 | 303,366 | 36% | P68366 | TBA4A_HUMAN | TBA4A_HUMAN | Tubulin alpha-4A chain (Alpha-tubulin 1) (Testis-specific alpha-tubulin) (Tubulin H2-alpha) (Tubulin alpha-1 chain) | TUBA4A TUBA1 |
| sp\|O14908\|GIPC1_HUMAN | 344,524 | 400,213 | 310,915 | 653,911 | sp\|O14908\|GIPC1_HUMAN | 490,160 | 137,554 | 28% | O14908 | GIPC1_HUMAN | GIPC1_HUMAN | PDZ domain-containing protein GIPC1 (GAIP C-terminus-interacting protein) (RGS-GAIP-interacting protein) (RGS19-interacting protein 1) (Synectin) (Tax interaction protein 2) (TIP-2) | GIPC1 C19orf3 GIPC RGS19IP1 |

| | | | | | | | | | | | | | |
|---|---|---|---|---|---|---|---|---|---|---|---|---|---|
| sp\|P12235\|ADT1_HUMAN | 82,945 | 100,161 | 23,382 | 71,789 | sp\|P12235\|ADT1_HUMAN | 58,475 | 26,459 | 45% | P12235 | ADT1_HUMAN | ADT1_HUMAN | ADP/ATP translocase 1 (ADP,ATP carrier protein 1) (ADP,ATP carrier protein, heart/skeletal muscle isoform T1) (Adenine nucleotide translocator 1) (ANT 1) (Solute carrier family 25 member 4) | SLC25A4 AAC1 ANT1 |
| sp\|P51153\|RAB13_HUMAN | 449,761 | 492,114 | 280,582 | 206,471 | sp\|P51153\|RAB13_HUMAN | 337,838 | 102,168 | 30% | P51153 | RAB13_HUMAN | RAB13_HUMAN | Ras-related protein Rab-13 (Cell growth-inhibiting gene 4 protein) | RAB13 GIG4 |
| sp\|Q9NX63\|MIC19_HUMAN | 82,817 | 88,274 | 178,521 | 129,953 | sp\|Q9NX63\|MIC19_HUMAN | 124,990 | 48,781 | 39% | Q9NX63 | MIC19_HUMAN | MIC19_HUMAN | MICOS complex subunit MIC19 (Coiled-coil-helix-coiled-coil-helix domain-containing protein 3) | CHCHD3 MIC19 MINOS3 |
| sp\|Q9Y6M5\|ZNT1_HUMAN | 208,871 | 207,585 | 167,509 | 119,660 | sp\|Q9Y6M5\|ZNT1_HUMAN | 151,785 | 47,684 | 31% | Q9Y6M5 | ZNT1_HUMAN | ZNT1_HUMAN | Zinc transporter 1 (ZnT-1) (Solute carrier family 30 member 1) | SLC30A1 ZNT1 |
| sp\|P62979\|RS27A_HUMAN | 22,235,169 | 14,706,050 | 7,276,671 | 14,296,793 | sp\|P62979\|RS27A_HUMAN | 12,477,176 | 6,205,785 | 50% | P62979 | RS27A_HUMAN | RS27A_HUMAN | Ubiquitin-40S ribosomal protein S27a (Ubiquitin carboxyl extension protein 80) [Cleaved into: Ubiquitin; 40S ribosomal protein S27a (Small ribosomal subunit protein eS31)] | RPS27A UBA80 UBCEP1 |
| sp\|Q96HE7\|ERO1A_HUMAN | 53,688 | 42,817 | 44,884 | 61,025 | sp\|Q96HE7\|ERO1A_HUMAN | 55,629 | 33,054 | 59% | Q96HE7 | ERO1A_HUMAN | ERO1A_HUMAN | ERO1-like protein alpha (ERO1-L) (ERO1-L-alpha) (EC 1.8.4.-) (Endoplasmic oxidoreductin-1-like protein) (Endoplasmic reticulum oxidoreductase alpha) (Oxidoreductin-1-L-alpha) | ERO1A ERO1L UNQ434/PRO865 |
| sp\|P60900\|PSA6_HUMAN | 421,423 | 432,489 | 170,703 | 388,917 | sp\|P60900\|PSA6_HUMAN | 331,903 | 90,259 | 27% | P60900 | PSA6_HUMAN | PSA6_HUMAN | Proteasome subunit alpha type-6 (27 kDa prosomal protein) (PROS-27) (p27K) (Macropain iota chain) (Multicatalytic endopeptidase complex iota chain) (Proteasome iota chain) | PSMA6 PROS27 |
| sp\|Q99460\|PSMD1_HUMAN | 262,429 | 180,377 | 69,100 | 255,938 | sp\|Q99460\|PSMD1_HUMAN | 181,590 | 61,527 | 34% | Q99460 | PSMD1_HUMAN | PSMD1_HUMAN | 26S proteasome non-ATPase regulatory subunit 1 (26S proteasome regulatory subunit RPN2) (26S proteasome regulatory subunit S1) (26S proteasome subunit p112) | PSMD1 |
| sp\|P60953\|CDC42_HUMAN | 1,107,784 | 1,554,247 | 227,502 | 676,766 | sp\|P60953\|CDC42_HUMAN | 855,786 | 474,935 | 55% | P60953 | CDC42_HUMAN | CDC42_HUMAN | Cell division control protein 42 homolog (EC 3.6.5.2) (G25K GTP-binding protein) | CDC42 |
| sp\|P12955\|PEPD_HUMAN | 165,504 | 206,217 | 170,964 | 236,721 | sp\|P12955\|PEPD_HUMAN | 216,123 | 49,817 | 23% | P12955 | PEPD_HUMAN | PEPD_HUMAN | Xaa-Pro dipeptidase (X-Pro dipeptidase) (EC 3.4.13.9) (Imidodipeptidase) (Peptidase D) (Proline dipeptidase) (Prolidase) | PEPD PRD |
| sp\|O15371\|EIF3D_HUMAN | 182,976 | 212,509 | 384,378 | 298,331 | sp\|O15371\|EIF3D_HUMAN | 237,946 | 71,662 | 30% | O15371 | EIF3D_HUMAN | EIF3D_HUMAN | Eukaryotic translation initiation factor 3 subunit D (eIF3d) (Eukaryotic translation initiation factor 3 subunit 7) (eIF-3-zeta) (eIF3 p66) | EIF3D EIF3S7 |
| sp\|Q53GQ0\|DHB12_HUMAN | 176,807 | 200,953 | 95,562 | 134,138 | sp\|Q53GQ0\|DHB12_HUMAN | 131,368 | 41,352 | 31% | Q53GQ0 | DHB12_HUMAN | DHB12_HUMAN | Very-long-chain 3-oxoacyl-CoA reductase (EC 1.1.1.330) (17-beta-hydroxysteroid dehydrogenase 12) (17-beta-HSD 12) (3-ketoacyl-CoA reductase) (KAR) (Estradiol 17-beta-dehydrogenase 12) (EC 1.1.1.62) (Short chain dehydrogenase/reductase family 12C member 1) | HSD17B12 SDR12C1 |

| | | | | | | | | | | | | | |
|---|---|---|---|---|---|---|---|---|---|---|---|---|---|
| sp\|Q9H4A4\|AMPB_HUMAN | 159,065 | 192,644 | 352,103 | 336,478 | sp\|Q9H4A4\|AMPB_HUMAN | 249,104 | 85,537 | 34% | Q9H4A4 | AMPB_HUMAN | AMPB_HUMAN | Aminopeptidase B (AP-B) (EC 3.4.11.6) (Arginine aminopeptidase) (Arginyl aminopeptidase) | RNPEP APB |
| sp\|P83731\|RL24_HUMAN | 1,386,280 | 1,197,543 | 934,265 | 818,383 | sp\|P83731\|RL24_HUMAN | 773,925 | 374,757 | 48% | P83731 | RL24_HUMAN | RL24_HUMAN | 60S ribosomal protein L24 (60S ribosomal protein L30) (Large ribosomal subunit protein eL24) | RPL24 |
| sp\|P27635\|RL10_HUMAN | 215,984 | 263,270 | 96,972 | 75,580 | sp\|P27635\|RL10_HUMAN | 108,325 | 88,698 | 82% | P27635 | RL10_HUMAN | RL10_HUMAN | 60S ribosomal protein L10 (Laminin receptor homolog) (Large ribosomal subunit protein uL16) (Protein QM) (Ribosomal protein L10) (Tumor suppressor QM) | RPL10 DXS648E QM |
| sp\|P60201\|MYPR_HUMAN | 4,096,485 | 4,094,199 | 2,891,742 | 3,135,206 | sp\|P60201\|MYPR_HUMAN | 3,028,901 | 855,144 | 28% | P60201 | MYPR_HUMAN | MYPR_HUMAN | Myelin proteolipid protein (PLP) (Lipophilin) | PLP1 PLP |
| sp\|P13804\|ETFA_HUMAN | 80,312 | 112,212 | 30,969 | 33,468 | sp\|P13804\|ETFA_HUMAN | 64,393 | 30,940 | 48% | P13804 | ETFA_HUMAN | ETFA_HUMAN | Electron transfer flavoprotein subunit alpha, mitochondrial (Alpha-ETF) | ETFA |
| sp\|Q12765\|SCRN1_HUMAN | 166,107 | 219,989 | 239,785 | 272,120 | sp\|Q12765\|SCRN1_HUMAN | 209,037 | 40,188 | 19% | Q12765 | SCRN1_HUMAN | SCRN1_HUMAN | Secernin-1 | SCRN1 KIAA0193 |
| sp\|Q8WTV0\|SCRB1_HUMAN | 1,045,016 | 1,606,105 | 545,294 | 1,004,496 | sp\|Q8WTV0\|SCRB1_HUMAN | 947,464 | 369,520 | 39% | Q8WTV0 | SCRB1_HUMAN | SCRB1_HUMAN | Scavenger receptor class B member 1 (SRB1) (CD36 and LIMPII analogous 1) (CLA-1) (CD36 antigen-like 1) (Collagen type I receptor, thrombospondin receptor-like 1) (SR-BI) (CD antigen CD36) | SCARB1 CD36L1 CLA1 |
| sp\|Q04917\|1433F_HUMAN | 506,157 | 636,943 | 328,156 | 404,925 | sp\|Q04917\|1433F_HUMAN | 440,025 | 150,771 | 34% | Q04917 | 1433F_HUMAN | 1433F_HUMAN | 14-3-3 protein eta (Protein AS1) | YWHAH YWHA1 |
| sp\|Q96AY3\|FKB10_HUMAN | 48,919 | 69,047 | 58,142 | 67,282 | sp\|Q96AY3\|FKB10_HUMAN | 55,892 | 15,379 | 28% | Q96AY3 | FKB10_HUMAN | FKB10_HUMAN | Peptidyl-prolyl cis-trans isomerase FKBP10 (PPIase FKBP10) (EC 5.2.1.8) (65 kDa FK506-binding protein) (65 kDa FKBP) (FKBP-65) (FK506-binding protein 10) (FKBP-10) (Immunophilin FKBP65) (Rotamase) | FKBP10 FKBP65 PSEC0056 |
| sp\|Q9Y277\|VDAC3_HUMAN | 55,960 | 85,335 | 38,130 | 43,679 | sp\|Q9Y277\|VDAC3_HUMAN | 55,810 | 23,659 | 42% | Q9Y277 | VDAC3_HUMAN | VDAC3_HUMAN | Voltage-dependent anion-selective channel protein 3 (VDAC-3) (hVDAC3) (Outer mitochondrial membrane protein porin 3) | VDAC3 |
| sp\|P14314\|GLU2B_HUMAN | 25,575 | 57,523 | 114,200 | 118,503 | sp\|P14314\|GLU2B_HUMAN | 55,670 | 40,485 | 73% | P14314 | GLU2B_HUMAN | GLU2B_HUMAN | Glucosidase 2 subunit beta (80K-H protein) (Glucosidase II subunit beta) (Protein kinase C substrate 60.1 kDa protein heavy chain) (PKCSH) | PRKCSH G19P1 |
| sp\|P10644\|KAP0_HUMAN | 137,391 | 140,169 | 93,132 | 118,900 | sp\|P10644\|KAP0_HUMAN | 146,191 | 38,447 | 26% | P10644 | KAP0_HUMAN | KAP0_HUMAN | cAMP-dependent protein kinase type I-alpha regulatory subunit (Tissue-specific extinguisher 1) (TSE1) | PRKAR1A PKR1 PRKAR1 TSE1 |
| sp\|P35613\|BASI_HUMAN | 997,293 | 1,646,519 | 1,230,508 | 1,509,721 | sp\|P35613\|BASI_HUMAN | 1,287,668 | 257,292 | 20% | P35613 | BASI_HUMAN | BASI_HUMAN | Basigin (5F7) (Collagenase stimulatory factor) (Extracellular matrix metalloproteinase inducer) (EMMPRIN) (Hepatoma-associated antigen) (HAb18G) (Leukocyte activation antigen M6) (OK blood group antigen) (Tumor cell-derived collagenase stimulatory factor) (TCSF) (CD antigen CD147) | BSG UNQ6505/PRO21383 |
| sp\|Q9Y6E2\|BZW2_HUMAN | 207,795 | 185,104 | 207,374 | 229,867 | sp\|Q9Y6E2\|BZW2_HUMAN | 242,482 | 85,076 | 35% | Q9Y6E2 | BZW2_HUMAN | 5MP1_HUMAN | eIF5-mimic protein 1 (Basic leucine zipper and W2 domain-containing protein 2) | BZW2 5MP1 HSPC028 MSTP017 |

| | | | | | | | | | | | | | |
|---|---|---|---|---|---|---|---|---|---|---|---|---|---|
| sp\|P30041\|PRDX6_HUMAN | 620,683 | 726,706 | 533,167 | 428,611 | sp\|P30041\|PRDX6_HUMAN | 545,245 | 140,030 | 26% | P30041 | PRDX6_HUMAN | PRDX6_HUMAN | Peroxiredoxin-6 (EC 1.11.1.27) (1-Cys peroxiredoxin) (1-Cys PRX) (24 kDa protein) (Acidic calcium-independent phospholipase A2) (aiPLA2) (EC 3.1.1.4) (Antioxidant protein 2) (Glutathione-dependent peroxiredoxin) (Liver 2D page spot 40) (Lysophosphatidylcholine acyltransferase 5) (LPC acyltransferase 5) (LPCAT-5) (Lyso-PC acyltransferase 5) (EC 2.3.1.23) (Non-selenium glutathione peroxidase) (NSGPx) (Red blood cells page spot 12) | PRDX6 AOP2 KIAA0106 |
| sp\|P54578\|UBP14_HUMAN | 245,182 | 306,984 | 196,521 | 303,186 | sp\|P54578\|UBP14_HUMAN | 250,584 | 51,926 | 21% | P54578 | UBP14_HUMAN | UBP14_HUMAN | Ubiquitin carboxyl-terminal hydrolase 14 (EC 3.4.19.12) (Deubiquitinating enzyme 14) (Ubiquitin thioesterase 14) (Ubiquitin-specific-processing protease 14) | USP14 TGT |
| sp\|P26599\|PTBP1_HUMAN | 43,522 | 64,797 | 22,863 | 87,151 | sp\|P26599\|PTBP1_HUMAN | 41,356 | 24,474 | 59% | P26599 | PTBP1_HUMAN | PTBP1_HUMAN | Polypyrimidine tract-binding protein 1 (PTB) (57 kDa RNA-binding protein PPTB-1) (Heterogeneous nuclear ribonucleoprotein I) (hnRNP I) | PTBP1 PTB |
| sp\|P11233\|RALA_HUMAN | 1,301,538 | 1,706,795 | 706,169 | 883,870 | sp\|P11233\|RALA_HUMAN | 1,021,895 | 405,370 | 40% | P11233 | RALA_HUMAN | RALA_HUMAN | Ras-related protein Ral-A (EC 3.6.5.2) | RALA RAL |
| sp\|P28074\|PSB5_HUMAN | 289,792 | 258,022 | 538,141 | 354,047 | sp\|P28074\|PSB5_HUMAN | 313,653 | 106,403 | 34% | P28074 | PSB5_HUMAN | PSB5_HUMAN | Proteasome subunit beta type-5 (EC 3.4.25.1) (Macropain epsilon chain) (Multicatalytic endopeptidase complex epsilon chain) (Proteasome chain 6) (Proteasome epsilon chain) (Proteasome subunit MB1) (Proteasome subunit X) | PSMB5 LMPX MB1 X |
| sp\|Q04837\|SSBP_HUMAN | 140,590 | 131,080 | 216,203 | 162,177 | sp\|Q04837\|SSBP_HUMAN | 145,022 | 39,862 | 27% | Q04837 | SSBP_HUMAN | SSBP_HUMAN | Single-stranded DNA-binding protein, mitochondrial (Mt-SSB) (MtSSB) (PWP1-interacting protein 17) | SSBP1 SSBP |
| sp\|P08708\|RS17_HUMAN | 252,453 | 336,614 | 212,800 | 212,178 | sp\|P08708\|RS17_HUMAN | 230,052 | 54,189 | 24% | P08708 | RS17_HUMAN | RS17_HUMAN | 40S ribosomal protein S17 (Small ribosomal subunit protein eS17) | RPS17 RPS17L |
| sp\|Q07960\|RHG01_HUMAN | 93,849 | 141,677 | 16,984 | 58,026 | sp\|Q07960\|RHG01_HUMAN | 82,953 | 38,135 | 46% | Q07960 | RHG01_HUMAN | RHG01_HUMAN | Rho GTPase-activating protein 1 (CDC42 GTPase-activating protein) (GTPase-activating protein rhoGAP) (Rho-related small GTPase protein activator) (Rho-type GTPase-activating protein 1) (p50-RhoGAP) | ARHGAP1 CDC42GAP RHOGAP1 |
| sp\|O14817\|TSN4_HUMAN | 3,251,886 | 3,580,106 | 3,496,598 | 2,970,202 | sp\|O14817\|TSN4_HUMAN | 2,519,595 | 967,508 | 38% | O14817 | TSN4_HUMAN | TSN4_HUMAN | Tetraspanin-4 (Tspan-4) (Novel antigen 2) (NAG-2) (Transmembrane 4 superfamily member 7) | TSPAN4 NAG2 TM4SF7 |
| sp\|P02768\|ALBU_HUMAN | 3,927,617 | 2,357,180 | 3,312,054 | 1,085,146 | sp\|P02768\|ALBU_HUMAN | 32,480,109 | 31,922,249 | 98% | P02768 | ALBU_HUMAN | ALBU_HUMAN | Albumin | ALB GIG20 GIG42 PRO0903 PRO1708 PRO2044 PRO2619 PRO2675 UNQ696/PRO1341 |
| sp\|P62330\|ARF6_HUMAN | 840,559 | 1,221,793 | 773,148 | 1,121,136 | sp\|P62330\|ARF6_HUMAN | 992,424 | 200,406 | 20% | P62330 | ARF6_HUMAN | ARF6_HUMAN | ADP-ribosylation factor 6 (EC 3.6.5.2) | ARF6 |
| sp\|P27708\|PYR1_HUMAN | 191,077 | 164,620 | 337,207 | 234,849 | sp\|P27708\|PYR1_HUMAN | 220,093 | 75,720 | 34% | P27708 | PYR1_HUMAN | PYR1_HUMAN | CAD protein [Includes: Glutamine-dependent carbamoyl-phosphate synthase (EC 6.3.5.5); Aspartate carbamoyltransferase (EC 2.1.3.2); Dihydroorotase (EC 3.5.2.3)] | CAD |

| | | | | | | | | | | | | | |
|---|---|---|---|---|---|---|---|---|---|---|---|---|---|
| sp\|P43490\|NAMPT_HUMAN | 68,682 | 94,851 | 42,293 | 41,643 | sp\|P43490\|NAMPT_HUMAN | 55,898 | 23,550 | 42% | P43490 | NAMPT_HUMAN | NAMPT_HUMAN | Nicotinamide phosphoribosyltransferase (NAmPRTase) (Nampt) (EC 2.4.2.12) (Pre-B-cell colony-enhancing factor 1) (Pre-B cell-enhancing factor) (Visfatin) | NAMPT PBEF PBEF1 |
| sp\|P61353\|RL27_HUMAN | 1,562,605 | 1,167,567 | 1,507,513 | 1,266,168 | sp\|P61353\|RL27_HUMAN | 937,626 | 492,282 | 53% | P61353 | RL27_HUMAN | RL27_HUMAN | 60S ribosomal protein L27 (Large ribosomal subunit protein eL27) | RPL27 |
| sp\|P46977\|STT3A_HUMAN | 6,689 | 5,902 | 10,820 | 2,140 | sp\|P46977\|STT3A_HUMAN | 4,456 | 3,143 | 71% | P46977 | STT3A_HUMAN | STT3A_HUMAN | Dolichyl-diphosphooligosaccharide--protein glycosyltransferase subunit STT3A (Oligosaccharyl transferase subunit STT3A) (STT3-A) (EC 2.4.99.18) (B5) (Integral membrane protein 1) (Transmembrane protein TMC) | STT3A ITM1 TMC |
| sp\|P62873\|GBB1_HUMAN | 2,340,385 | 2,495,482 | 2,165,833 | 2,305,527 | sp\|P62873\|GBB1_HUMAN | 2,181,511 | 191,424 | 9% | P62873 | GBB1_HUMAN | GBB1_HUMAN | Guanine nucleotide-binding protein G(I)/G(S)/G(T) subunit beta-1 (Transducin beta chain 1) | GNB1 |
| sp\|P78324\|SHPS1_HUMAN | 711,859 | 680,835 | 1,258,333 | 1,003,529 | sp\|P78324\|SHPS1_HUMAN | 716,030 | 290,689 | 41% | P78324 | SHPS1_HUMAN | SHPS1_HUMAN | Tyrosine-protein phosphatase non-receptor type substrate 1 (SHP substrate 1) (SHPS-1) (Brain Ig-like molecule with tyrosine-based activation motifs) (Bit) (CD172 antigen-like family member A) (Inhibitory receptor SHPS-1) (Macrophage fusion receptor) (MyD-1 antigen) (Signal-regulatory protein alpha-1) (Sirp-alpha-1) (Signal-regulatory protein alpha-2) (Sirp-alpha-2) (Signal-regulatory protein alpha-3) (Sirp-alpha-3) (p84) (CD antigen CD172a) | SIRPA BIT MFR MYD1 PTPNS1 SHPS1 SIRP |
| sp\|P53004\|BIEA_HUMAN | 285,294 | 382,060 | 238,043 | 138,676 | sp\|P53004\|BIEA_HUMAN | 287,909 | 84,336 | 29% | P53004 | BIEA_HUMAN | BIEA_HUMAN | Biliverdin reductase A (BVR A) (EC 1.3.1.24) (Biliverdin-IX alpha-reductase) | BLVRA BLVR BVR |
| sp\|P46783\|RS10_HUMAN | 185,383 | 660,067 | 244,426 | 795,113 | sp\|P46783\|RS10_HUMAN | 390,128 | 237,802 | 61% | P46783 | RS10_HUMAN | RS10_HUMAN | 40S ribosomal protein S10 (Small ribosomal subunit protein eS10) | RPS10 |
| sp\|P61026\|RAB10_HUMAN | 843,351 | 1,289,619 | 748,511 | 1,232,110 | sp\|P61026\|RAB10_HUMAN | 974,606 | 210,822 | 22% | P61026 | RAB10_HUMAN | RAB10_HUMAN | Ras-related protein Rab-10 (EC 3.6.5.2) | RAB10 |
| sp\|P61764\|STXB1_HUMAN | 122,188 | 136,635 | 140,134 | 199,340 | sp\|P61764\|STXB1_HUMAN | 158,479 | 49,996 | 32% | P61764 | STXB1_HUMAN | STXB1_HUMAN | Syntaxin-binding protein 1 (MUNC18-1) (N-Sec1) (Protein unc-18 homolog 1) (Unc18-1) (Protein unc-18 homolog A) (Unc-18A) (p67) | STXBP1 UNC18A |
| sp\|Q9UL46\|PSME2_HUMAN | 182,384 | 267,954 | 227,632 | 466,844 | sp\|Q9UL46\|PSME2_HUMAN | 306,290 | 120,547 | 39% | Q9UL46 | PSME2_HUMAN | PSME2_HUMAN | Proteasome activator complex subunit 2 (11S regulator complex subunit beta) (REG-beta) (Activator of multicatalytic protease subunit 2) (Proteasome activator 28 subunit beta) (PA28b) (PA28beta) | PSME2 |
| sp\|Q9BSJ8\|ESYT1_HUMAN | 206,088 | 190,233 | 117,693 | 253,538 | sp\|Q9BSJ8\|ESYT1_HUMAN | 218,920 | 85,771 | 39% | Q9BSJ8 | ESYT1_HUMAN | ESYT1_HUMAN | Extended synaptotagmin-1 (E-Syt1) (Membrane-bound C2 domain-containing protein) | ESYT1 FAM62A KIAA0747 MBC2 |
| sp\|P52565\|GDIR1_HUMAN | 403,917 | 534,877 | 266,670 | 362,861 | sp\|P52565\|GDIR1_HUMAN | 354,793 | 116,696 | 33% | P52565 | GDIR1_HUMAN | GDIR1_HUMAN | Rho GDP-dissociation inhibitor 1 (Rho GDI 1) (Rho-GDI alpha) | ARHGDIA GDIA1 |
| sp\|Q13492\|PICAL_HUMAN | 336,825 | 441,872 | 178,493 | 242,941 | sp\|Q13492\|PICAL_HUMAN | 277,649 | 82,180 | 30% | Q13492 | PICAL_HUMAN | PICAL_HUMAN | Phosphatidylinositol-binding clathrin assembly protein (Clathrin assembly lymphoid myeloid leukemia protein) | PICALM CALM |

| | | | | | | | | | | | | | |
|---|---|---|---|---|---|---|---|---|---|---|---|---|---|
| sp\|P60891\|PRPS1_HUMAN | 27,621 | 27,170 | 31,539 | 27,052 | sp\|P60891\|PRPS1_HUMAN | 19,302 | 11,601 | 60% | P60891 | PRPS1_HUMAN | PRPS1_HUMAN | Ribose-phosphate pyrophosphokinase 1 (EC 2.7.6.1) (PPRibP) (Phosphoribosyl pyrophosphate synthase I) (PRS-I) | PRPS1 |
| sp\|P20042\|IF2B_HUMAN | 303,734 | 319,977 | 336,694 | 358,891 | sp\|P20042\|IF2B_HUMAN | 327,815 | 33,266 | 10% | P20042 | IF2B_HUMAN | IF2B_HUMAN | Eukaryotic translation initiation factor 2 subunit 2 (Eukaryotic translation initiation factor 2 subunit beta) (eIF-2-beta) | EIF2S2 EIF2B |
| sp\|P07437\|TBB5_HUMAN | 806,912 | 2,199,827 | 1,539,274 | 1,742,924 | sp\|P07437\|TBB5_HUMAN | 1,177,883 | 638,028 | 54% | P07437 | TBB5_HUMAN | TBB5_HUMAN | Tubulin beta chain (Tubulin beta-5 chain) | TUBB TUBB5 OK/SW-cl.56 |
| sp\|Q99523\|SORT_HUMAN | 562,428 | 556,122 | 612,130 | 543,512 | sp\|Q99523\|SORT_HUMAN | 421,624 | 160,450 | 38% | Q99523 | SORT_HUMAN | SORT_HUMAN | Sortilin (100 kDa NT receptor) (Glycoprotein 95) (Gp95) (Neurotensin receptor 3) (NT3) (NTR3) | SORT1 |
| sp\|P67809\|YBOX1_HUMAN | 131,681 | 154,402 | 125,930 | 121,506 | sp\|P67809\|YBOX1_HUMAN | 103,862 | 40,550 | 39% | P67809 | YBOX1_HUMAN | YBOX1_HUMAN | Y-box-binding protein 1 (YB-1) (CCAAT-binding transcription factor I subunit A) (CBF-A) (DNA-binding protein B) (DBPB) (Enhancer factor I subunit A) (EFI-A) (Nuclease-sensitive element-binding protein 1) (Y-box transcription factor) | YBX1 NSEP1 YB1 |
| sp\|P62826\|RAN_HUMAN | 1,544,492 | 1,526,464 | 782,191 | 719,696 | sp\|P62826\|RAN_HUMAN | 883,259 | 453,189 | 51% | P62826 | RAN_HUMAN | RAN_HUMAN | GTP-binding nuclear protein Ran (EC 3.6.5.-) (Androgen receptor-associated protein 24) (GTPase Ran) (Ras-like protein TC4) (Ras-related nuclear protein) | RAN ARA24 OK/SW-cl.81 |
| sp\|P25398\|RS12_HUMAN | 506,004 | 535,541 | 179,709 | 147,765 | sp\|P25398\|RS12_HUMAN | 264,158 | 192,490 | 73% | P25398 | RS12_HUMAN | RS12_HUMAN | 40S ribosomal protein S12 (Small ribosomal subunit protein eS12) | RPS12 |
| sp\|P02538\|K2C6A_HUMAN | 471,468 | 543,637 | 789,745 | 576,605 | sp\|P02538\|K2C6A_HUMAN | 839,169 | 428,300 | 51% | P02538 | K2C6A_HUMAN | K2C6A_HUMAN | Keratin, type II cytoskeletal 6A (Cytokeratin-6A) (CK-6A) (Cytokeratin-6D) (CK-6D) (Keratin-6A) (K6A) (Type-II keratin Kb6) (allergen Hom s 5) | KRT6A K6A KRT6D |
| sp\|P36957\|ODO2_HUMAN | 174,711 | 175,937 | 292,607 | 304,736 | sp\|P36957\|ODO2_HUMAN | 188,697 | 77,199 | 41% | P36957 | ODO2_HUMAN | ODO2_HUMAN | Dihydrolipoyllysine-residue succinyltransferase component of 2-oxoglutarate dehydrogenase complex, mitochondrial (EC 2.3.1.61) (2-oxoglutarate dehydrogenase complex component E2) (OGDC-E2) (Dihydrolipoamide succinyltransferase component of 2-oxoglutarate dehydrogenase complex) (E2K) | DLST DLTS |
| sp\|P50281\|MMP14_HUMAN | 310,898 | 320,780 | 375,413 | 340,922 | sp\|P50281\|MMP14_HUMAN | 306,803 | 56,365 | 18% | P50281 | MMP14_HUMAN | MMP14_HUMAN | Matrix metalloproteinase-14 (MMP-14) (EC 3.4.24.80) (MMP-X1) (Membrane-type matrix metalloproteinase 1) (MT-MMP 1) (MTMMP1) (Membrane-type-1 matrix metalloproteinase) (MT1-MMP) (MT1MMP) | MMP14 |
| sp\|Q9Y512\|SAM50_HUMAN | 18,281 | 19,445 | 9,286 | 20,457 | sp\|Q9Y512\|SAM50_HUMAN | 19,437 | 10,821 | 56% | Q9Y512 | SAM50_HUMAN | SAM50_HUMAN | Sorting and assembly machinery component 50 homolog (Transformation-related gene 3 protein) (TRG-3) | SAMM50 SAM50 CGI-51 TRG3 |

| | | | | | | | | | | | | | |
|---|---|---|---|---|---|---|---|---|---|---|---|---|---|
| sp\|Q9UGH3\|S23A2_HUMAN | 682,618 | 850,054 | 532,934 | 752,746 | sp\|Q9UGH3\|S23A2_HUMAN | 676,201 | 102,042 | 15% | Q9UGH3 | S23A2_HUMAN | S23A2_HUMAN | Solute carrier family 23 member 2 (Na(+)/L-ascorbic acid transporter 2) (Nucleobase transporter-like 1 protein) (Sodium-dependent vitamin C transporter 2) (hSVCT2) (Yolk sac permease-like molecule 2) | SLC23A2 KIAA0238 NBTL1 SLC23A1 SVCT2 YSPL2 |
| sp\|Q99733\|NP1L4_HUMAN | 130,513 | 147,010 | 246,832 | 201,335 | sp\|Q99733\|NP1L4_HUMAN | 171,443 | 47,621 | 28% | Q99733 | NP1L4_HUMAN | NP1L4_HUMAN | Nucleosome assembly protein 1-like 4 (Nucleosome assembly protein 2) (NAP-2) | NAP1L4 NAP2 |
| sp\|P07954\|FUMH_HUMAN | 62,821 | 76,424 | 66,044 | 87,645 | sp\|P07954\|FUMH_HUMAN | 65,214 | 16,656 | 26% | P07954 | FUMH_HUMAN | FUMH_HUMAN | Fumarate hydratase, mitochondrial (Fumarase) (HsFH) (EC 4.2.1.2) | FH |
| sp\|Q7L2H7\|EIF3M_HUMAN | 271,733 | 377,607 | 206,876 | 147,216 | sp\|Q7L2H7\|EIF3M_HUMAN | 279,714 | 93,428 | 33% | Q7L2H7 | EIF3M_HUMAN | EIF3M_HUMAN | Eukaryotic translation initiation factor 3 subunit M (eIF3m) (Fetal lung protein B5) (hFL-B5) (PCI domain-containing protein 1) | EIF3M HFLB5 PCID1 GA17 PNAS-125 |
| sp\|Q9UP95\|S12A4_HUMAN | 301,134 | 297,691 | 264,283 | 380,263 | sp\|Q9UP95\|S12A4_HUMAN | 249,596 | 86,236 | 35% | Q9UP95 | S12A4_HUMAN | S12A4_HUMAN | Solute carrier family 12 member 4 (Electroneutral potassium-chloride cotransporter 1) (Erythroid K-Cl cotransporter 1) (hKCC1) | SLC12A4 KCC1 |
| sp\|P17980\|PRS6A_HUMAN | 291,046 | 294,418 | 330,054 | 294,956 | sp\|P17980\|PRS6A_HUMAN | 258,966 | 51,954 | 20% | P17980 | PRS6A_HUMAN | PRS6A_HUMAN | 26S proteasome regulatory subunit 6A (26S proteasome AAA-ATPase subunit RPT5) (Proteasome 26S subunit ATPase 3) (Proteasome subunit P50) (Tat-binding protein 1) (TBP-1) | PSMC3 TBP1 |
| sp\|P62191\|PRS4_HUMAN | 278,357 | 196,046 | 146,625 | 131,954 | sp\|P62191\|PRS4_HUMAN | 204,976 | 57,920 | 28% | P62191 | PRS4_HUMAN | PRS4_HUMAN | 26S proteasome regulatory subunit 4 (P26s4) (26S proteasome AAA-ATPase subunit RPT2) (Proteasome 26S subunit ATPase 1) | PSMC1 |
| sp\|Q9H3N1\|TMX1_HUMAN | 67,305 | 69,898 | 51,192 | 63,149 | sp\|Q9H3N1\|TMX1_HUMAN | 62,374 | 28,340 | 45% | Q9H3N1 | TMX1_HUMAN | TMX1_HUMAN | Thioredoxin-related transmembrane protein 1 (Thioredoxin domain-containing protein 1) (Transmembrane Trx-related protein) | TMX1 TMX TXNDC TXNDC1 PSEC0085 UNQ235/PRO268 |
| sp\|P11279\|LAMP1_HUMAN | 979,296 | 794,157 | 1,215,457 | 985,915 | sp\|P11279\|LAMP1_HUMAN | 733,887 | 302,178 | 41% | P11279 | LAMP1_HUMAN | LAMP1_HUMAN | Lysosome-associated membrane glycoprotein 1 (LAMP-1) (Lysosome-associated membrane protein 1) (CD107 antigen-like family member A) (CD antigen CD107a) | LAMP1 |
| sp\|O43488\|ARK72_HUMAN | 246,295 | 312,711 | 309,467 | 407,418 | sp\|O43488\|ARK72_HUMAN | 279,832 | 64,163 | 23% | O43488 | ARK72_HUMAN | ARK72_HUMAN | Aflatoxin B1 aldehyde reductase member 2 (EC 1.1.1.n11) (AFB1 aldehyde reductase 1) (AFB1-AR 1) (Aldoketoreductase 7) (Succinic semialdehyde reductase) (SSA reductase) | AKR7A2 AFAR AFAR1 AKR7 |
| sp\|P35222\|CTNB1_HUMAN | 151,754 | 148,363 | 257,607 | 287,620 | sp\|P35222\|CTNB1_HUMAN | 336,242 | 335,025 | 100% | P35222 | CTNB1_HUMAN | CTNB1_HUMAN | Catenin beta-1 (Beta-catenin) | CTNNB1 CTNNB OK/SW-cl.35 PRO2286 |
| sp\|P69905\|HBA_HUMAN | 2,115,605 | 1,352,928 | 1,665,807 | 798,977 | sp\|P69905\|HBA_HUMAN | 8,780,001 | 8,431,053 | 96% | P69905 | HBA_HUMAN | HBA_HUMAN | Hemoglobin subunit alpha (Alpha-globin) (Hemoglobin alpha chain) | HBA1; HBA2 |
| sp\|Q92499\|DDX1_HUMAN | 377,032 | 375,403 | 567,320 | 514,019 | sp\|Q92499\|DDX1_HUMAN | 394,245 | 111,522 | 28% | Q92499 | DDX1_HUMAN | DDX1_HUMAN | ATP-dependent RNA helicase DDX1 (EC 3.6.4.13) (DEAD box protein 1) (DEAD box protein retinoblastoma) (DBP-RB) | DDX1 |
| sp\|Q9NVJ2\|ARL8B_HUMAN | 403,052 | 391,636 | 308,871 | 228,696 | sp\|Q9NVJ2\|ARL8B_HUMAN | 272,258 | 96,897 | 36% | Q9NVJ2 | ARL8B_HUMAN | ARL8B_HUMAN | ADP-ribosylation factor-like protein 8B (EC 3.6.5.2) (ADP-ribosylation factor-like protein 10C) (Novel small G protein indispensable for equal chromosome segregation 1) | ARL8B ARL10C GIE1 |

| | | | | | | | | | | | | | |
|---|---|---|---|---|---|---|---|---|---|---|---|---|---|
| sp\|P09651\|ROA1_HUMAN | 361,601 | 370,165 | 224,348 | 208,992 | sp\|P09651\|ROA1_HUMAN | 201,684 | 112,640 | 56% | P09651 | ROA1_HUMAN | ROA1_HUMAN | Heterogeneous nuclear ribonucleoprotein A1 (hnRNP A1) (Helix-destabilizing protein) (Single-strand RNA-binding protein) (hnRNP core protein A1) [Cleaved into: Heterogeneous nuclear ribonucleoprotein A1, N-terminally processed] | HNRNPA1 HNRPA1 |
| sp\|P48449\|ERG7_HUMAN | 85,591 | 82,795 | 94,774 | 147,581 | sp\|P48449\|ERG7_HUMAN | 117,712 | 57,867 | 49% | P48449 | ERG7_HUMAN | LSS_HUMAN | Lanosterol synthase (EC 5.4.99.7) (2,3-epoxysqualene--lanosterol cyclase) (Oxidosqualene--lanosterol cyclase) (OSC) (hOSC) | LSS OSC |
| sp\|O75578\|ITA10_HUMAN | 398,092 | 411,194 | 577,151 | 646,465 | sp\|O75578\|ITA10_HUMAN | 489,531 | 136,660 | 28% | O75578 | ITA10_HUMAN | ITA10_HUMAN | Integrin alpha-10 | ITGA10 UNQ468/PRO827 |
| sp\|P25789\|PSA4_HUMAN | 515,617 | 622,592 | 362,894 | 339,445 | sp\|P25789\|PSA4_HUMAN | 408,691 | 134,435 | 33% | P25789 | PSA4_HUMAN | PSA4_HUMAN | Proteasome subunit alpha type-4 (Macropain subunit C9) (Multicatalytic endopeptidase complex subunit C9) (Proteasome component C9) (Proteasome subunit L) | PSMA4 HC9 PSC9 |
| sp\|P61604\|CH10_HUMAN | 531,065 | 465,092 | 206,899 | 268,721 | sp\|P61604\|CH10_HUMAN | 269,810 | 159,851 | 59% | P61604 | CH10_HUMAN | CH10_HUMAN | 10 kDa heat shock protein, mitochondrial (Hsp10) (10 kDa chaperonin) (Chaperonin 10) (CPN10) (Early-pregnancy factor) (EPF) | HSPE1 |
| sp\|P18077\|RL35A_HUMAN | 973,102 | 706,032 | 1,266,140 | 1,586,678 | sp\|P18077\|RL35A_HUMAN | 799,228 | 436,582 | 55% | P18077 | RL35A_HUMAN | RL35A_HUMAN | 60S ribosomal protein L35a (Cell growth-inhibiting gene 33 protein) (Large ribosomal subunit protein eL33) | RPL35A GIG33 |
| sp\|Q9NR31\|SAR1A_HUMAN | 72,522 | 75,600 | 65,815 | 70,554 | sp\|Q9NR31\|SAR1A_HUMAN | 64,862 | 22,930 | 35% | Q9NR31 | SAR1A_HUMAN | SAR1A_HUMAN | GTP-binding protein SAR1a (COPII-associated small GTPase) | SAR1A SAR1 SARA SARA1 |
| sp\|Q9UHB9\|SRP68_HUMAN | 108,051 | 95,169 | 181,517 | 162,891 | sp\|Q9UHB9\|SRP68_HUMAN | 108,105 | 43,643 | 40% | Q9UHB9 | SRP68_HUMAN | SRP68_HUMAN | Signal recognition particle subunit SRP68 (SRP68) (Signal recognition particle 68 kDa protein) | SRP68 |
| sp\|P07947\|YES_HUMAN | 133,705 | 201,804 | 91,863 | 134,713 | sp\|P07947\|YES_HUMAN | 121,506 | 44,257 | 36% | P07947 | YES_HUMAN | YES_HUMAN | Tyrosine-protein kinase Yes (EC 2.7.10.2) (Proto-oncogene c-Yes) (p61-Yes) | YES1 YES |
| sp\|P60033\|CD81_HUMAN | 18,409,524 | 20,983,440 | 3,497,475 | 7,125,343 | sp\|P60033\|CD81_HUMAN | 9,680,779 | 7,457,270 | 77% | P60033 | CD81_HUMAN | CD81_HUMAN | CD81 antigen (26 kDa cell surface protein TAPA-1) (Target of the antiproliferative antibody 1) (Tetraspanin-28) (Tspan-28) (CD antigen CD81) | CD81 TAPA1 TSPAN28 |
| sp\|P30566\|PUR8_HUMAN | 300,530 | 389,578 | 320,392 | 397,064 | sp\|P30566\|PUR8_HUMAN | 354,873 | 67,185 | 19% | P30566 | PUR8_HUMAN | PUR8_HUMAN | Adenylosuccinate lyase (ADSL) (ASL) (EC 4.3.2.2) (Adenylosuccinase) (ASase) | ADSL AMPS |
| sp\|Q9HB71\|CYBP_HUMAN | 344,156 | 354,968 | 263,183 | 260,934 | sp\|Q9HB71\|CYBP_HUMAN | 279,571 | 49,821 | 18% | Q9HB71 | CYBP_HUMAN | CYBP_HUMAN | Calcyclin-binding protein (CacyBP) (hCacyBP) (S100A6-binding protein) (Siah-interacting protein) | CACYBP S100A6BP SIP PNAS-107 |
| sp\|P06748\|NPM_HUMAN | 226,922 | 286,138 | 165,284 | 222,840 | sp\|P06748\|NPM_HUMAN | 172,549 | 65,618 | 38% | P06748 | NPM_HUMAN | NPM_HUMAN | Nucleophosmin (NPM) (Nucleolar phosphoprotein B23) (Nucleolar protein NO38) (Numatrin) | NPM1 NPM |
| sp\|P48637\|GSHB_HUMAN | 8,330 | 77,524 | 111,727 | 112,464 | sp\|P48637\|GSHB_HUMAN | 85,375 | 48,190 | 56% | P48637 | GSHB_HUMAN | GSHB_HUMAN | Glutathione synthetase (GSH synthetase) (GSH-S) (EC 6.3.2.3) (Glutathione synthase) | GSS |

| | | | | | | | | | | | | | |
|---|---|---|---|---|---|---|---|---|---|---|---|---|---|
| sp\|Q9UBS4\|DJB11_HUMAN | 127,210 | 38,081 | 177,551 | 78,440 | sp\|Q9UBS4\|DJB11_HUMAN | 239,513 | 157,237 | 66% | Q9UBS4 | DJB11_HUMAN | DJB11_HUMAN | DnaJ homolog subfamily B member 11 (APOBEC1-binding protein 2) (ABBP-2) (DnaJ protein homolog 9) (ER-associated DNAJ) (ER-associated Hsp40 co-chaperone) (Endoplasmic reticulum DNA J domain-containing protein 3) (ER-resident protein ERdj3) (ERdj3) (ERj3p) (HEDJ) (Human DnaJ protein 9) (hDj-9) (PWP1-interacting protein 4) | DNAJB11 EDJ ERJ3 HDJ9 PSEC0121 UNQ537/PRO1080 |
| sp\|O96008\|TOM40_HUMAN | 59,631 | 21,280 | 19,184 | 75,110 | sp\|O96008\|TOM40_HUMAN | 39,254 | 24,308 | 62% | O96008 | TOM40_HUMAN | TOM40_HUMAN | Mitochondrial import receptor subunit TOM40 homolog (Protein Haymaker) (Translocase of outer membrane 40 kDa subunit homolog) (p38.5) | TOMM40 C19orf1 PEREC1 TOM40 |
| sp\|Q9NSE4\|SYIM_HUMAN | 28,872 | 40,331 | 30,871 | 17,293 | sp\|Q9NSE4\|SYIM_HUMAN | 42,382 | 26,747 | 63% | Q9NSE4 | SYIM_HUMAN | SYIM_HUMAN | Isoleucine--tRNA ligase, mitochondrial (EC 6.1.1.5) (Isoleucyl-tRNA synthetase) (IleRS) | IARS2 |
| sp\|P54819\|KAD2_HUMAN | 204,311 | 296,158 | 198,832 | 297,806 | sp\|P54819\|KAD2_HUMAN | 191,965 | 75,229 | 39% | P54819 | KAD2_HUMAN | KAD2_HUMAN | Adenylate kinase 2, mitochondrial (AK 2) (EC 2.7.4.3) (ATP-AMP transphosphorylase 2) (ATP:AMP phosphotransferase) (Adenylate monophosphate kinase) [Cleaved into: Adenylate kinase 2, mitochondrial, N-terminally processed] | AK2 ADK2 |
| sp\|P20340\|RAB6A_HUMAN | 461,272 | 568,002 | 372,893 | 344,742 | sp\|P20340\|RAB6A_HUMAN | 456,229 | 72,418 | 16% | P20340 | RAB6A_HUMAN | RAB6A_HUMAN | Ras-related protein Rab-6A (Rab-6) | RAB6A RAB6 |
| sp\|O75396\|SC22B_HUMAN | 164,634 | 161,122 | 230,892 | 200,622 | sp\|O75396\|SC22B_HUMAN | 194,099 | 50,400 | 26% | O75396 | SC22B_HUMAN | SC22B_HUMAN | Vesicle-trafficking protein SEC22b (ER-Golgi SNARE of 24 kDa) (ERS-24) (ERS24) (SEC22 vesicle-trafficking protein homolog B) (SEC22 vesicle-trafficking protein-like 1) | SEC22B SEC22L1 |
| sp\|P12956\|XRCC6_HUMAN | 372,877 | 367,701 | 320,299 | 366,932 | sp\|P12956\|XRCC6_HUMAN | 264,073 | 102,129 | 39% | P12956 | XRCC6_HUMAN | XRCC6_HUMAN | X-ray repair cross-complementing protein 6 (EC 3.6.4.-) (EC 4.2.99.-) (5'-deoxyribose-5-phosphate lyase Ku70) (5'-dRP lyase Ku70) (70 kDa subunit of Ku antigen) (ATP-dependent DNA helicase 2 subunit 1) (ATP-dependent DNA helicase II 70 kDa subunit) (CTC box-binding factor 75 kDa subunit) (CTC75) (CTCBF) (DNA repair protein XRCC6) (Lupus Ku autoantigen protein p70) (Ku70) (Thyroid-lupus autoantigen) (TLAA) (X-ray repair complementing defective repair in Chinese hamster cells 6) | XRCC6 G22P1 |
| sp\|P62888\|RL30_HUMAN | 1,934,855 | 1,609,227 | 1,969,496 | 1,524,667 | sp\|P62888\|RL30_HUMAN | 1,225,729 | 594,116 | 48% | P62888 | RL30_HUMAN | RL30_HUMAN | 60S ribosomal protein L30 (Large ribosomal subunit protein eL30) | RPL30 |
| sp\|P56537\|IF6_HUMAN | 324,524 | 365,643 | 37,143 | 106,570 | sp\|P56537\|IF6_HUMAN | 236,560 | 112,113 | 47% | P56537 | IF6_HUMAN | IF6_HUMAN | Eukaryotic translation initiation factor 6 (eIF-6) (B(2)GCN homolog) (B4 integrin interactor) (CAB) (p27(BBP)) | EIF6 EIF3A ITGB4BP OK/SW-cl.27 |
| sp\|Q92973\|TNPO1_HUMAN | 77,617 | 91,846 | 30,619 | 59,471 | sp\|Q92973\|TNPO1_HUMAN | 53,944 | 25,699 | 48% | Q92973 | TNPO1_HUMAN | TNPO1_HUMAN | Transportin-1 (Importin beta-2) (Karyopherin beta-2) (M9 region interaction protein) (MIP) | TNPO1 KPNB2 MIP1 TRN |

| | | | | | | | | | | | | | |
|---|---|---|---|---|---|---|---|---|---|---|---|---|---|
| sp\|Q99873\|ANM1_HUMAN | 364,687 | 368,414 | 335,921 | 373,465 | sp\|Q99873\|ANM1_HUMAN | 296,272 | 70,373 | 24% | Q99873 | ANM1_HUMAN | ANM1_HUMAN | Protein arginine N-methyltransferase 1 (EC 2.1.1.319) (Histone-arginine N-methyltransferase PRMT1) (Interferon receptor 1-bound protein 4) | PRMT1 HMT2 HRMT1L2 IR1B4 |
| sp\|P20645\|MPRD_HUMAN | 908,102 | 631,185 | 1,135,588 | 812,456 | sp\|P20645\|MPRD_HUMAN | 602,325 | 322,577 | 54% | P20645 | MPRD_HUMAN | MPRD_HUMAN | Cation-dependent mannose-6-phosphate receptor (CD Man-6-P receptor) (CD-MPR) (46 kDa mannose 6-phosphate receptor) (MPR 46) | M6PR MPR46 MPRD |
| sp\|Q9UNF1\|MAGD2_HUMAN | 586,143 | 665,081 | 458,818 | 641,782 | sp\|Q9UNF1\|MAGD2_HUMAN | 495,475 | 121,219 | 24% | Q9UNF1 | MAGD2_HUMAN | MAGD2_HUMAN | Melanoma-associated antigen D2 (11B6) (Breast cancer-associated gene 1 protein) (BCG-1) (Hepatocellular carcinoma-associated protein JCL-1) (MAGE-D2 antigen) | MAGED2 BCG1 |
| sp\|P07686\|HEXB_HUMAN | 209,380 | 228,262 | 184,599 | 153,494 | sp\|P07686\|HEXB_HUMAN | 140,672 | 62,758 | 45% | P07686 | HEXB_HUMAN | HEXB_HUMAN | Beta-hexosaminidase subunit beta (EC 3.2.1.52) (Beta-N-acetylhexosaminidase subunit beta) (Hexosaminidase subunit B) (Cervical cancer proto-oncogene 7 protein) (HCC-7) (N-acetyl-beta-glucosaminidase subunit beta) [Cleaved into: Beta-hexosaminidase subunit beta chain B; Beta-hexosaminidase subunit beta chain A] | HEXB HCC7 |
| sp\|P30084\|ECHM_HUMAN | 73,515 | 100,539 | 128,274 | 31,418 | sp\|P30084\|ECHM_HUMAN | 84,452 | 32,757 | 39% | P30084 | ECHM_HUMAN | ECHM_HUMAN | Enoyl-CoA hydratase, mitochondrial (mECH) (mECH1) (EC 4.2.1.17) (EC 5.3.3.8) (Enoyl-CoA hydratase 1) (ECHS1) (Short-chain enoyl-CoA hydratase) (SCEH) | ECHS1 |
| sp\|P62829\|RL23_HUMAN | 439,833 | 466,630 | 244,201 | 282,221 | sp\|P62829\|RL23_HUMAN | 268,715 | 134,953 | 50% | P62829 | RL23_HUMAN | RL23_HUMAN | 60S ribosomal protein L23 (60S ribosomal protein L17) (Large ribosomal subunit protein uL14) | RPL23 |
| sp\|P25788\|PSA3_HUMAN | 348,825 | 334,249 | 280,559 | 332,152 | sp\|P25788\|PSA3_HUMAN | 288,043 | 68,028 | 24% | P25788 | PSA3_HUMAN | PSA3_HUMAN | Proteasome subunit alpha type-3 (Macropain subunit C8) (Multicatalytic endopeptidase complex subunit C8) (Proteasome component C8) | PSMA3 HC8 PSC8 |
| sp\|P09471\|GNAO_HUMAN | 353,993 | 361,460 | 576,916 | 656,612 | sp\|P09471\|GNAO_HUMAN | 424,266 | 146,646 | 35% | P09471 | GNAO_HUMAN | GNAO_HUMAN | Guanine nucleotide-binding protein G(o) subunit alpha | GNAO1 |
| sp\|Q9UN37\|VPS4A_HUMAN | 788,666 | 830,385 | 959,436 | 1,161,015 | sp\|Q9UN37\|VPS4A_HUMAN | 813,719 | 225,673 | 28% | Q9UN37 | VPS4A_HUMAN | VPS4A_HUMAN | Vacuolar protein sorting-associated protein 4A (EC 3.6.4.6) (Protein SKD2) (VPS4-1) (hVPS4) | VPS4A VPS4 |
| sp\|O15498\|YKT6_HUMAN | 294,647 | 355,330 | 127,269 | 298,527 | sp\|O15498\|YKT6_HUMAN | 273,426 | 68,863 | 25% | O15498 | YKT6_HUMAN | YKT6_HUMAN | Synaptobrevin homolog YKT6 (EC 2.3.1.-) | YKT6 |
| sp\|O14980\|XPO1_HUMAN | 257,415 | 416,815 | 219,641 | 278,654 | sp\|O14980\|XPO1_HUMAN | 241,469 | 97,236 | 40% | O14980 | XPO1_HUMAN | XPO1_HUMAN | Exportin-1 (Exp1) (Chromosome region maintenance 1 protein homolog) | XPO1 CRM1 |
| sp\|P59998\|ARPC4_HUMAN | 358,085 | 459,860 | 525,286 | 755,357 | sp\|P59998\|ARPC4_HUMAN | 546,854 | 151,520 | 28% | P59998 | ARPC4_HUMAN | ARPC4_HUMAN | Actin-related protein 2/3 complex subunit 4 (Arp2/3 complex 20 kDa subunit) (p20-ARC) | ARPC4 ARC20 |

| | | | | | | | | | | | | | |
|---|---|---|---|---|---|---|---|---|---|---|---|---|---|
| sp\|P11766\|ADHX_HUMAN | 337,802 | 394,966 | 229,713 | 111,455 | sp\|P11766\|ADHX_HUMAN | 247,402 | 100,460 | 41% | P11766 | ADHX_HUMAN | ADHX_HUMAN | Alcohol dehydrogenase class-3 (EC 1.1.1.1) (Alcohol dehydrogenase 5) (Alcohol dehydrogenase class chi chain) (Alcohol dehydrogenase class-III) (Glutathione-dependent formaldehyde dehydrogenase) (FALDH) (FDH) (GSH-FDH) (EC 1.1.1.-) (S-(hydroxymethyl)glutathione dehydrogenase) (EC 1.1.1.284) | ADH5 ADHX FDH |
| sp\|P19961\|AMY2B_HUMAN | 55,570 | 30,039 | 53,900 | 62,228 | sp\|P19961\|AMY2B_HUMAN | 768,026 | 830,747 | 108% | P19961 | AMY2B_HUMAN | AMY2B_HUMAN | Alpha-amylase 2B (EC 3.2.1.1) (1,4-alpha-D-glucan glucanohydrolase 2B) (Carcinoid alpha-amylase) | AMY2B |
| sp\|Q07954\|LRP1_HUMAN | 79,640 | 79,313 | 58,132 | 72,860 | sp\|Q07954\|LRP1_HUMAN | 151,238 | 91,578 | 61% | Q07954 | LRP1_HUMAN | LRP1_HUMAN | Prolow-density lipoprotein receptor-related protein 1 (LRP-1) (Alpha-2-macroglobulin receptor) (A2MR) (Apolipoprotein E receptor) (APOER) (CD antigen CD91) [Cleaved into: Low-density lipoprotein receptor-related protein 1 85 kDa subunit (LRP-85); Low-density lipoprotein receptor-related protein 1 515 kDa subunit (LRP-515); Low-density lipoprotein receptor-related protein 1 intracellular domain (LRPICD)] | LRP1 A2MR APR |
| sp\|Q13310\|PABP4_HUMAN | 70,108 | 69,503 | 71,913 | 35,367 | sp\|Q13310\|PABP4_HUMAN | 62,501 | 17,847 | 29% | Q13310 | PABP4_HUMAN | PABP4_HUMAN | Polyadenylate-binding protein 4 (PABP-4) (Poly(A)-binding protein 4) (Activated-platelet protein 1) (APP-1) (Inducible poly(A)-binding protein) (iPABP) | PABPC4 APP1 PABP4 |
| sp\|P15170\|ERF3A_HUMAN | 53,732 | 45,264 | 8,709 | 83,346 | sp\|P15170\|ERF3A_HUMAN | 45,175 | 29,934 | 66% | P15170 | ERF3A_HUMAN | ERF3A_HUMAN | Eukaryotic peptide chain release factor GTP-binding subunit ERF3A (Eukaryotic peptide chain release factor subunit 3a) (eRF3a) (G1 to S phase transition protein 1 homolog) | GSPT1 ERF3A |
| sp\|O15031\|PLXB2_HUMAN | 535,874 | 546,923 | 506,826 | 670,737 | sp\|O15031\|PLXB2_HUMAN | 561,460 | 88,842 | 16% | O15031 | PLXB2_HUMAN | PLXB2_HUMAN | Plexin-B2 (MM1) | PLXNB2 KIAA0315 |
| sp\|P62195\|PRS8_HUMAN | 267,314 | 333,944 | 283,306 | 330,466 | sp\|P62195\|PRS8_HUMAN | 292,797 | 60,085 | 21% | P62195 | PRS8_HUMAN | PRS8_HUMAN | 26S proteasome regulatory subunit 8 (26S proteasome AAA-ATPase subunit RPT6) (Proteasome 26S subunit ATPase 5) (Proteasome subunit p45) (Thyroid hormone receptor-interacting protein 1) (TRIP1) (p45/SUG) | PSMC5 SUG1 |
| sp\|O15162\|PLS1_HUMAN | 540,143 | 601,018 | 843,174 | 795,046 | sp\|O15162\|PLS1_HUMAN | 602,678 | 228,181 | 38% | O15162 | PLS1_HUMAN | PLS1_HUMAN | Phospholipid scramblase 1 (PL scramblase 1) (Ca(2+)-dependent phospholipid scramblase 1) (Erythrocyte phospholipid scramblase) (Mg(2+)-dependent nuclease) (EC 3.1.-.-) (MmTRA1b) | PLSCR1 |
| sp\|Q86VE9\|SERC5_HUMAN | 3,283,282 | 2,885,096 | 3,664,409 | 3,455,426 | sp\|Q86VE9\|SERC5_HUMAN | 3,060,245 | 479,726 | 16% | Q86VE9 | SERC5_HUMAN | SERC5_HUMAN | Serine incorporator 5 | SERINC5 C5orf12 |
| sp\|O00625\|PIR_HUMAN | 225,921 | 363,356 | 190,958 | 254,821 | sp\|O00625\|PIR_HUMAN | 234,717 | 74,531 | 32% | O00625 | PIR_HUMAN | PIR_HUMAN | Pirin (EC 1.13.11.24) (Probable quercetin 2,3-dioxygenase PIR) (Probable quercetinase) | PIR |
| sp\|P07910\|HNRPC_HUMAN | 433,863 | 347,345 | 415,499 | 276,771 | sp\|P07910\|HNRPC_HUMAN | 239,628 | 147,833 | 62% | P07910 | HNRPC_HUMAN | HNRPC_HUMAN | Heterogeneous nuclear ribonucleoproteins C1/C2 (hnRNP C1/C2) | HNRNPC HNRPC |

| | | | | | | | | | | | | | |
|---|---|---|---|---|---|---|---|---|---|---|---|---|---|
| sp\|Q9Y653\|AGRG1_HUMAN | 476,063 | 565,342 | 318,092 | 456,532 | sp\|Q9Y653\|AGRG1_HUMAN | 417,539 | 83,105 | 20% | Q9Y653 | AGRG1_HUMAN | AGRG1_HUMAN | Adhesion G-protein coupled receptor G1 (G-protein coupled receptor 56) (Protein TM7XN1) [Cleaved into: ADGRG1 N-terminal fragment (ADGRG1 NT) (GPR56 N-terminal fragment) (GPR56 NT) (GPR56(N)) (GPR56 extracellular subunit) (GPR56 subunit alpha); ADGRG1 C-terminal fragment (ADGRG1 CT) (GPR56 C-terminal fragment) (GPR56 CT) (GPR56(C)) (GPR56 seven-transmembrane subunit) (GPR56 7TM) (GPR56 subunit beta)] | ADGRG1 GPR56 TM7LN4 TM7XN1 UNQ540/PRO1083 |
| sp\|Q9C0H2\|TTYH3_HUMAN | 526,563 | 600,403 | 549,917 | 501,335 | sp\|Q9C0H2\|TTYH3_HUMAN | 465,188 | 176,293 | 38% | Q9C0H2 | TTYH3_HUMAN | TTYH3_HUMAN | Protein tweety homolog 3 (hTTY3) | TTYH3 KIAA1691 |
| sp\|Q9H5V8\|CDCP1_HUMAN | 531,967 | 522,419 | 444,895 | 370,442 | sp\|Q9H5V8\|CDCP1_HUMAN | 450,430 | 60,938 | 14% | Q9H5V8 | CDCP1_HUMAN | CDCP1_HUMAN | CUB domain-containing protein 1 (Membrane glycoprotein gp140) (Subtractive immunization M plus HEp3-associated 135 kDa protein) (SIMA135) (Transmembrane and associated with src kinases) (CD antigen CD318) | CDCP1 TRASK UNQ2486/PRO5773 |
| sp\|P30043\|BLVRB_HUMAN | 297,925 | 363,303 | 365,953 | 415,907 | sp\|P30043\|BLVRB_HUMAN | 334,130 | 48,528 | 15% | P30043 | BLVRB_HUMAN | BLVRB_HUMAN | Flavin reductase (NADPH) (FR) (EC 1.5.1.30) (Biliverdin reductase B) (BVR-B) (EC 1.3.1.24) (Biliverdin-IX beta-reductase) (Green heme-binding protein) (GHBP) (NADPH-dependent diaphorase) (NADPH-flavin reductase) (FLR) | BLVRB FLR |
| sp\|P31946\|1433B_HUMAN | 724,366 | 1,035,104 | 427,809 | 490,778 | sp\|P31946\|1433B_HUMAN | 629,571 | 262,842 | 42% | P31946 | 1433B_HUMAN | 1433B_HUMAN | 14-3-3 protein beta/alpha (Protein 1054) (Protein kinase C inhibitor protein 1) (KCIP-1) [Cleaved into: 14-3-3 protein beta/alpha, N-terminally processed] | YWHAB |
| sp\|P61020\|RAB5B_HUMAN | 274,283 | 308,913 | 196,079 | 265,808 | sp\|P61020\|RAB5B_HUMAN | 228,686 | 72,447 | 32% | P61020 | RAB5B_HUMAN | RAB5B_HUMAN | Ras-related protein Rab-5B (EC 3.6.5.2) | RAB5B |
| sp\|Q13765\|NACA_HUMAN | 104,724 | 108,941 | 86,620 | 81,220 | sp\|Q13765\|NACA_HUMAN | 103,006 | 25,461 | 25% | Q13765 | NACA_HUMAN | NACA_HUMAN | Nascent polypeptide-associated complex subunit alpha (NAC-alpha) (Alpha-NAC) (allergen Hom s 2) | NACA HSD48 |
| sp\|Q07812\|BAX_HUMAN | 227,360 | 287,227 | 262,860 | 391,233 | sp\|Q07812\|BAX_HUMAN | 255,116 | 78,175 | 31% | Q07812 | BAX_HUMAN | BAX_HUMAN | Apoptosis regulator BAX (Bcl-2-like protein 4) (Bcl2-L-4) | BAX BCL2L4 |
| sp\|Q07021\|C1QBP_HUMAN | 82,755 | 114,813 | 65,133 | 108,467 | sp\|Q07021\|C1QBP_HUMAN | 85,306 | 42,972 | 50% | Q07021 | C1QBP_HUMAN | C1QBP_HUMAN | Complement component 1 Q subcomponent-binding protein, mitochondrial (ASF/SF2-associated protein p32) (Glycoprotein gC1qBP) (C1qBP) (Hyaluronan-binding protein 1) (Mitochondrial matrix protein p32) (gC1q-R protein) (p33) (SF2AP32) | C1QBP GC1QBP HABP1 SF2P32 |
| sp\|P51571\|SSRD_HUMAN | 30,040 | 35,331 | 6,579 | 32,803 | sp\|P51571\|SSRD_HUMAN | 22,723 | 12,014 | 53% | P51571 | SSRD_HUMAN | SSRD_HUMAN | Translocon-associated protein subunit delta (TRAP-delta) (Signal sequence receptor subunit delta) (SSR-delta) | SSR4 TRAPD |

| | | | | | | | | | | | | | |
|---|---|---|---|---|---|---|---|---|---|---|---|---|---|
| sp\|P30825\|CTR1_HUMAN | 452,895 | 595,492 | 196,323 | 392,770 | sp\|P30825\|CTR1_HUMAN | 390,602 | 146,678 | 38% | P30825 | CTR1_HUMAN | SL7A1_HUMAN | High affinity cationic amino acid transporter 1 (CAT-1) (CAT1) (Ecotropic retroviral leukemia receptor homolog) (Ecotropic retrovirus receptor homolog) (Solute carrier family 7 member 1) (System Y+ basic amino acid transporter) | SLC7A1 ATRC1 ERR REC1L |
| sp\|P28066\|PSA5_HUMAN | 419,376 | 443,097 | 442,457 | 524,075 | sp\|P28066\|PSA5_HUMAN | 430,559 | 62,831 | 15% | P28066 | PSA5_HUMAN | PSA5_HUMAN | Proteasome subunit alpha type-5 (Macropain zeta chain) (Multicatalytic endopeptidase complex zeta chain) (Proteasome zeta chain) | PSMA5 |
| sp\|P19623\|SPEE_HUMAN | 1,056,658 | 1,068,921 | 1,041,216 | 1,243,865 | sp\|P19623\|SPEE_HUMAN | 985,946 | 189,215 | 19% | P19623 | SPEE_HUMAN | SPEE_HUMAN | Spermidine synthase (SPDSY) (EC 2.5.1.16) (Putrescine aminopropyltransferase) | SRM SPS1 SRML1 |
| sp\|P30520\|PURA2_HUMAN | 183,164 | 189,005 | 125,762 | 163,644 | sp\|P30520\|PURA2_HUMAN | 174,219 | 49,485 | 28% | P30520 | PURA2_HUMAN | PURA2_HUMAN | Adenylosuccinate synthetase isozyme 2 (AMPSase 2) (AdSS 2) (EC 6.3.4.4) (Adenylosuccinate synthetase, acidic isozyme) (Adenylosuccinate synthetase, liver isozyme) (L-type adenylosuccinate synthetase) (IMP--aspartate ligase 2) | ADSS2 ADSS |
| sp\|Q9Y3F4\|STRAP_HUMAN | 157,923 | 207,414 | 141,130 | 139,961 | sp\|Q9Y3F4\|STRAP_HUMAN | 162,316 | 40,392 | 25% | Q9Y3F4 | STRAP_HUMAN | STRAP_HUMAN | Serine-threonine kinase receptor-associated protein (MAP activator with WD repeats) (UNR-interacting protein) (WD-40 repeat protein PT-WD) | STRAP MAWD UNRIP |
| sp\|O14495\|PLPP3_HUMAN | 979,226 | 994,801 | 740,421 | 756,491 | sp\|O14495\|PLPP3_HUMAN | 760,168 | 165,907 | 22% | O14495 | PLPP3_HUMAN | PLPP3_HUMAN | Phospholipid phosphatase 3 (EC 3.1.3.-) (EC 3.1.3.4) (Lipid phosphate phosphohydrolase 3) (PAP2-beta) (Phosphatidate phosphohydrolase type 2b) (Phosphatidic acid phosphatase 2b) (PAP-2b) (PAP2b) (Vascular endothelial growth factor and type I collagen-inducible protein) (VCIP) | PLPP3 LPP3 PPAP2B |
| sp\|P50502\|F10A1_HUMAN | 453,598 | 481,210 | 416,354 | 403,968 | sp\|P50502\|F10A1_HUMAN | 390,035 | 106,027 | 27% | P50502 | F10A1_HUMAN | F10A1_HUMAN | Hsc70-interacting protein (Hip) (Aging-associated protein 2) (Progesterone receptor-associated p48 protein) (Protein FAM10A1) (Putative tumor suppressor ST13) (Renal carcinoma antigen NY-REN-33) (Suppression of tumorigenicity 13 protein) | ST13 AAG2 FAM10A1 HIP SNC6 |
| sp\|Q9NVA2\|SEP11_HUMAN | 46,828 | 99,849 | 37,873 | 72,230 | sp\|Q9NVA2\|SEP11_HUMAN | 60,813 | 24,161 | 40% | Q9NVA2 | SEP11_HUMAN | SEP11_HUMAN | Septin-11 | SEPTIN11 SEPT11 |
| sp\|O43747\|AP1G1_HUMAN | 119,684 | 179,279 | 297,459 | 386,005 | sp\|O43747\|AP1G1_HUMAN | 239,292 | 87,391 | 37% | O43747 | AP1G1_HUMAN | AP1G1_HUMAN | AP-1 complex subunit gamma-1 (Adaptor protein complex AP-1 subunit gamma-1) (Adaptor-related protein complex 1 subunit gamma-1) (Clathrin assembly protein complex 1 gamma-1 large chain) (Gamma1-adaptin) (Golgi adaptor HA1/AP1 adaptin subunit gamma-1) | AP1G1 ADTG CLAPG1 |
| sp\|P29992\|GNA11_HUMAN | 154,346 | 318,413 | 187,093 | 291,911 | sp\|P29992\|GNA11_HUMAN | 256,657 | 62,682 | 24% | P29992 | GNA11_HUMAN | GNA11_HUMAN | Guanine nucleotide-binding protein subunit alpha-11 (G alpha-11) (G-protein subunit alpha-11) (Guanine nucleotide-binding protein G(y) subunit alpha) | GNA11 GA11 |

| | | | | | | | | | | | | | |
|---|---|---|---|---|---|---|---|---|---|---|---|---|---|
| sp\|Q8TEX9\|IPO4_HUMAN | 205,670 | 192,367 | 225,644 | 310,704 | sp\|Q8TEX9\|IPO4_HUMAN | 219,944 | 57,268 | 26% | Q8TEX9 | IPO4_HUMAN | IPO4_HUMAN | Importin-4 (Imp4) (Importin-4b) (Imp4b) (Ran-binding protein 4) (RanBP4) | IPO4 IMP4B RANBP4 |
| sp\|P30533\|AMRP_HUMAN | 13,323 | 26,703 | 91,531 | 77,036 | sp\|P30533\|AMRP_HUMAN | 39,348 | 30,935 | 79% | P30533 | AMRP_HUMAN | AMRP_HUMAN | Alpha-2-macroglobulin receptor-associated protein (Alpha-2-MRAP) (Low density lipoprotein receptor-related protein-associated protein 1) (RAP) | LRPAP1 A2MRAP |
| sp\|P17174\|AATC_HUMAN | 175,770 | 160,046 | 159,061 | 146,577 | sp\|P17174\|AATC_HUMAN | 169,811 | 31,225 | 18% | P17174 | AATC_HUMAN | AATC_HUMAN | Aspartate aminotransferase, cytoplasmic (cAspAT) (EC 2.6.1.1) (EC 2.6.1.3) (Cysteine aminotransferase, cytoplasmic) (Cysteine transaminase, cytoplasmic) (cCAT) (Glutamate oxaloacetate transaminase 1) (Transaminase A) | GOT1 |
| sp\|Q9Y3A5\|SBDS_HUMAN | 118,889 | 41,192 | 42,995 | 97,625 | sp\|Q9Y3A5\|SBDS_HUMAN | 86,634 | 38,659 | 45% | Q9Y3A5 | SBDS_HUMAN | SBDS_HUMAN | Ribosome maturation protein SBDS (Shwachman-Bodian-Diamond syndrome protein) | SBDS CGI-97 |
| sp\|P0DP25\|CALM3_HUMAN | 610,190 | 723,970 | 1,204,563 | 1,560,363 | sp\|P0DP25\|CALM3_HUMAN | 1,021,237 | 326,237 | 32% | P0DP25 | CALM3_HUMAN | CALM3_HUMAN | Calmodulin-3 | CALM3 CALML2 CAM3 CAMC CAMIII |
| sp\|P61254\|RL26_HUMAN | 1,027,889 | 844,263 | 473,738 | 790,739 | sp\|P61254\|RL26_HUMAN | 613,666 | 243,437 | 40% | P61254 | RL26_HUMAN | RL26_HUMAN | 60S ribosomal protein L26 (Large ribosomal subunit protein uL24) | RPL26 |
| sp\|P47755\|CAZA2_HUMAN | 224,501 | 369,837 | 427,921 | 600,631 | sp\|P47755\|CAZA2_HUMAN | 417,987 | 126,012 | 30% | P47755 | CAZA2_HUMAN | CAZA2_HUMAN | F-actin-capping protein subunit alpha-2 (CapZ alpha-2) | CAPZA2 |
| sp\|P53990\|IST1_HUMAN | 1,412,282 | 1,645,040 | 1,109,393 | 1,176,005 | sp\|P53990\|IST1_HUMAN | 1,159,496 | 278,905 | 24% | P53990 | IST1_HUMAN | IST1_HUMAN | IST1 homolog (hIST1) (Charged multivesicular body protein 8) (CHMP8) (Putative MAPK-activating protein PM28) | IST1 KIAA0174 |
| sp\|Q8NBJ7\|SUMF2_HUMAN | 114,471 | 103,882 | 85,889 | 86,416 | sp\|Q8NBJ7\|SUMF2_HUMAN | 88,536 | 22,823 | 26% | Q8NBJ7 | SUMF2_HUMAN | SUMF2_HUMAN | Inactive C-alpha-formylglycine-generating enzyme 2 (Paralog of formylglycine-generating enzyme) (pFGE) (Sulfatase-modifying factor 2) | SUMF2 PSEC0171 UNQ1968/PRO4500 |
| sp\|P40261\|NNMT_HUMAN | 60,482 | 116,501 | 90,311 | 46,942 | sp\|P40261\|NNMT_HUMAN | 81,900 | 21,976 | 27% | P40261 | NNMT_HUMAN | NNMT_HUMAN | Nicotinamide N-methyltransferase (EC 2.1.1.1) | NNMT |
| sp\|P50443\|S26A2_HUMAN | 849,285 | 878,558 | 539,346 | 836,823 | sp\|P50443\|S26A2_HUMAN | 723,361 | 175,021 | 24% | P50443 | S26A2_HUMAN | S26A2_HUMAN | Sulfate transporter (Diastrophic dysplasia protein) (Solute carrier family 26 member 2) | SLC26A2 DTD DTDST |
| sp\|P13693\|TCTP_HUMAN | 407,336 | 501,444 | 358,801 | 273,699 | sp\|P13693\|TCTP_HUMAN | 359,160 | 105,344 | 29% | P13693 | TCTP_HUMAN | TCTP_HUMAN | Translationally-controlled tumor protein (TCTP) (Fortilin) (Histamine-releasing factor) (HRF) (p23) | TPT1 |
| sp\|P30085\|KCY_HUMAN | 142,505 | 150,167 | 51,746 | 93,408 | sp\|P30085\|KCY_HUMAN | 106,358 | 31,526 | 30% | P30085 | KCY_HUMAN | KCY_HUMAN | UMP-CMP kinase (EC 2.7.4.14) (Deoxycytidylate kinase) (CK) (dCMP kinase) (Nucleoside-diphosphate kinase) (EC 2.7.4.6) (Uridine monophosphate/cytidine monophosphate kinase) (UMP/CMP kinase) (UMP/CMPK) | CMPK1 CMK CMPK UCK UMK UMPK |
| sp\|Q9NP72\|RAB18_HUMAN | 155,891 | 141,097 | 316,618 | 291,564 | sp\|Q9NP72\|RAB18_HUMAN | 205,614 | 72,629 | 35% | Q9NP72 | RAB18_HUMAN | RAB18_HUMAN | Ras-related protein Rab-18 | RAB18 |
| sp\|Q96PU5\|NED4L_HUMAN | 239,774 | 368,556 | 458,066 | 629,950 | sp\|Q96PU5\|NED4L_HUMAN | 439,807 | 169,574 | 39% | Q96PU5 | NED4L_HUMAN | NED4L_HUMAN | E3 ubiquitin-protein ligase NEDD4-like (EC 2.3.2.26) (EC 2.3.2.36) (HECT-type E3 ubiquitin transferase NED4L) (NEDD4.2) (Nedd4-2) | NEDD4L KIAA0439 NEDL3 |
| sp\|Q02978\|M2OM_HUMAN | 18,356 | 26,158 | 2,193 | 13,274 | sp\|Q02978\|M2OM_HUMAN | 12,058 | 8,281 | 69% | Q02978 | M2OM_HUMAN | M2OM_HUMAN | Mitochondrial 2-oxoglutarate/malate carrier protein (OGCP) (Solute carrier family 25 member 11) | SLC25A11 SLC20A4 |
| sp\|Q9BWM7\|SFXN3_HUMAN | 88,303 | 56,186 | 297,012 | 172,441 | sp\|Q9BWM7\|SFXN3_HUMAN | 178,848 | 113,493 | 63% | Q9BWM7 | SFXN3_HUMAN | SFXN3_HUMAN | Sideroflexin-3 | SFXN3 |
| sp\|Q13045\|FLII_HUMAN | 168,904 | 167,876 | 195,132 | 298,119 | sp\|Q13045\|FLII_HUMAN | 270,758 | 110,090 | 41% | Q13045 | FLII_HUMAN | FLII_HUMAN | Protein flightless-1 homolog | FLII FLIL |

| | | | | | | | | | | | | | |
|---|---|---|---|---|---|---|---|---|---|---|---|---|---|
| sp\|Q16555\|DPYL2_HUMAN | 130,231 | 143,955 | 195,155 | 142,190 | sp\|Q16555\|DPYL2_HUMAN | 163,911 | 26,794 | 16% | Q16555 | DPYL2_HUMAN | DPYL2_HUMAN | Dihydropyrimidinase-related protein 2 (DRP-2) (Collapsin response mediator protein 2) (CRMP-2) (N2A3) (Unc-33-like phosphoprotein 2) (ULIP-2) | DPYSL2 CRMP2 ULIP2 |
| sp\|P00403\|COX2_HUMAN | 113,562 | 111,832 | 142,789 | 136,931 | sp\|P00403\|COX2_HUMAN | 131,313 | 57,036 | 43% | P00403 | COX2_HUMAN | COX2_HUMAN | Cytochrome c oxidase subunit 2 (EC 7.1.1.9) (Cytochrome c oxidase polypeptide II) | MT-CO2 COII COX2 COXII MTCO2 |
| sp\|Q15293\|RCN1_HUMAN | 9,489 | 9,880 | 6,060 | 5,431 | sp\|Q15293\|RCN1_HUMAN | 10,240 | 6,051 | 59% | Q15293 | RCN1_HUMAN | RCN1_HUMAN | Reticulocalbin-1 | RCN1 RCN |
| sp\|P84077\|ARF1_HUMAN | 1,009,971 | 1,279,273 | 332,634 | 372,845 | sp\|P84077\|ARF1_HUMAN | 630,335 | 451,822 | 72% | P84077 | ARF1_HUMAN | ARF1_HUMAN | ADP-ribosylation factor 1 | ARF1 |
| sp\|P51572\|BAP31_HUMAN | 35,739 | 27,035 | 27,285 | 9,359 | sp\|P51572\|BAP31_HUMAN | 27,436 | 13,228 | 48% | P51572 | BAP31_HUMAN | BAP31_HUMAN | B-cell receptor-associated protein 31 (BCR-associated protein 31) (Bap31) (6C6-AG tumor-associated antigen) (Protein CDM) (p28) | BCAP31 BAP31 DXS1357E |
| sp\|Q6NUK1\|SCMC1_HUMAN | 112,073 | 43,876 | 202,395 | 232,705 | sp\|Q6NUK1\|SCMC1_HUMAN | 145,872 | 73,255 | 50% | Q6NUK1 | SCMC1_HUMAN | SCMC1_HUMAN | Calcium-binding mitochondrial carrier protein SCaMC-1 (Mitochondrial ATP-Mg/Pi carrier protein 1) (Mitochondrial Ca(2+)-dependent solute carrier protein 1) (Small calcium-binding mitochondrial carrier protein 1) (Solute carrier family 25 member 24) | SLC25A24 APC1 MCSC1 SCAMC1 |
| sp\|P01008\|ANT3_HUMAN | 76,813 | 111,928 | 125,703 | 103,981 | sp\|P01008\|ANT3_HUMAN | 1,581,283 | 1,582,223 | 100% | P01008 | ANT3_HUMAN | ANT3_HUMAN | Antithrombin-III (ATIII) (Serpin C1) | SERPINC1 AT3 PRO0309 |
| sp\|Q15181\|IPYR_HUMAN | 109,340 | 324,592 | 266,522 | 299,236 | sp\|Q15181\|IPYR_HUMAN | 241,113 | 66,167 | 27% | Q15181 | IPYR_HUMAN | IPYR_HUMAN | Inorganic pyrophosphatase (EC 3.6.1.1) (Pyrophosphate phospho-hydrolase) (PPase) | PPA1 IOPPP PP |
| sp\|P53794\|SC5A3_HUMAN | 515,797 | 489,963 | 356,430 | 544,654 | sp\|P53794\|SC5A3_HUMAN | 446,401 | 66,472 | 15% | P53794 | SC5A3_HUMAN | SC5A3_HUMAN | Sodium/myo-inositol cotransporter (Na(+)/myo-inositol cotransporter) (Sodium/myo-inositol transporter 1) (SMIT1) (Solute carrier family 5 member 3) | SLC5A3 |
| sp\|P61619\|S61A1_HUMAN | 54,024 | 95,253 | 89,619 | 70,828 | sp\|P61619\|S61A1_HUMAN | 76,631 | 18,207 | 24% | P61619 | S61A1_HUMAN | S61A1_HUMAN | Protein transport protein Sec61 subunit alpha isoform 1 (Sec61 alpha-1) | SEC61A1 SEC61A |
| sp\|P49207\|RL34_HUMAN | 1,006,657 | 941,335 | 817,774 | 888,454 | sp\|P49207\|RL34_HUMAN | 607,138 | 333,794 | 55% | P49207 | RL34_HUMAN | RL34_HUMAN | 60S ribosomal protein L34 (Large ribosomal subunit protein eL34) | RPL34 |
| sp\|P53007\|TXTP_HUMAN | 51,835 | 64,727 | 54,242 | 125,070 | sp\|P53007\|TXTP_HUMAN | 79,129 | 34,673 | 44% | P53007 | TXTP_HUMAN | TXTP_HUMAN | Tricarboxylate transport protein, mitochondrial (Citrate transport protein) (CTP) (Solute carrier family 25 member 1) (Tricarboxylate carrier protein) | SLC25A1 SLC20A3 |
| sp\|O95433\|AHSA1_HUMAN | 137,913 | 134,766 | 225,659 | 215,824 | sp\|O95433\|AHSA1_HUMAN | 175,682 | 40,589 | 23% | O95433 | AHSA1_HUMAN | AHSA1_HUMAN | Activator of 90 kDa heat shock protein ATPase homolog 1 (AHA1) (p38) | AHSA1 C14orf3 HSPC322 |
| sp\|Q8WV92\|MITD1_HUMAN | 430,551 | 495,747 | 494,371 | 641,588 | sp\|Q8WV92\|MITD1_HUMAN | 510,472 | 95,004 | 19% | Q8WV92 | MITD1_HUMAN | MITD1_HUMAN | MIT domain-containing protein 1 | MITD1 |
| sp\|Q9BS26\|ERP44_HUMAN | 116,071 | 84,790 | 146,423 | 139,266 | sp\|Q9BS26\|ERP44_HUMAN | 95,138 | 37,818 | 40% | Q9BS26 | ERP44_HUMAN | ERP44_HUMAN | Endoplasmic reticulum resident protein 44 (ER protein 44) (ERp44) (Thioredoxin domain-containing protein 4) | ERP44 KIAA0573 TXNDC4 UNQ532/PRO1075 |

| | | | | | | | | | | | | | |
|---|---|---|---|---|---|---|---|---|---|---|---|---|---|
| sp\|P11717\|MPRI_HUMAN | 306,485 | 229,393 | 246,137 | 171,991 | sp\|P11717\|MPRI_HUMAN | 289,700 | 67,451 | 23% | P11717 | MPRI_HUMAN | MPRI_HUMAN | Cation-independent mannose-6-phosphate receptor (CI Man-6-P receptor) (CI-MPR) (M6PR) (300 kDa mannose 6-phosphate receptor) (MPR 300) (Insulin-like growth factor 2 receptor) (Insulin-like growth factor II receptor) (IGF-II receptor) (M6P/IGF2 receptor) (M6P/IGF2R) (CD antigen CD222) | IGF2R MPRI |
| sp\|Q03135\|CAV1_HUMAN | 207,554 | 172,479 | 71,380 | 239,114 | sp\|Q03135\|CAV1_HUMAN | 157,960 | 60,236 | 38% | Q03135 | CAV1_HUMAN | CAV1_HUMAN | Caveolin-1 | CAV1 CAV |
| sp\|P57088\|TMM33_HUMAN | 151,767 | 89,055 | 106,270 | 277,153 | sp\|P57088\|TMM33_HUMAN | 166,563 | 80,408 | 48% | P57088 | TMM33_HUMAN | TMM33_HUMAN | Transmembrane protein 33 (Protein DB83) (SHINC-3) | TMEM33 DB83 |
| sp\|O00186\|STXB3_HUMAN | 415,362 | 556,061 | 438,483 | 469,206 | sp\|O00186\|STXB3_HUMAN | 420,967 | 69,918 | 17% | O00186 | STXB3_HUMAN | STXB3_HUMAN | Syntaxin-binding protein 3 (Platelet Sec1 protein) (PSP) (Protein unc-18 homolog 3) (Unc18-3) (Protein unc-18 homolog C) (Unc-18C) | STXBP3 |
| sp\|Q04637\|IF4G1_HUMAN | 3,343 | 6,829 | 12,433 | 9,879 | sp\|Q04637\|IF4G1_HUMAN | 8,908 | 3,897 | 44% | Q04637 | IF4G1_HUMAN | IF4G1_HUMAN | Eukaryotic translation initiation factor 4 gamma 1 (eIF-4-gamma 1) (eIF-4G 1) (eIF-4G1) (p220) | EIF4G1 EIF4F EIF4G EIF4GI |
| sp\|P42765\|THIM_HUMAN | 80,037 | 90,484 | 49,755 | 102,930 | sp\|P42765\|THIM_HUMAN | 82,899 | 24,680 | 30% | P42765 | THIM_HUMAN | THIM_HUMAN | 3-ketoacyl-CoA thiolase, mitochondrial (EC 2.3.1.16) (Acetyl-CoA acetyltransferase) (EC 2.3.1.9) (Acetyl-CoA acyltransferase) (Acyl-CoA hydrolase, mitochondrial) (EC 3.1.2.-) (EC 3.1.2.1) (EC 3.1.2.2) (Beta-ketothiolase) (Mitochondrial 3-oxoacyl-CoA thiolase) (T1) | ACAA2 |
| sp\|O75821\|EIF3G_HUMAN | 71,448 | 67,087 | 72,413 | 106,550 | sp\|O75821\|EIF3G_HUMAN | 81,863 | 22,932 | 28% | O75821 | EIF3G_HUMAN | EIF3G_HUMAN | Eukaryotic translation initiation factor 3 subunit G (eIF3g) (Eukaryotic translation initiation factor 3 RNA-binding subunit) (eIF-3 RNA-binding subunit) (Eukaryotic translation initiation factor 3 subunit 4) (eIF-3-delta) (eIF3 p42) (eIF3 p44) | EIF3G EIF3S4 |
| sp\|P55209\|NP1L1_HUMAN | 105,002 | 153,739 | 378,292 | 412,206 | sp\|P55209\|NP1L1_HUMAN | 301,404 | 125,441 | 42% | P55209 | NP1L1_HUMAN | NP1L1_HUMAN | Nucleosome assembly protein 1-like 1 (NAP-1-related protein) (hNRP) | NAP1L1 NRP |
| sp\|Q16666\|IF16_HUMAN | 11,723 | 13,899 | 18,787 | 2,552 | sp\|Q16666\|IF16_HUMAN | 9,269 | 5,507 | 59% | Q16666 | IF16_HUMAN | IF16_HUMAN | Gamma-interferon-inducible protein 16 (Ifi-16) (Interferon-inducible myeloid differentiation transcriptional activator) | IFI16 IFNGIP1 |
| sp\|O00468\|AGRIN_HUMAN | 869,014 | 688,410 | 1,180,699 | 1,074,148 | sp\|O00468\|AGRIN_HUMAN | 786,653 | 239,607 | 30% | O00468 | AGRIN_HUMAN | AGRIN_HUMAN | Agrin [Cleaved into: Agrin N-terminal 110 kDa subunit; Agrin C-terminal 110 kDa subunit; Agrin C-terminal 90 kDa fragment (C90); Agrin C-terminal 22 kDa fragment (C22)] | AGRN AGRIN |
| sp\|O00231\|PSD11_HUMAN | 253,138 | 254,751 | 170,930 | 163,264 | sp\|O00231\|PSD11_HUMAN | 173,271 | 61,935 | 36% | O00231 | PSD11_HUMAN | PSD11_HUMAN | 26S proteasome non-ATPase regulatory subunit 11 (26S proteasome regulatory subunit RPN6) (26S proteasome regulatory subunit S9) (26S proteasome regulatory subunit p44.5) | PSMD11 |
| sp\|P10599\|THIO_HUMAN | 467,428 | 466,829 | 364,264 | 349,885 | sp\|P10599\|THIO_HUMAN | 393,271 | 100,756 | 26% | P10599 | THIO_HUMAN | THIO_HUMAN | Thioredoxin (Trx) (ATL-derived factor) (ADF) (Surface-associated sulphydryl protein) (SASP) (allergen Hom s Trx) | TXN TRDX TRX TRX1 |

| | | | | | | | | | | | | | |
|---|---|---|---|---|---|---|---|---|---|---|---|---|---|
| sp\|P48509\|CD151_HUMAN | 1,782,712 | 2,067,381 | 1,424,956 | 1,747,769 | sp\|P48509\|CD151_HUMAN | 1,319,136 | 659,120 | 50% | P48509 | CD151_HUMAN | CD151_HUMAN | CD151 antigen (GP27) (Membrane glycoprotein SFA-1) (Platelet-endothelial tetraspan antigen 3) (PETA-3) (Tetraspanin-24) (Tspan-24) (CD antigen CD151) | CD151 TSPAN24 |
| sp\|Q8TC12\|RDH11_HUMAN | 20,897 | 21,011 | 13,343 | 15,983 | sp\|Q8TC12\|RDH11_HUMAN | 14,726 | 5,468 | 37% | Q8TC12 | RDH11_HUMAN | RDH11_HUMAN | Retinol dehydrogenase 11 (EC 1.1.1.300) (Androgen-regulated short-chain dehydrogenase/reductase 1) (HCV core-binding protein HCBP12) (Prostate short-chain dehydrogenase/reductase 1) (Retinal reductase 1) (RalR1) (Short chain dehydrogenase/reductase family 7C member 1) | RDH11 ARSDR1 PSDR1 SDR7C1 CGI-82 |
| sp\|P62266\|RS23_HUMAN | 526,807 | 588,850 | 388,816 | 315,773 | sp\|P62266\|RS23_HUMAN | 326,597 | 166,912 | 51% | P62266 | RS23_HUMAN | RS23_HUMAN | 40S ribosomal protein S23 (Small ribosomal subunit protein uS12) | RPS23 |
| sp\|P36955\|PEDF_HUMAN | 150,518 | 102,980 | 85,856 | 133,255 | sp\|P36955\|PEDF_HUMAN | 1,704,930 | 1,743,661 | 102% | P36955 | PEDF_HUMAN | PEDF_HUMAN | Pigment epithelium-derived factor (PEDF) (Cell proliferation-inducing gene 35 protein) (EPC-1) (Serpin F1) | SERPINF1 PEDF PIG35 |
| sp\|P00492\|HPRT_HUMAN | 102,183 | 226,826 | 92,988 | 79,135 | sp\|P00492\|HPRT_HUMAN | 106,997 | 69,949 | 65% | P00492 | HPRT_HUMAN | HPRT_HUMAN | Hypoxanthine-guanine phosphoribosyltransferase (HGPRT) (HGPRTase) (EC 2.4.2.8) | HPRT1 HPRT |
| sp\|P46926\|GNPI1_HUMAN | 129,915 | 209,337 | 105,386 | 133,256 | sp\|P46926\|GNPI1_HUMAN | 125,212 | 39,095 | 31% | P46926 | GNPI1_HUMAN | GNPI1_HUMAN | Glucosamine-6-phosphate isomerase 1 (EC 3.5.99.6) (Glucosamine-6-phosphate deaminase 1) (GNPDA 1) (GlcN6P deaminase 1) (Oscillin) | GNPDA1 GNPI HLN KIAA0060 |
| sp\|Q13510\|ASAH1_HUMAN | 133,776 | 148,912 | 191,478 | 193,129 | sp\|Q13510\|ASAH1_HUMAN | 133,206 | 46,104 | 35% | Q13510 | ASAH1_HUMAN | ASAH1_HUMAN | Acid ceramidase (AC) (ACDase) (Acid CDase) (EC 3.5.1.23) (Acylsphingosine deacylase) (N-acylethanolamine hydrolase ASAH1) (EC 3.5.1.-) (N-acylsphingosine amidohydrolase) (Putative 32 kDa heart protein) (PHP32) [Cleaved into: Acid ceramidase subunit alpha; Acid ceramidase subunit beta] | ASAH1 ASAH HSD-33 HSD33 |
| sp\|Q13509\|TBB3_HUMAN | 275,036 | 344,280 | 95,832 | 202,569 | sp\|Q13509\|TBB3_HUMAN | 178,759 | 95,855 | 54% | Q13509 | TBB3_HUMAN | TBB3_HUMAN | Tubulin beta-3 chain (Tubulin beta-4 chain) (Tubulin beta-III) | TUBB3 TUBB4 |
| sp\|Q9Y376\|CAB39_HUMAN | 275,072 | 482,465 | 332,944 | 538,224 | sp\|Q9Y376\|CAB39_HUMAN | 431,852 | 127,101 | 29% | Q9Y376 | CAB39_HUMAN | CAB39_HUMAN | Calcium-binding protein 39 (MO25alpha) (Protein Mo25) | CAB39 MO25 CGI-66 |
| sp\|Q9H0X4\|F234A_HUMAN | 123,816 | 126,401 | 181,951 | 226,836 | sp\|Q9H0X4\|F234A_HUMAN | 178,064 | 55,766 | 31% | Q9H0X4 | F234A_HUMAN | F234A_HUMAN | Protein FAM234A (Protein ITFG3) | FAM234A C16orf9 ITFG3 |
| sp\|Q07020\|RL18_HUMAN | 1,505,226 | 1,423,488 | 1,182,480 | 919,315 | sp\|Q07020\|RL18_HUMAN | 857,448 | 481,034 | 56% | Q07020 | RL18_HUMAN | RL18_HUMAN | 60S ribosomal protein L18 (Large ribosomal subunit protein eL18) | RPL18 |
| sp\|Q6FI13\|H2A2A_HUMAN | 1,638,216 | 850,964 | 592,920 | 260,798 | sp\|Q6FI13\|H2A2A_HUMAN | 587,098 | 481,401 | 82% | Q6FI13 | H2A2A_HUMAN | H2A2A_HUMAN | Histone H2A type 2-A (H2A-clustered histone 18) (H2A-clustered histone 19) (Histone H2A.2) (Histone H2A/o) | H2AC18 H2AFO HIST2H2AA HIST2H2AA3; H2AC19 HIST2H2AA4 |
| sp\|P49721\|PSB2_HUMAN | 555,992 | 620,852 | 859,315 | 976,399 | sp\|P49721\|PSB2_HUMAN | 814,231 | 183,948 | 23% | P49721 | PSB2_HUMAN | PSB2_HUMAN | Proteasome subunit beta type-2 (Macropain subunit C7-I) (Multicatalytic endopeptidase complex subunit C7-I) (Proteasome component C7-I) | PSMB2 |

| | | | | | | | | | | | | | |
|---|---|---|---|---|---|---|---|---|---|---|---|---|---|
| sp\|O60884\|DNJA2_HUMAN | 227,094 | 256,280 | 134,095 | 152,974 | sp\|O60884\|DNJA2_HUMAN | 181,864 | 51,110 | 28% | O60884 | DNJA2_HUMAN | DNJA2_HUMAN | DnaJ homolog subfamily A member 2 (Cell cycle progression restoration gene 3 protein) (Dnj3) (Dj3) (HIRA-interacting protein 4) (Renal carcinoma antigen NY-REN-14) | DNAJA2 CPR3 HIRIP4 |
| sp\|Q15008\|PSMD6_HUMAN | 200,799 | 251,672 | 186,717 | 409,930 | sp\|Q15008\|PSMD6_HUMAN | 228,106 | 86,558 | 38% | Q15008 | PSMD6_HUMAN | PSMD6_HUMAN | 26S proteasome non-ATPase regulatory subunit 6 (26S proteasome regulatory subunit RPN7) (26S proteasome regulatory subunit S10) (Breast cancer-associated protein SGA-113M) (Phosphonoformate immuno-associated protein 4) (Proteasome regulatory particle subunit p44S10) (p42A) | PSMD6 KIAA0107 PFAAP4 |
| sp\|Q00796\|DHSO_HUMAN | 288,227 | 365,654 | 444,381 | 453,722 | sp\|Q00796\|DHSO_HUMAN | 390,335 | 58,048 | 15% | Q00796 | DHSO_HUMAN | DHSO_HUMAN | Sorbitol dehydrogenase (SDH) (EC 1.1.1.-) ((R,R)-butanediol dehydrogenase) (EC 1.1.1.4) (L-iditol 2-dehydrogenase) (EC 1.1.1.14) (Polyol dehydrogenase) (Ribitol dehydrogenase) (RDH) (EC 1.1.1.56) (Xylitol dehydrogenase) (XDH) (EC 1.1.1.9) | SORD |
| sp\|P23919\|KTHY_HUMAN | 231,885 | 237,021 | 231,835 | 347,800 | sp\|P23919\|KTHY_HUMAN | 229,741 | 51,229 | 22% | P23919 | KTHY_HUMAN | KTHY_HUMAN | Thymidylate kinase (EC 2.7.4.9) (dTMP kinase) | DTYMK CDC8 TMPK TYMK |
| sp\|Q9P2R7\|SUCB1_HUMAN | 28,663 | 34,408 | 10,995 | 52,605 | sp\|Q9P2R7\|SUCB1_HUMAN | 52,220 | 47,323 | 91% | Q9P2R7 | SUCB1_HUMAN | SUCB1_HUMAN | Succinate--CoA ligase [ADP-forming] subunit beta, mitochondrial (EC 6.2.1.5) (ATP-specific succinyl-CoA synthetase subunit beta) (A-SCS) (Succinyl-CoA synthetase beta-A chain) (SCS-betaA) | SUCLA2 |
| sp\|Q8NG11\|TSN14_HUMAN | 1,346,966 | 1,583,473 | 658,591 | 834,372 | sp\|Q8NG11\|TSN14_HUMAN | 896,658 | 389,504 | 43% | Q8NG11 | TSN14_HUMAN | TSN14_HUMAN | Tetraspanin-14 (Tspan-14) (DC-TM4F2) (Transmembrane 4 superfamily member 14) | TSPAN14 TM4SF14 |
| sp\|Q969Z0\|TBRG4_HUMAN | 10,749 | 14,047 | 13,462 | 5,964 | sp\|Q969Z0\|TBRG4_HUMAN | 19,770 | 14,995 | 76% | Q969Z0 | TBRG4_HUMAN | FAKD4_HUMAN | FAST kinase domain-containing protein 4 (Cell cycle progression restoration protein 2) (Cell cycle progression protein 2) (Protein TBRG4) (Transforming growth factor beta regulator 4) | TBRG4 CPR2 FASTKD4 KIAA0948 |
| sp\|O15260\|SURF4_HUMAN | 347,578 | 365,867 | 188,094 | 488,852 | sp\|O15260\|SURF4_HUMAN | 282,007 | 111,635 | 40% | O15260 | SURF4_HUMAN | SURF4_HUMAN | Surfeit locus protein 4 | SURF4 SURF-4 |
| sp\|Q16698\|DECR_HUMAN | 27,083 | 30,725 | 28,440 | 13,081 | sp\|Q16698\|DECR_HUMAN | 24,551 | 12,828 | 52% | Q16698 | DECR_HUMAN | DECR_HUMAN | 2,4-dienoyl-CoA reductase [(3E)-enoyl-CoA-producing], mitochondrial (EC 1.3.1.124) (2,4-dienoyl-CoA reductase [NADPH]) (4-enoyl-CoA reductase [NADPH]) (Short chain dehydrogenase/reductase family 18C member 1) | DECR1 DECR SDR18C1 |
| sp\|P61019\|RAB2A_HUMAN | 731,946 | 679,204 | 457,961 | 690,292 | sp\|P61019\|RAB2A_HUMAN | 594,885 | 103,323 | 17% | P61019 | RAB2A_HUMAN | RAB2A_HUMAN | Ras-related protein Rab-2A (EC 3.6.5.2) | RAB2A RAB2 |
| sp\|Q12792\|TWF1_HUMAN | 96,679 | 123,378 | 103,790 | 122,648 | sp\|Q12792\|TWF1_HUMAN | 111,580 | 40,333 | 36% | Q12792 | TWF1_HUMAN | TWF1_HUMAN | Twinfilin-1 (Protein A6) (Protein tyrosine kinase 9) | TWF1 PTK9 |
| sp\|P00750\|TPA_HUMAN | 161,498 | 175,089 | 189,374 | 199,734 | sp\|P00750\|TPA_HUMAN | 302,059 | 135,814 | 45% | P00750 | TPA_HUMAN | TPA_HUMAN | Tissue-type plasminogen activator (t-PA) (t-plasminogen activator) (tPA) (EC 3.4.21.68) (Alteplase) (Reteplase) [Cleaved into: Tissue-type plasminogen activator chain A; Tissue-type plasminogen activator chain B] | PLAT |

| | | | | | | | | | | | | | |
|---|---|---|---|---|---|---|---|---|---|---|---|---|---|
| sp\|P00488\|F13A_HUMAN | 23,965 | 6,219 | 50,326 | 42,272 | sp\|P00488\|F13A_HUMAN | 139,059 | 127,958 | 92% | P00488 | F13A_HUMAN | F13A_HUMAN | Coagulation factor XIII A chain (Coagulation factor XIIIa) (EC 2.3.2.13) (Protein-glutamine gamma-glutamyltransferase A chain) (Transglutaminase A chain) | F13A1 F13A |
| sp\|P36776\|LONM_HUMAN | 32,734 | 38,209 | 52,893 | 64,012 | sp\|P36776\|LONM_HUMAN | 54,478 | 31,926 | 59% | P36776 | LONM_HUMAN | LONM_HUMAN | Lon protease homolog, mitochondrial (EC 3.4.21.53) (LONHs) (Lon protease-like protein) (LONP) (Mitochondrial ATP-dependent protease Lon) (Serine protease 15) | LONP1 PRSS15 |
| sp\|Q15738\|NSDHL_HUMAN | 10,615 | 50,771 | 78,723 | 71,494 | sp\|Q15738\|NSDHL_HUMAN | 44,522 | 23,179 | 52% | Q15738 | NSDHL_HUMAN | NSDHL_HUMAN | Sterol-4-alpha-carboxylate 3-dehydrogenase, decarboxylating (EC 1.1.1.170) (Protein H105e3) | NSDHL H105E3 |
| sp\|P62913\|RL11_HUMAN | 121,397 | 135,053 | 67,811 | 41,526 | sp\|P62913\|RL11_HUMAN | 75,244 | 37,248 | 50% | P62913 | RL11_HUMAN | RL11_HUMAN | 60S ribosomal protein L11 (CLL-associated antigen KW-12) (Large ribosomal subunit protein uL5) | RPL11 |
| sp\|P23142\|FBLN1_HUMAN | 99,061 | 70,732 | 169,395 | 113,422 | sp\|P23142\|FBLN1_HUMAN | 832,473 | 811,364 | 97% | P23142 | FBLN1_HUMAN | FBLN1_HUMAN | Fibulin-1 (FIBL-1) | FBLN1 PP213 |
| sp\|Q9UBQ5\|EIF3K_HUMAN | 150,217 | 187,101 | 41,391 | 207,020 | sp\|Q9UBQ5\|EIF3K_HUMAN | 126,178 | 56,126 | 44% | Q9UBQ5 | EIF3K_HUMAN | EIF3K_HUMAN | Eukaryotic translation initiation factor 3 subunit K (eIF3k) (Eukaryotic translation initiation factor 3 subunit 12) (Muscle-specific gene M9 protein) (PLAC-24) (eIF-3 p25) (eIF-3 p28) | EIF3K EIF3S12 ARG134 HSPC029 MSTP001 PTD001 |
| sp\|Q96S86\|HPLN3_HUMAN | 94,551 | 71,619 | 57,701 | 37,717 | sp\|Q96S86\|HPLN3_HUMAN | 82,952 | 31,656 | 38% | Q96S86 | HPLN3_HUMAN | HPLN3_HUMAN | Hyaluronan and proteoglycan link protein 3 | HAPLN3 EXLD1 UNQ238/PRO271 |
| sp\|O94925\|GLSK_HUMAN | 20,446 | 41,508 | 23,233 | 26,775 | sp\|O94925\|GLSK_HUMAN | 53,850 | 43,562 | 81% | O94925 | GLSK_HUMAN | GLSK_HUMAN | Glutaminase kidney isoform, mitochondrial (GLS) (EC 3.5.1.2) (K-glutaminase) (L-glutamine amidohydrolase) [Cleaved into: Glutaminase kidney isoform, mitochondrial 68 kDa chain; Glutaminase kidney isoform, mitochondrial 65 kDa chain] | GLS GLS1 KIAA0838 |
| sp\|P62851\|RS25_HUMAN | 77,296 | 137,177 | 83,741 | 51,265 | sp\|P62851\|RS25_HUMAN | 66,199 | 40,110 | 61% | P62851 | RS25_HUMAN | RS25_HUMAN | 40S ribosomal protein S25 (Small ribosomal subunit protein eS25) | RPS25 |
| sp\|O94919\|ENDD1_HUMAN | 211,463 | 154,203 | 204,713 | 203,819 | sp\|O94919\|ENDD1_HUMAN | 153,790 | 56,103 | 36% | O94919 | ENDD1_HUMAN | ENDD1_HUMAN | Endonuclease domain-containing 1 protein (EC 3.1.30.-) | ENDOD1 KIAA0830 |
| sp\|P51159\|RB27A_HUMAN | 279,463 | 313,291 | 139,837 | 166,664 | sp\|P51159\|RB27A_HUMAN | 216,096 | 68,071 | 32% | P51159 | RB27A_HUMAN | RB27A_HUMAN | Ras-related protein Rab-27A (Rab-27) (EC 3.6.5.2) (GTP-binding protein Ram) | RAB27A RAB27 |
| sp\|Q92542\|NICA_HUMAN | 234,715 | 289,117 | 216,589 | 109,702 | sp\|Q92542\|NICA_HUMAN | 190,907 | 78,334 | 41% | Q92542 | NICA_HUMAN | NICA_HUMAN | Nicastrin | NCSTN KIAA0253 UNQ1874/PRO4317 |
| sp\|P18206\|VINC_HUMAN | 67,954 | 150,821 | 41,957 | 76,842 | sp\|P18206\|VINC_HUMAN | 118,668 | 70,004 | 59% | P18206 | VINC_HUMAN | VINC_HUMAN | Vinculin (Metavinculin) (MV) | VCL |
| sp\|Q02809\|PLOD1_HUMAN | 113,092 | 64,504 | 244,709 | 163,037 | sp\|Q02809\|PLOD1_HUMAN | 134,874 | 61,386 | 46% | Q02809 | PLOD1_HUMAN | PLOD1_HUMAN | Procollagen-lysine,2-oxoglutarate 5-dioxygenase 1 (EC 1.14.11.4) (Lysyl hydroxylase 1) (LH1) | PLOD1 LLH PLOD |
| sp\|P21926\|CD9_HUMAN | 4,942,783 | 4,957,254 | 2,757,413 | 2,820,870 | sp\|P21926\|CD9_HUMAN | 3,243,729 | 1,252,299 | 39% | P21926 | CD9_HUMAN | CD9_HUMAN | CD9 antigen (5H9 antigen) (Cell growth-inhibiting gene 2 protein) (Leukocyte antigen MIC3) (Motility-related protein) (MRP-1) (Tetraspanin-29) (Tspan-29) (p24) (CD antigen CD9) | CD9 MIC3 TSPAN29 GIG2 |
| sp\|O75054\|IGSF3_HUMAN | 143,328 | 86,661 | 176,515 | 167,854 | sp\|O75054\|IGSF3_HUMAN | 131,221 | 60,105 | 46% | O75054 | IGSF3_HUMAN | IGSF3_HUMAN | Immunoglobulin superfamily member 3 (IgSF3) (Glu-Trp-Ile EWI motif-containing protein 3) (EWI-3) | IGSF3 EWI3 KIAA0466 |
| sp\|Q32P28\|P3H1_HUMAN | 51,408 | 41,490 | 42,390 | 65,078 | sp\|Q32P28\|P3H1_HUMAN | 52,510 | 8,258 | 16% | Q32P28 | P3H1_HUMAN | P3H1_HUMAN | Prolyl 3-hydroxylase 1 (EC 1.14.11.7) (Growth suppressor 1) (Leucine- and proline-enriched proteoglycan 1) (Leprecan-1) | P3H1 GROS1 LEPRE1 PSEC0109 |

| | | | | | | | | | | | | | |
|---|---|---|---|---|---|---|---|---|---|---|---|---|---|
| sp\|P21964\|COMT_HUMAN | 52,579 | 73,457 | 4,334 | 86,135 | sp\|P21964\|COMT_HUMAN | 42,316 | 27,695 | 65% | P21964 | COMT_HUMAN | COMT_HUMAN | Catechol O-methyltransferase (EC 2.1.1.6) | COMT |
| sp\|Q9UKS6\|PACN3_HUMAN | 216,880 | 275,757 | 402,883 | 426,201 | sp\|Q9UKS6\|PACN3_HUMAN | 307,692 | 83,326 | 27% | Q9UKS6 | PACN3_HUMAN | PACN3_HUMAN | Protein kinase C and casein kinase substrate in neurons protein 3 (SH3 domain-containing protein 6511) | PACSIN3 |
| sp\|Q9Y678\|COPG1_HUMAN | 89,191 | 70,736 | 173,159 | 110,151 | sp\|Q9Y678\|COPG1_HUMAN | 97,730 | 36,869 | 38% | Q9Y678 | COPG1_HUMAN | COPG1_HUMAN | Coatomer subunit gamma-1 (Gamma-1-coat protein) (Gamma-1-COP) | COPG1 COPG |
| sp\|O95084\|PRS23_HUMAN | 234,874 | 248,888 | 194,101 | 226,085 | sp\|O95084\|PRS23_HUMAN | 207,216 | 36,029 | 17% | O95084 | PRS23_HUMAN | PRS23_HUMAN | Serine protease 23 (EC 3.4.21.-) (Putative secreted protein Zsig13) | PRSS23 ZSIG13 UNQ270/PRO307 |
| sp\|Q99880\|H2B1L_HUMAN | 2,200,356 | 1,574,793 | 4,355,583 | 1,499,348 | sp\|Q99880\|H2B1L_HUMAN | 1,916,624 | 1,220,009 | 64% | Q99880 | H2B1L_HUMAN | H2B1L_HUMAN | Histone H2B type 1-L (Histone H2B.c) (H2B/c) | H2BC13 H2BFC HIST1H2BL |
| sp\|P62633\|CNBP_HUMAN | 114,655 | 116,647 | 110,473 | 152,524 | sp\|P62633\|CNBP_HUMAN | 112,005 | 34,721 | 31% | P62633 | CNBP_HUMAN | CNBP_HUMAN | CCHC-type zinc finger nucleic acid binding protein (Cellular nucleic acid-binding protein) (CNBP) (Zinc finger protein 9) | CNBP RNF163 ZNF9 |
| sp\|O43390\|HNRPR_HUMAN | 448,225 | 331,845 | 217,854 | 189,617 | sp\|O43390\|HNRPR_HUMAN | 200,193 | 129,782 | 65% | O43390 | HNRPR_HUMAN | HNRPR_HUMAN | Heterogeneous nuclear ribonucleoprotein R (hnRNP R) | HNRNPR HNRPR |
| sp\|Q96FN4\|CPNE2_HUMAN | 291,615 | 279,155 | 267,717 | 305,078 | sp\|Q96FN4\|CPNE2_HUMAN | 267,337 | 36,637 | 14% | Q96FN4 | CPNE2_HUMAN | CPNE2_HUMAN | Copine-2 (Copine II) | CPNE2 |
| sp\|Q9BRX8\|F213A_HUMAN | 139,670 | 152,904 | 92,146 | 150,301 | sp\|Q9BRX8\|F213A_HUMAN | 133,110 | 39,814 | 30% | Q9BRX8 | F213A_HUMAN | PXL2A_HUMAN | Peroxiredoxin-like 2A (Peroxiredoxin-like 2 activated in M-CSF stimulated monocytes) (Protein PAMM) (Redox-regulatory protein FAM213A) | PRXL2A C10orf58 FAM213A PAMM PRO2290 PSEC0139 UNQ611/PRO1198 |
| sp\|Q8TCT9\|HM13_HUMAN | 198,377 | 236,131 | 90,745 | 155,560 | sp\|Q8TCT9\|HM13_HUMAN | 129,917 | 65,172 | 50% | Q8TCT9 | HM13_HUMAN | HM13_HUMAN | Minor histocompatibility antigen H13 (EC 3.4.23.-) (Intramembrane protease 1) (IMP-1) (IMPAS-1) (hIMP1) (Presenilin-like protein 3) (Signal peptide peptidase) | HM13 H13 IMP1 PSL3 SPP MSTP086 |
| sp\|Q16543\|CDC37_HUMAN | 85,925 | 114,743 | 113,378 | 138,775 | sp\|Q16543\|CDC37_HUMAN | 103,507 | 21,329 | 21% | Q16543 | CDC37_HUMAN | CDC37_HUMAN | Hsp90 co-chaperone Cdc37 (Hsp90 chaperone protein kinase-targeting subunit) (p50Cdc37) [Cleaved into: Hsp90 co-chaperone Cdc37, N-terminally processed] | CDC37 CDC37A |
| sp\|Q8NC51\|PAIRB_HUMAN | 62,378 | 70,291 | 35,370 | 38,758 | sp\|Q8NC51\|PAIRB_HUMAN | 59,670 | 33,846 | 57% | Q8NC51 | PAIRB_HUMAN | PAIRB_HUMAN | Plasminogen activator inhibitor 1 RNA-binding protein (PAI1 RNA-binding protein 1) (PAI-RBP1) (SERPINE1 mRNA-binding protein 1) | SERBP1 PAIRBP1 CGI-55 |
| sp\|Q9BVA1\|TBB2B_HUMAN | 102,989 | 97,468 | 80,880 | 77,691 | sp\|Q9BVA1\|TBB2B_HUMAN | 69,645 | 24,822 | 36% | Q9BVA1 | TBB2B_HUMAN | TBB2B_HUMAN | Tubulin beta-2B chain | TUBB2B |
| sp\|Q9H4B7\|TBB1_HUMAN | 452,337 | 633,694 | 75,721 | 268,642 | sp\|Q9H4B7\|TBB1_HUMAN | 467,653 | 200,990 | 43% | Q9H4B7 | TBB1_HUMAN | TBB1_HUMAN | Tubulin beta-1 chain | TUBB1 |
| sp\|Q15366\|PCBP2_HUMAN | 284,269 | 573,585 | 344,017 | 558,212 | sp\|Q15366\|PCBP2_HUMAN | 346,361 | 152,618 | 44% | Q15366 | PCBP2_HUMAN | PCBP2_HUMAN | Poly(rC)-binding protein 2 (Alpha-CP2) (Heterogeneous nuclear ribonucleoprotein E2) (hnRNP E2) | PCBP2 |
| sp\|P32322\|P5CR1_HUMAN | 37,967 | 35,970 | 41,846 | 26,644 | sp\|P32322\|P5CR1_HUMAN | 41,733 | 22,811 | 55% | P32322 | P5CR1_HUMAN | P5CR1_HUMAN | Pyrroline-5-carboxylate reductase 1, mitochondrial (P5C reductase 1) (P5CR 1) (EC 1.5.1.2) | PYCR1 |
| sp\|P19827\|ITIH1_HUMAN | 28,993 | 18,002 | 17,857 | 30,929 | sp\|P19827\|ITIH1_HUMAN | 205,487 | 197,345 | 96% | P19827 | ITIH1_HUMAN | ITIH1_HUMAN | Inter-alpha-trypsin inhibitor heavy chain H1 (ITI heavy chain H1) (ITI-HC1) (Inter-alpha-inhibitor heavy chain 1) (Inter-alpha-trypsin inhibitor complex component III) (Serum-derived hyaluronan-associated protein) (SHAP) | ITIH1 IGHEP1 |

| | | | | | | | | | | | | | |
|---|---|---|---|---|---|---|---|---|---|---|---|---|---|
| sp\|P28072\|PSB6_HUMAN | 413,861 | 353,199 | 595,934 | 641,186 | sp\|P28072\|PSB6_HUMAN | 468,818 | 125,889 | 27% | P28072 | PSB6_HUMAN | PSB6_HUMAN | Proteasome subunit beta type-6 (EC 3.4.25.1) (Macropain delta chain) (Multicatalytic endopeptidase complex delta chain) (Proteasome delta chain) (Proteasome subunit Y) | PSMB6 LMPY Y |
| sp\|Q07866\|KLC1_HUMAN | 67,253 | 80,989 | 91,231 | 138,030 | sp\|Q07866\|KLC1_HUMAN | 78,630 | 32,224 | 41% | Q07866 | KLC1_HUMAN | KLC1_HUMAN | Kinesin light chain 1 (KLC 1) | KLC1 KLC KNS2 |
| sp\|P61006\|RAB8A_HUMAN | 85,873 | 124,427 | 43,517 | 120,986 | sp\|P61006\|RAB8A_HUMAN | 79,097 | 44,526 | 56% | P61006 | RAB8A_HUMAN | RAB8A_HUMAN | Ras-related protein Rab-8A (EC 3.6.5.2) (Oncogene c-mel) | RAB8A MEL RAB8 |
| sp\|P43487\|RANG_HUMAN | 48,799 | 133,454 | 57,692 | 98,566 | sp\|P43487\|RANG_HUMAN | 73,351 | 30,949 | 42% | P43487 | RANG_HUMAN | RANG_HUMAN | Ran-specific GTPase-activating protein (Ran-binding protein 1) (RanBP1) | RANBP1 |
| sp\|Q96S97\|MYADM_HUMAN | 1,065,693 | 1,339,520 | 1,456,512 | 1,814,649 | sp\|Q96S97\|MYADM_HUMAN | 1,230,323 | 292,766 | 24% | Q96S97 | MYADM_HUMAN | MYADM_HUMAN | Myeloid-associated differentiation marker (Protein SB135) | MYADM UNQ553/PRO1110 |
| sp\|P49189\|AL9A1_HUMAN | 125,301 | 132,724 | 76,812 | 111,624 | sp\|P49189\|AL9A1_HUMAN | 113,375 | 19,734 | 17% | P49189 | AL9A1_HUMAN | AL9A1_HUMAN | 4-trimethylaminobutyraldehyde dehydrogenase (TMABA-DH) (TMABALDH) (EC 1.2.1.47) (Aldehyde dehydrogenase E3 isozyme) (Aldehyde dehydrogenase family 9 member A1) (EC 1.2.1.3) (Gamma-aminobutyraldehyde dehydrogenase) (EC 1.2.1.19) (R-aminobutyraldehyde dehydrogenase) [Cleaved into: 4-trimethylaminobutyraldehyde dehydrogenase, N-terminally processed] | ALDH9A1 ALDH4 ALDH7 ALDH9 |
| sp\|P10586\|PTPRF_HUMAN | 248,133 | 276,090 | 343,944 | 479,983 | sp\|P10586\|PTPRF_HUMAN | 401,508 | 141,799 | 35% | P10586 | PTPRF_HUMAN | PTPRF_HUMAN | Receptor-type tyrosine-protein phosphatase F (EC 3.1.3.48) (Leukocyte common antigen related) (LAR) | PTPRF LAR |
| sp\|P05543\|THBG_HUMAN | 54,375 | 22,028 | 50,287 | 56,854 | sp\|P05543\|THBG_HUMAN | 240,952 | 300,594 | 125% | P05543 | THBG_HUMAN | THBG_HUMAN | Thyroxine-binding globulin (Serpin A7) (T4-binding globulin) | SERPINA7 TBG |
| sp\|P46779\|RL28_HUMAN | 369,335 | 326,748 | 178,922 | 197,525 | sp\|P46779\|RL28_HUMAN | 187,468 | 109,761 | 59% | P46779 | RL28_HUMAN | RL28_HUMAN | 60S ribosomal protein L28 (Large ribosomal subunit protein eL28) | RPL28 |
| sp\|P04114\|APOB_HUMAN | 53,154 | 50,595 | 57,073 | 68,164 | sp\|P04114\|APOB_HUMAN | 315,800 | 307,922 | 98% | P04114 | APOB_HUMAN | APOB_HUMAN | Apolipoprotein B-100 (Apo B-100) [Cleaved into: Apolipoprotein B-48 (Apo B-48)] | APOB |
| sp\|O94973\|AP2A2_HUMAN | 140,304 | 257,000 | 215,951 | 186,403 | sp\|O94973\|AP2A2_HUMAN | 215,654 | 134,812 | 63% | O94973 | AP2A2_HUMAN | AP2A2_HUMAN | AP-2 complex subunit alpha-2 (100 kDa coated vesicle protein C) (Adaptor protein complex AP-2 subunit alpha-2) (Adaptor-related protein complex 2 subunit alpha-2) (Alpha-adaptin C) (Alpha2-adaptin) (Clathrin assembly protein complex 2 alpha-C large chain) (Huntingtin yeast partner J) (Huntingtin-interacting protein 9) (HIP-9) (Huntingtin-interacting protein J) (Plasma membrane adaptor HA2/AP2 adaptin alpha C subunit) | AP2A2 ADTAB CLAPA2 HIP9 HYPJ KIAA0899 |
| sp\|Q99816\|TS101_HUMAN | 466,854 | 643,423 | 969,276 | 871,395 | sp\|Q99816\|TS101_HUMAN | 698,765 | 189,252 | 27% | Q99816 | TS101_HUMAN | TS101_HUMAN | Tumor susceptibility gene 101 protein (ESCRT-I complex subunit TSG101) | TSG101 |
| sp\|P52788\|SPSY_HUMAN | 127,169 | 142,263 | 96,228 | 86,813 | sp\|P52788\|SPSY_HUMAN | 101,143 | 30,770 | 30% | P52788 | SPSY_HUMAN | SPSY_HUMAN | Spermine synthase (SPMSY) (EC 2.5.1.22) (Spermidine aminopropyltransferase) | SMS |

| | | | | | | | | | | | | | |
|---|---|---|---|---|---|---|---|---|---|---|---|---|---|
| sp\|Q9Y320\|TMX2_HUMAN | 30,961 | 21,700 | 39,467 | 17,129 | sp\|Q9Y320\|TMX2_HUMAN | 68,741 | 62,637 | 91% | Q9Y320 | TMX2_HUMAN | TMX2_HUMAN | Thioredoxin-related transmembrane protein 2 (Cell proliferation-inducing gene 26 protein) (Thioredoxin domain-containing protein 14) | TMX2 TXNDC14 CGI-31 My009 PIG26 PSEC0045 UNQ237/PRO270 |
| sp\|O43633\|CHM2A_HUMAN | 376,071 | 426,672 | 347,379 | 347,291 | sp\|O43633\|CHM2A_HUMAN | 346,810 | 44,333 | 13% | O43633 | CHM2A_HUMAN | CHM2A_HUMAN | Charged multivesicular body protein 2a (Chromatin-modifying protein 2a) (CHMP2a) (Putative breast adenocarcinoma marker BC-2) (Vacuolar protein sorting-associated protein 2-1) (Vps2-1) (hVps2-1) | CHMP2A BC2 CHMP2 |
| sp\|P50914\|RL14_HUMAN | 1,224,174 | 1,057,522 | 986,663 | 937,176 | sp\|P50914\|RL14_HUMAN | 750,195 | 336,895 | 45% | P50914 | RL14_HUMAN | RL14_HUMAN | 60S ribosomal protein L14 (CAG-ISL 7) (Large ribosomal subunit protein eL14) | RPL14 |
| sp\|Q8NBJ5\|GT251_HUMAN | 19,128 | 11,897 | 27,004 | 9,202 | sp\|Q8NBJ5\|GT251_HUMAN | 14,152 | 7,459 | 53% | Q8NBJ5 | GT251_HUMAN | GT251_HUMAN | Procollagen galactosyltransferase 1 (EC 2.4.1.50) (Collagen beta(1-O)galactosyltransferase 1) (ColGalT 1) (Glycosyltransferase 25 family member 1) (Hydroxylysine galactosyltransferase 1) | COLGALT1 GLT25D1 PSEC0241 |
| sp\|Q9BWD1\|THIC_HUMAN | 27,761 | 191,030 | 83,049 | 65,563 | sp\|Q9BWD1\|THIC_HUMAN | 95,844 | 48,887 | 51% | Q9BWD1 | THIC_HUMAN | THIC_HUMAN | Acetyl-CoA acetyltransferase, cytosolic (EC 2.3.1.9) (Acetyl-CoA transferase-like protein) (Cytosolic acetoacetyl-CoA thiolase) | ACAT2 ACTL |
| sp\|Q9HAV0\|GBB4_HUMAN | 319,455 | 372,612 | 292,665 | 304,864 | sp\|Q9HAV0\|GBB4_HUMAN | 349,280 | 84,206 | 24% | Q9HAV0 | GBB4_HUMAN | GBB4_HUMAN | Guanine nucleotide-binding protein subunit beta-4 (Transducin beta chain 4) | GNB4 |
| sp\|P10301\|RRAS_HUMAN | 245,480 | 329,858 | 205,203 | 382,174 | sp\|P10301\|RRAS_HUMAN | 279,510 | 59,583 | 21% | P10301 | RRAS_HUMAN | RRAS_HUMAN | Ras-related protein R-Ras (EC 3.6.5.-) (p23) | RRAS |
| sp\|P31949\|S10AB_HUMAN | 215,815 | 344,136 | 282,777 | 245,340 | sp\|P31949\|S10AB_HUMAN | 277,656 | 59,030 | 21% | P31949 | S10AB_HUMAN | S10AB_HUMAN | Protein S100-A11 (Calgizzarin) (Metastatic lymph node gene 70 protein) (MLN 70) (Protein S100-C) (S100 calcium-binding protein A11) [Cleaved into: Protein S100-A11, N-terminally processed] | S100A11 MLN70 S100C |
| sp\|P39060\|COIA1_HUMAN | 393,362 | 250,658 | 688,729 | 501,107 | sp\|P39060\|COIA1_HUMAN | 571,109 | 273,418 | 48% | P39060 | COIA1_HUMAN | COIA1_HUMAN | Collagen alpha-1(XVIII) chain [Cleaved into: Endostatin; Non-collagenous domain 1 (NC1)] | COL18A1 |
| sp\|P62495\|ERF1_HUMAN | 110,415 | 110,078 | 87,781 | 104,033 | sp\|P62495\|ERF1_HUMAN | 81,787 | 29,656 | 36% | P62495 | ERF1_HUMAN | ERF1_HUMAN | Eukaryotic peptide chain release factor subunit 1 (Eukaryotic release factor 1) (eRF1) (Protein Cl1) (TB3-1) | ETF1 ERF1 RF1 SUP45L1 |
| sp\|Q9ULZ3\|ASC_HUMAN | 193,532 | 344,131 | 177,876 | 249,743 | sp\|Q9ULZ3\|ASC_HUMAN | 213,387 | 67,658 | 32% | Q9ULZ3 | ASC_HUMAN | ASC_HUMAN | Apoptosis-associated speck-like protein containing a CARD (hASC) (Caspase recruitment domain-containing protein 5) (PYD and CARD domain-containing protein) (Target of methylation-induced silencing 1) | PYCARD ASC CARD5 TMS1 |
| sp\|P48739\|PIPNB_HUMAN | 237,409 | 204,510 | 75,803 | 66,972 | sp\|P48739\|PIPNB_HUMAN | 145,428 | 73,297 | 50% | P48739 | PIPNB_HUMAN | PIPNB_HUMAN | Phosphatidylinositol transfer protein beta isoform (PI-TP-beta) (PtdIns transfer protein beta) (PtdInsTP beta) | PITPNB |
| sp\|Q96EY5\|MB12A_HUMAN | 314,869 | 385,693 | 186,361 | 308,360 | sp\|Q96EY5\|MB12A_HUMAN | 283,007 | 74,426 | 26% | Q96EY5 | MB12A_HUMAN | MB12A_HUMAN | Multivesicular body subunit 12A (CIN85/CD2AP family-binding protein) (ESCRT-I complex subunit MVB12A) (Protein FAM125A) | MVB12A CFBP FAM125A |

| | | | | | | | | | | | | | |
|---|---|---|---|---|---|---|---|---|---|---|---|---|---|
| sp\|P61421\|VA0D1_HUMAN | 217,608 | 193,551 | 213,190 | 366,829 | sp\|P61421\|VA0D1_HUMAN | 226,288 | 85,265 | 38% | P61421 | VA0D1_HUMAN | VA0D1_HUMAN | V-type proton ATPase subunit d 1 (V-ATPase subunit d 1) (32 kDa accessory protein) (V-ATPase 40 kDa accessory protein) (V-ATPase AC39 subunit) (p39) (Vacuolar proton pump subunit d 1) | ATP6V0D1 ATP6D VPATPD |
| sp\|P49720\|PSB3_HUMAN | 209,187 | 231,448 | 98,311 | 222,560 | sp\|P49720\|PSB3_HUMAN | 197,965 | 41,808 | 21% | P49720 | PSB3_HUMAN | PSB3_HUMAN | Proteasome subunit beta type-3 (Proteasome chain 13) (Proteasome component C10-II) (Proteasome theta chain) | PSMB3 |
| sp\|O94804\|STK10_HUMAN | 250,710 | 266,100 | 324,544 | 258,490 | sp\|O94804\|STK10_HUMAN | 229,432 | 60,259 | 26% | O94804 | STK10_HUMAN | STK10_HUMAN | Serine/threonine-protein kinase 10 (EC 2.7.11.1) (Lymphocyte-oriented kinase) | STK10 LOK |
| sp\|Q9Y2A7\|NCKP1_HUMAN | 270,407 | 332,974 | 37,384 | 309,867 | sp\|Q9Y2A7\|NCKP1_HUMAN | 197,820 | 102,398 | 52% | Q9Y2A7 | NCKP1_HUMAN | NCKP1_HUMAN | Nck-associated protein 1 (NAP 1) (Membrane-associated protein HEM-2) (p125Nap1) | NCKAP1 HEM2 KIAA0587 NAP1 |
| sp\|Q9UK41\|VPS28_HUMAN | 518,248 | 628,625 | 769,346 | 857,527 | sp\|Q9UK41\|VPS28_HUMAN | 677,983 | 142,977 | 21% | Q9UK41 | VPS28_HUMAN | VPS28_HUMAN | Vacuolar protein sorting-associated protein 28 homolog (H-Vps28) (ESCRT-I complex subunit VPS28) | VPS28 |
| sp\|P02771\|FETA_HUMAN | 34,208 | 35,454 | 48,117 | 56,083 | sp\|P02771\|FETA_HUMAN | 280,982 | 320,011 | 114% | P02771 | FETA_HUMAN | FETA_HUMAN | Alpha-fetoprotein (Alpha-1-fetoprotein) (Alpha-fetoglobulin) | AFP HPAFP |
| sp\|Q12907\|LMAN2_HUMAN | 154,042 | 108,439 | 168,517 | 129,241 | sp\|Q12907\|LMAN2_HUMAN | 113,592 | 43,252 | 38% | Q12907 | LMAN2_HUMAN | LMAN2_HUMAN | Vesicular integral-membrane protein VIP36 (Glycoprotein GP36b) (Lectin mannose-binding 2) (Vesicular integral-membrane protein 36) (VIP36) | LMAN2 C5orf8 |
| sp\|Q15582\|BGH3_HUMAN | 72,409 | 17,727 | 63,999 | 60,564 | sp\|Q15582\|BGH3_HUMAN | 129,104 | 95,090 | 74% | Q15582 | BGH3_HUMAN | BGH3_HUMAN | Transforming growth factor-beta-induced protein ig-h3 (Beta ig-h3) (Kerato-epithelin) (RGD-containing collagen-associated protein) (RGD-CAP) | TGFBI BIGH3 |
| sp\|P62899\|RL31_HUMAN | 465,862 | 594,284 | 509,819 | 557,283 | sp\|P62899\|RL31_HUMAN | 453,679 | 106,856 | 24% | P62899 | RL31_HUMAN | RL31_HUMAN | 60S ribosomal protein L31 (Large ribosomal subunit protein eL31) | RPL31 |
| sp\|P39059\|COFA1_HUMAN | 1,111,364 | 813,668 | 1,558,277 | 1,100,163 | sp\|P39059\|COFA1_HUMAN | 1,028,171 | 248,275 | 24% | P39059 | COFA1_HUMAN | COFA1_HUMAN | Collagen alpha-1(XV) chain [Cleaved into: Restin (Endostatin-XV) (Related to endostatin) (Restin-I); Restin-2 (Restin-II); Restin-3 (Restin-III); Restin-4 (Restin-IV)] | COL15A1 |
| sp\|Q5VW32\|BROX_HUMAN | 279,992 | 369,474 | 135,812 | 95,333 | sp\|Q5VW32\|BROX_HUMAN | 182,870 | 106,934 | 58% | Q5VW32 | BROX_HUMAN | BROX_HUMAN | BRO1 domain-containing protein BROX (BRO1 domain- and CAAX motif-containing protein) | BROX BROFTI C1orf58 |
| sp\|P37802\|TAGL2_HUMAN | 136,050 | 253,607 | 223,866 | 304,712 | sp\|P37802\|TAGL2_HUMAN | 304,951 | 98,525 | 32% | P37802 | TAGL2_HUMAN | TAGL2_HUMAN | Transgelin-2 (Epididymis tissue protein Li 7e) (SM22-alpha homolog) | TAGLN2 KIAA0120 CDABP0035 |
| sp\|P13987\|CD59_HUMAN | 3,534,658 | 3,552,245 | 2,623,526 | 2,528,950 | sp\|P13987\|CD59_HUMAN | 2,312,568 | 907,329 | 39% | P13987 | CD59_HUMAN | CD59_HUMAN | CD59 glycoprotein (1F5 antigen) (20 kDa homologous restriction factor) (HRF-20) (HRF20) (MAC-inhibitory protein) (MAC-IP) (MEM43 antigen) (Membrane attack complex inhibition factor) (MACIF) (Membrane inhibitor of reactive lysis) (MIRL) (Protectin) (CD antigen CD59) | CD59 MIC11 MIN1 MIN2 MIN3 MSK21 |
| sp\|P13473\|LAMP2_HUMAN | 865,439 | 950,655 | 1,034,662 | 1,074,945 | sp\|P13473\|LAMP2_HUMAN | 753,931 | 250,928 | 33% | P13473 | LAMP2_HUMAN | LAMP2_HUMAN | Lysosome-associated membrane glycoprotein 2 (LAMP-2) (Lysosome-associated membrane protein 2) (CD107 antigen-like family member B) (LGP-96) (CD antigen CD107b) | LAMP2 |

| | | | | | | | | | | | | | |
|---|---|---|---|---|---|---|---|---|---|---|---|---|---|
| sp\|Q9P035\|HACD3_HUMAN | 64,015 | 68,190 | 64,946 | 86,021 | sp\|Q9P035\|HACD3_HUMAN | 61,924 | 19,338 | 31% | Q9P035 | HACD3_HUMAN | HACD3_HUMAN | Very-long-chain (3R)-3-hydroxyacyl-CoA dehydratase 3 (EC 4.2.1.134) (3-hydroxyacyl-CoA dehydratase 3) (HACD3) (Butyrate-induced protein 1) (B-ind1) (hB-ind1) (Protein-tyrosine phosphatase-like A domain-containing protein 1) | HACD3 BIND1 PTPLAD1 |
| sp\|P31943\|HNRH1_HUMAN | 142,216 | 119,838 | 127,719 | 110,856 | sp\|P31943\|HNRH1_HUMAN | 86,533 | 42,508 | 49% | P31943 | HNRH1_HUMAN | HNRH1_HUMAN | Heterogeneous nuclear ribonucleoprotein H (hnRNP H) [Cleaved into: Heterogeneous nuclear ribonucleoprotein H, N-terminally processed] | HNRNPH1 HNRPH HNRPH1 |
| sp\|Q99426\|TBCB_HUMAN | 56,697 | 54,923 | 78,073 | 83,341 | sp\|Q99426\|TBCB_HUMAN | 55,902 | 16,596 | 30% | Q99426 | TBCB_HUMAN | TBCB_HUMAN | Tubulin-folding cofactor B (Cytoskeleton-associated protein 1) (Cytoskeleton-associated protein CKAPI) (Tubulin-specific chaperone B) | TBCB CG22 CKAP1 |
| sp\|Q96P70\|IPO9_HUMAN | 23,446 | 49,090 | 30,064 | 64,722 | sp\|Q96P70\|IPO9_HUMAN | 51,122 | 21,770 | 43% | Q96P70 | IPO9_HUMAN | IPO9_HUMAN | Importin-9 (Imp9) (Ran-binding protein 9) (RanBP9) | IPO9 IMP9 KIAA1192 RANBP9 HSPC273 |
| sp\|P12109\|CO6A1_HUMAN | 4,874 | 1,733 | 15,331 | 12,774 | sp\|P12109\|CO6A1_HUMAN | 88,575 | 98,259 | 111% | P12109 | CO6A1_HUMAN | CO6A1_HUMAN | Collagen alpha-1(VI) chain | COL6A1 |
| sp\|Q99808\|S29A1_HUMAN | 516,418 | 732,369 | 595,971 | 969,399 | sp\|Q99808\|S29A1_HUMAN | 600,091 | 272,417 | 45% | Q99808 | S29A1_HUMAN | S29A1_HUMAN | Equilibrative nucleoside transporter 1 (Equilibrative nitrobenzylmercaptopurine riboside-sensitive nucleoside transporter) (Equilibrative NBMPR-sensitive nucleoside transporter) (Nucleoside transporter, es-type) (Solute carrier family 29 member 1) | SLC29A1 ENT1 |
| sp\|Q9BXS5\|AP1M1_HUMAN | 80,169 | 156,549 | 118,658 | 153,614 | sp\|Q9BXS5\|AP1M1_HUMAN | 123,489 | 24,862 | 20% | Q9BXS5 | AP1M1_HUMAN | AP1M1_HUMAN | AP-1 complex subunit mu-1 (AP-mu chain family member mu1A) (Adaptor protein complex AP-1 subunit mu-1) (Adaptor-related protein complex 1 subunit mu-1) (Clathrin assembly protein complex 1 mu-1 medium chain 1) (Clathrin coat assembly protein AP47) (Clathrin coat-associated protein AP47) (Golgi adaptor HA1/AP1 adaptin mu-1 subunit) (Mu-adaptin 1) (Mu1A-adaptin) | AP1M1 CLTNM |
| sp\|P33121\|ACSL1_HUMAN | 29,278 | 23,246 | 17,003 | 22,079 | sp\|P33121\|ACSL1_HUMAN | 29,061 | 10,988 | 38% | P33121 | ACSL1_HUMAN | ACSL1_HUMAN | Long-chain-fatty-acid--CoA ligase 1 (EC 6.2.1.3) (Acyl-CoA synthetase 1) (ACS1) (Arachidonate--CoA ligase) (EC 6.2.1.15) (Long-chain acyl-CoA synthetase 1) (LACS 1) (Long-chain acyl-CoA synthetase 2) (LACS 2) (Long-chain fatty acid-CoA ligase 2) (Palmitoyl-CoA ligase 1) (Palmitoyl-CoA ligase 2) (Phytanate--CoA ligase) (EC 6.2.1.24) | ACSL1 FACL1 FACL2 LACS LACS1 LACS2 |
| sp\|O15127\|SCAM2_HUMAN | 442,178 | 389,275 | 306,229 | 325,070 | sp\|O15127\|SCAM2_HUMAN | 339,457 | 74,762 | 22% | O15127 | SCAM2_HUMAN | SCAM2_HUMAN | Secretory carrier-associated membrane protein 2 (Secretory carrier membrane protein 2) | SCAMP2 |
| sp\|Q13425\|SNTB2_HUMAN | 383,146 | 183,502 | 449,397 | 331,887 | sp\|Q13425\|SNTB2_HUMAN | 327,667 | 89,927 | 27% | Q13425 | SNTB2_HUMAN | SNTB2_HUMAN | Beta-2-syntrophin (59 kDa dystrophin-associated protein A1 basic component 2) (Syntrophin-3) (SNT3) (Syntrophin-like) (SNTL) | SNTB2 D16S2531E SNT2B2 SNTL |

| | | | | | | | | | | | | | |
|---|---|---|---|---|---|---|---|---|---|---|---|---|---|
| sp\|Q15392\|DHC24_HUMAN | 17,426 | 8,173 | 14,873 | 8,901 | sp\|Q15392\|DHC24_HUMAN | 13,645 | 7,926 | 58% | Q15392 | DHC24_HUMAN | DHC24_HUMAN | Delta(24)-sterol reductase (EC 1.3.1.72) (24-dehydrocholesterol reductase) (3-beta-hydroxysterol Delta-24-reductase) (Diminuto/dwarf1 homolog) (Seladin-1) | DHCR24 KIAA0018 |
| sp\|Q9Y289\|SC5A6_HUMAN | 597,842 | 659,020 | 552,333 | 435,175 | sp\|Q9Y289\|SC5A6_HUMAN | 516,635 | 98,973 | 19% | Q9Y289 | SC5A6_HUMAN | SC5A6_HUMAN | Sodium-dependent multivitamin transporter (Na(+)-dependent multivitamin transporter) (Solute carrier family 5 member 6) | SLC5A6 SMVT |
| sp\|Q9Y2H2\|SAC2_HUMAN | 81,690 | 117,446 | 125,562 | 161,671 | sp\|Q9Y2H2\|SAC2_HUMAN | 135,534 | 39,179 | 29% | Q9Y2H2 | SAC2_HUMAN | SAC2_HUMAN | Phosphatidylinositide phosphatase SAC2 (EC 3.1.3.25) (Inositol polyphosphate 5-phosphatase F) (Sac domain-containing inositol phosphatase 2) (Sac domain-containing phosphoinositide 4-phosphatase 2) (hSAC2) | INPP5F KIAA0966 SAC2 MSTP007 MSTP047 |
| sp\|P28070\|PSB4_HUMAN | 172,978 | 196,883 | 110,017 | 141,274 | sp\|P28070\|PSB4_HUMAN | 143,686 | 30,995 | 22% | P28070 | PSB4_HUMAN | PSB4_HUMAN | Proteasome subunit beta type-4 (26 kDa prosomal protein) (HsBPROS26) (PROS-26) (Macropain beta chain) (Multicatalytic endopeptidase complex beta chain) (Proteasome beta chain) (Proteasome chain 3) (HsN3) | PSMB4 PROS26 |
| sp\|P35221\|CTNA1_HUMAN | 42,691 | 70,027 | 21,997 | 50,206 | sp\|P35221\|CTNA1_HUMAN | 63,355 | 32,544 | 51% | P35221 | CTNA1_HUMAN | CTNA1_HUMAN | Catenin alpha-1 (Alpha E-catenin) (Cadherin-associated protein) (Renal carcinoma antigen NY-REN-13) | CTNNA1 |
| sp\|P78539\|SRPX_HUMAN | 1,015,761 | 1,161,196 | 1,250,293 | 1,209,466 | sp\|P78539\|SRPX_HUMAN | 947,606 | 247,685 | 26% | P78539 | SRPX_HUMAN | SRPX_HUMAN | Sushi repeat-containing protein SRPX | SRPX ETX1 |
| sp\|Q12849\|GRSF1_HUMAN | 67,853 | 70,942 | 117,950 | 97,240 | sp\|Q12849\|GRSF1_HUMAN | 90,096 | 36,587 | 41% | Q12849 | GRSF1_HUMAN | GRSF1_HUMAN | G-rich sequence factor 1 (GRSF-1) | GRSF1 |
| sp\|Q05655\|KPCD_HUMAN | 292,846 | 317,845 | 244,950 | 182,441 | sp\|Q05655\|KPCD_HUMAN | 199,725 | 79,792 | 40% | Q05655 | KPCD_HUMAN | KPCD_HUMAN | Protein kinase C delta type (EC 2.7.11.13) (Tyrosine-protein kinase PRKCD) (EC 2.7.10.2) (nPKC-delta) [Cleaved into: Protein kinase C delta type regulatory subunit; Protein kinase C delta type catalytic subunit (Sphingosine-dependent protein kinase-1) (SDK1)] | PRKCD PKCD |
| sp\|Q15233\|NONO_HUMAN | 115,017 | 104,152 | 172,039 | 116,155 | sp\|Q15233\|NONO_HUMAN | 132,122 | 73,019 | 55% | Q15233 | NONO_HUMAN | NONO_HUMAN | Non-POU domain-containing octamer-binding protein (NonO protein) (54 kDa nuclear RNA- and DNA-binding protein) (p54(nrb)) (p54nrb) (55 kDa nuclear protein) (NMT55) (DNA-binding p52/p100 complex, 52 kDa subunit) | NONO NRB54 |
| sp\|P31937\|3HIDH_HUMAN | 51,592 | 61,759 | 28,157 | 30,085 | sp\|P31937\|3HIDH_HUMAN | 42,614 | 12,365 | 29% | P31937 | 3HIDH_HUMAN | 3HIDH_HUMAN | 3-hydroxyisobutyrate dehydrogenase, mitochondrial (HIBADH) (EC 1.1.1.31) | HIBADH |
| sp\|P55786\|PSA_HUMAN | 92,995 | 123,655 | 76,503 | 158,205 | sp\|P55786\|PSA_HUMAN | 121,085 | 31,668 | 26% | P55786 | PSA_HUMAN | PSA_HUMAN | Puromycin-sensitive aminopeptidase (PSA) (EC 3.4.11.14) (Cytosol alanyl aminopeptidase) (AAP-S) | NPEPPS PSA |
| sp\|Q05682\|CALD1_HUMAN | 8,726 | 20,573 | 22,595 | 31,712 | sp\|Q05682\|CALD1_HUMAN | 23,047 | 9,599 | 42% | Q05682 | CALD1_HUMAN | CALD1_HUMAN | Caldesmon (CDM) | CALD1 CAD CDM |
| sp\|O75964\|ATP5L_HUMAN | 18,409 | 26,699 | 26,337 | 6,032 | sp\|O75964\|ATP5L_HUMAN | 18,860 | 10,051 | 53% | O75964 | ATP5L_HUMAN | ATP5L_HUMAN | ATP synthase subunit g, mitochondrial (ATPase subunit g) (ATP synthase membrane subunit g) | ATP5MG ATP5L |

| | | | | | | | | | | | | | |
|---|---|---|---|---|---|---|---|---|---|---|---|---|---|
| sp\|Q9NUQ9\|FA49B_HUMAN | 87,002 | 119,306 | 82,955 | 110,005 | sp\|Q9NUQ9\|FA49B_HUMAN | 107,008 | 45,692 | 43% | Q9NUQ9 | FA49B_HUMAN | CYRIB_HUMAN | CYFIP-related Rac1 interactor B (L1) | CYRIB CYRI FAM49B BM-009 |
| sp\|O60664\|PLIN3_HUMAN | 54,501 | 34,839 | 58,572 | 42,788 | sp\|O60664\|PLIN3_HUMAN | 53,888 | 9,946 | 18% | O60664 | PLIN3_HUMAN | PLIN3_HUMAN | Perilipin-3 (47 kDa mannose 6-phosphate receptor-binding protein) (47 kDa MPR-binding protein) (Cargo selection protein TIP47) (Mannose-6-phosphate receptor-binding protein 1) (Placental protein 17) (PP17) | PLIN3 M6PRBP1 TIP47 |
| sp\|Q9UBV8\|PEF1_HUMAN | 390,484 | 486,112 | 813,975 | 732,338 | sp\|Q9UBV8\|PEF1_HUMAN | 667,760 | 199,063 | 30% | Q9UBV8 | PEF1_HUMAN | PEF1_HUMAN | Peflin (PEF protein with a long N-terminal hydrophobic domain) (Penta-EF hand domain-containing protein 1) | PEF1 ABP32 UNQ1845/PRO3573 |
| sp\|Q562R1\|ACTBL_HUMAN | 116,600 | 127,235 | 592,298 | 587,354 | sp\|Q562R1\|ACTBL_HUMAN | 289,252 | 220,968 | 76% | Q562R1 | ACTBL_HUMAN | ACTBL_HUMAN | Beta-actin-like protein 2 (Kappa-actin) | ACTBL2 |
| sp\|P56192\|SYMC_HUMAN | 61,580 | 107,154 | 47,916 | 111,311 | sp\|P56192\|SYMC_HUMAN | 84,528 | 27,114 | 32% | P56192 | SYMC_HUMAN | SYMC_HUMAN | Methionine--tRNA ligase, cytoplasmic (EC 6.1.1.10) (Methionyl-tRNA synthetase) (MetRS) | MARS1 MARS |
| sp\|P24534\|EF1B_HUMAN | 156,525 | 180,404 | 102,638 | 97,574 | sp\|P24534\|EF1B_HUMAN | 112,525 | 40,173 | 36% | P24534 | EF1B_HUMAN | EF1B_HUMAN | Elongation factor 1-beta (EF-1-beta) | EEF1B2 EEF1B EF1B |
| sp\|P62910\|RL32_HUMAN | 399,996 | 524,896 | 482,441 | 394,522 | sp\|P62910\|RL32_HUMAN | 350,839 | 128,127 | 37% | P62910 | RL32_HUMAN | RL32_HUMAN | 60S ribosomal protein L32 (Large ribosomal subunit protein eL32) | RPL32 PP9932 |
| sp\|P20036\|DPA1_HUMAN | 1,287,647 | 2,386,036 | 651,632 | 1,636,747 | sp\|P20036\|DPA1_HUMAN | 1,654,568 | 532,639 | 32% | P20036 | DPA1_HUMAN | DPA1_HUMAN | HLA class II histocompatibility antigen, DP alpha 1 chain (DP(W3)) (DP(W4)) (HLA-SB alpha chain) (MHC class II DP3-alpha) (MHC class II DPA1) | HLA-DPA1 HLA-DP1A HLASB |
| sp\|Q8N0U8\|VKORL_HUMAN | 10,268 | 17,317 | 2,721 | 9,481 | sp\|Q8N0U8\|VKORL_HUMAN | 8,890 | 4,859 | 55% | Q8N0U8 | VKORL_HUMAN | VKORL_HUMAN | Vitamin K epoxide reductase complex subunit 1-like protein 1 (VKORC1-like protein 1) (EC 1.17.4.4) | VKORC1L1 |
| sp\|P30046\|DOPD_HUMAN | 149,291 | 234,496 | 60,810 | 130,154 | sp\|P30046\|DOPD_HUMAN | 120,044 | 71,707 | 60% | P30046 | DOPD_HUMAN | DOPD_HUMAN | D-dopachrome decarboxylase (EC 4.1.1.84) (D-dopachrome tautomerase) (Phenylpyruvate tautomerase II) | DDT |
| sp\|P15531\|NDKA_HUMAN | 150,406 | 284,990 | 116,531 | 44,476 | sp\|P15531\|NDKA_HUMAN | 157,076 | 78,675 | 50% | P15531 | NDKA_HUMAN | NDKA_HUMAN | Nucleoside diphosphate kinase A (NDK A) (NDP kinase A) (EC 2.7.4.6) (Granzyme A-activated DNase) (GAAD) (Metastasis inhibition factor nm23) (NM23-H1) (Tumor metastatic process-associated protein) | NME1 NDPKA NM23 |
| sp\|P62841\|RS15_HUMAN | 84,816 | 112,646 | 52,939 | 72,827 | sp\|P62841\|RS15_HUMAN | 86,276 | 24,195 | 28% | P62841 | RS15_HUMAN | RS15_HUMAN | 40S ribosomal protein S15 (RIG protein) (Small ribosomal subunit protein uS19) | RPS15 RIG |
| sp\|P36507\|MP2K2_HUMAN | 72,608 | 69,098 | 71,288 | 35,906 | sp\|P36507\|MP2K2_HUMAN | 64,625 | 23,344 | 36% | P36507 | MP2K2_HUMAN | MP2K2_HUMAN | Dual specificity mitogen-activated protein kinase kinase 2 (MAP kinase kinase 2) (MAPKK 2) (EC 2.7.12.2) (ERK activator kinase 2) (MAPK/ERK kinase 2) (MEK 2) | MAP2K2 MEK2 MKK2 PRKMK2 |
| sp\|Q15185\|TEBP_HUMAN | 133,893 | 173,092 | 65,435 | 95,693 | sp\|Q15185\|TEBP_HUMAN | 109,226 | 44,456 | 41% | Q15185 | TEBP_HUMAN | TEBP_HUMAN | Prostaglandin E synthase 3 (EC 5.3.99.3) (Cytosolic prostaglandin E2 synthase) (cPGES) (Hsp90 co-chaperone) (Progesterone receptor complex p23) (Telomerase-binding protein p23) | PTGES3 P23 TEBP |
| sp\|P61513\|RL37A_HUMAN | 162,769 | 190,627 | 94,584 | 71,522 | sp\|P61513\|RL37A_HUMAN | 99,343 | 52,586 | 53% | P61513 | RL37A_HUMAN | RL37A_HUMAN | 60S ribosomal protein L37a (Large ribosomal subunit protein eL43) | RPL37A |

| | | | | | | | | | | | | | |
|---|---|---|---|---|---|---|---|---|---|---|---|---|---|
| sp\|Q9BUL8\|PDC10_HUMAN | 47,003 | 63,094 | 63,580 | 30,025 | sp\|Q9BUL8\|PDC10_HUMAN | 51,911 | 12,063 | 23% | Q9BUL8 | PDC10_HUMAN | PDC10_HUMAN | Programmed cell death protein 10 (Cerebral cavernous malformations 3 protein) (TF-1 cell apoptosis-related protein 15) | PDCD10 CCM3 TFAR15 |
| sp\|P35268\|RL22_HUMAN | 577,557 | 626,007 | 416,963 | 392,956 | sp\|P35268\|RL22_HUMAN | 391,380 | 150,372 | 38% | P35268 | RL22_HUMAN | RL22_HUMAN | 60S ribosomal protein L22 (EBER-associated protein) (EAP) (Epstein-Barr virus small RNA-associated protein) (Heparin-binding protein HBp15) (Large ribosomal subunit protein eL22) | RPL22 |
| sp\|Q16186\|ADRM1_HUMAN | 98,239 | 109,078 | 109,542 | 140,813 | sp\|Q16186\|ADRM1_HUMAN | 106,451 | 15,795 | 15% | Q16186 | ADRM1_HUMAN | ADRM1_HUMAN | Proteasomal ubiquitin receptor ADRM1 (110 kDa cell membrane glycoprotein) (Gp110) (Adhesion-regulating molecule 1) (ARM-1) (Proteasome regulatory particle non-ATPase 13) (hRpn13) (Rpn13 homolog) | ADRM1 GP110 |
| sp\|Q13308\|PTK7_HUMAN | 223,028 | 228,774 | 419,693 | 289,906 | sp\|Q13308\|PTK7_HUMAN | 333,417 | 108,404 | 33% | Q13308 | PTK7_HUMAN | PTK7_HUMAN | Inactive tyrosine-protein kinase 7 (Colon carcinoma kinase 4) (CCK-4) (Protein-tyrosine kinase 7) (Pseudo tyrosine kinase receptor 7) (Tyrosine-protein kinase-like 7) | PTK7 CCK4 |
| sp\|P02765\|FETUA_HUMAN | 462,669 | 218,220 | 512,543 | 153,629 | sp\|P02765\|FETUA_HUMAN | 2,225,514 | 2,167,861 | 97% | P02765 | FETUA_HUMAN | FETUA_HUMAN | Alpha-2-HS-glycoprotein (Alpha-2-Z-globulin) (Ba-alpha-2-glycoprotein) (Fetuin-A) [Cleaved into: Alpha-2-HS-glycoprotein chain A; Alpha-2-HS-glycoprotein chain B] | AHSG FETUA PRO2743 |
| sp\|Q9BVK6\|TMED9_HUMAN | 61,882 | 56,854 | 41,695 | 67,325 | sp\|Q9BVK6\|TMED9_HUMAN | 48,794 | 25,694 | 53% | Q9BVK6 | TMED9_HUMAN | TMED9_HUMAN | Transmembrane emp24 domain-containing protein 9 (GMP25) (Glycoprotein 25L2) (p24 family protein alpha-2) (p24alpha2) (p25) | TMED9 GP25L2 |
| sp\|P14550\|AK1A1_HUMAN | 122,739 | 153,536 | 41,354 | 122,250 | sp\|P14550\|AK1A1_HUMAN | 161,039 | 65,117 | 40% | P14550 | AK1A1_HUMAN | AK1A1_HUMAN | Aldo-keto reductase family 1 member A1 (EC 1.1.1.2) (EC 1.1.1.372) (EC 1.1.1.54) (Alcohol dehydrogenase [NADP(+)]) (Aldehyde reductase) (Glucuronate reductase) (EC 1.1.1.19) (Glucuronolactone reductase) (EC 1.1.1.20) | AKR1A1 ALDR1 ALR |
| sp\|P13073\|COX41_HUMAN | 43,742 | 44,795 | 28,460 | 16,711 | sp\|P13073\|COX41_HUMAN | 25,793 | 13,272 | 51% | P13073 | COX41_HUMAN | COX41_HUMAN | Cytochrome c oxidase subunit 4 isoform 1, mitochondrial (Cytochrome c oxidase polypeptide IV) (Cytochrome c oxidase subunit IV isoform 1) (COX IV-1) | COX4I1 COX4 |
| sp\|P04156\|PRIO_HUMAN | 248,340 | 237,677 | 457,790 | 358,601 | sp\|P04156\|PRIO_HUMAN | 264,292 | 112,484 | 43% | P04156 | PRIO_HUMAN | PRIO_HUMAN | Major prion protein (PrP) (ASCR) (PrP27-30) (PrP33-35C) (CD antigen CD230) | PRNP ALTPRP PRIP PRP |
| sp\|Q05639\|EF1A2_HUMAN | 134,275 | 66,127 | 68,254 | 92,195 | sp\|Q05639\|EF1A2_HUMAN | 85,482 | 36,172 | 42% | Q05639 | EF1A2_HUMAN | EF1A2_HUMAN | Elongation factor 1-alpha 2 (EF-1-alpha-2) (Eukaryotic elongation factor 1 A-2) (eEF1A-2) (Statin-S1) | EEF1A2 EEF1AL STN |
| sp\|P60866\|RS20_HUMAN | 1,372,458 | 1,418,076 | 1,066,173 | 776,379 | sp\|P60866\|RS20_HUMAN | 819,429 | 421,389 | 51% | P60866 | RS20_HUMAN | RS20_HUMAN | 40S ribosomal protein S20 (Small ribosomal subunit protein uS10) | RPS20 |
| sp\|P46776\|RL27A_HUMAN | 1,621,897 | 1,378,655 | 1,186,221 | 1,028,757 | sp\|P46776\|RL27A_HUMAN | 892,032 | 476,628 | 53% | P46776 | RL27A_HUMAN | RL27A_HUMAN | 60S ribosomal protein L27a (Large ribosomal subunit protein uL15) | RPL27A |
| sp\|P26196\|DDX6_HUMAN | 137,143 | 175,048 | 142,361 | 199,951 | sp\|P26196\|DDX6_HUMAN | 182,415 | 65,547 | 36% | P26196 | DDX6_HUMAN | DDX6_HUMAN | Probable ATP-dependent RNA helicase DDX6 (EC 3.6.4.13) (ATP-dependent RNA helicase p54) (DEAD box protein 6) (Oncogene RCK) | DDX6 HLR2 RCK |

| | | | | | | | | | | | | | |
|---|---|---|---|---|---|---|---|---|---|---|---|---|---|
| sp\|P57729\|RAB38_HUMAN | 560,844 | 603,463 | 505,124 | 599,570 | sp\|P57729\|RAB38_HUMAN | 439,377 | 142,740 | 32% | P57729 | RAB38_HUMAN | RAB38_HUMAN | Ras-related protein Rab-38 (Melanoma antigen NY-MEL-1) | RAB38 |
| sp\|Q9NZW5\|MPP6_HUMAN | 368,890 | 386,539 | 383,962 | 426,634 | sp\|Q9NZW5\|MPP6_HUMAN | 471,653 | 224,833 | 48% | Q9NZW5 | MPP6_HUMAN | PALS2_HUMAN | Protein PALS2 (MAGUK p55 subfamily member 6) (Membrane protein, palmitoylated 6) (Veli-associated MAGUK 1) (VAM-1) | PALS2 MPP6 VAM1 |
| sp\|O00487\|PSDE_HUMAN | 150,081 | 139,690 | 111,185 | 115,922 | sp\|O00487\|PSDE_HUMAN | 104,497 | 29,520 | 28% | O00487 | PSDE_HUMAN | PSDE_HUMAN | 26S proteasome non-ATPase regulatory subunit 14 (EC 3.4.19.-) (26S proteasome regulatory subunit RPN11) (26S proteasome-associated PAD1 homolog 1) | PSMD14 POH1 |
| sp\|P78344\|IF4G2_HUMAN | 51,128 | 74,309 | 110,246 | 36,424 | sp\|P78344\|IF4G2_HUMAN | 57,891 | 32,864 | 57% | P78344 | IF4G2_HUMAN | IF4G2_HUMAN | Eukaryotic translation initiation factor 4 gamma 2 (eIF-4-gamma 2) (eIF-4G 2) (eIF4G 2) (Death-associated protein 5) (DAP-5) (p97) | EIF4G2 DAP5 OK/SW-cl.75 |
| sp\|P25391\|LAMA1_HUMAN | 195,158 | 171,454 | 158,025 | 302,264 | sp\|P25391\|LAMA1_HUMAN | 231,939 | 76,247 | 33% | P25391 | LAMA1_HUMAN | LAMA1_HUMAN | Laminin subunit alpha-1 (Laminin A chain) (Laminin-1 subunit alpha) (Laminin-3 subunit alpha) (S-laminin subunit alpha) (S-LAM alpha) | LAMA1 LAMA |
| sp\|Q8NBI5\|S43A3_HUMAN | 1,401,575 | 1,701,725 | 1,383,630 | 2,014,809 | sp\|Q8NBI5\|S43A3_HUMAN | 1,599,694 | 346,671 | 22% | Q8NBI5 | S43A3_HUMAN | S43A3_HUMAN | Equilibrative nucleobase transporter 1 (Protein FOAP-13) (Solute carrier family 43 member 3) | SLC43A3 ENBT1 PSEC0252 |
| sp\|P09486\|SPRC_HUMAN | 60,090 | 34,671 | 103,362 | 61,914 | sp\|P09486\|SPRC_HUMAN | 52,332 | 23,232 | 44% | P09486 | SPRC_HUMAN | SPRC_HUMAN | SPARC (Basement-membrane protein 40) (BM-40) (Osteonectin) (ON) (Secreted protein acidic and rich in cysteine) | SPARC ON |
| sp\|Q9BXW7\|HDHD5_HUMAN | 963 | 13,376 | 4,272 | 23,824 | sp\|Q9BXW7\|HDHD5_HUMAN | 14,392 | 11,376 | 79% | Q9BXW7 | HDHD5_HUMAN | HDHD5_HUMAN | Haloacid dehalogenase-like hydrolase domain-containing 5 (Cat eye syndrome critical region protein 5) | HDHD5 CECR5 |
| sp\|Q04760\|LGUL_HUMAN | 257,604 | 296,689 | 213,680 | 283,669 | sp\|Q04760\|LGUL_HUMAN | 233,962 | 107,687 | 46% | Q04760 | LGUL_HUMAN | LGUL_HUMAN | Lactoylglutathione lyase (EC 4.4.1.5) (Aldoketomutase) (Glyoxalase I) (Glx I) (Ketone-aldehyde mutase) (Methylglyoxalase) (S-D-lactoylglutathione methylglyoxal lyase) | GLO1 |
| sp\|P26006\|ITA3_HUMAN | 487,831 | 438,700 | 625,532 | 618,464 | sp\|P26006\|ITA3_HUMAN | 590,839 | 232,223 | 39% | P26006 | ITA3_HUMAN | ITA3_HUMAN | Integrin alpha-3 (CD49 antigen-like family member C) (FRP-2) (Galactoprotein B3) (GAPB3) (VLA-3 subunit alpha) (CD antigen CD49c) [Cleaved into: Integrin alpha-3 heavy chain; Integrin alpha-3 light chain] | ITGA3 MSK18 |
| sp\|O75954\|TSN9_HUMAN | 552,905 | 539,884 | 266,613 | 439,518 | sp\|O75954\|TSN9_HUMAN | 379,458 | 118,627 | 31% | O75954 | TSN9_HUMAN | TSN9_HUMAN | Tetraspanin-9 (Tspan-9) (Tetraspan NET-5) | TSPAN9 NET5 |
| sp\|P68431\|H31_HUMAN | 1,473 | 558 | 1,507 | 5,664 | sp\|P68431\|H31_HUMAN | 2,982 | 2,788 | 93% | P68431 | H31_HUMAN | H31_HUMAN | Histone H3.1 (Histone H3/a) (Histone H3/b) (Histone H3/c) (Histone H3/d) (Histone H3/f) (Histone H3/h) (Histone H3/i) (Histone H3/j) (Histone H3/k) (Histone H3/l) | H3C1 H3FA HIST1H3A; H3C2 H3FL HIST1H3B; H3C3 H3FC HIST1H3C; H3C4 H3FB HIST1H3D; H3C6 H3FD HIST1H3E; H3C7 H3FI HIST1H3F; H3C8 H3FH HIST1H3G; H3C10 H3FK HIST1H3H; H3C11 H3FF HIST1H3I; H3C12 H3FJ HIST1H3J |
| sp\|P42224\|STAT1_HUMAN | 118,145 | 136,521 | 78,327 | 106,786 | sp\|P42224\|STAT1_HUMAN | 117,685 | 32,850 | 28% | P42224 | STAT1_HUMAN | STAT1_HUMAN | Signal transducer and activator of transcription 1-alpha/beta (Transcription factor ISGF-3 components p91/p84) | STAT1 |
| sp\|Q8NE71\|ABCF1_HUMAN | 146,752 | 126,699 | 249,366 | 206,641 | sp\|Q8NE71\|ABCF1_HUMAN | 152,204 | 51,219 | 34% | Q8NE71 | ABCF1_HUMAN | ABCF1_HUMAN | ATP-binding cassette sub-family F member 1 (ATP-binding cassette 50) (TNF-alpha-stimulated ABC protein) | ABCF1 ABC50 |

| | | | | | | | | | | | | | |
|---|---|---|---|---|---|---|---|---|---|---|---|---|---|
| sp\|P49755\|TMEDA_HUMAN | 68,777 | 55,923 | 58,552 | 28,343 | sp\|P49755\|TMEDA_HUMAN | 35,915 | 22,277 | 62% | P49755 | TMEDA_HUMAN | TMEDA_HUMAN | Transmembrane emp24 domain-containing protein 10 (Protein TMED10) (21 kDa transmembrane-trafficking protein) (S31I125) (S31III125) (Tmp-21-I) (Transmembrane protein Tmp21) (p23) (p24 family protein delta-1) (p24delta1) (p24delta) | TMED10 TMP21 |
| sp\|Q9ULF5\|S39AA_HUMAN | 282,430 | 308,736 | 226,559 | 255,917 | sp\|Q9ULF5\|S39AA_HUMAN | 250,086 | 51,597 | 21% | Q9ULF5 | S39AA_HUMAN | S39AA_HUMAN | Zinc transporter ZIP10 (Solute carrier family 39 member 10) (Zrt- and Irt-like protein 10) (ZIP-10) | SLC39A10 KIAA1265 ZIP10 |
| sp\|O43837\|IDH3B_HUMAN | 32,521 | 25,270 | 38,460 | 70,687 | sp\|O43837\|IDH3B_HUMAN | 36,022 | 17,738 | 49% | O43837 | IDH3B_HUMAN | IDH3B_HUMAN | Isocitrate dehydrogenase [NAD] subunit beta, mitochondrial (Isocitric dehydrogenase subunit beta) (NAD(+)-specific ICDH subunit beta) | IDH3B |
| sp\|Q92890\|UFD1_HUMAN | 126,765 | 152,795 | 89,124 | 86,984 | sp\|Q92890\|UFD1_HUMAN | 153,415 | 55,543 | 36% | Q92890 | UFD1_HUMAN | UFD1_HUMAN | Ubiquitin recognition factor in ER-associated degradation protein 1 (Ubiquitin fusion degradation protein 1) (UB fusion protein 1) | UFD1 UFD1L |
| sp\|Q9UBB4\|ATX10_HUMAN | 48,015 | 106,557 | 62,538 | 177,287 | sp\|Q9UBB4\|ATX10_HUMAN | 93,369 | 50,425 | 54% | Q9UBB4 | ATX10_HUMAN | ATX10_HUMAN | Ataxin-10 (Brain protein E46 homolog) (Spinocerebellar ataxia type 10 protein) | ATXN10 SCA10 |
| sp\|Q9UBR2\|CATZ_HUMAN | 191,410 | 127,984 | 372,352 | 192,971 | sp\|Q9UBR2\|CATZ_HUMAN | 180,443 | 82,781 | 46% | Q9UBR2 | CATZ_HUMAN | CATZ_HUMAN | Cathepsin Z (EC 3.4.18.1) (Cathepsin P) (Cathepsin X) | CTSZ |
| sp\|P47813\|IF1AX_HUMAN | 13,856 | 33,384 | 44,662 | 63,929 | sp\|P47813\|IF1AX_HUMAN | 45,902 | 25,323 | 55% | P47813 | IF1AX_HUMAN | IF1AX_HUMAN | Eukaryotic translation initiation factor 1A, X-chromosomal (eIF-1A X isoform) (Eukaryotic translation initiation factor 4C) (eIF-4C) | EIF1AX EIF1A EIF4C |
| sp\|P31150\|GDIA_HUMAN | 196,187 | 219,102 | 156,958 | 244,857 | sp\|P31150\|GDIA_HUMAN | 212,424 | 34,231 | 16% | P31150 | GDIA_HUMAN | GDIA_HUMAN | Rab GDP dissociation inhibitor alpha (Rab GDI alpha) (Guanosine diphosphate dissociation inhibitor 1) (GDI-1) (Oligophrenin-2) (Protein XAP-4) | GDI1 GDIL OPHN2 RABGDIA XAP4 |
| sp\|Q14696\|MESD_HUMAN | 26,128 | 28,986 | 63,930 | 28,009 | sp\|Q14696\|MESD_HUMAN | 39,343 | 23,595 | 60% | Q14696 | MESD_HUMAN | MESD_HUMAN | LRP chaperone MESD (LDLR chaperone MESD) (Mesoderm development LRP chaperone MESD) (Mesoderm development candidate 2) (Mesoderm development protein) (Renal carcinoma antigen NY-REN-61) | MESD KIAA0081 MESDC2 MESDM UNQ1911/PRO4369 |
| sp\|O14773\|TPP1_HUMAN | 40,068 | 31,694 | 40,234 | 65,452 | sp\|O14773\|TPP1_HUMAN | 37,968 | 25,051 | 66% | O14773 | TPP1_HUMAN | TPP1_HUMAN | Tripeptidyl-peptidase 1 (TPP-1) (EC 3.4.14.9) (Cell growth-inhibiting gene 1 protein) (Lysosomal pepstatin-insensitive protease) (LPIC) (Tripeptidyl aminopeptidase) (Tripeptidyl-peptidase I) (TPP-I) | TPP1 CLN2 GIG1 UNQ267/PRO304 |
| sp\|Q9H0U4\|RAB1B_HUMAN | 242,440 | 271,137 | 102,512 | 209,688 | sp\|Q9H0U4\|RAB1B_HUMAN | 206,222 | 51,148 | 25% | Q9H0U4 | RAB1B_HUMAN | RAB1B_HUMAN | Ras-related protein Rab-1B (EC 3.6.5.2) | RAB1B |
| sp\|Q9NS69\|TOM22_HUMAN | 4,890 | 50,206 | 8,393 | 66,054 | sp\|Q9NS69\|TOM22_HUMAN | 23,343 | 22,820 | 98% | Q9NS69 | TOM22_HUMAN | TOM22_HUMAN | Mitochondrial import receptor subunit TOM22 homolog (hTom22) (1C9-2) (Translocase of outer membrane 22 kDa subunit homolog) | TOMM22 TOM22 |
| sp\|Q9NQ88\|TIGAR_HUMAN | 1,252 | 62,880 | 23,090 | 55,888 | sp\|Q9NQ88\|TIGAR_HUMAN | 36,354 | 20,738 | 57% | Q9NQ88 | TIGAR_HUMAN | TIGAR_HUMAN | Fructose-2,6-bisphosphatase TIGAR (EC 3.1.3.46) (TP53-induced glycolysis and apoptosis regulator) (TP53-induced glycolysis regulatory phosphatase) | TIGAR C12orf5 |

| | | | | | | | | | | | | | |
|---|---|---|---|---|---|---|---|---|---|---|---|---|---|
| sp\|Q96C19\|EFHD2_HUMAN | 50,995 | 35,704 | 52,537 | 65,668 | sp\|Q96C19\|EFHD2_HUMAN | 41,940 | 15,721 | 37% | Q96C19 | EFHD2_HUMAN | EFHD2_HUMAN | EF-hand domain-containing protein D2 (Swiprosin-1) | EFHD2 SWS1 |
| sp\|Q14103\|HNRPD_HUMAN | 314,806 | 272,454 | 168,064 | 49,835 | sp\|Q14103\|HNRPD_HUMAN | 141,625 | 101,535 | 72% | Q14103 | HNRPD_HUMAN | HNRPD_HUMAN | Heterogeneous nuclear ribonucleoprotein D0 (hnRNP D0) (AU-rich element RNA-binding protein 1) | HNRNPD AUF1 HNRPD |
| sp\|P67812\|SC11A_HUMAN | 48,768 | 13,369 | 21,459 | 123,670 | sp\|P67812\|SC11A_HUMAN | 41,830 | 36,661 | 88% | P67812 | SC11A_HUMAN | SC11A_HUMAN | Signal peptidase complex catalytic subunit SEC11A (EC 3.4.21.89) (Endopeptidase SP18) (Microsomal signal peptidase 18 kDa subunit) (SPase 18 kDa subunit) (SEC11 homolog A) (SEC11-like protein 1) (SPC18) | SEC11A SEC11L1 SPC18 SPCS4A |
| sp\|P62942\|FKB1A_HUMAN | 424,899 | 391,173 | 541,571 | 728,484 | sp\|P62942\|FKB1A_HUMAN | 488,847 | 118,852 | 24% | P62942 | FKB1A_HUMAN | FKB1A_HUMAN | Peptidyl-prolyl cis-trans isomerase FKBP1A (PPIase FKBP1A) (EC 5.2.1.8) (12 kDa FK506-binding protein) (12 kDa FKBP) (FKBP-12) (Calstabin-1) (FK506-binding protein 1A) (FKBP-1A) (Immunophilin FKBP12) (Rotamase) | FKBP1A FKBP1 FKBP12 |
| sp\|P62857\|RS28_HUMAN | 156,418 | 173,516 | 390,928 | 497,971 | sp\|P62857\|RS28_HUMAN | 295,975 | 142,277 | 48% | P62857 | RS28_HUMAN | RS28_HUMAN | 40S ribosomal protein S28 (Small ribosomal subunit protein eS28) | RPS28 |
| sp\|P62847\|RS24_HUMAN | 497,798 | 492,304 | 112,468 | 362,568 | sp\|P62847\|RS24_HUMAN | 264,225 | 169,085 | 64% | P62847 | RS24_HUMAN | RS24_HUMAN | 40S ribosomal protein S24 (Small ribosomal subunit protein eS24) | RPS24 |
| sp\|P61803\|DAD1_HUMAN | 38,080 | 35,575 | 37,784 | 65,025 | sp\|P61803\|DAD1_HUMAN | 37,761 | 15,108 | 40% | P61803 | DAD1_HUMAN | DAD1_HUMAN | Dolichyl-diphosphooligosaccharide--protein glycosyltransferase subunit DAD1 (Oligosaccharyl transferase subunit DAD1) (Defender against cell death 1) (DAD-1) | DAD1 |
| sp\|P43307\|SSRA_HUMAN | 97,518 | 75,779 | 115,056 | 71,600 | sp\|P43307\|SSRA_HUMAN | 82,347 | 18,722 | 23% | P43307 | SSRA_HUMAN | SSRA_HUMAN | Translocon-associated protein subunit alpha (TRAP-alpha) (Signal sequence receptor subunit alpha) (SSR-alpha) | SSR1 TRAPA PSEC0262 |
| sp\|P04080\|CYTB_HUMAN | 230,330 | 464,584 | 134,487 | 305,781 | sp\|P04080\|CYTB_HUMAN | 275,297 | 146,035 | 53% | P04080 | CYTB_HUMAN | CYTB_HUMAN | Cystatin-B (CPI-B) (Liver thiol proteinase inhibitor) (Stefin-B) | CSTB CST6 STFB |
| sp\|O14880\|MGST3_HUMAN | 47,765 | 60,428 | 18,272 | 157,124 | sp\|O14880\|MGST3_HUMAN | 60,182 | 44,417 | 74% | O14880 | MGST3_HUMAN | MGST3_HUMAN | Microsomal glutathione S-transferase 3 (Microsomal GST-3) (Glutathione peroxidase MGST3) (EC 1.11.1.-) (LTC4 synthase MGST3) (EC 4.4.1.20) (Microsomal glutathione S-transferase III) (Microsomal GST-III) | MGST3 |
| sp\|O14735\|CDIPT_HUMAN | 78,856 | 86,393 | 35,159 | 62,372 | sp\|O14735\|CDIPT_HUMAN | 60,621 | 16,720 | 28% | O14735 | CDIPT_HUMAN | CDIPT_HUMAN | CDP-diacylglycerol--inositol 3-phosphatidyltransferase (EC 2.7.8.11) (Phosphatidylinositol synthase) (PI synthase) (PtdIns synthase) | CDIPT PIS PIS1 |
| sp\|P10768\|ESTD_HUMAN | 157,964 | 175,195 | 92,126 | 97,228 | sp\|P10768\|ESTD_HUMAN | 117,735 | 36,862 | 31% | P10768 | ESTD_HUMAN | ESTD_HUMAN | S-formylglutathione hydrolase (FGH) (EC 3.1.2.12) (Esterase D) (Methylumbelliferyl-acetate deacetylase) (EC 3.1.1.56) | ESD |
| sp\|Q9UBI6\|GBG12_HUMAN | 420,689 | 776,415 | 337,570 | 486,083 | sp\|Q9UBI6\|GBG12_HUMAN | 445,331 | 146,864 | 33% | Q9UBI6 | GBG12_HUMAN | GBG12_HUMAN | Guanine nucleotide-binding protein G(I)/G(S)/G(O) subunit gamma-12 | GNG12 |
| sp\|P27701\|CD82_HUMAN | 345,853 | 587,291 | 261,502 | 323,104 | sp\|P27701\|CD82_HUMAN | 356,154 | 129,906 | 36% | P27701 | CD82_HUMAN | CD82_HUMAN | CD82 antigen (C33 antigen) (IA4) (Inducible membrane protein R2) (Metastasis suppressor Kangai-1) (Suppressor of tumorigenicity 6 protein) (Tetraspanin-27) (Tspan-27) (CD antigen CD82) | CD82 KAI1 SAR2 ST6 TSPAN27 |

| | | | | | | | | | | | | | |
|---|---|---|---|---|---|---|---|---|---|---|---|---|---|
| sp\|P55735\|SEC13_HUMAN | 124,848 | 103,220 | 78,468 | 90,450 | sp\|P55735\|SEC13_HUMAN | 87,259 | 24,429 | 28% | P55735 | SEC13_HUMAN | SEC13_HUMAN | Protein SEC13 homolog (GATOR complex protein SEC13) (SEC13-like protein 1) (SEC13-related protein) | SEC13 D3S1231E SEC13A SEC13L1 SEC13R |
| sp\|O60841\|IF2P_HUMAN | 63,879 | 100,861 | 91,609 | 100,058 | sp\|O60841\|IF2P_HUMAN | 94,613 | 20,080 | 21% | O60841 | IF2P_HUMAN | IF2P_HUMAN | Eukaryotic translation initiation factor 5B (eIF-5B) (EC 3.6.5.3) (Translation initiation factor IF-2) | EIF5B IF2 KIAA0741 |
| sp\|P61221\|ABCE1_HUMAN | 10,374 | 30,550 | 22,038 | 45,620 | sp\|P61221\|ABCE1_HUMAN | 25,478 | 10,156 | 40% | P61221 | ABCE1_HUMAN | ABCE1_HUMAN | ATP-binding cassette sub-family E member 1 (2'-5'-oligoadenylate-binding protein) (HuHP68) (RNase L inhibitor) (Ribonuclease 4 inhibitor) (RNS4I) | ABCE1 RLI RNASEL1 RNASELI RNS4I OK/SW-cl.40 |
| sp\|P49591\|SYSC_HUMAN | 177,691 | 185,238 | 124,314 | 214,140 | sp\|P49591\|SYSC_HUMAN | 140,373 | 62,184 | 44% | P49591 | SYSC_HUMAN | SYSC_HUMAN | Serine--tRNA ligase, cytoplasmic (EC 6.1.1.11) (Seryl-tRNA synthetase) (SerRS) (Seryl-tRNA(Ser/Sec) synthetase) | SARS1 SARS SERS |
| sp\|P50416\|CPT1A_HUMAN | 14,864 | 14,979 | 133,208 | 14,052 | sp\|P50416\|CPT1A_HUMAN | 62,955 | 53,127 | 84% | P50416 | CPT1A_HUMAN | CPT1A_HUMAN | Carnitine O-palmitoyltransferase 1, liver isoform (CPT1-L) (EC 2.3.1.21) (Carnitine O-palmitoyltransferase I, liver isoform) (CPT I) (CPTI-L) (Carnitine palmitoyltransferase 1A) | CPT1A CPT1 |
| sp\|P40967\|PMEL_HUMAN | 23,709 | 52,398 | 54,366 | 22,939 | sp\|P40967\|PMEL_HUMAN | 212,607 | 236,640 | 111% | P40967 | PMEL_HUMAN | PMEL_HUMAN | Melanocyte protein PMEL (ME20-M) (ME20M) (Melanocyte protein Pmel 17) (Melanocytes lineage-specific antigen GP100) (Melanoma-associated ME20 antigen) (P1) (P100) (Premelanosome protein) (Silver locus protein homolog) [Cleaved into: M-alpha (95 kDa melanocyte-specific secreted glycoprotein) (P26) (Secreted melanoma-associated ME20 antigen) (ME20-S) (ME20S); M-beta] | PMEL D12S53E PMEL17 SILV |
| sp\|P62750\|RL23A_HUMAN | 177,672 | 260,245 | 111,848 | 125,552 | sp\|P62750\|RL23A_HUMAN | 134,152 | 62,945 | 47% | P62750 | RL23A_HUMAN | RL23A_HUMAN | 60S ribosomal protein L23a (Large ribosomal subunit protein uL23) | RPL23A |
| sp\|P10515\|ODP2_HUMAN | 6,460 | 10,171 | 7,616 | 4,307 | sp\|P10515\|ODP2_HUMAN | 12,006 | 7,738 | 64% | P10515 | ODP2_HUMAN | ODP2_HUMAN | Dihydrolipoyllysine-residue acetyltransferase component of pyruvate dehydrogenase complex, mitochondrial (EC 2.3.1.12) (70 kDa mitochondrial autoantigen of primary biliary cirrhosis) (PBC) (Dihydrolipoamide acetyltransferase component of pyruvate dehydrogenase complex) (M2 antigen complex 70 kDa subunit) (Pyruvate dehydrogenase complex component E2) (PDC-E2) (PDCE2) | DLAT DLTA |
| sp\|P35270\|SPRE_HUMAN | 117,250 | 114,419 | 151,927 | 247,507 | sp\|P35270\|SPRE_HUMAN | 176,636 | 56,683 | 32% | P35270 | SPRE_HUMAN | SPRE_HUMAN | Sepiapterin reductase (SPR) (EC 1.1.1.153) | SPR |
| sp\|Q9Y2B0\|CNPY2_HUMAN | 10,782 | 26,963 | 16,153 | 53,307 | sp\|Q9Y2B0\|CNPY2_HUMAN | 22,951 | 13,808 | 60% | Q9Y2B0 | CNPY2_HUMAN | CNPY2_HUMAN | Protein canopy homolog 2 (MIR-interacting saposin-like protein) (Putative secreted protein Zsig9) (Transmembrane protein 4) | CNPY2 MSAP TMEM4 ZSIG9 UNQ1943/PRO4426 |
| sp\|Q13404\|UB2V1_HUMAN | 303,701 | 352,407 | 263,052 | 331,805 | sp\|Q13404\|UB2V1_HUMAN | 316,127 | 54,182 | 17% | Q13404 | UB2V1_HUMAN | UB2V1_HUMAN | Ubiquitin-conjugating enzyme E2 variant 1 (UEV-1) (CROC-1) (TRAF6-regulated IKK activator 1 beta Uev1A) | UBE2V1 CROC1 UBE2V UEV1 P/OKcl.19 |

| | | | | | | | | | | | | | |
|---|---|---|---|---|---|---|---|---|---|---|---|---|---|
| sp\|P31942\|HNRH3_HUMAN | 52,457 | 55,898 | 35,020 | 19,249 | sp\|P31942\|HNRH3_HUMAN | 34,156 | 13,805 | 40% | P31942 | HNRH3_HUMAN | HNRH3_HUMAN | Heterogeneous nuclear ribonucleoprotein H3 (hnRNP H3) (Heterogeneous nuclear ribonucleoprotein 2H9) (hnRNP 2H9) | HNRNPH3 HNRPH3 |
| sp\|Q9Y295\|DRG1_HUMAN | 134,935 | 136,232 | 49,937 | 111,837 | sp\|Q9Y295\|DRG1_HUMAN | 95,782 | 36,022 | 38% | Q9Y295 | DRG1_HUMAN | DRG1_HUMAN | Developmentally-regulated GTP-binding protein 1 (DRG-1) (Neural precursor cell expressed developmentally down-regulated protein 3) (NEDD-3) (Translation factor GTPase DRG1) (TRAFAC GTPase DRG1) (EC 3.6.5.-) | DRG1 NEDD3 |
| sp\|Q96RP9\|EFGM_HUMAN | 5,661 | 21,560 | 23,890 | 159,994 | sp\|Q96RP9\|EFGM_HUMAN | 63,136 | 61,113 | 97% | Q96RP9 | EFGM_HUMAN | EFGM_HUMAN | Elongation factor G, mitochondrial (EF-Gmt) (Elongation factor G 1, mitochondrial) (mEF-G 1) (Elongation factor G1) (hEFG1) | GFM1 EFG EFG1 GFM |
| sp\|O95197\|RTN3_HUMAN | 171,737 | 207,111 | 148,587 | 201,679 | sp\|O95197\|RTN3_HUMAN | 166,071 | 40,026 | 24% | O95197 | RTN3_HUMAN | RTN3_HUMAN | Reticulon-3 (Homolog of ASY protein) (HAP) (Neuroendocrine-specific protein-like 2) (NSP-like protein 2) (Neuroendocrine-specific protein-like II) (NSP-like protein II) (NSPLII) | RTN3 ASYIP NSPL2 |
| sp\|O43399\|TPD54_HUMAN | 109,516 | 204,642 | 155,627 | 172,455 | sp\|O43399\|TPD54_HUMAN | 154,018 | 37,577 | 24% | O43399 | TPD54_HUMAN | TPD54_HUMAN | Tumor protein D54 (hD54) (Tumor protein D52-like 2) | TPD52L2 |
| sp\|P61081\|UBC12_HUMAN | 164,523 | 167,088 | 268,330 | 283,768 | sp\|P61081\|UBC12_HUMAN | 196,534 | 53,325 | 27% | P61081 | UBC12_HUMAN | UBC12_HUMAN | NEDD8-conjugating enzyme Ubc12 (EC 2.3.2.34) (NEDD8 carrier protein) (Ubiquitin-conjugating enzyme E2 M) | UBE2M UBC12 |
| sp\|P08134\|RHOC_HUMAN | 607,210 | 828,015 | 429,107 | 512,745 | sp\|P08134\|RHOC_HUMAN | 567,610 | 158,460 | 28% | P08134 | RHOC_HUMAN | RHOC_HUMAN | Rho-related GTP-binding protein RhoC (Rho cDNA clone 9) (h9) | RHOC ARH9 ARHC |
| sp\|Q9Y285\|SYFA_HUMAN | 85,795 | 74,549 | 19,205 | 48,140 | sp\|Q9Y285\|SYFA_HUMAN | 51,095 | 20,860 | 41% | Q9Y285 | SYFA_HUMAN | SYFA_HUMAN | Phenylalanine--tRNA ligase alpha subunit (EC 6.1.1.20) (CML33) (Phenylalanyl-tRNA synthetase alpha subunit) (PheRS) | FARSA FARS FARSL FARSLA |
| sp\|Q9H299\|SH3L3_HUMAN | 144,711 | 172,042 | 206,537 | 268,035 | sp\|Q9H299\|SH3L3_HUMAN | 189,294 | 67,349 | 36% | Q9H299 | SH3L3_HUMAN | SH3L3_HUMAN | SH3 domain-binding glutamic acid-rich-like protein 3 (SH3 domain-binding protein 1) (SH3BP-1) | SH3BGRL3 P1725 |
| sp\|P06493\|CDK1_HUMAN | 72,787 | 97,864 | 95,486 | 90,756 | sp\|P06493\|CDK1_HUMAN | 86,803 | 11,352 | 13% | P06493 | CDK1_HUMAN | CDK1_HUMAN | Cyclin-dependent kinase 1 (CDK1) (EC 2.7.11.22) (EC 2.7.11.23) (Cell division control protein 2 homolog) (Cell division protein kinase 1) (p34 protein kinase) | CDK1 CDC2 CDC28A CDKN1 P34CDC2 |
| sp\|P22061\|PIMT_HUMAN | 141,769 | 156,419 | 301,737 | 316,959 | sp\|P22061\|PIMT_HUMAN | 226,412 | 78,944 | 35% | P22061 | PIMT_HUMAN | PIMT_HUMAN | Protein-L-isoaspartate(D-aspartate) O-methyltransferase (PIMT) (EC 2.1.1.77) (L-isoaspartyl protein carboxyl methyltransferase) (Protein L-isoaspartyl/D-aspartyl methyltransferase) (Protein-beta-aspartate methyltransferase) | PCMT1 |
| sp\|P49184\|DNSL1_HUMAN | 467,831 | 490,827 | 749,555 | 810,709 | sp\|P49184\|DNSL1_HUMAN | 557,081 | 148,866 | 27% | P49184 | DNSL1_HUMAN | DNSL1_HUMAN | Deoxyribonuclease-1-like 1 (EC 3.1.21.-) (DNase X) (Deoxyribonuclease I-like 1) (DNase I-like 1) (Muscle-specific DNase I-like) (XIB) | DNASE1L1 DNAS1L1 DNL1L |
| sp\|O14936\|CSKP_HUMAN | 131,221 | 139,153 | 96,138 | 98,832 | sp\|O14936\|CSKP_HUMAN | 109,688 | 24,463 | 22% | O14936 | CSKP_HUMAN | CSKP_HUMAN | Peripheral plasma membrane protein CASK (hCASK) (EC 2.7.11.1) (Calcium/calmodulin-dependent serine protein kinase) (Protein lin-2 homolog) | CASK LIN2 |
| sp\|Q9Y5M8\|SRPRB_HUMAN | 23,032 | 29,764 | 28,301 | 23,891 | sp\|Q9Y5M8\|SRPRB_HUMAN | 29,726 | 14,135 | 48% | Q9Y5M8 | SRPRB_HUMAN | SRPRB_HUMAN | Signal recognition particle receptor subunit beta (SR-beta) (Protein APMCF1) | SRPRB PSEC0230 |

| | | | | | | | | | | | | | |
|---|---|---|---|---|---|---|---|---|---|---|---|---|---|
| sp\|P55145\|MANF_HUMAN | 23,273 | 61,394 | 27,677 | 46,382 | sp\|P55145\|MANF_HUMAN | 112,485 | 116,056 | 103% | P55145 | MANF_HUMAN | MANF_HUMAN | Mesencephalic astrocyte-derived neurotrophic factor (Arginine-rich protein) (Protein ARMET) | MANF ARMET ARP |
| sp\|Q99720\|SGMR1_HUMAN | 43,514 | 33,003 | 16,968 | 64,176 | sp\|Q99720\|SGMR1_HUMAN | 35,493 | 22,287 | 63% | Q99720 | SGMR1_HUMAN | SGMR1_HUMAN | Sigma non-opioid intracellular receptor 1 (Aging-associated gene 8 protein) (SR31747-binding protein) (SR-BP) (Sigma 1-type opioid receptor) (SIG-1R) (Sigma1-receptor) (Sigma1R) (hSigmaR1) | SIGMAR1 OPRS1 SRBP AAG8 |
| sp\|P09110\|THIK_HUMAN | 41,803 | 46,808 | 43,029 | 71,414 | sp\|P09110\|THIK_HUMAN | 42,567 | 17,577 | 41% | P09110 | THIK_HUMAN | THIK_HUMAN | 3-ketoacyl-CoA thiolase, peroxisomal (EC 2.3.1.16) (Acetyl-CoA C-myristoyltransferase) (EC 2.3.1.155) (Acetyl-CoA acyltransferase) (EC 2.3.1.9) (Beta-ketothiolase) (Peroxisomal 3-oxoacyl-CoA thiolase) | ACAA1 ACAA PTHIO |
| sp\|P00742\|FA10_HUMAN | 82,424 | 46,430 | 51,371 | 38,935 | sp\|P00742\|FA10_HUMAN | 392,797 | 407,320 | 104% | P00742 | FA10_HUMAN | FA10_HUMAN | Coagulation factor X (EC 3.4.21.6) (Stuart factor) (Stuart-Prower factor) [Cleaved into: Factor X light chain; Factor X heavy chain; Activated factor Xa heavy chain] | F10 |
| sp\|O95571\|ETHE1_HUMAN | 136,063 | 147,335 | 169,462 | 322,952 | sp\|O95571\|ETHE1_HUMAN | 169,989 | 78,469 | 46% | O95571 | ETHE1_HUMAN | ETHE1_HUMAN | Persulfide dioxygenase ETHE1, mitochondrial (EC 1.13.11.18) (Ethylmalonic encephalopathy protein 1) (Hepatoma subtracted clone one protein) (Sulfur dioxygenase ETHE1) | ETHE1 HSCO |
| sp\|Q7L513\|FCRLA_HUMAN | 34,005 | 30,167 | 75,589 | 77,044 | sp\|Q7L513\|FCRLA_HUMAN | 37,956 | 24,950 | 66% | Q7L513 | FCRLA_HUMAN | FCRLA_HUMAN | Fc receptor-like A (Fc receptor homolog expressed in B-cells) (Fc receptor-like and mucin-like protein 1) (Fc receptor-like protein) (Fc receptor-related protein X) (FcRX) | FCRLA FCRL FCRL1 FCRLM1 FCRX FREB UNQ291/PRO329 |
| sp\|P23588\|IF4B_HUMAN | 492 | 810 | 1,540 | 1,961 | sp\|P23588\|IF4B_HUMAN | 1,430 | 604 | 42% | P23588 | IF4B_HUMAN | IF4B_HUMAN | Eukaryotic translation initiation factor 4B (eIF-4B) | EIF4B |
| sp\|Q86YQ8\|CPNE8_HUMAN | 190,943 | 187,754 | 134,502 | 112,238 | sp\|Q86YQ8\|CPNE8_HUMAN | 192,809 | 49,257 | 26% | Q86YQ8 | CPNE8_HUMAN | CPNE8_HUMAN | Copine-8 (Copine VIII) | CPNE8 |
| sp\|Q6NZI2\|CAVN1_HUMAN | 69,723 | 121,824 | 62,589 | 97,680 | sp\|Q6NZI2\|CAVN1_HUMAN | 146,405 | 87,170 | 60% | Q6NZI2 | CAVN1_HUMAN | CAVN1_HUMAN | Caveolae-associated protein 1 (Cavin-1) (Polymerase I and transcript release factor) | CAVIN1 PTRF FKSG13 |
| sp\|O43760\|SNG2_HUMAN | 617,888 | 586,640 | 684,817 | 493,648 | sp\|O43760\|SNG2_HUMAN | 483,913 | 137,088 | 28% | O43760 | SNG2_HUMAN | SNG2_HUMAN | Synaptogyrin-2 (Cellugyrin) | SYNGR2 UNQ352/PRO615 |
| sp\|Q9ULC4\|MCTS1_HUMAN | 23,333 | 58,730 | 9,189 | 6,480 | sp\|Q9ULC4\|MCTS1_HUMAN | 25,170 | 22,026 | 88% | Q9ULC4 | MCTS1_HUMAN | MCTS1_HUMAN | Malignant T-cell-amplified sequence 1 (MCT-1) (Multiple copies T-cell malignancies) | MCTS1 MCT1 |
| sp\|Q71UM5\|RS27L_HUMAN | 460,559 | 423,619 | 751,015 | 691,977 | sp\|Q71UM5\|RS27L_HUMAN | 446,856 | 188,976 | 42% | Q71UM5 | RS27L_HUMAN | RS27L_HUMAN | 40S ribosomal protein S27-like (Small ribosomal subunit protein eS27-like) | RPS27L |
| sp\|Q9Y6M7\|S4A7_HUMAN | 126,606 | 177,097 | 164,444 | 242,595 | sp\|Q9Y6M7\|S4A7_HUMAN | 195,645 | 69,985 | 36% | Q9Y6M7 | S4A7_HUMAN | S4A7_HUMAN | Sodium bicarbonate cotransporter 3 (Electroneutral Na/HCO(3) cotransporter) (Sodium bicarbonate cotransporter 2) (Sodium bicarbonate cotransporter 2b) (Bicarbonate transporter) (Solute carrier family 4 member 7) | SLC4A7 BT NBC2 NBC2B NBC3 NBCn1 SBC2 SLC4A6 |
| sp\|P84098\|RL19_HUMAN | 70,233 | 5,884 | 57,061 | 13,383 | sp\|P84098\|RL19_HUMAN | 27,892 | 23,560 | 84% | P84098 | RL19_HUMAN | RL19_HUMAN | 60S ribosomal protein L19 (Large ribosomal subunit protein eL19) | RPL19 |
| sp\|Q00341\|VIGLN_HUMAN | 33,690 | 29,050 | 15,198 | 50,110 | sp\|Q00341\|VIGLN_HUMAN | 25,748 | 14,006 | 54% | Q00341 | VIGLN_HUMAN | VIGLN_HUMAN | Vigilin (High density lipoprotein-binding protein) (HDL-binding protein) | HDLBP HBP VGL |

| | | | | | | | | | | | | | |
|---|---|---|---|---|---|---|---|---|---|---|---|---|---|
| sp\|P46778\|RL21_HUMAN | 200,270 | 168,517 | 171,714 | 142,436 | sp\|P46778\|RL21_HUMAN | 95,327 | 82,657 | 87% | P46778 | RL21_HUMAN | RL21_HUMAN | 60S ribosomal protein L21 (Large ribosomal subunit protein eL21) | RPL21 |
| sp\|P00747\|PLMN_HUMAN | 28,102 | 18,426 | 73,639 | 45,596 | sp\|P00747\|PLMN_HUMAN | 145,924 | 119,066 | 82% | P00747 | PLMN_HUMAN | PLMN_HUMAN | Plasminogen (EC 3.4.21.7) [Cleaved into: Plasmin heavy chain A; Activation peptide; Angiostatin; Plasmin heavy chain A, short form; Plasmin light chain B] | PLG |
| sp\|O43865\|SAHH2_HUMAN | 145,506 | 100,624 | 222,836 | 271,445 | sp\|O43865\|SAHH2_HUMAN | 200,281 | 56,938 | 28% | O43865 | SAHH2_HUMAN | SAHH2_HUMAN | S-adenosylhomocysteine hydrolase-like protein 1 (DC-expressed AHCY-like molecule) (IP(3)Rs binding protein released with IP(3)) (IRBIT) (Putative adenosylhomocysteinase 2) (S-adenosyl-L-homocysteine hydrolase 2) (AdoHcyase 2) | AHCYL1 DCAL IRBIT XPVKONA |
| sp\|P54852\|EMP3_HUMAN | 1,475,926 | 2,724,795 | 278,758 | 482,106 | sp\|P54852\|EMP3_HUMAN | 976,406 | 959,484 | 98% | P54852 | EMP3_HUMAN | EMP3_HUMAN | Epithelial membrane protein 3 (EMP-3) (Hematopoietic neural membrane protein 1) (HNMP-1) (Protein YMP) | EMP3 YMP |
| sp\|P00441\|SODC_HUMAN | 102,828 | 96,328 | 60,235 | 30,908 | sp\|P00441\|SODC_HUMAN | 110,819 | 83,147 | 75% | P00441 | SODC_HUMAN | SODC_HUMAN | Superoxide dismutase [Cu-Zn] (EC 1.15.1.1) (Superoxide dismutase 1) (hSod1) | SOD1 |
| sp\|Q969Q0\|RL36L_HUMAN | 265,433 | 213,503 | 473,961 | 264,081 | sp\|Q969Q0\|RL36L_HUMAN | 211,304 | 126,584 | 60% | Q969Q0 | RL36L_HUMAN | RL36L_HUMAN | 60S ribosomal protein L36a-like (Large ribosomal subunit protein eL42-like) | RPL36AL |
| sp\|P30519\|HMOX2_HUMAN | 31,631 | 32,463 | 33,210 | 32,231 | sp\|P30519\|HMOX2_HUMAN | 30,531 | 6,935 | 23% | P30519 | HMOX2_HUMAN | HMOX2_HUMAN | Heme oxygenase 2 (HO-2) (EC 1.14.14.18) [Cleaved into: Heme oxygenase 2 soluble form] | HMOX2 HO2 |
| sp\|P51884\|LUM_HUMAN | 7,999 | 36,310 | 26,868 | 21,958 | sp\|P51884\|LUM_HUMAN | 263,357 | 370,441 | 141% | P51884 | LUM_HUMAN | LUM_HUMAN | Lumican (Keratan sulfate proteoglycan lumican) (KSPG lumican) | LUM LDC SLRR2D |
| sp\|Q15056\|IF4H_HUMAN | 1,937 | 2,675 | 3,943 | 12,494 | sp\|Q15056\|IF4H_HUMAN | 10,709 | 16,478 | 154% | Q15056 | IF4H_HUMAN | IF4H_HUMAN | Eukaryotic translation initiation factor 4H (eIF-4H) (Williams-Beuren syndrome chromosomal region 1 protein) | EIF4H KIAA0038 WBSCR1 WSCR1 |
| sp\|O95183\|VAMP5_HUMAN | 348,627 | 440,103 | 404,665 | 331,474 | sp\|O95183\|VAMP5_HUMAN | 337,429 | 74,810 | 22% | O95183 | VAMP5_HUMAN | VAMP5_HUMAN | Vesicle-associated membrane protein 5 (VAMP-5) (Myobrevin) | VAMP5 HSPC191 |
| sp\|Q14254\|FLOT2_HUMAN | 314,762 | 329,424 | 496,369 | 480,353 | sp\|Q14254\|FLOT2_HUMAN | 373,113 | 111,964 | 30% | Q14254 | FLOT2_HUMAN | FLOT2_HUMAN | Flotillin-2 (Epidermal surface antigen) (ESA) (Membrane component chromosome 17 surface marker 1) | FLOT2 ESA1 M17S1 |
| sp\|Q9NUU7\|DD19A_HUMAN | 34,546 | 60,761 | 57,383 | 40,876 | sp\|Q9NUU7\|DD19A_HUMAN | 50,017 | 10,781 | 22% | Q9NUU7 | DD19A_HUMAN | DD19A_HUMAN | ATP-dependent RNA helicase DDX19A (EC 3.6.4.13) (DDX19-like protein) (DEAD box protein 19A) | DDX19A DDX19L |
| sp\|P06737\|PYGL_HUMAN | 94,326 | 89,133 | 271,555 | 222,839 | sp\|P06737\|PYGL_HUMAN | 207,859 | 79,922 | 38% | P06737 | PYGL_HUMAN | PYGL_HUMAN | Glycogen phosphorylase, liver form (EC 2.4.1.1) | PYGL |
| sp\|P24752\|THIL_HUMAN | 3,892 | 10,190 | 3,073 | 9,231 | sp\|P24752\|THIL_HUMAN | 13,112 | 19,186 | 146% | P24752 | THIL_HUMAN | THIL_HUMAN | Acetyl-CoA acetyltransferase, mitochondrial (EC 2.3.1.9) (Acetoacetyl-CoA thiolase) (T2) | ACAT1 ACAT MAT |
| sp\|Q9Y5X1\|SNX9_HUMAN | 129,259 | 196,338 | 347,489 | 371,918 | sp\|Q9Y5X1\|SNX9_HUMAN | 206,992 | 98,945 | 48% | Q9Y5X1 | SNX9_HUMAN | SNX9_HUMAN | Sorting nexin-9 (SH3 and PX domain-containing protein 1) (Protein SDP1) (SH3 and PX domain-containing protein 3A) | SNX9 SH3PX1 SH3PXD3A |

| | | | | | | | | | | | | | |
|---|---|---|---|---|---|---|---|---|---|---|---|---|---|
| sp\|P22413\|ENPP1_HUMAN | 608 | 1,018 | 9,399 | 1,313 | sp\|P22413\|ENPP1_HUMAN | 3,054 | 2,839 | 93% | P22413 | ENPP1_HUMAN | ENPP1_HUMAN | Ectonucleotide pyrophosphatase/phosphodiesterase family member 1 (E-NPP 1) (Membrane component chromosome 6 surface marker 1) (Phosphodiesterase I/nucleotide pyrophosphatase 1) (Plasma-cell membrane glycoprotein PC-1) [Cleaved into: Ectonucleotide pyrophosphatase/phosphodiesterase family member 1, secreted form] [Includes: Alkaline phosphodiesterase I (EC 3.1.4.1); Nucleotide pyrophosphatase (NPPase) (EC 3.6.1.9) (Nucleotide diphosphatase)] | ENPP1 M6S1 NPPS PC1 PDNP1 |
| sp\|P61769\|B2MG_HUMAN | 187,190 | 361,130 | 122,312 | 148,448 | sp\|P61769\|B2MG_HUMAN | 165,146 | 99,076 | 60% | P61769 | B2MG_HUMAN | B2MG_HUMAN | Beta-2-microglobulin [Cleaved into: Beta-2-microglobulin form pI 5.3] | B2M CDABP0092 HDCMA22P |
| sp\|P37268\|FDFT_HUMAN | 34,583 | 41,294 | 35,657 | 18,070 | sp\|P37268\|FDFT_HUMAN | 34,524 | 18,361 | 53% | P37268 | FDFT_HUMAN | FDFT_HUMAN | Squalene synthase (SQS) (SS) (EC 2.5.1.21) (FPP:FPP farnesyltransferase) (Farnesyl-diphosphate farnesyltransferase) (Farnesyl-diphosphate farnesyltransferase 1) | FDFT1 |
| sp\|P63010\|AP2B1_HUMAN | 222,003 | 455,500 | 199,112 | 258,110 | sp\|P63010\|AP2B1_HUMAN | 234,587 | 112,194 | 48% | P63010 | AP2B1_HUMAN | AP2B1_HUMAN | AP-2 complex subunit beta (AP105B) (Adaptor protein complex AP-2 subunit beta) (Adaptor-related protein complex 2 subunit beta) (Beta-2-adaptin) (Beta-adaptin) (Clathrin assembly protein complex 2 beta large chain) (Plasma membrane adaptor HA2/AP2 adaptin beta subunit) | AP2B1 ADTB2 CLAPB1 |
| sp\|P62854\|RS26_HUMAN | 1,162,599 | 1,078,050 | 2,377,235 | 1,715,572 | sp\|P62854\|RS26_HUMAN | 1,103,670 | 653,519 | 59% | P62854 | RS26_HUMAN | RS26_HUMAN | 40S ribosomal protein S26 (Small ribosomal subunit protein eS26) | RPS26 |
| sp\|Q9H8S9\|MOB1A_HUMAN | 291,854 | 312,222 | 149,841 | 267,593 | sp\|Q9H8S9\|MOB1A_HUMAN | 284,568 | 63,562 | 22% | Q9H8S9 | MOB1A_HUMAN | MOB1A_HUMAN | MOB kinase activator 1A (Mob1 alpha) (Mob1A) (Mob1 homolog 1B) (Mps one binder kinase activator-like 1B) | MOB1A C2orf6 MOB4B MOBK1B MOBKL1B |
| sp\|Q9UBG0\|MRC2_HUMAN | 104,022 | 78,096 | 110,138 | 116,722 | sp\|Q9UBG0\|MRC2_HUMAN | 182,572 | 89,111 | 49% | Q9UBG0 | MRC2_HUMAN | MRC2_HUMAN | C-type mannose receptor 2 (C-type lectin domain family 13 member E) (Endocytic receptor 180) (Macrophage mannose receptor 2) (Urokinase-type plasminogen activator receptor-associated protein) (UPAR-associated protein) (Urokinase receptor-associated protein) (CD antigen CD280) | MRC2 CLEC13E ENDO180 KIAA0709 UPARAP |
| sp\|P31153\|METK2_HUMAN | 160,409 | 158,363 | 140,473 | 145,833 | sp\|P31153\|METK2_HUMAN | 180,551 | 40,612 | 22% | P31153 | METK2_HUMAN | METK2_HUMAN | S-adenosylmethionine synthase isoform type-2 (AdoMet synthase 2) (EC 2.5.1.6) (Methionine adenosyltransferase 2) (MAT 2) (Methionine adenosyltransferase II) (MAT-II) | MAT2A AMS2 MATA2 |
| sp\|Q15836\|VAMP3_HUMAN | 176,895 | 161,061 | 190,117 | 130,278 | sp\|Q15836\|VAMP3_HUMAN | 154,963 | 54,151 | 35% | Q15836 | VAMP3_HUMAN | VAMP3_HUMAN | Vesicle-associated membrane protein 3 (VAMP-3) (Cellubrevin) (CEB) (Synaptobrevin-3) | VAMP3 SYB3 |
| sp\|Q8IUW5\|RELL1_HUMAN | 92,016 | 91,063 | 70,032 | 75,527 | sp\|Q8IUW5\|RELL1_HUMAN | 68,979 | 23,591 | 34% | Q8IUW5 | RELL1_HUMAN | RELL1_HUMAN | RELT-like protein 1 | RELL1 PSEC0162 |

| | | | | | | | | | | | | | |
|---|---|---|---|---|---|---|---|---|---|---|---|---|---|
| sp\|Q9BUP3\|HTAI2_HUMAN | 109,179 | 106,696 | 42,877 | 177,366 | sp\|Q9BUP3\|HTAI2_HUMAN | 85,567 | 49,617 | 58% | Q9BUP3 | HTAI2_HUMAN | HTAI2_HUMAN | Oxidoreductase HTATIP2 (EC 1.1.1.-) (30 kDa HIV-1 TAT-interacting protein) (HIV-1 TAT-interactive protein 2) | HTATIP2 CC3 TIP30 |
| sp\|Q9Y625\|GPC6_HUMAN | 145,508 | 213,838 | 67,930 | 105,243 | sp\|Q9Y625\|GPC6_HUMAN | 124,373 | 58,560 | 47% | Q9Y625 | GPC6_HUMAN | GPC6_HUMAN | Glypican-6 [Cleaved into: Secreted glypican-6] | GPC6 UNQ369/PRO705 |
| sp\|Q16831\|UPP1_HUMAN | 125,545 | 139,497 | 146,254 | 180,626 | sp\|Q16831\|UPP1_HUMAN | 125,906 | 36,298 | 29% | Q16831 | UPP1_HUMAN | UPP1_HUMAN | Uridine phosphorylase 1 (UPase 1) (UrdPase 1) (EC 2.4.2.3) | UPP1 UP |
| sp\|P61088\|UBE2N_HUMAN | 148,164 | 222,421 | 97,346 | 98,985 | sp\|P61088\|UBE2N_HUMAN | 136,511 | 50,377 | 37% | P61088 | UBE2N_HUMAN | UBE2N_HUMAN | Ubiquitin-conjugating enzyme E2 N (EC 2.3.2.23) (Bendless-like ubiquitin-conjugating enzyme) (E2 ubiquitin-conjugating enzyme N) (Ubc13) (UbcH13) (Ubiquitin carrier protein N) (Ubiquitin-protein ligase N) | UBE2N BLU |
| sp\|Q16531\|DDB1_HUMAN | 110,667 | 87,933 | 44,109 | 136,482 | sp\|Q16531\|DDB1_HUMAN | 104,262 | 64,024 | 61% | Q16531 | DDB1_HUMAN | DDB1_HUMAN | DNA damage-binding protein 1 (DDB p127 subunit) (DNA damage-binding protein a) (DDBa) (Damage-specific DNA-binding protein 1) (HBV X-associated protein 1) (XAP-1) (UV-damaged DNA-binding factor) (UV-damaged DNA-binding protein 1) (UV-DDB 1) (XPE-binding factor) (XPE-BF) (Xeroderma pigmentosum group E-complementing protein) (XPCe) | DDB1 XAP1 |
| sp\|O00429\|DNM1L_HUMAN | 81,096 | 86,084 | 105,312 | 90,245 | sp\|O00429\|DNM1L_HUMAN | 81,589 | 16,430 | 20% | O00429 | DNM1L_HUMAN | DNM1L_HUMAN | Dynamin-1-like protein (EC 3.6.5.5) (Dnm1p/Vps1p-like protein) (DVLP) (Dynamin family member proline-rich carboxyl-terminal domain less) (Dymple) (Dynamin-like protein) (Dynamin-like protein 4) (Dynamin-like protein IV) (HdynIV) (Dynamin-related protein 1) | DNM1L DLP1 DRP1 |
| sp\|P49419\|AL7A1_HUMAN | 131,980 | 139,272 | 102,042 | 130,488 | sp\|P49419\|AL7A1_HUMAN | 131,576 | 58,981 | 45% | P49419 | AL7A1_HUMAN | AL7A1_HUMAN | Alpha-aminoadipic semialdehyde dehydrogenase (Alpha-AASA dehydrogenase) (EC 1.2.1.31) (Aldehyde dehydrogenase family 7 member A1) (EC 1.2.1.3) (Antiquitin-1) (Betaine aldehyde dehydrogenase) (EC 1.2.1.8) (Delta1-piperideine-6-carboxylate dehydrogenase) (P6c dehydrogenase) | ALDH7A1 ATQ1 |
| sp\|P08962\|CD63_HUMAN | 4,577,473 | 4,656,620 | 3,733,542 | 3,342,308 | sp\|P08962\|CD63_HUMAN | 3,176,941 | 1,116,310 | 35% | P08962 | CD63_HUMAN | CD63_HUMAN | CD63 antigen (Granulophysin) (Lysosomal-associated membrane protein 3) (LAMP-3) (Lysosome integral membrane protein 1) (Limp1) (Melanoma-associated antigen ME491) (OMA81H) (Ocular melanoma-associated antigen) (Tetraspanin-30) (Tspan-30) (CD antigen CD63) | CD63 MLA1 TSPAN30 |
| sp\|Q8TCJ2\|STT3B_HUMAN | 3,700 | 15,930 | 51,702 | 20,304 | sp\|Q8TCJ2\|STT3B_HUMAN | 24,196 | 17,697 | 73% | Q8TCJ2 | STT3B_HUMAN | STT3B_HUMAN | Dolichyl-diphosphooligosaccharide--protein glycosyltransferase subunit STT3B (Oligosaccharyl transferase subunit STT3B) (STT3-B) (EC 2.4.99.18) (Source of immunodominant MHC-associated peptides homolog) | STT3B SIMP |

| | | | | | | | | | | | | | |
|---|---|---|---|---|---|---|---|---|---|---|---|---|---|
| sp\|O95297\|MPZL1_HUMAN | 783,735 | 791,995 | 603,507 | 593,505 | sp\|O95297\|MPZL1_HUMAN | 610,577 | 125,287 | 21% | O95297 | MPZL1_HUMAN | MPZL1_HUMAN | Myelin protein zero-like protein 1 (Protein zero-related) | MPZL1 PZR UNQ849/PRO1787 |
| sp\|Q9Y2V2\|CHSP1_HUMAN | 577,833 | 495,954 | 868,013 | 1,018,634 | sp\|Q9Y2V2\|CHSP1_HUMAN | 530,519 | 303,511 | 57% | Q9Y2V2 | CHSP1_HUMAN | CHSP1_HUMAN | Calcium-regulated heat-stable protein 1 (Calcium-regulated heat-stable protein of 24 kDa) (CRHSP-24) | CARHSP1 |
| sp\|Q10471\|GALT2_HUMAN | 329,313 | 186,072 | 679,843 | 496,582 | sp\|Q10471\|GALT2_HUMAN | 309,090 | 194,011 | 63% | Q10471 | GALT2_HUMAN | GALT2_HUMAN | Polypeptide N-acetylgalactosaminyltransferase 2 (EC 2.4.1.41) (Polypeptide GalNAc transferase 2) (GalNAc-T2) (pp-GaNTase 2) (Protein-UDP acetylgalactosaminyltransferase 2) (UDP-GalNAc:polypeptide N-acetylgalactosaminyltransferase 2) [Cleaved into: Polypeptide N-acetylgalactosaminyltransferase 2 soluble form] | GALNT2 |
| sp\|Q9UN86\|G3BP2_HUMAN | 37,029 | 45,152 | 35,938 | 45,589 | sp\|Q9UN86\|G3BP2_HUMAN | 36,131 | 12,748 | 35% | Q9UN86 | G3BP2_HUMAN | G3BP2_HUMAN | Ras GTPase-activating protein-binding protein 2 (G3BP-2) (GAP SH3 domain-binding protein 2) | G3BP2 KIAA0660 |
| sp\|P12081\|SYHC_HUMAN | 113,532 | 121,415 | 207,772 | 190,001 | sp\|P12081\|SYHC_HUMAN | 140,924 | 38,741 | 27% | P12081 | SYHC_HUMAN | HARS1_HUMAN | Histidine--tRNA ligase, cytoplasmic (EC 6.1.1.21) (Histidyl-tRNA synthetase) (HisRS) | HARS1 HARS HRS |
| sp\|Q96FW1\|OTUB1_HUMAN | 116,812 | 119,684 | 141,572 | 124,298 | sp\|Q96FW1\|OTUB1_HUMAN | 105,491 | 26,965 | 26% | Q96FW1 | OTUB1_HUMAN | OTUB1_HUMAN | Ubiquitin thioesterase OTUB1 (EC 3.4.19.12) (Deubiquitinating enzyme OTUB1) (OTU domain-containing ubiquitin aldehyde-binding protein 1) (Otubain-1) (hOTU1) (Ubiquitin-specific-processing protease OTUB1) | OTUB1 OTB1 OTU1 HSPC263 |
| sp\|P16402\|H13_HUMAN | 2,549,715 | 910,967 | 2,512,180 | 741,725 | sp\|P16402\|H13_HUMAN | 1,074,645 | 935,214 | 87% | P16402 | H13_HUMAN | H13_HUMAN | Histone H1.3 (Histone H1c) (Histone H1s-2) | H1-3 H1F3 HIST1H1D |
| sp\|Q15631\|TSN_HUMAN | 64,316 | 112,095 | 56,481 | 62,937 | sp\|Q15631\|TSN_HUMAN | 73,483 | 18,336 | 25% | Q15631 | TSN_HUMAN | TSN_HUMAN | Translin (EC 3.1.-.-) (Component 3 of promoter of RISC) (C3PO) | TSN |
| sp\|Q96AZ6\|ISG20_HUMAN | 62,973 | 40,611 | 36,399 | 49,093 | sp\|Q96AZ6\|ISG20_HUMAN | 48,979 | 8,332 | 17% | Q96AZ6 | ISG20_HUMAN | ISG20_HUMAN | Interferon-stimulated gene 20 kDa protein (EC 3.1.13.1) (Estrogen-regulated transcript 45 protein) (Promyelocytic leukemia nuclear body-associated protein ISG20) | ISG20 HEM45 |
| sp\|Q15102\|PA1B3_HUMAN | 183,590 | 195,623 | 317,861 | 279,365 | sp\|Q15102\|PA1B3_HUMAN | 239,967 | 53,217 | 22% | Q15102 | PA1B3_HUMAN | PA1B3_HUMAN | Platelet-activating factor acetylhydrolase IB subunit alpha1 (EC 3.1.1.47) (PAF acetylhydrolase 29 kDa subunit) (PAF-AH 29 kDa subunit) (PAF-AH subunit gamma) (PAFAH subunit gamma) | PAFAH1B3 PAFAHG |
| sp\|Q15005\|SPCS2_HUMAN | 42,738 | 75,351 | 38,297 | 37,201 | sp\|Q15005\|SPCS2_HUMAN | 109,303 | 146,147 | 134% | Q15005 | SPCS2_HUMAN | SPCS2_HUMAN | Signal peptidase complex subunit 2 (Microsomal signal peptidase 25 kDa subunit) (SPase 25 kDa subunit) | SPCS2 KIAA0102 SPC25 |
| sp\|O75947\|ATP5H_HUMAN | 4,430 | 30,576 | 14,018 | 16,233 | sp\|O75947\|ATP5H_HUMAN | 31,903 | 35,271 | 111% | O75947 | ATP5H_HUMAN | ATP5H_HUMAN | ATP synthase subunit d, mitochondrial (ATPase subunit d) (ATP synthase peripheral stalk subunit d) | ATP5PD ATP5H My032 |
| sp\|Q7Z5G4\|GOGA7_HUMAN | 145,515 | 167,904 | 112,870 | 191,742 | sp\|Q7Z5G4\|GOGA7_HUMAN | 169,284 | 46,357 | 27% | Q7Z5G4 | GOGA7_HUMAN | GOGA7_HUMAN | Golgin subfamily A member 7 (Golgi complex-associated protein of 16 kDa) | GOLGA7 GCP16 HDCKB03P HSPC041 |
| sp\|P06730\|IF4E_HUMAN | 64,755 | 63,983 | 38,205 | 45,187 | sp\|P06730\|IF4E_HUMAN | 44,403 | 17,660 | 40% | P06730 | IF4E_HUMAN | IF4E_HUMAN | Eukaryotic translation initiation factor 4E (eIF-4E) (eIF4E) (eIF-4F 25 kDa subunit) (mRNA cap-binding protein) | EIF4E EIF4EL1 EIF4F |

| | | | | | | | | | | | | | |
|---|---|---|---|---|---|---|---|---|---|---|---|---|---|
| sp\|P35237\|SPB6_HUMAN | 108,909 | 125,906 | 19,299 | 43,097 | sp\|P35237\|SPB6_HUMAN | 71,066 | 41,757 | 59% | P35237 | SPB6_HUMAN | SPB6_HUMAN | Serpin B6 (Cytoplasmic antiproteinase) (CAP) (Peptidase inhibitor 6) (PI-6) (Placental thrombin inhibitor) | SERPINB6 PI6 PTI |
| sp\|Q14444\|CAPR1_HUMAN | 19,308 | 34,748 | 19,346 | 28,945 | sp\|Q14444\|CAPR1_HUMAN | 26,449 | 9,943 | 38% | Q14444 | CAPR1_HUMAN | CAPR1_HUMAN | Caprin-1 (Cell cycle-associated protein 1) (Cytoplasmic activation- and proliferation-associated protein 1) (GPI-anchored membrane protein 1) (GPI-anchored protein p137) (GPI-p137) (p137GPI) (Membrane component chromosome 11 surface marker 1) (RNA granule protein 105) | CAPRIN1 GPIAP1 GPIP137 M11S1 RNG105 |
| sp\|O75477\|ERLN1_HUMAN | 53,010 | 47,554 | 54,484 | 76,269 | sp\|O75477\|ERLN1_HUMAN | 44,552 | 19,830 | 45% | O75477 | ERLN1_HUMAN | ERLN1_HUMAN | Erlin-1 (Endoplasmic reticulum lipid raft-associated protein 1) (Protein KE04) (Stomatin-prohibitin-flotillin-HflC/K domain-containing protein 1) (SPFH domain-containing protein 1) | ERLIN1 C10orf69 KE04 KEO4 SPFH1 |
| sp\|P08174\|DAF_HUMAN | 679,431 | 643,540 | 722,668 | 836,068 | sp\|P08174\|DAF_HUMAN | 555,962 | 187,694 | 34% | P08174 | DAF_HUMAN | DAF_HUMAN | Complement decay-accelerating factor (CD antigen CD55) | CD55 CR DAF |
| sp\|P35080\|PROF2_HUMAN | 255,330 | 279,178 | 187,591 | 318,730 | sp\|P35080\|PROF2_HUMAN | 223,689 | 59,293 | 27% | P35080 | PROF2_HUMAN | PROF2_HUMAN | Profilin-2 (Profilin II) | PFN2 |
| sp\|Q86UE4\|LYRIC_HUMAN | 4,670 | 14,779 | 16,746 | 36,429 | sp\|Q86UE4\|LYRIC_HUMAN | 16,152 | 9,892 | 61% | Q86UE4 | LYRIC_HUMAN | LYRIC_HUMAN | Protein LYRIC (3D3/LYRIC) (Astrocyte elevated gene-1 protein) (AEG-1) (Lysine-rich CEACAM1 co-isolated protein) (Metadherin) (Metastasis adhesion protein) | MTDH AEG1 LYRIC |
| sp\|O75663\|TIPRL_HUMAN | 35,124 | 38,618 | 22,650 | 25,161 | sp\|O75663\|TIPRL_HUMAN | 31,167 | 7,437 | 24% | O75663 | TIPRL_HUMAN | TIPRL_HUMAN | TIP41-like protein (Putative MAPK-activating protein PM10) (Type 2A-interacting protein) (TIP) | TIPRL |
| sp\|Q96PY5\|FMNL2_HUMAN | 151,730 | 208,083 | 37,241 | 86,823 | sp\|Q96PY5\|FMNL2_HUMAN | 97,618 | 68,976 | 71% | Q96PY5 | FMNL2_HUMAN | FMNL2_HUMAN | Formin-like protein 2 (Formin homology 2 domain-containing protein 2) | FMNL2 FHOD2 KIAA1902 |
| sp\|P49006\|MRP_HUMAN | 873,546 | 755,738 | 1,361,615 | 1,314,743 | sp\|P49006\|MRP_HUMAN | 919,914 | 280,079 | 30% | P49006 | MRP_HUMAN | MRP_HUMAN | MARCKS-related protein (MARCKS-like protein 1) (Macrophage myristoylated alanine-rich C kinase substrate) (Mac-MARCKS) (MacMARCKS) | MARCKSL1 MLP MRP |
| sp\|P49747\|COMP_HUMAN | 11,970 | 11,210 | 12,342 | 60,000 | sp\|P49747\|COMP_HUMAN | 141,066 | 156,814 | 111% | P49747 | COMP_HUMAN | COMP_HUMAN | Cartilage oligomeric matrix protein (COMP) (Thrombospondin-5) (TSP5) | COMP |
| sp\|P13798\|ACPH_HUMAN | 117,163 | 142,675 | 104,771 | 147,216 | sp\|P13798\|ACPH_HUMAN | 112,457 | 33,464 | 30% | P13798 | ACPH_HUMAN | ACPH_HUMAN | Acylamino-acid-releasing enzyme (AARE) (EC 3.4.19.1) (Acyl-peptide hydrolase) (APH) (Acylaminoacyl-peptidase) (Oxidized protein hydrolase) (OPH) | APEH D3F15S2 D3S48E DNF15S2 |
| sp\|Q9Y3U8\|RL36_HUMAN | 297,553 | 253,130 | 367,408 | 382,305 | sp\|Q9Y3U8\|RL36_HUMAN | 262,302 | 87,227 | 33% | Q9Y3U8 | RL36_HUMAN | RL36_HUMAN | 60S ribosomal protein L36 (Large ribosomal subunit protein eL36) | RPL36 |
| sp\|P01116\|RASK_HUMAN | 180,141 | 72,842 | 43,884 | 110,394 | sp\|P01116\|RASK_HUMAN | 88,709 | 42,850 | 48% | P01116 | RASK_HUMAN | RASK_HUMAN | GTPase KRas (EC 3.6.5.2) (K-Ras 2) (Ki-Ras) (c-K-ras) (c-Ki-ras) [Cleaved into: GTPase KRas, N-terminally processed] | KRAS KRAS2 RASK2 |
| sp\|O75351\|VPS4B_HUMAN | 381,826 | 452,448 | 122,422 | 352,930 | sp\|O75351\|VPS4B_HUMAN | 281,958 | 116,172 | 41% | O75351 | VPS4B_HUMAN | VPS4B_HUMAN | Vacuolar protein sorting-associated protein 4B (EC 3.6.4.6) (Cell migration-inducing gene 1 protein) (Suppressor of K(+) transport growth defect 1) (Protein SKD1) | VPS4B SKD1 VPS42 MIG1 |

| | | | | | | | | | | | | | |
|---|---|---|---|---|---|---|---|---|---|---|---|---|---|
| sp\|P36383\|CXG1_HUMAN | 177,439 | 463,592 | 248,374 | 644,814 | sp\|P36383\|CXG1_HUMAN | 415,306 | 144,734 | 35% | P36383 | CXG1_HUMAN | CXG1_HUMAN | Gap junction gamma-1 protein (Connexin-45) (Cx45) (Gap junction alpha-7 protein) | GJC1 GJA7 |
| sp\|Q92820\|GGH_HUMAN | 58,073 | 46,872 | 80,804 | 22,747 | sp\|Q92820\|GGH_HUMAN | 73,363 | 40,237 | 55% | Q92820 | GGH_HUMAN | GGH_HUMAN | Gamma-glutamyl hydrolase (EC 3.4.19.9) (Conjugase) (GH) (Gamma-Glu-X carboxypeptidase) | GGH |
| sp\|P07099\|HYEP_HUMAN | 17,742 | 54,454 | 41,170 | 8,673 | sp\|P07099\|HYEP_HUMAN | 29,407 | 14,740 | 50% | P07099 | HYEP_HUMAN | HYEP_HUMAN | Epoxide hydrolase 1 (EC 3.3.2.9) (Epoxide hydratase) (Microsomal epoxide hydrolase) (mEH) | EPHX1 EPHX EPOX |
| sp\|O15145\|ARPC3_HUMAN | 81,234 | 142,900 | 21,528 | 78,805 | sp\|O15145\|ARPC3_HUMAN | 76,626 | 40,368 | 53% | O15145 | ARPC3_HUMAN | ARPC3_HUMAN | Actin-related protein 2/3 complex subunit 3 (Arp2/3 complex 21 kDa subunit) (p21-ARC) | ARPC3 ARC21 |
| sp\|Q9NY33\|DPP3_HUMAN | 41,506 | 32,706 | 88,142 | 125,637 | sp\|Q9NY33\|DPP3_HUMAN | 86,231 | 46,510 | 54% | Q9NY33 | DPP3_HUMAN | DPP3_HUMAN | Dipeptidyl peptidase 3 (EC 3.4.14.4) (Dipeptidyl aminopeptidase III) (Dipeptidyl arylamidase III) (Dipeptidyl peptidase III) (DPP III) (Enkephalinase B) | DPP3 |
| sp\|O43237\|DC1L2_HUMAN | 36,615 | 60,750 | 6,151 | 47,584 | sp\|O43237\|DC1L2_HUMAN | 35,851 | 16,550 | 46% | O43237 | DC1L2_HUMAN | DC1L2_HUMAN | Cytoplasmic dynein 1 light intermediate chain 2 (Dynein light intermediate chain 2, cytosolic) (LIC-2) (LIC53/55) | DYNC1LI2 DNCLI2 LIC2 |
| sp\|O95302\|FKBP9_HUMAN | 35,420 | 19,585 | 53,457 | 14,486 | sp\|O95302\|FKBP9_HUMAN | 36,131 | 27,259 | 75% | O95302 | FKBP9_HUMAN | FKBP9_HUMAN | Peptidyl-prolyl cis-trans isomerase FKBP9 (PPIase FKBP9) (EC 5.2.1.8) (63 kDa FK506-binding protein) (63 kDa FKBP) (FKBP-63) (FK506-binding protein 9) (FKBP-9) (Rotamase) | FKBP9 FKBP60 FKBP63 |
| sp\|P51991\|ROA3_HUMAN | 152,290 | 132,082 | 102,612 | 87,005 | sp\|P51991\|ROA3_HUMAN | 81,107 | 44,390 | 55% | P51991 | ROA3_HUMAN | ROA3_HUMAN | Heterogeneous nuclear ribonucleoprotein A3 (hnRNP A3) | HNRNPA3 HNRPA3 |
| sp\|P53680\|AP2S1_HUMAN | 588,307 | 576,381 | 943,426 | 855,692 | sp\|P53680\|AP2S1_HUMAN | 676,892 | 158,681 | 23% | P53680 | AP2S1_HUMAN | AP2S1_HUMAN | AP-2 complex subunit sigma (Adaptor protein complex AP-2 subunit sigma) (Adaptor-related protein complex 2 subunit sigma) (Clathrin assembly protein 2 sigma small chain) (Clathrin coat assembly protein AP17) (Clathrin coat-associated protein AP17) (HA2 17 kDa subunit) (Plasma membrane adaptor AP-2 17 kDa protein) (Sigma2-adaptin) | AP2S1 AP17 CLAPS2 |
| sp\|Q10713\|MPPA_HUMAN | 5,491 | 13,299 | 14,241 | 18,057 | sp\|Q10713\|MPPA_HUMAN | 18,268 | 8,186 | 45% | Q10713 | MPPA_HUMAN | MPPA_HUMAN | Mitochondrial-processing peptidase subunit alpha (Alpha-MPP) (Inactive zinc metalloprotease alpha) (P-55) | PMPCA INPP5E KIAA0123 MPPA |
| sp\|O95861\|BPNT1_HUMAN | 39,743 | 26,882 | 8,867 | 37,248 | sp\|O95861\|BPNT1_HUMAN | 30,112 | 9,787 | 33% | O95861 | BPNT1_HUMAN | BPNT1_HUMAN | 3'(2'),5'-bisphosphate nucleotidase 1 (EC 3.1.3.7) (Bisphosphate 3'-nucleotidase 1) (PAP-inositol 1,4-phosphatase) (PIP) | BPNT1 |
| sp\|P08493\|MGP_HUMAN | 446,465 | 697,410 | 1,972,593 | 1,936,033 | sp\|P08493\|MGP_HUMAN | 904,444 | 665,750 | 74% | P08493 | MGP_HUMAN | MGP_HUMAN | Matrix Gla protein (MGP) (Cell growth-inhibiting gene 36 protein) | MGP MGLAP GIG36 |
| sp\|O15372\|EIF3H_HUMAN | 6,447 | 5,764 | 1,478 | 1,947 | sp\|O15372\|EIF3H_HUMAN | 3,192 | 2,095 | 66% | O15372 | EIF3H_HUMAN | EIF3H_HUMAN | Eukaryotic translation initiation factor 3 subunit H (eIF3h) (Eukaryotic translation initiation factor 3 subunit 3) (eIF-3-gamma) (eIF3 p40 subunit) | EIF3H EIF3S3 |
| sp\|P14866\|HNRPL_HUMAN | 134,323 | 96,154 | 82,917 | 32,312 | sp\|P14866\|HNRPL_HUMAN | 88,642 | 45,413 | 51% | P14866 | HNRPL_HUMAN | HNRPL_HUMAN | Heterogeneous nuclear ribonucleoprotein L (hnRNP L) | HNRNPL HNRPL P/OKcl.14 |
| sp\|Q9Y639\|NPTN_HUMAN | 262,417 | 329,513 | 102,337 | 279,460 | sp\|Q9Y639\|NPTN_HUMAN | 220,401 | 90,056 | 41% | Q9Y639 | NPTN_HUMAN | NPTN_HUMAN | Neuroplastin (Stromal cell-derived receptor 1) (SDR-1) | NPTN SDFR1 SDR1 |

| | | | | | | | | | | | | | |
|---|---|---|---|---|---|---|---|---|---|---|---|---|---|
| sp\|P01031\|CO5_HUMAN | 7,481 | 4,967 | 24,506 | 8,354 | sp\|P01031\|CO5_HUMAN | 111,418 | 110,551 | 99% | P01031 | CO5_HUMAN | CO5_HUMAN | Complement C5 (C3 and PZP-like alpha-2-macroglobulin domain-containing protein 4) [Cleaved into: Complement C5 beta chain; Complement C5 alpha chain; C5a anaphylatoxin; Complement C5 alpha' chain] | C5 CPAMD4 |
| sp\|O95202\|LETM1_HUMAN | 7,374 | 9,132 | 8,994 | 13,641 | sp\|O95202\|LETM1_HUMAN | 18,749 | 11,093 | 59% | O95202 | LETM1_HUMAN | LETM1_HUMAN | Mitochondrial proton/calcium exchanger protein (Leucine zipper-EF-hand-containing transmembrane protein 1) | LETM1 |
| sp\|P27816\|MAP4_HUMAN | 5,844 | 25,869 | 5,522 | 22,704 | sp\|P27816\|MAP4_HUMAN | 16,612 | 9,269 | 56% | P27816 | MAP4_HUMAN | MAP4_HUMAN | Microtubule-associated protein 4 (MAP-4) | MAP4 |
| sp\|P33316\|DUT_HUMAN | 95,877 | 102,438 | 129,218 | 197,654 | sp\|P33316\|DUT_HUMAN | 142,055 | 53,715 | 38% | P33316 | DUT_HUMAN | DUT_HUMAN | Deoxyuridine 5'-triphosphate nucleotidohydrolase, mitochondrial (dUTPase) (EC 3.6.1.23) (dUTP pyrophosphatase) | DUT |
| sp\|P45974\|UBP5_HUMAN | 197,210 | 215,480 | 188,789 | 205,387 | sp\|P45974\|UBP5_HUMAN | 196,748 | 17,297 | 9% | P45974 | UBP5_HUMAN | UBP5_HUMAN | Ubiquitin carboxyl-terminal hydrolase 5 (EC 3.4.19.12) (Deubiquitinating enzyme 5) (Isopeptidase T) (Ubiquitin thioesterase 5) (Ubiquitin-specific-processing protease 5) | USP5 ISOT |
| sp\|Q9Y6E0\|STK24_HUMAN | 88,232 | 110,510 | 75,303 | 77,151 | sp\|Q9Y6E0\|STK24_HUMAN | 71,364 | 34,727 | 49% | Q9Y6E0 | STK24_HUMAN | STK24_HUMAN | Serine/threonine-protein kinase 24 (EC 2.7.11.1) (Mammalian STE20-like protein kinase 3) (MST-3) (STE20-like kinase MST3) [Cleaved into: Serine/threonine-protein kinase 24 36 kDa subunit (Mammalian STE20-like protein kinase 3 N-terminal) (MST3/N); Serine/threonine-protein kinase 24 12 kDa subunit (Mammalian STE20-like protein kinase 3 C-terminal) (MST3/C)] | STK24 MST3 STK3 |
| sp\|P62070\|RRAS2_HUMAN | 158,710 | 134,211 | 272,955 | 210,764 | sp\|P62070\|RRAS2_HUMAN | 203,692 | 95,762 | 47% | P62070 | RRAS2_HUMAN | RRAS2_HUMAN | Ras-related protein R-Ras2 (EC 3.6.5.-) (Ras-like protein TC21) (Teratocarcinoma oncogene) | RRAS2 TC21 |
| sp\|O95292\|VAPB_HUMAN | 42,655 | 41,312 | 20,569 | 30,146 | sp\|O95292\|VAPB_HUMAN | 36,825 | 16,532 | 45% | O95292 | VAPB_HUMAN | VAPB_HUMAN | Vesicle-associated membrane protein-associated protein B/C (VAMP-B/VAMP-C) (VAMP-associated protein B/C) (VAP-B/VAP-C) | VAPB UNQ484/PRO983 |
| sp\|Q14627\|I13R2_HUMAN | 287,239 | 296,327 | 214,558 | 162,028 | sp\|Q14627\|I13R2_HUMAN | 212,874 | 63,464 | 30% | Q14627 | I13R2_HUMAN | I13R2_HUMAN | Interleukin-13 receptor subunit alpha-2 (IL-13 receptor subunit alpha-2) (IL-13R subunit alpha-2) (IL-13R-alpha-2) (IL-13RA2) (Interleukin-13-binding protein) (CD antigen CD213a2) | IL13RA2 IL13R |
| sp\|O43617\|TPPC3_HUMAN | 28,507 | 33,988 | 11,239 | 23,078 | sp\|O43617\|TPPC3_HUMAN | 27,149 | 17,438 | 64% | O43617 | TPPC3_HUMAN | TPPC3_HUMAN | Trafficking protein particle complex subunit 3 (BET3 homolog) | TRAPPC3 BET3 CDABP0066 |
| sp\|P78310\|CXAR_HUMAN | 177,539 | 407,257 | 415,375 | 399,626 | sp\|P78310\|CXAR_HUMAN | 353,317 | 86,445 | 24% | P78310 | CXAR_HUMAN | CXAR_HUMAN | Coxsackievirus and adenovirus receptor (CAR) (hCAR) (CVB3-binding protein) (Coxsackievirus B-adenovirus receptor) (HCVADR) | CXADR CAR |

| | | | | | | | | | | | | | |
|---|---|---|---|---|---|---|---|---|---|---|---|---|---|
| sp\|Q99436\|PSB7_HUMAN | 253,639 | 259,983 | 302,811 | 268,730 | sp\|Q99436\|PSB7_HUMAN | 250,628 | 26,402 | 11% | Q99436 | PSB7_HUMAN | PSB7_HUMAN | Proteasome subunit beta type-7 (EC 3.4.25.1) (Macropain chain Z) (Multicatalytic endopeptidase complex chain Z) (Proteasome subunit Z) | PSMB7 Z |
| sp\|Q9H0E2\|TOLIP_HUMAN | 293,300 | 338,339 | 429,079 | 505,610 | sp\|Q9H0E2\|TOLIP_HUMAN | 411,335 | 98,419 | 24% | Q9H0E2 | TOLIP_HUMAN | TOLIP_HUMAN | Toll-interacting protein | TOLLIP |
| sp\|Q93096\|TP4A1_HUMAN | 162,597 | 163,404 | 140,195 | 204,800 | sp\|Q93096\|TP4A1_HUMAN | 161,870 | 62,121 | 38% | Q93096 | TP4A1_HUMAN | TP4A1_HUMAN | Protein tyrosine phosphatase type IVA 1 (EC 3.1.3.48) (PTP(CAAXI)) (Protein-tyrosine phosphatase 4a1) (Protein-tyrosine phosphatase of regenerating liver 1) (PRL-1) | PTP4A1 PRL1 PTPCAAX1 |
| sp\|P10643\|CO7_HUMAN | 69,978 | 30,532 | 99,990 | 74,257 | sp\|P10643\|CO7_HUMAN | 400,954 | 383,265 | 96% | P10643 | CO7_HUMAN | CO7_HUMAN | Complement component C7 | C7 |
| sp\|Q9H488\|OFUT1_HUMAN | 21,375 | 24,422 | 22,522 | 14,427 | sp\|Q9H488\|OFUT1_HUMAN | 18,948 | 8,053 | 42% | Q9H488 | OFUT1_HUMAN | OFUT1_HUMAN | GDP-fucose protein O-fucosyltransferase 1 (EC 2.4.1.221) (Peptide-O-fucosyltransferase 1) (O-FucT-1) | POFUT1 FUT12 KIAA0180 |
| sp\|P13929\|ENOB_HUMAN | 21,740 | 28,652 | 11,294 | 13,368 | sp\|P13929\|ENOB_HUMAN | 56,837 | 57,058 | 100% | P13929 | ENOB_HUMAN | ENOB_HUMAN | Beta-enolase (EC 4.2.1.11) (2-phospho-D-glycerate hydro-lyase) (Enolase 3) (Muscle-specific enolase) (MSE) (Skeletal muscle enolase) | ENO3 |
| sp\|P14174\|MIF_HUMAN | 409,199 | 799,796 | 311,595 | 562,808 | sp\|P14174\|MIF_HUMAN | 480,445 | 158,201 | 33% | P14174 | MIF_HUMAN | MIF_HUMAN | Macrophage migration inhibitory factor (MIF) (EC 5.3.2.1) (Glycosylation-inhibiting factor) (GIF) (L-dopachrome isomerase) (L-dopachrome tautomerase) (EC 5.3.3.12) (Phenylpyruvate tautomerase) | MIF GLIF MMIF |
| sp\|P55010\|IF5_HUMAN | 60,562 | 99,159 | 87,659 | 63,094 | sp\|P55010\|IF5_HUMAN | 68,793 | 15,714 | 23% | P55010 | IF5_HUMAN | IF5_HUMAN | Eukaryotic translation initiation factor 5 (eIF-5) | EIF5 |
| sp\|Q14257\|RCN2_HUMAN | 19,986 | 22,358 | 20,041 | 60,667 | sp\|Q14257\|RCN2_HUMAN | 32,888 | 23,065 | 70% | Q14257 | RCN2_HUMAN | RCN2_HUMAN | Reticulocalbin-2 (Calcium-binding protein ERC-55) (E6-binding protein) (E6BP) | RCN2 ERC55 |
| sp\|P07858\|CATB_HUMAN | 35,997 | 26,922 | 47,817 | 51,948 | sp\|P07858\|CATB_HUMAN | 27,545 | 16,426 | 60% | P07858 | CATB_HUMAN | CATB_HUMAN | Cathepsin B (EC 3.4.22.1) (APP secretase) (APPS) (Cathepsin B1) [Cleaved into: Cathepsin B light chain; Cathepsin B heavy chain] | CTSB CPSB |
| sp\|P63220\|RS21_HUMAN | 102,715 | 91,145 | 152,999 | 172,592 | sp\|P63220\|RS21_HUMAN | 114,150 | 47,144 | 41% | P63220 | RS21_HUMAN | RS21_HUMAN | 40S ribosomal protein S21 (Small ribosomal subunit protein eS21) | RPS21 |
| sp\|Q9Y240\|CLC11_HUMAN | 13,651 | 12,610 | 52,135 | 62,224 | sp\|Q9Y240\|CLC11_HUMAN | 134,004 | 147,422 | 110% | Q9Y240 | CLC11_HUMAN | CLC11_HUMAN | C-type lectin domain family 11 member A (C-type lectin superfamily member 3) (Lymphocyte secreted C-type lectin) (Osteolectin) (Stem cell growth factor) (p47) | CLEC11A CLECSF3 LSLCL SCGF |
| sp\|Q5ZPR3\|CD276_HUMAN | 188,288 | 170,405 | 327,825 | 437,990 | sp\|Q5ZPR3\|CD276_HUMAN | 270,073 | 106,761 | 40% | Q5ZPR3 | CD276_HUMAN | CD276_HUMAN | CD276 antigen (4Ig-B7-H3) (B7 homolog 3) (B7-H3) (Costimulatory molecule) (CD antigen CD276) | CD276 B7H3 PSEC0249 UNQ309/PRO352 |
| sp\|Q9Y4W6\|AFG32_HUMAN | 8,619 | 27,478 | 8,451 | 15,952 | sp\|Q9Y4W6\|AFG32_HUMAN | 17,741 | 6,587 | 37% | Q9Y4W6 | AFG32_HUMAN | AFG32_HUMAN | AFG3-like protein 2 (EC 3.4.24.-) (Paraplegin-like protein) | AFG3L2 |
| sp\|Q9UNH7\|SNX6_HUMAN | 40,648 | 61,204 | 37,217 | 32,462 | sp\|Q9UNH7\|SNX6_HUMAN | 37,373 | 11,025 | 30% | Q9UNH7 | SNX6_HUMAN | SNX6_HUMAN | Sorting nexin-6 (TRAF4-associated factor 2) [Cleaved into: Sorting nexin-6, N-terminally processed] | SNX6 |
| sp\|P20339\|RAB5A_HUMAN | 246,657 | 254,020 | 409,595 | 319,678 | sp\|P20339\|RAB5A_HUMAN | 313,788 | 72,001 | 23% | P20339 | RAB5A_HUMAN | RAB5A_HUMAN | Ras-related protein Rab-5A (EC 3.6.5.2) | RAB5A RAB5 |
| sp\|Q86UX7\|URP2_HUMAN | 40,201 | 241,843 | 107,492 | 50,550 | sp\|Q86UX7\|URP2_HUMAN | 265,348 | 191,082 | 72% | Q86UX7 | URP2_HUMAN | URP2_HUMAN | Fermitin family homolog 3 (Kindlin-3) (MIG2-like protein) (Unc-112-related protein 2) | FERMT3 KIND3 MIG2B URP2 |
| sp\|P15151\|PVR_HUMAN | 623,350 | 556,186 | 643,785 | 751,123 | sp\|P15151\|PVR_HUMAN | 565,580 | 133,222 | 24% | P15151 | PVR_HUMAN | PVR_HUMAN | Poliovirus receptor (Nectin-like protein 5) (NECL-5) (CD antigen CD155) | PVR PVS |

| | | | | | | | | | | | | | |
|---|---|---|---|---|---|---|---|---|---|---|---|---|---|
| sp\|O75251\|NDUS7_HUMAN | 11,584 | 22,283 | 34,493 | 9,195 | sp\|O75251\|NDUS7_HUMAN | 25,622 | 16,667 | 65% | O75251 | NDUS7_HUMAN | NDUS7_HUMAN | NADH dehydrogenase [ubiquinone] iron-sulfur protein 7, mitochondrial (EC 7.1.1.2) (Complex I-20kD) (CI-20kD) (NADH-ubiquinone oxidoreductase 20 kDa subunit) (PSST subunit) | NDUFS7 |
| sp\|P04271\|S100B_HUMAN | 4,496 | 10,675 | 122,833 | 12,548 | sp\|P04271\|S100B_HUMAN | 66,746 | 117,662 | 176% | P04271 | S100B_HUMAN | S100B_HUMAN | Protein S100-B (S-100 protein beta chain) (S-100 protein subunit beta) (S100 calcium-binding protein B) | S100B |
| sp\|Q96ND0\|F210A_HUMAN | 33,608 | 35,347 | 22,788 | 37,224 | sp\|Q96ND0\|F210A_HUMAN | 29,249 | 7,591 | 26% | Q96ND0 | F210A_HUMAN | F210A_HUMAN | Protein FAM210A | FAM210A C18orf19 |
| sp\|Q06323\|PSME1_HUMAN | 2,928 | 33,328 | 11,578 | 40,696 | sp\|Q06323\|PSME1_HUMAN | 30,899 | 17,222 | 56% | Q06323 | PSME1_HUMAN | PSME1_HUMAN | Proteasome activator complex subunit 1 (11S regulator complex subunit alpha) (REG-alpha) (Activator of multicatalytic protease subunit 1) (Interferon gamma up-regulated I-5111 protein) (IGUP I-5111) (Proteasome activator 28 subunit alpha) (PA28a) (PA28alpha) | PSME1 IFI5111 |
| sp\|Q12904\|AIMP1_HUMAN | 54,072 | 78,348 | 93,931 | 132,515 | sp\|Q12904\|AIMP1_HUMAN | 83,716 | 24,556 | 29% | Q12904 | AIMP1_HUMAN | AIMP1_HUMAN | Aminoacyl tRNA synthase complex-interacting multifunctional protein 1 (Multisynthase complex auxiliary component p43) [Cleaved into: Endothelial monocyte-activating polypeptide 2 (EMAP-2) (Endothelial monocyte-activating polypeptide II) (EMAP-II) (Small inducible cytokine subfamily E member 1)] | AIMP1 EMAP2 SCYE1 |
| sp\|P02788\|TRFL_HUMAN | 6,326 | 10,819 | 55,476 | 7,841 | sp\|P02788\|TRFL_HUMAN | 631,606 | 1,228,289 | 194% | P02788 | TRFL_HUMAN | TRFL_HUMAN | Lactotransferrin (Lactoferrin) (EC 3.4.21.-) (Growth-inhibiting protein 12) (Talalactoferrin) [Cleaved into: Lactoferricin-H (Lfcin-H); Kaliocin-1; Lactoferroxin-A; Lactoferroxin-B; Lactoferroxin-C] | LTF GIG12 LF |
| sp\|P24666\|PPAC_HUMAN | 212,978 | 274,984 | 294,588 | 319,624 | sp\|P24666\|PPAC_HUMAN | 260,606 | 48,488 | 19% | P24666 | PPAC_HUMAN | PPAC_HUMAN | Low molecular weight phosphotyrosine protein phosphatase (LMW-PTP) (LMW-PTPase) (EC 3.1.3.48) (Adipocyte acid phosphatase) (Low molecular weight cytosolic acid phosphatase) (EC 3.1.3.2) (Red cell acid phosphatase 1) | ACP1 |
| sp\|O75436\|VP26A_HUMAN | 123,918 | 154,402 | 166,022 | 165,553 | sp\|O75436\|VP26A_HUMAN | 142,057 | 26,002 | 18% | O75436 | VP26A_HUMAN | VP26A_HUMAN | Vacuolar protein sorting-associated protein 26A (Vesicle protein sorting 26A) (hVPS26) | VPS26A VPS26 |
| sp\|O15554\|KCNN4_HUMAN | 156,917 | 171,940 | 76,348 | 141,212 | sp\|O15554\|KCNN4_HUMAN | 125,329 | 35,771 | 29% | O15554 | KCNN4_HUMAN | KCNN4_HUMAN | Intermediate conductance calcium-activated potassium channel protein 4 (SK4) (SKCa 4) (SKCa4) (IKCa1) (IK1) (KCa3.1) (KCa4) (Putative Gardos channel) | KCNN4 IK1 IKCA1 KCA4 SK4 |
| sp\|O95299\|NDUAA_HUMAN | 19,612 | 9,141 | 17,336 | 24,328 | sp\|O95299\|NDUAA_HUMAN | 20,216 | 12,198 | 60% | O95299 | NDUAA_HUMAN | NDUAA_HUMAN | NADH dehydrogenase [ubiquinone] 1 alpha subcomplex subunit 10, mitochondrial (Complex I-42kD) (CI-42kD) (NADH-ubiquinone oxidoreductase 42 kDa subunit) | NDUFA10 |

| | | | | | | | | | | | | | |
|---|---|---|---|---|---|---|---|---|---|---|---|---|---|
| sp\|Q6IBS0\|TWF2_HUMAN | 23,056 | 20,639 | 45,891 | 45,808 | sp\|Q6IBS0\|TWF2_HUMAN | 48,949 | 23,626 | 48% | Q6IBS0 | TWF2_HUMAN | TWF2_HUMAN | Twinfilin-2 (A6-related protein) (hA6RP) (Protein tyrosine kinase 9-like) (Twinfilin-1-like protein) | TWF2 PTK9L MSTP011 |
| sp\|O75915\|PRAF3_HUMAN | 230,002 | 187,994 | 149,858 | 232,818 | sp\|O75915\|PRAF3_HUMAN | 145,866 | 66,056 | 45% | O75915 | PRAF3_HUMAN | PRAF3_HUMAN | PRA1 family protein 3 (ADP-ribosylation factor-like protein 6-interacting protein 5) (ARL-6-interacting protein 5) (Aip-5) (Cytoskeleton-related vitamin A-responsive protein) (Dermal papilla-derived protein 11) (GTRAP3-18) (Glutamate transporter EAAC1-interacting protein) (JM5) (Prenylated Rab acceptor protein 2) (Protein JWa) (Putative MAPK-activating protein PM27) | ARL6IP5 DERP11 JWA PRA2 PRAF3 HSPC127 |
| sp\|P68036\|UB2L3_HUMAN | 59,593 | 126,997 | 86,830 | 69,239 | sp\|P68036\|UB2L3_HUMAN | 71,525 | 31,764 | 44% | P68036 | UB2L3_HUMAN | UB2L3_HUMAN | Ubiquitin-conjugating enzyme E2 L3 (EC 2.3.2.23) (E2 ubiquitin-conjugating enzyme L3) (L-UBC) (UbcH7) (Ubiquitin carrier protein L3) (Ubiquitin-conjugating enzyme E2-F1) (Ubiquitin-protein ligase L3) | UBE2L3 UBCE7 UBCH7 |
| sp\|P20290\|BTF3_HUMAN | 28,580 | 65,959 | 10,111 | 31,610 | sp\|P20290\|BTF3_HUMAN | 28,506 | 21,492 | 75% | P20290 | BTF3_HUMAN | BTF3_HUMAN | Transcription factor BTF3 (Nascent polypeptide-associated complex subunit beta) (NAC-beta) (RNA polymerase B transcription factor 3) | BTF3 NACB OK/SW-cl.8 |
| sp\|P68032\|ACTC_HUMAN | 2,974,513 | 3,568,000 | 544,932 | 839,007 | sp\|P68032\|ACTC_HUMAN | 1,522,316 | 1,312,836 | 86% | P68032 | ACTC_HUMAN | ACTC_HUMAN | Actin, alpha cardiac muscle 1 (Alpha-cardiac actin) [Cleaved into: Actin, alpha cardiac muscle 1, intermediate form] | ACTC1 ACTC |
| sp\|P05026\|AT1B1_HUMAN | 76,328 | 114,981 | 72,411 | 32,004 | sp\|P05026\|AT1B1_HUMAN | 84,477 | 26,678 | 32% | P05026 | AT1B1_HUMAN | AT1B1_HUMAN | Sodium/potassium-transporting ATPase subunit beta-1 (Sodium/potassium-dependent ATPase subunit beta-1) | ATP1B1 ATP1B |
| sp\|P53985\|MOT1_HUMAN | 354,130 | 357,432 | 371,451 | 522,667 | sp\|P53985\|MOT1_HUMAN | 403,247 | 58,444 | 14% | P53985 | MOT1_HUMAN | MOT1_HUMAN | Monocarboxylate transporter 1 (MCT 1) (Solute carrier family 16 member 1) | SLC16A1 MCT1 |
| sp\|P53602\|MVD1_HUMAN | 53,535 | 55,522 | 83,378 | 81,643 | sp\|P53602\|MVD1_HUMAN | 58,861 | 23,182 | 39% | P53602 | MVD1_HUMAN | MVD1_HUMAN | Diphosphomevalonate decarboxylase (EC 4.1.1.33) (Mevalonate (diphospho)decarboxylase) (MDDase) (Mevalonate pyrophosphate decarboxylase) | MVD MPD |
| sp\|O75955\|FLOT1_HUMAN | 244,522 | 248,239 | 465,510 | 257,958 | sp\|O75955\|FLOT1_HUMAN | 223,658 | 133,972 | 60% | O75955 | FLOT1_HUMAN | FLOT1_HUMAN | Flotillin-1 | FLOT1 |
| sp\|Q15257\|PTPA_HUMAN | 52,454 | 109,755 | 40,500 | 112,843 | sp\|Q15257\|PTPA_HUMAN | 87,725 | 29,598 | 34% | Q15257 | PTPA_HUMAN | PTPA_HUMAN | Serine/threonine-protein phosphatase 2A activator (EC 5.2.1.8) (PP2A, subunit B', PR53 isoform) (Phosphotyrosyl phosphatase activator) (PTPA) (Serine/threonine-protein phosphatase 2A regulatory subunit 4) (Serine/threonine-protein phosphatase 2A regulatory subunit B') | PTPA PPP2R4 |
| sp\|P49748\|ACADV_HUMAN | 19,252 | 12,912 | 86,469 | 19,166 | sp\|P49748\|ACADV_HUMAN | 47,732 | 42,207 | 88% | P49748 | ACADV_HUMAN | ACADV_HUMAN | Very long-chain specific acyl-CoA dehydrogenase, mitochondrial (VLCAD) (EC 1.3.8.9) | ACADVL VLCAD |

| | | | | | | | | | | | | | |
|---|---|---|---|---|---|---|---|---|---|---|---|---|---|
| sp\|Q9NZL4\|HPBP1_HUMAN | 31,534 | 32,287 | 26,217 | 10,376 | sp\|Q9NZL4\|HPBP1_HUMAN | 29,113 | 14,645 | 50% | Q9NZL4 | HPBP1_HUMAN | HPBP1_HUMAN | Hsp70-binding protein 1 (HspBP1) (Heat shock protein-binding protein 1) (Hsp70-binding protein 2) (HspBP2) (Hsp70-interacting protein 1) (Hsp70-interacting protein 2) | HSPBP1 HSPBP PP1845 |
| sp\|Q9C004\|SPY4_HUMAN | 93,222 | 129,554 | 231,588 | 185,128 | sp\|Q9C004\|SPY4_HUMAN | 128,089 | 54,399 | 42% | Q9C004 | SPY4_HUMAN | SPY4_HUMAN | Protein sprouty homolog 4 (Spry-4) | SPRY4 |
| sp\|Q14956\|GPNMB_HUMAN | 446,357 | 406,568 | 329,603 | 234,863 | sp\|Q14956\|GPNMB_HUMAN | 249,377 | 130,354 | 52% | Q14956 | GPNMB_HUMAN | GPNMB_HUMAN | Transmembrane glycoprotein NMB (Hematopoietic growth factor inducible neurokinin-1 type) | GPNMB HGFIN NMB UNQ1725/PRO9925 |
| sp\|Q14166\|TTL12_HUMAN | 46,565 | 58,879 | 45,102 | 44,847 | sp\|Q14166\|TTL12_HUMAN | 34,962 | 20,417 | 58% | Q14166 | TTL12_HUMAN | TTL12_HUMAN | Tubulin--tyrosine ligase-like protein 12 (Inactive tubulin--tyrosine ligase-like protein 12) | TTLL12 KIAA0153 |
| sp\|P56134\|ATPK_HUMAN | 84,783 | 96,999 | 105,334 | 158,060 | sp\|P56134\|ATPK_HUMAN | 85,299 | 38,116 | 45% | P56134 | ATPK_HUMAN | ATPK_HUMAN | ATP synthase subunit f, mitochondrial (ATP synthase membrane subunit f) | ATP5MF ATP5J2 ATP5JL |
| sp\|P00390\|GSHR_HUMAN | 54,149 | 85,366 | 68,611 | 88,146 | sp\|P00390\|GSHR_HUMAN | 66,436 | 16,557 | 25% | P00390 | GSHR_HUMAN | GSHR_HUMAN | Glutathione reductase, mitochondrial (GR) (GRase) (EC 1.8.1.7) | GSR GLUR GRD1 |
| sp\|P62834\|RAP1A_HUMAN | 695,597 | 758,335 | 243,855 | 218,014 | sp\|P62834\|RAP1A_HUMAN | 403,521 | 270,969 | 67% | P62834 | RAP1A_HUMAN | RAP1A_HUMAN | Ras-related protein Rap-1A (EC 3.6.5.2) (C21KG) (G-22K) (GTP-binding protein smg p21A) (Ras-related protein Krev-1) | RAP1A KREV1 |
| sp\|P49821\|NDUV1_HUMAN | 18,548 | 17,360 | 31,870 | 13,611 | sp\|P49821\|NDUV1_HUMAN | 23,652 | 22,142 | 94% | P49821 | NDUV1_HUMAN | NDUV1_HUMAN | NADH dehydrogenase [ubiquinone] flavoprotein 1, mitochondrial (EC 7.1.1.2) (Complex I-51kD) (CI-51kD) (NADH dehydrogenase flavoprotein 1) (NADH-ubiquinone oxidoreductase 51 kDa subunit) | NDUFV1 UQOR1 |
| sp\|Q9Y5X2\|SNX8_HUMAN | 199,518 | 61,865 | 115,620 | 166,104 | sp\|Q9Y5X2\|SNX8_HUMAN | 118,585 | 51,619 | 44% | Q9Y5X2 | SNX8_HUMAN | SNX8_HUMAN | Sorting nexin-8 | SNX8 |
| sp\|Q9UBQ7\|GRHPR_HUMAN | 57,380 | 71,627 | 26,969 | 46,296 | sp\|Q9UBQ7\|GRHPR_HUMAN | 56,083 | 24,429 | 44% | Q9UBQ7 | GRHPR_HUMAN | GRHPR_HUMAN | Glyoxylate reductase/hydroxypyruvate reductase (EC 1.1.1.79) (EC 1.1.1.81) | GRHPR GLXR MSTP035 |
| sp\|Q9UNN8\|EPCR_HUMAN | 523,868 | 480,113 | 771,950 | 837,117 | sp\|Q9UNN8\|EPCR_HUMAN | 584,405 | 185,863 | 32% | Q9UNN8 | EPCR_HUMAN | EPCR_HUMAN | Endothelial protein C receptor (Activated protein C receptor) (APC receptor) (Endothelial cell protein C receptor) (CD antigen CD201) | PROCR EPCR |
| sp\|Q96DH6\|MSI2H_HUMAN | 13,147 | 27,278 | 21,744 | 14,969 | sp\|Q96DH6\|MSI2H_HUMAN | 17,417 | 7,712 | 44% | Q96DH6 | MSI2H_HUMAN | MSI2H_HUMAN | RNA-binding protein Musashi homolog 2 (Musashi-2) | MSI2 |
| sp\|Q16563\|SYPL1_HUMAN | 1,005,512 | 717,552 | 592,360 | 563,790 | sp\|Q16563\|SYPL1_HUMAN | 530,785 | 243,603 | 46% | Q16563 | SYPL1_HUMAN | SYPL1_HUMAN | Synaptophysin-like protein 1 (Pantophysin) | SYPL1 SYPL |
| sp\|Q15125\|EBP_HUMAN | 6,031 | 56,498 | 79,341 | 18,436 | sp\|Q15125\|EBP_HUMAN | 40,522 | 26,499 | 65% | Q15125 | EBP_HUMAN | EBP_HUMAN | 3-beta-hydroxysteroid-Delta(8),Delta(7)-isomerase (EC 5.3.3.5) (Cholestenol Delta-isomerase) (Delta(8)-Delta(7) sterol isomerase) (D8-D7 sterol isomerase) (Emopamil-binding protein) | EBP |
| sp\|Q04941\|PLP2_HUMAN | 1,354,850 | 1,034,579 | 2,155,279 | 2,316,077 | sp\|Q04941\|PLP2_HUMAN | 1,402,819 | 575,115 | 41% | Q04941 | PLP2_HUMAN | PLP2_HUMAN | Proteolipid protein 2 (Differentiation-dependent protein A4) (Intestinal membrane A4 protein) | PLP2 A4 |
| sp\|P68871\|HBB_HUMAN | 1,721,085 | 1,036,666 | 2,939,484 | 1,247,033 | sp\|P68871\|HBB_HUMAN | 21,570,890 | 22,522,436 | 104% | P68871 | HBB_HUMAN | HBB_HUMAN | Hemoglobin subunit beta (Beta-globin) (Hemoglobin beta chain) [Cleaved into: LVV-hemorphin-7; Spinorphin] | HBB |

| | | | | | | | | | | | | | |
|---|---|---|---|---|---|---|---|---|---|---|---|---|---|
| sp\|P68402\|PA1B2_HUMAN | 119,835 | 100,230 | 76,247 | 122,931 | sp\|P68402\|PA1B2_HUMAN | 100,054 | 16,863 | 17% | P68402 | PA1B2_HUMAN | PA1B2_HUMAN | Platelet-activating factor acetylhydrolase IB subunit alpha2 (EC 3.1.1.47) (PAF acetylhydrolase 30 kDa subunit) (PAF-AH 30 kDa subunit) (PAF-AH subunit beta) (PAFAH subunit beta) | PAFAH1B2 PAFAHB |
| sp\|P63173\|RL38_HUMAN | 3,590 | 68,255 | 30,101 | 35,076 | sp\|P63173\|RL38_HUMAN | 26,039 | 23,911 | 92% | P63173 | RL38_HUMAN | RL38_HUMAN | 60S ribosomal protein L38 (Large ribosomal subunit protein eL38) | RPL38 |
| sp\|P62318\|SMD3_HUMAN | 335,849 | 314,529 | 233,889 | 273,670 | sp\|P62318\|SMD3_HUMAN | 193,299 | 108,174 | 56% | P62318 | SMD3_HUMAN | SMD3_HUMAN | Small nuclear ribonucleoprotein Sm D3 (Sm-D3) (snRNP core protein D3) | SNRPD3 |
| sp\|P61916\|NPC2_HUMAN | 44,521 | 35,743 | 44,285 | 61,994 | sp\|P61916\|NPC2_HUMAN | 40,517 | 17,234 | 43% | P61916 | NPC2_HUMAN | NPC2_HUMAN | NPC intracellular cholesterol transporter 2 (Epididymal secretory protein E1) (Human epididymis-specific protein 1) (He1) (Niemann-Pick disease type C2 protein) | NPC2 HE1 |
| sp\|P46976\|GLYG_HUMAN | 852,612 | 482,824 | 951,099 | 448,469 | sp\|P46976\|GLYG_HUMAN | 536,488 | 400,263 | 75% | P46976 | GLYG_HUMAN | GLYG_HUMAN | Glycogenin-1 (GN-1) (GN1) (EC 2.4.1.186) | GYG1 GYG |
| sp\|P14209\|CD99_HUMAN | 629,615 | 450,312 | 1,544,087 | 1,562,522 | sp\|P14209\|CD99_HUMAN | 840,282 | 468,445 | 56% | P14209 | CD99_HUMAN | CD99_HUMAN | CD99 antigen (12E7) (E2 antigen) (Protein MIC2) (T-cell surface glycoprotein E2) (CD antigen CD99) | CD99 MIC2 MIC2X MIC2Y |
| sp\|P11166\|GTR1_HUMAN | 315,693 | 261,092 | 315,444 | 386,771 | sp\|P11166\|GTR1_HUMAN | 508,370 | 252,712 | 50% | P11166 | GTR1_HUMAN | GTR1_HUMAN | Solute carrier family 2, facilitated glucose transporter member 1 (Glucose transporter type 1, erythrocyte/brain) (GLUT-1) (HepG2 glucose transporter) | SLC2A1 GLUT1 |
| sp\|O43169\|CYB5B_HUMAN | 49,844 | 75,163 | 45,206 | 19,356 | sp\|O43169\|CYB5B_HUMAN | 47,331 | 16,462 | 35% | O43169 | CYB5B_HUMAN | CYB5B_HUMAN | Cytochrome b5 type B (Cytochrome b5 outer mitochondrial membrane isoform) | CYB5B CYB5M OMB5 |
| sp\|O14828\|SCAM3_HUMAN | 233,110 | 111,298 | 409,821 | 332,016 | sp\|O14828\|SCAM3_HUMAN | 197,369 | 117,105 | 59% | O14828 | SCAM3_HUMAN | SCAM3_HUMAN | Secretory carrier-associated membrane protein 3 (Secretory carrier membrane protein 3) | SCAMP3 C1orf3 PROPIN1 |
| sp\|Q92930\|RAB8B_HUMAN | 112,188 | 166,371 | 154,495 | 213,003 | sp\|Q92930\|RAB8B_HUMAN | 159,959 | 45,844 | 29% | Q92930 | RAB8B_HUMAN | RAB8B_HUMAN | Ras-related protein Rab-8B | RAB8B |
| sp\|P30536\|TSPO_HUMAN | 23,370 | 29,106 | 34,224 | 56,292 | sp\|P30536\|TSPO_HUMAN | 26,368 | 18,155 | 69% | P30536 | TSPO_HUMAN | TSPO_HUMAN | Translocator protein (Mitochondrial benzodiazepine receptor) (PKBS) (Peripheral-type benzodiazepine receptor) (PBR) | TSPO BZRP MBR |
| sp\|P49773\|HINT1_HUMAN | 47,833 | 55,366 | 44,239 | 35,333 | sp\|P49773\|HINT1_HUMAN | 39,471 | 11,096 | 28% | P49773 | HINT1_HUMAN | HINT1_HUMAN | Adenosine 5'-monophosphoramidase HINT1 (EC 3.9.1.-) (Desumoylating isopeptidase HINT1) (EC 3.4.22.-) (Histidine triad nucleotide-binding protein 1) (Protein kinase C inhibitor 1) (Protein kinase C-interacting protein 1) (PKCI-1) | HINT1 HINT PKCI1 PRKCNH1 |
| sp\|Q9H1E5\|TMX4_HUMAN | 2,210 | 264 | 1,492 | 1,315 | sp\|Q9H1E5\|TMX4_HUMAN | 8,421 | 11,307 | 134% | Q9H1E5 | TMX4_HUMAN | TMX4_HUMAN | Thioredoxin-related transmembrane protein 4 (Thioredoxin domain-containing protein 13) | TMX4 KIAA1162 TXNDC13 PSEC0095 UNQ475/PRO938 |
| sp\|O75718\|CRTAP_HUMAN | 12,884 | 23,480 | 7,041 | 18,862 | sp\|O75718\|CRTAP_HUMAN | 15,426 | 8,549 | 55% | O75718 | CRTAP_HUMAN | CRTAP_HUMAN | Cartilage-associated protein | CRTAP CASP |
| sp\|Q86TX2\|ACOT1_HUMAN | 54,231 | 61,260 | 87,871 | 108,219 | sp\|Q86TX2\|ACOT1_HUMAN | 69,421 | 24,973 | 36% | Q86TX2 | ACOT1_HUMAN | ACOT1_HUMAN | Acyl-coenzyme A thioesterase 1 (Acyl-CoA thioesterase 1) (EC 3.1.2.-) (CTE-I) (CTE-Ib) (Inducible cytosolic acyl-coenzyme A thioester hydrolase) (Long chain acyl-CoA thioester hydrolase) (Long chain acyl-CoA hydrolase) (Palmitoyl-coenzyme A thioesterase) (EC 3.1.2.2) | ACOT1 CTE1 |

| | | | | | | | | | | | | | |
|---|---|---|---|---|---|---|---|---|---|---|---|---|---|
| sp\|Q9BRQ8\|AIFM2_HUMAN | 35,621 | 38,105 | 7,052 | 65,261 | sp\|Q9BRQ8\|AIFM2_HUMAN | 25,147 | 20,122 | 80% | Q9BRQ8 | AIFM2_HUMAN | FSP1_HUMAN | Ferroptosis suppressor protein 1 (FSP1) (EC 1.6.5.-) (Apoptosis-inducing factor homologous mitochondrion-associated inducer of death) (AMID) (p53-responsive gene 3 protein) | AIFM2 AMID PRG3 |
| sp\|P04233\|HG2A_HUMAN | 61,125 | 66,792 | 116,457 | 96,024 | sp\|P04233\|HG2A_HUMAN | 60,591 | 35,027 | 58% | P04233 | HG2A_HUMAN | HG2A_HUMAN | HLA class II histocompatibility antigen gamma chain (HLA-DR antigens-associated invariant chain) (Ia antigen-associated invariant chain) (Ii) (CD antigen CD74) [Cleaved into: Class-II-associated invariant chain peptide (CLIP)] | CD74 DHLAG |
| sp\|P02675\|FIBB_HUMAN | 31,657 | 12,013 | 87,199 | 46,798 | sp\|P02675\|FIBB_HUMAN | 347,022 | 339,977 | 98% | P02675 | FIBB_HUMAN | FIBB_HUMAN | Fibrinogen beta chain [Cleaved into: Fibrinopeptide B; Fibrinogen beta chain] | FGB |
| sp\|Q15436\|SC23A_HUMAN | 28,043 | 28,045 | 58,842 | 49,283 | sp\|Q15436\|SC23A_HUMAN | 33,993 | 17,759 | 52% | Q15436 | SC23A_HUMAN | SC23A_HUMAN | Protein transport protein Sec23A (hSec23A) (SEC23-related protein A) | SEC23A |
| sp\|Q9NZ01\|TECR_HUMAN | 3,396 | 5,453 | 1,588 | 1,680 | sp\|Q9NZ01\|TECR_HUMAN | 2,326 | 1,440 | 62% | Q9NZ01 | TECR_HUMAN | TECR_HUMAN | Very-long-chain enoyl-CoA reductase (EC 1.3.1.93) (Synaptic glycoprotein SC2) (Trans-2,3-enoyl-CoA reductase) (TER) | TECR GPSN2 SC2 |
| sp\|P63167\|DYL1_HUMAN | 48,237 | 91,792 | 4,947 | 12,273 | sp\|P63167\|DYL1_HUMAN | 32,486 | 31,891 | 98% | P63167 | DYL1_HUMAN | DYL1_HUMAN | Dynein light chain 1, cytoplasmic (8 kDa dynein light chain) (DLC8) (Dynein light chain LC8-type 1) (Protein inhibitor of neuronal nitric oxide synthase) (PIN) | DYNLL1 DLC1 DNCL1 DNCLC1 HDLC1 |
| sp\|P36404\|ARL2_HUMAN | 190,039 | 196,857 | 219,123 | 158,835 | sp\|P36404\|ARL2_HUMAN | 199,792 | 61,330 | 31% | P36404 | ARL2_HUMAN | ARL2_HUMAN | ADP-ribosylation factor-like protein 2 | ARL2 |
| sp\|Q70UQ0\|IKIP_HUMAN | 11,188 | 9,557 | 14,434 | 5,736 | sp\|Q70UQ0\|IKIP_HUMAN | 11,655 | 3,851 | 33% | Q70UQ0 | IKIP_HUMAN | IKIP_HUMAN | Inhibitor of nuclear factor kappa-B kinase-interacting protein (I kappa-B kinase-interacting protein) (IKBKB-interacting protein) (IKK-interacting protein) | IKBIP IKIP |
| sp\|P01011\|AACT_HUMAN | 95,257 | 89,995 | 88,468 | 89,929 | sp\|P01011\|AACT_HUMAN | 80,996 | 34,568 | 43% | P01011 | AACT_HUMAN | AACT_HUMAN | Alpha-1-antichymotrypsin (ACT) (Cell growth-inhibiting gene 24/25 protein) (Serpin A3) [Cleaved into: Alpha-1-antichymotrypsin His-Pro-less] | SERPINA3 AACT GIG24 GIG25 |
| sp\|O43143\|DHX15_HUMAN | 146,706 | 97,094 | 102,969 | 57,219 | sp\|O43143\|DHX15_HUMAN | 83,883 | 33,566 | 40% | O43143 | DHX15_HUMAN | DHX15_HUMAN | ATP-dependent RNA helicase DHX15 (EC 3.6.4.13) (ATP-dependent RNA helicase #46) (DEAH box protein 15) (Splicing factor Prp43) (hPrp43) | DHX15 DBP1 DDX15 |
| sp\|P02679\|FIBG_HUMAN | 7,613 | 3,808 | 16,986 | 38,411 | sp\|P02679\|FIBG_HUMAN | 112,145 | 117,124 | 104% | P02679 | FIBG_HUMAN | FIBG_HUMAN | Fibrinogen gamma chain | FGG PRO2061 |
| sp\|Q16576\|RBBP7_HUMAN | 191,904 | 238,003 | 172,092 | 193,847 | sp\|Q16576\|RBBP7_HUMAN | 136,145 | 77,009 | 57% | Q16576 | RBBP7_HUMAN | RBBP7_HUMAN | Histone-binding protein RBBP7 (Histone acetyltransferase type B subunit 2) (Nucleosome-remodeling factor subunit RBAP46) (Retinoblastoma-binding protein 7) (RBBP-7) (Retinoblastoma-binding protein p46) | RBBP7 RBAP46 |
| sp\|P55011\|S12A2_HUMAN | 94,614 | 167,022 | 261,060 | 266,716 | sp\|P55011\|S12A2_HUMAN | 261,970 | 143,300 | 55% | P55011 | S12A2_HUMAN | S12A2_HUMAN | Solute carrier family 12 member 2 (Basolateral Na-K-Cl symporter) (Bumetanide-sensitive sodium-(potassium)-chloride cotransporter 2) | SLC12A2 NKCC1 |
| sp\|Q969H8\|MYDGF_HUMAN | 41,717 | 49,514 | 45,636 | 9,205 | sp\|Q969H8\|MYDGF_HUMAN | 32,719 | 18,638 | 57% | Q969H8 | MYDGF_HUMAN | MYDGF_HUMAN | Myeloid-derived growth factor (MYDGF) | MYDGF C19orf10 |

| | | | | | | | | | | | | | |
|---|---|---|---|---|---|---|---|---|---|---|---|---|---|
| sp\|P10155\|RO60_HUMAN | 57,961 | 86,056 | 36,432 | 67,692 | sp\|P10155\|RO60_HUMAN | 44,069 | 24,371 | 55% | P10155 | RO60_HUMAN | RO60_HUMAN | RNA-binding protein RO60 (60 kDa SS-A/Ro ribonucleoprotein) (60 kDa Ro protein) (60 kDa ribonucleoprotein Ro) (RoRNP) (Ro 60 kDa autoantigen) (Ro60 autoantigen) (Sjoegren syndrome antigen A2) (Sjoegren syndrome type A antigen) (SS-A) (TROVE domain family member 2) | RO60 SSA2 TROVE2 |
| sp\|Q92882\|OSTF1_HUMAN | 106,250 | 140,171 | 154,693 | 164,133 | sp\|Q92882\|OSTF1_HUMAN | 133,701 | 20,839 | 16% | Q92882 | OSTF1_HUMAN | OSTF1_HUMAN | Osteoclast-stimulating factor 1 | OSTF1 |
| sp\|Q9BVC6\|TM109_HUMAN | 141,418 | 88,546 | 192,963 | 168,689 | sp\|Q9BVC6\|TM109_HUMAN | 141,228 | 37,820 | 27% | Q9BVC6 | TM109_HUMAN | TM109_HUMAN | Transmembrane protein 109 (Mitsugumin-23) (Mg23) | TMEM109 |
| sp\|Q00839\|HNRPU_HUMAN | 399,626 | 277,781 | 330,476 | 222,639 | sp\|Q00839\|HNRPU_HUMAN | 207,719 | 127,225 | 61% | Q00839 | HNRPU_HUMAN | HNRPU_HUMAN | Heterogeneous nuclear ribonucleoprotein U (hnRNP U) (GRIP120) (Nuclear p120 ribonucleoprotein) (Scaffold-attachment factor A) (SAF-A) (p120) (pp120) | HNRNPU C1orf199 HNRPU SAFA U21.1 |
| sp\|Q9Y5S1\|TRPV2_HUMAN | 39,318 | 34,305 | 73,850 | 79,015 | sp\|Q9Y5S1\|TRPV2_HUMAN | 49,774 | 19,281 | 39% | Q9Y5S1 | TRPV2_HUMAN | TRPV2_HUMAN | Transient receptor potential cation channel subfamily V member 2 (TrpV2) (Osm-9-like TRP channel 2) (OTRPC2) (Vanilloid receptor-like protein 1) (VRL-1) | TRPV2 VRL |
| sp\|Q9UKM9\|RALY_HUMAN | 54,641 | 47,345 | 47,161 | 17,823 | sp\|Q9UKM9\|RALY_HUMAN | 32,484 | 17,315 | 53% | Q9UKM9 | RALY_HUMAN | RALY_HUMAN | RNA-binding protein Raly (Autoantigen p542) (Heterogeneous nuclear ribonucleoprotein C-like 2) (hnRNP core protein C-like 2) (hnRNP associated with lethal yellow protein homolog) | RALY HNRPCL2 P542 |
| sp\|P16435\|NCPR_HUMAN | 12,492 | 15,165 | 17,502 | 14,069 | sp\|P16435\|NCPR_HUMAN | 16,546 | 4,418 | 27% | P16435 | NCPR_HUMAN | NCPR_HUMAN | NADPH--cytochrome P450 reductase (CPR) (P450R) (EC 1.6.2.4) | POR CYPOR |
| sp\|O95834\|EMAL2_HUMAN | 13,815 | 29,410 | 22,281 | 58,972 | sp\|O95834\|EMAL2_HUMAN | 76,532 | 56,682 | 74% | O95834 | EMAL2_HUMAN | EMAL2_HUMAN | Echinoderm microtubule-associated protein-like 2 (EMAP-2) (HuEMAP-2) | EML2 EMAP2 EMAPL2 |
| sp\|Q14558\|KPRA_HUMAN | 24,693 | 28,732 | 63,894 | 102,617 | sp\|Q14558\|KPRA_HUMAN | 105,471 | 69,671 | 66% | Q14558 | KPRA_HUMAN | KPRA_HUMAN | Phosphoribosyl pyrophosphate synthase-associated protein 1 (PRPP synthase-associated protein 1) (39 kDa phosphoribosypyrophosphate synthase-associated protein) (PAP39) | PRPSAP1 |
| sp\|Q3LXA3\|TKFC_HUMAN | 5,029 | 13,052 | 3,021 | 20,795 | sp\|Q3LXA3\|TKFC_HUMAN | 10,437 | 6,873 | 66% | Q3LXA3 | TKFC_HUMAN | TKFC_HUMAN | Triokinase/FMN cyclase (Bifunctional ATP-dependent dihydroxyacetone kinase/FAD-AMP lyase (cyclizing)) [Includes: ATP-dependent dihydroxyacetone kinase (DHA kinase) (EC 2.7.1.28) (EC 2.7.1.29) (Glycerone kinase) (Triokinase) (Triose kinase); FAD-AMP lyase (cyclizing) (EC 4.6.1.15) (FAD-AMP lyase (cyclic FMN forming)) (FMN cyclase)] | TKFC DAK |
| sp\|P61009\|SPCS3_HUMAN | 34,828 | 34,521 | 15,656 | 13,352 | sp\|P61009\|SPCS3_HUMAN | 25,123 | 13,465 | 54% | P61009 | SPCS3_HUMAN | SPCS3_HUMAN | Signal peptidase complex subunit 3 (Microsomal signal peptidase 22/23 kDa subunit) (SPC22/23) (SPase 22/23 kDa subunit) | SPCS3 SPC22 UNQ1841/PRO3567 |

| | | | | | | | | | | | | | |
|---|---|---|---|---|---|---|---|---|---|---|---|---|---|
| sp\|P02774\|VTDB_HUMAN | 42,745 | 22,644 | 16,035 | 38,686 | sp\|P02774\|VTDB_HUMAN | 245,907 | 275,392 | 112% | P02774 | VTDB_HUMAN | VTDB_HUMAN | Vitamin D-binding protein (DBP) (VDB) (Gc protein-derived macrophage activating factor) (Gc-MAF) (GcMAF) (Gc-globulin) (Group-specific component) (Gc) (Vitamin D-binding protein-macrophage activating factor) (DBP-maf) | GC |
| sp\|Q99988\|GDF15_HUMAN | 85,320 | 136,454 | 53,197 | 244,815 | sp\|Q99988\|GDF15_HUMAN | 140,749 | 57,906 | 41% | Q99988 | GDF15_HUMAN | GDF15_HUMAN | Growth/differentiation factor 15 (GDF-15) (Macrophage inhibitory cytokine 1) (MIC-1) (NSAID-activated gene 1 protein) (NAG-1) (NSAID-regulated gene 1 protein) (NRG-1) (Placental TGF-beta) (Placental bone morphogenetic protein) (Prostate differentiation factor) | GDF15 MIC1 PDF PLAB PTGFB |
| sp\|P16035\|TIMP2_HUMAN | 43,025 | 34,287 | 49,799 | 35,533 | sp\|P16035\|TIMP2_HUMAN | 42,953 | 20,911 | 49% | P16035 | TIMP2_HUMAN | TIMP2_HUMAN | Metalloproteinase inhibitor 2 (CSC-21K) (Tissue inhibitor of metalloproteinases 2) (TIMP-2) | TIMP2 |
| sp\|O43598\|DNPH1_HUMAN | 5,913 | 35,737 | 15,419 | 82,192 | sp\|O43598\|DNPH1_HUMAN | 45,248 | 30,166 | 67% | O43598 | DNPH1_HUMAN | DNPH1_HUMAN | 2'-deoxynucleoside 5'-phosphate N-hydrolase 1 (EC 3.2.2.-) (c-Myc-responsive protein RCL) | DNPH1 C6orf108 RCL |
| sp\|O75695\|XRP2_HUMAN | 119,017 | 155,834 | 81,593 | 76,925 | sp\|O75695\|XRP2_HUMAN | 92,550 | 34,970 | 38% | O75695 | XRP2_HUMAN | XRP2_HUMAN | Protein XRP2 | RP2 |
| sp\|P22307\|NLTP_HUMAN | 30,862 | 34,647 | 12,103 | 54,000 | sp\|P22307\|NLTP_HUMAN | 23,710 | 15,538 | 66% | P22307 | NLTP_HUMAN | SCP2_HUMAN | Sterol carrier protein 2 (SCP-2) (Acetyl-CoA C-myristoyltransferase) (EC 2.3.1.155) (Non-specific lipid-transfer protein) (NSL-TP) (Propanoyl-CoA C-acyltransferase) (EC 2.3.1.176) (SCP-2/3-oxoacyl-CoA thiolase) (SCP-2/thiolase) (EC 2.3.1.16) (SCP-chi) (SCPX) (Sterol carrier protein X) (SCP-X) | SCP2 |
| sp\|Q15041\|AR6P1_HUMAN | 23,915 | 19,886 | 28,140 | 25,691 | sp\|Q15041\|AR6P1_HUMAN | 16,050 | 9,794 | 61% | Q15041 | AR6P1_HUMAN | AR6P1_HUMAN | ADP-ribosylation factor-like protein 6-interacting protein 1 (ARL-6-interacting protein 1) (Aip-1) (Apoptotic regulator in the membrane of the endoplasmic reticulum) | ARL6IP1 ARL6IP ARMER KIAA0069 |
| sp\|P42766\|RL35_HUMAN | 221,420 | 263,548 | 206,355 | 204,766 | sp\|P42766\|RL35_HUMAN | 159,580 | 71,742 | 45% | P42766 | RL35_HUMAN | RL35_HUMAN | 60S ribosomal protein L35 (Large ribosomal subunit protein uL29) | RPL35 |
| sp\|P52272\|HNRPM_HUMAN | 78,473 | 68,970 | 108,526 | 54,099 | sp\|P52272\|HNRPM_HUMAN | 84,618 | 72,536 | 86% | P52272 | HNRPM_HUMAN | HNRPM_HUMAN | Heterogeneous nuclear ribonucleoprotein M (hnRNP M) | HNRNPM HNRPM NAGR1 |
| sp\|P15559\|NQO1_HUMAN | 11,895 | 35,926 | 11,358 | 23,776 | sp\|P15559\|NQO1_HUMAN | 19,353 | 9,117 | 47% | P15559 | NQO1_HUMAN | NQO1_HUMAN | NAD(P)H dehydrogenase [quinone] 1 (EC 1.6.5.2) (Azoreductase) (DT-diaphorase) (DTD) (Menadione reductase) (NAD(P)H:quinone oxidoreductase 1) (Phylloquinone reductase) (Quinone reductase 1) (QR1) | NQO1 DIA4 NMOR1 |
| sp\|Q15126\|PMVK_HUMAN | 49,509 | 48,179 | 53,828 | 10,911 | sp\|Q15126\|PMVK_HUMAN | 67,326 | 47,548 | 71% | Q15126 | PMVK_HUMAN | PMVK_HUMAN | Phosphomevalonate kinase (PMKase) (hPMK) (EC 2.7.4.2) | PMVK PMKI |

| | | | | | | | | | | | | | |
|---|---|---|---|---|---|---|---|---|---|---|---|---|---|
| sp\|P63208\|SKP1_HUMAN | 2,553 | 120,340 | 31,360 | 136,222 | sp\|P63208\|SKP1_HUMAN | 67,491 | 46,498 | 69% | P63208 | SKP1_HUMAN | SKP1_HUMAN | S-phase kinase-associated protein 1 (Cyclin-A/CDK2-associated protein p19) (p19A) (Organ of Corti protein 2) (OCP-2) (Organ of Corti protein II) (OCP-II) (RNA polymerase II elongation factor-like protein) (SIII) (Transcription elongation factor B polypeptide 1-like) (p19skp1) | SKP1 EMC19 OCP2 SKP1A TCEB1L |
| sp\|Q9HD42\|CHM1A_HUMAN | 121,894 | 175,577 | 93,149 | 67,335 | sp\|Q9HD42\|CHM1A_HUMAN | 106,263 | 35,940 | 34% | Q9HD42 | CHM1A_HUMAN | CHM1A_HUMAN | Charged multivesicular body protein 1a (Chromatin-modifying protein 1a) (CHMP1a) (Vacuolar protein sorting-associated protein 46-1) (Vps46-1) (hVps46-1) | CHMP1A CHMP1 KIAA0047 PCOLN3 PRSM1 |
| sp\|P24530\|EDNRB_HUMAN | 173,273 | 278,577 | 31,643 | 33,991 | sp\|P24530\|EDNRB_HUMAN | 150,750 | 130,760 | 87% | P24530 | EDNRB_HUMAN | EDNRB_HUMAN | Endothelin receptor type B (ET-B) (ET-BR) (Endothelin receptor non-selective type) | EDNRB ETRB |
| sp\|Q9BU23\|LMF2_HUMAN | 15,991 | 14,625 | 34,500 | 37,648 | sp\|Q9BU23\|LMF2_HUMAN | 29,331 | 17,660 | 60% | Q9BU23 | LMF2_HUMAN | LMF2_HUMAN | Lipase maturation factor 2 (Transmembrane protein 112B) (Transmembrane protein 153) | LMF2 TMEM112B TMEM153 |
| sp\|P20742\|PZP_HUMAN | 25,518 | 11,914 | 15,034 | 32,316 | sp\|P20742\|PZP_HUMAN | 817,786 | 914,721 | 112% | P20742 | PZP_HUMAN | PZP_HUMAN | Pregnancy zone protein (C3 and PZP-like alpha-2-macroglobulin domain-containing protein 6) | PZP CPAMD6 |
| sp\|P49593\|PPM1F_HUMAN | 51,622 | 51,690 | 73,130 | 86,085 | sp\|P49593\|PPM1F_HUMAN | 78,772 | 28,446 | 36% | P49593 | PPM1F_HUMAN | PPM1F_HUMAN | Protein phosphatase 1F (EC 3.1.3.16) (Ca(2+)/calmodulin-dependent protein kinase phosphatase) (CaM-kinase phosphatase) (CaMKPase) (Partner of PIX 2) (Protein fem-2 homolog) (hFem-2) | PPM1F KIAA0015 POPX2 |
| sp\|P61758\|PFD3_HUMAN | 9,684 | 58,535 | 11,978 | 13,707 | sp\|P61758\|PFD3_HUMAN | 26,883 | 20,990 | 78% | P61758 | PFD3_HUMAN | PFD3_HUMAN | Prefoldin subunit 3 (HIBBJ46) (von Hippel-Lindau-binding protein 1) (VBP-1) (VHL-binding protein 1) | VBP1 PFDN3 |
| sp\|O15075\|DCLK1_HUMAN | 96,255 | 129,755 | 27,119 | 120,414 | sp\|O15075\|DCLK1_HUMAN | 81,620 | 39,019 | 48% | O15075 | DCLK1_HUMAN | DCLK1_HUMAN | Serine/threonine-protein kinase DCLK1 (EC 2.7.11.1) (Doublecortin domain-containing protein 3A) (Doublecortin-like and CAM kinase-like 1) (Doublecortin-like kinase 1) | DCLK1 DCAMKL1 DCDC3A KIAA0369 |
| sp\|P04818\|TYSY_HUMAN | 251,749 | 50,289 | 48,341 | 84,446 | sp\|P04818\|TYSY_HUMAN | 110,806 | 80,866 | 73% | P04818 | TYSY_HUMAN | TYSY_HUMAN | Thymidylate synthase (TS) (TSase) (EC 2.1.1.45) | TYMS TS OK/SW-cl.29 |
| sp\|P00491\|PNPH_HUMAN | 15,082 | 23,922 | 6,501 | 25,183 | sp\|P00491\|PNPH_HUMAN | 26,102 | 15,758 | 60% | P00491 | PNPH_HUMAN | PNPH_HUMAN | Purine nucleoside phosphorylase (PNP) (EC 2.4.2.1) (Inosine phosphorylase) (Inosine-guanosine phosphorylase) | PNP NP |
| sp\|Q13838\|DX39B_HUMAN | 165,181 | 156,041 | 273,917 | 275,672 | sp\|Q13838\|DX39B_HUMAN | 149,017 | 90,636 | 61% | Q13838 | DX39B_HUMAN | DX39B_HUMAN | Spliceosome RNA helicase DDX39B (EC 3.6.4.13) (56 kDa U2AF65-associated protein) (ATP-dependent RNA helicase p47) (DEAD box protein UAP56) (HLA-B-associated transcript 1 protein) | DDX39B BAT1 UAP56 |
| sp\|Q76LX8\|ATS13_HUMAN | 14,200 | 5,298 | 260,571 | 242,737 | sp\|Q76LX8\|ATS13_HUMAN | 216,047 | 143,547 | 66% | Q76LX8 | ATS13_HUMAN | ATS13_HUMAN | A disintegrin and metalloproteinase with thrombospondin motifs 13 (ADAM-TS 13) (ADAM-TS13) (ADAMTS-13) (EC 3.4.24.87) (von Willebrand factor-cleaving protease) (vWF-CP) (vWF-cleaving protease) | ADAMTS13 C9orf8 UNQ6102/PRO20085 |
| sp\|Q9NY35\|CLDN1_HUMAN | 208,491 | 232,659 | 130,392 | 107,428 | sp\|Q9NY35\|CLDN1_HUMAN | 156,445 | 51,783 | 33% | Q9NY35 | CLDN1_HUMAN | CLDN1_HUMAN | Claudin domain-containing protein 1 (Membrane protein GENX-3745) | CLDND1 C3orf4 HSPC174 PSEC0054 UNQ2511/PRO6000 |
| sp\|Q92734\|TFG_HUMAN | 50,810 | 4,009 | 25,781 | 94,156 | sp\|Q92734\|TFG_HUMAN | 79,757 | 53,196 | 67% | Q92734 | TFG_HUMAN | TFG_HUMAN | Protein TFG (TRK-fused gene protein) | TFG |

| | | | | | | | | | | | | | |
|---|---|---|---|---|---|---|---|---|---|---|---|---|---|
| sp\|Q96EY1\|DNJA3_HUMAN | 8,306 | 22,120 | 32,021 | 30,813 | sp\|Q96EY1\|DNJA3_HUMAN | 35,658 | 23,632 | 66% | Q96EY1 | DNJA3_HUMAN | DNJA3_HUMAN | DnaJ homolog subfamily A member 3, mitochondrial (DnaJ protein Tid-1) (hTid-1) (Hepatocellular carcinoma-associated antigen 57) (Tumorous imaginal discs protein Tid56 homolog) | DNAJA3 HCA57 TID1 |
| sp\|Q9UG63\|ABCF2_HUMAN | 41,324 | 61,822 | 20,727 | 45,515 | sp\|Q9UG63\|ABCF2_HUMAN | 36,601 | 13,357 | 36% | Q9UG63 | ABCF2_HUMAN | ABCF2_HUMAN | ATP-binding cassette sub-family F member 2 (Iron-inhibited ABC transporter 2) | ABCF2 HUSSY-18 |
| sp\|Q969X1\|LFG3_HUMAN | 537,019 | 555,779 | 780,238 | 809,887 | sp\|Q969X1\|LFG3_HUMAN | 604,835 | 138,260 | 23% | Q969X1 | LFG3_HUMAN | LFG3_HUMAN | Protein lifeguard 3 (Protein RECS1 homolog) (Transmembrane BAX inhibitor motif-containing protein 1) | TMBIM1 LFG3 RECS1 PP1201 PSEC0158 |
| sp\|P12236\|ADT3_HUMAN | 104,737 | 142,752 | 47,429 | 21,942 | sp\|P12236\|ADT3_HUMAN | 54,426 | 47,628 | 88% | P12236 | ADT3_HUMAN | ADT3_HUMAN | ADP/ATP translocase 3 (ADP,ATP carrier protein 3) (ADP,ATP carrier protein, isoform T2) (ANT 2) (Adenine nucleotide translocator 3) (ANT 3) (Solute carrier family 25 member 6) [Cleaved into: ADP/ATP translocase 3, N-terminally processed] | SLC25A6 AAC3 ANT3 CDABP0051 |
| sp\|Q96QR8\|PURB_HUMAN | 89,901 | 56,315 | 80,936 | 18,214 | sp\|Q96QR8\|PURB_HUMAN | 55,438 | 23,206 | 42% | Q96QR8 | PURB_HUMAN | PURB_HUMAN | Transcriptional activator protein Pur-beta (Purine-rich element-binding protein B) | PURB |
| sp\|O75165\|DJC13_HUMAN | 34,525 | 53,743 | 57,549 | 86,432 | sp\|O75165\|DJC13_HUMAN | 45,040 | 24,418 | 54% | O75165 | DJC13_HUMAN | DJC13_HUMAN | DnaJ homolog subfamily C member 13 (Required for receptor-mediated endocytosis 8) (RME-8) | DNAJC13 KIAA0678 RME8 |
| sp\|Q9NQW7\|XPP1_HUMAN | 26,716 | 6,718 | 5,438 | 13,001 | sp\|Q9NQW7\|XPP1_HUMAN | 23,088 | 17,275 | 75% | Q9NQW7 | XPP1_HUMAN | XPP1_HUMAN | Xaa-Pro aminopeptidase 1 (EC 3.4.11.9) (Aminoacylproline aminopeptidase) (Cytosolic aminopeptidase P) (Soluble aminopeptidase P) (sAmp) (X-Pro aminopeptidase 1) (X-prolyl aminopeptidase 1, soluble) | XPNPEP1 XPNPEPL XPNPEPL1 |
| sp\|O76071\|CIAO1_HUMAN | 48,023 | 52,509 | 55,249 | 35,481 | sp\|O76071\|CIAO1_HUMAN | 47,417 | 13,176 | 28% | O76071 | CIAO1_HUMAN | CIAO1_HUMAN | Probable cytosolic iron-sulfur protein assembly protein CIAO1 (WD repeat-containing protein 39) | CIAO1 CIA1 WDR39 |
| sp\|Q13491\|GPM6B_HUMAN | 461,514 | 489,044 | 360,424 | 484,294 | sp\|Q13491\|GPM6B_HUMAN | 315,799 | 164,969 | 52% | Q13491 | GPM6B_HUMAN | GPM6B_HUMAN | Neuronal membrane glycoprotein M6-b (M6b) | GPM6B M6B |
| sp\|P23246\|SFPQ_HUMAN | 54,398 | 53,132 | 98,343 | 12,528 | sp\|P23246\|SFPQ_HUMAN | 36,314 | 31,334 | 86% | P23246 | SFPQ_HUMAN | SFPQ_HUMAN | Splicing factor, proline- and glutamine-rich (100 kDa DNA-pairing protein) (hPOMp100) (DNA-binding p52/p100 complex, 100 kDa subunit) (Polypyrimidine tract-binding protein-associated-splicing factor) (PSF) (PTB-associated-splicing factor) | SFPQ PSF |
| sp\|A0FGR8\|ESYT2_HUMAN | 42,710 | 23,991 | 64,196 | 52,049 | sp\|A0FGR8\|ESYT2_HUMAN | 42,360 | 14,878 | 35% | A0FGR8 | ESYT2_HUMAN | ESYT2_HUMAN | Extended synaptotagmin-2 (E-Syt2) (Chr2Syt) | ESYT2 FAM62B KIAA1228 |
| sp\|Q08722\|CD47_HUMAN | 603,851 | 644,286 | 190,951 | 273,478 | sp\|Q08722\|CD47_HUMAN | 332,610 | 189,949 | 57% | Q08722 | CD47_HUMAN | CD47_HUMAN | Leukocyte surface antigen CD47 (Antigenic surface determinant protein OA3) (Integrin-associated protein) (IAP) (Protein MER6) (CD antigen CD47) | CD47 MER6 |
| sp\|Q03154\|ACY1_HUMAN | 4,525 | 72,116 | 69,763 | 85,243 | sp\|Q03154\|ACY1_HUMAN | 47,575 | 28,933 | 61% | Q03154 | ACY1_HUMAN | ACY1_HUMAN | Aminoacylase-1 (ACY-1) (EC 3.5.1.14) (N-acyl-L-amino-acid amidohydrolase) | ACY1 |
| sp\|P78527\|PRKDC_HUMAN | 10,767 | 19,025 | 23,663 | 10,685 | sp\|P78527\|PRKDC_HUMAN | 18,033 | 11,451 | 64% | P78527 | PRKDC_HUMAN | PRKDC_HUMAN | DNA-dependent protein kinase catalytic subunit (DNA-PK catalytic subunit) (DNA-PKcs) (EC 2.7.11.1) (DNPK1) (p460) | PRKDC HYRC HYRC1 |
| sp\|O60749\|SNX2_HUMAN | 6,042 | 19,274 | 30,009 | 30,172 | sp\|O60749\|SNX2_HUMAN | 18,494 | 9,852 | 53% | O60749 | SNX2_HUMAN | SNX2_HUMAN | Sorting nexin-2 (Transformation-related gene 9 protein) (TRG-9) | SNX2 TRG9 |

| | | | | | | | | | | | | | |
|---|---|---|---|---|---|---|---|---|---|---|---|---|---|
| sp\|Q9NP79\|VTA1_HUMAN | 107,914 | 167,052 | 93,372 | 102,844 | sp\|Q9NP79\|VTA1_HUMAN | 110,981 | 26,167 | 24% | Q9NP79 | VTA1_HUMAN | VTA1_HUMAN | Vacuolar protein sorting-associated protein VTA1 homolog (Dopamine-responsive gene 1 protein) (DRG-1) (LYST-interacting protein 5) (LIP5) (SKD1-binding protein 1) (SBP1) | VTA1 C6orf55 HSPC228 My012 |
| sp\|O43795\|MYO1B_HUMAN | 220,042 | 342,193 | 246,753 | 405,566 | sp\|O43795\|MYO1B_HUMAN | 283,195 | 64,981 | 23% | O43795 | MYO1B_HUMAN | MYO1B_HUMAN | Unconventional myosin-Ib (MYH-1c) (Myosin I alpha) (MMI-alpha) (MMIa) | MYO1B |
| sp\|Q14624\|ITIH4_HUMAN | 82,716 | 17,396 | 102,884 | 58,611 | sp\|Q14624\|ITIH4_HUMAN | 165,940 | 131,857 | 79% | Q14624 | ITIH4_HUMAN | ITIH4_HUMAN | Inter-alpha-trypsin inhibitor heavy chain H4 (ITI heavy chain H4) (ITI-HC4) (Inter-alpha-inhibitor heavy chain 4) (Inter-alpha-trypsin inhibitor family heavy chain-related protein) (IHRP) (Plasma kallikrein sensitive glycoprotein 120) (Gp120) (PK-120) [Cleaved into: 70 kDa inter-alpha-trypsin inhibitor heavy chain H4; 35 kDa inter-alpha-trypsin inhibitor heavy chain H4] | ITIH4 IHRP ITIHL1 PK120 PRO1851 |
| sp\|P82930\|RT34_HUMAN | 15,444 | 17,963 | 3,715 | 9,416 | sp\|P82930\|RT34_HUMAN | 10,093 | 4,967 | 49% | P82930 | RT34_HUMAN | RT34_HUMAN | 28S ribosomal protein S34, mitochondrial (MRP-S34) (S34mt) (Mitochondrial small ribosomal subunit protein mS34) | MRPS34 |
| sp\|P56937\|DHB7_HUMAN | 16,458 | 17,811 | 5,523 | 25,860 | sp\|P56937\|DHB7_HUMAN | 13,843 | 7,309 | 53% | P56937 | DHB7_HUMAN | DHB7_HUMAN | 3-keto-steroid reductase/17-beta-hydroxysteroid dehydrogenase 7 (17-beta-hydroxysteroid dehydrogenase 7) (17-beta-HSD 7) (3-keto-steroid reductase) (EC 1.1.1.270) (Dihydrotestosterone oxidoreductase) (EC 1.1.1.210) (Estradiol 17-beta-dehydrogenase 7) (EC 1.1.1.62) (Short chain dehydrogenase/reductase family 37C member 1) | HSD17B7 17HSD7 SDR37C1 UNQ2563/PRO6243 |
| sp\|O43505\|B4GA1_HUMAN | 50,616 | 20,368 | 88,327 | 47,346 | sp\|O43505\|B4GA1_HUMAN | 186,298 | 168,737 | 91% | O43505 | B4GA1_HUMAN | B4GA1_HUMAN | Beta-1,4-glucuronyltransferase 1 (EC 2.4.1.-) (I-beta-1,3-N-acetylglucosaminyltransferase) (iGnT) (N-acetyllactosaminide beta-1,3-N-acetylglucosaminyltransferase) (Poly-N-acetyllactosamine extension enzyme) (UDP-GlcNAc:betaGal beta-1,3-N-acetylglucosaminyltransferase 1) | B4GAT1 B3GNT1 B3GNT6 |
| sp\|Q8NFJ5\|RAI3_HUMAN | 151,696 | 167,662 | 252,023 | 232,067 | sp\|Q8NFJ5\|RAI3_HUMAN | 220,531 | 63,150 | 29% | Q8NFJ5 | RAI3_HUMAN | RAI3_HUMAN | Retinoic acid-induced protein 3 (G-protein coupled receptor family C group 5 member A) (Phorbol ester induced gene 1) (PEIG-1) (Retinoic acid-induced gene 1 protein) (RAIG-1) | GPRC5A GPCR5A RAI3 RAIG1 |
| sp\|Q14192\|FHL2_HUMAN | 19,987 | 62,304 | 138,124 | 32,506 | sp\|Q14192\|FHL2_HUMAN | 59,417 | 42,749 | 72% | Q14192 | FHL2_HUMAN | FHL2_HUMAN | Four and a half LIM domains protein 2 (FHL-2) (LIM domain protein DRAL) (Skeletal muscle LIM-protein 3) (SLIM-3) | FHL2 DRAL SLIM3 |
| sp\|Q969V3\|NCLN_HUMAN | 14,545 | 19,309 | 69,560 | 87,913 | sp\|Q969V3\|NCLN_HUMAN | 70,749 | 37,070 | 52% | Q969V3 | NCLN_HUMAN | NCLN_HUMAN | Nicalin (Nicastrin-like protein) | NCLN |

| | | | | | | | | | | | | | |
|---|---|---|---|---|---|---|---|---|---|---|---|---|---|
| sp\|P15509\|CSF2R_HUMAN | 106,526 | 162,049 | 46,373 | 121,105 | sp\|P15509\|CSF2R_HUMAN | 104,654 | 35,754 | 34% | P15509 | CSF2R_HUMAN | CSF2R_HUMAN | Granulocyte-macrophage colony-stimulating factor receptor subunit alpha (GM-CSF-R-alpha) (GMCSFR-alpha) (GMR-alpha) (CDw116) (CD antigen CD116) | CSF2RA CSF2R CSF2RY |
| sp\|Q00688\|FKBP3_HUMAN | 15,691 | 31,392 | 17,629 | 16,551 | sp\|Q00688\|FKBP3_HUMAN | 16,871 | 9,780 | 58% | Q00688 | FKBP3_HUMAN | FKBP3_HUMAN | Peptidyl-prolyl cis-trans isomerase FKBP3 (PPIase FKBP3) (EC 5.2.1.8) (25 kDa FK506-binding protein) (25 kDa FKBP) (FKBP-25) (FK506-binding protein 3) (FKBP-3) (Immunophilin FKBP25) (Rapamycin-selective 25 kDa immunophilin) (Rotamase) | FKBP3 FKBP25 |
| sp\|P07358\|CO8B_HUMAN | 9,627 | 7,248 | 9,019 | 3,245 | sp\|P07358\|CO8B_HUMAN | 116,272 | 130,943 | 113% | P07358 | CO8B_HUMAN | CO8B_HUMAN | Complement component C8 beta chain (Complement component 8 subunit beta) | C8B |
| sp\|P48556\|PSMD8_HUMAN | 114,743 | 103,641 | 78,635 | 74,255 | sp\|P48556\|PSMD8_HUMAN | 65,721 | 34,220 | 52% | P48556 | PSMD8_HUMAN | PSMD8_HUMAN | 26S proteasome non-ATPase regulatory subunit 8 (26S proteasome regulatory subunit RPN12) (26S proteasome regulatory subunit S14) (p31) | PSMD8 |
| sp\|Q9NTJ5\|SAC1_HUMAN | 54,101 | 60,826 | 21,889 | 61,731 | sp\|Q9NTJ5\|SAC1_HUMAN | 61,136 | 29,387 | 48% | Q9NTJ5 | SAC1_HUMAN | SAC1_HUMAN | Phosphatidylinositol-3-phosphatase SAC1 (EC 3.1.3.64) (Phosphatidylinositol-4-phosphate phosphatase) (Suppressor of actin mutations 1-like protein) | SACM1L KIAA0851 SAC1 |
| sp\|Q16851\|UGPA_HUMAN | 61,549 | 75,158 | 73,682 | 74,260 | sp\|Q16851\|UGPA_HUMAN | 61,531 | 29,441 | 48% | Q16851 | UGPA_HUMAN | UGPA_HUMAN | UTP--glucose-1-phosphate uridylyltransferase (EC 2.7.7.9) (UDP-glucose pyrophosphorylase) (UDPGP) (UGPase) | UGP2 UGP1 |
| sp\|O60831\|PRAF2_HUMAN | 4,171 | 6,029 | 20,748 | 18,223 | sp\|O60831\|PRAF2_HUMAN | 9,792 | 7,309 | 75% | O60831 | PRAF2_HUMAN | PRAF2_HUMAN | PRA1 family protein 2 | PRAF2 JM4 |
| sp\|Q02241\|KIF23_HUMAN | 136,879 | 168,659 | 161,016 | 178,567 | sp\|Q02241\|KIF23_HUMAN | 138,243 | 31,435 | 23% | Q02241 | KIF23_HUMAN | KIF23_HUMAN | Kinesin-like protein KIF23 (Kinesin-like protein 5) (Mitotic kinesin-like protein 1) | KIF23 KNSL5 MKLP1 |
| sp\|Q9Y3I1\|FBX7_HUMAN | 66,525 | 82,215 | 57,344 | 18,711 | sp\|Q9Y3I1\|FBX7_HUMAN | 69,425 | 37,588 | 54% | Q9Y3I1 | FBX7_HUMAN | FBX7_HUMAN | F-box only protein 7 | FBXO7 FBX7 |
| sp\|Q9Y3L5\|RAP2C_HUMAN | 67,016 | 117,594 | 46,086 | 100,691 | sp\|Q9Y3L5\|RAP2C_HUMAN | 85,181 | 27,334 | 32% | Q9Y3L5 | RAP2C_HUMAN | RAP2C_HUMAN | Ras-related protein Rap-2c (EC 3.6.5.2) | RAP2C |
| sp\|Q96GW7\|PGCB_HUMAN | 344,649 | 378,174 | 595,361 | 743,267 | sp\|Q96GW7\|PGCB_HUMAN | 429,147 | 187,299 | 44% | Q96GW7 | PGCB_HUMAN | PGCB_HUMAN | Brevican core protein (Brain-enriched hyaluronan-binding protein) (BEHAB) (Chondroitin sulfate proteoglycan 7) | BCAN BEHAB CSPG7 UNQ2525/PRO6018 |
| sp\|Q9BXJ4\|C1QT3_HUMAN | 10,211 | 8,393 | 30,336 | 28,308 | sp\|Q9BXJ4\|C1QT3_HUMAN | 150,805 | 162,882 | 108% | Q9BXJ4 | C1QT3_HUMAN | C1QT3_HUMAN | Complement C1q tumor necrosis factor-related protein 3 (Collagenous repeat-containing sequence 26 kDa protein) (CORS26) (Secretory protein CORS26) | C1QTNF3 CTRP3 UNQ753/PRO1484 |
| sp\|Q99653\|CHP1_HUMAN | 110,423 | 42,480 | 123,421 | 135,376 | sp\|Q99653\|CHP1_HUMAN | 108,422 | 36,880 | 34% | Q99653 | CHP1_HUMAN | CHP1_HUMAN | Calcineurin B homologous protein 1 (Calcineurin B-like protein) (Calcium-binding protein CHP) (Calcium-binding protein p22) (EF-hand calcium-binding domain-containing protein p22) | CHP1 CHP |
| sp\|P02787\|TRFE_HUMAN | 5,206 | 9,523 | 5,396 | 12,570 | sp\|P02787\|TRFE_HUMAN | 69,415 | 69,357 | 100% | P02787 | TRFE_HUMAN | TRFE_HUMAN | Serotransferrin (Transferrin) (Beta-1 metal-binding globulin) (Siderophilin) | TF PRO1400 |

| | | | | | | | | | | | | | |
|---|---|---|---|---|---|---|---|---|---|---|---|---|---|
| sp\|P04062\|GLCM_HUMAN | 148,400 | 16,875 | 99,988 | 109,602 | sp\|P04062\|GLCM_HUMAN | 71,307 | 48,928 | 69% | P04062 | GLCM_HUMAN | GLCM_HUMAN | Lysosomal acid glucosylceramidase (Lysosomal acid GCase) (EC 3.2.1.45) (Acid beta-glucosidase) (Alglucerase) (Beta-glucocerebrosidase) (Beta-GC) (Cholesterol glucosyltransferase) (SGTase) (EC 2.4.1.-) (Cholesteryl-beta-glucosidase) (EC 3.2.1.-) (D-glucosyl-N-acylsphingosine glucohydrolase) (Imiglucerase) (Lysosomal cholesterol glycosyltransferase) (Lysosomal galactosylceramidase) (EC 3.2.1.46) (Lysosomal glycosylceramidase) | GBA GC GLUC |
| sp\|O76003\|GLRX3_HUMAN | 63,961 | 74,615 | 8,303 | 33,250 | sp\|O76003\|GLRX3_HUMAN | 36,102 | 24,206 | 67% | O76003 | GLRX3_HUMAN | GLRX3_HUMAN | Glutaredoxin-3 (PKC-interacting cousin of thioredoxin) (PICOT) (PKC-theta-interacting protein) (PKCq-interacting protein) (Thioredoxin-like protein 2) | GLRX3 PICOT TXNL2 HUSSY-22 |
| sp\|P51511\|MMP15_HUMAN | 168,149 | 158,195 | 202,881 | 175,742 | sp\|P51511\|MMP15_HUMAN | 191,591 | 32,927 | 17% | P51511 | MMP15_HUMAN | MMP15_HUMAN | Matrix metalloproteinase-15 (MMP-15) (EC 3.4.24.-) (Membrane-type matrix metalloproteinase 2) (MT-MMP 2) (MTMMP2) (Membrane-type-2 matrix metalloproteinase) (MT2-MMP) (MT2MMP) (SMCP-2) | MMP15 |
| sp\|Q96HC4\|PDLI5_HUMAN | 42,301 | 54,658 | 24,382 | 28,633 | sp\|Q96HC4\|PDLI5_HUMAN | 36,708 | 14,506 | 40% | Q96HC4 | PDLI5_HUMAN | PDLI5_HUMAN | PDZ and LIM domain protein 5 (Enigma homolog) (Enigma-like PDZ and LIM domains protein) | PDLIM5 ENH L9 |
| sp\|P02748\|CO9_HUMAN | 56,551 | 25,484 | 73,317 | 35,885 | sp\|P02748\|CO9_HUMAN | 697,306 | 792,359 | 114% | P02748 | CO9_HUMAN | CO9_HUMAN | Complement component C9 [Cleaved into: Complement component C9a; Complement component C9b] | C9 |
| sp\|P50552\|VASP_HUMAN | 24,656 | 19,465 | 52,357 | 33,605 | sp\|P50552\|VASP_HUMAN | 20,447 | 16,239 | 79% | P50552 | VASP_HUMAN | VASP_HUMAN | Vasodilator-stimulated phosphoprotein (VASP) | VASP |
| sp\|P52597\|HNRPF_HUMAN | 96,262 | 86,505 | 96,489 | 39,468 | sp\|P52597\|HNRPF_HUMAN | 122,329 | 111,245 | 91% | P52597 | HNRPF_HUMAN | HNRPF_HUMAN | Heterogeneous nuclear ribonucleoprotein F (hnRNP F) (Nucleolin-like protein mcs94-1) [Cleaved into: Heterogeneous nuclear ribonucleoprotein F, N-terminally processed] | HNRNPF HNRPF |
| sp\|Q92743\|HTRA1_HUMAN | 91,868 | 57,934 | 56,285 | 42,897 | sp\|Q92743\|HTRA1_HUMAN | 70,473 | 20,010 | 28% | Q92743 | HTRA1_HUMAN | HTRA1_HUMAN | Serine protease HTRA1 (EC 3.4.21.-) (High-temperature requirement A serine peptidase 1) (L56) (Serine protease 11) | HTRA1 HTRA PRSS11 |
| sp\|P20020\|AT2B1_HUMAN | 61,856 | 59,516 | 34,885 | 47,629 | sp\|P20020\|AT2B1_HUMAN | 46,436 | 21,319 | 46% | P20020 | AT2B1_HUMAN | AT2B1_HUMAN | Plasma membrane calcium-transporting ATPase 1 (EC 7.2.2.10) (Plasma membrane calcium ATPase isoform 1) (PMCA1) (Plasma membrane calcium pump isoform 1) | ATP2B1 PMCA1 |
| sp\|O15431\|COPT1_HUMAN | 37,135 | 60,062 | 22,902 | 17,958 | sp\|O15431\|COPT1_HUMAN | 35,839 | 13,762 | 38% | O15431 | COPT1_HUMAN | COPT1_HUMAN | High affinity copper uptake protein 1 (Copper transporter 1) (hCTR1) (Solute carrier family 31 member 1) | SLC31A1 COPT1 CTR1 |

| | | | | | | | | | | | | | |
|---|---|---|---|---|---|---|---|---|---|---|---|---|---|
| sp\|P00751\|CFAB_HUMAN | 26,938 | 17,937 | 33,342 | 27,164 | sp\|P00751\|CFAB_HUMAN | 305,991 | 328,130 | 107% | P00751 | CFAB_HUMAN | CFAB_HUMAN | Complement factor B (EC 3.4.21.47) (C3/C5 convertase) (Glycine-rich beta glycoprotein) (GBG) (PBF2) (Properdin factor B) [Cleaved into: Complement factor B Ba fragment; Complement factor B Bb fragment] | CFB BF BFD |
| sp\|Q15063\|POSTN_HUMAN | 13,558 | 11,156 | 53,453 | 10,555 | sp\|Q15063\|POSTN_HUMAN | 49,088 | 43,602 | 89% | Q15063 | POSTN_HUMAN | POSTN_HUMAN | Periostin (PN) (Osteoblast-specific factor 2) (OSF-2) | POSTN OSF2 |
| sp\|Q15417\|CNN3_HUMAN | 9,290 | 32,979 | 3,469 | 8,883 | sp\|Q15417\|CNN3_HUMAN | 18,921 | 12,122 | 64% | Q15417 | CNN3_HUMAN | CNN3_HUMAN | Calponin-3 (Calponin, acidic isoform) | CNN3 |
| sp\|P36543\|VATE1_HUMAN | 77,010 | 100,965 | 3,418 | 90,198 | sp\|P36543\|VATE1_HUMAN | 58,355 | 36,910 | 63% | P36543 | VATE1_HUMAN | VATE1_HUMAN | V-type proton ATPase subunit E 1 (V-ATPase subunit E 1) (V-ATPase 31 kDa subunit) (p31) (Vacuolar proton pump subunit E 1) | ATP6V1E1 ATP6E ATP6E2 |
| sp\|P05362\|ICAM1_HUMAN | 184,615 | 230,668 | 323,498 | 355,701 | sp\|P05362\|ICAM1_HUMAN | 263,481 | 77,927 | 30% | P05362 | ICAM1_HUMAN | ICAM1_HUMAN | Intercellular adhesion molecule 1 (ICAM-1) (Major group rhinovirus receptor) (CD antigen CD54) | ICAM1 |
| sp\|Q4KMQ2\|ANO6_HUMAN | 76,801 | 90,539 | 133,722 | 224,231 | sp\|Q4KMQ2\|ANO6_HUMAN | 140,100 | 64,507 | 46% | Q4KMQ2 | ANO6_HUMAN | ANO6_HUMAN | Anoctamin-6 (Small-conductance calcium-activated nonselective cation channel) (SCAN channel) (Transmembrane protein 16F) | ANO6 TMEM16F |
| sp\|Q9UJU6\|DBNL_HUMAN | 21,257 | 15,872 | 46,571 | 45,752 | sp\|Q9UJU6\|DBNL_HUMAN | 34,082 | 12,458 | 37% | Q9UJU6 | DBNL_HUMAN | DBNL_HUMAN | Drebrin-like protein (Cervical SH3P7) (Cervical mucin-associated protein) (Drebrin-F) (HPK1-interacting protein of 55 kDa) (HIP-55) (SH3 domain-containing protein 7) | DBNL CMAP SH3P7 PP5423 |
| sp\|O43175\|SERA_HUMAN | 42,650 | 57,633 | 53,492 | 76,038 | sp\|O43175\|SERA_HUMAN | 53,332 | 12,104 | 23% | O43175 | SERA_HUMAN | SERA_HUMAN | D-3-phosphoglycerate dehydrogenase (3-PGDH) (EC 1.1.1.95) (2-oxoglutarate reductase) (EC 1.1.1.399) (Malate dehydrogenase) (EC 1.1.1.37) | PHGDH PGDH3 |
| sp\|P08034\|CXB1_HUMAN | 476 | 54,766 | 15,964 | 35,369 | sp\|P08034\|CXB1_HUMAN | 22,120 | 18,258 | 83% | P08034 | CXB1_HUMAN | CXB1_HUMAN | Gap junction beta-1 protein (Connexin-32) (Cx32) (GAP junction 28 kDa liver protein) | GJB1 CX32 |
| sp\|Q9Y5J5\|PHLA3_HUMAN | 89,537 | 95,071 | 43,011 | 70,501 | sp\|Q9Y5J5\|PHLA3_HUMAN | 63,710 | 21,403 | 34% | Q9Y5J5 | PHLA3_HUMAN | PHLA3_HUMAN | Pleckstrin homology-like domain family A member 3 (TDAG51/Ipl homolog 1) | PHLDA3 TIH1 |
| sp\|P25685\|DNJB1_HUMAN | 16,023 | 39,733 | 6,606 | 16,512 | sp\|P25685\|DNJB1_HUMAN | 17,587 | 12,417 | 71% | P25685 | DNJB1_HUMAN | DNJB1_HUMAN | DnaJ homolog subfamily B member 1 (DnaJ protein homolog 1) (Heat shock 40 kDa protein 1) (HSP40) (Heat shock protein 40) (Human DnaJ protein 1) (hDj-1) | DNAJB1 DNAJ1 HDJ1 HSPF1 |
| sp\|Q9UL26\|RB22A_HUMAN | 129,533 | 146,394 | 80,743 | 60,026 | sp\|Q9UL26\|RB22A_HUMAN | 101,679 | 33,306 | 33% | Q9UL26 | RB22A_HUMAN | RB22A_HUMAN | Ras-related protein Rab-22A (Rab-22) | RAB22A RAB22 |
| sp\|Q9Y5Y2\|NUBP2_HUMAN | 17,959 | 31,825 | 38,037 | 10,849 | sp\|Q9Y5Y2\|NUBP2_HUMAN | 22,168 | 13,279 | 60% | Q9Y5Y2 | NUBP2_HUMAN | NUBP2_HUMAN | Cytosolic Fe-S cluster assembly factor NUBP2 (Nucleotide-binding protein 2) (NBP 2) | NUBP2 |
| sp\|P30626\|SORCN_HUMAN | 102,737 | 6,044 | 125,243 | 121,161 | sp\|P30626\|SORCN_HUMAN | 70,289 | 47,304 | 67% | P30626 | SORCN_HUMAN | SORCN_HUMAN | Sorcin (22 kDa protein) (CP-22) (CP22) (V19) | SRI |
| sp\|Q92692\|NECT2_HUMAN | 41,415 | 37,512 | 87,816 | 75,355 | sp\|Q92692\|NECT2_HUMAN | 64,908 | 26,208 | 40% | Q92692 | NECT2_HUMAN | NECT2_HUMAN | Nectin-2 (Herpes virus entry mediator B) (Herpesvirus entry mediator B) (HveB) (Nectin cell adhesion molecule 2) (Poliovirus receptor-related protein 2) (CD antigen CD112) | NECTIN2 HVEB PRR2 PVRL2 |
| sp\|P29317\|EPHA2_HUMAN | 97,325 | 134,006 | 37,142 | 48,710 | sp\|P29317\|EPHA2_HUMAN | 71,318 | 37,443 | 53% | P29317 | EPHA2_HUMAN | EPHA2_HUMAN | Ephrin type-A receptor 2 (EC 2.7.10.1) (Epithelial cell kinase) (Tyrosine-protein kinase receptor ECK) | EPHA2 ECK |

| | | | | | | | | | | | | | |
|---|---|---|---|---|---|---|---|---|---|---|---|---|---|
| sp\|Q15393\|SF3B3_HUMAN | 91,547 | 66,258 | 161,248 | 145,920 | sp\|Q15393\|SF3B3_HUMAN | 86,927 | 46,315 | 53% | Q15393 | SF3B3_HUMAN | SF3B3_HUMAN | Splicing factor 3B subunit 3 (Pre-mRNA-splicing factor SF3b 130 kDa subunit) (SF3b130) (STAF130) (Spliceosome-associated protein 130) (SAP 130) | SF3B3 KIAA0017 SAP130 |
| sp\|O94887\|FARP2_HUMAN | 71,651 | 91,477 | 8,723 | 7,551 | sp\|O94887\|FARP2_HUMAN | 56,312 | 31,549 | 56% | O94887 | FARP2_HUMAN | FARP2_HUMAN | FERM, ARHGEF and pleckstrin domain-containing protein 2 (FERM domain-including RhoGEF) (FIR) (FERM, RhoGEF and pleckstrin domain-containing protein 2) (Pleckstrin homology domain-containing family C member 3) (PH domain-containing family C member 3) | FARP2 KIAA0793 PLEKHC3 |
| sp\|Q13561\|DCTN2_HUMAN | 51,047 | 56,983 | 284,809 | 23,208 | sp\|Q13561\|DCTN2_HUMAN | 122,260 | 100,050 | 82% | Q13561 | DCTN2_HUMAN | DCTN2_HUMAN | Dynactin subunit 2 (50 kDa dynein-associated polypeptide) (Dynactin complex 50 kDa subunit) (DCTN-50) (p50 dynamitin) | DCTN2 DCTN50 |
| sp\|Q9Y5X3\|SNX5_HUMAN | 41,340 | 44,239 | 17,313 | 4,867 | sp\|Q9Y5X3\|SNX5_HUMAN | 27,899 | 15,057 | 54% | Q9Y5X3 | SNX5_HUMAN | SNX5_HUMAN | Sorting nexin-5 | SNX5 |
| sp\|P35443\|TSP4_HUMAN | 13,608 | 13,600 | 26,040 | 41,933 | sp\|P35443\|TSP4_HUMAN | 94,385 | 90,917 | 96% | P35443 | TSP4_HUMAN | TSP4_HUMAN | Thrombospondin-4 | THBS4 TSP4 |
| sp\|Q15404\|RSU1_HUMAN | 71,894 | 130,704 | 12,869 | 66,795 | sp\|Q15404\|RSU1_HUMAN | 114,164 | 57,447 | 50% | Q15404 | RSU1_HUMAN | RSU1_HUMAN | Ras suppressor protein 1 (RSP-1) (Rsu-1) | RSU1 RSP1 |
| sp\|P84095\|RHOG_HUMAN | 73,110 | 82,223 | 84,620 | 76,378 | sp\|P84095\|RHOG_HUMAN | 73,184 | 10,055 | 14% | P84095 | RHOG_HUMAN | RHOG_HUMAN | Rho-related GTP-binding protein RhoG | RHOG ARHG |
| sp\|Q13724\|MOGS_HUMAN | 3,663 | 19,210 | 16,996 | 3,037 | sp\|Q13724\|MOGS_HUMAN | 12,916 | 11,615 | 90% | Q13724 | MOGS_HUMAN | MOGS_HUMAN | Mannosyl-oligosaccharide glucosidase (EC 3.2.1.106) (Processing A-glucosidase I) | MOGS GCS1 |
| sp\|O75044\|SRGP2_HUMAN | 33,488 | 33,297 | 47,755 | 63,165 | sp\|O75044\|SRGP2_HUMAN | 42,503 | 17,152 | 40% | O75044 | SRGP2_HUMAN | SRGP2_HUMAN | SLIT-ROBO Rho GTPase-activating protein 2 (srGAP2) (Formin-binding protein 2) (Rho GTPase-activating protein 34) | SRGAP2 ARHGAP34 FNBP2 KIAA0456 SRGAP2A |
| sp\|P60983\|GMFB_HUMAN | 69,906 | 82,444 | 19,919 | 116,118 | sp\|P60983\|GMFB_HUMAN | 70,380 | 32,867 | 47% | P60983 | GMFB_HUMAN | GMFB_HUMAN | Glia maturation factor beta (GMF-beta) | GMFB |
| sp\|O95747\|OXSR1_HUMAN | 57,588 | 66,346 | 8,685 | 49,217 | sp\|O95747\|OXSR1_HUMAN | 34,164 | 24,472 | 72% | O95747 | OXSR1_HUMAN | OXSR1_HUMAN | Serine/threonine-protein kinase OSR1 (EC 2.7.11.1) (Oxidative stress-responsive 1 protein) | OXSR1 KIAA1101 OSR1 |
| sp\|O43813\|LANC1_HUMAN | 51,458 | 64,303 | 44,863 | 46,507 | sp\|O43813\|LANC1_HUMAN | 58,436 | 29,904 | 51% | O43813 | LANC1_HUMAN | LANC1_HUMAN | Glutathione S-transferase LANCL1 (EC 2.5.1.18) (40 kDa erythrocyte membrane protein) (p40) (LanC-like protein 1) | LANCL1 GPR69A |
| sp\|Q13144\|EI2BE_HUMAN | 12,756 | 17,730 | 1,902 | 15,492 | sp\|Q13144\|EI2BE_HUMAN | 11,043 | 8,356 | 76% | Q13144 | EI2BE_HUMAN | EI2BE_HUMAN | Translation initiation factor eIF-2B subunit epsilon (eIF-2B GDP-GTP exchange factor subunit epsilon) | EIF2B5 EIF2BE |
| sp\|P51570\|GALK1_HUMAN | 45,030 | 34,347 | 7,405 | 10,190 | sp\|P51570\|GALK1_HUMAN | 43,858 | 25,540 | 58% | P51570 | GALK1_HUMAN | GALK1_HUMAN | Galactokinase (EC 2.7.1.6) (Galactose kinase) | GALK1 GALK |
| sp\|P15144\|AMPN_HUMAN | 155,155 | 111,366 | 69,449 | 135,902 | sp\|P15144\|AMPN_HUMAN | 85,233 | 51,428 | 60% | P15144 | AMPN_HUMAN | AMPN_HUMAN | Aminopeptidase N (AP-N) (hAPN) (EC 3.4.11.2) (Alanyl aminopeptidase) (Aminopeptidase M) (AP-M) (Microsomal aminopeptidase) (Myeloid plasma membrane glycoprotein CD13) (gp150) (CD antigen CD13) | ANPEP APN CD13 PEPN |
| sp\|Q9NX76\|CKLF6_HUMAN | 47,716 | 66,931 | 70,901 | 337,687 | sp\|Q9NX76\|CKLF6_HUMAN | 180,854 | 118,087 | 65% | Q9NX76 | CKLF6_HUMAN | CKLF6_HUMAN | CKLF-like MARVEL transmembrane domain-containing protein 6 (Chemokine-like factor superfamily member 6) | CMTM6 CKLFSF6 |

| | | | | | | | | | | | | | |
|---|---|---|---|---|---|---|---|---|---|---|---|---|---|
| sp\|O95757\|HS74L_HUMAN | 43,096 | 49,848 | 18,194 | 17,266 | sp\|O95757\|HS74L_HUMAN | 86,903 | 78,614 | 90% | O95757 | HS74L_HUMAN | HS74L_HUMAN | Heat shock 70 kDa protein 4L (Heat shock 70-related protein APG-1) (Heat-shock protein family A member 4-like protein) (HSPA4-like protein) (Osmotic stress protein 94) | HSPA4L APG1 OSP94 |
| sp\|Q00577\|PURA_HUMAN | 39,712 | 58,875 | 16,488 | 80,744 | sp\|Q00577\|PURA_HUMAN | 51,403 | 27,657 | 54% | Q00577 | PURA_HUMAN | PURA_HUMAN | Transcriptional activator protein Pur-alpha (Purine-rich single-stranded DNA-binding protein alpha) | PURA PUR1 |
| sp\|Q15382\|RHEB_HUMAN | 94,834 | 110,243 | 100,390 | 93,392 | sp\|Q15382\|RHEB_HUMAN | 88,538 | 14,920 | 17% | Q15382 | RHEB_HUMAN | RHEB_HUMAN | GTP-binding protein Rheb (Ras homolog enriched in brain) | RHEB RHEB2 |
| sp\|Q9NUP9\|LIN7C_HUMAN | 73,160 | 75,495 | 168,412 | 192,027 | sp\|Q9NUP9\|LIN7C_HUMAN | 127,899 | 52,661 | 41% | Q9NUP9 | LIN7C_HUMAN | LIN7C_HUMAN | Protein lin-7 homolog C (Lin-7C) (Mammalian lin-seven protein 3) (MALS-3) (Vertebrate lin-7 homolog 3) (Veli-3) | LIN7C MALS3 VELI3 |
| sp\|P20774\|MIME_HUMAN | 6,812 | 2,512 | 6,630 | 7,471 | sp\|P20774\|MIME_HUMAN | 42,485 | 50,563 | 119% | P20774 | MIME_HUMAN | MIME_HUMAN | Mimecan (Osteoglycin) (Osteoinductive factor) (OIF) | OGN OIF SLRR3A |
| sp\|O14744\|ANM5_HUMAN | 92,113 | 65,051 | 166,780 | 27,639 | sp\|O14744\|ANM5_HUMAN | 109,823 | 61,366 | 56% | O14744 | ANM5_HUMAN | ANM5_HUMAN | Protein arginine N-methyltransferase 5 (PRMT5) (EC 2.1.1.320) (72 kDa ICln-binding protein) (Histone-arginine N-methyltransferase PRMT5) (Jak-binding protein 1) (Shk1 kinase-binding protein 1 homolog) (SKB1 homolog) (SKB1Hs) [Cleaved into: Protein arginine N-methyltransferase 5, N-terminally processed] | PRMT5 HRMT1L5 IBP72 JBP1 SKB1 |
| sp\|Q8IVM0\|CCD50_HUMAN | 114,418 | 115,648 | 417,877 | 422,971 | sp\|Q8IVM0\|CCD50_HUMAN | 230,606 | 139,448 | 60% | Q8IVM0 | CCD50_HUMAN | CCD50_HUMAN | Coiled-coil domain-containing protein 50 (Protein Ymer) | CCDC50 C3orf6 |
| sp\|Q6P2Q9\|PRP8_HUMAN | 3,097 | 1,530 | 3,563 | 3,820 | sp\|Q6P2Q9\|PRP8_HUMAN | 6,267 | 4,061 | 65% | Q6P2Q9 | PRP8_HUMAN | PRP8_HUMAN | Pre-mRNA-processing-splicing factor 8 (220 kDa U5 snRNP-specific protein) (PRP8 homolog) (Splicing factor Prp8) (p220) | PRPF8 PRPC8 |
| sp\|P30490\|1B52_HUMAN | 104,803 | 78,633 | 63,052 | 166,508 | sp\|P30490\|1B52_HUMAN | 77,379 | 41,455 | 54% | P30490 | 1B52_HUMAN | HLAB_HUMAN | HLA class I histocompatibility antigen, B alpha chain (Human leukocyte antigen B) (HLA-B) | HLA-B HLAB |
| sp\|Q14203\|DCTN1_HUMAN | 38,603 | 52,215 | 36,069 | 18,861 | sp\|Q14203\|DCTN1_HUMAN | 34,325 | 12,335 | 36% | Q14203 | DCTN1_HUMAN | DCTN1_HUMAN | Dynactin subunit 1 (150 kDa dynein-associated polypeptide) (DAP-150) (DP-150) (p135) (p150-glued) | DCTN1 |
| sp\|Q06828\|FMOD_HUMAN | 34,524 | 25,685 | 24,222 | 40,665 | sp\|Q06828\|FMOD_HUMAN | 228,241 | 216,209 | 95% | Q06828 | FMOD_HUMAN | FMOD_HUMAN | Fibromodulin (FM) (Collagen-binding 59 kDa protein) (Keratan sulfate proteoglycan fibromodulin) (KSPG fibromodulin) | FMOD FM SLRR2E |
| sp\|Q96JB6\|LOXL4_HUMAN | 6,699 | 28,967 | 20,726 | 9,375 | sp\|Q96JB6\|LOXL4_HUMAN | 13,949 | 8,952 | 64% | Q96JB6 | LOXL4_HUMAN | LOXL4_HUMAN | Lysyl oxidase homolog 4 (EC 1.4.3.-) (Lysyl oxidase-like protein 4) (Lysyl oxidase-related protein C) | LOXL4 LOXC |
| sp\|Q9NR50\|EI2BG_HUMAN | 29,185 | 29,924 | 48,762 | 9,487 | sp\|Q9NR50\|EI2BG_HUMAN | 27,292 | 15,865 | 58% | Q9NR50 | EI2BG_HUMAN | EI2BG_HUMAN | Translation initiation factor eIF-2B subunit gamma (eIF-2B GDP-GTP exchange factor subunit gamma) | EIF2B3 |
| sp\|P47914\|RL29_HUMAN | 79,078 | 92,630 | 60,388 | 47,273 | sp\|P47914\|RL29_HUMAN | 45,068 | 30,634 | 68% | P47914 | RL29_HUMAN | RL29_HUMAN | 60S ribosomal protein L29 (Cell surface heparin-binding protein HIP) (Large ribosomal subunit protein eL29) | RPL29 |

| | | | | | | | | | | | | | |
|---|---|---|---|---|---|---|---|---|---|---|---|---|---|
| sp\|Q96N66\|MBOA7_HUMAN | 16,068 | 15,560 | 12,296 | 13,669 | sp\|Q96N66\|MBOA7_HUMAN | 18,566 | 9,087 | 49% | Q96N66 | MBOA7_HUMAN | MBOA7_HUMAN | Lysophospholipid acyltransferase 7 (LPLAT 7) (EC 2.3.1.-) (1-acylglycerophosphatidylinositol O-acyltransferase) (Bladder and breast carcinoma-overexpressed gene 1 protein) (Leukocyte receptor cluster member 4) (Lysophosphatidylinositol acyltransferase) (LPIAT) (Lyso-PI acyltransferase) (Membrane-bound O-acyltransferase domain-containing protein 7) (O-acyltransferase domain-containing protein 7) (h-mboa-7) | MBOAT7 BB1 LENG4 OACT7 |
| sp\|Q9Y2Q3\|GSTK1_HUMAN | 18,511 | 23,226 | 6,511 | 6,403 | sp\|Q9Y2Q3\|GSTK1_HUMAN | 10,253 | 7,210 | 70% | Q9Y2Q3 | GSTK1_HUMAN | GSTK1_HUMAN | Glutathione S-transferase kappa 1 (EC 2.5.1.18) (GST 13-13) (GST class-kappa) (GSTK1-1) (hGSTK1) (Glutathione S-transferase subunit 13) | GSTK1 HDCMD47P |
| sp\|Q9UKR5\|ERG28_HUMAN | 52,568 | 49,272 | 52,135 | 75,530 | sp\|Q9UKR5\|ERG28_HUMAN | 38,894 | 22,567 | 58% | Q9UKR5 | ERG28_HUMAN | ERG28_HUMAN | Ergosterol biosynthetic protein 28 homolog | ERG28 C14orf1 AD-011 HSPC288 x0006 |
| sp\|P16949\|STMN1_HUMAN | 52,459 | 53,980 | 142,764 | 151,213 | sp\|P16949\|STMN1_HUMAN | 77,700 | 49,852 | 64% | P16949 | STMN1_HUMAN | STMN1_HUMAN | Stathmin (Leukemia-associated phosphoprotein p18) (Metablastin) (Oncoprotein 18) (Op18) (Phosphoprotein p19) (pp19) (Prosolin) (Protein Pr22) (pp17) | STMN1 C1orf215 LAP18 OP18 |
| sp\|Q9BQA1\|MEP50_HUMAN | 76,830 | 74,684 | 131,323 | 145,486 | sp\|Q9BQA1\|MEP50_HUMAN | 87,469 | 38,083 | 44% | Q9BQA1 | MEP50_HUMAN | MEP50_HUMAN | Methylosome protein 50 (MEP-50) (Androgen receptor cofactor p44) (WD repeat-containing protein 77) (p44/Mep50) | WDR77 MEP50 WD45 HKMT1069 Nbla10071 |
| sp\|O75116\|ROCK2_HUMAN | 63,456 | 127,397 | 123,043 | 131,776 | sp\|O75116\|ROCK2_HUMAN | 99,464 | 38,186 | 38% | O75116 | ROCK2_HUMAN | ROCK2_HUMAN | Rho-associated protein kinase 2 (EC 2.7.11.1) (Rho kinase 2) (Rho-associated, coiled-coil-containing protein kinase 2) (Rho-associated, coiled-coil-containing protein kinase II) (ROCK-II) (p164 ROCK-2) | ROCK2 KIAA0619 |
| sp\|Q9Y5K5\|UCHL5_HUMAN | 33,851 | 33,695 | 54,618 | 10,721 | sp\|Q9Y5K5\|UCHL5_HUMAN | 89,804 | 135,484 | 151% | Q9Y5K5 | UCHL5_HUMAN | UCHL5_HUMAN | Ubiquitin carboxyl-terminal hydrolase isozyme L5 (UCH-L5) (EC 3.4.19.12) (Ubiquitin C-terminal hydrolase UCH37) (Ubiquitin thioesterase L5) | UCHL5 UCH37 AD-019 CGI-70 |
| sp\|Q9UL25\|RAB21_HUMAN | 102,924 | 88,664 | 160,302 | 158,108 | sp\|Q9UL25\|RAB21_HUMAN | 106,677 | 35,410 | 33% | Q9UL25 | RAB21_HUMAN | RAB21_HUMAN | Ras-related protein Rab-21 | RAB21 KIAA0118 |
| sp\|Q9P2E9\|RRBP1_HUMAN | 16,895 | 15,709 | 77,892 | 70,547 | sp\|Q9P2E9\|RRBP1_HUMAN | 32,631 | 27,927 | 86% | Q9P2E9 | RRBP1_HUMAN | RRBP1_HUMAN | Ribosome-binding protein 1 (180 kDa ribosome receptor homolog) (RRp) (ES/130-related protein) (Ribosome receptor protein) | RRBP1 KIAA1398 |
| sp\|Q99447\|PCY2_HUMAN | 27,592 | 30,385 | 112,043 | 81,841 | sp\|Q99447\|PCY2_HUMAN | 46,339 | 35,390 | 76% | Q99447 | PCY2_HUMAN | PCY2_HUMAN | Ethanolamine-phosphate cytidylyltransferase (EC 2.7.7.14) (CTP:phosphoethanolamine cytidylyltransferase) (Phosphorylethanolamine transferase) | PCYT2 |
| sp\|Q14165\|MLEC_HUMAN | 17,298 | 17,794 | 29,982 | 26,369 | sp\|Q14165\|MLEC_HUMAN | 22,745 | 12,491 | 55% | Q14165 | MLEC_HUMAN | MLEC_HUMAN | Malectin | MLEC KIAA0152 |
| sp\|P19256\|LFA3_HUMAN | 292,972 | 290,775 | 296,232 | 251,807 | sp\|P19256\|LFA3_HUMAN | 224,051 | 69,528 | 31% | P19256 | LFA3_HUMAN | LFA3_HUMAN | Lymphocyte function-associated antigen 3 (Ag3) (Surface glycoprotein LFA-3) (CD antigen CD58) | CD58 LFA3 |

| | | | | | | | | | | | | | |
|---|---|---|---|---|---|---|---|---|---|---|---|---|---|
| sp\|P22352\|GPX3_HUMAN | 13,423 | 12,823 | 5,680 | 27,091 | sp\|P22352\|GPX3_HUMAN | 48,739 | 42,842 | 88% | P22352 | GPX3_HUMAN | GPX3_HUMAN | Glutathione peroxidase 3 (GPx-3) (GSHPx-3) (EC 1.11.1.9) (Extracellular glutathione peroxidase) (Plasma glutathione peroxidase) (GPx-P) (GSHPx-P) | GPX3 GPXP |
| sp\|P60903\|S10AA_HUMAN | 155,059 | 242,149 | 66,840 | 27,711 | sp\|P60903\|S10AA_HUMAN | 132,336 | 73,868 | 56% | P60903 | S10AA_HUMAN | S10AA_HUMAN | Protein S100-A10 (Calpactin I light chain) (Calpactin-1 light chain) (Cellular ligand of annexin II) (S100 calcium-binding protein A10) (p10 protein) (p11) | S100A10 ANX2LG CAL1L CLP11 |
| sp\|O60684\|IMA7_HUMAN | 10,605 | 44,055 | 55,834 | 21,214 | sp\|O60684\|IMA7_HUMAN | 46,581 | 22,040 | 47% | O60684 | IMA7_HUMAN | IMA7_HUMAN | Importin subunit alpha-7 (Karyopherin subunit alpha-6) | KPNA6 IPOA7 |
| sp\|P50148\|GNAQ_HUMAN | 88,776 | 123,769 | 128,382 | 134,690 | sp\|P50148\|GNAQ_HUMAN | 122,775 | 15,120 | 12% | P50148 | GNAQ_HUMAN | GNAQ_HUMAN | Guanine nucleotide-binding protein G(q) subunit alpha (Guanine nucleotide-binding protein alpha-q) | GNAQ GAQ |
| sp\|Q9BXI6\|TB10A_HUMAN | 30,193 | 28,423 | 65,619 | 50,891 | sp\|Q9BXI6\|TB10A_HUMAN | 49,718 | 21,532 | 43% | Q9BXI6 | TB10A_HUMAN | TB10A_HUMAN | TBC1 domain family member 10A (EBP50-PDX interactor of 64 kDa) (EPI64 protein) (Rab27A-GAP-alpha) | TBC1D10A EPI64 TBC1D10 |
| sp\|Q13409\|DC1I2_HUMAN | 32,778 | 34,304 | 58,859 | 61,144 | sp\|Q13409\|DC1I2_HUMAN | 39,172 | 14,422 | 37% | Q13409 | DC1I2_HUMAN | DC1I2_HUMAN | Cytoplasmic dynein 1 intermediate chain 2 (Cytoplasmic dynein intermediate chain 2) (Dynein intermediate chain 2, cytosolic) (DH IC-2) | DYNC1I2 DNCI2 DNCIC2 |
| sp\|Q96BJ3\|AIDA_HUMAN | 52,714 | 72,060 | 99,911 | 99,813 | sp\|Q96BJ3\|AIDA_HUMAN | 62,093 | 33,251 | 54% | Q96BJ3 | AIDA_HUMAN | AIDA_HUMAN | Axin interactor, dorsalization-associated protein (Axin interaction partner and dorsalization antagonist) | AIDA C1orf80 |
| sp\|P35606\|COPB2_HUMAN | 71,767 | 51,042 | 35,193 | 14,488 | sp\|P35606\|COPB2_HUMAN | 55,084 | 49,322 | 90% | P35606 | COPB2_HUMAN | COPB2_HUMAN | Coatomer subunit beta' (Beta'-coat protein) (Beta'-COP) (p102) | COPB2 |
| sp\|Q8N9N7\|LRC57_HUMAN | 128,072 | 126,215 | 63,653 | 10,942 | sp\|Q8N9N7\|LRC57_HUMAN | 74,875 | 47,539 | 63% | Q8N9N7 | LRC57_HUMAN | LRC57_HUMAN | Leucine-rich repeat-containing protein 57 | LRRC57 |
| sp\|Q9UNE7\|CHIP_HUMAN | 78,023 | 114,326 | 126,884 | 158,188 | sp\|Q9UNE7\|CHIP_HUMAN | 85,704 | 48,209 | 56% | Q9UNE7 | CHIP_HUMAN | CHIP_HUMAN | E3 ubiquitin-protein ligase CHIP (EC 2.3.2.27) (Antigen NY-CO-7) (CLL-associated antigen KW-8) (Carboxy terminus of Hsp70-interacting protein) (RING-type E3 ubiquitin transferase CHIP) (STIP1 homology and U box-containing protein 1) | STUB1 CHIP PP1131 |
| sp\|P55036\|PSMD4_HUMAN | 4,645 | 16,378 | 13,053 | 44,300 | sp\|P55036\|PSMD4_HUMAN | 21,564 | 13,412 | 62% | P55036 | PSMD4_HUMAN | PSMD4_HUMAN | 26S proteasome non-ATPase regulatory subunit 4 (26S proteasome regulatory subunit RPN10) (26S proteasome regulatory subunit S5A) (Antisecretory factor 1) (AF) (ASF) (Multiubiquitin chain-binding protein) | PSMD4 MCB1 |
| sp\|P04004\|VTNC_HUMAN | 530,108 | 270,368 | 747,347 | 452,686 | sp\|P04004\|VTNC_HUMAN | 3,473,862 | 3,419,156 | 98% | P04004 | VTNC_HUMAN | VTNC_HUMAN | Vitronectin (VN) (S-protein) (Serum-spreading factor) (V75) [Cleaved into: Vitronectin V65 subunit; Vitronectin V10 subunit; Somatomedin-B] | VTN |
| sp\|Q9NZJ7\|MTCH1_HUMAN | 14,998 | 17,099 | 17,590 | 50,444 | sp\|Q9NZJ7\|MTCH1_HUMAN | 22,717 | 14,875 | 65% | Q9NZJ7 | MTCH1_HUMAN | MTCH1_HUMAN | Mitochondrial carrier homolog 1 (Presenilin-associated protein) | MTCH1 PSAP CGI-64 UNQ1871/PRO4314 |
| sp\|P09496\|CLCA_HUMAN | 236,952 | 98,664 | 320,091 | 320,520 | sp\|P09496\|CLCA_HUMAN | 201,906 | 99,430 | 49% | P09496 | CLCA_HUMAN | CLCA_HUMAN | Clathrin light chain A (Lca) | CLTA |
| sp\|Q15363\|TMED2_HUMAN | 13,472 | 16,458 | 3,162 | 18,843 | sp\|Q15363\|TMED2_HUMAN | 8,852 | 6,733 | 76% | Q15363 | TMED2_HUMAN | TMED2_HUMAN | Transmembrane emp24 domain-containing protein 2 (Membrane protein p24A) (p24) (p24 family protein beta-1) (p24beta1) | TMED2 RNP24 |

| | | | | | | | | | | | | | |
|---|---|---|---|---|---|---|---|---|---|---|---|---|---|
| sp\|P46459\|NSF_HUMAN | 6,540 | 1,584 | 4,948 | 4,171 | sp\|P46459\|NSF_HUMAN | 7,070 | 6,529 | 92% | P46459 | NSF_HUMAN | NSF_HUMAN | Vesicle-fusing ATPase (EC 3.6.4.6) (N-ethylmaleimide-sensitive fusion protein) (NEM-sensitive fusion protein) (Vesicular-fusion protein NSF) | NSF |
| sp\|Q9Y570\|PPME1_HUMAN | 24,917 | 12,114 | 15,660 | 4,420 | sp\|Q9Y570\|PPME1_HUMAN | 30,884 | 30,580 | 99% | Q9Y570 | PPME1_HUMAN | PPME1_HUMAN | Protein phosphatase methylesterase 1 (PME-1) (EC 3.1.1.89) | PPME1 PME1 PP2593 PRO0750 |
| sp\|P05387\|RLA2_HUMAN | 114,520 | 112,924 | 71,518 | 66,811 | sp\|P05387\|RLA2_HUMAN | 78,131 | 30,664 | 39% | P05387 | RLA2_HUMAN | RLA2_HUMAN | 60S acidic ribosomal protein P2 (Large ribosomal subunit protein P2) (Renal carcinoma antigen NY-REN-44) | RPLP2 D11S2243E RPP2 |
| sp\|Q9H3K6\|BOLA2_HUMAN | 25,008 | 27,931 | 100,089 | 123,791 | sp\|Q9H3K6\|BOLA2_HUMAN | 52,755 | 42,036 | 80% | Q9H3K6 | BOLA2_HUMAN | BOLA2_HUMAN | BolA-like protein 2 | BOLA2 BOLA2A My016; BOLA2B |
| sp\|Q9BQB6\|VKOR1_HUMAN | 45,061 | 43,350 | 71,049 | 72,234 | sp\|Q9BQB6\|VKOR1_HUMAN | 43,143 | 21,658 | 50% | Q9BQB6 | VKOR1_HUMAN | VKOR1_HUMAN | Vitamin K epoxide reductase complex subunit 1 (EC 1.17.4.4) (Vitamin K1 2,3-epoxide reductase subunit 1) | VKORC1 VKOR MSTP134 MSTP576 UNQ308/PRO351 |
| sp\|P42574\|CASP3_HUMAN | 22,065 | 20,457 | 58,738 | 50,868 | sp\|P42574\|CASP3_HUMAN | 38,367 | 14,661 | 38% | P42574 | CASP3_HUMAN | CASP3_HUMAN | Caspase-3 (CASP-3) (EC 3.4.22.56) (Apopain) (Cysteine protease CPP32) (CPP-32) (Protein Yama) (SREBP cleavage activity 1) (SCA-1) [Cleaved into: Caspase-3 subunit p17; Caspase-3 subunit p12] | CASP3 CPP32 |
| sp\|P20338\|RAB4A_HUMAN | 55,835 | 58,621 | 90,284 | 121,502 | sp\|P20338\|RAB4A_HUMAN | 77,539 | 26,209 | 34% | P20338 | RAB4A_HUMAN | RAB4A_HUMAN | Ras-related protein Rab-4A (EC 3.6.5.2) | RAB4A RAB4 |
| sp\|Q9NRV9\|HEBP1_HUMAN | 31,553 | 39,908 | 77,980 | 65,602 | sp\|Q9NRV9\|HEBP1_HUMAN | 40,089 | 22,957 | 57% | Q9NRV9 | HEBP1_HUMAN | HEBP1_HUMAN | Heme-binding protein 1 (p22HBP) | HEBP1 HBP |
| sp\|Q6XQN6\|PNCB_HUMAN | 20,717 | 39,553 | 10,730 | 31,057 | sp\|Q6XQN6\|PNCB_HUMAN | 37,480 | 36,123 | 96% | Q6XQN6 | PNCB_HUMAN | PNCB_HUMAN | Nicotinate phosphoribosyltransferase (NAPRTase) (EC 6.3.4.21) (FHA-HIT-interacting protein) (Nicotinate phosphoribosyltransferase domain-containing protein 1) | NAPRT FHIP NAPRT1 |
| sp\|P36551\|HEM6_HUMAN | 19,140 | 26,575 | 17,001 | 57,301 | sp\|P36551\|HEM6_HUMAN | 24,498 | 14,835 | 61% | P36551 | HEM6_HUMAN | HEM6_HUMAN | Oxygen-dependent coproporphyrinogen-III oxidase, mitochondrial (COX) (Coprogen oxidase) (Coproporphyrinogenase) (EC 1.3.3.3) | CPOX CPO CPX |
| sp\|P56377\|AP1S2_HUMAN | 69,968 | 68,252 | 87,781 | 133,514 | sp\|P56377\|AP1S2_HUMAN | 86,252 | 24,240 | 28% | P56377 | AP1S2_HUMAN | AP1S2_HUMAN | AP-1 complex subunit sigma-2 (Adaptor protein complex AP-1 subunit sigma-1B) (Adaptor-related protein complex 1 subunit sigma-1B) (Clathrin assembly protein complex 1 sigma-1B small chain) (Golgi adaptor HA1/AP1 adaptin sigma-1B subunit) (Sigma 1B subunit of AP-1 clathrin) (Sigma-adaptin 1B) (Sigma1B-adaptin) | AP1S2 DC22 |
| sp\|P08697\|A2AP_HUMAN | 29,243 | 15,732 | 38,520 | 20,582 | sp\|P08697\|A2AP_HUMAN | 391,079 | 423,424 | 108% | P08697 | A2AP_HUMAN | A2AP_HUMAN | Alpha-2-antiplasmin (Alpha-2-AP) (Alpha-2-plasmin inhibitor) (Alpha-2-PI) (Serpin F2) | SERPINF2 AAP PLI |
| sp\|O75674\|TM1L1_HUMAN | 118,841 | 170,584 | 270,971 | 291,496 | sp\|O75674\|TM1L1_HUMAN | 233,674 | 64,463 | 28% | O75674 | TM1L1_HUMAN | TM1L1_HUMAN | TOM1-like protein 1 (Src-activating and signaling molecule protein) (Target of Myb-like protein 1) | TOM1L1 SRCASM |
| sp\|Q13885\|TBB2A_HUMAN | 90,626 | 83,809 | 84,890 | 81,113 | sp\|Q13885\|TBB2A_HUMAN | 61,714 | 25,423 | 41% | Q13885 | TBB2A_HUMAN | TBB2A_HUMAN | Tubulin beta-2A chain (Tubulin beta class IIa) | TUBB2A TUBB2 |
| sp\|Q71U36\|TBA1A_HUMAN | 76,601 | 515,737 | 92,405 | 602,653 | sp\|Q71U36\|TBA1A_HUMAN | 341,097 | 184,915 | 54% | Q71U36 | TBA1A_HUMAN | TBA1A_HUMAN | Tubulin alpha-1A chain (Alpha-tubulin 3) (Tubulin B-alpha-1) (Tubulin alpha-3 chain) [Cleaved into: Detyrosinated tubulin alpha-1A chain] | TUBA1A TUBA3 |

| | | | | | | | | | | | | | |
|---|---|---|---|---|---|---|---|---|---|---|---|---|---|
| sp\|P20039\|2B1B_HUMAN | 722,090 | 736,816 | 398,333 | 363,661 | sp\|P20039\|2B1B_HUMAN | 454,280 | 193,659 | 43% | P20039 | 2B1B_HUMAN | DRB1_HUMAN | HLA class II histocompatibility antigen, DRB1 beta chain (Human leukocyte antigen DRB1) (HLA-DRB1) | HLA-DRB1 |
| sp\|P84085\|ARF5_HUMAN | 26,925 | 26,485 | 23,962 | 8,085 | sp\|P84085\|ARF5_HUMAN | 18,730 | 8,605 | 46% | P84085 | ARF5_HUMAN | ARF5_HUMAN | ADP-ribosylation factor 5 | ARF5 |
| sp\|P06241\|FYN_HUMAN | 13,429 | 907 | 16,588 | 2,927 | sp\|P06241\|FYN_HUMAN | 5,861 | 6,184 | 106% | P06241 | FYN_HUMAN | FYN_HUMAN | Tyrosine-protein kinase Fyn (EC 2.7.10.2) (Proto-oncogene Syn) (Proto-oncogene c-Fyn) (Src-like kinase) (SLK) (p59-Fyn) | FYN |
| sp\|P11908\|PRPS2_HUMAN | 33,948 | 30,599 | 15,382 | 8,377 | sp\|P11908\|PRPS2_HUMAN | 15,330 | 11,876 | 77% | P11908 | PRPS2_HUMAN | PRPS2_HUMAN | Ribose-phosphate pyrophosphokinase 2 (EC 2.7.6.1) (PPRibP) (Phosphoribosyl pyrophosphate synthase II) (PRS-II) | PRPS2 |
| sp\|P08727\|K1C19_HUMAN | 668 | 332 | 1,336 | 1,254 | sp\|P08727\|K1C19_HUMAN | 1,513 | 1,453 | 96% | P08727 | K1C19_HUMAN | K1C19_HUMAN | Keratin, type I cytoskeletal 19 (Cytokeratin-19) (CK-19) (Keratin-19) (K19) | KRT19 |
| sp\|P10114\|RAP2A_HUMAN | 57,870 | 85,758 | 23,001 | 100,808 | sp\|P10114\|RAP2A_HUMAN | 77,332 | 30,282 | 39% | P10114 | RAP2A_HUMAN | RAP2A_HUMAN | Ras-related protein Rap-2a (EC 3.6.5.2) (RbBP-30) | RAP2A |
| sp\|Q71UI9\|H2AV_HUMAN | 1,056 | 1,101 | 2,276 | 3,078 | sp\|Q71UI9\|H2AV_HUMAN | 3,220 | 2,209 | 69% | Q71UI9 | H2AV_HUMAN | H2AV_HUMAN | Histone H2A.V (H2A.F/Z) (H2A.Z variant histone 2) | H2AZ2 H2AFV H2AV |
| sp\|Q9BY32\|ITPA_HUMAN | 16,286 | 23,643 | 15,775 | 31,980 | sp\|Q9BY32\|ITPA_HUMAN | 19,567 | 7,091 | 36% | Q9BY32 | ITPA_HUMAN | ITPA_HUMAN | Inosine triphosphate pyrophosphatase (ITPase) (Inosine triphosphatase) (EC 3.6.1.9) (Non-canonical purine NTP pyrophosphatase) (Non-standard purine NTP pyrophosphatase) (Nucleoside-triphosphate diphosphatase) (Nucleoside-triphosphate pyrophosphatase) (NTPase) (Putative oncogene protein hlc14-06-p) | ITPA C20orf37 My049 OK/SW-cl.9 |
| sp\|O95336\|6PGL_HUMAN | 28,951 | 25,457 | 31,005 | 25,806 | sp\|O95336\|6PGL_HUMAN | 36,316 | 10,036 | 28% | O95336 | 6PGL_HUMAN | 6PGL_HUMAN | 6-phosphogluconolactonase (6PGL) (EC 3.1.1.31) | PGLS |
| sp\|Q9Y5A9\|YTHD2_HUMAN | 12,574 | 11,645 | 12,486 | 8,641 | sp\|Q9Y5A9\|YTHD2_HUMAN | 14,322 | 5,985 | 42% | Q9Y5A9 | YTHD2_HUMAN | YTHD2_HUMAN | YTH domain-containing family protein 2 (DF2) (CLL-associated antigen KW-14) (High-glucose-regulated protein 8) (Renal carcinoma antigen NY-REN-2) | YTHDF2 HGRG8 |
| sp\|Q9NPQ8\|RIC8A_HUMAN | 3,920 | 15,481 | 9,518 | 5,304 | sp\|Q9NPQ8\|RIC8A_HUMAN | 9,316 | 5,227 | 56% | Q9NPQ8 | RIC8A_HUMAN | RIC8A_HUMAN | Synembryn-A (Protein Ric-8A) | RIC8A |
| sp\|Q92982\|NINJ1_HUMAN | 26,099 | 29,300 | 10,819 | 40,970 | sp\|Q92982\|NINJ1_HUMAN | 23,253 | 14,055 | 60% | Q92982 | NINJ1_HUMAN | NINJ1_HUMAN | Ninjurin-1 (Nerve injury-induced protein 1) [Cleaved into: Secreted ninjurin-1 (Soluble ninjurin-1)] | NINJ1 |
| sp\|Q13153\|PAK1_HUMAN | 7,547 | 7,943 | 37,731 | 27,304 | sp\|Q13153\|PAK1_HUMAN | 23,825 | 13,813 | 58% | Q13153 | PAK1_HUMAN | PAK1_HUMAN | Serine/threonine-protein kinase PAK 1 (EC 2.7.11.1) (Alpha-PAK) (p21-activated kinase 1) (PAK-1) (p65-PAK) | PAK1 |
| sp\|Q03692\|COAA1_HUMAN | 91,442 | 8,043 | 14,086 | 14,275 | sp\|Q03692\|COAA1_HUMAN | 48,176 | 44,168 | 92% | Q03692 | COAA1_HUMAN | COAA1_HUMAN | Collagen alpha-1(X) chain | COL10A1 |
| sp\|P53618\|COPB_HUMAN | 70,041 | 69,429 | 28,686 | 81,576 | sp\|P53618\|COPB_HUMAN | 50,279 | 25,320 | 50% | P53618 | COPB_HUMAN | COPB_HUMAN | Coatomer subunit beta (Beta-coat protein) (Beta-COP) | COPB1 COPB MSTP026 |
| sp\|O43708\|MAAI_HUMAN | 20,711 | 33,967 | 6,152 | 6,020 | sp\|O43708\|MAAI_HUMAN | 11,152 | 10,688 | 96% | O43708 | MAAI_HUMAN | MAAI_HUMAN | Maleylacetoacetate isomerase (MAAI) (EC 5.2.1.2) (GSTZ1-1) (Glutathione S-transferase zeta 1) (EC 2.5.1.18) | GSTZ1 MAAI |
| sp\|Q9Y3D6\|FIS1_HUMAN | 17,368 | 26,133 | 1,851 | 4,158 | sp\|Q9Y3D6\|FIS1_HUMAN | 10,851 | 10,396 | 96% | Q9Y3D6 | FIS1_HUMAN | FIS1_HUMAN | Mitochondrial fission 1 protein (FIS1 homolog) (hFis1) (Tetratricopeptide repeat protein 11) (TPR repeat protein 11) | FIS1 TTC11 CGI-135 |

| | | | | | | | | | | | | | |
|---|---|---|---|---|---|---|---|---|---|---|---|---|---|
| sp\|Q9P0J0\|NDUAD_HUMAN | 4,491 | 1,570 | 4,997 | 8,183 | sp\|Q9P0J0\|NDUAD_HUMAN | 4,394 | 2,242 | 51% | Q9P0J0 | NDUAD_HUMAN | NDUAD_HUMAN | NADH dehydrogenase [ubiquinone] 1 alpha subcomplex subunit 13 (Cell death regulatory protein GRIM-19) (Complex I-B16.6) (CI-B16.6) (Gene associated with retinoic and interferon-induced mortality 19 protein) (GRIM-19) (Gene associated with retinoic and IFN-induced mortality 19 protein) (NADH-ubiquinone oxidoreductase B16.6 subunit) | NDUFA13 GRIM19 CDA016 CGI-39 |
| sp\|Q9NRP0\|OSTC_HUMAN | 61,117 | 92,467 | 19,892 | 56,988 | sp\|Q9NRP0\|OSTC_HUMAN | 43,099 | 27,929 | 65% | Q9NRP0 | OSTC_HUMAN | OSTC_HUMAN | Oligosaccharyltransferase complex subunit OSTC (Hydrophobic protein HSF-28) | OSTC DC2 HDCMD45P HSPC307 |
| sp\|Q9H6V9\|LDAH_HUMAN | 45,866 | 62,543 | 35,913 | 107,751 | sp\|Q9H6V9\|LDAH_HUMAN | 49,755 | 29,908 | 60% | Q9H6V9 | LDAH_HUMAN | LDAH_HUMAN | Lipid droplet-associated hydrolase (EC 3.1.1.-) (Lipid droplet-associated serine hydrolase) (hLDAH) | LDAH C2orf43 |
| sp\|Q9BRA2\|TXD17_HUMAN | 37,806 | 42,885 | 18,400 | 32,820 | sp\|Q9BRA2\|TXD17_HUMAN | 34,694 | 20,823 | 60% | Q9BRA2 | TXD17_HUMAN | TXD17_HUMAN | Thioredoxin domain-containing protein 17 (14 kDa thioredoxin-related protein) (TRP14) (Protein 42-9-9) (Thioredoxin-like protein 5) | TXNDC17 TXNL5 |
| sp\|Q96RD7\|PANX1_HUMAN | 23,893 | 2,215 | 2,279 | 2,617 | sp\|Q96RD7\|PANX1_HUMAN | 8,298 | 7,803 | 94% | Q96RD7 | PANX1_HUMAN | PANX1_HUMAN | Pannexin-1 | PANX1 MRS1 UNQ2529/PRO6028 |
| sp\|Q969T9\|WBP2_HUMAN | 59,537 | 61,083 | 121,260 | 21,560 | sp\|Q969T9\|WBP2_HUMAN | 51,750 | 35,832 | 69% | Q969T9 | WBP2_HUMAN | WBP2_HUMAN | WW domain-binding protein 2 (WBP-2) | WBP2 |
| sp\|Q969E2\|SCAM4_HUMAN | 89,570 | 99,288 | 66,141 | 50,417 | sp\|Q969E2\|SCAM4_HUMAN | 54,127 | 30,009 | 55% | Q969E2 | SCAM4_HUMAN | SCAM4_HUMAN | Secretory carrier-associated membrane protein 4 (Secretory carrier membrane protein 4) | SCAMP4 |
| sp\|Q92485\|ASM3B_HUMAN | 59,475 | 62,487 | 113,997 | 130,800 | sp\|Q92485\|ASM3B_HUMAN | 73,914 | 33,157 | 45% | Q92485 | ASM3B_HUMAN | ASM3B_HUMAN | Acid sphingomyelinase-like phosphodiesterase 3b (ASM-like phosphodiesterase 3b) (EC 3.1.4.-) | SMPDL3B ASML3B ASMLPD |
| sp\|Q8WUM9\|S20A1_HUMAN | 134,890 | 152,195 | 93,927 | 11,380 | sp\|Q8WUM9\|S20A1_HUMAN | 98,798 | 58,515 | 59% | Q8WUM9 | S20A1_HUMAN | S20A1_HUMAN | Sodium-dependent phosphate transporter 1 (Gibbon ape leukemia virus receptor 1) (GLVR-1) (Leukemia virus receptor 1 homolog) (Phosphate transporter 1) (PiT-1) (Solute carrier family 20 member 1) | SLC20A1 GLVR1 PIT1 |
| sp\|Q5SRD1\|TI23B_HUMAN | 8,596 | 7,340 | 8,521 | 78,972 | sp\|Q5SRD1\|TI23B_HUMAN | 57,097 | 48,680 | 85% | Q5SRD1 | TI23B_HUMAN | TI23B_HUMAN | Mitochondrial import inner membrane translocase subunit Tim23B (TIMM23B) | TIMM23B |
| sp\|Q15717\|ELAV1_HUMAN | 40,857 | 36,474 | 10,649 | 68,018 | sp\|Q15717\|ELAV1_HUMAN | 31,000 | 20,464 | 66% | Q15717 | ELAV1_HUMAN | ELAV1_HUMAN | ELAV-like protein 1 (Hu-antigen R) (HuR) | ELAVL1 HUR |
| sp\|Q15388\|TOM20_HUMAN | 4,412 | 3,741 | 15,311 | 4,760 | sp\|Q15388\|TOM20_HUMAN | 10,311 | 6,884 | 67% | Q15388 | TOM20_HUMAN | TOM20_HUMAN | Mitochondrial import receptor subunit TOM20 homolog (Mitochondrial 20 kDa outer membrane protein) (Outer mitochondrial membrane receptor Tom20) | TOMM20 KIAA0016 |
| sp\|Q15043\|S39AE_HUMAN | 113,009 | 110,037 | 79,926 | 190,558 | sp\|Q15043\|S39AE_HUMAN | 125,031 | 38,982 | 31% | Q15043 | S39AE_HUMAN | S39AE_HUMAN | Metal cation symporter ZIP14 (LIV-1 subfamily of ZIP zinc transporter 4) (LZT-Hs4) (Solute carrier family 39 member 14) (Zrt- and Irt-like protein 14) (ZIP-14) | SLC39A14 KIAA0062 ZIP14 |
| sp\|Q14728\|MFS10_HUMAN | 9,114 | 12,453 | 2,607 | 3,429 | sp\|Q14728\|MFS10_HUMAN | 7,398 | 3,673 | 50% | Q14728 | MFS10_HUMAN | MFS10_HUMAN | Major facilitator superfamily domain-containing protein 10 (Tetracycline transporter-like protein) | MFSD10 TETRAN |
| sp\|Q13907\|IDI1_HUMAN | 18,895 | 39,725 | 39,655 | 61,524 | sp\|Q13907\|IDI1_HUMAN | 41,595 | 32,144 | 77% | Q13907 | IDI1_HUMAN | IDI1_HUMAN | Isopentenyl-diphosphate Delta-isomerase 1 (EC 5.3.3.2) (Isopentenyl pyrophosphate isomerase 1) (IPP isomerase 1) (IPPI1) | IDI1 |

| | | | | | | | | | | | | | |
|---|---|---|---|---|---|---|---|---|---|---|---|---|---|
| sp\|Q13148\|TADBP_HUMAN | 23,540 | 35,571 | 4,322 | 7,597 | sp\|Q13148\|TADBP_HUMAN | 15,826 | 9,927 | 63% | Q13148 | TADBP_HUMAN | TADBP_HUMAN | TAR DNA-binding protein 43 (TDP-43) | TARDBP TDP43 |
| sp\|Q01629\|IFM2_HUMAN | 64,618 | 67,701 | 5,126 | 27,358 | sp\|Q01629\|IFM2_HUMAN | 31,800 | 25,090 | 79% | Q01629 | IFM2_HUMAN | IFM2_HUMAN | Interferon-induced transmembrane protein 2 (Dispanin subfamily A member 2c) (DSPA2c) (Interferon-inducible protein 1-8D) | IFITM2 |
| sp\|P80297\|MT1X_HUMAN | 33,567 | 21,335 | 18,013 | 6,590 | sp\|P80297\|MT1X_HUMAN | 16,293 | 12,707 | 78% | P80297 | MT1X_HUMAN | MT1X_HUMAN | Metallothionein-1X (MT-1X) (Metallothionein-IX) (MT-IX) | MT1X |
| sp\|P62861\|RS30_HUMAN | 115,917 | 192,544 | 107,117 | 122,499 | sp\|P62861\|RS30_HUMAN | 117,599 | 54,983 | 47% | P62861 | RS30_HUMAN | RS30_HUMAN | FAU ubiquitin-like and ribosomal protein S30 [Cleaved into: Ubiquitin-like protein FUBI; 40S ribosomal protein S30 (Small ribosomal subunit protein eS30)] | FAU |
| sp\|P62304\|RUXE_HUMAN | 95,915 | 163,258 | 51,922 | 109,834 | sp\|P62304\|RUXE_HUMAN | 70,432 | 50,413 | 72% | P62304 | RUXE_HUMAN | RUXE_HUMAN | Small nuclear ribonucleoprotein E (snRNP-E) (Sm protein E) (Sm-E) (SmE) | SNRPE |
| sp\|P52306\|GDS1_HUMAN | 23,963 | 29,254 | 139,935 | 54,860 | sp\|P52306\|GDS1_HUMAN | 71,566 | 51,666 | 72% | P52306 | GDS1_HUMAN | GDS1_HUMAN | Rap1 GTPase-GDP dissociation stimulator 1 (Exchange factor smgGDS) (SMG GDS protein) (SMG P21 stimulatory GDP/GTP exchange protein) | RAP1GDS1 SMGGDS |
| sp\|P41240\|CSK_HUMAN | 18,976 | 63,569 | 30,723 | 105,533 | sp\|P41240\|CSK_HUMAN | 47,099 | 32,195 | 68% | P41240 | CSK_HUMAN | CSK_HUMAN | Tyrosine-protein kinase CSK (EC 2.7.10.2) (C-Src kinase) (Protein-tyrosine kinase CYL) | CSK |
| sp\|P39210\|MPV17_HUMAN | 4,944 | 13,228 | 7,256 | 7,247 | sp\|P39210\|MPV17_HUMAN | 9,259 | 4,426 | 48% | P39210 | MPV17_HUMAN | MPV17_HUMAN | Protein Mpv17 | MPV17 |
| sp\|P30049\|ATPD_HUMAN | 11,065 | 15,505 | 10,203 | 8,982 | sp\|P30049\|ATPD_HUMAN | 10,070 | 3,780 | 38% | P30049 | ATPD_HUMAN | ATPD_HUMAN | ATP synthase subunit delta, mitochondrial (ATP synthase F1 subunit delta) (F-ATPase delta subunit) | ATP5F1D ATP5D |
| sp\|P28065\|PSB9_HUMAN | 18,227 | 22,773 | 61,345 | 118,662 | sp\|P28065\|PSB9_HUMAN | 49,977 | 44,852 | 90% | P28065 | PSB9_HUMAN | PSB9_HUMAN | Proteasome subunit beta type-9 (EC 3.4.25.1) (Low molecular mass protein 2) (Macropain chain 7) (Multicatalytic endopeptidase complex chain 7) (Proteasome chain 7) (Proteasome subunit beta-1i) (Really interesting new gene 12 protein) | PSMB9 LMP2 PSMB6i RING12 |
| sp\|P23297\|S10A1_HUMAN | 88,885 | 193,637 | 23,334 | 29,306 | sp\|P23297\|S10A1_HUMAN | 91,315 | 71,132 | 78% | P23297 | S10A1_HUMAN | S10A1_HUMAN | Protein S100-A1 (S-100 protein alpha chain) (S-100 protein subunit alpha) (S100 calcium-binding protein A1) | S100A1 S100A |
| sp\|P12429\|ANXA3_HUMAN | 1,658 | 2,476 | 1,891 | 1,621 | sp\|P12429\|ANXA3_HUMAN | 3,011 | 2,541 | 84% | P12429 | ANXA3_HUMAN | ANXA3_HUMAN | Annexin A3 (35-alpha calcimedin) (Annexin III) (Annexin-3) (Inositol 1,2-cyclic phosphate 2-phosphohydrolase) (Lipocortin III) (Placental anticoagulant protein III) (PAP-III) | ANXA3 ANX3 |
| sp\|P09960\|LKHA4_HUMAN | 0 | 155 | 0 | 2,644 | sp\|P09960\|LKHA4_HUMAN | 1,072 | 1,198 | 112% | P09960 | LKHA4_HUMAN | LKHA4_HUMAN | Leukotriene A-4 hydrolase (LTA-4 hydrolase) (EC 3.3.2.6) (Leukotriene A(4) hydrolase) (Tripeptide aminopeptidase LTA4H) (EC 3.4.11.4) | LTA4H LTA4 |
| sp\|P02647\|APOA1_HUMAN | 46,582 | 47,781 | 20,312 | 14,376 | sp\|P02647\|APOA1_HUMAN | 418,364 | 440,574 | 105% | P02647 | APOA1_HUMAN | APOA1_HUMAN | Apolipoprotein A-I (Apo-AI) (ApoA-I) (Apolipoprotein A1) [Cleaved into: Proapolipoprotein A-I (ProapoA-I); Truncated apolipoprotein A-I (Apolipoprotein A-I(1-242))] | APOA1 |
| sp\|P00846\|ATP6_HUMAN | 47,562 | 4,206 | 28,615 | 16,595 | sp\|P00846\|ATP6_HUMAN | 19,118 | 13,472 | 70% | P00846 | ATP6_HUMAN | ATP6_HUMAN | ATP synthase subunit a (F-ATPase protein 6) | MT-ATP6 ATP6 ATPASE6 MTATP6 |
| sp\|O95219\|SNX4_HUMAN | 6,849 | 32,049 | 50,420 | 26,884 | sp\|O95219\|SNX4_HUMAN | 25,015 | 15,527 | 62% | O95219 | SNX4_HUMAN | SNX4_HUMAN | Sorting nexin-4 | SNX4 |

| | | | | | | | | | | | | | |
|---|---|---|---|---|---|---|---|---|---|---|---|---|---|
| sp\|O95164\|UBL3_HUMAN | 82,485 | 79,046 | 127,903 | 105,408 | sp\|O95164\|UBL3_HUMAN | 91,227 | 24,931 | 27% | O95164 | UBL3_HUMAN | UBL3_HUMAN | Ubiquitin-like protein 3 (Membrane-anchored ubiquitin-fold protein) (HsMUB) (MUB) (Protein HCG-1) | UBL3 PNSC1 |
| sp\|O75352\|MPU1_HUMAN | 1,153 | 24,699 | 12,554 | 17,670 | sp\|O75352\|MPU1_HUMAN | 10,023 | 8,334 | 83% | O75352 | MPU1_HUMAN | MPU1_HUMAN | Mannose-P-dolichol utilization defect 1 protein (Suppressor of Lec15 and Lec35 glycosylation mutation homolog) (SL15) | MPDU1 |
| sp\|O15511\|ARPC5_HUMAN | 172,358 | 227,677 | 169,038 | 200,340 | sp\|O15511\|ARPC5_HUMAN | 157,050 | 62,447 | 40% | O15511 | ARPC5_HUMAN | ARPC5_HUMAN | Actin-related protein 2/3 complex subunit 5 (Arp2/3 complex 16 kDa subunit) (p16-ARC) | ARPC5 ARC16 |
| sp\|O15258\|RER1_HUMAN | 50,301 | 42,072 | 22,333 | 11,591 | sp\|O15258\|RER1_HUMAN | 32,356 | 16,656 | 51% | O15258 | RER1_HUMAN | RER1_HUMAN | Protein RER1 | RER1 |
| sp\|O00764\|PDXK_HUMAN | 11,921 | 21,159 | 13,783 | 20,559 | sp\|O00764\|PDXK_HUMAN | 24,391 | 31,382 | 129% | O00764 | PDXK_HUMAN | PDXK_HUMAN | Pyridoxal kinase (EC 2.7.1.35) (Pyridoxine kinase) | PDXK C21orf124 C21orf97 PKH PNK PRED79 |
| sp\|Q16650\|TBR1_HUMAN | 152,391 | 132,060 | 244,657 | 235,971 | sp\|Q16650\|TBR1_HUMAN | 143,161 | 64,920 | 45% | Q16650 | TBR1_HUMAN | TBR1_HUMAN | T-box brain protein 1 (T-brain-1) (TBR-1) (TES-56) | TBR1 |
| sp\|P48444\|COPD_HUMAN | 54,865 | 51,241 | 69,955 | 5,258 | sp\|P48444\|COPD_HUMAN | 59,881 | 44,401 | 74% | P48444 | COPD_HUMAN | COPD_HUMAN | Coatomer subunit delta (Archain) (Delta-coat protein) (Delta-COP) | ARCN1 COPD |
| sp\|Q8IX04\|UEVLD_HUMAN | 74,935 | 72,245 | 120,878 | 132,178 | sp\|Q8IX04\|UEVLD_HUMAN | 88,790 | 38,001 | 43% | Q8IX04 | UEVLD_HUMAN | UEVLD_HUMAN | Ubiquitin-conjugating enzyme E2 variant 3 (UEV-3) (EV and lactate/malate dehydrogenase domain-containing protein) | UEVLD UEV3 |
| sp\|P62306\|RUXF_HUMAN | 36,671 | 37,171 | 23,867 | 60,483 | sp\|P62306\|RUXF_HUMAN | 27,542 | 17,847 | 65% | P62306 | RUXF_HUMAN | RUXF_HUMAN | Small nuclear ribonucleoprotein F (snRNP-F) (Sm protein F) (Sm-F) (SmF) | SNRPF PBSCF |
| sp\|P22303\|ACES_HUMAN | 3,088 | 2,343 | 6,549 | 2,350 | sp\|P22303\|ACES_HUMAN | 30,300 | 32,591 | 108% | P22303 | ACES_HUMAN | ACES_HUMAN | Acetylcholinesterase (AChE) (EC 3.1.1.7) | ACHE |
| sp\|P11172\|UMPS_HUMAN | 24,129 | 25,750 | 34,887 | 17,110 | sp\|P11172\|UMPS_HUMAN | 35,223 | 23,617 | 67% | P11172 | UMPS_HUMAN | UMPS_HUMAN | Uridine 5'-monophosphate synthase (UMP synthase) [Includes: Orotate phosphoribosyltransferase (OPRT) (OPRTase) (EC 2.4.2.10); Orotidine 5'-phosphate decarboxylase (ODC) (OMPD) (EC 4.1.1.23) (OMPdecase)] | UMPS OK/SW-cl.21 |
| sp\|Q16674\|MIA_HUMAN | 157,065 | 182,383 | 243,840 | 257,310 | sp\|Q16674\|MIA_HUMAN | 161,137 | 62,401 | 39% | Q16674 | MIA_HUMAN | MIA_HUMAN | Melanoma-derived growth regulatory protein (Melanoma inhibitory activity protein) | MIA |
| sp\|Q07955\|SRSF1_HUMAN | 78,246 | 49,474 | 134,600 | 117,649 | sp\|Q07955\|SRSF1_HUMAN | 182,414 | 145,974 | 80% | Q07955 | SRSF1_HUMAN | SRSF1_HUMAN | Serine/arginine-rich splicing factor 1 (Alternative-splicing factor 1) (ASF-1) (Splicing factor, arginine/serine-rich 1) (pre-mRNA-splicing factor SF2, P33 subunit) | SRSF1 ASF SF2 SF2P33 SFRS1 OK/SW-cl.3 |
| sp\|O75153\|CLU_HUMAN | 19,811 | 4,052 | 44,053 | 47,704 | sp\|O75153\|CLU_HUMAN | 30,468 | 18,415 | 60% | O75153 | CLU_HUMAN | CLU_HUMAN | Clustered mitochondria protein homolog | CLUH KIAA0664 |
| sp\|Q08170\|SRSF4_HUMAN | 92,015 | 89,764 | 149,716 | 130,189 | sp\|Q08170\|SRSF4_HUMAN | 77,941 | 45,018 | 58% | Q08170 | SRSF4_HUMAN | SRSF4_HUMAN | Serine/arginine-rich splicing factor 4 (Pre-mRNA-splicing factor SRP75) (SRP001LB) (Splicing factor, arginine/serine-rich 4) | SRSF4 SFRS4 SRP75 |
| sp\|O75347\|TBCA_HUMAN | 70,973 | 65,782 | 183,121 | 218,755 | sp\|O75347\|TBCA_HUMAN | 143,018 | 91,620 | 64% | O75347 | TBCA_HUMAN | TBCA_HUMAN | Tubulin-specific chaperone A (TCP1-chaperonin cofactor A) (Tubulin-folding cofactor A) (CFA) | TBCA |
| sp\|Q01415\|GALK2_HUMAN | 69,802 | 64,028 | 61,612 | 95,119 | sp\|Q01415\|GALK2_HUMAN | 58,257 | 22,074 | 38% | Q01415 | GALK2_HUMAN | GALK2_HUMAN | N-acetylgalactosamine kinase (EC 2.7.1.157) (GalNAc kinase) (Galactokinase 2) | GALK2 GK2 |
| sp\|P12111\|CO6A3_HUMAN | 2,184 | 1,768 | 16,492 | 6,148 | sp\|P12111\|CO6A3_HUMAN | 40,193 | 40,159 | 100% | P12111 | CO6A3_HUMAN | CO6A3_HUMAN | Collagen alpha-3(VI) chain | COL6A3 |
| sp\|Q9C0B5\|ZDHC5_HUMAN | 95,253 | 107,124 | 215,014 | 203,251 | sp\|Q9C0B5\|ZDHC5_HUMAN | 149,060 | 72,460 | 49% | Q9C0B5 | ZDHC5_HUMAN | ZDHC5_HUMAN | Palmitoyltransferase ZDHHC5 (EC 2.3.1.225) (Zinc finger DHHC domain-containing protein 5) (DHHC-5) (Zinc finger protein 375) | ZDHHC5 KIAA1748 ZNF375 |

| | | | | | | | | | | | | | |
|---|---|---|---|---|---|---|---|---|---|---|---|---|---|
| sp\|O14578\|CTRO_HUMAN | 10,567 | 6,778 | 7,819 | 20,696 | sp\|O14578\|CTRO_HUMAN | 12,735 | 5,510 | 43% | O14578 | CTRO_HUMAN | CTRO_HUMAN | Citron Rho-interacting kinase (CRIK) (EC 2.7.11.1) (Serine/threonine-protein kinase 21) | CIT CRIK KIAA0949 STK21 |
| sp\|Q8IVT2\|MISP_HUMAN | 361,149 | 318,373 | 1,258,438 | 1,316,598 | sp\|Q8IVT2\|MISP_HUMAN | 841,075 | 479,711 | 57% | Q8IVT2 | MISP_HUMAN | MISP_HUMAN | Mitotic interactor and substrate of PLK1 (Mitotic spindle positioning protein) | MISP C19orf21 |
| sp\|Q6PI48\|SYDM_HUMAN | 30,429 | 26,681 | 36,843 | 37,457 | sp\|Q6PI48\|SYDM_HUMAN | 28,234 | 12,032 | 43% | Q6PI48 | SYDM_HUMAN | SYDM_HUMAN | Aspartate--tRNA ligase, mitochondrial (EC 6.1.1.12) (Aspartyl-tRNA synthetase) (AspRS) | DARS2 |
| sp\|P08779\|K1C16_HUMAN | 107,526 | 74,181 | 74,900 | 87,316 | sp\|P08779\|K1C16_HUMAN | 132,997 | 91,622 | 69% | P08779 | K1C16_HUMAN | K1C16_HUMAN | Keratin, type I cytoskeletal 16 (Cytokeratin-16) (CK-16) (Keratin-16) (K16) | KRT16 KRT16A |
| sp\|Q9UH62\|ARMX3_HUMAN | 12,662 | 22,095 | 11,972 | 27,125 | sp\|Q9UH62\|ARMX3_HUMAN | 18,713 | 11,881 | 63% | Q9UH62 | ARMX3_HUMAN | ARMX3_HUMAN | Armadillo repeat-containing X-linked protein 3 (ARM protein lost in epithelial cancers on chromosome X 3) (Protein ALEX3) | ARMCX3 ALEX3 BM-017 UNQ2517/PRO6007 |
| sp\|P28062\|PSB8_HUMAN | 38,383 | 46,080 | 72,699 | 18,017 | sp\|P28062\|PSB8_HUMAN | 46,846 | 25,121 | 54% | P28062 | PSB8_HUMAN | PSB8_HUMAN | Proteasome subunit beta type-8 (EC 3.4.25.1) (Low molecular mass protein 7) (Macropain subunit C13) (Multicatalytic endopeptidase complex subunit C13) (Proteasome component C13) (Proteasome subunit beta-5i) (Really interesting new gene 10 protein) | PSMB8 LMP7 PSMB5i RING10 Y2 |
| sp\|Q9NZN4\|EHD2_HUMAN | 15,406 | 23,917 | 8,045 | 8,473 | sp\|Q9NZN4\|EHD2_HUMAN | 31,220 | 24,431 | 78% | Q9NZN4 | EHD2_HUMAN | EHD2_HUMAN | EH domain-containing protein 2 (PAST homolog 2) | EHD2 PAST2 |
| sp\|O43681\|ASNA_HUMAN | 51,665 | 55,767 | 90,586 | 99,357 | sp\|O43681\|ASNA_HUMAN | 61,607 | 25,439 | 41% | O43681 | ASNA_HUMAN | GET3_HUMAN | ATPase GET3 (EC 3.6.-.-) (Arsenical pump-driving ATPase) (Arsenite-stimulated ATPase) (Guided entry of tail-anchored proteins factor 3, ATPase) (Transmembrane domain recognition complex 40 kDa ATPase subunit) (hARSA-I) (hASNA-I) | GET3 ARSA ASNA1 TRC40 |
| sp\|O94979\|SC31A_HUMAN | 11,067 | 19,197 | 31,376 | 15,540 | sp\|O94979\|SC31A_HUMAN | 26,244 | 18,431 | 70% | O94979 | SC31A_HUMAN | SC31A_HUMAN | Protein transport protein Sec31A (ABP125) (ABP130) (SEC31-like protein 1) (SEC31-related protein A) (Web1-like protein) | SEC31A KIAA0905 SEC31L1 HSPC275 HSPC334 |
| sp\|P41217\|OX2G_HUMAN | 63,247 | 78,070 | 5,471 | 13,112 | sp\|P41217\|OX2G_HUMAN | 33,567 | 31,383 | 93% | P41217 | OX2G_HUMAN | OX2G_HUMAN | OX-2 membrane glycoprotein (CD antigen CD200) | CD200 MOX1 MOX2 My033 |
| sp\|P39748\|FEN1_HUMAN | 93,056 | 89,483 | 51,881 | 15,629 | sp\|P39748\|FEN1_HUMAN | 56,575 | 34,843 | 62% | P39748 | FEN1_HUMAN | FEN1_HUMAN | Flap endonuclease 1 (FEN-1) (EC 3.1.-.-) (DNase IV) (Flap structure-specific endonuclease 1) (Maturation factor 1) (MF1) (hFEN-1) | FEN1 RAD2 |
| sp\|Q9NRH3\|TBG2_HUMAN | 71,333 | 63,305 | 136,724 | 128,201 | sp\|Q9NRH3\|TBG2_HUMAN | 94,388 | 33,796 | 36% | Q9NRH3 | TBG2_HUMAN | TBG2_HUMAN | Tubulin gamma-2 chain (Gamma-2-tubulin) | TUBG2 |
| sp\|P57105\|SYJ2B_HUMAN | 11,056 | 13,717 | 4,512 | 103,988 | sp\|P57105\|SYJ2B_HUMAN | 39,330 | 36,863 | 94% | P57105 | SYJ2B_HUMAN | SYJ2B_HUMAN | Synaptojanin-2-binding protein (Mitochondrial outer membrane protein 25) | SYNJ2BP OMP25 |
| sp\|O15400\|STX7_HUMAN | 18,966 | 21,712 | 32,323 | 7,751 | sp\|O15400\|STX7_HUMAN | 19,638 | 9,108 | 46% | O15400 | STX7_HUMAN | STX7_HUMAN | Syntaxin-7 | STX7 |
| sp\|Q9Y3E7\|CHMP3_HUMAN | 36,953 | 48,035 | 10,923 | 23,115 | sp\|Q9Y3E7\|CHMP3_HUMAN | 31,816 | 17,356 | 55% | Q9Y3E7 | CHMP3_HUMAN | CHMP3_HUMAN | Charged multivesicular body protein 3 (Chromatin-modifying protein 3) (Neuroendocrine differentiation factor) (Vacuolar protein sorting-associated protein 24) (hVps24) | CHMP3 CGI149 NEDF VPS24 CGI-149 |

| | | | | | | | | | | | | | |
|---|---|---|---|---|---|---|---|---|---|---|---|---|---|
| sp\|P06703\|S10A6_HUMAN | 245,034 | 304,210 | 494,491 | 475,449 | sp\|P06703\|S10A6_HUMAN | 361,683 | 90,576 | 25% | P06703 | S10A6_HUMAN | S10A6_HUMAN | Protein S100-A6 (Calcyclin) (Growth factor-inducible protein 2A9) (MLN 4) (Prolactin receptor-associated protein) (PRA) (S100 calcium-binding protein A6) | S100A6 CACY |
| sp\|P84103\|SRSF3_HUMAN | 142,232 | 101,712 | 265,808 | 154,991 | sp\|P84103\|SRSF3_HUMAN | 117,129 | 80,445 | 69% | P84103 | SRSF3_HUMAN | SRSF3_HUMAN | Serine/arginine-rich splicing factor 3 (Pre-mRNA-splicing factor SRP20) (Splicing factor, arginine/serine-rich 3) | SRSF3 SFRS3 SRP20 |
| sp\|Q04695\|K1C17_HUMAN | 38,229 | 49,097 | 56,451 | 25,726 | sp\|Q04695\|K1C17_HUMAN | 65,419 | 34,981 | 53% | Q04695 | K1C17_HUMAN | K1C17_HUMAN | Keratin, type I cytoskeletal 17 (39.1) (Cytokeratin-17) (CK-17) (Keratin-17) (K17) | KRT17 |
| sp\|Q9UBC2\|EP15R_HUMAN | 40,529 | 30,156 | 49,817 | 35,962 | sp\|Q9UBC2\|EP15R_HUMAN | 36,548 | 15,526 | 42% | Q9UBC2 | EP15R_HUMAN | EP15R_HUMAN | Epidermal growth factor receptor substrate 15-like 1 (Eps15-related protein) (Eps15R) | EPS15L1 EPS15R |
| sp\|O00629\|IMA3_HUMAN | 49,022 | 69,609 | 19,835 | 57,795 | sp\|O00629\|IMA3_HUMAN | 45,107 | 23,402 | 52% | O00629 | IMA3_HUMAN | IMA3_HUMAN | Importin subunit alpha-3 (Importin alpha Q1) (Qip1) (Karyopherin subunit alpha-4) | KPNA4 QIP1 |
| sp\|Q96G97\|BSCL2_HUMAN | 27,450 | 30,200 | 29,364 | 18,459 | sp\|Q96G97\|BSCL2_HUMAN | 42,083 | 28,010 | 67% | Q96G97 | BSCL2_HUMAN | BSCL2_HUMAN | Seipin (Bernardinelli-Seip congenital lipodystrophy type 2 protein) | BSCL2 |
| sp\|P31997\|CEAM8_HUMAN | 428,426 | 566,204 | 12,490 | 1,417,471 | sp\|P31997\|CEAM8_HUMAN | 476,397 | 480,899 | 101% | P31997 | CEAM8_HUMAN | CEAM8_HUMAN | Carcinoembryonic antigen-related cell adhesion molecule 8 (CD67 antigen) (Carcinoembryonic antigen CGM6) (Non-specific cross-reacting antigen NCA-95) (CD antigen CD66b) | CEACAM8 CGM6 |
| sp\|Q00765\|REEP5_HUMAN | 1,940 | 3,409 | 2,873 | 3,111 | sp\|Q00765\|REEP5_HUMAN | 6,866 | 11,011 | 160% | Q00765 | REEP5_HUMAN | REEP5_HUMAN | Receptor expression-enhancing protein 5 (Polyposis locus protein 1) (Protein TB2) | REEP5 C5orf18 DP1 TB2 |
| sp\|Q9NVD7\|PARVA_HUMAN | 45,639 | 41,598 | 41,581 | 12,259 | sp\|Q9NVD7\|PARVA_HUMAN | 42,605 | 14,702 | 35% | Q9NVD7 | PARVA_HUMAN | PARVA_HUMAN | Alpha-parvin (Actopaxin) (CH-ILKBP) (Calponin-like integrin-linked kinase-binding protein) (Matrix-remodeling-associated protein 2) | PARVA MXRA2 |
| sp\|Q86Y82\|STX12_HUMAN | 27,404 | 30,347 | 9,601 | 42,948 | sp\|Q86Y82\|STX12_HUMAN | 34,728 | 15,876 | 46% | Q86Y82 | STX12_HUMAN | STX12_HUMAN | Syntaxin-12 | STX12 |
| sp\|Q9NQ34\|TMM9B_HUMAN | 46,656 | 50,656 | 34,997 | 30,653 | sp\|Q9NQ34\|TMM9B_HUMAN | 43,828 | 16,631 | 38% | Q9NQ34 | TMM9B_HUMAN | TMM9B_HUMAN | Transmembrane protein 9B | TMEM9B C11orf15 UNQ712/PRO1375 |
| sp\|P07602\|SAP_HUMAN | 71,496 | 60,385 | 79,414 | 26,459 | sp\|P07602\|SAP_HUMAN | 46,752 | 21,533 | 46% | P07602 | SAP_HUMAN | SAP_HUMAN | Prosaposin (Proactivator polypeptide) [Cleaved into: Saposin-A (Protein A); Saposin-B-Val; Saposin-B (Cerebroside sulfate activator) (CSAct) (Dispersin) (Sphingolipid activator protein 1) (SAP-1) (Sulfatide/GM1 activator); Saposin-C (A1 activator) (Co-beta-glucosidase) (Glucosylceramidase activator) (Sphingolipid activator protein 2) (SAP-2); Saposin-D (Component C) (Protein C)] | PSAP GLBA SAP1 |
| sp\|Q12846\|STX4_HUMAN | 142,387 | 156,301 | 115,135 | 110,113 | sp\|Q12846\|STX4_HUMAN | 114,780 | 23,162 | 20% | Q12846 | STX4_HUMAN | STX4_HUMAN | Syntaxin-4 (Renal carcinoma antigen NY-REN-31) | STX4 STX4A |
| sp\|P48059\|LIMS1_HUMAN | 39,260 | 46,357 | 38,991 | 43,191 | sp\|P48059\|LIMS1_HUMAN | 45,944 | 15,816 | 34% | P48059 | LIMS1_HUMAN | LIMS1_HUMAN | LIM and senescent cell antigen-like-containing domain protein 1 (Particularly interesting new Cys-His protein 1) (PINCH-1) (Renal carcinoma antigen NY-REN-48) | LIMS1 PINCH PINCH1 |

| | | | | | | | | | | | | | |
|---|---|---|---|---|---|---|---|---|---|---|---|---|---|
| sp\|Q96CC6\|RHDF1_HUMAN | 26,552 | 24,445 | 131,979 | 114,256 | sp\|Q96CC6\|RHDF1_HUMAN | 90,565 | 66,424 | 73% | Q96CC6 | RHDF1_HUMAN | RHDF1_HUMAN | Inactive rhomboid protein 1 (iRhom1) (Epidermal growth factor receptor-related protein) (Rhomboid 5 homolog 1) (Rhomboid family member 1) (p100hRho) | RHBDF1 C16orf8 DIST1 IRHOM1 |
| sp\|Q8TBQ9\|KISHA_HUMAN | 57,705 | 38,357 | 152,021 | 87,045 | sp\|Q8TBQ9\|KISHA_HUMAN | 77,567 | 40,092 | 52% | Q8TBQ9 | KISHA_HUMAN | KISHA_HUMAN | Protein kish-A (Transmembrane protein 167) (Transmembrane protein 167A) | TMEM167A TMEM167 |
| sp\|Q07157\|ZO1_HUMAN | 40,404 | 42,809 | 13,162 | 16,612 | sp\|Q07157\|ZO1_HUMAN | 26,078 | 13,292 | 51% | Q07157 | ZO1_HUMAN | ZO1_HUMAN | Tight junction protein ZO-1 (Tight junction protein 1) (Zona occludens protein 1) (Zonula occludens protein 1) | TJP1 ZO1 |
| sp\|Q93050\|VPP1_HUMAN | 68,566 | 60,047 | 105,107 | 94,938 | sp\|Q93050\|VPP1_HUMAN | 62,398 | 29,243 | 47% | Q93050 | VPP1_HUMAN | VPP1_HUMAN | V-type proton ATPase 116 kDa subunit a 1 (V-ATPase 116 kDa subunit a 1) (Clathrin-coated vesicle/synaptic vesicle proton pump 116 kDa subunit) (Vacuolar adenosine triphosphatase subunit Ac116) (Vacuolar proton pump subunit 1) (Vacuolar proton translocating ATPase 116 kDa subunit a isoform 1) | ATP6V0A1 ATP6N1 ATP6N1A VPP1 |
| sp\|Q6RW13\|ATRAP_HUMAN | 46,500 | 8,848 | 14,282 | 61,532 | sp\|Q6RW13\|ATRAP_HUMAN | 34,860 | 19,639 | 56% | Q6RW13 | ATRAP_HUMAN | ATRAP_HUMAN | Type-1 angiotensin II receptor-associated protein (AT1 receptor-associated protein) | AGTRAP ATRAP |
| sp\|Q8TCT8\|SPP2A_HUMAN | 76,347 | 87,825 | 4,610 | 10,549 | sp\|Q8TCT8\|SPP2A_HUMAN | 37,483 | 33,430 | 89% | Q8TCT8 | SPP2A_HUMAN | SPP2A_HUMAN | Signal peptide peptidase-like 2A (SPP-like 2A) (SPPL2a) (EC 3.4.23.-) (Intramembrane protease 3) (IMP-3) (Presenilin-like protein 2) | SPPL2A IMP3 PSL2 PSEC0147 |
| sp\|Q7Z2W9\|RM21_HUMAN | 70,927 | 54,520 | 6,432 | 15,960 | sp\|Q7Z2W9\|RM21_HUMAN | 27,768 | 24,144 | 87% | Q7Z2W9 | RM21_HUMAN | RM21_HUMAN | 39S ribosomal protein L21, mitochondrial (L21mt) (MRP-L21) (Mitochondrial large ribosomal subunit protein bL21m) | MRPL21 |
| sp\|Q53GA4\|PHLA2_HUMAN | 109,190 | 86,327 | 115,390 | 135,646 | sp\|Q53GA4\|PHLA2_HUMAN | 171,892 | 88,710 | 52% | Q53GA4 | PHLA2_HUMAN | PHLA2_HUMAN | Pleckstrin homology-like domain family A member 2 (Beckwith-Wiedemann syndrome chromosomal region 1 candidate gene C protein) (Imprinted in placenta and liver protein) (Tumor-suppressing STF cDNA 3 protein) (Tumor-suppressing subchromosomal transferable fragment candidate gene 3 protein) (p17-Beckwith-Wiedemann region 1 C) (p17-BWR1C) | PHLDA2 BWR1C HLDA2 IPL TSSC3 |
| sp\|P04003\|C4BPA_HUMAN | 18,707 | 2,513 | 16,945 | 24,259 | sp\|P04003\|C4BPA_HUMAN | 68,160 | 85,375 | 125% | P04003 | C4BPA_HUMAN | C4BPA_HUMAN | C4b-binding protein alpha chain (C4bp) (Proline-rich protein) (PRP) | C4BPA C4BP |
| sp\|Q9H0H5\|RGAP1_HUMAN | 95,808 | 173,775 | 167,056 | 301,858 | sp\|Q9H0H5\|RGAP1_HUMAN | 169,511 | 66,201 | 39% | Q9H0H5 | RGAP1_HUMAN | RGAP1_HUMAN | Rac GTPase-activating protein 1 (Male germ cell RacGap) (MgcRacGAP) (Protein CYK4 homolog) (CYK4) (HsCYK-4) | RACGAP1 KIAA1478 MGCRACGAP |
| sp\|Q9UEY8\|ADDG_HUMAN | 36,074 | 43,859 | 17,983 | 12,646 | sp\|Q9UEY8\|ADDG_HUMAN | 32,254 | 11,434 | 35% | Q9UEY8 | ADDG_HUMAN | ADDG_HUMAN | Gamma-adducin (Adducin-like protein 70) | ADD3 ADDL |
| sp\|Q13303\|KCAB2_HUMAN | 6,455 | 20,111 | 14,730 | 11,461 | sp\|Q13303\|KCAB2_HUMAN | 25,779 | 31,052 | 120% | Q13303 | KCAB2_HUMAN | KCAB2_HUMAN | Voltage-gated potassium channel subunit beta-2 (EC 1.1.1.-) (K(+) channel subunit beta-2) (Kv-beta-2) (hKvbeta2) | KCNAB2 KCNA2B KCNK2 |

| | | | | | | | | | | | | | |
|---|---|---|---|---|---|---|---|---|---|---|---|---|---|
| sp\|Q9BSJ2\|GCP2_HUMAN | 6,830 | 2,363 | 19,497 | 8,739 | sp\|Q9BSJ2\|GCP2_HUMAN | 41,988 | 59,097 | 141% | Q9BSJ2 | GCP2_HUMAN | GCP2_HUMAN | Gamma-tubulin complex component 2 (GCP-2) (hGCP2) (Gamma-ring complex protein 103 kDa) (h103p) (hGrip103) (Spindle pole body protein Spc97 homolog) (hSpc97) | TUBGCP2 GCP2 |
| sp\|P09874\|PARP1_HUMAN | 96,670 | 50,740 | 170,843 | 65,975 | sp\|P09874\|PARP1_HUMAN | 58,443 | 53,651 | 92% | P09874 | PARP1_HUMAN | PARP1_HUMAN | Poly [ADP-ribose] polymerase 1 (PARP-1) (EC 2.4.2.30) (ADP-ribosyltransferase diphtheria toxin-like 1) (ARTD1) (DNA ADP-ribosyltransferase PARP1) (EC 2.4.2.-) (NAD(+) ADP-ribosyltransferase 1) (ADPRT 1) (Poly[ADP-ribose] synthase 1) (Protein poly-ADP-ribosyltransferase PARP1) (EC 2.4.2.-) | PARP1 ADPRT PPOL |
| sp\|Q92629\|SGCD_HUMAN | 231,302 | 210,284 | 163,590 | 153,257 | sp\|Q92629\|SGCD_HUMAN | 161,275 | 44,396 | 28% | Q92629 | SGCD_HUMAN | SGCD_HUMAN | Delta-sarcoglycan (Delta-SG) (35 kDa dystrophin-associated glycoprotein) (35DAG) | SGCD |
| sp\|P25189\|MYP0_HUMAN | 173,272 | 128,058 | 218,863 | 208,163 | sp\|P25189\|MYP0_HUMAN | 146,389 | 55,554 | 38% | P25189 | MYP0_HUMAN | MYP0_HUMAN | Myelin protein P0 (Myelin peripheral protein) (MPP) (Myelin protein zero) | MPZ |
| sp\|P48740\|MASP1_HUMAN | 11,055 | 4,913 | 5,105 | 10,879 | sp\|P48740\|MASP1_HUMAN | 28,863 | 24,231 | 84% | P48740 | MASP1_HUMAN | MASP1_HUMAN | Mannan-binding lectin serine protease 1 (EC 3.4.21.-) (Complement factor MASP-3) (Complement-activating component of Ra-reactive factor) (Mannose-binding lectin-associated serine protease 1) (MASP-1) (Mannose-binding protein-associated serine protease) (Ra-reactive factor serine protease p100) (RaRF) (Serine protease 5) [Cleaved into: Mannan-binding lectin serine protease 1 heavy chain; Mannan-binding lectin serine protease 1 light chain] | MASP1 CRARF CRARF1 PRSS5 |
| sp\|Q86W92\|LIPB1_HUMAN | 30,759 | 66,571 | 22,822 | 7,561 | sp\|Q86W92\|LIPB1_HUMAN | 36,042 | 19,288 | 54% | Q86W92 | LIPB1_HUMAN | LIPB1_HUMAN | Liprin-beta-1 (Protein tyrosine phosphatase receptor type f polypeptide-interacting protein-binding protein 1) (PTPRF-interacting protein-binding protein 1) (hSGT2) | PPFIBP1 KIAA1230 |
